# A broadly resolved molecular phylogeny of New Zealand cheilostome bryozoans as a framework for hypotheses of morphological evolution

**DOI:** 10.1101/2020.12.08.415943

**Authors:** RJS Orr, E Di Martino, DP Gordon, MH Ramsfjell, HL Mello, AM Smith, LH Liow

**Affiliations:** Natural History Museum, University of Oslo, Oslo, Norway; National Institute of Water and Atmospheric Research, Wellington, New Zealand; Department of Marine Science, University of Otago, Dunedin, New Zealand; Centre for Ecological and Evolutionary Synthesis, Department of Biosciences, University of Oslo, Oslo, Norway

**Keywords:** High-throughput sequencing (HTS), genome-skimming, cheilostome bryozoans, phylogenetics, frontal shield, mitogenome

## Abstract

Larger molecular phylogenies based on ever more genes are becoming commonplace with the advent of cheaper and more streamlined sequencing and bioinformatics pipelines. However, many groups of inconspicuous but no less evolutionarily or ecologically important marine invertebrates are still neglected in the quest for understanding species- and higher-level phylogenetic relationships. Here, we alleviate this issue by presenting the molecular sequences of 165 cheilostome bryozoan species from New Zealand waters. New Zealand is our geographic region of choice as its cheilostome fauna is taxonomically, functionally and ecologically diverse, and better characterized than many other such faunas in the world. Using this most taxonomically broadly-sampled and statistically-supported cheilostome phylogeny comprising 214 species, when including previously published sequences, we tested several existing systematic hypotheses based solely on morphological observations. We find that lower taxonomic level hypotheses (species and genera) are robust while our inferred trees did not reflect current higher-level systematics (family and above), illustrating a general need for the rethinking of current hypotheses. To illustrate the utility of our new phylogeny, we reconstruct the evolutionary history of frontal shields (i.e., a calcified bodywall layer in ascus-bearing cheilostomes) and asked if its presence has any bearing on the diversification rates of cheilostomes.

## 1. Introduction

Large and broadly-sampled phylogenies are vital to robustly answering many different classes of evolutionary questions, including those involving trait evolution, origins and evolution of biogeographic distributions and rates of taxonomic diversification. While megaphylogenies with hundreds to thousands of species (Smith et al., 2009) are available for many groups of vertebrates (Meredith et al., 2011; Prum et al., 2015) and plants (Zanne et al., 2014), and also for some non-vertebrate terrestrial groups (Varga et al., 2019), the molecular phylogenetics of many marine invertebrate groups remains relatively neglected (Arrigoni et al., 2017; Kocot et al., 2018; O’Hara et al., 2017).

In this contribution, we begin to rectify the paucity of large and/or taxonomically broadly sampled molecular phylogenies for marine invertebrates, targeting a phylum whose rich fossil record can be subsequently integrated for evolutionary analyses. Our focal group is Cheilostomatida, the dominant living order of the colonial metazoan phylum Bryozoa, with c. 5200 described extant species, corresponding to > 80% of the living species diversity of the phylum (Bock and Gordon, 2013). Cheilostomes first appeared in the fossil record in the Late Jurassic (c. 160 million years ago) and then displayed a spectacular diversification c. 55 million years later in the mid-Cretaceous (Taylor, 2020). Cheilostomes, common in benthic marine habitats globally, are lightly- to heavily-calcified and largely sessile as adults. Most species are encrusting, while fewer are erect, with some forming robust structures whereas many are small and inconspicuous (Fig. 1). Although a number of cheilostome bryozoans have been sequenced and placed in a molecular phylogenetic context (Fuchs et al., 2009; Knight et al., 2011; Orr et al., 2019a; Waeschenbach et al., 2012) the systematics of cheilostome bryozoans aimed at reflecting their evolutionary relationships still remain largely based on morphological characters (Bock and Gordon, 2013; Taylor and Waeschenbach, 2015). This is in part because cheilostome phylogenetic relationships have only recently benefited from high-throughput sequencing (HTS) techniques and the increased phylogenetic support it provides (Orr et al., 2019a, b, 2000). HTS yields more sequence data with lesser effort compared with traditional PCR and Sanger sequencing techniques (Fuchs et al., 2009; Knight et al., 2011; Waeschenbach et al., 2012). By applying genome-skimming approaches to greatly expand on the taxonomic sampling of cheilostomes for molecular phylogenetics, we independently test phylogenetic hypotheses implicit in their current systematics (Bock, 2020), and also facilitate future studies.

**Fig. 1.**
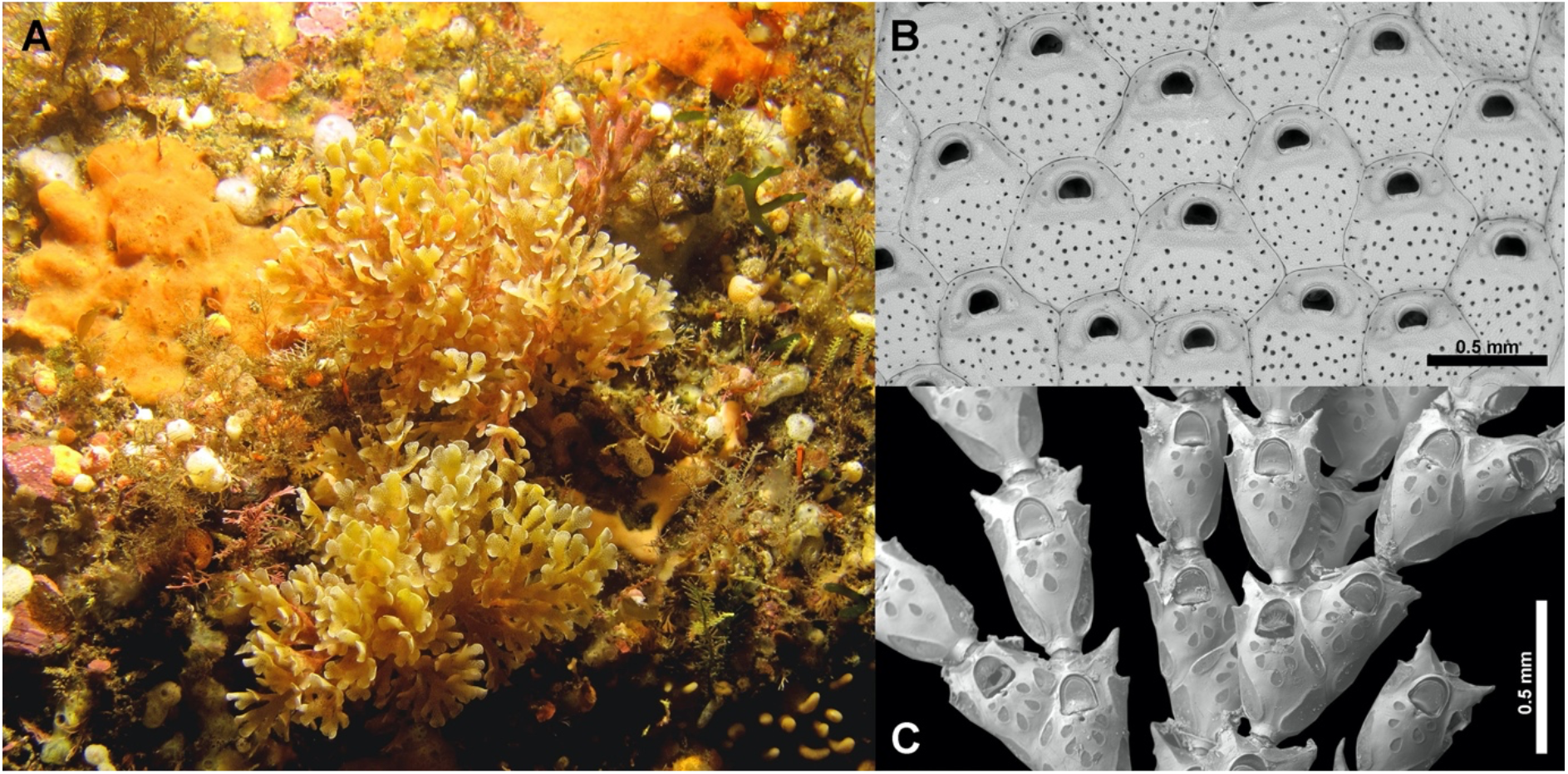
New Zealand Bryozoans. (A) Foliose branching colonies of the cheilostome bryozoan *Euthyroides episcopalis* from Fiordland, New Zealand (photo by Dr Mike Page, NIWA). (B) Scanning electron micrographs of an undescribed encrusting cheilostome, *Monoporella* n. sp. (BLEED 1360), from Three Kings Shelf, New Zealand. (C) The erect jointed catenicellid *Orthoscuticella fusiformis* (BLEED 1623).

We focus our sequencing effort in this contribution primarily on New Zealand cheilostomes for a number of reasons. Cheilostomes play a conspicuous role as habitat-building organisms in New Zealand as well as other temperate areas (Wood et al., 2012). In fact, some cheilostome thicket communities (Fig. 1) are protected in New Zealand because of their function as nurseries for commercial fish stocks (Bradstock and Gordon, 1983). As important components of marine communities, cheilostomes are crucial members of the marine food chain globally. This is because, like all bryozoans, they are efficient suspension-feeders (Gordon et al., 1987) while also providing food for other organismal groups (Lidgard, 2008). Cheilostomes are highly diverse in New Zealand, thanks to a combination of factors, including New Zealand’s geological and hydrographic setting, constituting the major part of the geological continent of Zealandia, which is 94% submerged (Campbell and Mortimer, 2014). Additionally, the New Zealand Exclusive Economic Zone, plus its extended continental shelf, is one of the largest in the world (5.7 million km^2^) with a wide latitudinal spread from subtropical to subantarctic (c. 23°-57.5° S). It also has varied seafloor topography, including extensive deep shelves, plateaus, ridges and seamounts (Gordon et al., 2010). Within this area, New Zealand has 359 genera and 1053 species of marine Bryozoa, including 867 cheilostomes (of which 285 species remain to be formally described). About 61% of New Zealand’s marine Bryozoa are endemic (Gordon et al., 2019), making New Zealand a doubtless diversity hotspot for cheilostome bryozoans. Complementing Recent diversity, the published Cenozoic record of cheilostome bryozoans is also rich, though relatively less studied (Brown, 1952; Gordon and Taylor, 2015; Rust and Gordon, 2011), comprising 531 species (of which 240 are in open nomenclature). This complementarity of living and fossil species renders a molecular phylogeny of New Zealand taxa amenable to modern statistical methods that integrate molecular and fossil data for inferring evolutionary processes (Heath et al., 2014). Last, but not least, New Zealand is one of the better-studied marine regions taxonomically and ecologically for Bryozoa (e.g. Gordon, 1984, 1986, 1989; Gordon et al., 2009; Schack et al., 2020), a phylum that is somewhat neglected in many other parts of the world. Bryozoan research has been continuously conducted in New Zealand since 1841 (Gordon et al., 2009) and a governmental agency, the National Institute of Water and Atmospheric Research (NIWA), is both the data manager and custodian for fisheries and invertebrate research data, hence assuring knowledge curation. All of this means that a cheilostome phylogeny with New Zealand species broadly represented allows us to begin to ask evolutionary and ecological questions while controlling for phylogenetic nonindependence.

Here we apply a genome-skimming approach to New Zealand cheilostome bryozoans and present a robustly supported molecular phylogeny based on 15 mitochondrial and 2 rRNA genes. The molecular sequences of 199 cheilostome colonies sampled in New Zealand are presented here for the first time. Using 180 species and 96 genera from New Zealand and previously sequenced, non-New Zealand species, we construct the largest and most taxonomically broadly sampled cheilostome phylogeny to date, with 263 in-group colonies, representing 214 species and 120 genera. The inclusion of non-New Zealand taxa allows us to explore the robustness of the inferred relationships among New Zealand species but also reduces phylogenetic inference errors by nature of a broader taxonomic sampling (Pollock et al., 2002). To illustrate the utility of our inferred tree for understanding cheilostome evolution, we reconstruct the evolutionary history of a morphological trait (the calcified frontal shield) and ask if its evolution might have changed the diversification rates of cheilostomes that have such a shield (ascophoran-grade) versus those that do not (anascan-grade). We also discuss several other key taxonomic traits widely thought to be evolutionarily stable and the consequences our highly resolved cheilostome phylogeny has for these. Our contribution is a first step towards a global cheilostome megaphylogeny, needed for answering biological questions that go beyond those probing genealogical relationships.

## 2. Methods

### 2.1. Sampling & SEM

Sequences are provided here for 207 New Zealand cheilostome colonies that were collected during several field expeditions by NIWA and University of Otago, New Zealand. While we have newly sequenced 199 colonies, we also supply unpublished sequences for 8 extra colonies we previously presented (see Supplementary Table S2). Samples were sorted, preserved in 70-96% ethanol, then shipped to the University of Oslo, Norway, for processing. Each bryozoan colony, preliminarily identified to the lowest possible taxonomic level (usually genus but sometimes species) using a stereoscope, was subsampled for DNA isolation, and also for scanning electron microscopy. The scanning electron micrographs (SEMs), taken with a Hitachi TM4040PLus after bleaching to remove tissue (where appropriate), are required for species-level confirmation. All SEM digital vouchers are supplied as a supplementary data file. Taxonomic identifications are made independently of the phylogenetic inference and metadata to avoid identification bias.

### 2.2. DNA isolation, sequencing and assembly

The 199 subsamples of colonies (henceforth “samples”) were dried before genomic DNA isolation using the DNeasy Blood and Tissue kit (QIAGEN, Germantown, MD, USA). Samples were homogenized in lysis buffer, using a pestle, in the presence of proteinase-K. Genomic DNA were sequenced at the Norwegian Sequencing Centre (Oslo, Norway) using Illumina HiSeq4000 150 bp paired-end (PE) sequencing with a 350 bp insert size. Approximately 20 samples (library preps) were genome-skimmed (multiplexed) on a single lane. Illumina HiSeq reads were quality checked using FastQC v.0.11.8 (Andrews, 2010), then quality- and adapter-trimmed using TrimGalore v0.4.4 with a Phred score cutoff of 30 (Krueger, 2015). Trimmed reads were *de novo* assembled with SPAdes 3.13 (Bankevich et al., 2012) using k-mers of 21, 33, 55, 77, 99 and 127. The mitogenome and rRNA operon of each sample were identified separately with blastn (Altschul et al., 1990) using blast+ against a database constructed from broadly sampled cheilostome sequences already deposited in NCBI (Orr et al., 2020). An E-value of 1.00e-185 and maximum target sequence of 1 were used to filter any blast hits of non-cheilostome origin.

### 2.3. Annotation

Mitogenomes for each of the samples were annotated with Mitos2 using a metazoan reference (RefSeq 89) and the invertebrate genetic code (Bernt et al., 2013) to identify two rRNA genes (rrnL and rrnS) and 13 protein coding genes *(atp6, atp8, cox1, cox2, cox3, cob, nad1, nad2, nad3, nad4, nad4l, nad5,* and *nad6*). In addition, two rRNA operon genes (ssu/18s and lsu/28s) were identified and annotated using RNAmmer (Lagesen et al., 2007). Eight published (Orr et al., 2019a, 2020) New Zealand samples (BLEED 48, 104, 127, 196, 344, 694, 1267 and 1687) were included in the subsequent workflow to bring the total number of samples to 229. Further, the mitogenomes and rRNA operons of 38 non-New Zealand bryozoans (Orr et al., 2020), were aligned with our samples to compile a broader cheilostome ingroup and ctenostome outgroup taxon sample.

### 2.4. Aligning

MAFFT (Katoh and Standley, 2013) was used for alignment with default parameters: for the four rRNA genes (nucleotide) the Q-INS-i model, considering secondary RNA structure, was utilized; for the 13 protein-coding genes, in amino acid format, the G-INS-I model was used. The 17 separate alignments were edited manually using Mesquite v3.61 to remove any uncertain characters (Maddison and Maddison, 2017). Ambiguously aligned characters were removed from each alignment using Gblocks (Talavera and Castresana, 2007) with least stringent parameters. The single-gene alignments were concatenated to a supermatrix using the catfasta2phyml perl script (Nylander, 2010). The alignments (both masked and unmasked) are available through Dryad (https://doi.org/10.5061/dryad.7pvmcvdrs)

### 2.5. Phylogenetic reconstruction

Maximum likelihood (ML) phylogenetic analyses were carried out for each single gene alignment using the “AUTO” parameter in RAxML v8.0.26 (Stamatakis, 2006) to establish the evolutionary model with the best fit. The general time reversible (GTR+G) was the preferred model for the four rRNA genes (18s, 28s, rrnS and rrnL), and MtZoa+G for all 13 protein coding genes. The two concatenated datasets (“New Zealand” and “global” = New Zealand + non-New Zealand, see section above), divided into rRNA and protein gene partitions, each with its own separate gamma distribution were analyzed using RAxML. The topology with the highest likelihood score of 100 heuristic searches was chosen. Bootstrap values were calculated from 500 pseudo-replicates.

Bayesian inference (BI) was performed using a modified version of MrBayes incorporating the MtZoa evolutionary model (Huelsenbeck and Ronquist, 2001; Tanabe, 2016). The datasets were executed, as before, with rRNA and protein gene partitions under their separate gamma distributions. Two independent runs, each with three heated and one cold Markov Chain Monte Carlo (MCMC) chain, were initiated from a random starting tree. The MCMC chains were run for 20,000,000 generations with trees sampled every 1,000^th^ generation. The posterior probabilities and mean marginal likelihood values of the trees were calculated after the burnin phase (5,000,000 generations). The average standard deviation of split frequencies between the two runs was <0.01, indicating convergence of the MCMC chains.

Congruence between the topological signal of the ML and Bayesian trees for both the New Zealand (Supplementary Figs. S1 and S2) and global (Supplementary Figs. S3 and S4) inferences was tested using the *I_cong_* index (de Vienne et al., 2007).

### 2.6. Ancestral state reconstruction and BiSSE analyses

The tips states of whether the sampled species is anascan (0, having a non-calcified frontal membrane) or ascophoran (1, having a calcified frontal shield), both states decipherable from SEMs, is given in in Fig. 3. We use a standard Markov model of binary character evolution (Pagel, 1994) implemented in ape (Paradis and Schliep, 2018) to estimate the ancestral states of the nodes on our inferred phylogeny. We use a standard binary state speciation and extinction model (Maddison et al., 2007) implemented in diversitree (FitzJohn, 2012) to investigate any differences in diversification rates due to the anascan or ascophoran frontal shield state of the species involved. As input for this latter analysis, we estimate that of the 1876 anascans and 3358 ascophoran species in Bock 2020, we have sampled 4.4% and 3.9% respectively to account for biases due to the sampling of species given the trait. We perform ancestral state reconstruction and BiSSE analyses for both ML and Bayesian “global” trees (Supplementary Figs. S3 and S4 respectively) to account for minor differences in the topological signal (see Results). In cases where there are multiple representatives within a species, we choose the colony with the highest number of nucleotides/amino-acids/genes to represent the species for these analyses.

## 3. Results

### 3.1. Sequencing and concatenation

We successfully sequenced and assembled 199 New Zealand cheilostome colonies, representing 165 species (SEM vouchers in Supplementary file) that have never been presented previously (Supplementary Table S1). We supply additional sequence data for a further eight species previously presented (Supplementary Table S2 and Orr et al., 2019b). The final 17 gene and 267 taxa “global” supermatrix constitutes 77% total character completeness for the dataset used to infer Fig. 2. For the convenience of future workers interested in only the New Zealand taxa, we supply also trees based on these data (Supplementary Figs. S1 (ML) and S2 (Bayesian), where character completeness is 78%). The assembled rRNA and mitogenomes are deposited at NCBI with accession numbers (Supplementary Table S2).

**Fig. 2.**
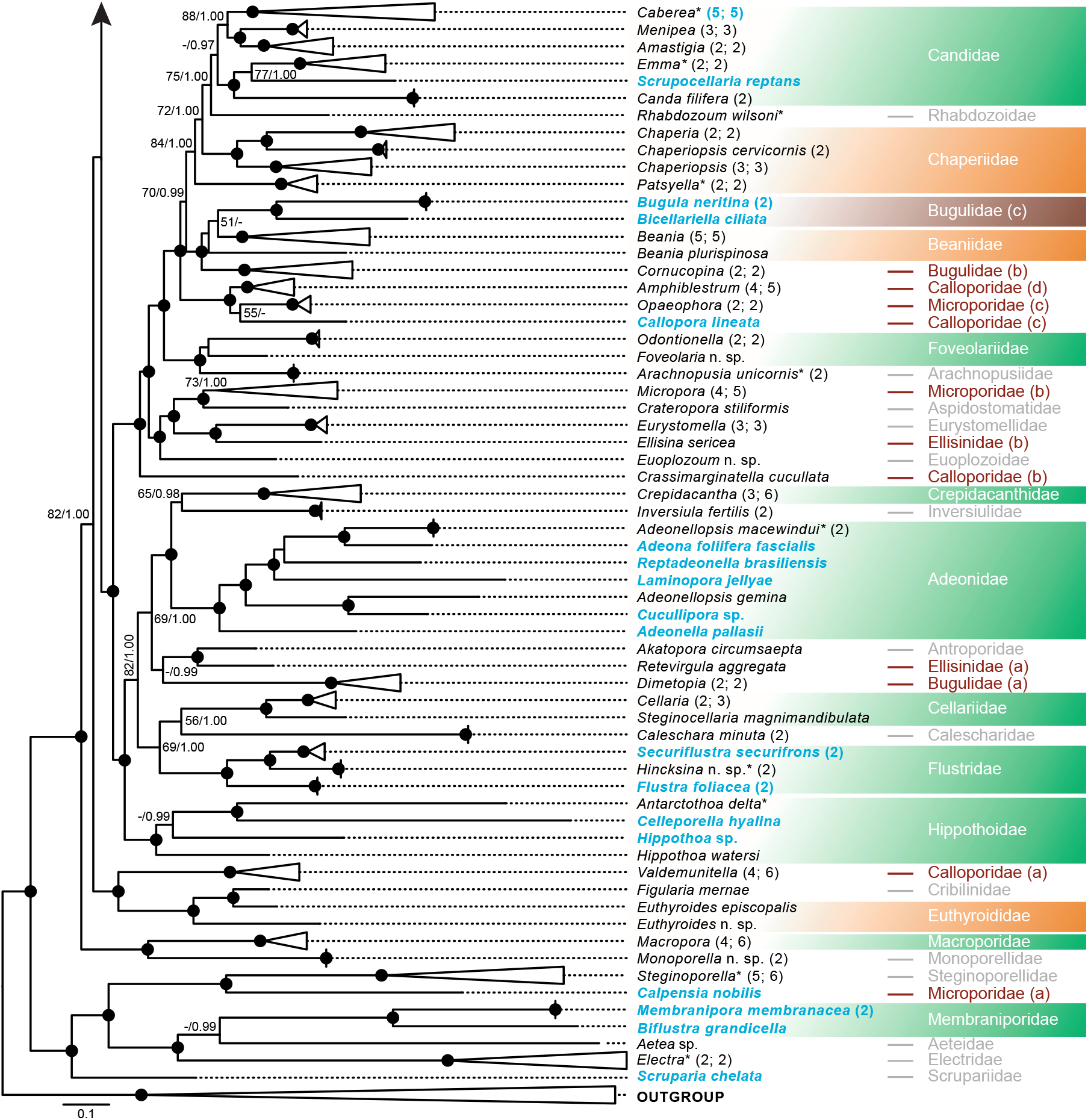
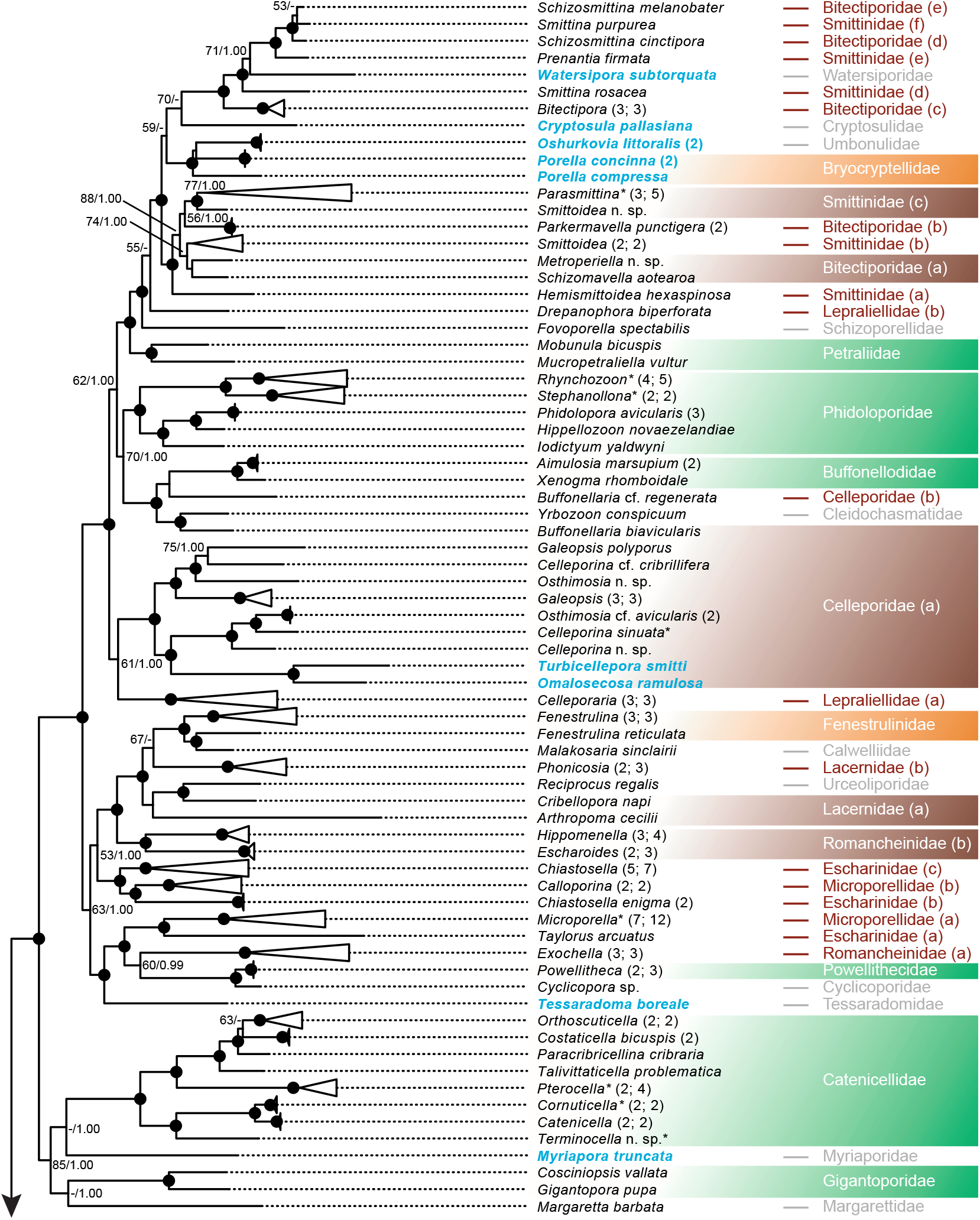
The inferred phylogeny of cheilostomes based on 17 genes including New Zealand and non-New Zealand data. Maximum likelihood topology of 263 cheilostome ingroup taxa and 4 ctenostome outgroup taxa with 9493 nucleotide and amino acid characters inferred using RAxML (100 heuristic searches and bootstrap of 500 pseudoreplicates). The tree branching has been collapsed at the genus level. The numbers on the internal nodes are ML bootstrap values (BS from RAxML) followed by posterior probabilities (PP from MrBayes). Circles indicate >90 BP and 0.99 PP, BS >50 and PP >0.95 are shown in numbers and others left out. Blue text are non-New Zealand taxa, none of which were generated in this study (see Supplementary table S2). Blue numbers dictate a collapsed genus that contains a mix of New Zealand and non-New Zealand taxa. Numbers in parentheses after branches show genus or species number followed by the number of species or colonies within the collapsed branch. * indicates taxa with sequence data generated from other studies but are also from New Zealand (see Supplementary Table S2). A green box highlights a monophyletic family (2 or more genera or in the case of monogeneric families, two or more species), an orange box a paraphyletic family, and a brown box, or brown family names, a polyphyletic family. The letter in brackets behind polyphyletic family names highlights the sub-clade. Grey family names indicate there are limited data to conclude any phylogenetic relationship (ancestry). i.e., families where only a single genus is represented, or monogeneric families where only a single species is represented. The tree is divided into two pages for ease of presentation; a) representing the basal groupings and b) the terminal groupings.

### 3.2. A global cheilostome phylogeny

#### 3.2.1. Broad taxon-sampling

Our inferred “global” cheilostome phylogeny, encompassing 214 species and 120 genera, from 56 families (Fig. 2) of which 229 colonies, 186 species and 96 genera, currently distributed in 48 families, are from New Zealand (Supplementary Figs. S1 and S2). The New Zealand and global trees represent c. 21% described species of cheilostomes from New Zealand and c. 15% of the described cheilostome genera globally, respectively. Both phylogenies (Fig. 2 and Supplementary Fig. S1) are robustly resolved with most branches and relationships receiving either high (>90 bootstrap (BS) / >0.99 Posterior Probability (PP)) or full support (100 BS / 1.00 PP). Our ingroup cheilostome taxa form a fully supported monophyletic clade, when we infer the global tree including a ctenostome outgroup (Fig. 2).

We summarize only general ingroup observations while referring the reader to topological details in Fig. 2 and Supplementary Fig. S3 that are not discussed here or in the Discussion. We also refrain from summarizing results above the family-level for reasons stated in the Discussion.

#### 3.2.2. Family relationships

Several families for which we have four or more genera represented form supported monophylies (Fig. 2), e.g. the fully supported Catenicellidae *(Orthoscuticella, Costaticella, Paracribricellina, Talivittaticella, Pterocella, Catenicella, Cornuticella* and *Terminocella),* Adeonidae *(Adeonellopsis, Adeona, Reptadeonella, Laminopora, Cucullipora, Adeonella),* Flustridae *(Flustra, Hincksina, Securiflustra)* and Hippothoidae *(Celleporella, Hippothoa, Antarctothoa*). Phidoloporidae *(Iodictyum, Hippellozoon, Phidolopora, Stephanollona, Rhynchozoon)* is highly supported while the Candidae *(Menipea, Amastigia, Caberea, Canda*, *Emma*) is less so.

Of the 29 families represented by two or more genera in our phylogeny, only 12 (c. 41%) are monophyletic in our inference (Fig. 2 green boxes). Families such as the Microporidae *(Micropora, Opaeophora, Calpensia),* Calloporidae *(Valdemunitella, Crassimarginatella, Callopora, Amphiblestrum),* Bugulidae *(Dimetopia, Bugula),* Romancheinidae (*Hippomenella*, *Escharoides* and *Exochella*), and Microporellidae (*Microporella*, *Calloporina*), all currently accepted in (Bock, 2020), are recovered as polyphyletic with high support (Fig. 2 brown boxes), while others such as Euthyroididae are paraphyletic (Fig. 2, orange boxes). Monogeneric families (e.g. Crepidacanthidae, Macroporidae and Powellithecidae) recovered as fully supported monophylies comprising multiple species are not considered here.

#### 3.2.3. Genus relationships

In contrast to family-level systematics, the 50 currently morphologically defined genera for which we have two or more representatives in general form monophyletic groupings (e.g. *Parasmittina, Bitectipora, Rhynchozoon, Microporella, Amphiblestrum, Micropora, Steginoporella* and 27 others) with either high or full support. A few genera are non-monophyletic (20% of those for which we have at least two representatives): several are recovered as paraphyletic in our tree *(Chiastosella, Fenestrulina, Smittoidea, Schizosmittina, Chaperiopsis* and *Valdemunitella),* while only a handful are polyphyletic *(Celleporina, Galeopsis* and *Osthimosia*).

Because there are indications that some species are phenotypically highly variable and others have morphologies that are not yet well-understood, we also sequenced multiple colonies of the same species in several cases even though our goal was to sequence one colony of each species. Morphologically identified species match genetic species inferred by phylogenetic inferences in these cases, including *Parasmittina aotea, Parkermavella punctigera, Chiastosella longaevitas*, *C. enigma*, *Microporella agonistes* and *M. intermedia.* For more details, please see individual SEM cards in the Supplementary data file.

#### 3.2.4. Congruent trees and a single incongruent branch

We show the ML and Bayesian trees for both the New Zealand (Supplementary Figs. S1 and S2) and global (Supplementary Figs. S3 and S4) inferences to have more congruent topologies than expected by chance with an *I_cong_* index of 38.72 and 10.05, respectively. The probability that they are topologically unrelated are 0 and 2.43e-124 for the New Zealand and global trees, respectively.

We highlight the incongruent placement of the *Euthyroides*, *Figularia* and *Valdemunitella* clade. Note that this clade is highly supported as a monophyly in both sets of trees, but its placement within the trees is contested; the ML trees, whether based only on the New Zealand taxa or all taxa (Fig. 2, Supplementary Figs. S1 and S3), place this clade in a basal position with an affinity to the *Macropora/Monoporella* grouping. The Bayes trees, however, infer a more derived position. In all instances (ML and Bayes), support for the inferred placement is lacking.

### 3.3. Ancestral state reconstruction and BiSSE

A different rates model for the transition of the anascan to ascophoran state has a less negative log-likelihood (−29.24) than that for an equal rates model (−35.16), suggesting that it describes our ML tree better. Parameter estimates indicate that the ascophoran state never goes to anascan, and anascan state goes to ascophoran at rate of 0.207 (std err 0.0273), in our ML tree. The estimated node states are shown in Fig. 3. Plots of posterior distributions of speciation and extinction rates show there is no detectable difference for either, given the frontal shield trait (Fig. 4). Note that BiSSE is prone to type II errors (Rabosky and Goldberg, 2015) but that we actually cannot reject the null hypothesis here and are hence on safe grounds. Ancestral state reconstruction for the frontal shield states and BiSSE analyses for the alternative Bayesian tree (Supplementary Figs. S5 and S6) are highly comparable with that estimates from the ML tree (Figs. 3 and 4).

**Fig. 3.**
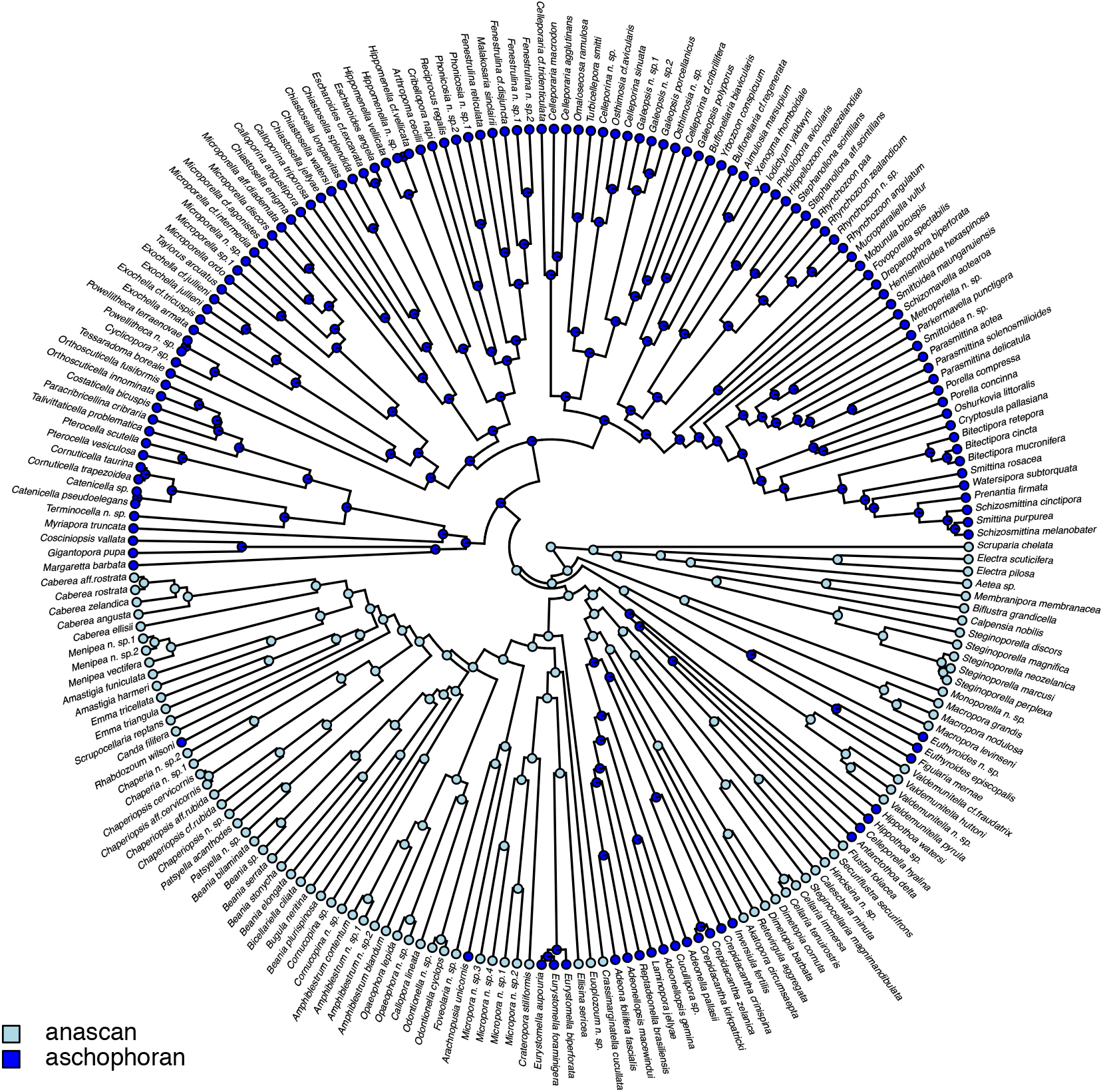
Inferred frontal shield states. Ancestral state reconstruction of anascan (light blue) versus ascophoran (blue) frontal shield states on the inferred global ML tree (see Supplementary Fig. S5 for the version based on the Bayesian tree).

**Fig. 4.**
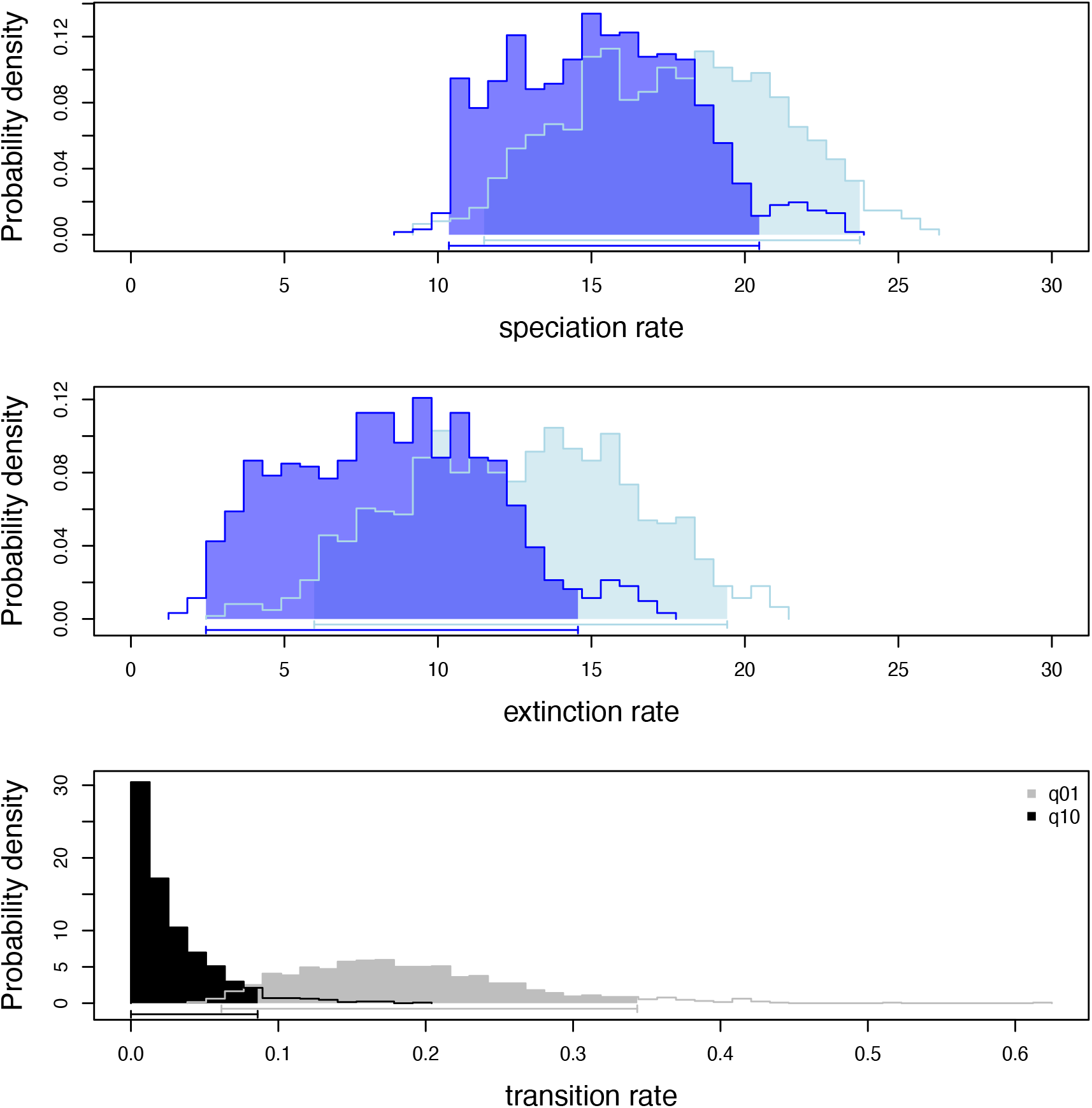
Parameter estimates from BiSSE analyses. Panels show the density of the posterior probabilities of speciation and extinction rates for anascan (light blue) versus ascophoran (blue), estimated from the global ML tree (see Supplementary Fig. S6 for Bayesian interpretation). Transition rates are also shown, where the transition from ascophoran to anascan is skewed towards zero, supporting the ancestral state reconstruction analyses.

## 4. Discussion

It has long been known that molecular and morphological approaches (the latter including fossil taxa) must be simultaneously embraced for robust phylogenetic inferences (Pyron, 2015). In this contribution, we have taken a substantial step in contributing new molecular data and a greatly expanded and robustly supported phylogeny for an understudied but ecologically and evolutionarily important phylum. Although we are interested primarily in New Zealand cheilostome bryozoans for reasons stated in our introduction, we have also now filled out numerous previously unsampled parts of the global cheilostome tree (compare Orr et al., 2020 with Fig. 2).

### 4.1. Higher-level cheilostome systematics needs revision

Cheilostome systematics is in a state of flux as molecular studies, coupled with the introduction of genome-skimming, are starting to take off for this diverse clade (Orr et al., 2019a, b, 2000). In providing a broadly sampled and robustly supported framework to evaluate evolutionary hypotheses we find that less than half of the 29 currently recognized families for which we have multiple genera represented are phylogenetically coherent. Our result emphasizes that much of the current family and higher-level bryozoan systematics, based largely on morphology, is unreliable, and further corroborates previous studies with statistically well-supported, but less broadly sampled, phylogenies (Orr et al., 2019a, b, 2000). One implication of this observation is that higher-level systematics (involving families, superfamilies and suborders) likely require substantial revision. We have hence refrained from detailing the mismatches of higher-level systematics (Bock, 2020) prematurely, but highlight new evolutionary hypotheses that have emerged, that are potentially supportable by morphological traits, given our molecular inferences (Fig. 2, Supplementary Figs. S3 and S4).

Notwithstanding some discrepancies between morphology-based hypotheses (Bock, 2020) and molecular data (this study), there is frequently mutual support. Take, for example, the basal grouping of Scrupariidae (*Scruparia*) as sister taxa to Electridae (*Electra*), Membraniporidae (*Biflustra* and *Membranipora*) and Aeteidae (*Aetea*) plus Steginoporellidae (*Steginoporella*) and Calpensiidae (*Calpensia*) (Fig. 2): all of these families are understood to have diverged prior to the evolution of ovicells (brooding structures) produced from a distal zooid (Ostrovsky, 2013). Our tree now corroborates this hypothesis with full statistical support.

A closely positioned clade formed by *Monoporella* (Monoporellidae) and *Macropora* (Macroporidae) shares the presence of large ooecia that evolved from basally articulated spines or costae (Ostrovsky, 2013) (but see next paragraph). The fully supported Arachnopusiidae (*Arachnopusia*) + Foveolariidae (*Foveolaria* and *Odontionella*) relationship is not indicated in present classification schemes (Bock, 2020), as species of *Arachnopusia* have an ascophoran state, while the Foveolariidae has an anascan state. However, we note that not only is the arachnopusiid frontal shield a straightforward structure to form (unlike other ascophoran structures), but some species in Arachnopusiidae (e.g. *A. gigantea*) are anascan-like, where the frontal shield is practically non-existent (Hayward, 1995).

### 4.2. A need for even broader taxon sampling to fill gaps

Our ML (Fig. 2, Supplementary Fig. S3) and Bayesian (Supplementary Fig. S4) trees are largely in agreement with only one clade demonstrating incongruence. This is the fully supported clade comprising *Valdemunitella* (currently Calloporidae), *Figularia* (currently Cribrilinidae) and *Euthyroides* (currently Euthyroididae). Based only on morphology, we might have hypothesized that the *Valdemunitella* clade (based on 4 species represented by 6 colonies) is closely associated with other representatives of the family Calloporidae (e.g. *Crassimarginatella* or *Callopora*), but neither of our trees inferred this position. Rather, our ML tree places this clade (including *Figularia* and *Euthyroides*, both currently belonging to other families) in a position close to Monoporellidae and Macroporidae (see paragraph above) and our Bayesian tree places it in a more derived position. However, note that nodes subtending this clade in both trees are poorly supported. Rather than speculating on evolutionary and/or morphological arguments for either or both of these placements, we argue, rather that this indicates that there are many crucial unsampled taxa that would potentially allow a more robust placement of this clade, such as other cribrilinids in addition to *Figularia* and other calloporids such as *Cauloramphus* which, similarly to *Valdemunitella,* has spines encircling the frontal uncalcified membrane, forming a costate shield in some species (Dick et al., 2011). In the event, *Valdemunitella, Figularia* and *Euthyroides are* morphologically united, not by a costate shield, but by identical bilobate ooecia with a median suture, and the presence of vicarious avicularia in most of their species.

### 4.3. The evolution of the cheilostome frontal-shield

Historical studies of cheilostome body-wall development and morphology led to the conclusion that ascophoran frontal shields were phylogenetically informative (Banta, 1970; Gordon and Voigt, 1996; Sandberg, 1977). Our results substantiate the observation that characters considered to have deep phylogenetic information such as frontal shields are more evolutionarily labile than previously thought, and sometimes may even be convergent rather than homologous traits (Knight et al., 2011; Orr et al., 2019a). It has already been suggested, for instance, that anascan and ascophoran states, respectively regarded as stemward and crownward, have evolved more than once (Dick et al., 2009; Gordon, 2000; Waeschenbach et al., 2012). We show here that the anascan state is very likely basal in the cheilostome tree and that the change from an anascan to ascophoran state has happened multiple times independently (seven times in Fig. 3), hence likely more times in the history of cheilostome evolution. It is also striking that an ascophoran-state never reverts back to the anascan-state, suggesting that it is evolutionarily unproblematic to evolve a more complex calcified skeleton, but that once this structure is in place, it has not been lost again (Fig. 3). This could be due to genetic or developmental constraints, and/or because the advantages conferred by a calcified frontal shield vastly outweighs its disadvantages. Testing a classic idea that morphological complexity may predict diversification rates (Schopf et al., 1975), we found that the (potentially) more morphologically complex ascophoran-grade cheilostomes have indistinguishable speciation and extinction rates compared with anascan-grade ones.

The frontal shield clearly contains phylogenetic information, but more research is needed to understand when it is informative, and why. As a further example, frontal shields produced by different developmental processes (e.g., lepralioid or umbonuloid (Hayward and Ryland, 1999; Taylor, 2020)) leave such distinct morphological tell-tale signs that it was commonly assumed that members within families constituted only a single type of frontal shield development. Our tree, however, places ascophoran taxa with lepralioid frontal shields (e.g., *Powellitheca/Cyclicopora; Celleporina, Galeopsis, Osthimosia*) and umbonuloid ones (*Exochella*; *Celleporaria*) in the same clades (Fig. 2), as already shown to a lesser extent in earlier extensive studies (Dick et al., 2009; Orr et al., 2019a; Waeschenbach et al., 2012). Yet, at the more derived part of our inferred tree, the structure of the frontal shield seems to be more phylogenetically informative than seemingly distinct features such as the lyrula (Berning et al., 2014). This is an anvil-shaped tooth-like structure projecting from the orifice that functions in water compensation. Specifically, the clade containing *Parasmittina* to *Hemismittoidea* (containing four and three genera of the families Smittinidae and Bitectiporidae respectively) has a non-pseudoporous umbonuloid frontal shield (Gordon, 2000), while the next one containing *Schizosmittina* to *Bitectipora* (containing two smittinid and two bitectiporid genera) has a pseudoporous lepralioid shield. The presence of a lyrula seems haphazard among these genera, where those in the Smittinidae have lyrula and those in the Bitectiporidae have a sinus. Our tree suggests new ways of partitioning some of the families and genera of Smittinoidea, which unexpectedly also includes the genera *Porella* (Bryocryptellidae) and *Oshurkovia* (Umbonulidae). To summarize, it is clear that a much more thorough and systematic investigation of the development and evolution of frontal shields is necessary for a deeper understanding of ascophoran cheilostomes.

### 4.4. Molecules suggest morphological hypotheses and pinpoint research needs

Another example of traits thought to be phylogenetically related and hence informative is the sinus versus the ascopore, pertaining to the ascophoran plumbing system. Because *Microporella, Fenestrulina* and *Calloporina* all have ascopores, they were historically united in the Microporellidae. A previous molecular study has clearly shown that *Fenestrulina* does not belong in the same clade as *Microporella* (Orr et al., 2019b). Here, we give molecular support to the hypothesis that *Calloporina* is not a microporellid and further suggest that *Chiastosella* (having a sinus, currently belonging to the Escharinidae) and *Calloporina* (having a slit-like ascopore) belong in the same clade, a relationship supported also by their shared distinctive ovicell (Brown, 1954; Cook et al., 2018). Supporting the long-held hypothesis that an ascopore should evolve by the cutting-off of a sinus, *Chiastosella* should be basalwards of *Calloporina* (Cook et al., 2018, p. 218). This is supported by our tree, which also suggests that *Chiastosella* may be paraphyletic with respect to *Calloporina*.

In multiple cases, taxa that are considered unique or unusual have placed in phylogenetic positions that suggest hypotheses of their evolutionary relationships based on morphology. For instance, *Rhabdozoum,* currently placed in its own family because of its highly distinctive morphology, is basal to Candidae, suggesting that they are closely related and that Candidae *sensu stricto* may have been derived from a *Rhabdozoum*-like ancestor. In fact, the initial zooid of the colony (ancestrula) of *Rhabdozoum* resembles those in some *Scrupocellaria* species and several of its mature zooidal features such as its ooecia, frontal avicularia and spines are reminiscent of species of *Amastigia* and *Menipea* (all Candidae *s.s.*). *Margaretta,* another rather distinct genus, is in a family with only one other monospecific genus (*Tubucella*). Here, *Margaretta* is inferred to be basal to Catenicellidae, suggesting that Catenicellidae *s.s.* may have been derived from a *Margaretta-like* ancestor, although it has always been thought that catenicellids are derived from cribrimorphs (Gordon, 2000; Gordon and Braga, 1994). Much research is required to unravel the mystery of this grouping, given that they are both so distinctive, sharing apparently only rhizoids, rootlets that attach the colony to the substrate. Note that we infer two distinct clades of Catenicellidae, one represented by *Catenicella* and *Cornuticella*, which are vittate (frontal pore chambers are long and narrow) and the second including *Orthoscuticella* and *Pterocella*, which are foraminate (frontal shield has numerous windows in the gymnocyst). Yet another example is the erect and branching calwelliid *Malakosaria* whose zooidal features resemble *Fenestrulina* (Fenestrulinidae), the genus in which *Malakosaria* nests in our tree.

One taxonomically challenging family deserves special mention. The speciose Celleporidae, with at least 252 described living taxa globally, is mostly characterized by nodular/massive colonies as a result of rapid frontal budding (the building of zooids on top of existing ones). As a consequence, autozooids are somewhat irregularly disposed and difficult to characterize morphologically. These genera are currently distinguished by the morphology of their ooecia (development of endooecium/tabula) and orifices (always sinuate but the sinus varies from a narrow slit to a broad and shallow concavity). Genus-level hypotheses based on these characters are problematic as indicated by our tree, in which *Celleporina*, *Galeopsis* and *Osthimosia* are non-monophyletic. *Buffonellaria* is excluded from the family and allied with Buffonellodidae, whereas *Celleporaria*, historically included in Celleporidae but subsequently split off because of its umbonuloid frontal shield (Harmer, 1957; Cook et al. 2018 p. 182), is reinstated.

### 4.5. Lower-level cheilostome systematics are very robust

Although higher-level systematics are in need of revision, we report that lower-level morphological hypotheses (i.e., species and genera) are very robust, supporting inferences based on common-garden experiments, to put forward the idea that “morphological species” are as good as “genetic species” in cheilostome bryozoans (Jackson and Cheetham, 1990). While Jackson and Cheetham experimented only with a handful of species, we now confirm their hard-earned insight implies that many more species and genera can be treated as distinct evolutionary lineages. This is an important result as many evolutionary and paleontological studies use morphospecies or even morpho-genera as the unit of analyses (Alroy, 2010; Heim et al., 2015). We also note that there are many New Zealand species in our tree that are yet undescribed (c. 20% of those newly sequenced here), indicating that continued exploration in the EEZ of New Zealand is crucial even for such a geographically well-characterized marine clade.

## 5. Conclusions

Our work shows that lower-level taxonomic sampling in phylogenetics is vital for understanding higher-level systematics, especially in an understudied group like cheilostome bryozoans. While we have contributed a substantial number of sequences from diverse species, many more must be included for the phylogenetic inferences and reliable systematic groupings for cheilostomes. By contributing molecular data and robustly supported phylogenetic inferences, we have supplied the basis for evolutionary (including phylogenetic) hypotheses that can be further examined. Once we are confident in the topology of at least parts of the cheilostome tree, we can start asking further questions on evolutionary processes.

### CRediT authorship contribution statement

**LHL** conceived the project, **RJSO** performed the lab, bioinformatic and phylogenetic inference. **LHL** performed the ancestral state and diversification analyses. **MHR** and **EDM** carried out the scanning electron microscopy, **EDM** and **DPG** identified the bryozoans, and **AMS** led the major ship-based collecting expeditions in which **HLM** and **DPG** took part in. **LHL, EDM** and **RJSO** drafted the paper. All coauthors contributed to editing and revisions.

## Acknowledgements

We thank Seabourne Rust for helping to sort and identify samples from the University of Otago, Sadie Mills for sending us samples from NIWA, Carolann Schack for her assistance during our Hawkes Bay field work and Kjetil L. Voje for identifying *Steginoporella* specimens. Specimens provided from the NIWA Invertebrate Collection were collected on numerous surveys including: Biodiversity survey of the western Ross Sea and Balleny Islands (TAN0402) undertaken by NIWA and financed by the former New Zealand Ministry of Fisheries (MFish); Oceans Survey 2020 Southern Colville Ridge (TAN1313) voyage, funded by Land Information New Zealand (LINZ) and GNS Science; Fisheries research trawl surveys conducted by NIWA and funded by Fisheries New Zealand (FNZ); Interdisciplinary New Zealand-Australian “MacRidge 2” research voyage (TAN0803), the biological component of which was part of NIWA’s research project “Seamounts: their importance to fisheries and marine ecosystems” funded by the New Zealand Foundation for Research, Science and Technology (FRST) and CSIRO’s Division of Marine and Atmospheric Research project “Biodiversity Voyages of Discovery” funded by the CSIRO Wealth from Oceans Flagship; Kerry Walton, University of Otago; Seamounts project (TAN0905) undertaken by NIWA and funded by FRST, with complementary funding from MFish; Scientific Observer Program funded by FNZ; Biogenic Habitats on the Continental Shelf project (voyages TAN1105 & TAN1108), funded by New Zealand Ministry for Primary Industry (MPI), FRST, NIWA and LINZ; Ocean Survey 20/20 Bay of Islands Coastal Biodiversity, Sediment and Seabed Habitat Project (TAN0906, KAH0907), funded and owned by LINZ; Ocean Survey 20/20 Mapping the Mineral Resources of the Kermadec Arc Project (TAN1104), funded by LINZ, GNS, NIWA and Woods Hole Oceanographic Institution; Oceans Survey 2020 Reinga (TAN1312) voyage, funded by LINZ and New Zealand Petroleum & Minerals; Impact of resource use on vulnerable deep-sea communities project (TAN1503), funded by the Ministry of Business, Innovation & Employment (MBIE) with support from MPI; Joint Japan-Tonga Trench leg of the Quelle 2013 Expedition (YK13-10), funded by JAMSTEC and supported by NIWA; Food-web dynamics of New Zealand marine ecosystems supported by the New Zealand government under “Coasts & Oceans” core funding from MBIE. We thank Lisbeth Thorsbek for help in the DNA isolation lab and the Norwegian Sequencing Centre (NSC) for library prep and sequencing and acknowledge Saga and Abel high-performance computing server time. This project has received funding from the European Research Council (ERC) under the European Union’s Horizon 2020 research and innovation programme (grant agreement No 724324 to L.H. Liow)

## Supplementary material

**Table S1:** Metadata spreadsheet for samples generated in this study

**Table S2:** Gene bank accessions for all samples used in this study

**Table S3:** Table with genes and inferred characters per taxon

**Figure S1: The inferred phylogeny of New Zealand cheilostomes based on 17 genes.** Maximum likelihood topology of 229 New Zealand cheilostome taxa with 9604 nucleotide and amino acid characters inferred using RAxML.

**Figure S2: The inferred phylogeny of New Zealand cheilostomes based on 17 genes.** Bayesian topology of 229 New Zealand cheilostome taxa with 9604 nucleotide and amino acid characters inferred using MrBayes.

**Figure S3: The inferred phylogeny of cheilostomes based on 17 genes including New Zealand and non-New Zealand data.** Maximum likelihood topology of 263 cheilostome ingroup taxa and 4 ctenostome outgroup taxa with 9493 nucleotide and amino acid characters inferred using RAxML.

**Figure S4: The inferred phylogeny of cheilostomes based on 17 genes including New Zealand and non-New Zealand data.** Bayesian topology of 263 cheilostome ingroup taxa and 4 ctenostome outgroup taxa with 9493 nucleotide and amino acid characters inferred using MrBayes.

**Figure S5: Inferred frontal shield states (Bayesian topology).** Ancestral state reconstruction of anascan (light blue) versus ascophoran (blue) frontal shield states on the inferred global Bayesian tree (see main text Fig. 3 for ML interpretation).

**Figure S6: Parameter estimates from BiSSE analyses (Bayesian topology). Panels** show the density of the posterior probabilities of speciation and extinction rates for anascan (light blue) versus ascophoran (blue), estimated from the global Bayesian tree (see main text Fig. 4 for version from the ML tree). Transition rates are also shown, where the transition from ascophoran to anascan is skewed towards zero, supporting the ancestral state reconstruction analyses.

**Supplementary PDF file:** SEM cards with taxonomic and collection information

## Notes

https://doi.org/10.5061/dryad.7pvmcvdrs

### Competing Interest Statement

The authors have declared no competing interest.

