## Supplementary figures and images for "A broadly resolved molecular phylogeny of New Zealand cheilostome bryozoans as a framework for hypotheses of morphological evolution"

### Fig. S1

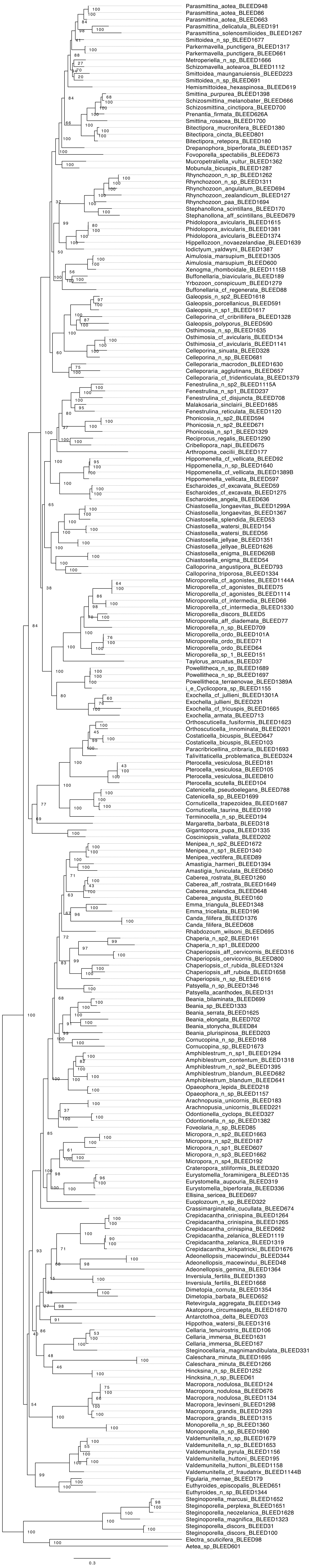

### Fig. S2

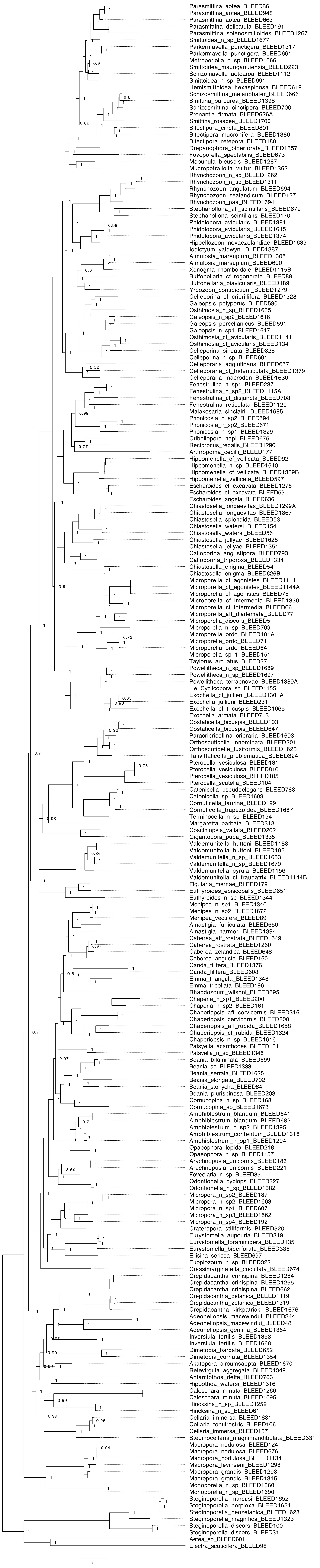

### Fig. S3

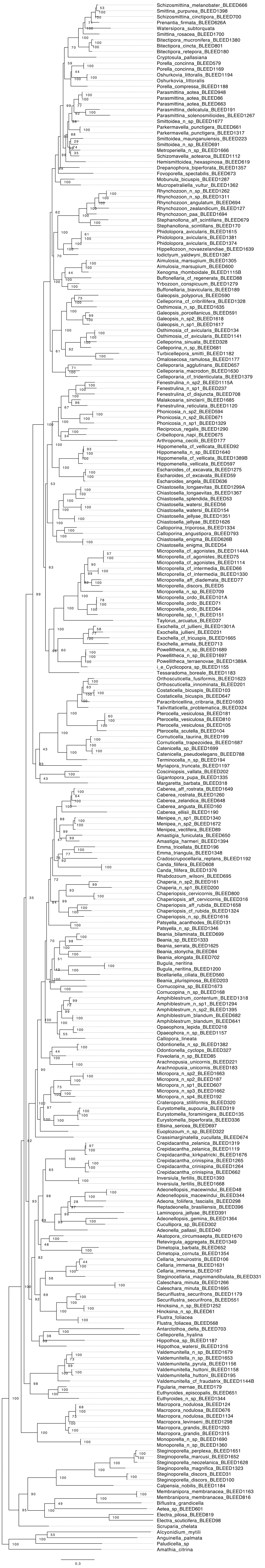

### Fig. S4

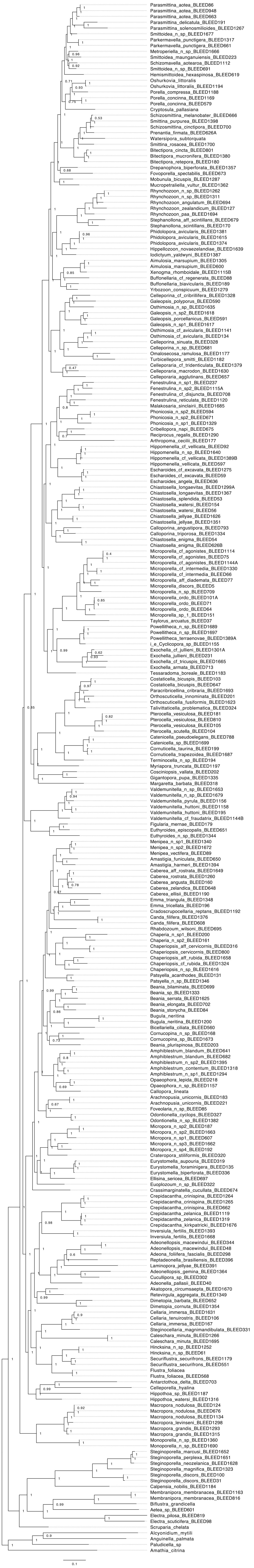

### Fig. S5

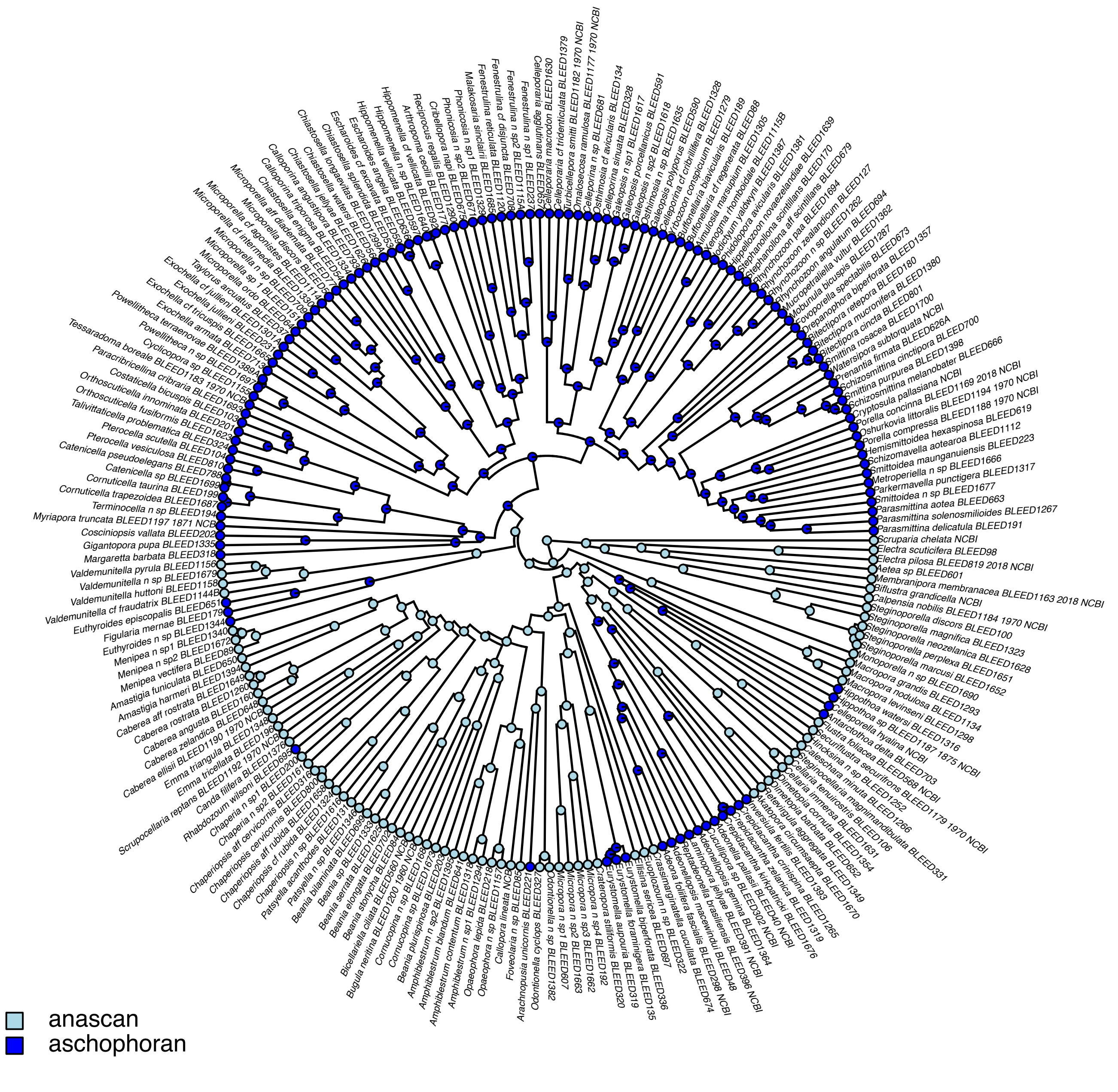

### Fig. S6

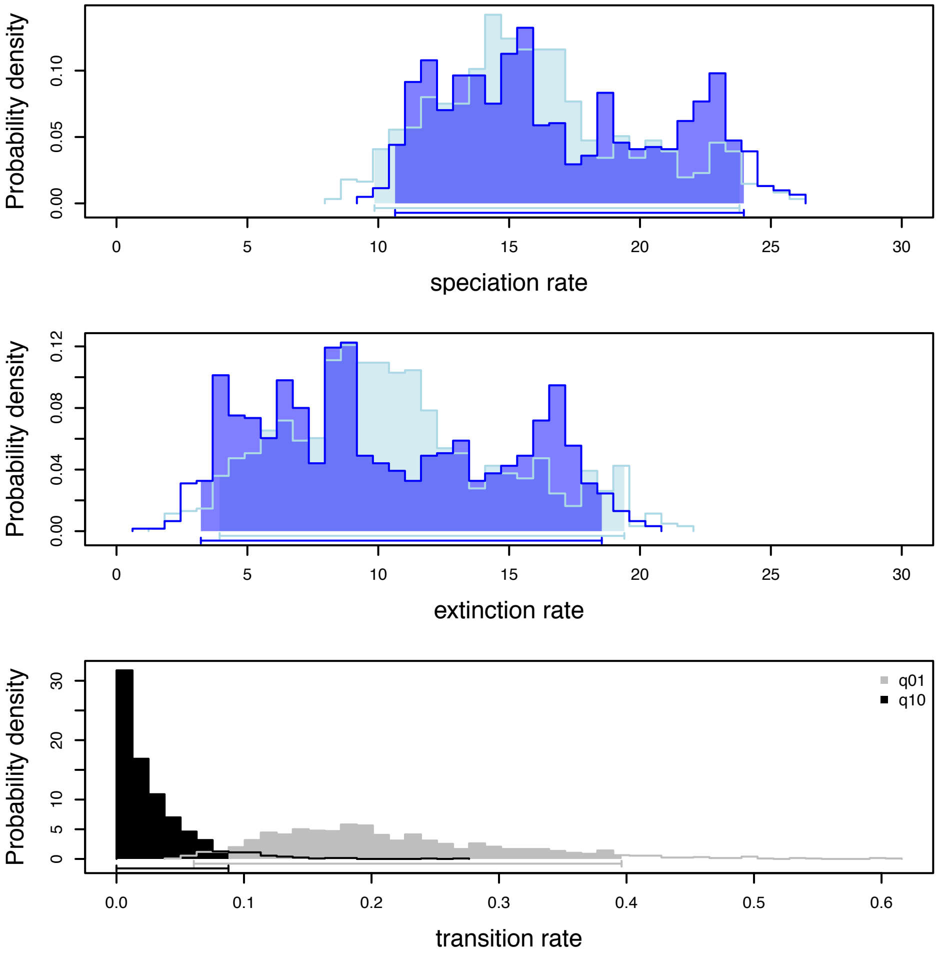
