## Supplementary data file for "A broadly resolved molecular phylogeny of New Zealand cheilostome bryozoans as a framework for hypotheses of morphological evolution"

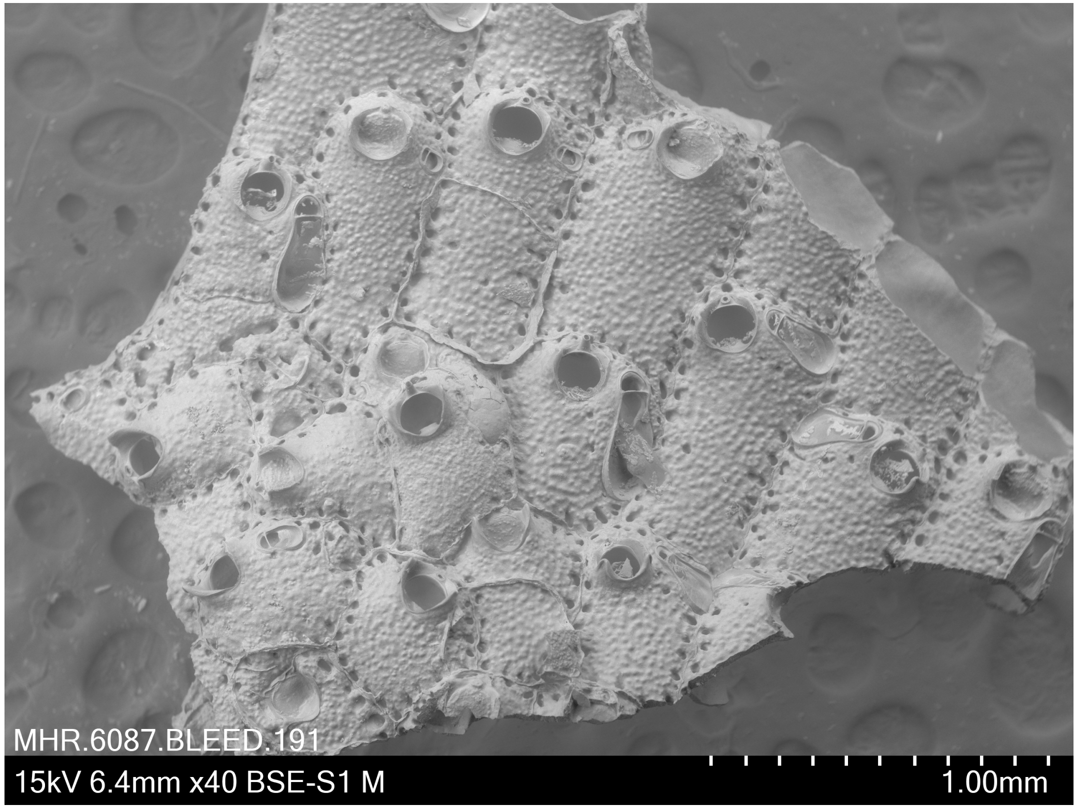


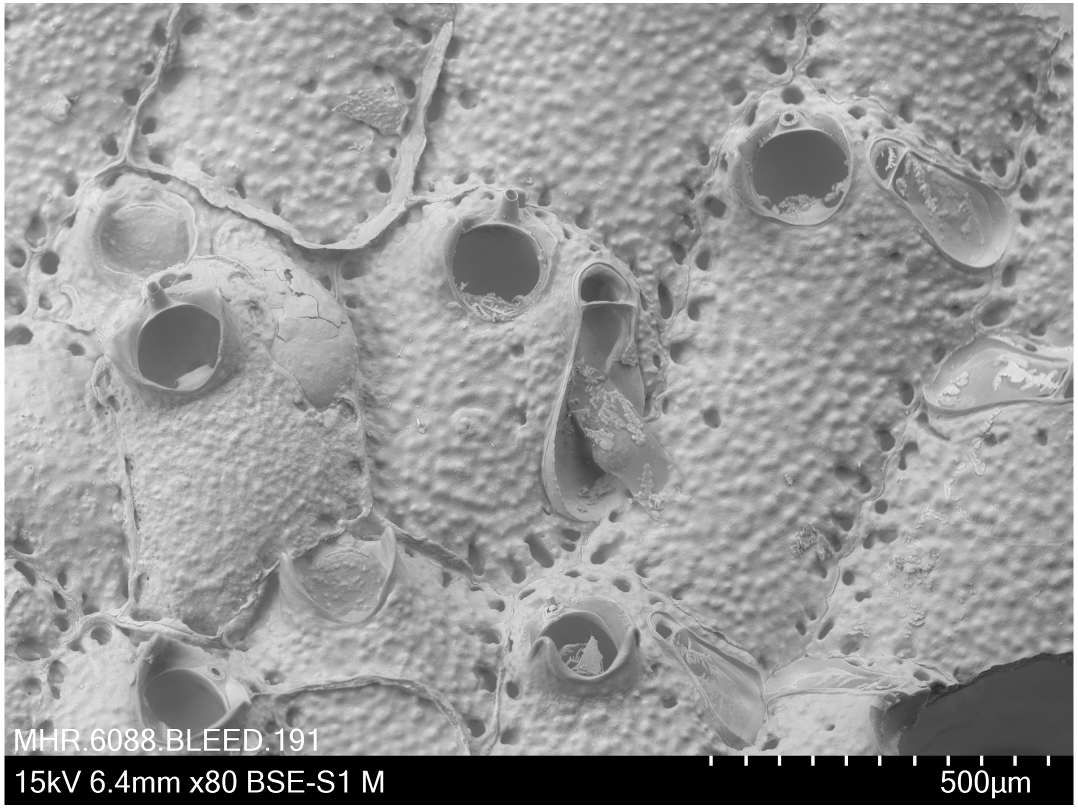


***Parasmittina delicatula*** **BLEED 191**
NIWA 73196 TAN1105/42
Coll: 28.03.2011
Lat: -33.9655
Long: 171.7723333
Depth: 96 m

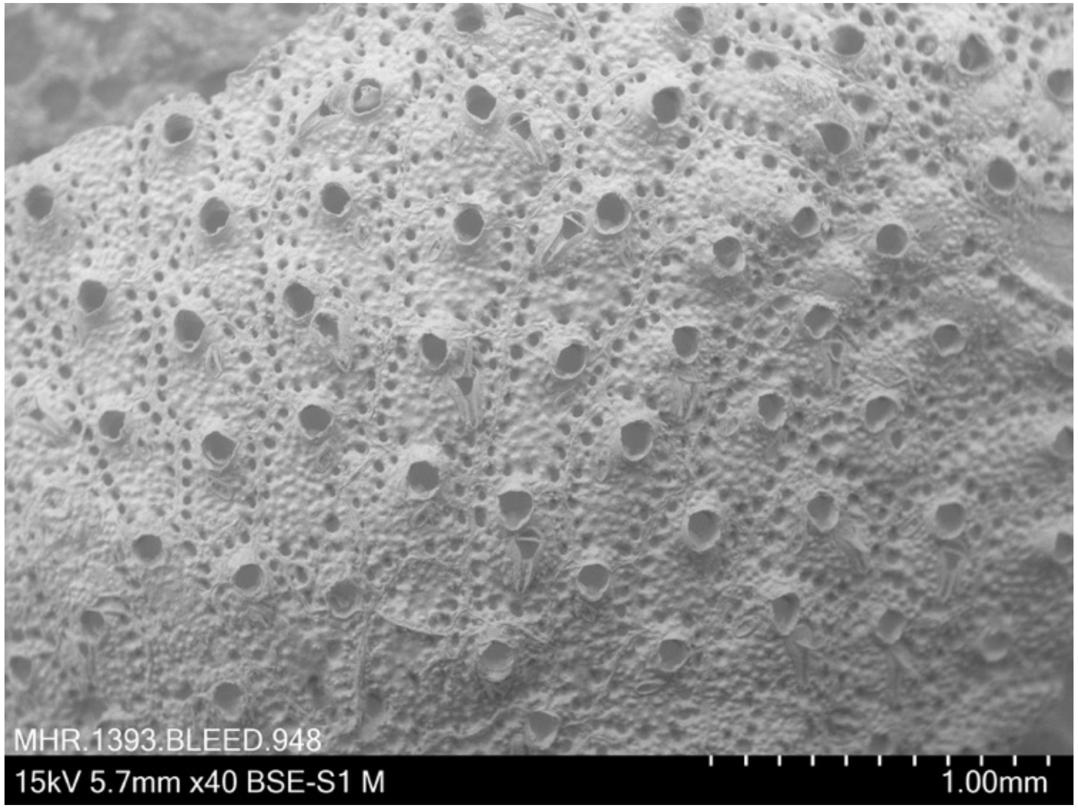


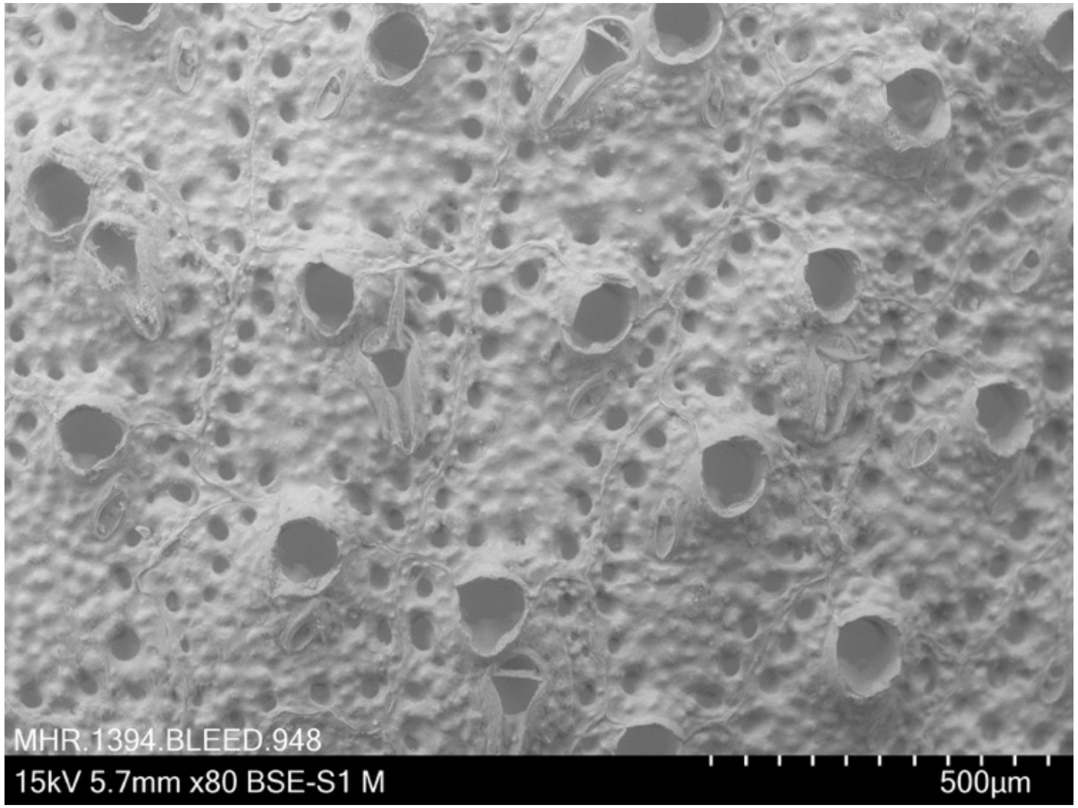


***Parasmittina aotea* BLEED 948**
Coll: 30.01.2008 (A. M. Smith)
Lat: -47.86
Long: 166.87
Depth: 157 m

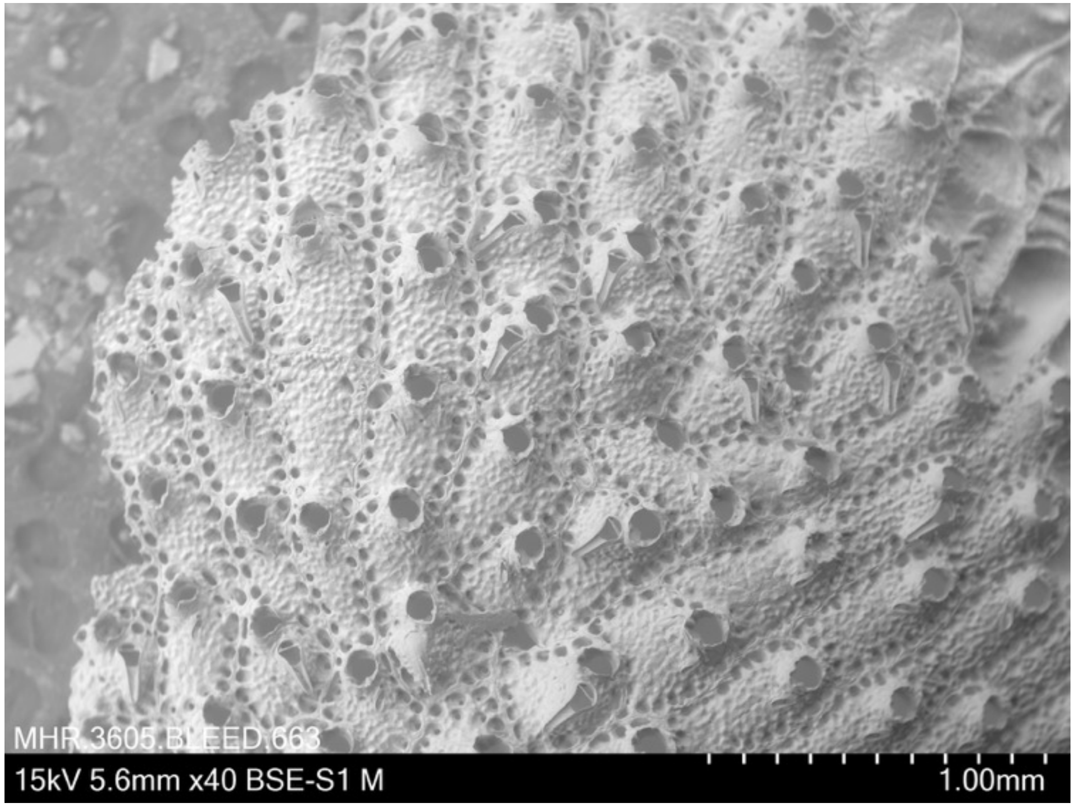


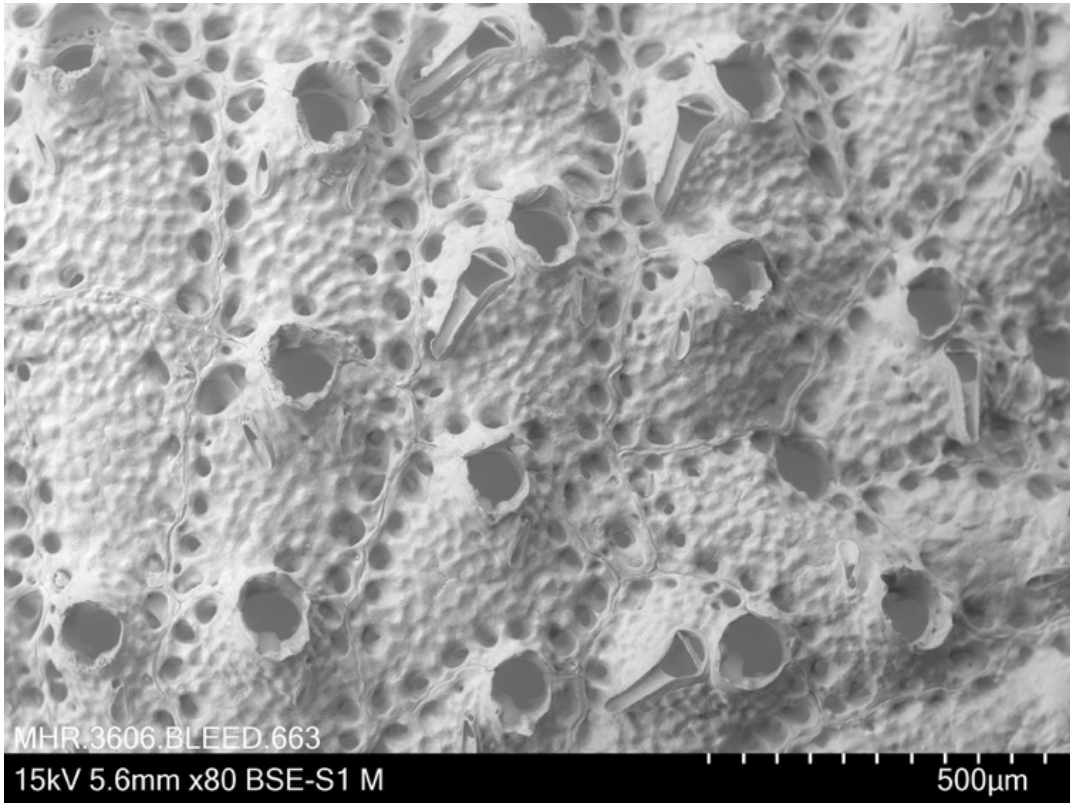


***Parasmittina aotea* BLEED 663**
Coll: 8.4.2018 (A. M. Smith, D. P. Gordon, H. Mello)
Lat: -47.13722
Long: 168.12583
Depth: 80-85 m

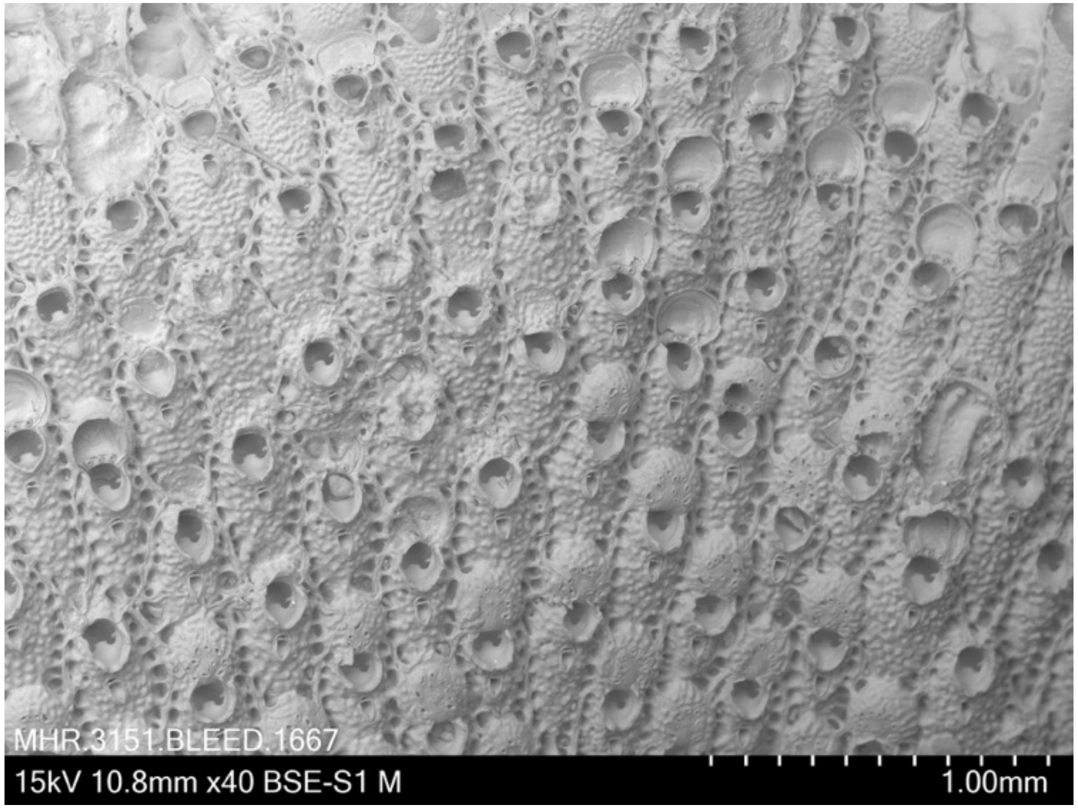

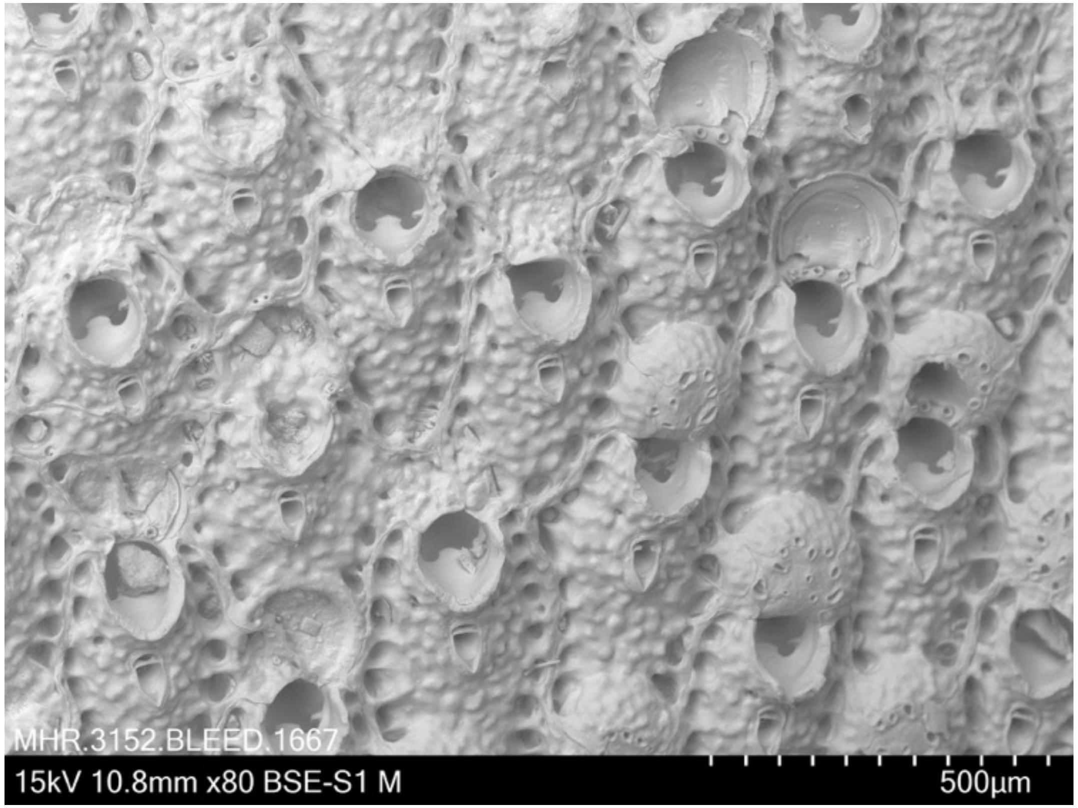


***Smittoidea* n. sp. BLEED 1677**

Coll: 3.18.2019 (A. M. Smith, D. P. Gordon)

Lat: -34.380000000000003

Long: 172.69

Depth 52.5 - 56.5m

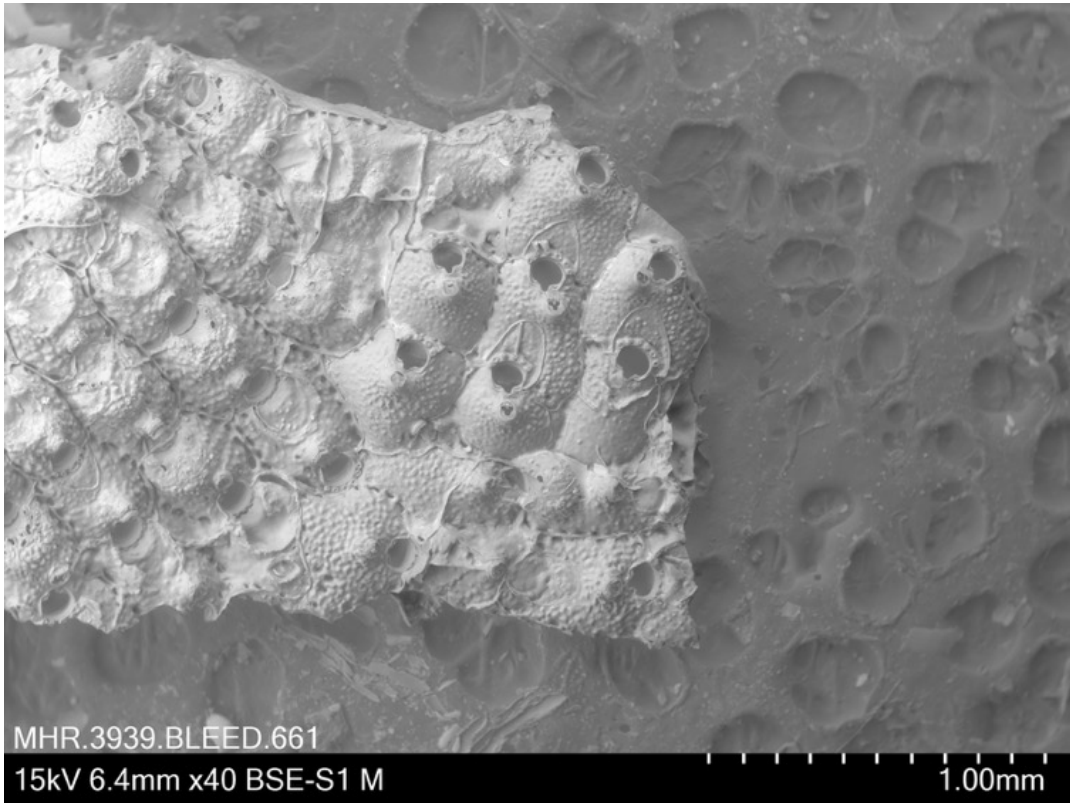


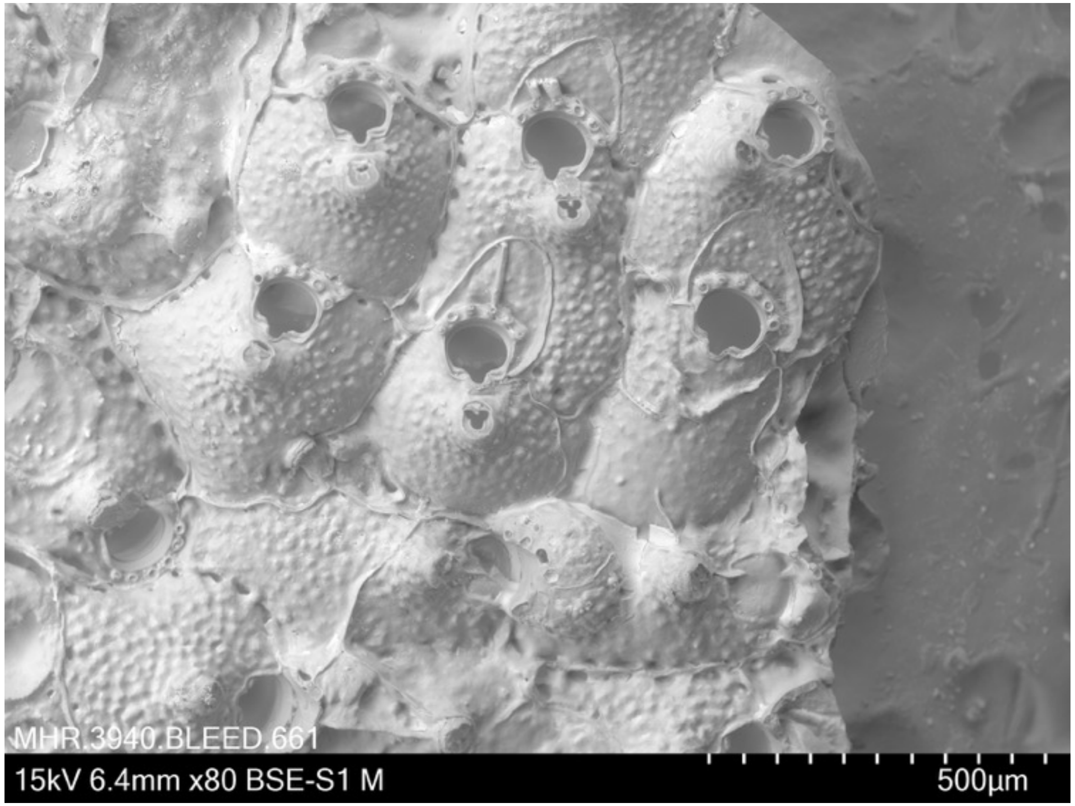


***Parkermavella punctigera*** **BLEED 661**
Coll: 8.4.2018 (A. M. Smith, D. P. Gordon, H. Mello)
Lat: -47.137222
Long: 168.12372
Depth: 80-85 m

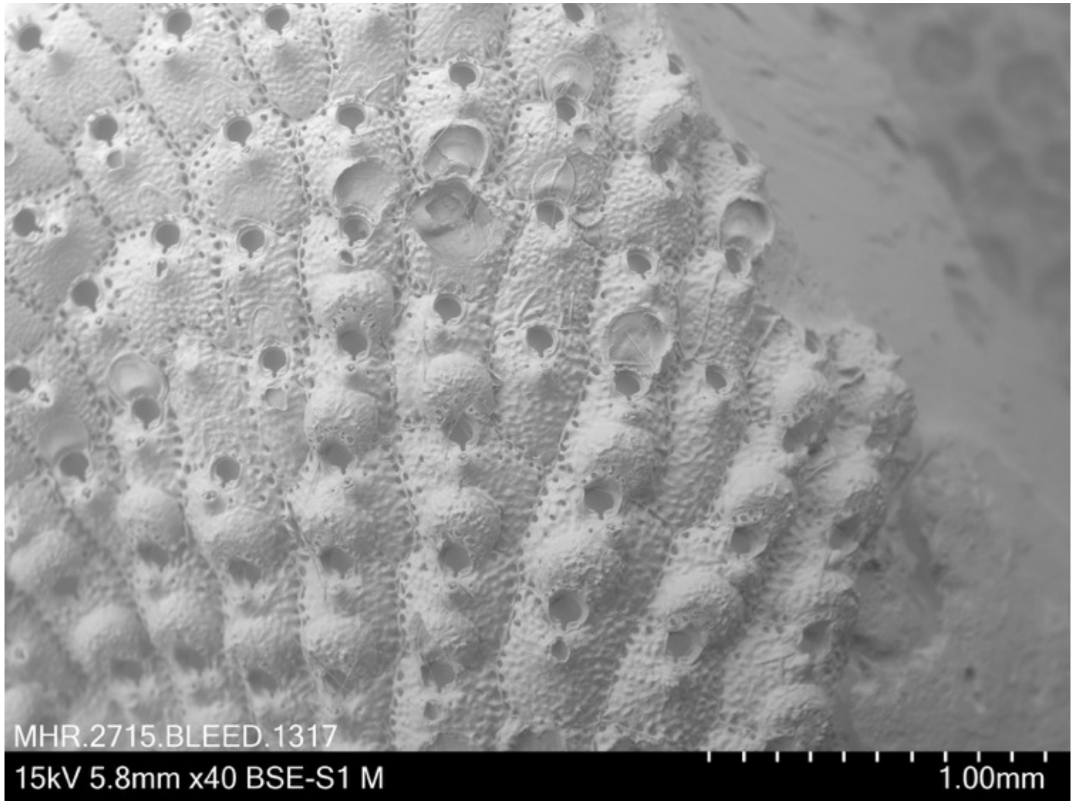


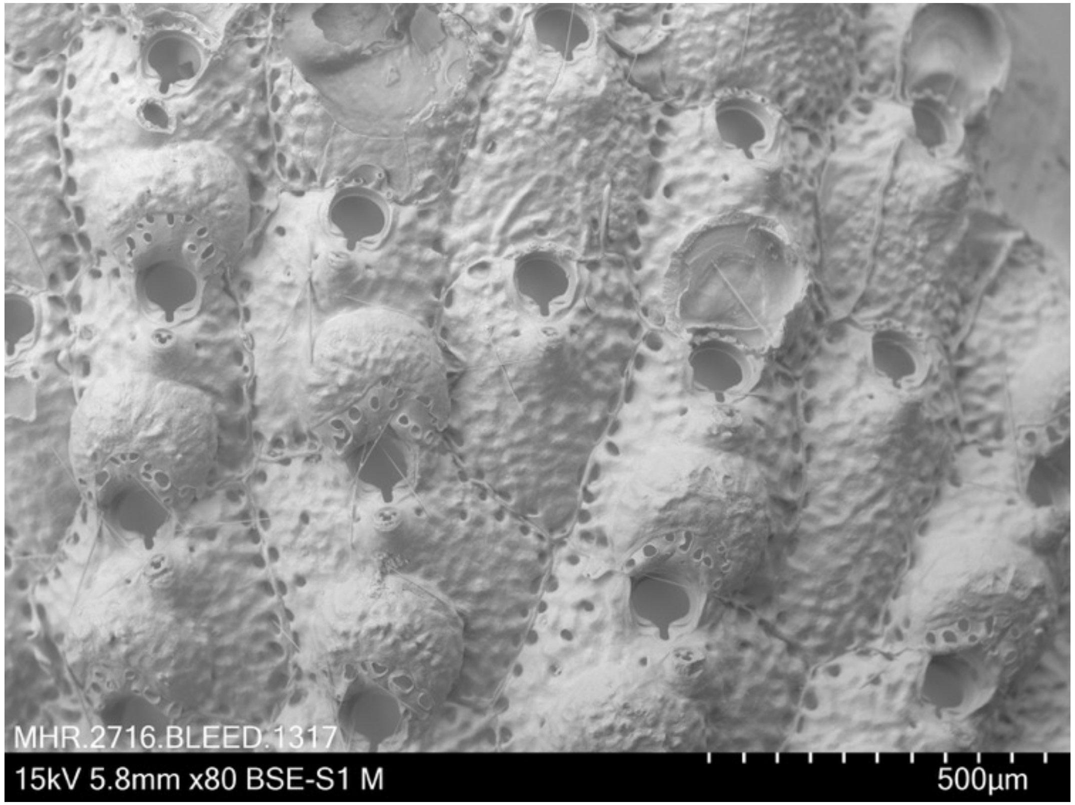


***Parkermavella punctigera*** **BLEED 1317**
Coll: 15.3.2019 (A. M. Smith; D. P. Gordon)
Lat: -36.46
Long: 174.9
Depth: 37 m

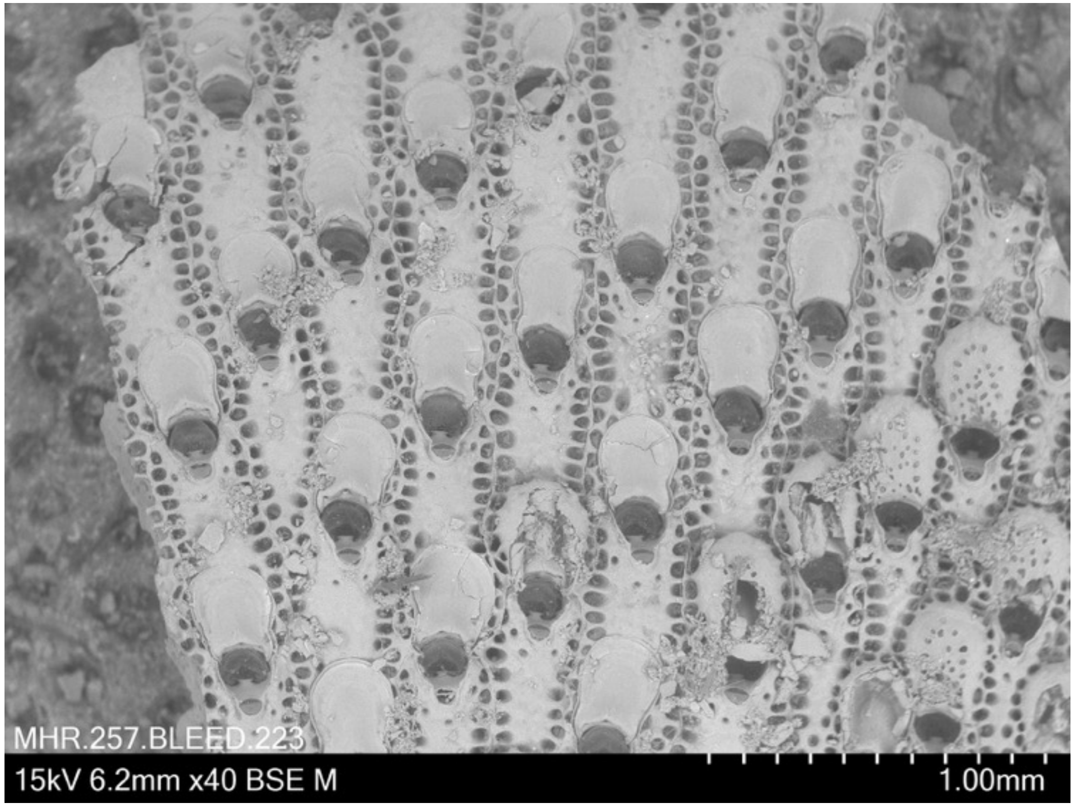


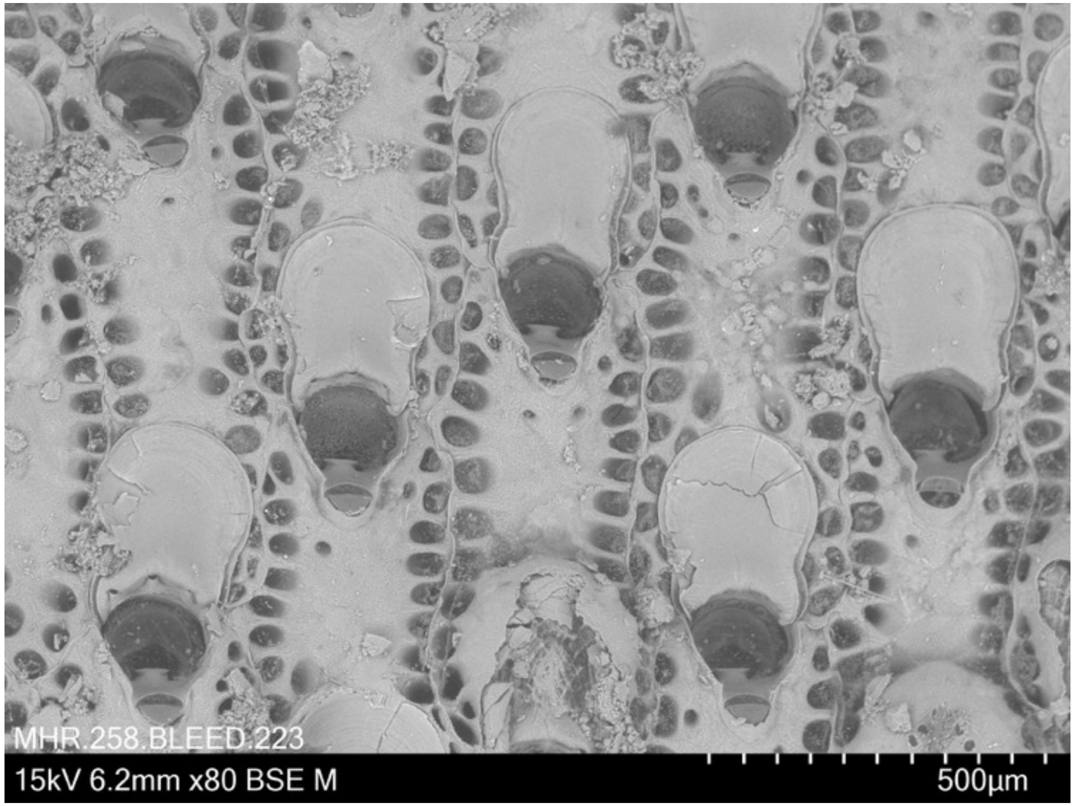


***Smittoidea maunganuiensis* BLEED 223**
Coll: 18.01.2009 (A. M. Smith)
Lat: -46.17
Long: 166.23
Depth: 136 m

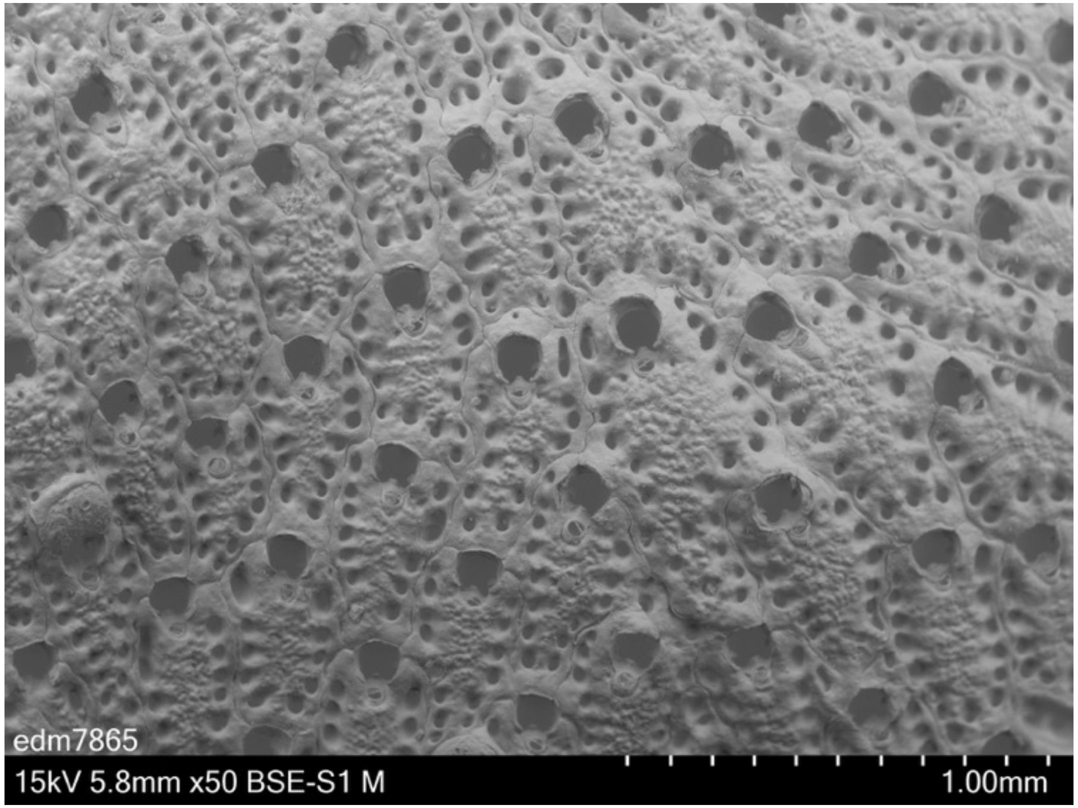


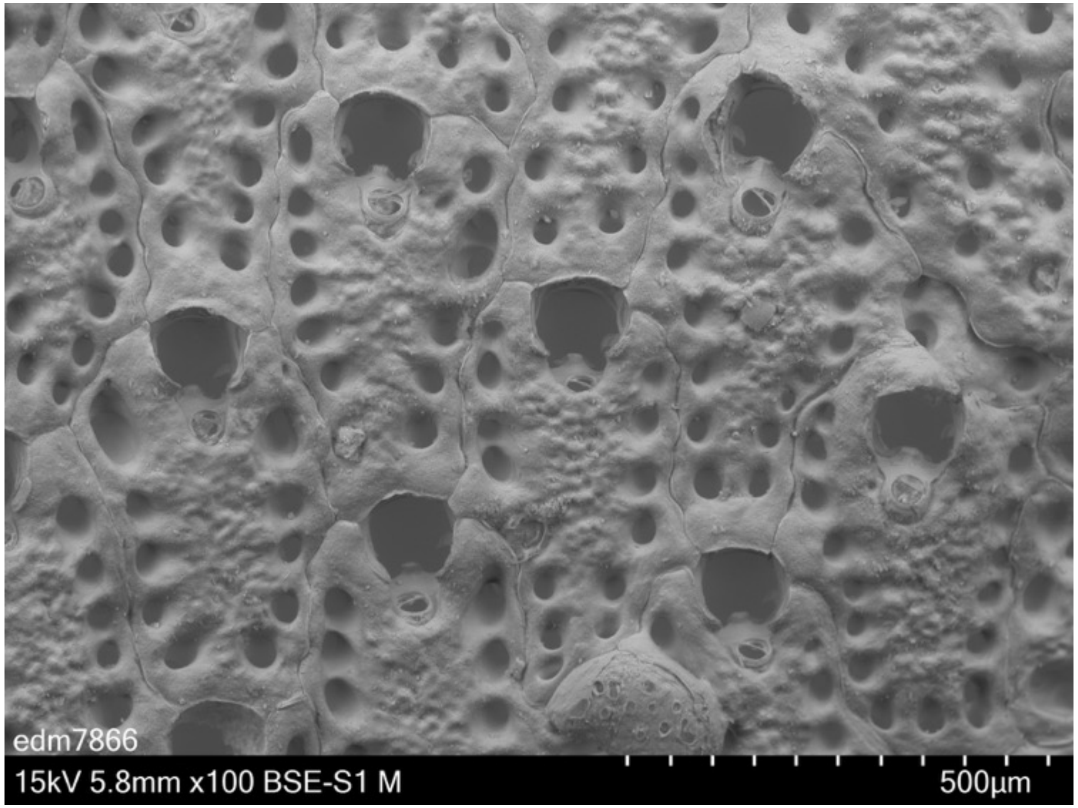


***Smittoidea* n. sp. BLEED 691**
Coll: 8.4.2018 (A. M. Smith, D. P. Gordon, H. Mello)
Lat: -47.136111
Long: 169.18861
Depth: 77 m

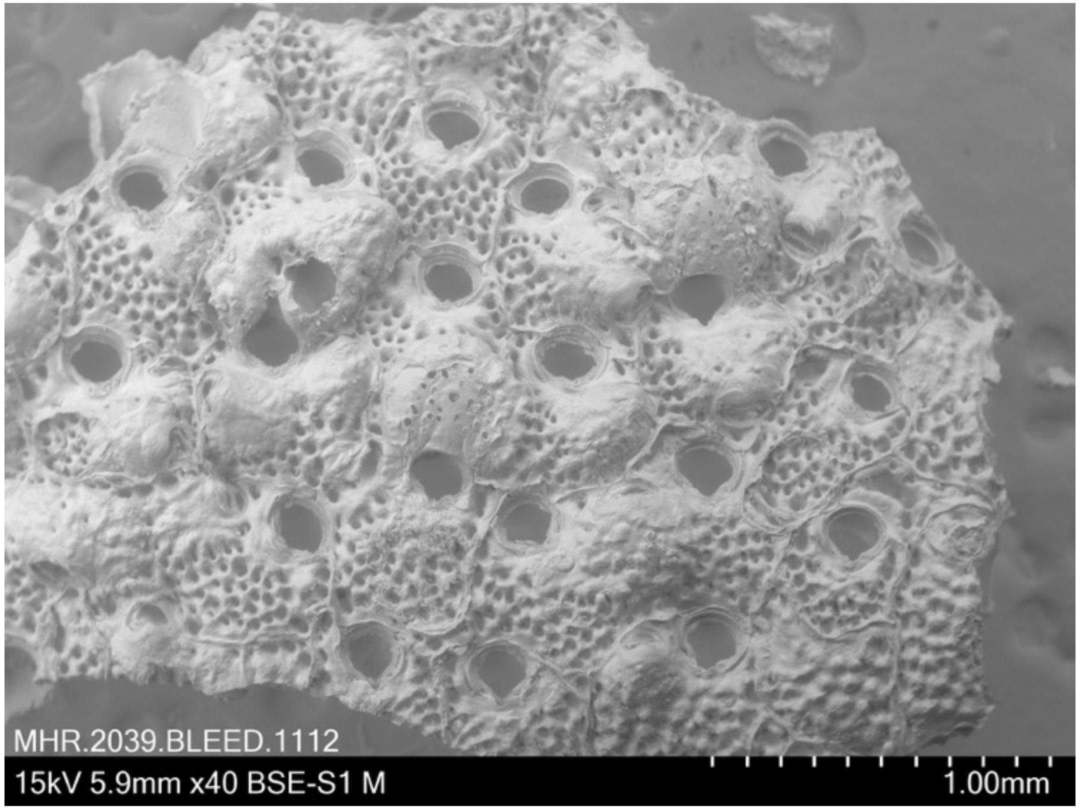


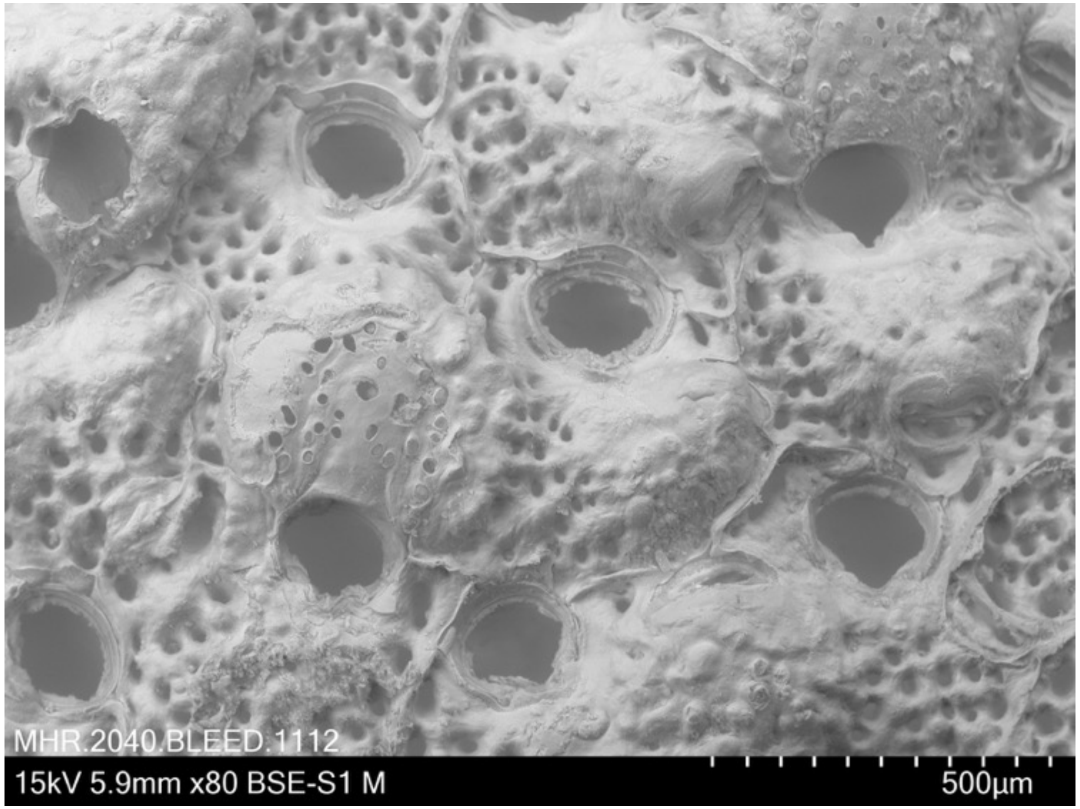


***Schizomavella aotearoa* BLEED 1112**
Coll: 10.11.18 (A. M. Smith)
Lat: -45.822221999999996
Long: 170.880833
Depth: 80 m

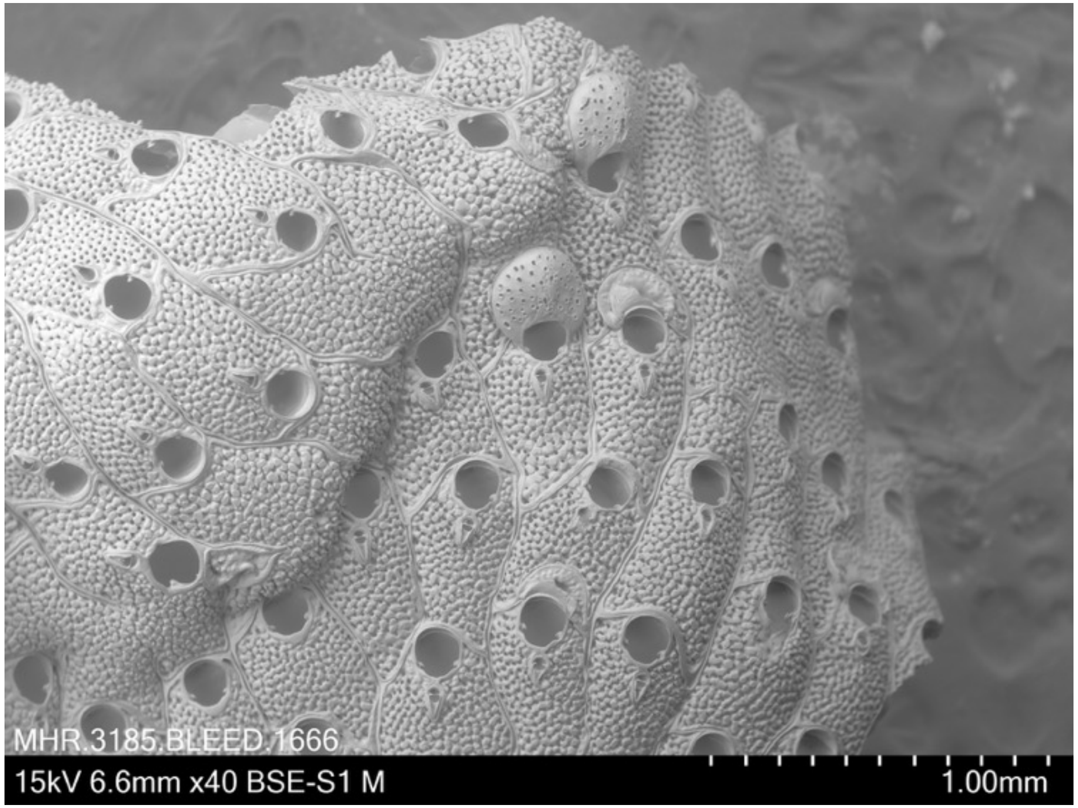


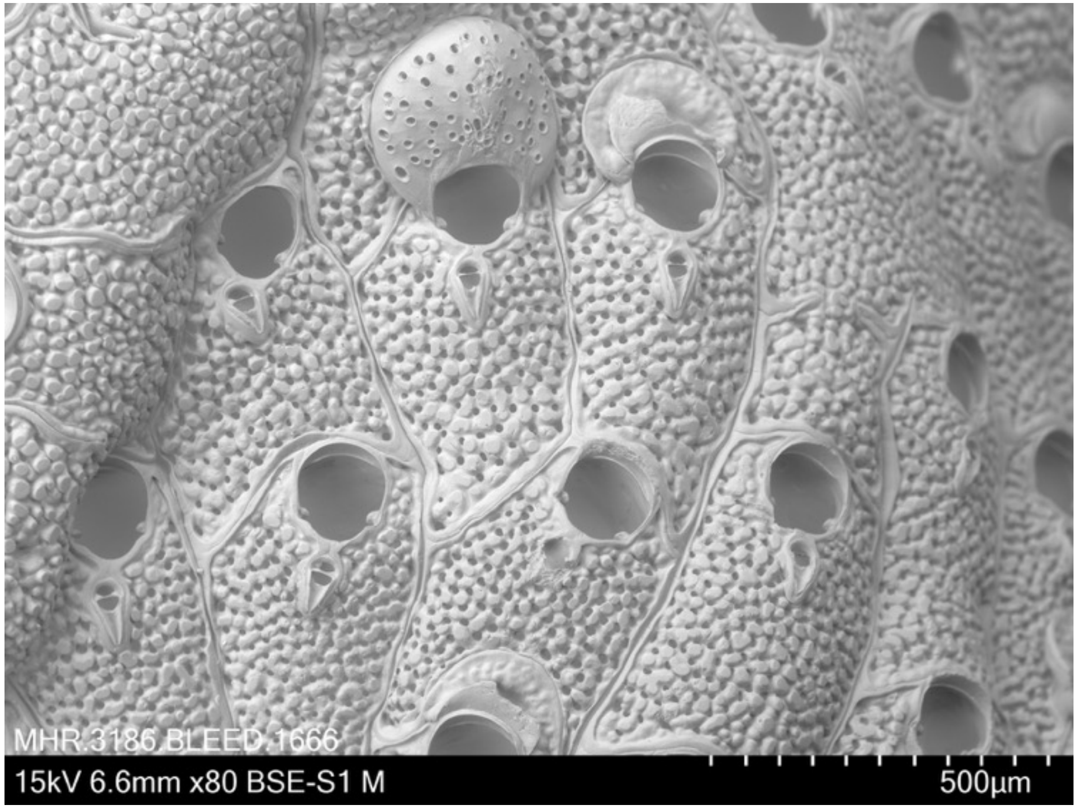


***Metroperiella* n. sp. BLEED 1666**
Coll: 18.3.2019 (A. M. Smith, D.P. Gordon)
Lat: -34.380000000000003
Long: 172.69
Depth: 52.5-56.5m

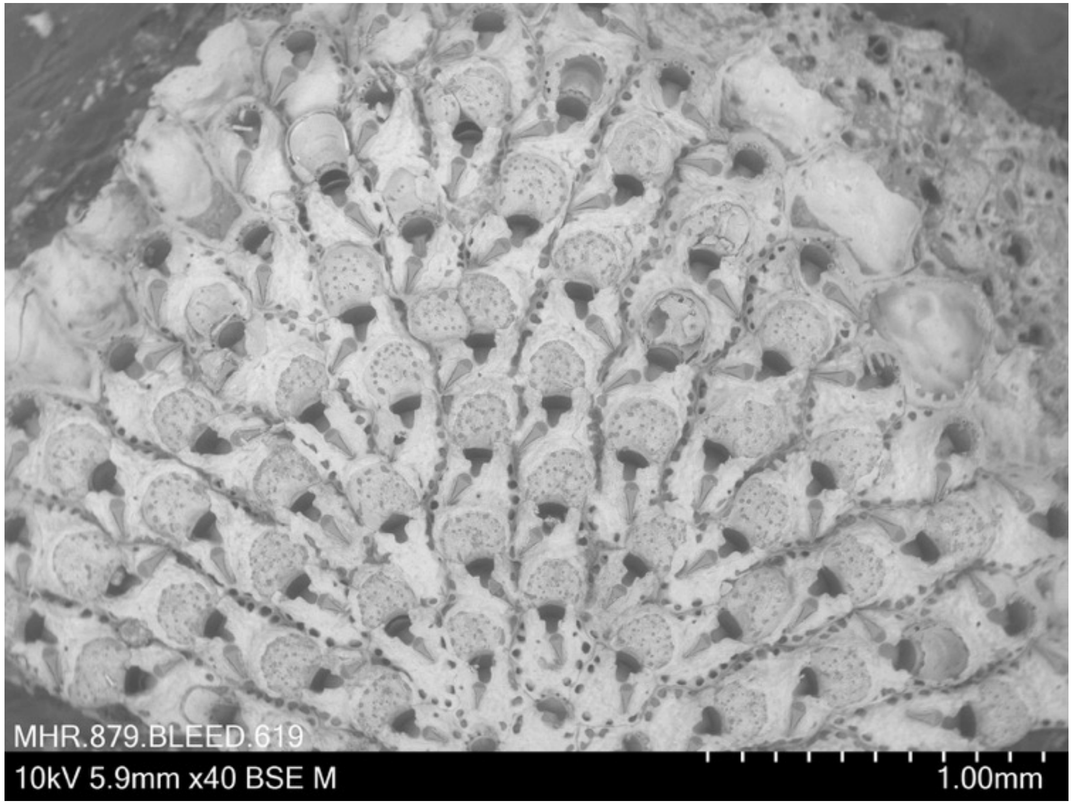


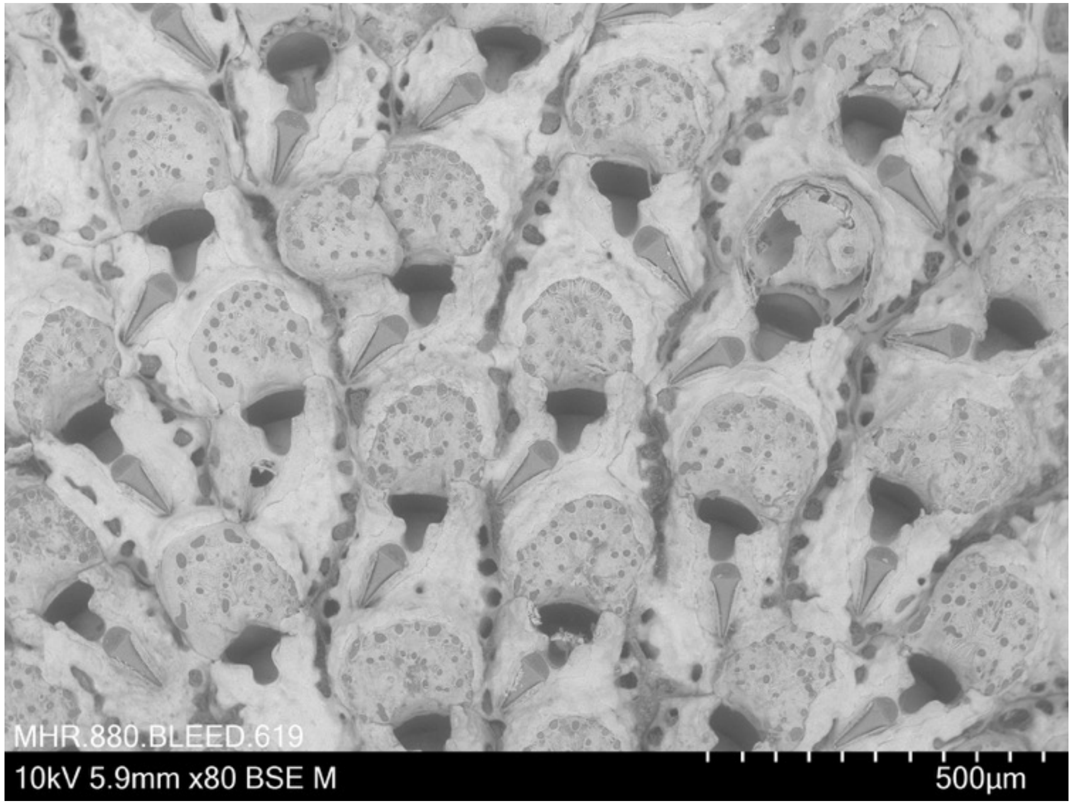


***Hemismittoidea hexaspinosa*** **BLEED 619**
Coll: 10.4.2018 (A. M.  Smith, D. P. Gordon, H. Mello)
Lat: -46.9450
Long: 168.1583
Depth: 42 m

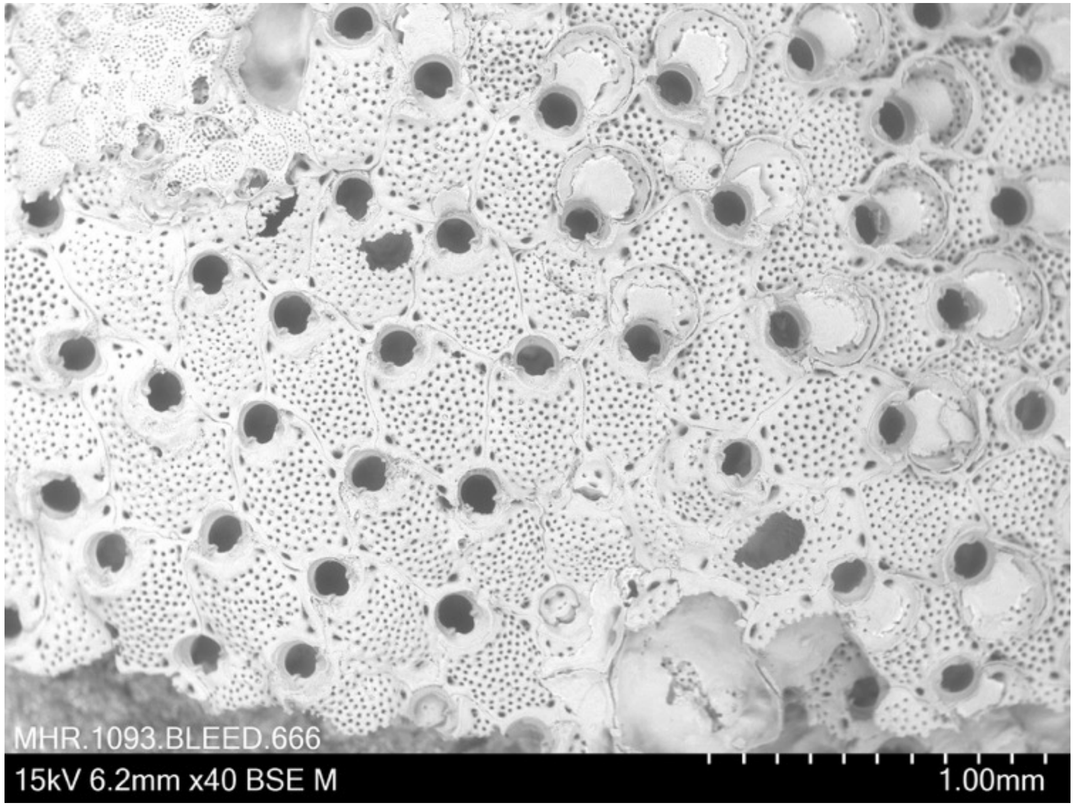


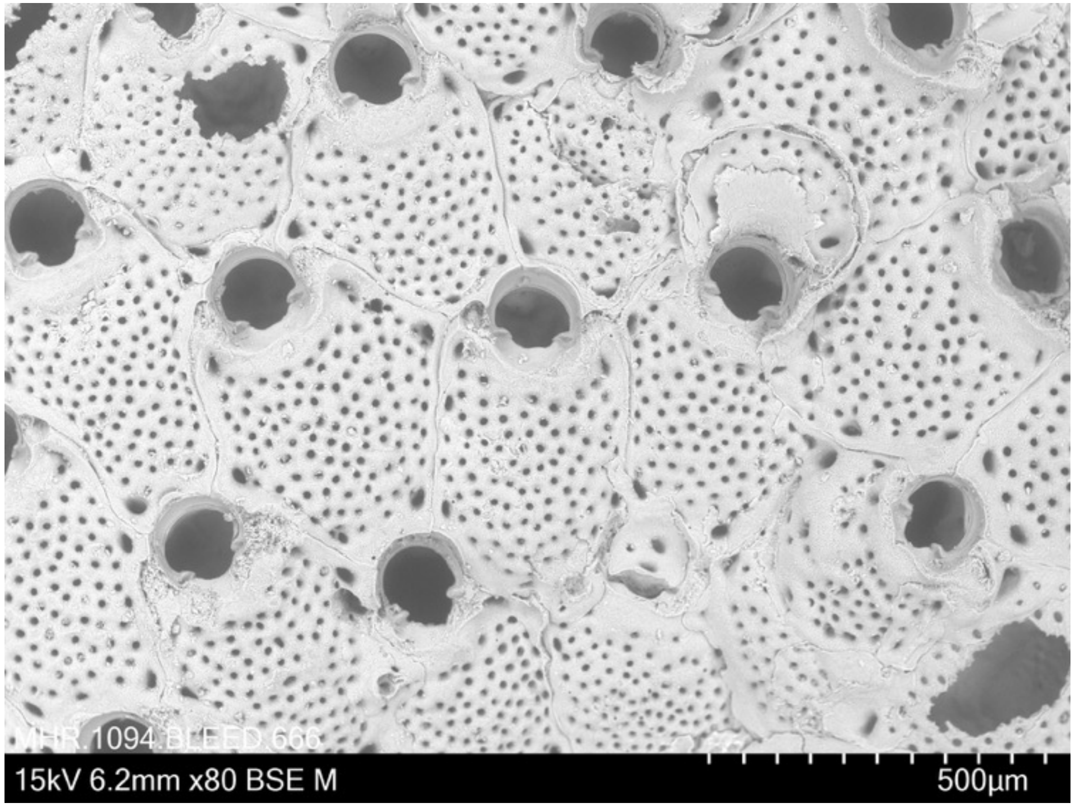


***Schizosmittina melanobater*** **BLEED 666**
Coll: 8.4.2018 (A. M. Smith, D. P. Gordon, H. Mello)
Lat: -47.1372
Long: 168.1258
Depth: 80-85 m

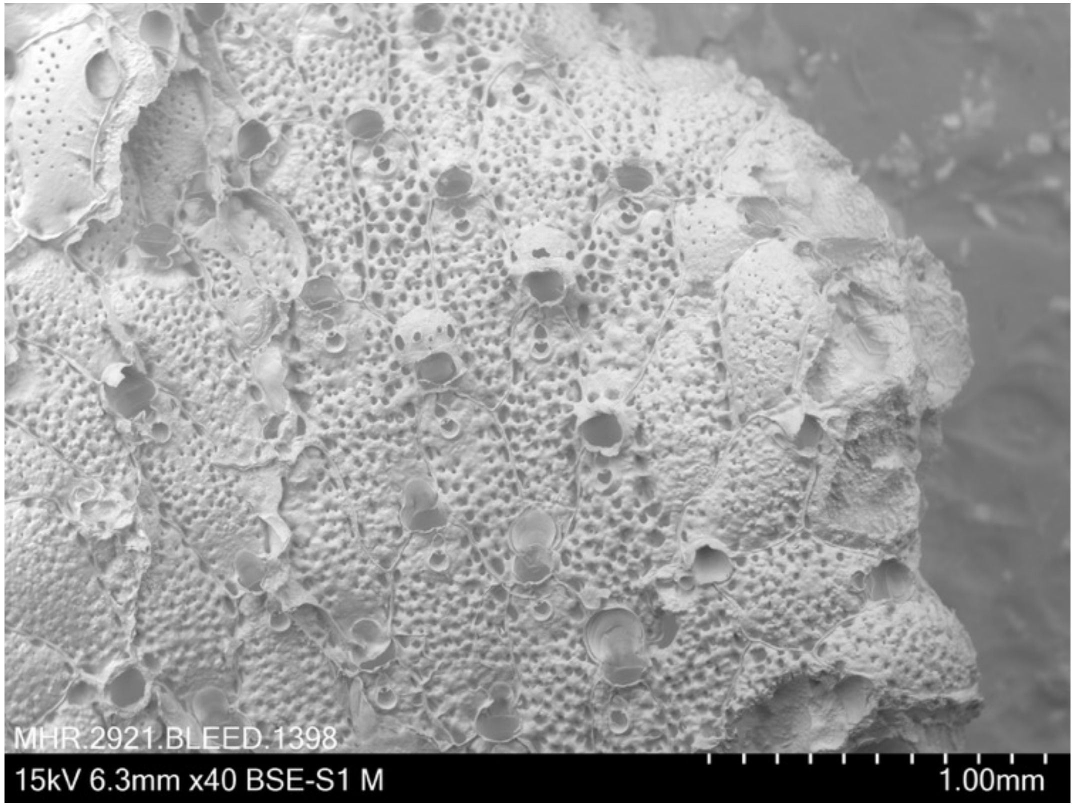


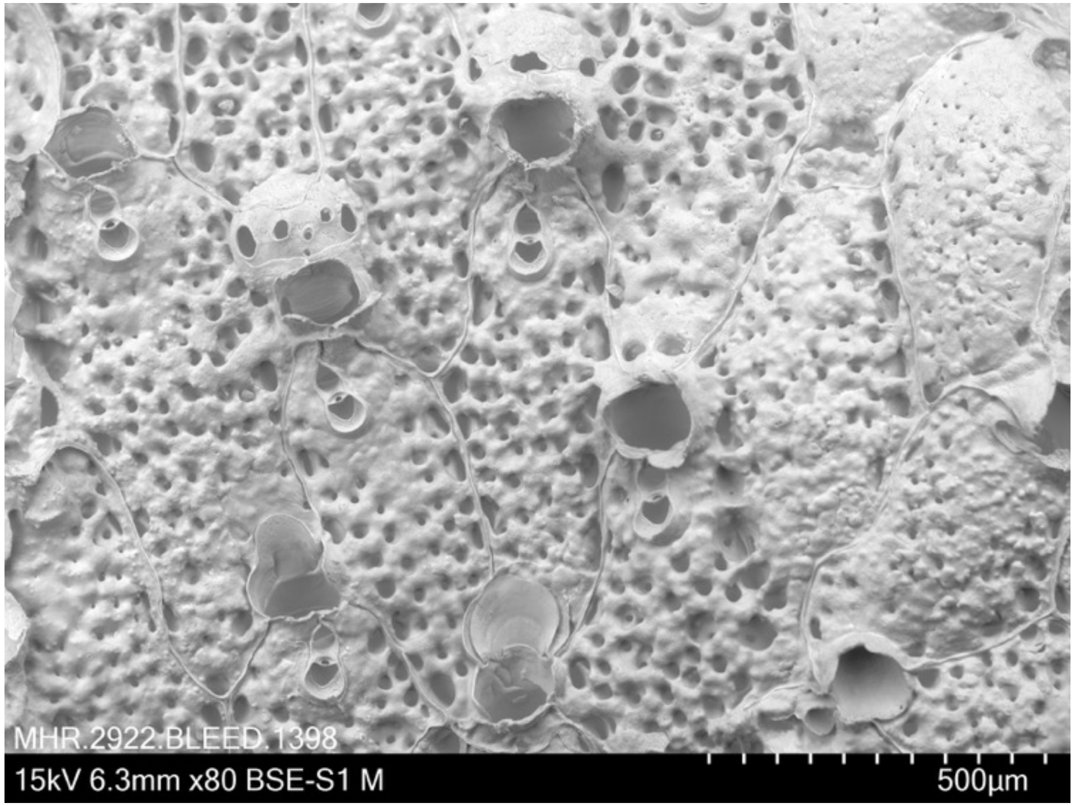


***Smittina purpurea* BLEED 1398**
Coll: 17.3.2019 (A. M. Smith, D. P. Gordon)
Lat: -34.159999999999997
Long: 172.09
Depth: 82.9 m

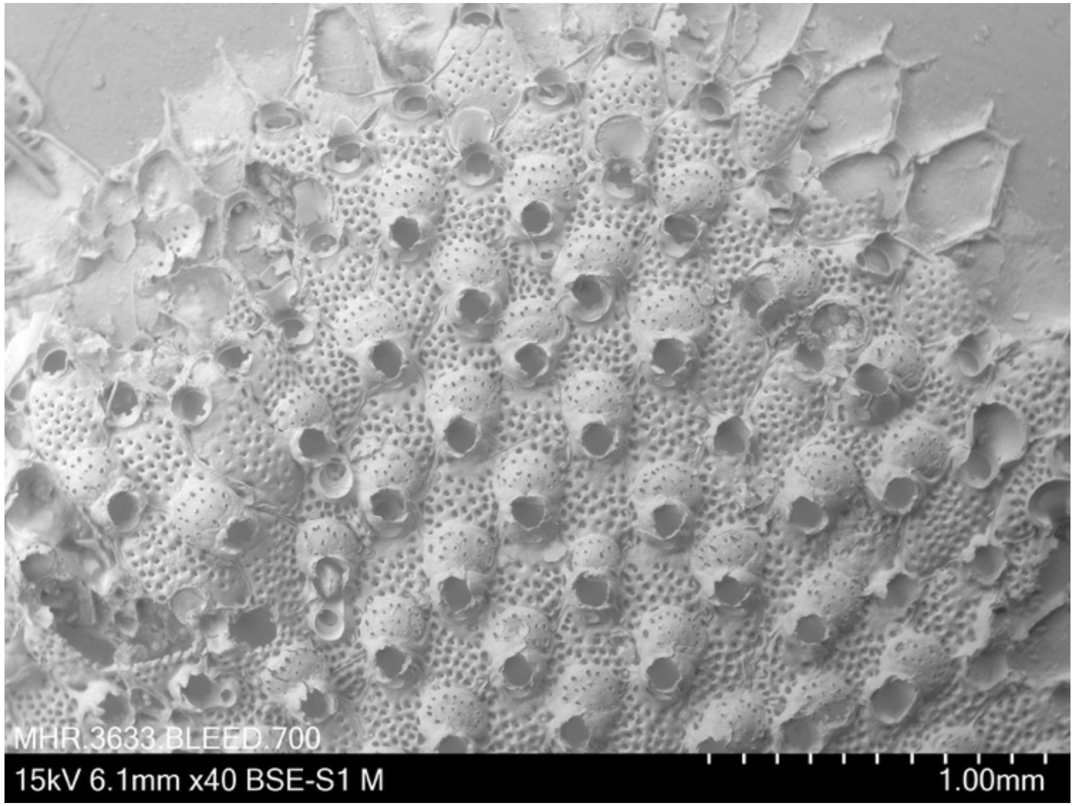


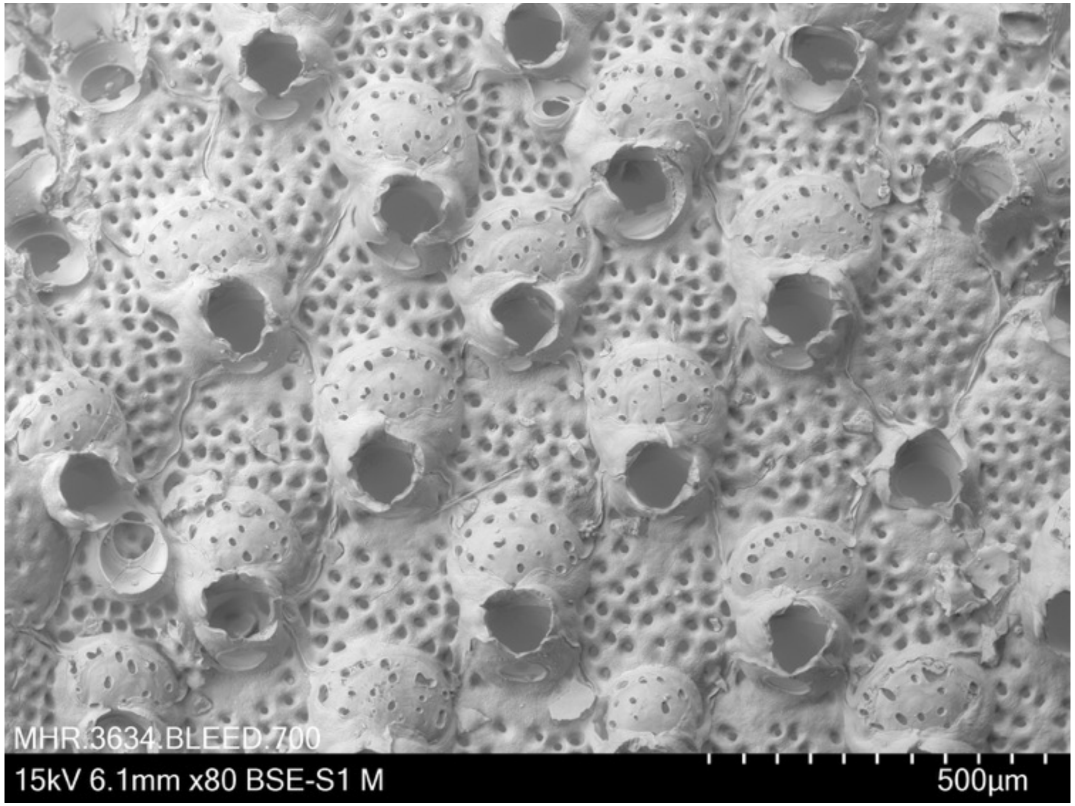


***Schizosmittina cinctipora* BLEED 700**
Coll: 8.4.2018 (A. M. Smith, D. P. Gordon, H. Mello)
Lat: -47.13583
Long: 180.01056
Depth: 72 m

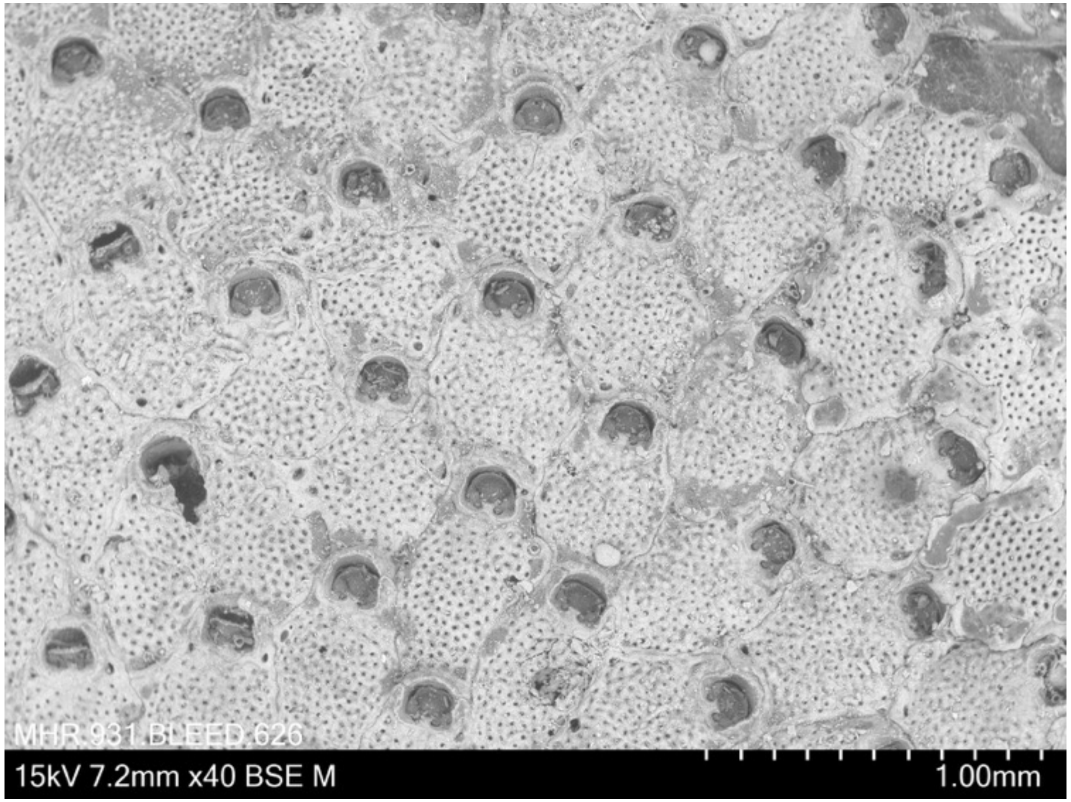


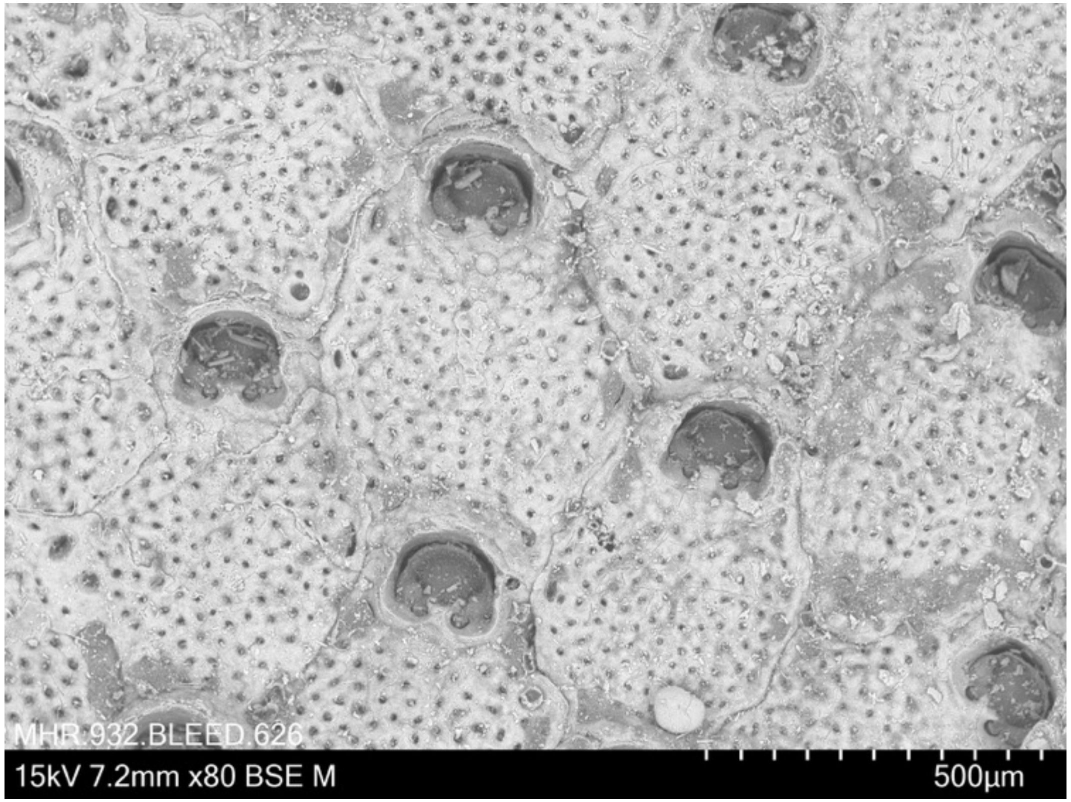


***Prenantia firmata* BLEED 626A**
Coll: 10.4.2018 (A. M. Smith, D. P. Gordon, H. Mello)
Lat: -46.945
Long: 168.1583
Depth: 42 m

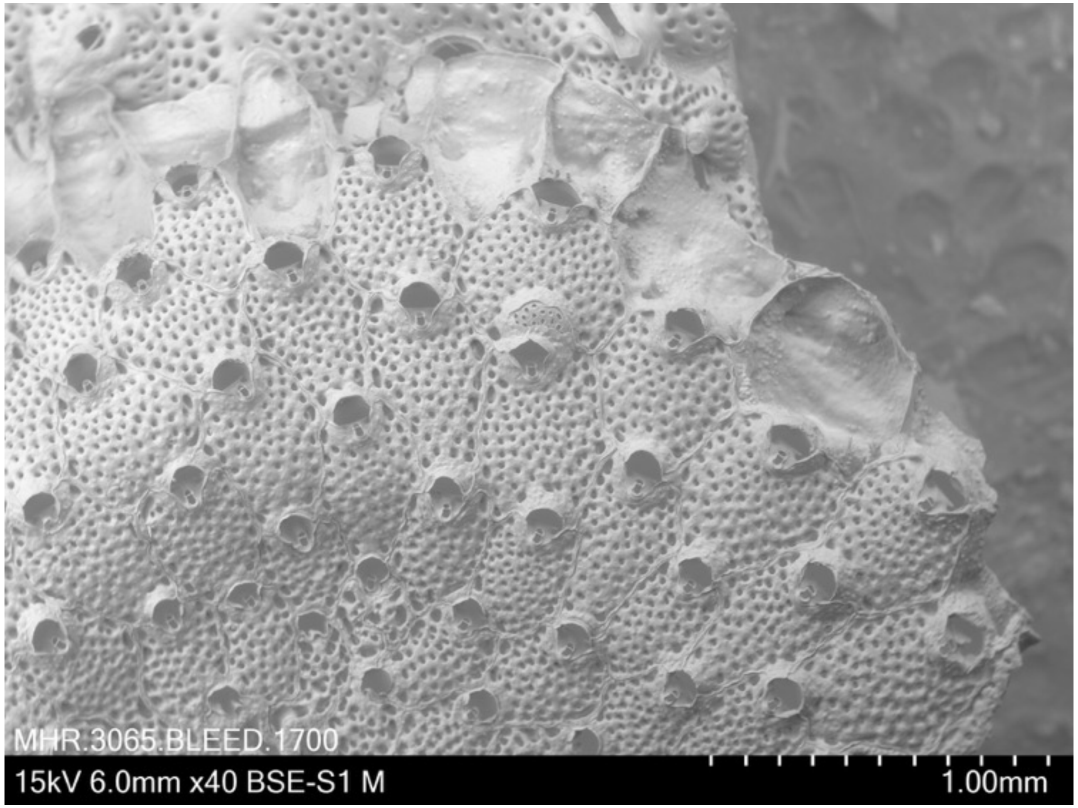


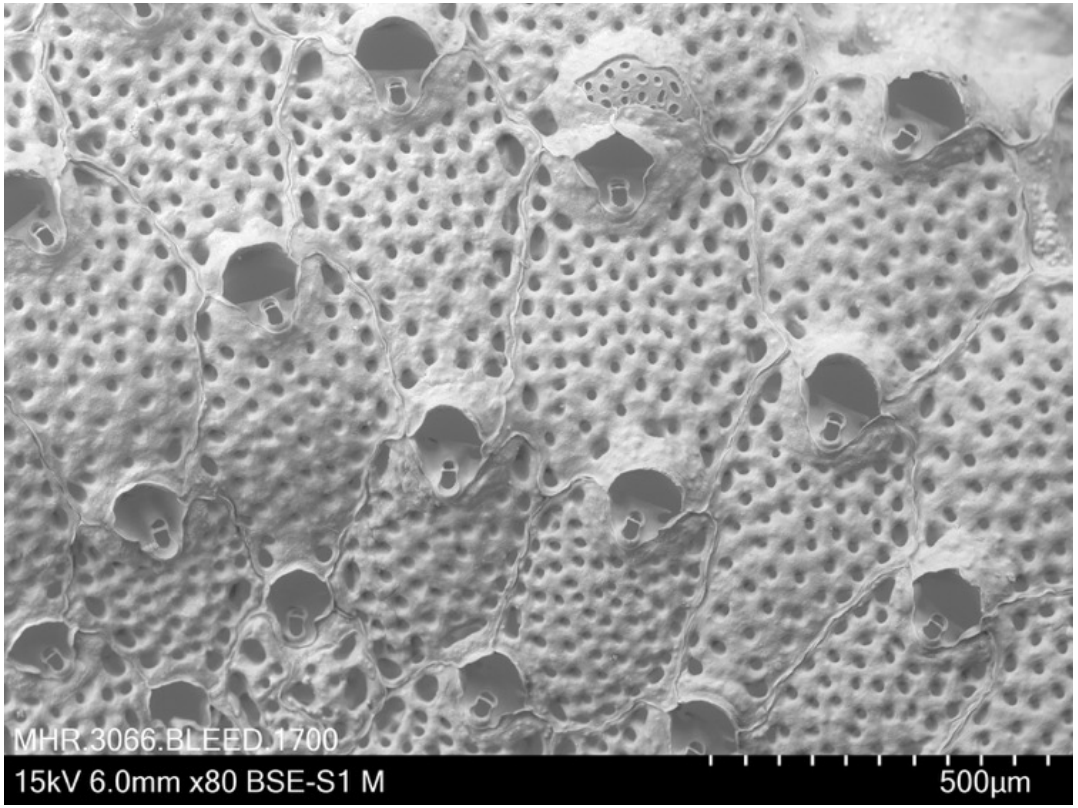


***Smittina rosacea*** **BLEED 1700**
Coll: 17.3.2019 (A. M. Smith, D. P. Gordon)
Lat: -34.16
Long: 172.09
Depth: 82.9 m

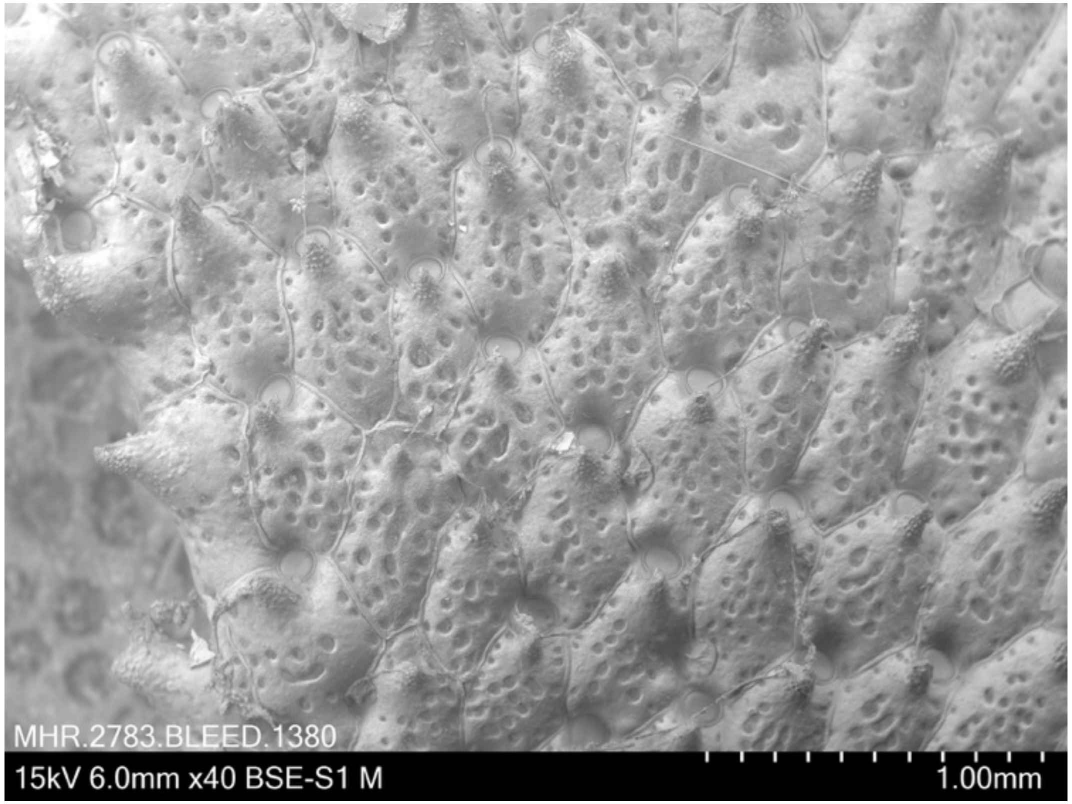

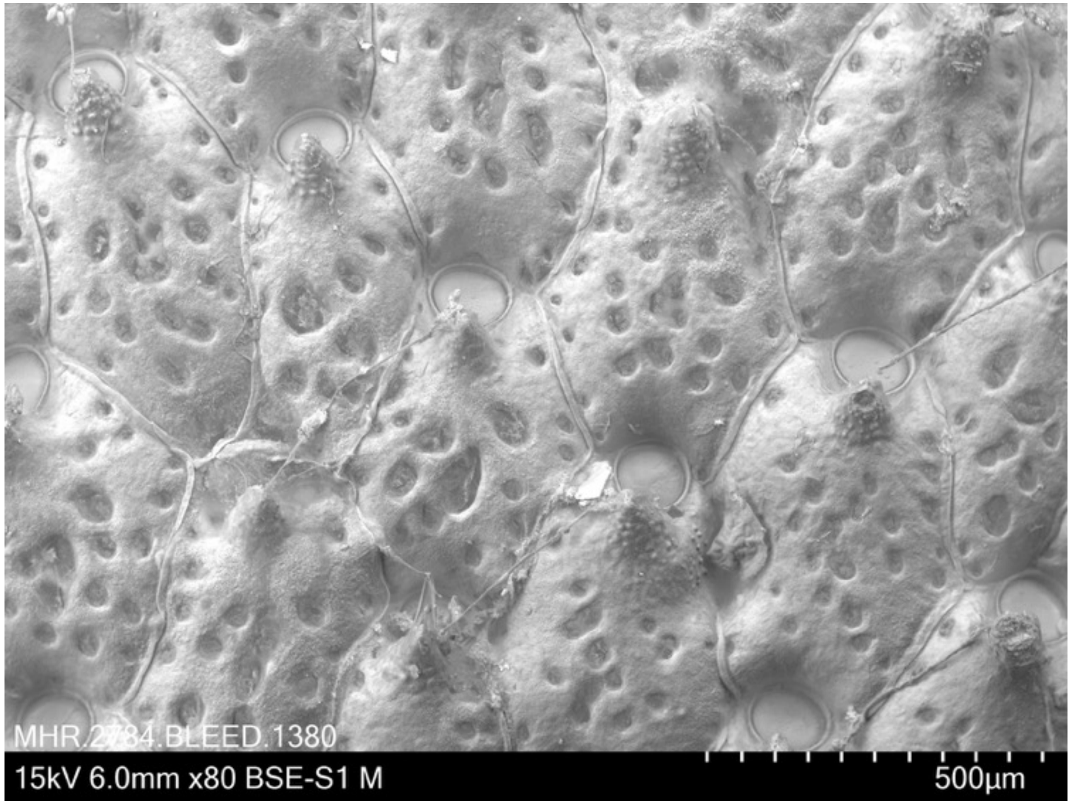


***Bitectipora mucronifera*** **BLEED 1380**
Coll: 17.3.2019 (A. M. Smith, D. P. Gordon)
Lat: -34.16
Long: 172.09
Depth: 82.9 m

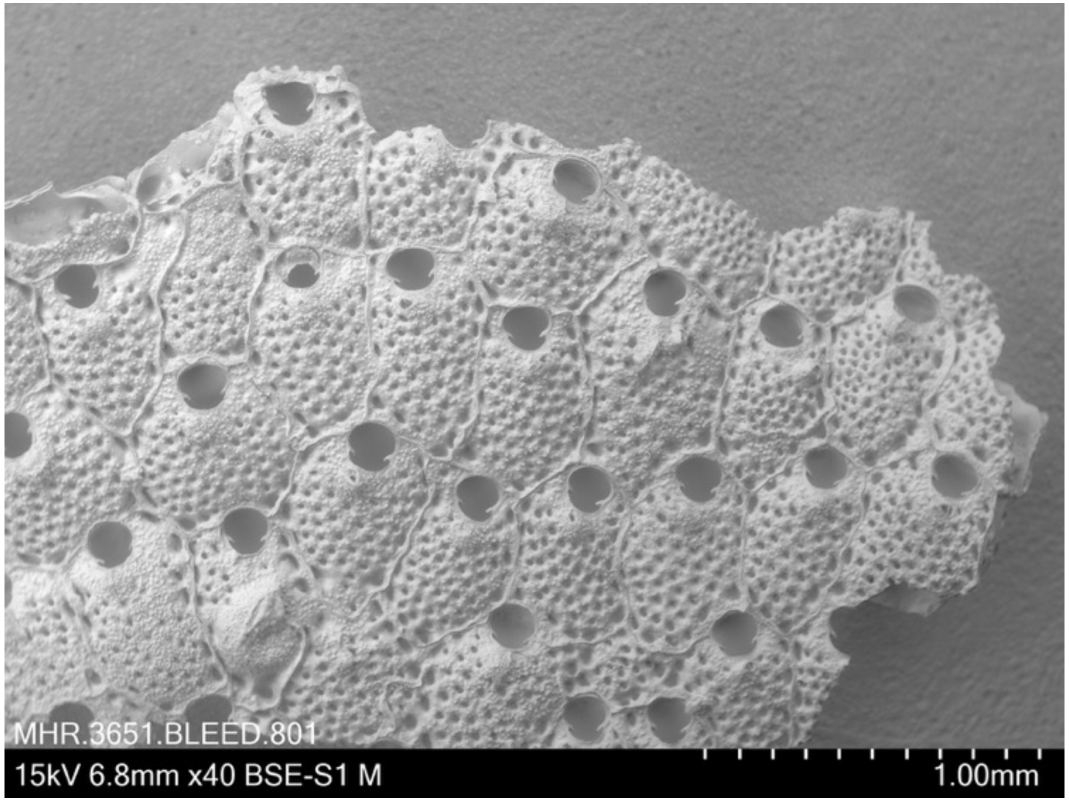


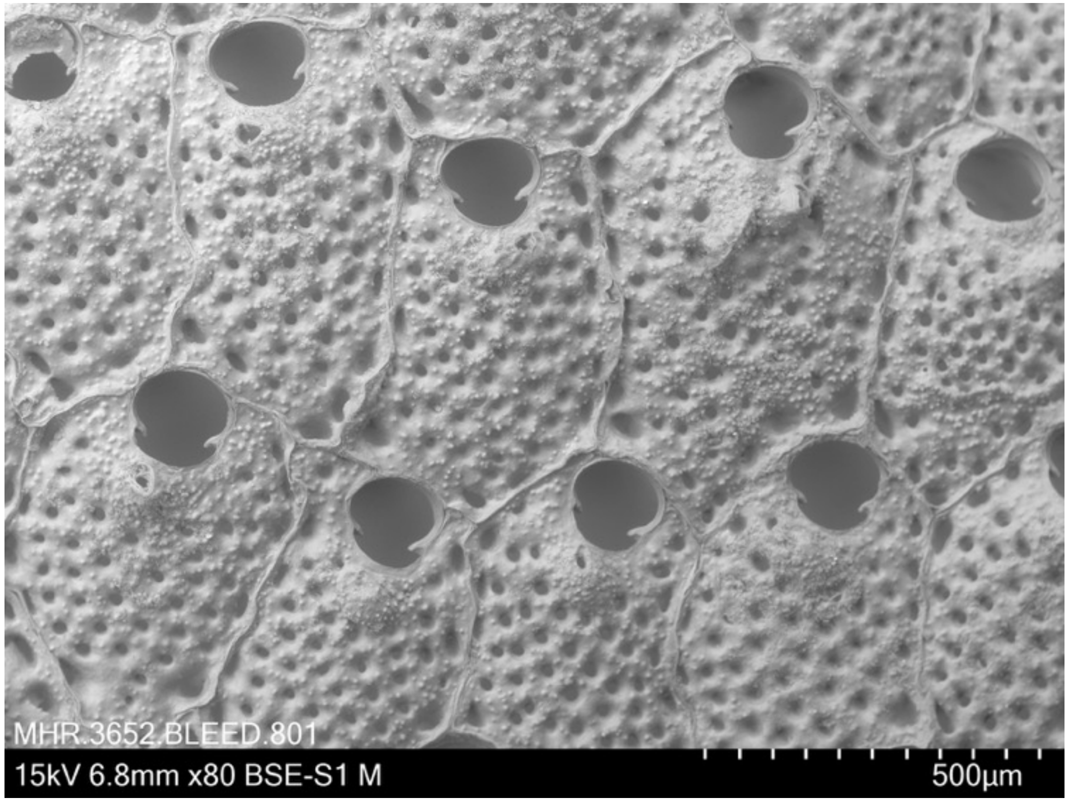


***Bitectipora cincta*** **BLEED 801**
Coll: 4.3.2018 (D. P.  Gordon, L .H. Liow, C. Schack)
Lat: -39.639352
Long: 177.008705
Depth: intertidal

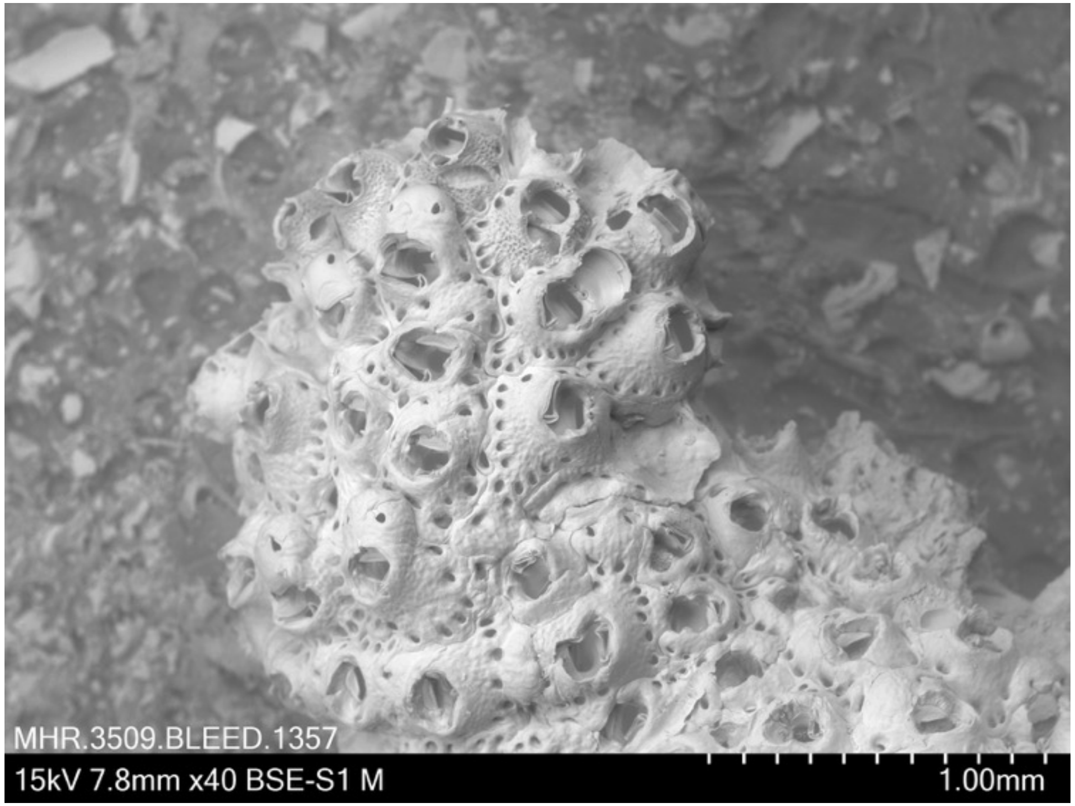


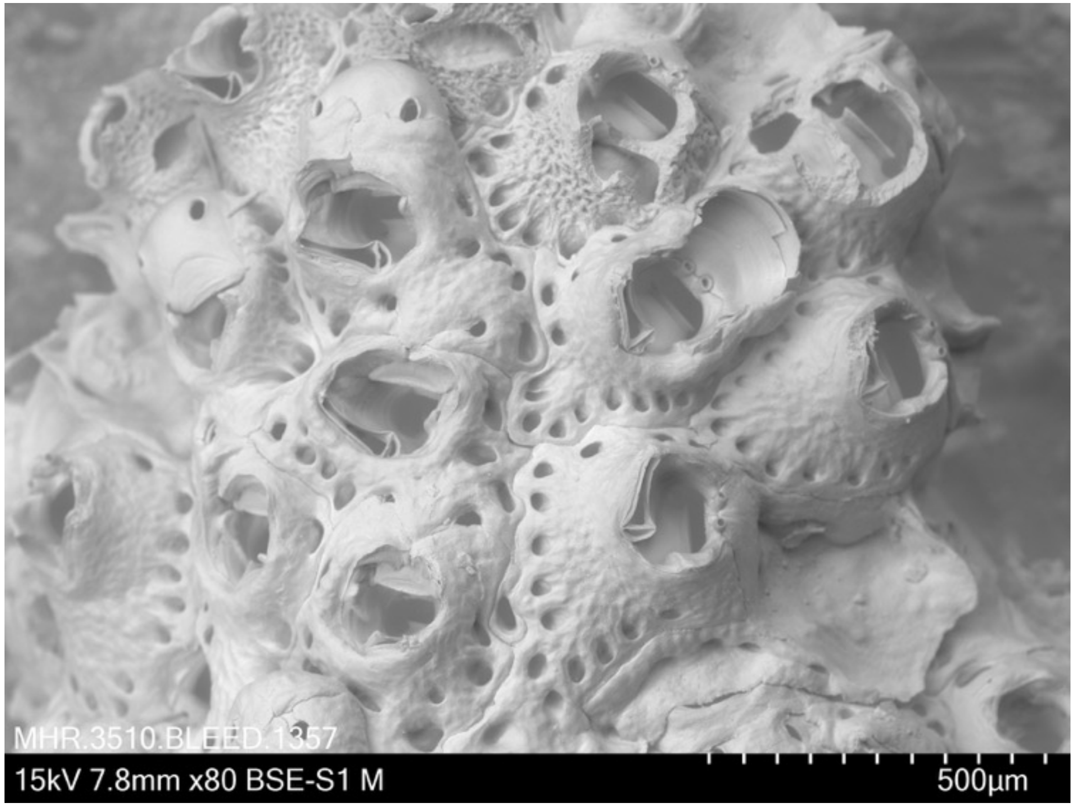


***Drepanophora biperforata* BLEED 1357**
Coll: 19.3.2019 (A. M. Smith, D. P. Gordon)
Lat: -34.99
Long: 173.97
Depth: 61.6 m

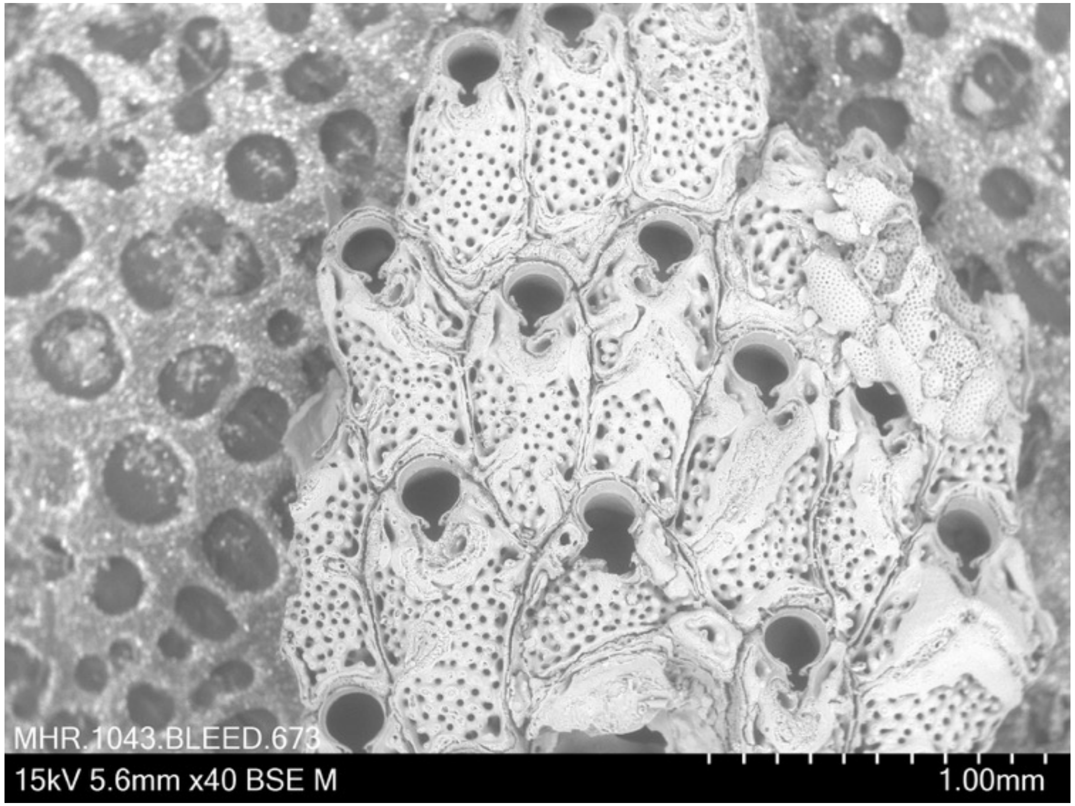


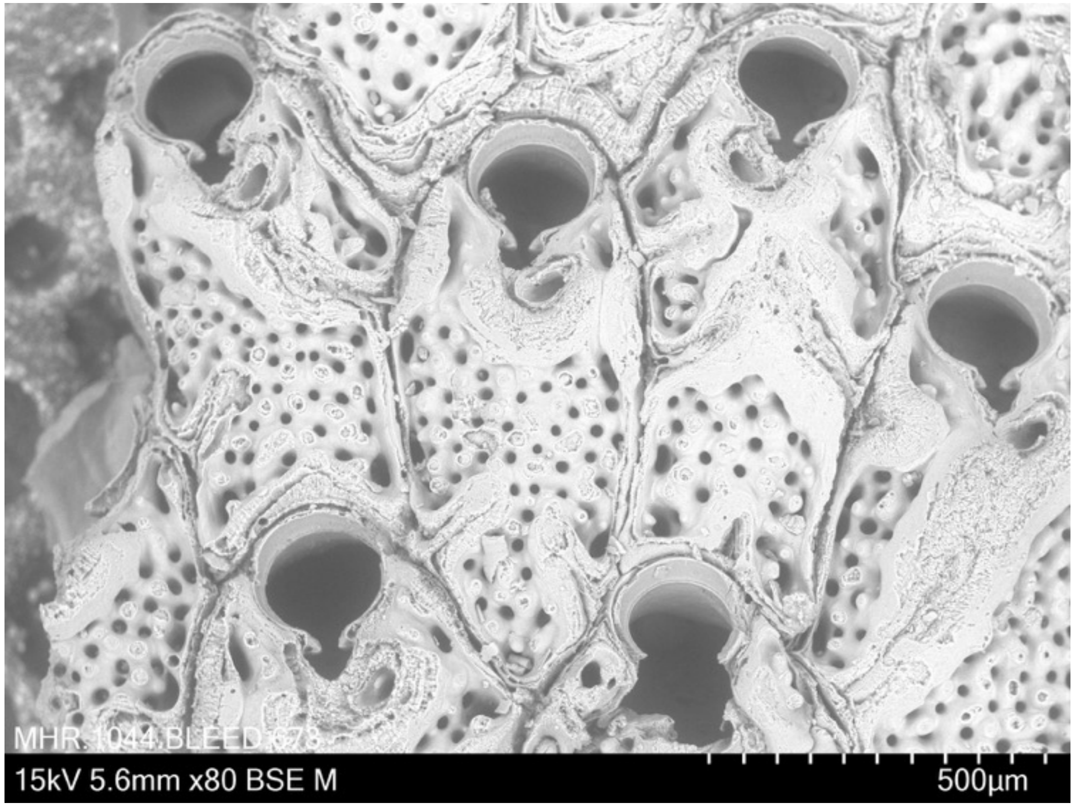


***Fovoporella spectabilis*** **BLEED 673**
Coll: 8.4.2018 (A. M. Smith, D. P. Gordon, H. Mello)
Lat: -47.1372
Long: 168.12583
Depth: 80-85m

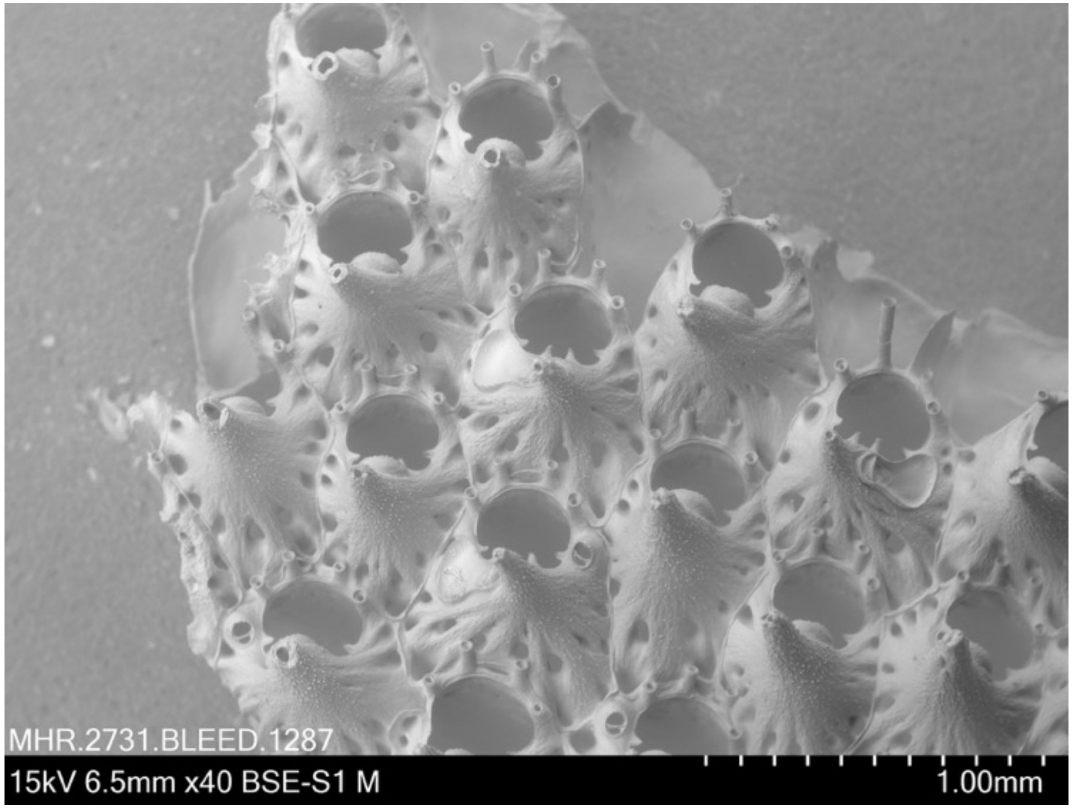


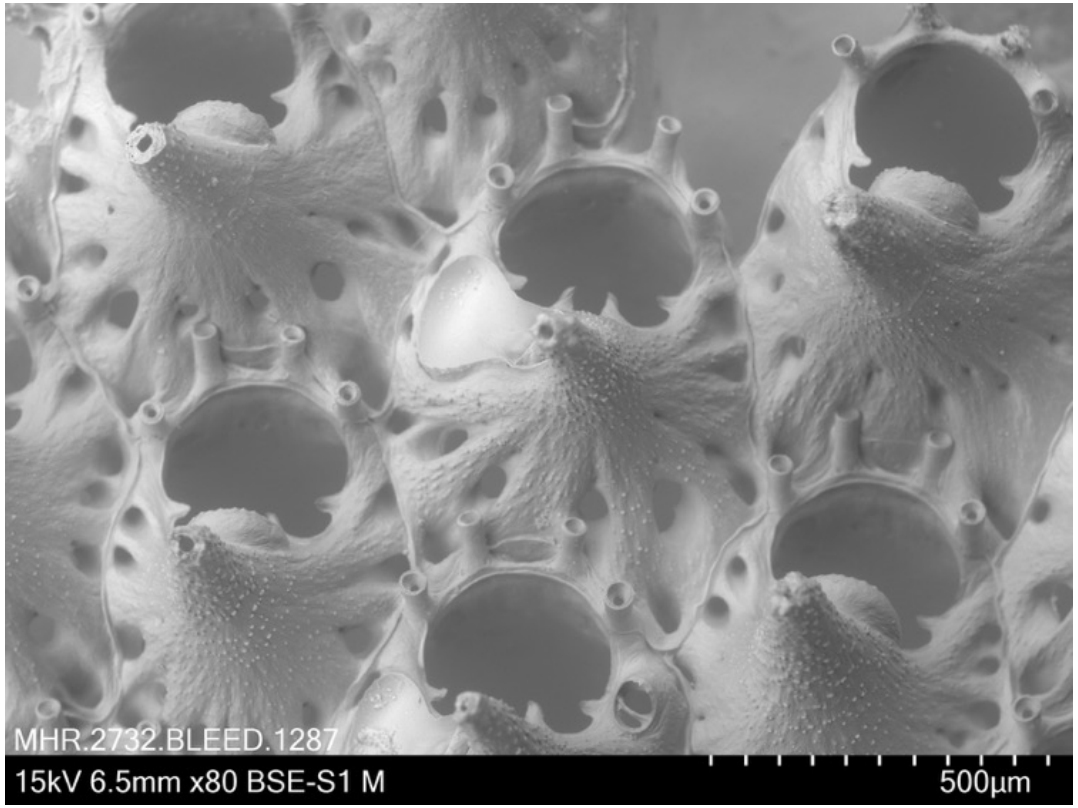


***Mobunula bicuspis* BLEED 1287**
Coll: 17.3.2019 (A. M. Smith, D. P. Gordon)
Lat: -34.17
Long: 172.04
Depth: 85.4 m

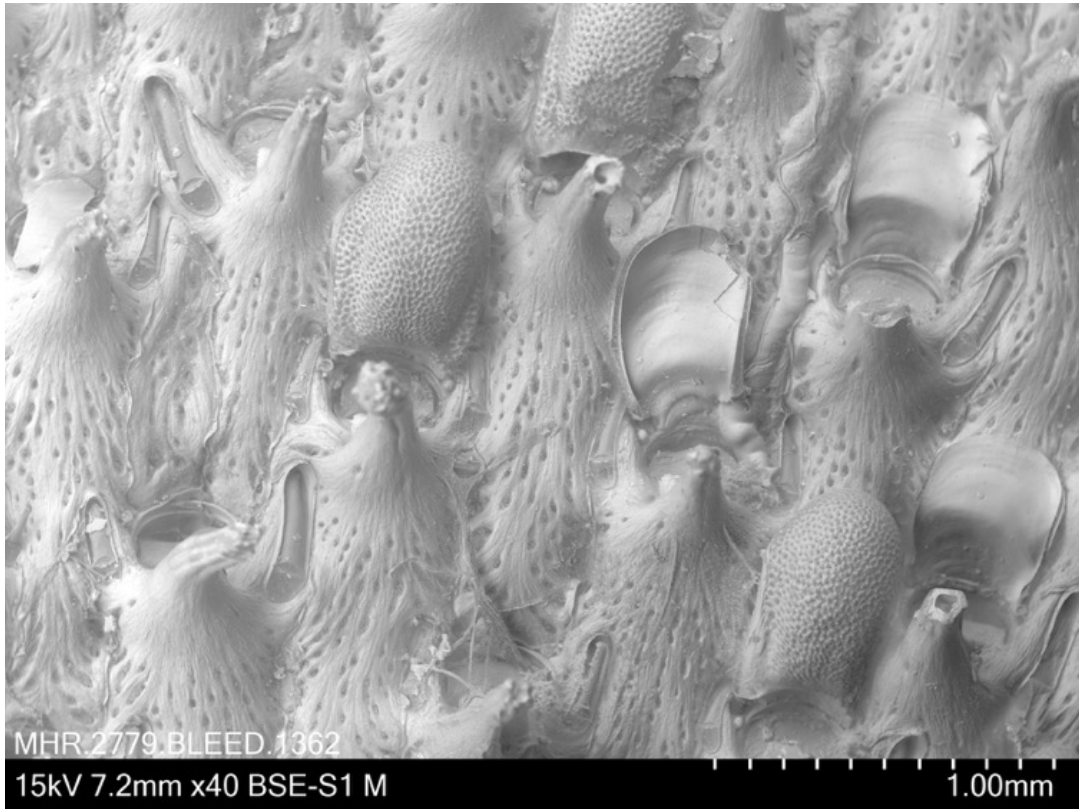


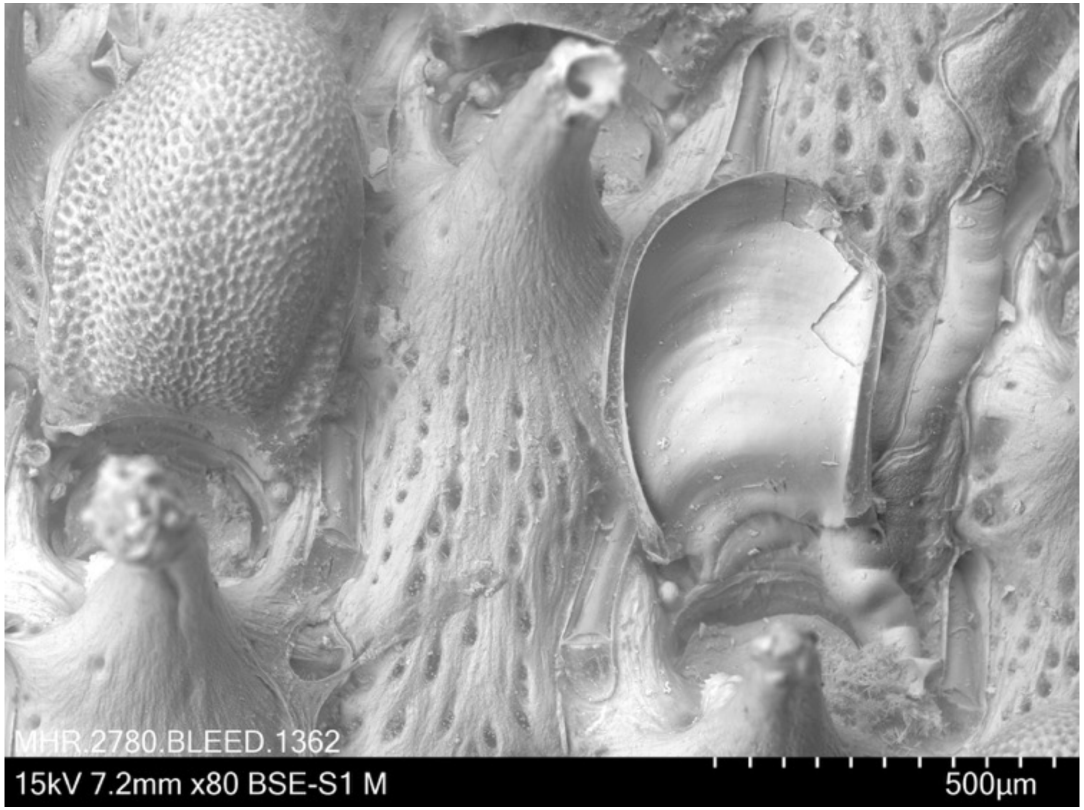


***Mucropetraliella vultur* BLEED 1362**
Coll: 17.3.2019 (A. M. Smith, D. P. Gordon)
Lat: -34.21
Long: 172.11
Depth: 104 m

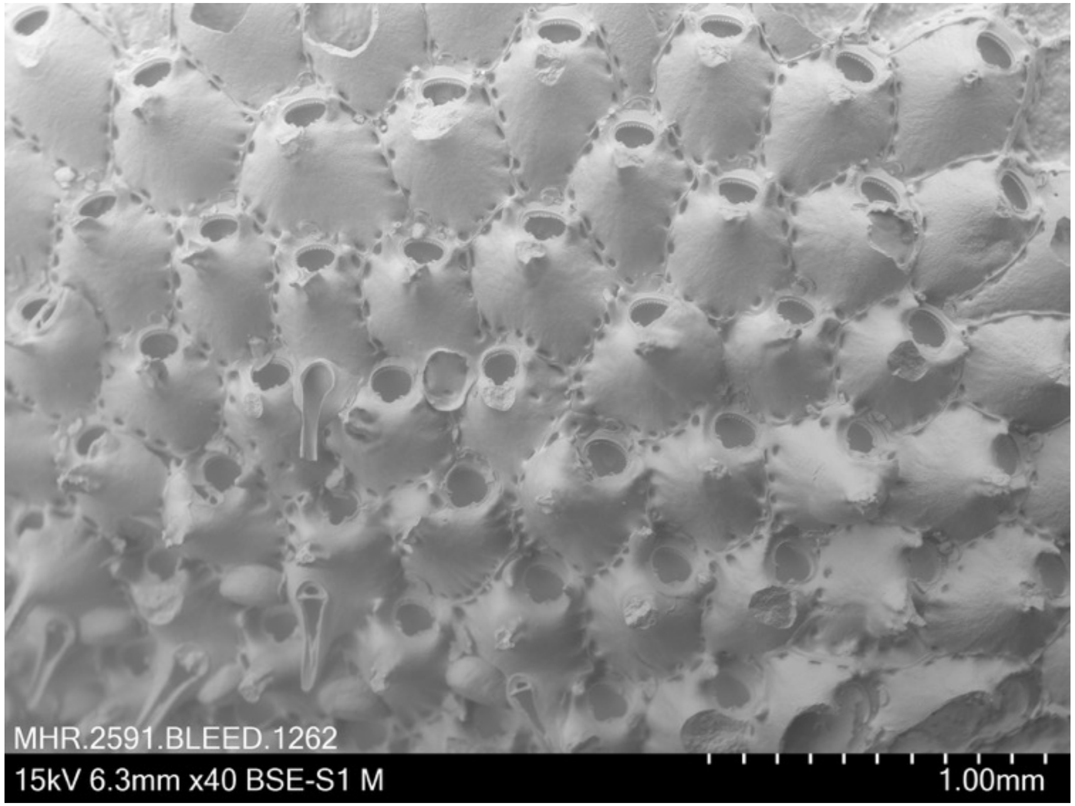


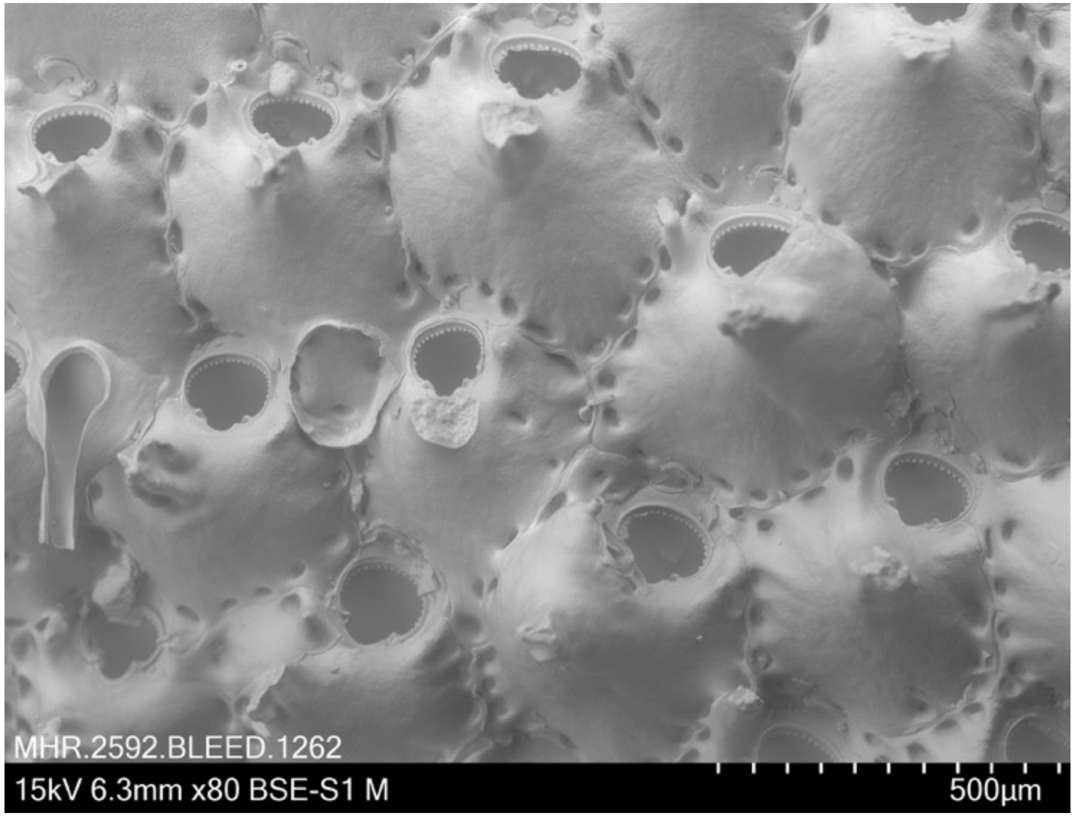


***Rhynchozoon* n. sp. BLEED 1262**
Coll: 16.3.2019 (A. M. Smith, D. P. Gordon)
Lat: -35.01
Long: 173.92
Depth: 17 m

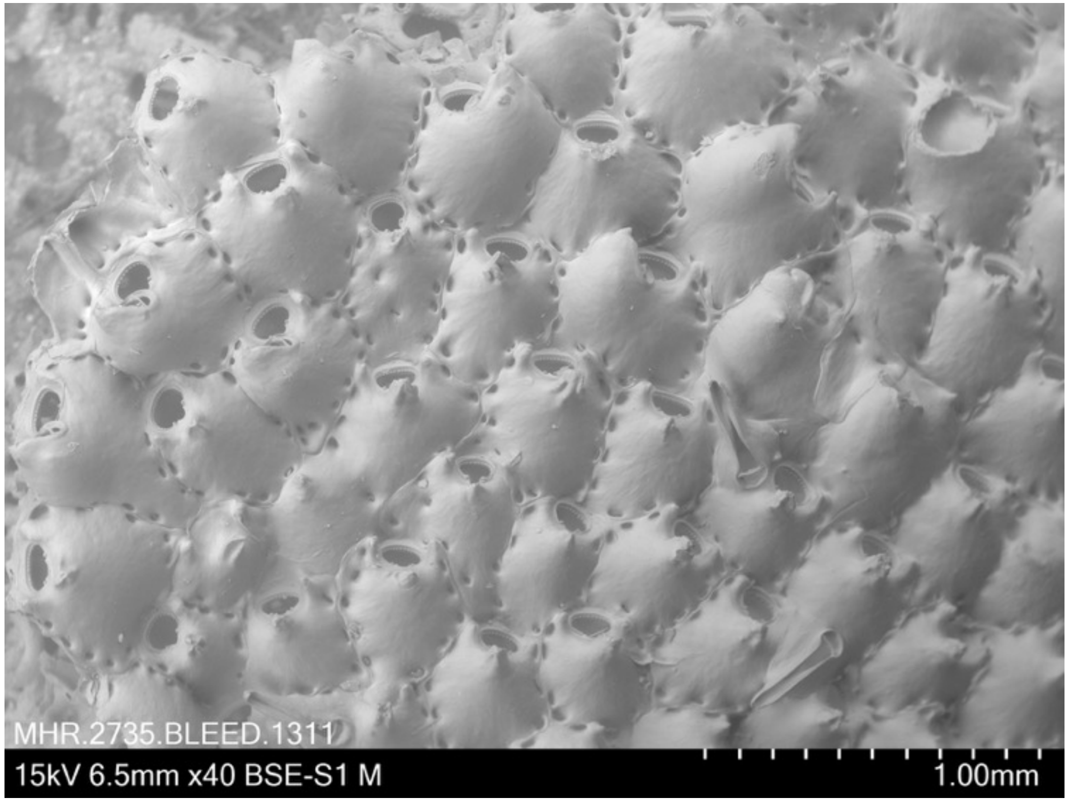


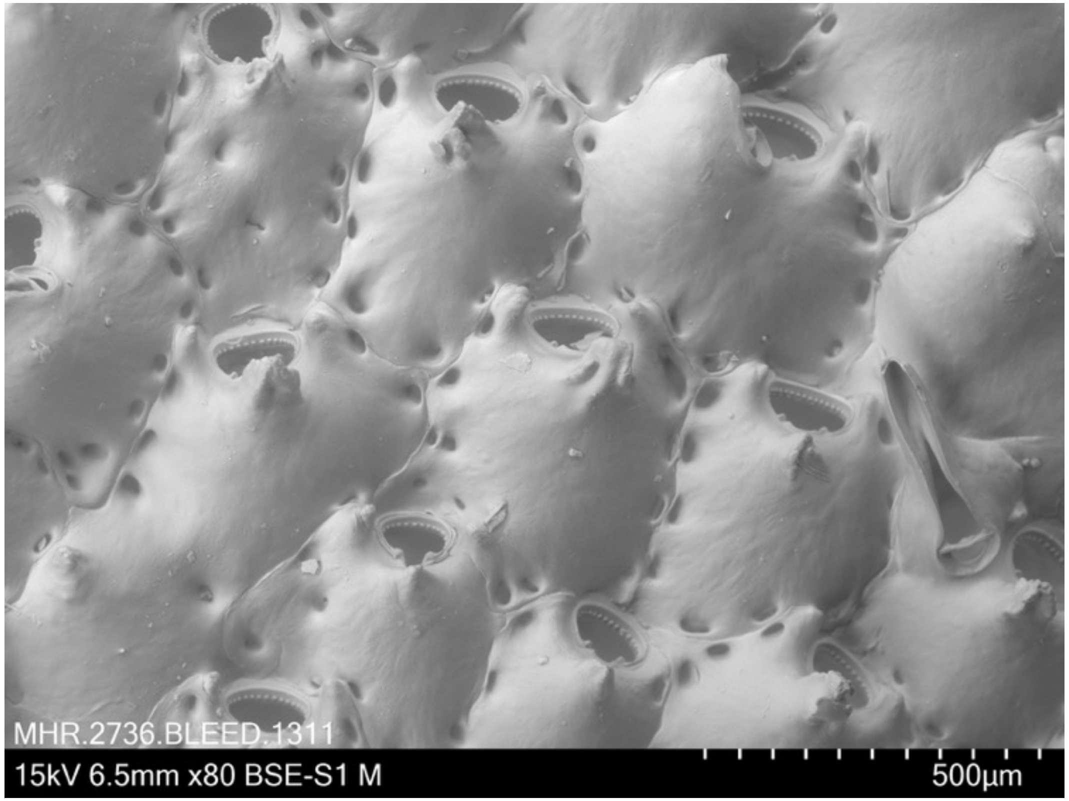


***Rhynchozoon* n. sp. BLEED 1311**
Coll: 15.3.2019 (A. M. Smith, D. P. Gordon)
Lat: -36.46
Long: 174.9
Depth: 37 m

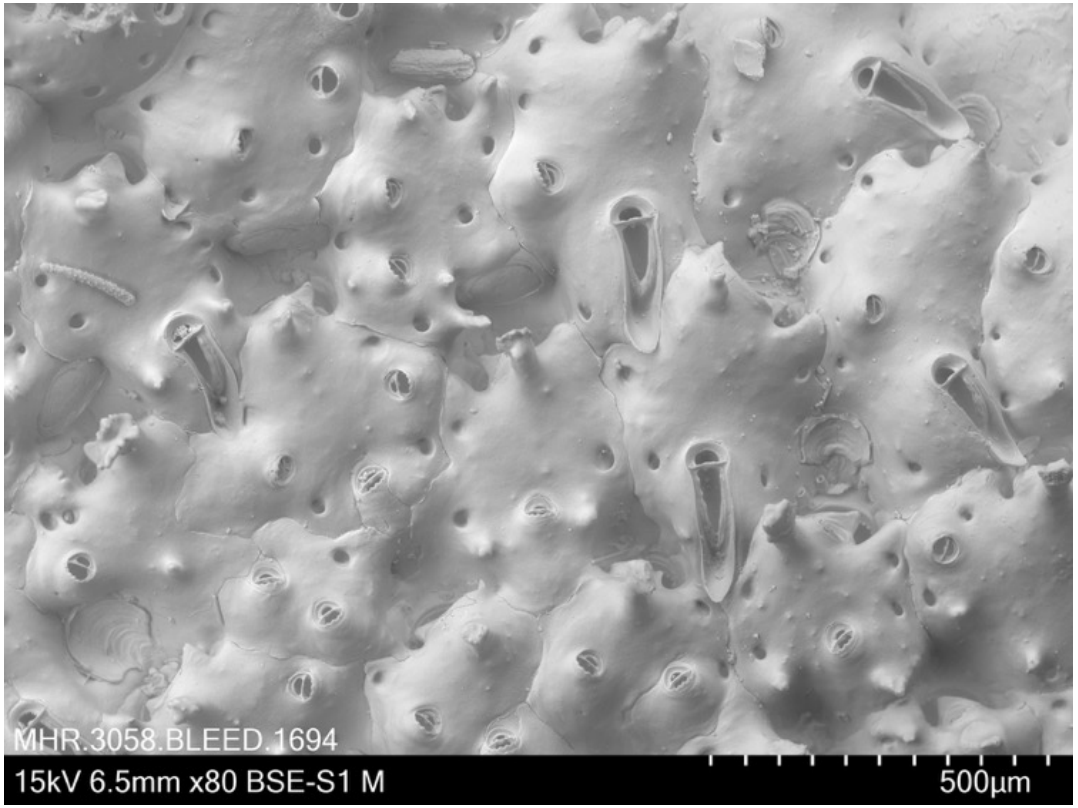

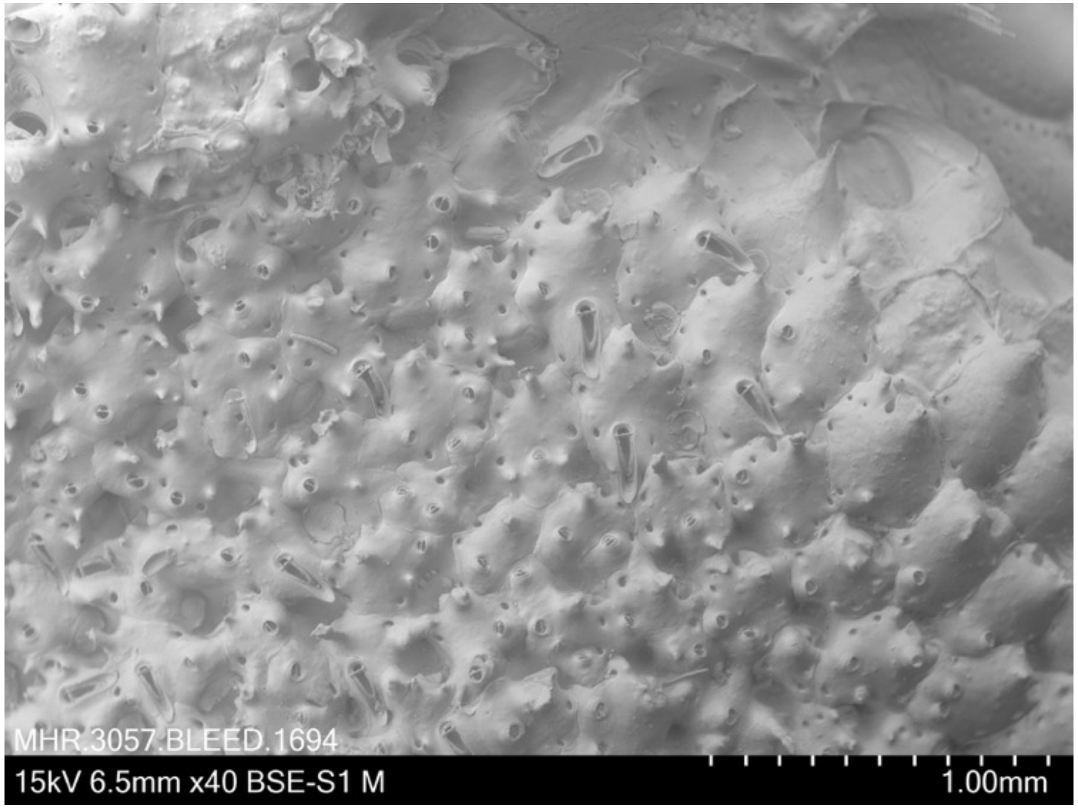


***Rhynchozoon paa* BLEED 1694**
Coll: 19.3.2019 (A. M. Smith, D. P. Gordon)
Lat: -34.99
Long: 173.97
Depth: 61.6 m

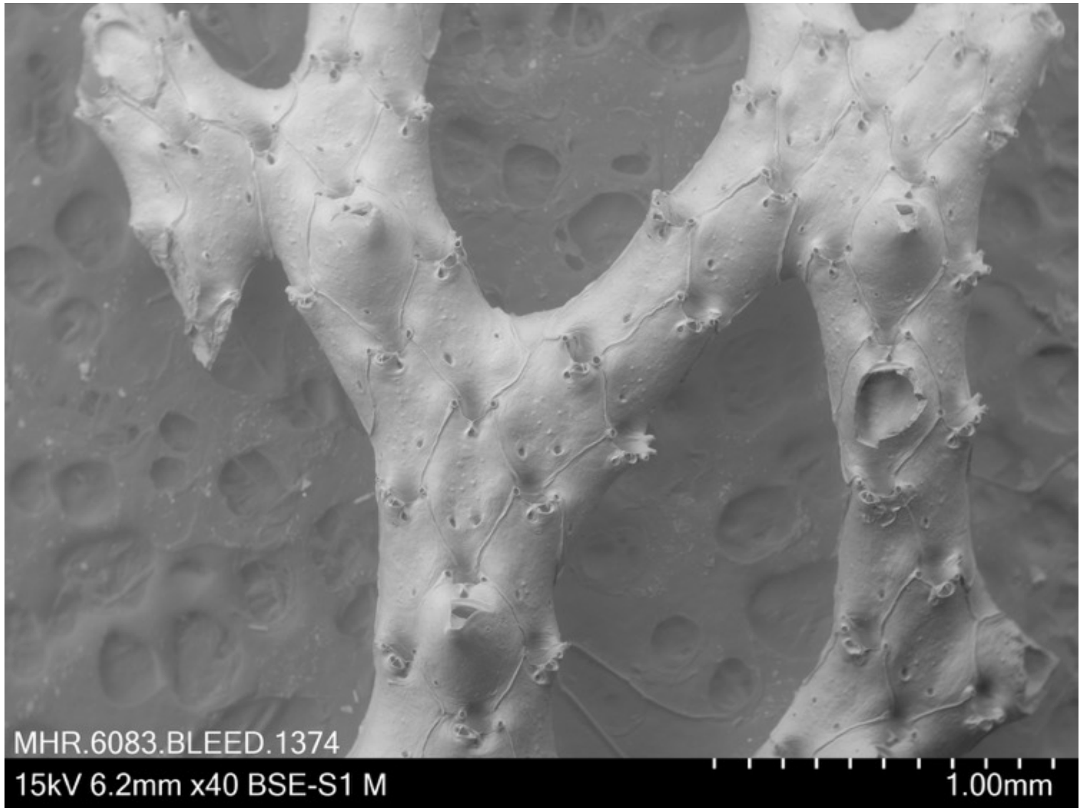

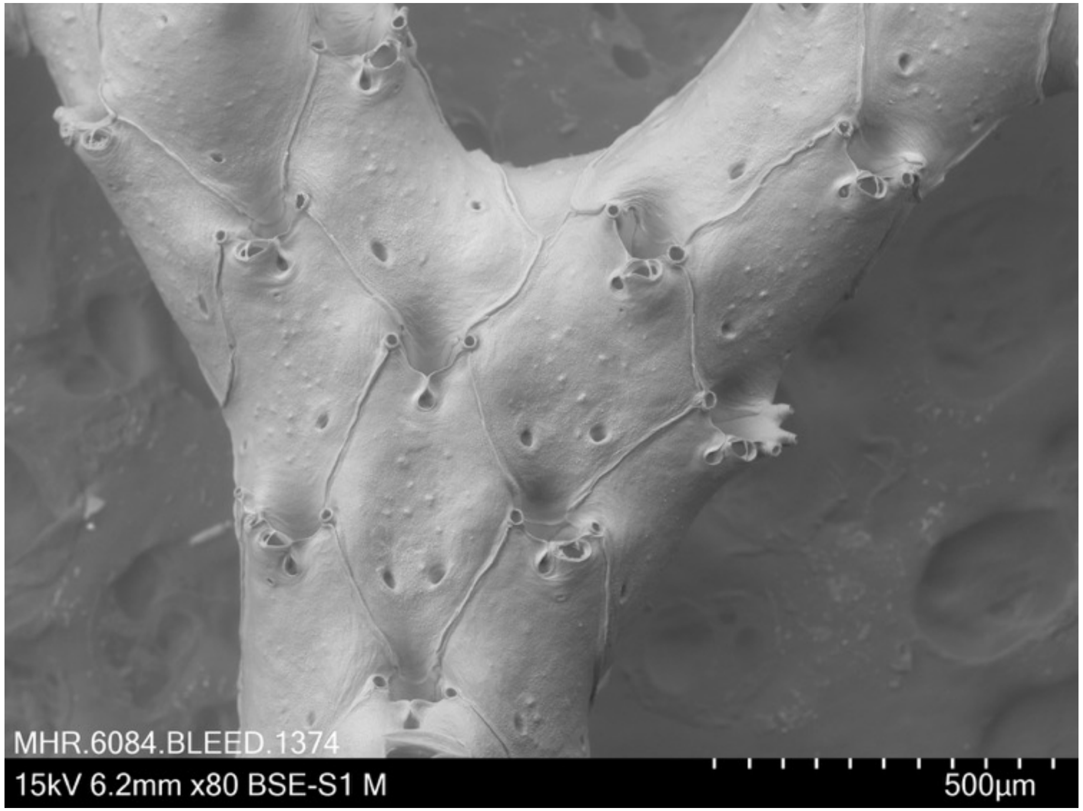


***Phidolopora avicularis* BLEED 1374**
Coll: 17.3.2019 (A. M. Smith, D. P .Gordon)
Lat: -34.159999999999997
Long: 172.09
Depth: 82.9 m

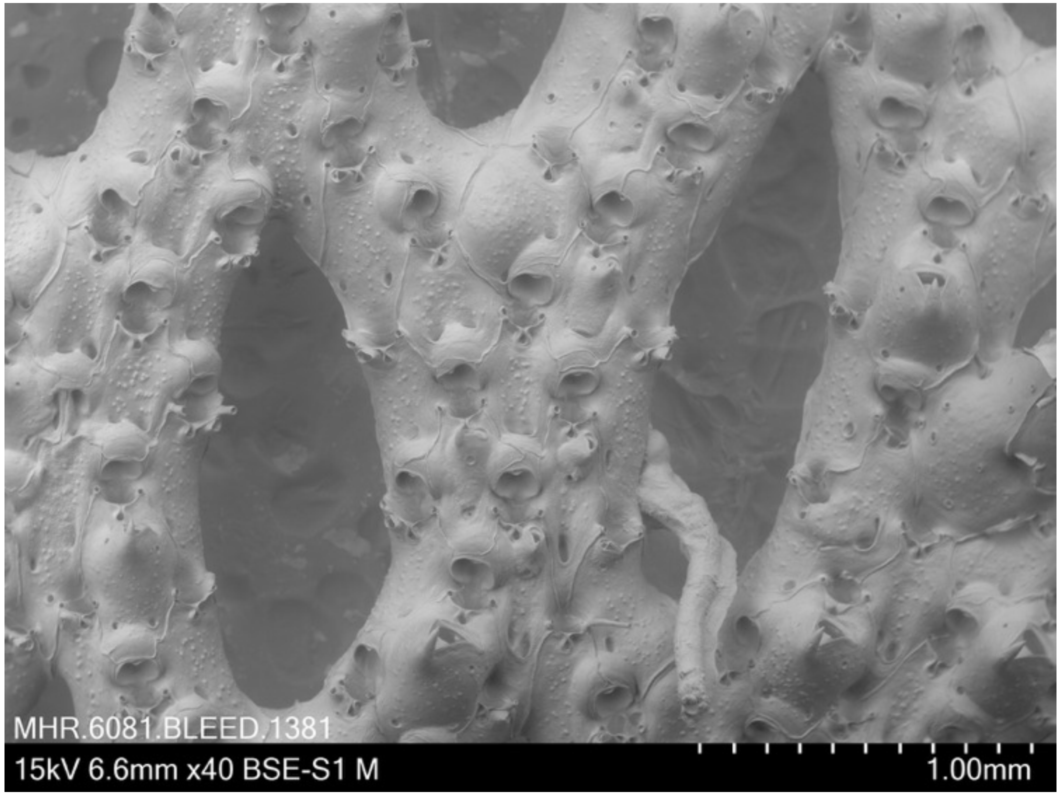


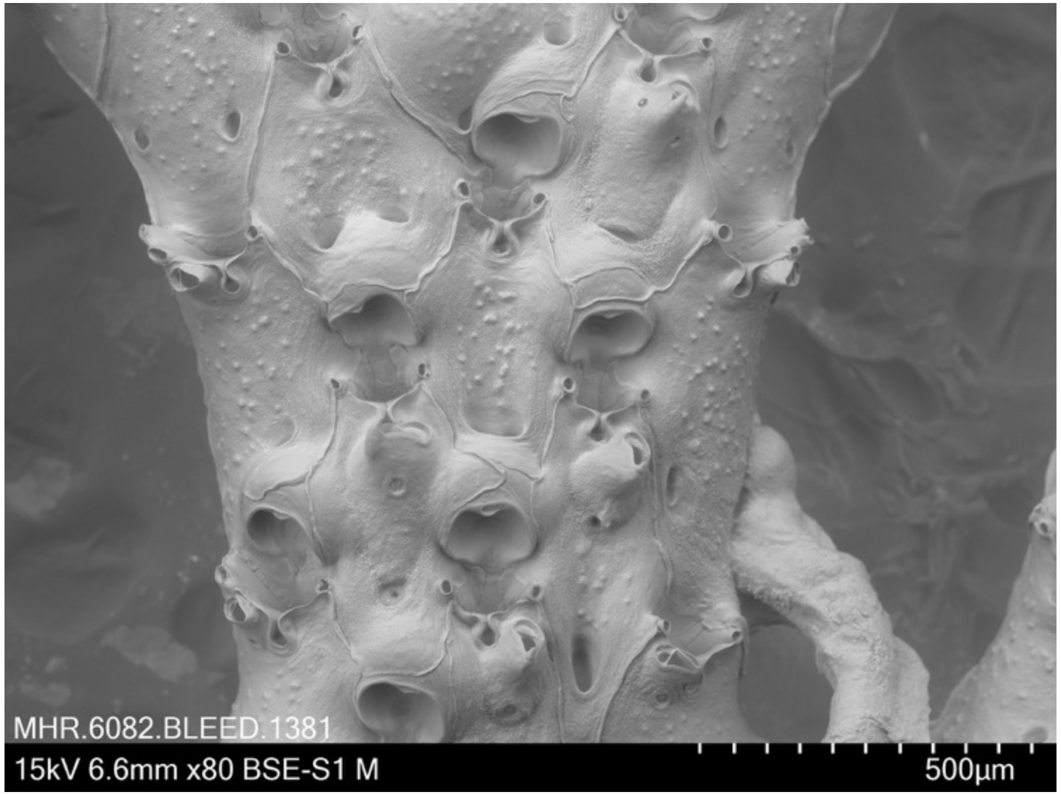


***Phidolopora avicularis* BLEED 1381**
Coll: 17.3.2019 (A. M. Smith, D. P. Gordon)
Lat: -34.159999999999997
Long: 172.09
Depth: 82.9 m

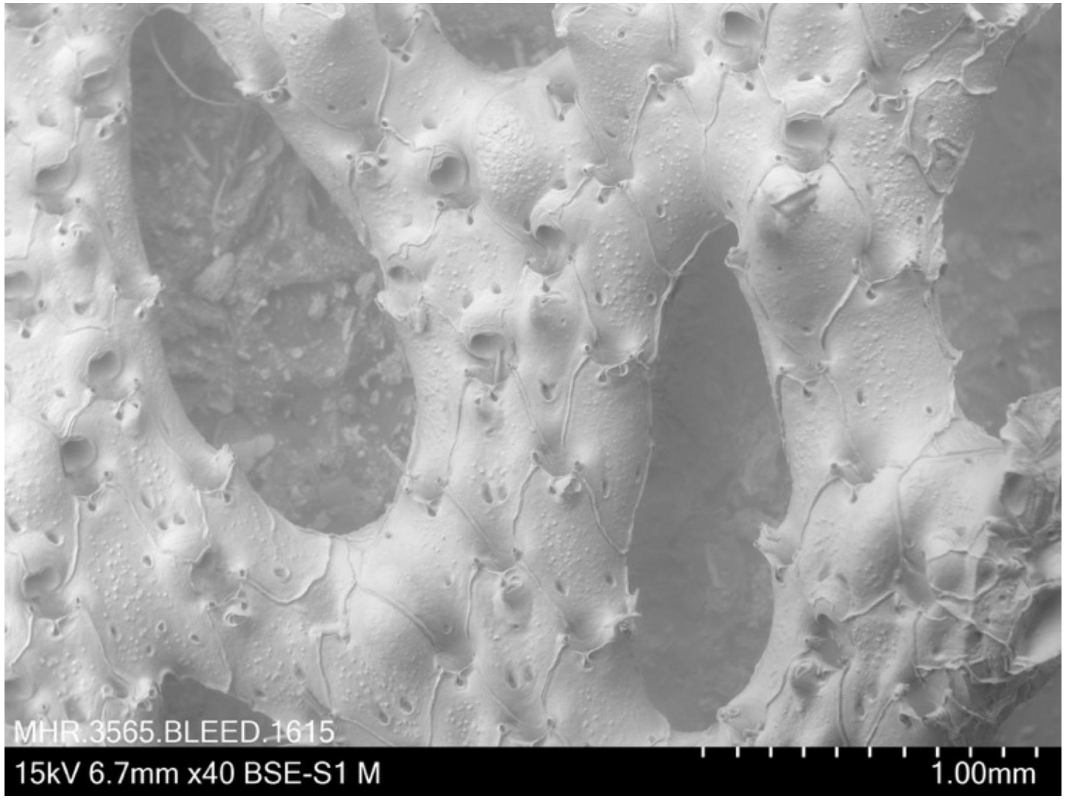


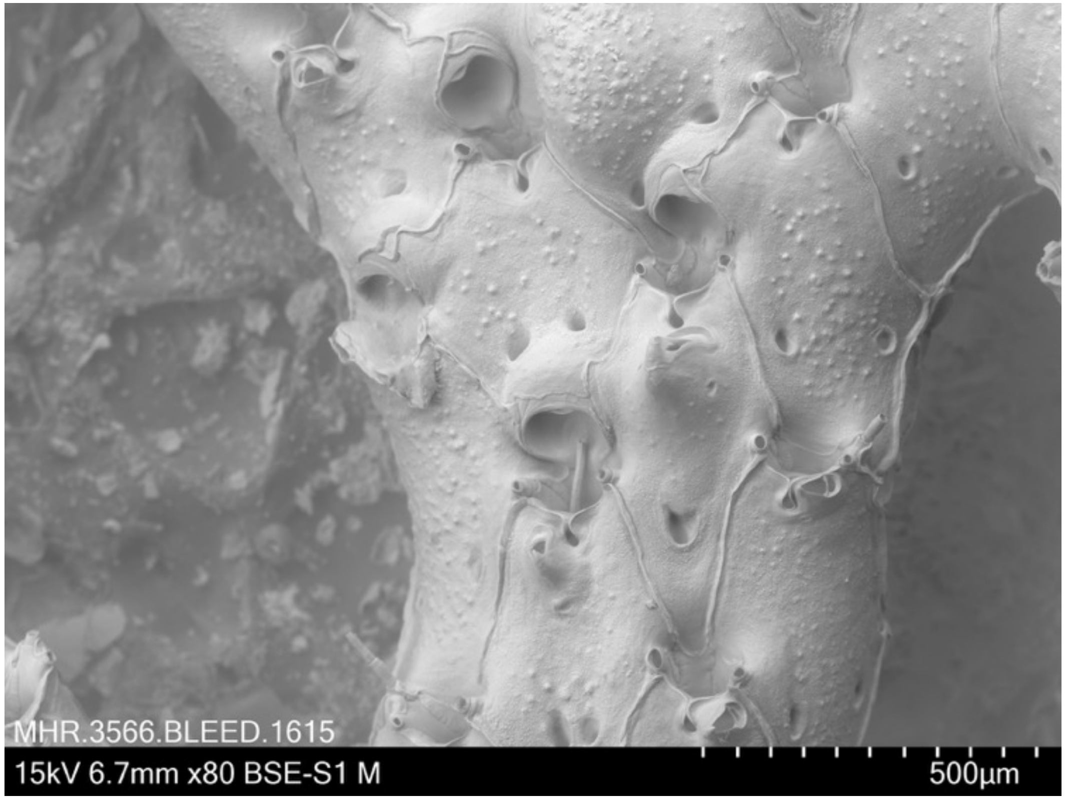


***Phidolopora avicularis* BLEED 1615**
Coll: 17.3.2019 (A. M. Smith, D. P. Gordon)
Lat: -34.159999999999997
Long: 172.09
Depth: 82.9 m

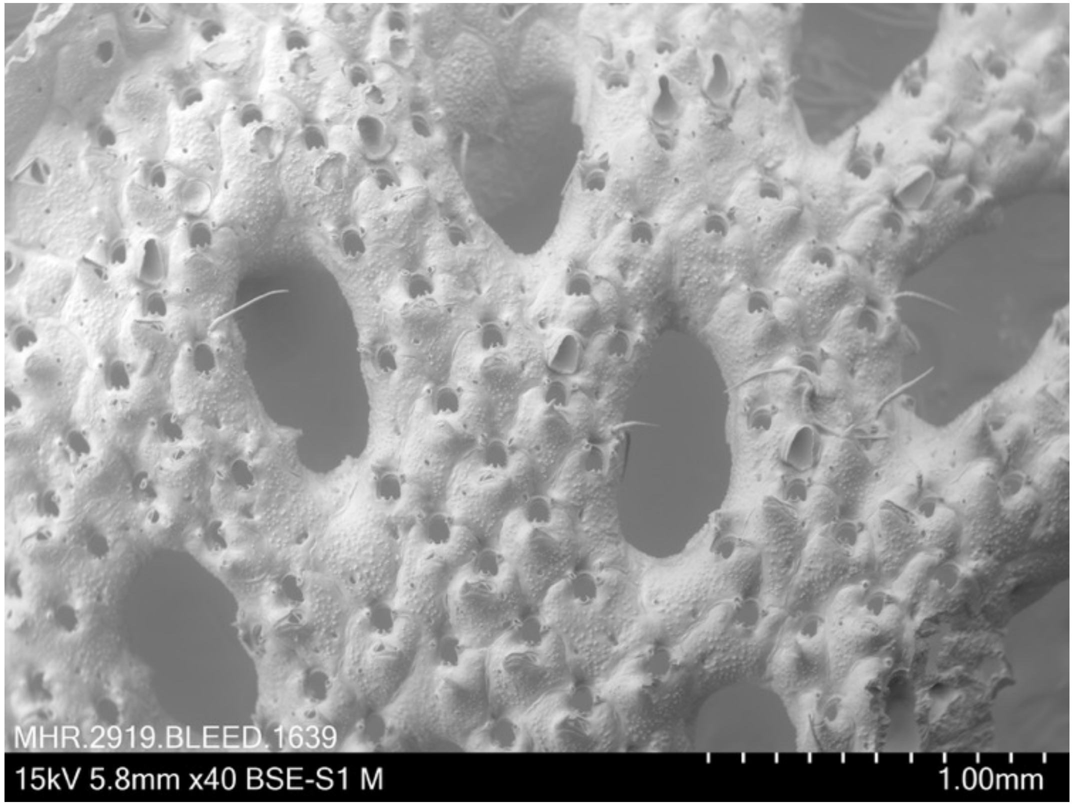


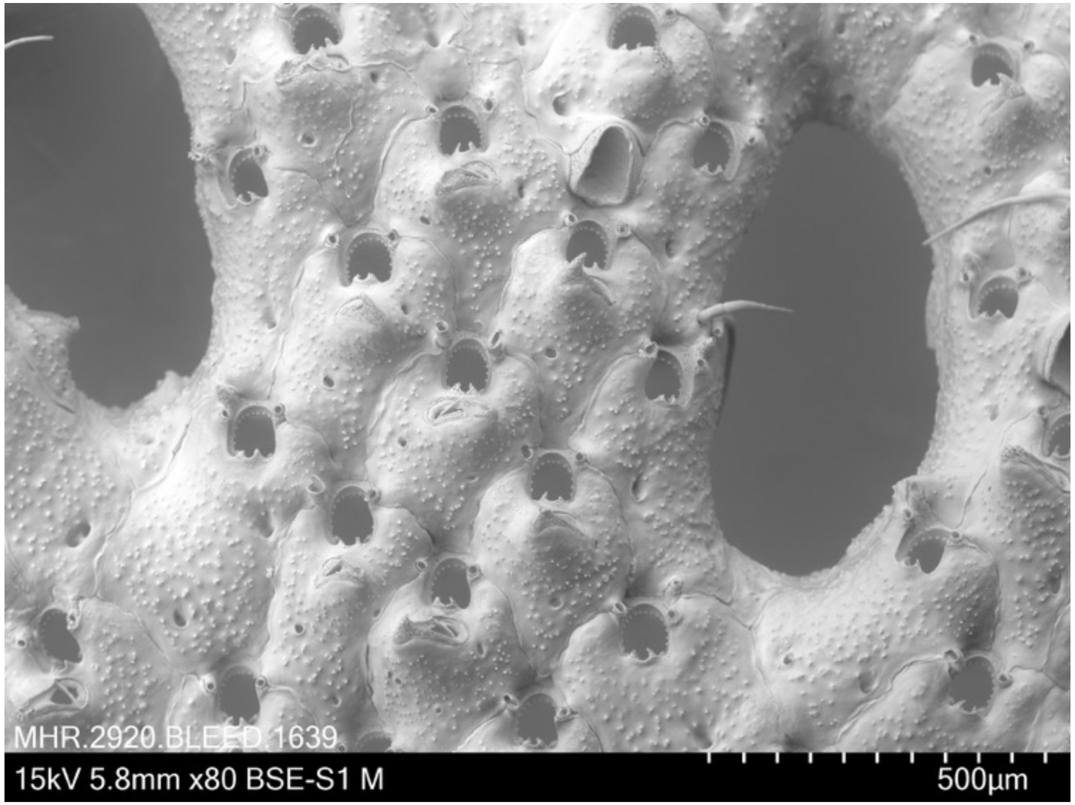


***Hippellozoon novaezelandiae* BLEED 1639**
Coll: 17.3.2019 (A. M. Smith, D. P. Gordon)
Lat: -34.16
Long: 172.09
Depth: 82.9 m

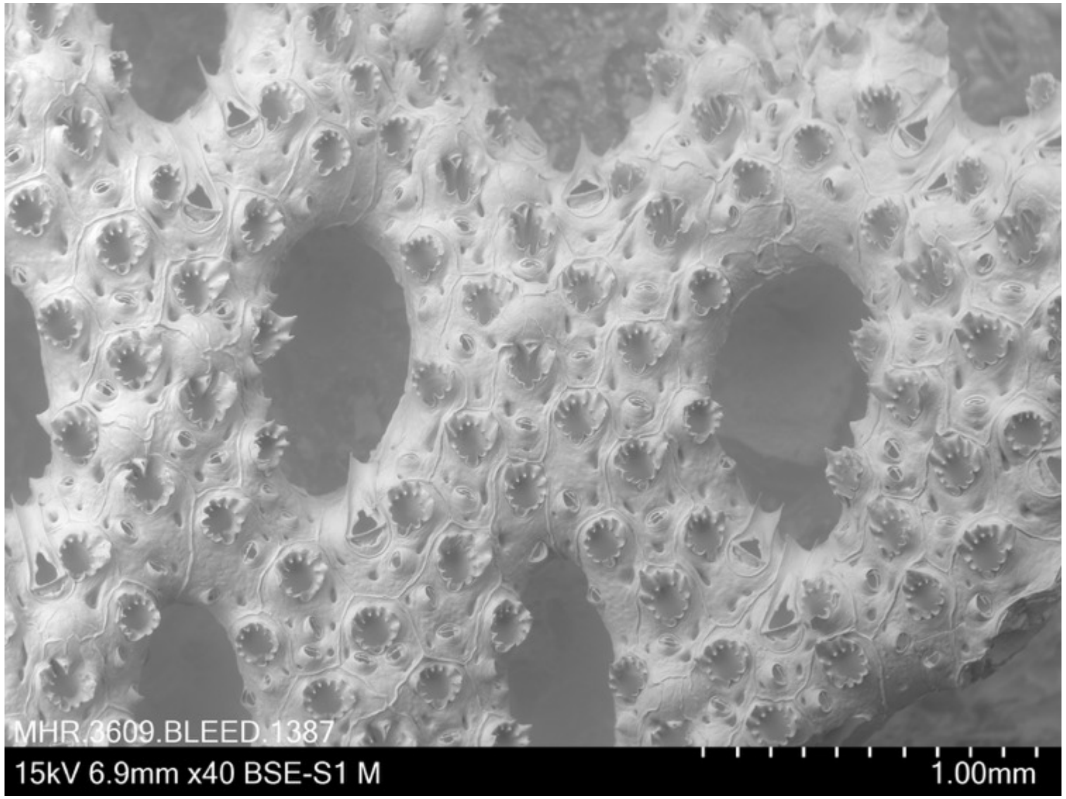


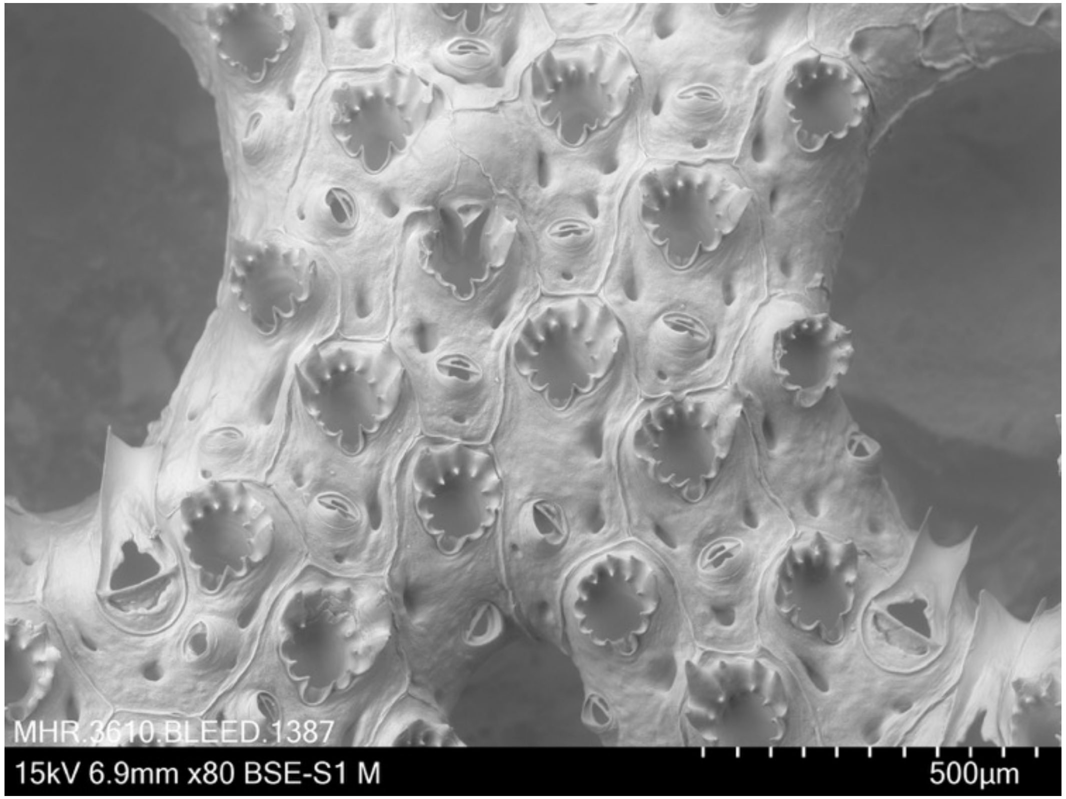


***Iodictyum yaldwynii*** **BLEED 1387**
Coll: 17.3.2019 (A. M. Smith, D. P. Gordon )
Lat: -34.159999999999997
Long: 172.09
Depth: 82.9 m

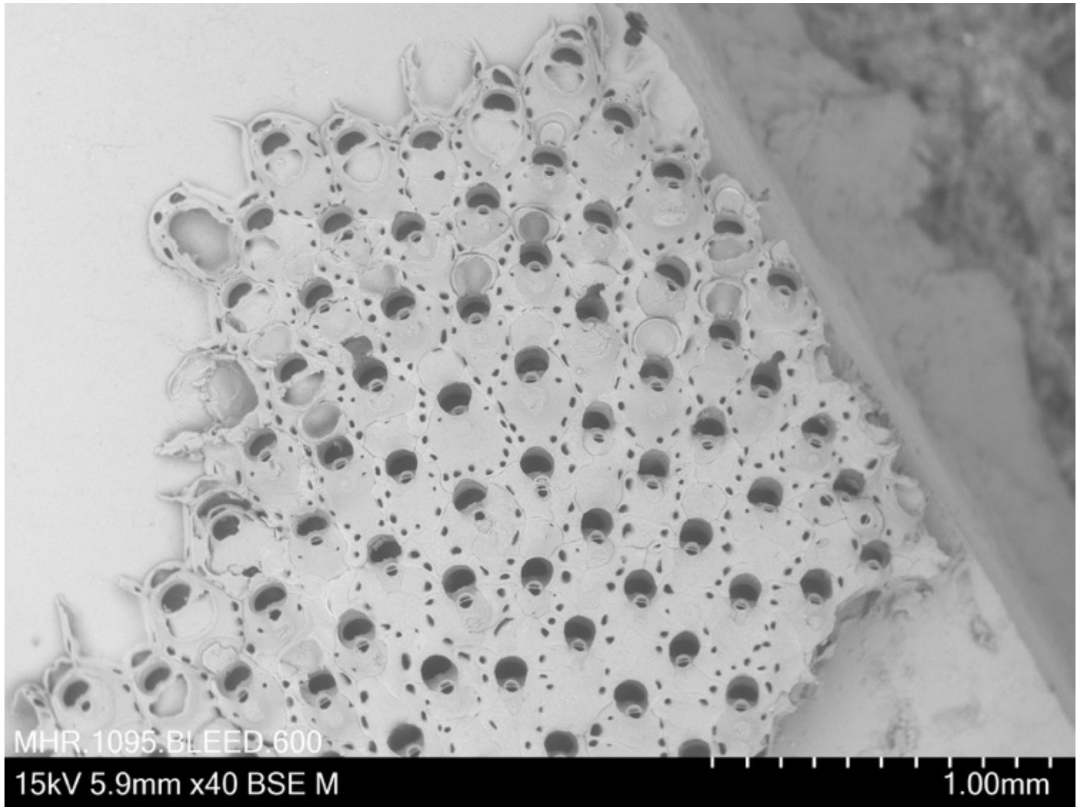


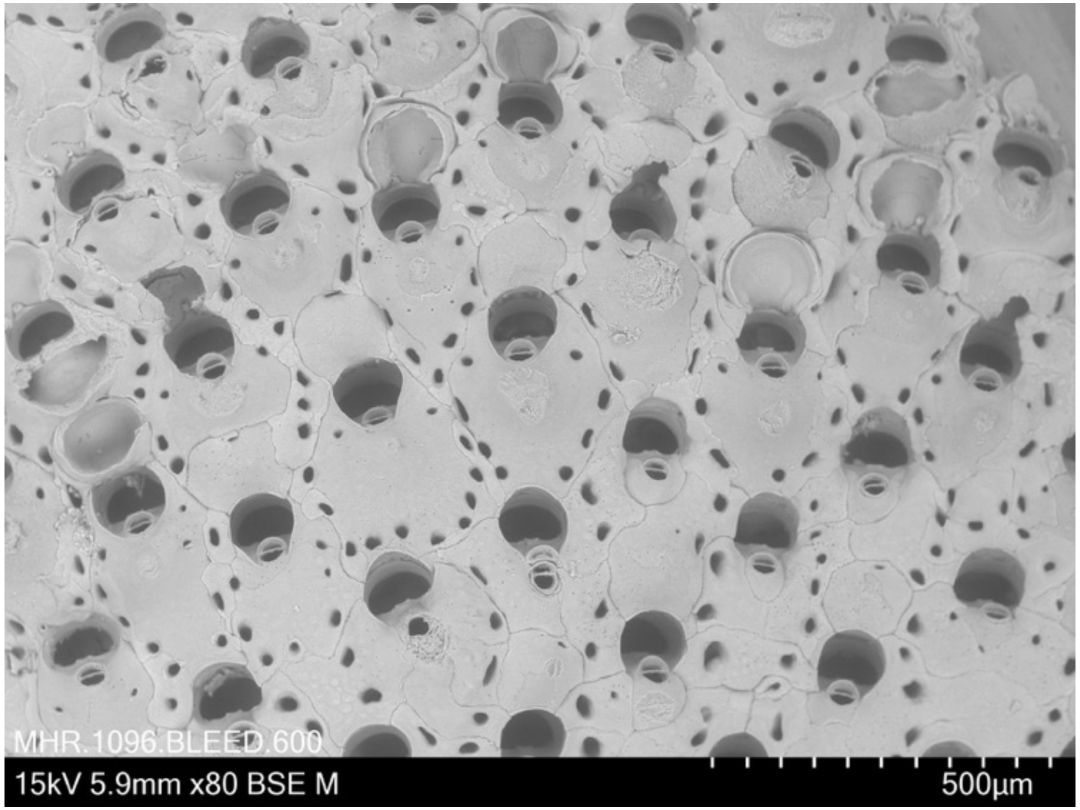


***Aimulosia marsupium*** **BLEED 600**
Coll: 11.4.2018 (A. M. Smith, D. P. Gordon, H. Mello)
Lat: -46.74417
Long: 168.3256
Depth: 29 m

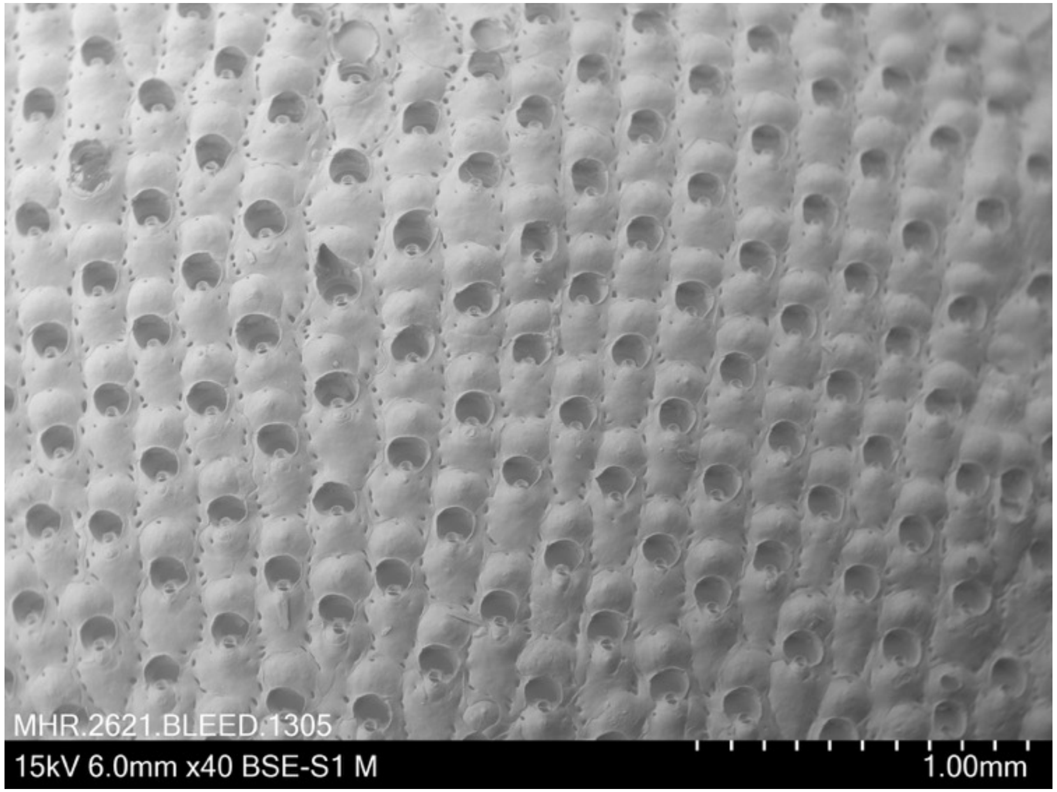


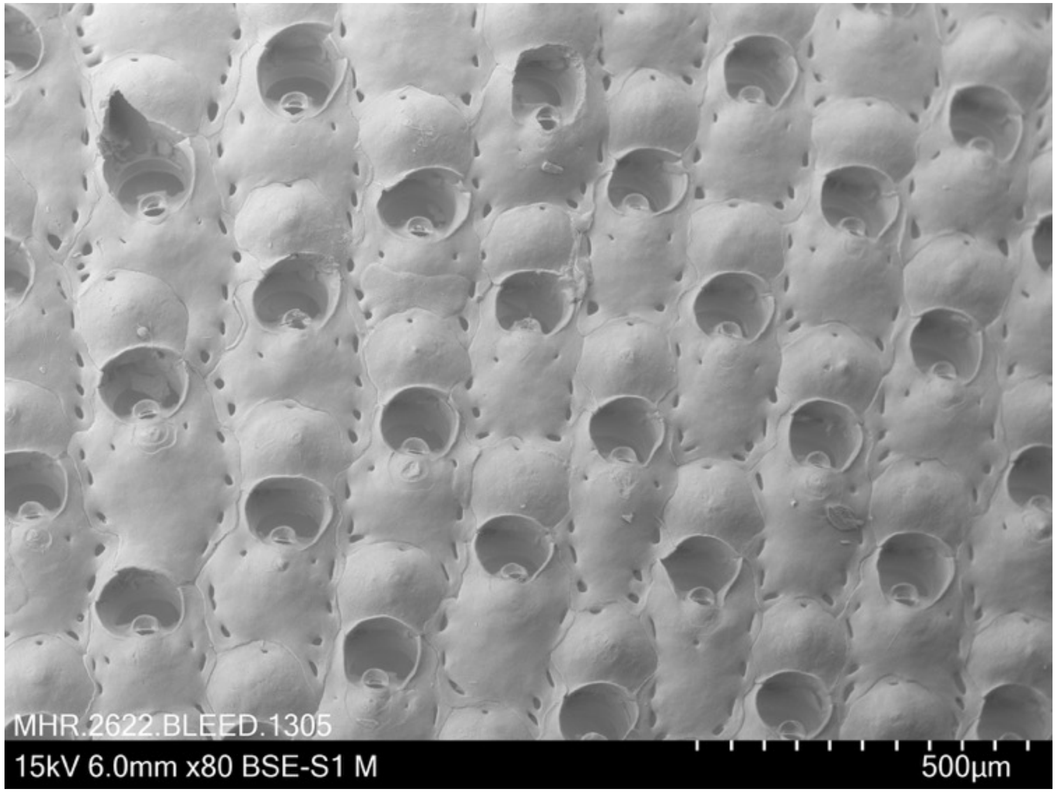


***Aimulosia marsupium* BLEED 1305**
Coll: 15.3.2019 (A. M. Smith)
Lat: -36.46
Long: 174.9
Depth: 37 m

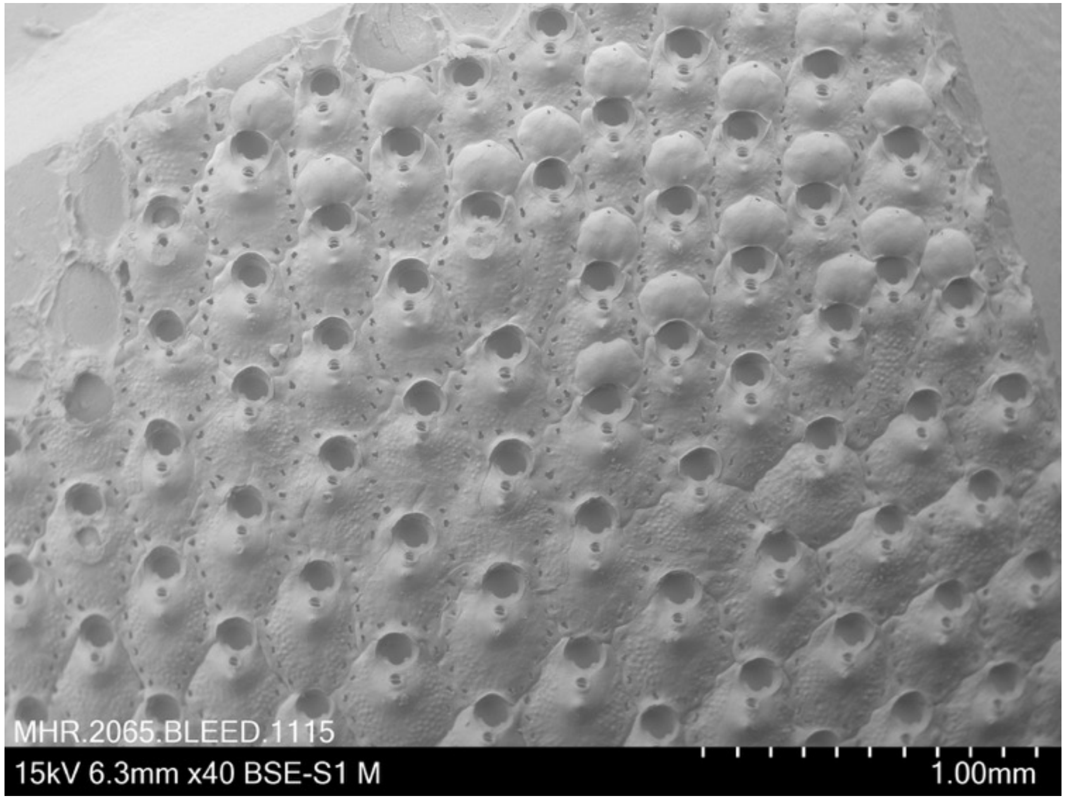


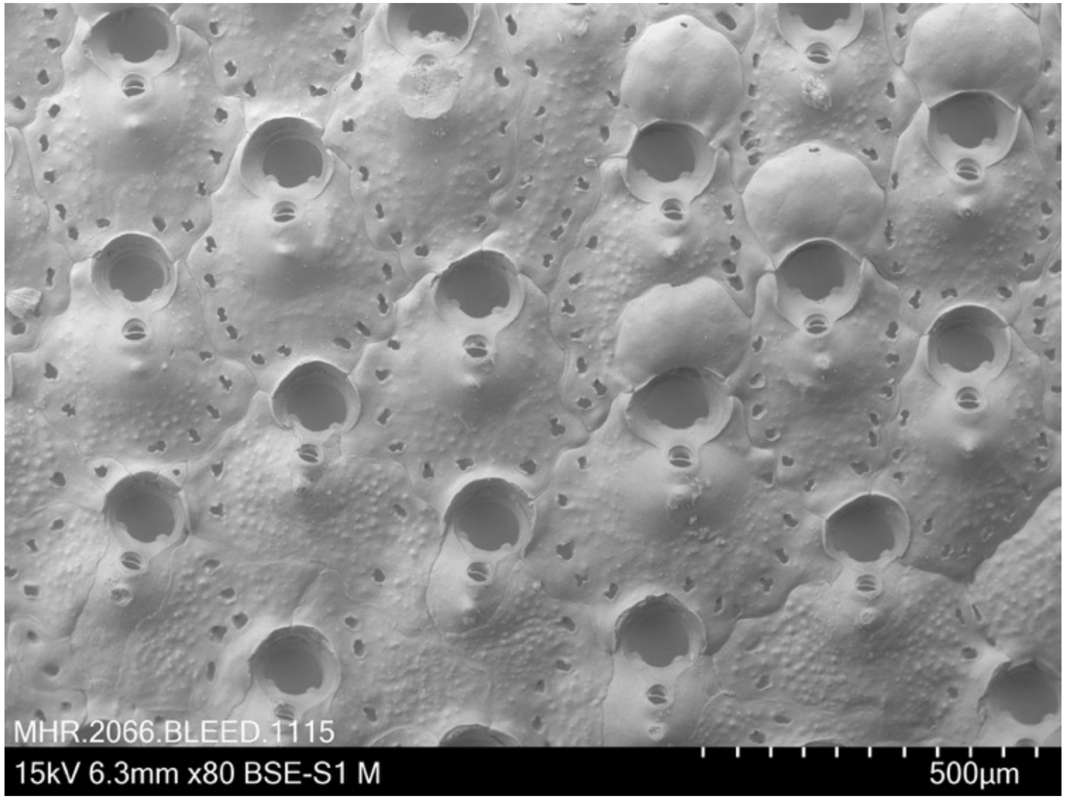


***Xenogma rhomboidale*** **BLEED 1115B**
Coll: 11.10.2018 (A. M. Smith)
Lat: -45.835000000000001
Long: 170.94916699999999
Depth: 111 m

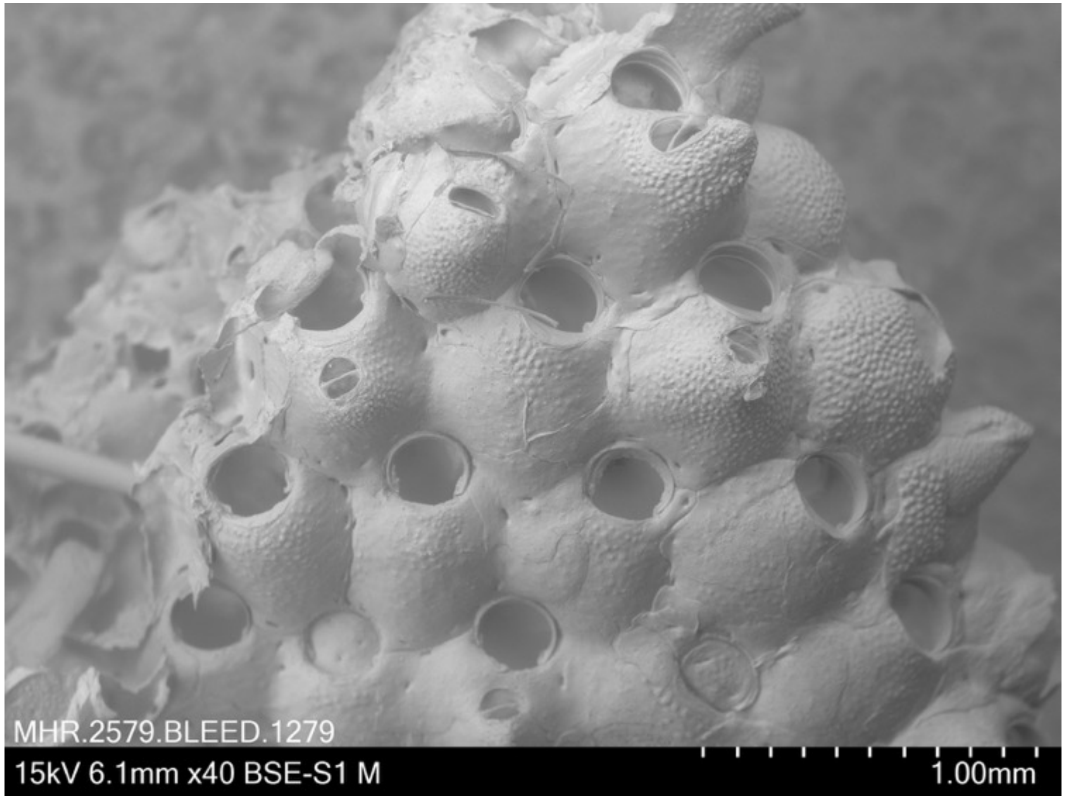


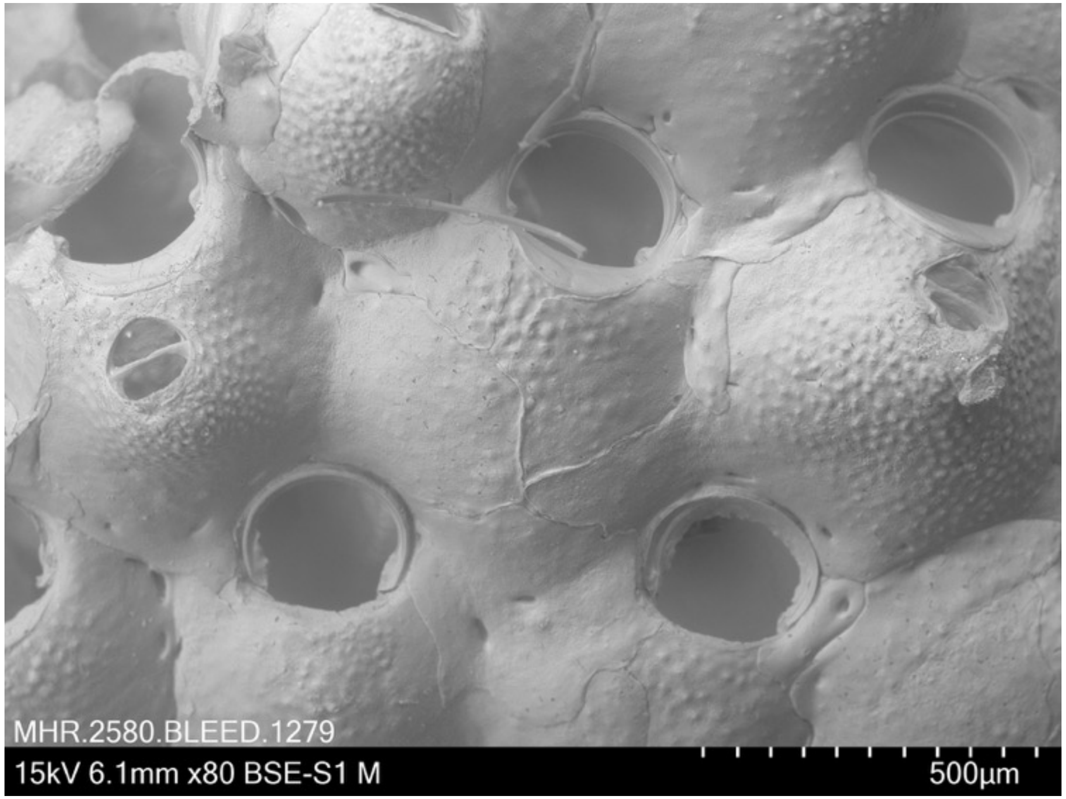


***Yrbozoon conspicuum* BLEED 1279**
Coll: 17.3.2019 (A. M. Smith, D. P. Gordon)
Lat: -34.17
Long: 172.04
Depth: 85.4 m

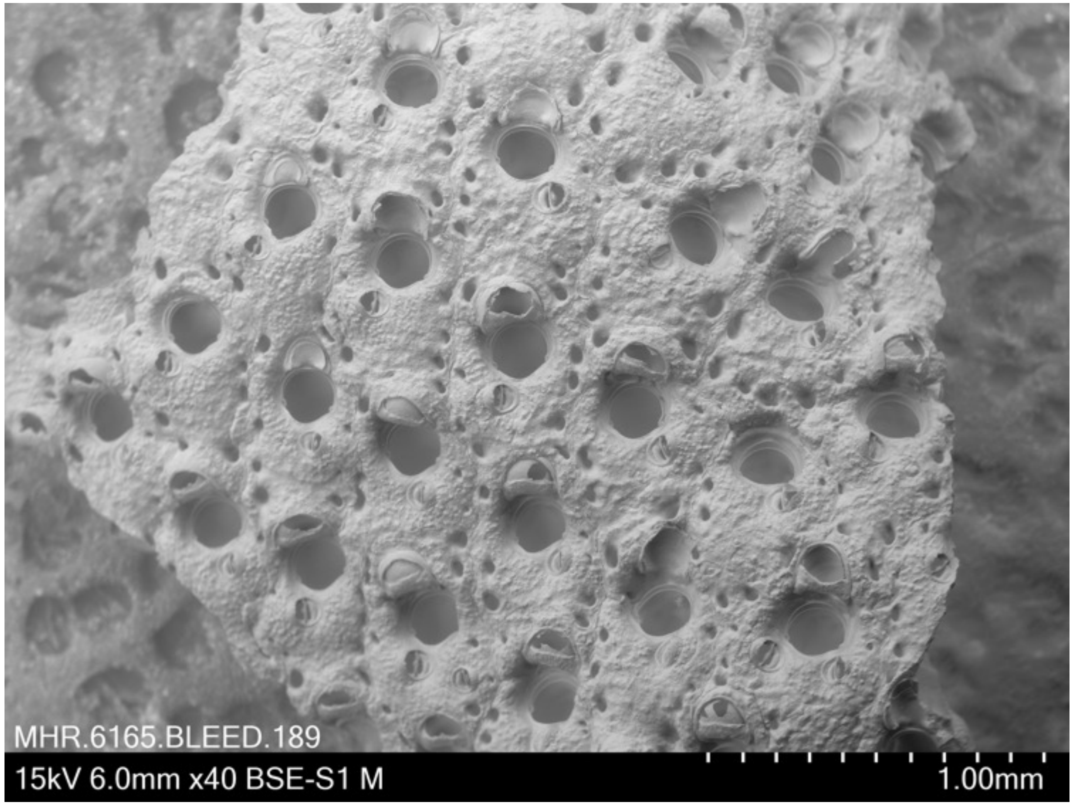


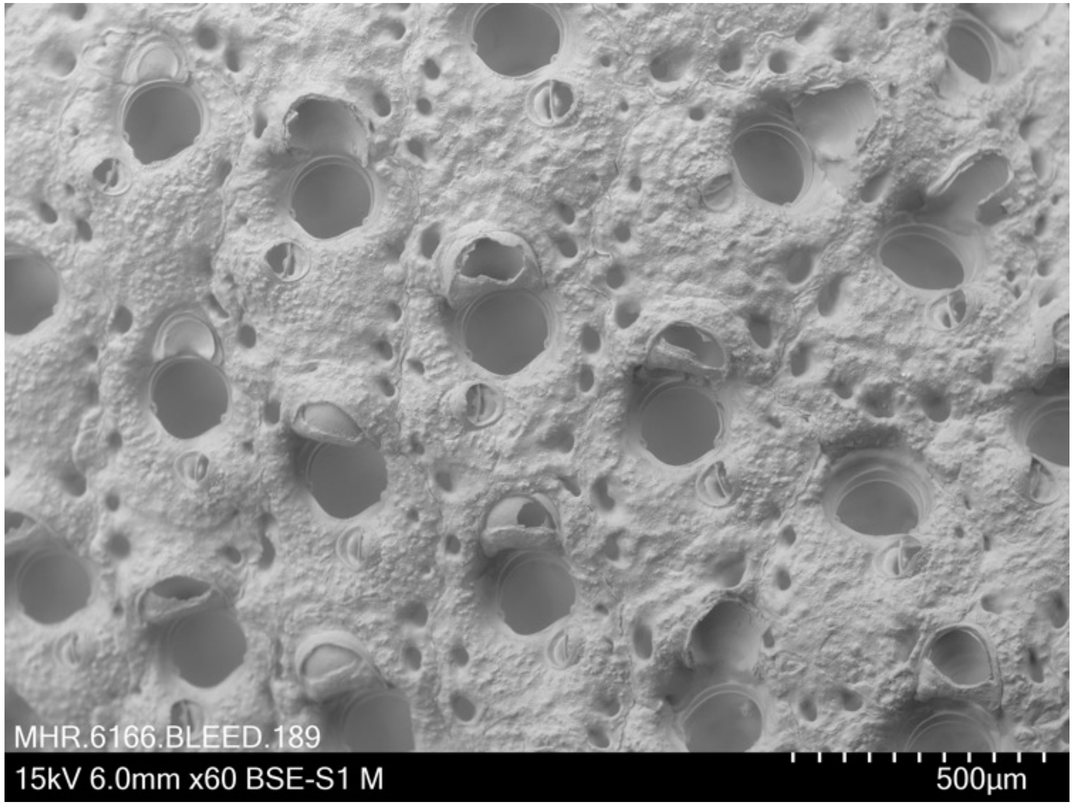


***Buffonellaria biavicularis* BLEED 189**
NIWA 56624 (TAN0906/159)
Coll: 14.0.2009
Lat: -34.388
Long: 173.041
Depth: 89 m

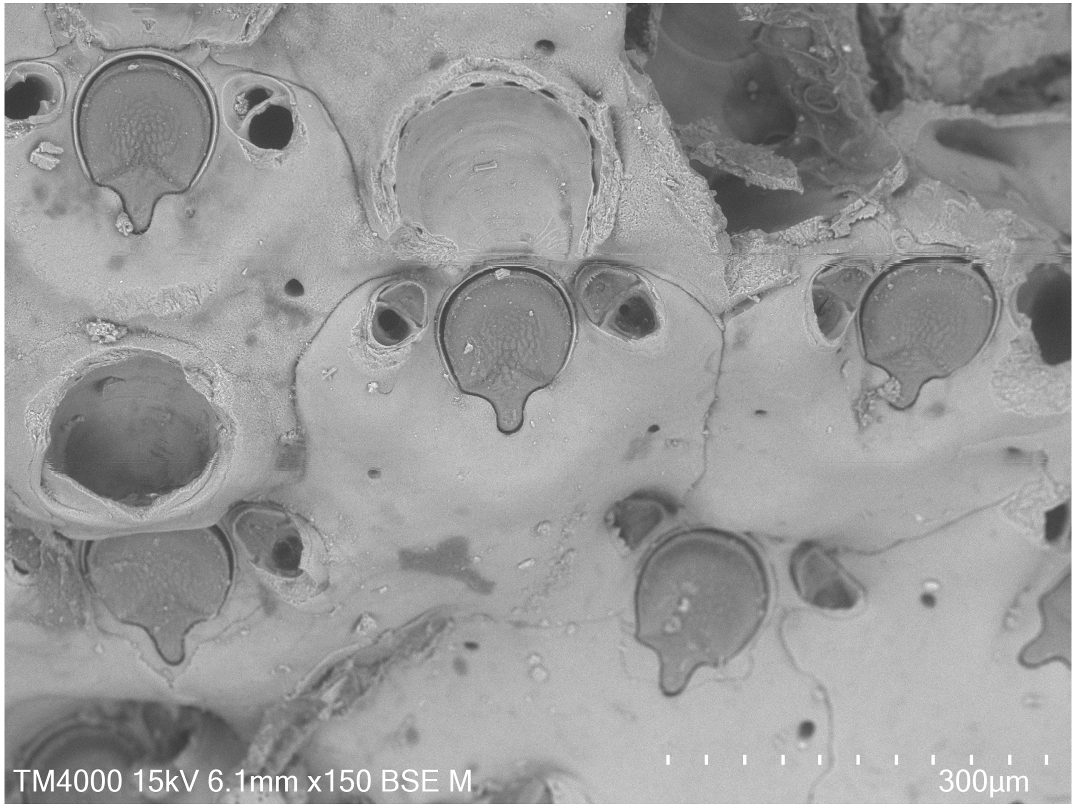


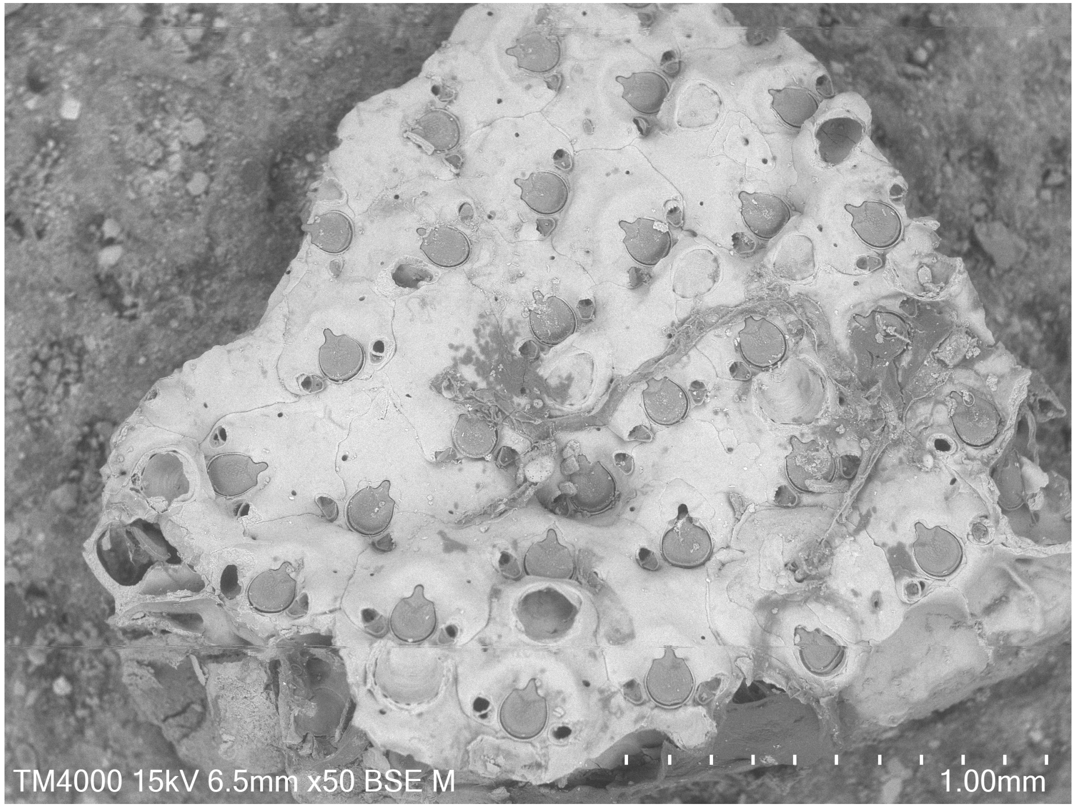


***Buffonellaria* cf. *regenerata* BLEED 88**
NIWA 98541 KAH0907/15
Coll: 11.8.2019
Lat: -35.18683
Long: 174.1173
Depth: 34 m

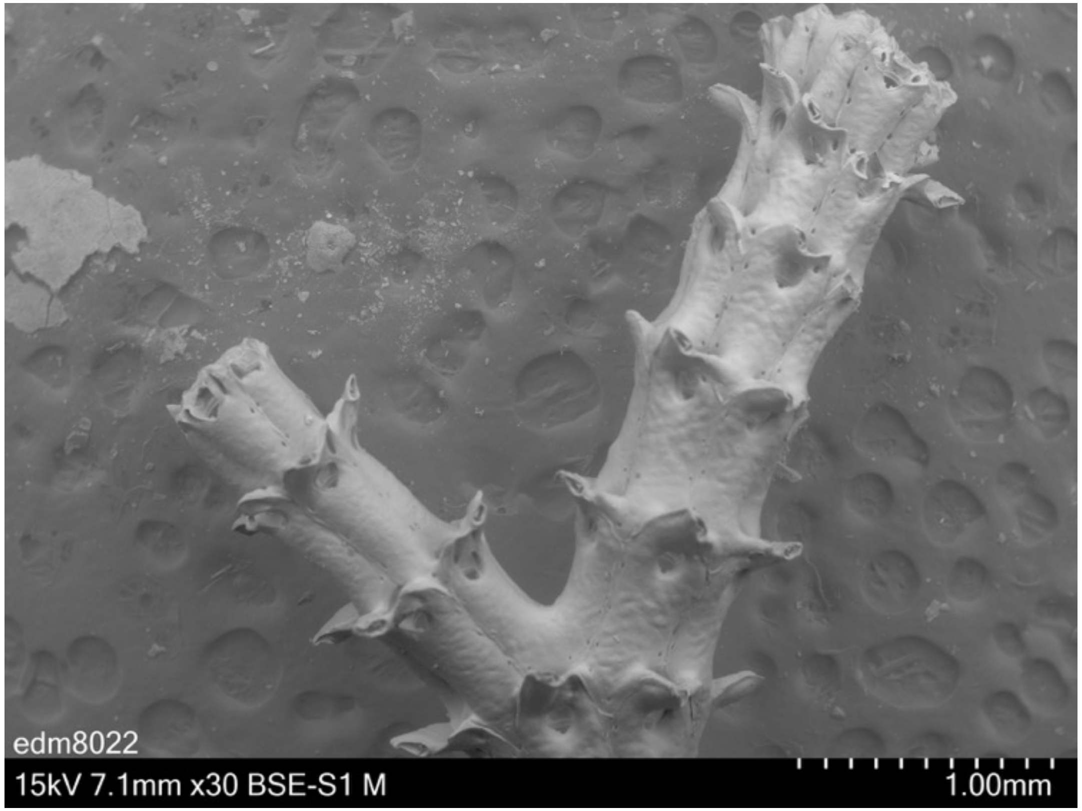


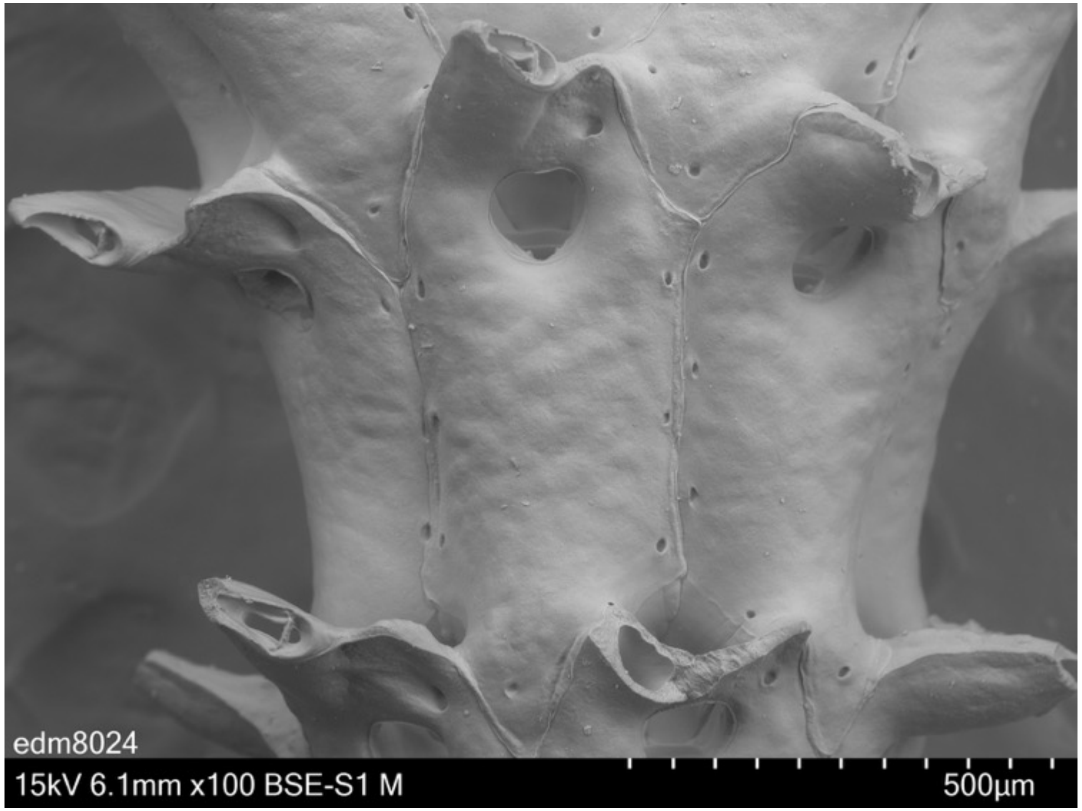


***Galeopsis* n. sp. 1 BLEED 1617**
Coll: 17.3.2019 (A. M. Smith, D. P. Gordon)
Lat: -34.16
Long: 172.09
Depth: 82.9 m

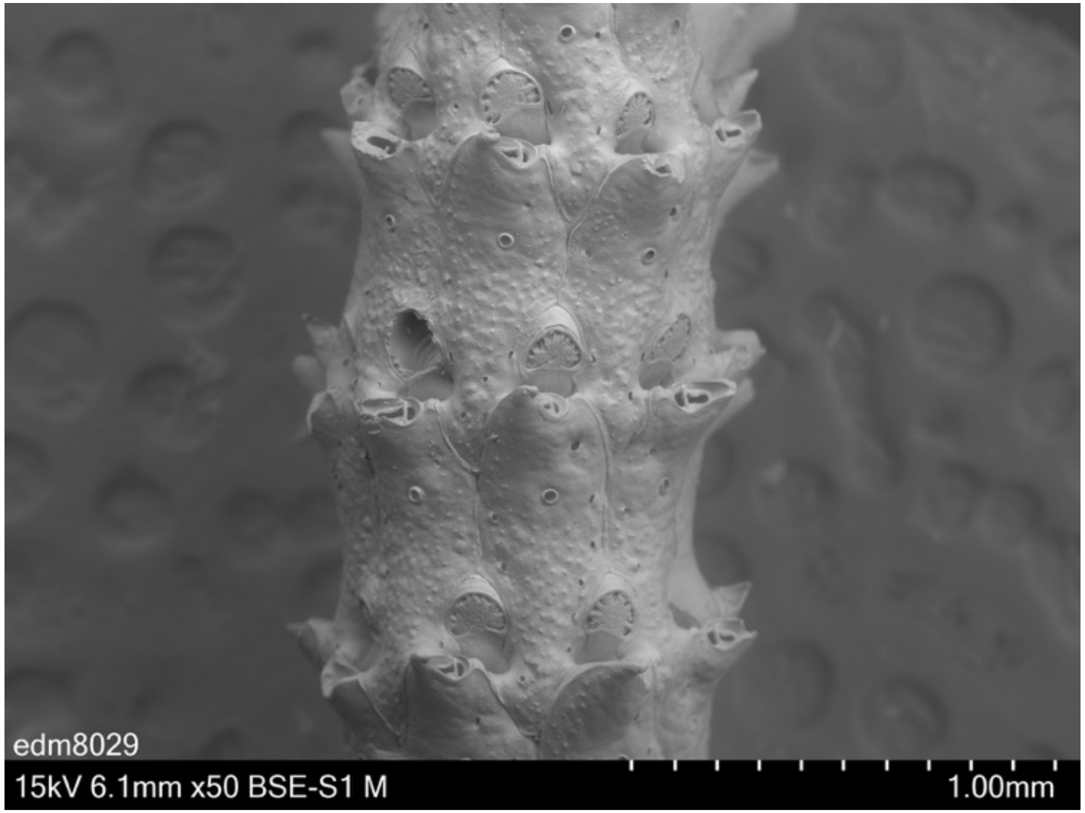

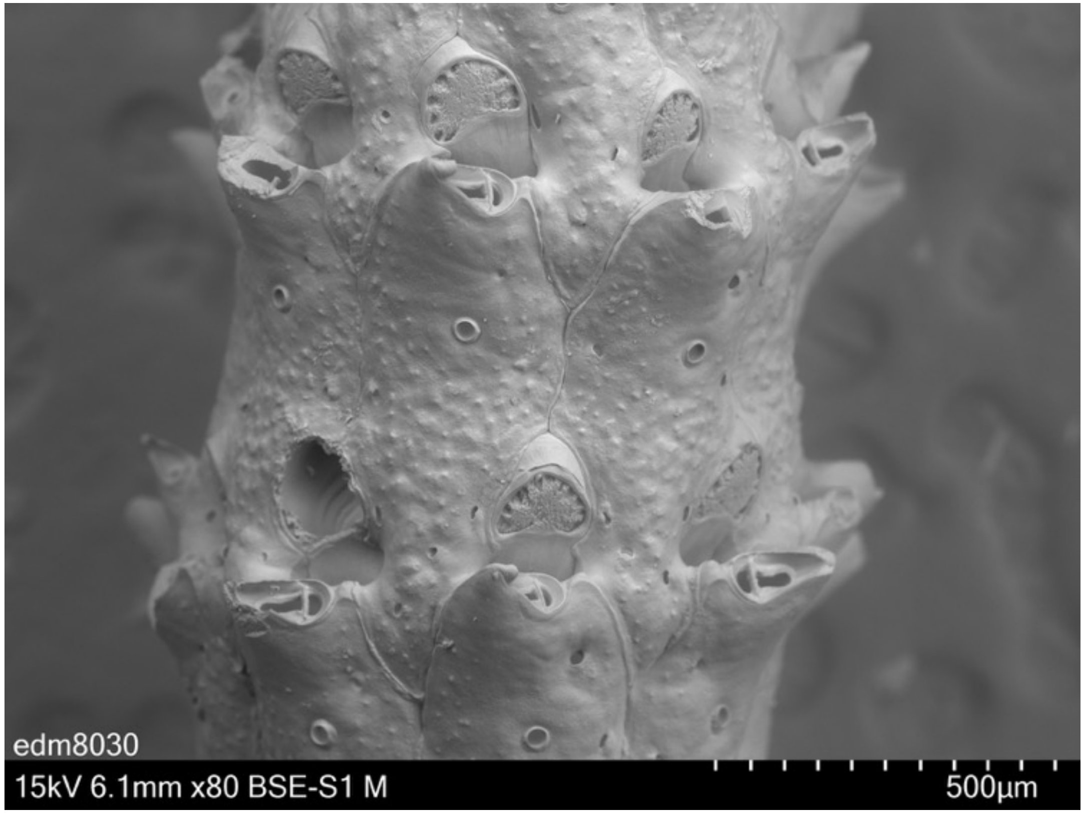


***Galeopsis*** **n. sp. 2 BLEED 1618**
Coll: 17.3. 2019 (A. M. Smith, D. P. Gordon)
Lat: -34.16
Long: 172.09
Depth: 82.9 m

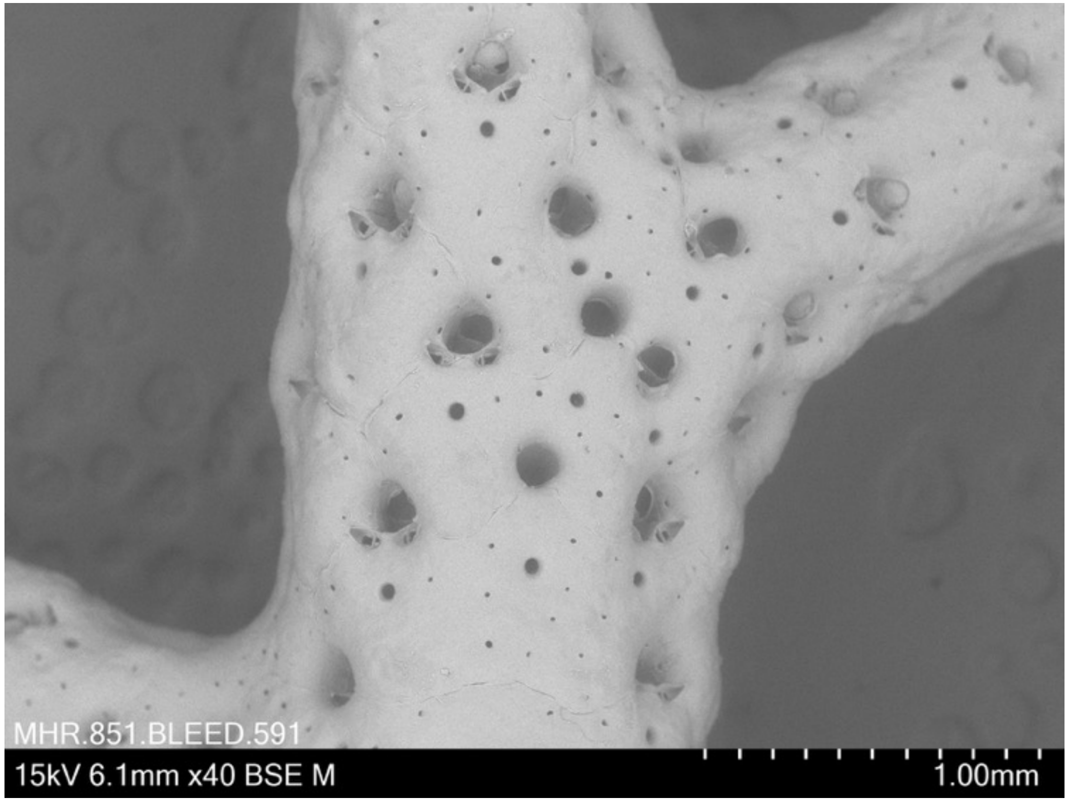


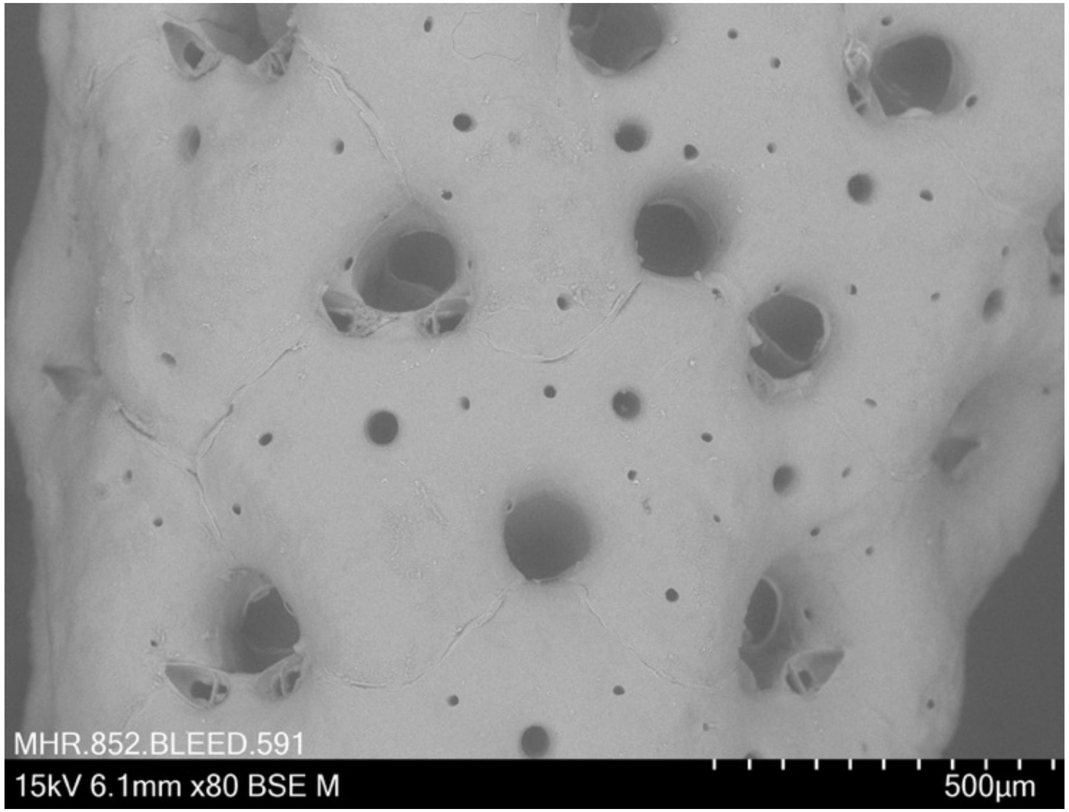


***Galeopsis porcellanicus*** **BLEED 591**
Coll: 10.4.2018 (A. M. Smith, D. P. Gordon, H. Mello)
Lat: -46.92417
Long: 168.2183
Depth: 58 m

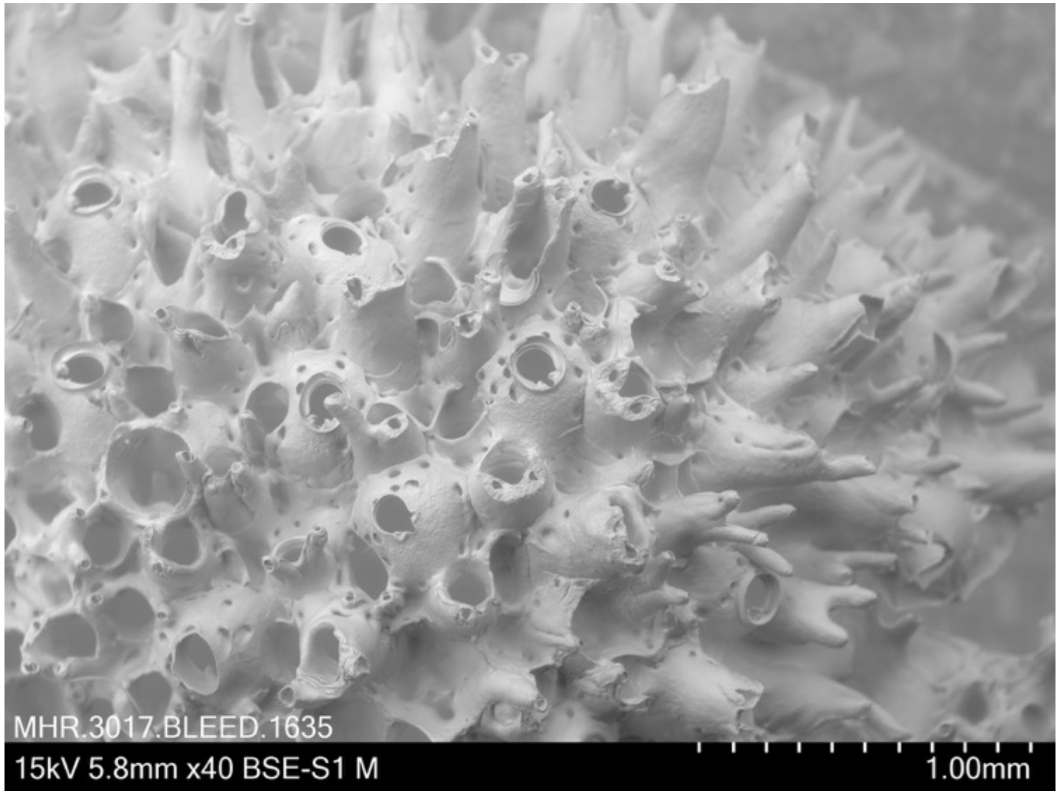


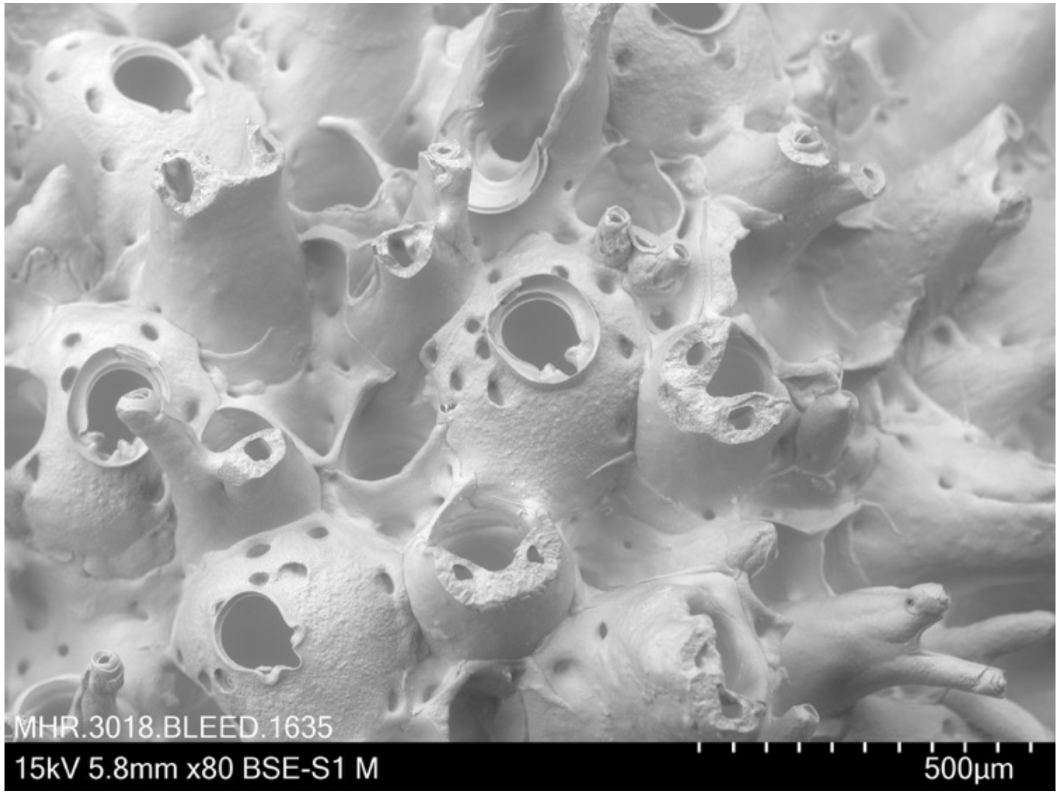


***Osthimosia* n. sp. BLEED 1635**
Coll: 17.3.2019 (A. M. Smith, D. P. Gordon)
Lat: -34.159999999999997
Long: 172.09
Depth: 82.9 m

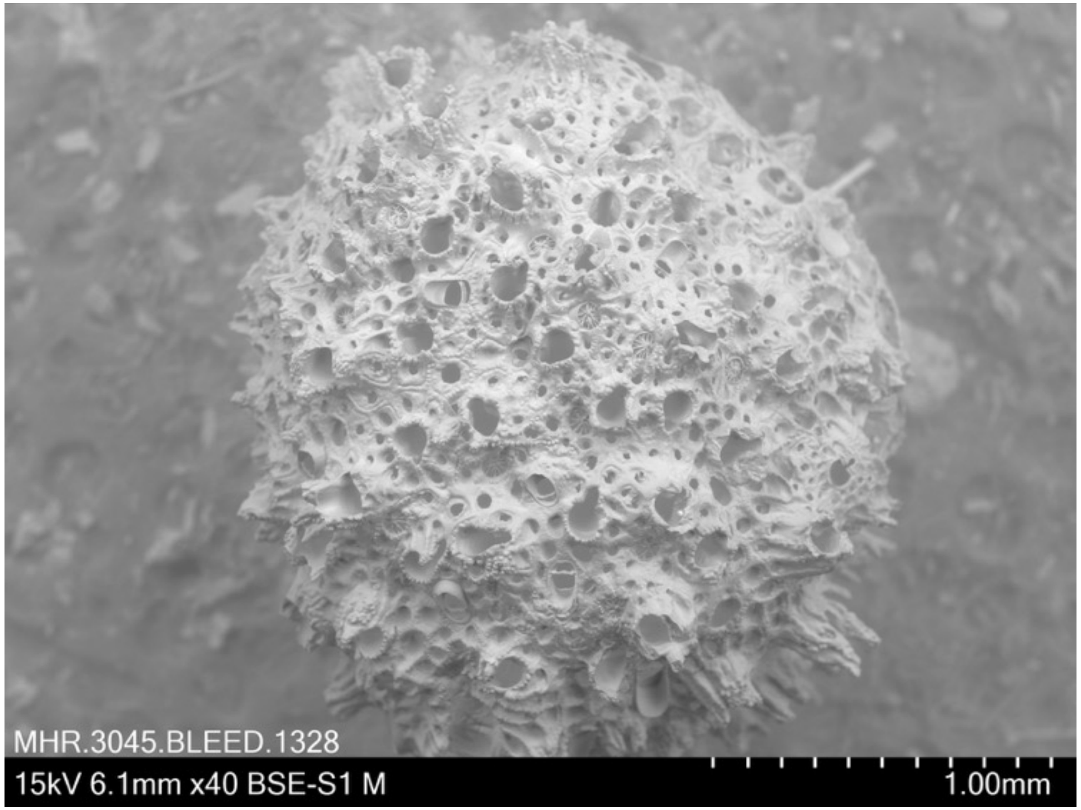


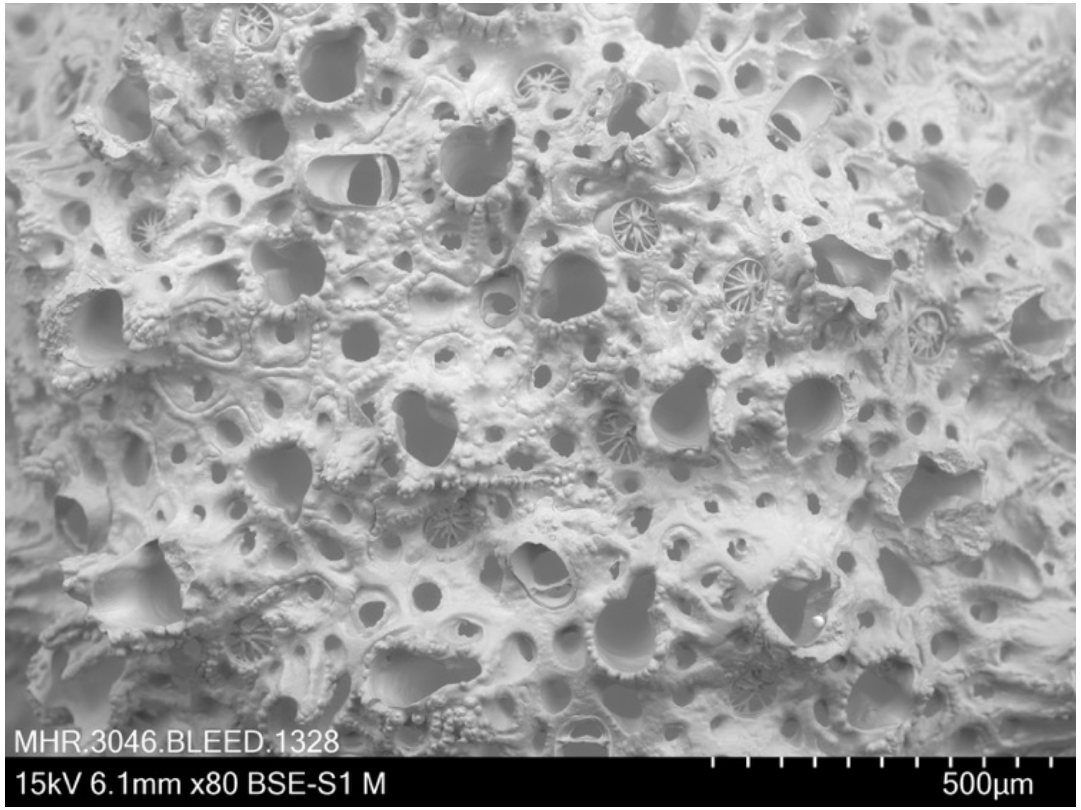


***Celleporina* cf. *cribrillifera* BLEED 1328**
Coll: 17.3.2019 (A. M. Smith, D. P. Gordon)
Lat: -34.12
Long: 172.21
Depth: 241 m

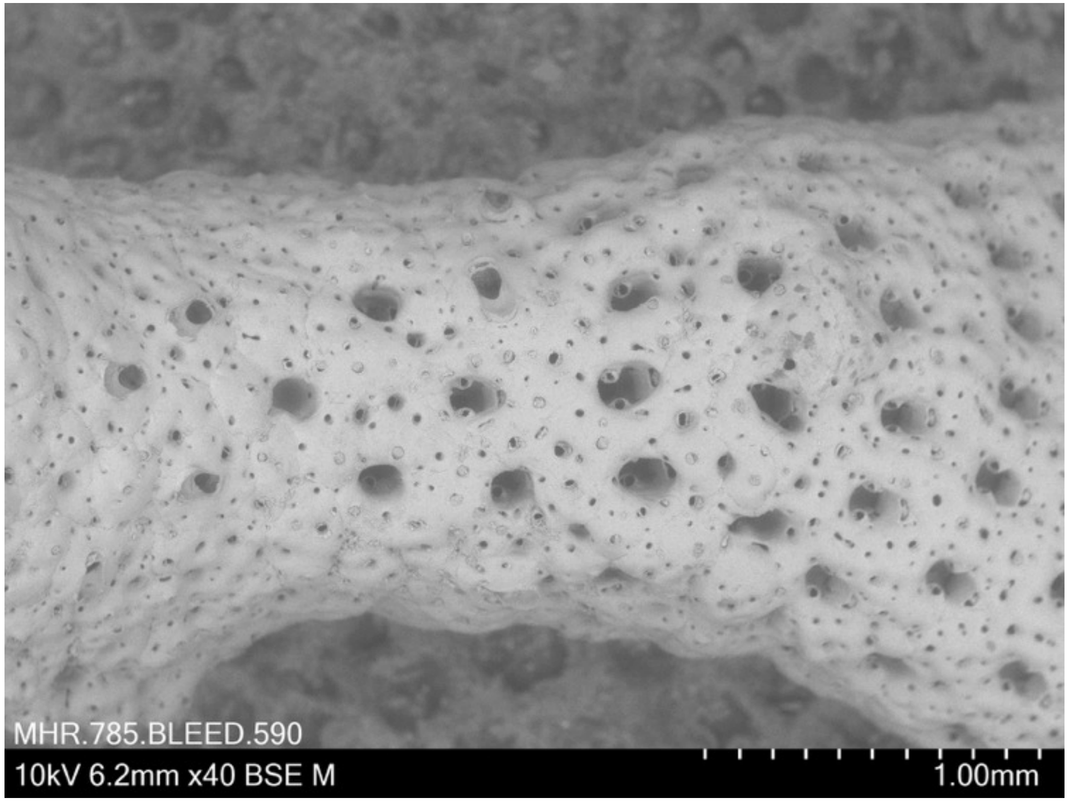


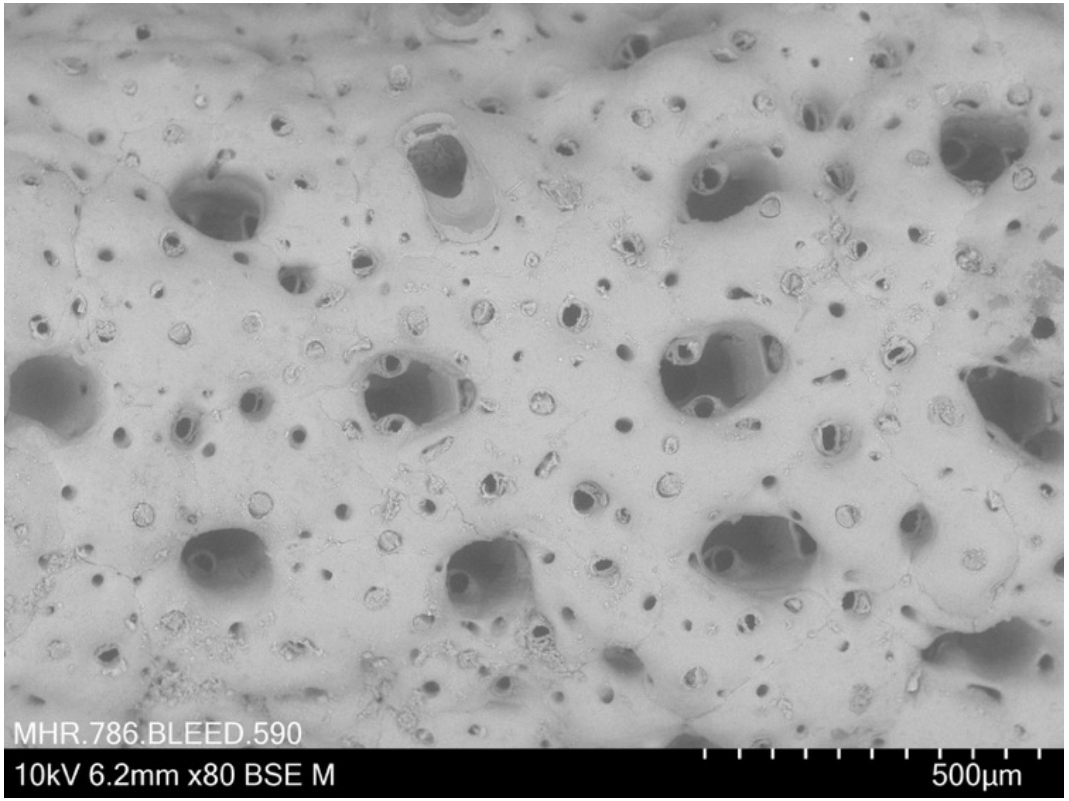


***Galeopsis polyporus* BLEED 590**
Coll: 10.4.2018 (A. M. Smith, D. P. Gordon, H. Mello)
Lat: -46.92417
Long: 168.2183
Depth: 58 m

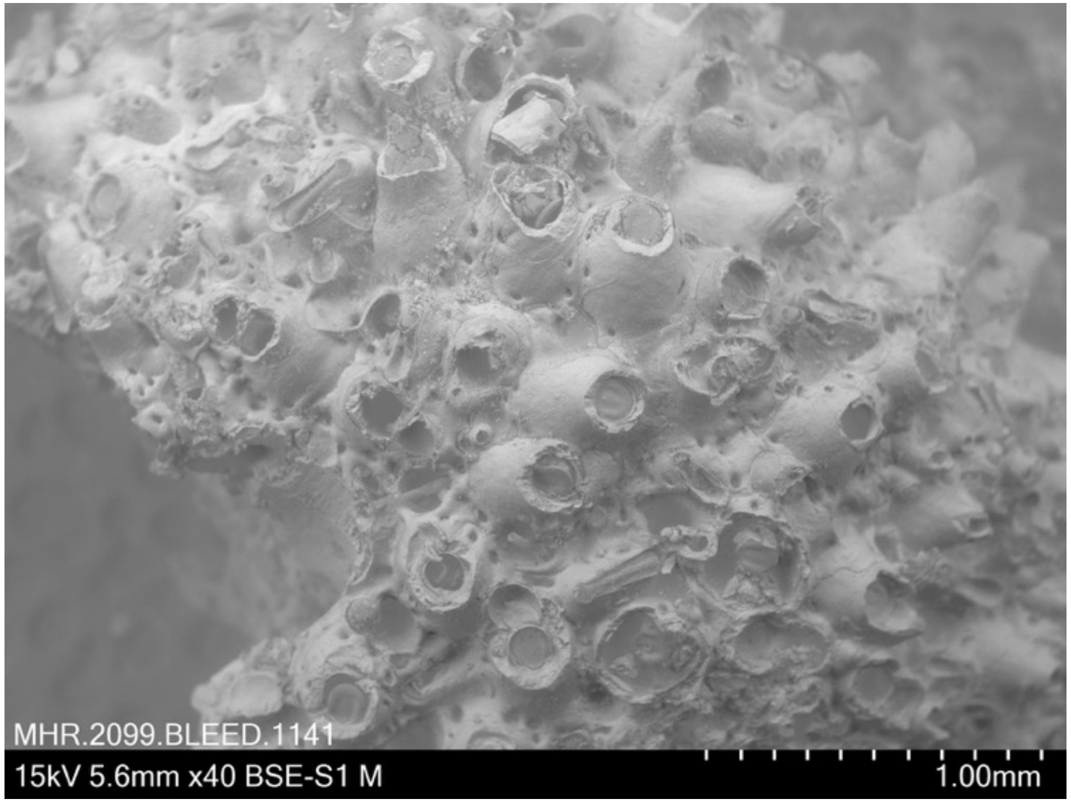


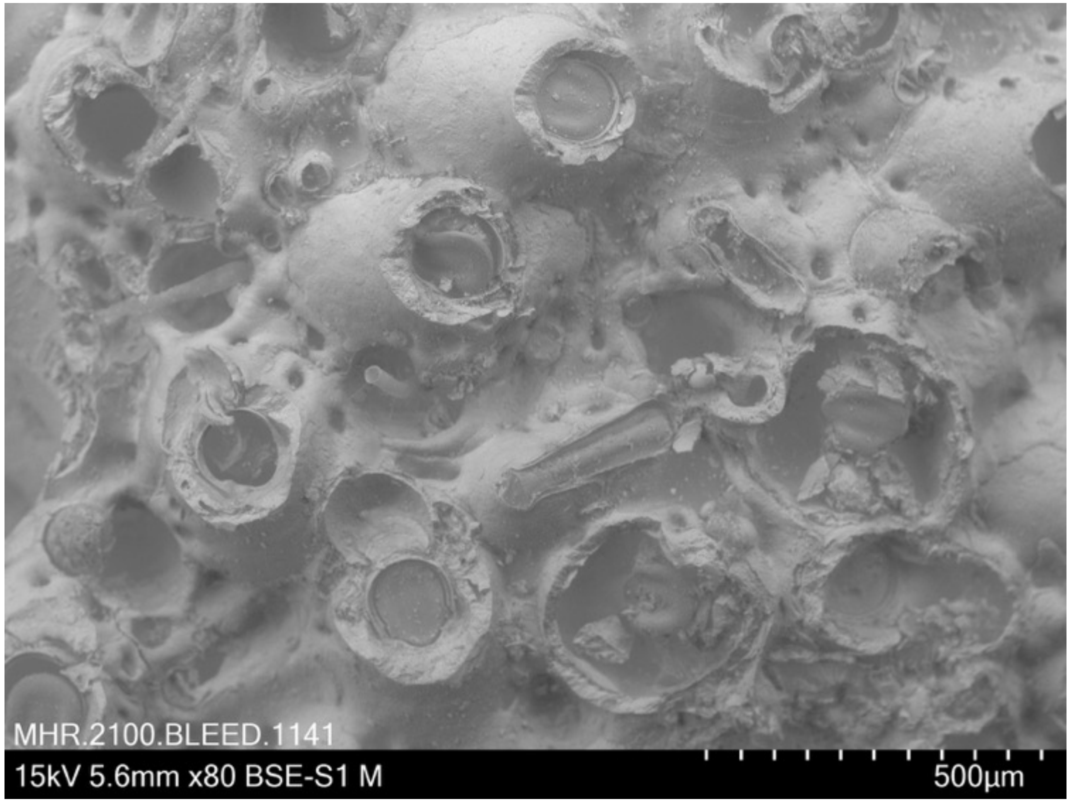


***Osthimosia* cf. *avicularis* BLEED 1141**
Coll: 10.11.2018 (A. M. Smith)
Lat: -45.7856
Long: 170.9219
Depth: 80.1 m

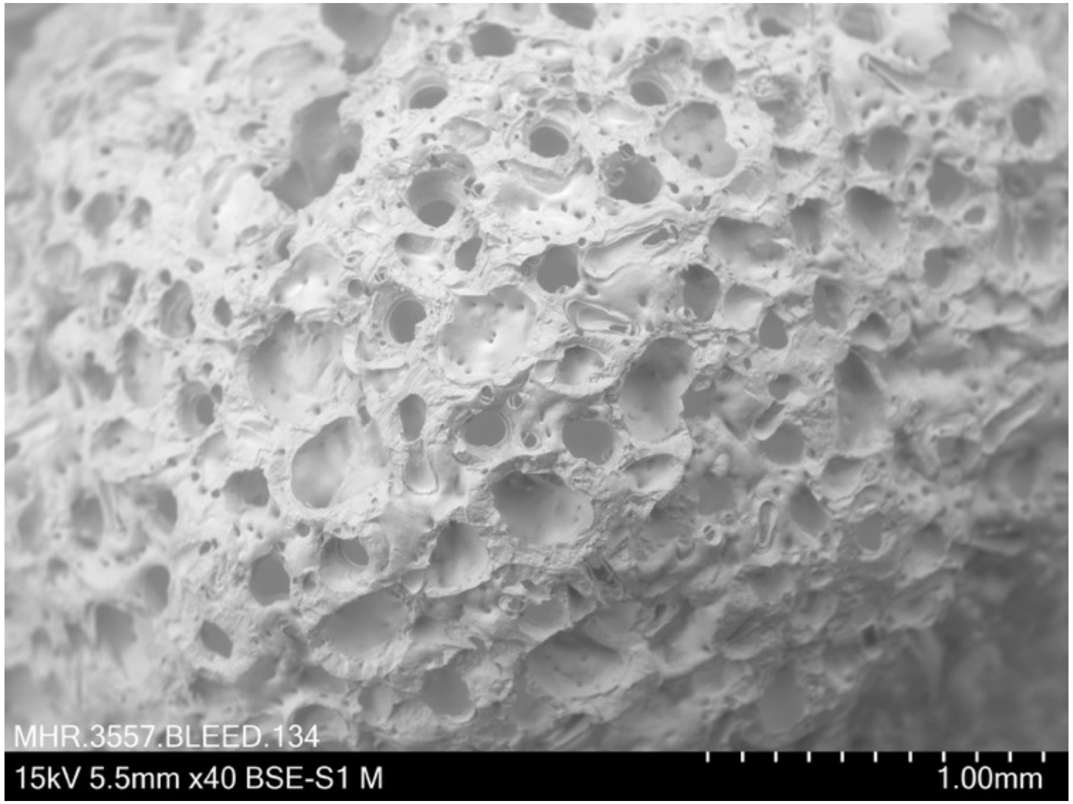


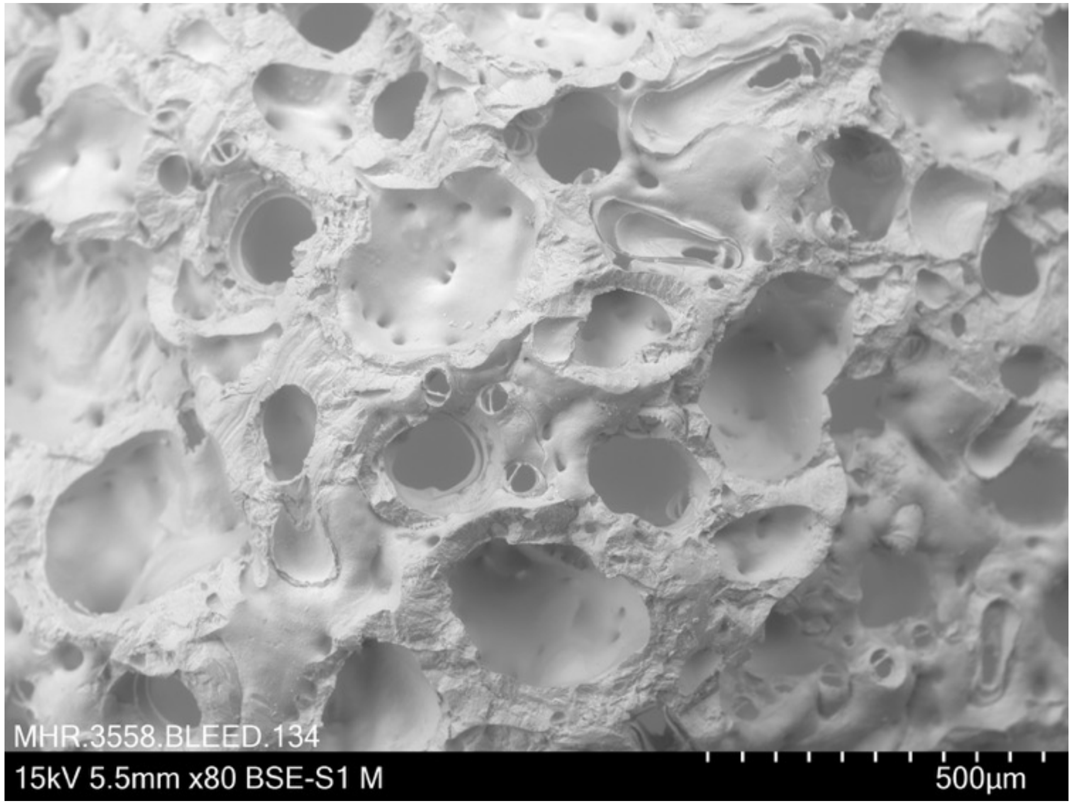


***Osthimosia* cf. *avicularis* BLEED 134**
NIWA 69631 (TRIP3298/36)
Coll: 12.4.2011
Lat: -46.283332999999999
Long: 166.133333
Depth: 270 m

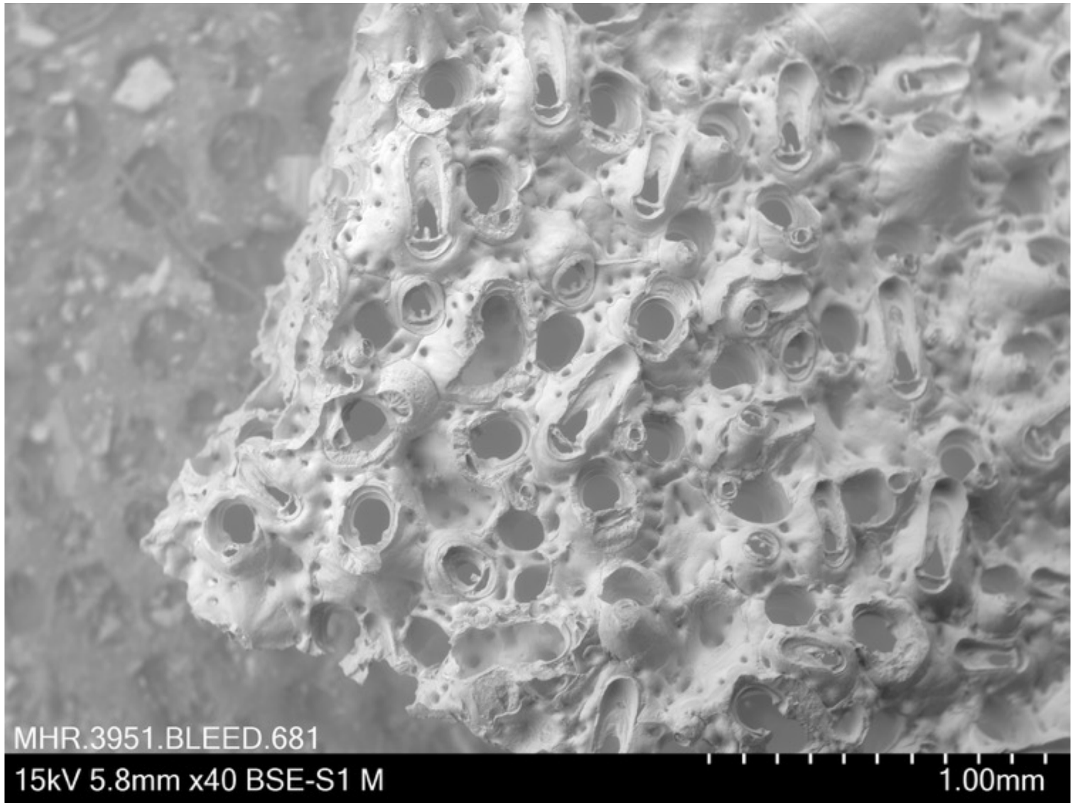


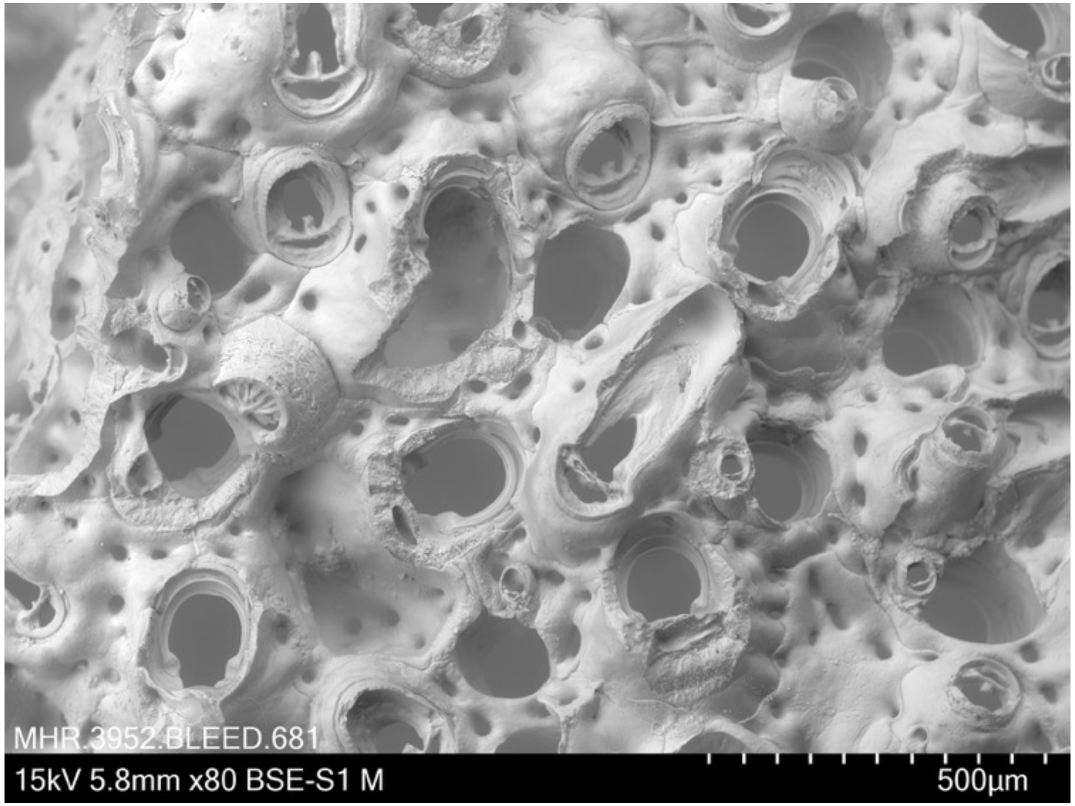


***Celleporina* n. sp. BLEED 681**
Coll: 8.4.2018 (A. M. Smith, D. P. Gordon, H. Mello)
Lat: -47.1361
Long: 169.189
Depth: 77 m

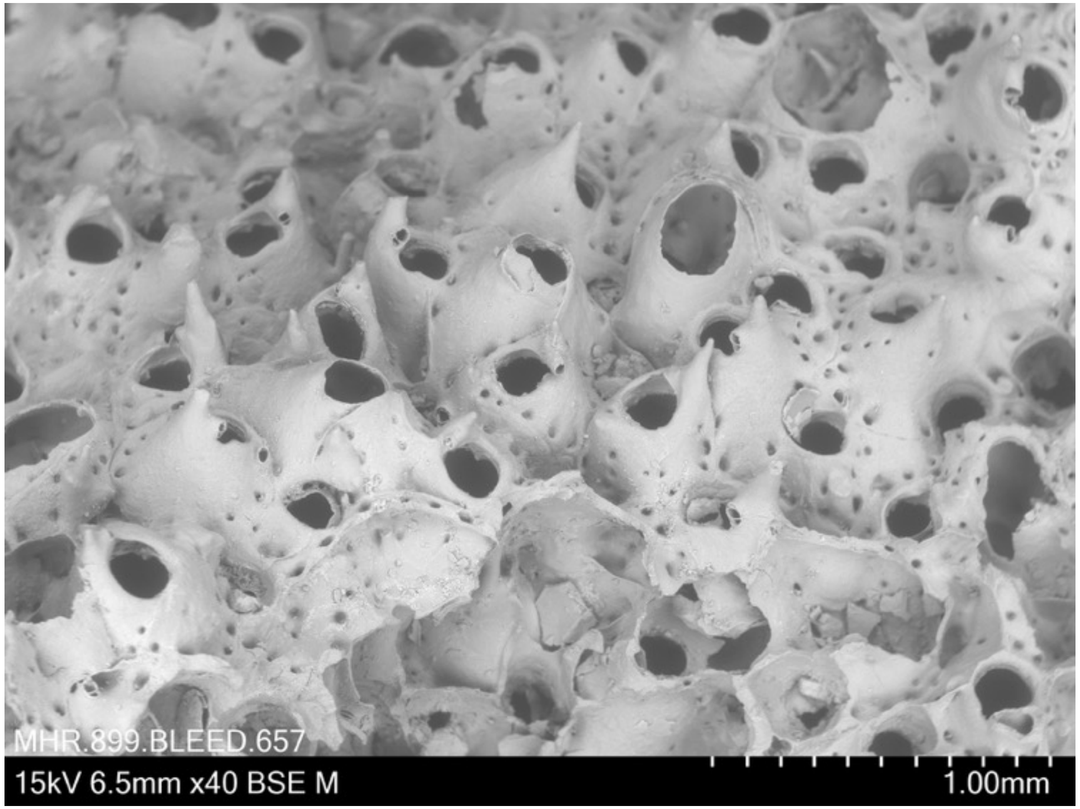


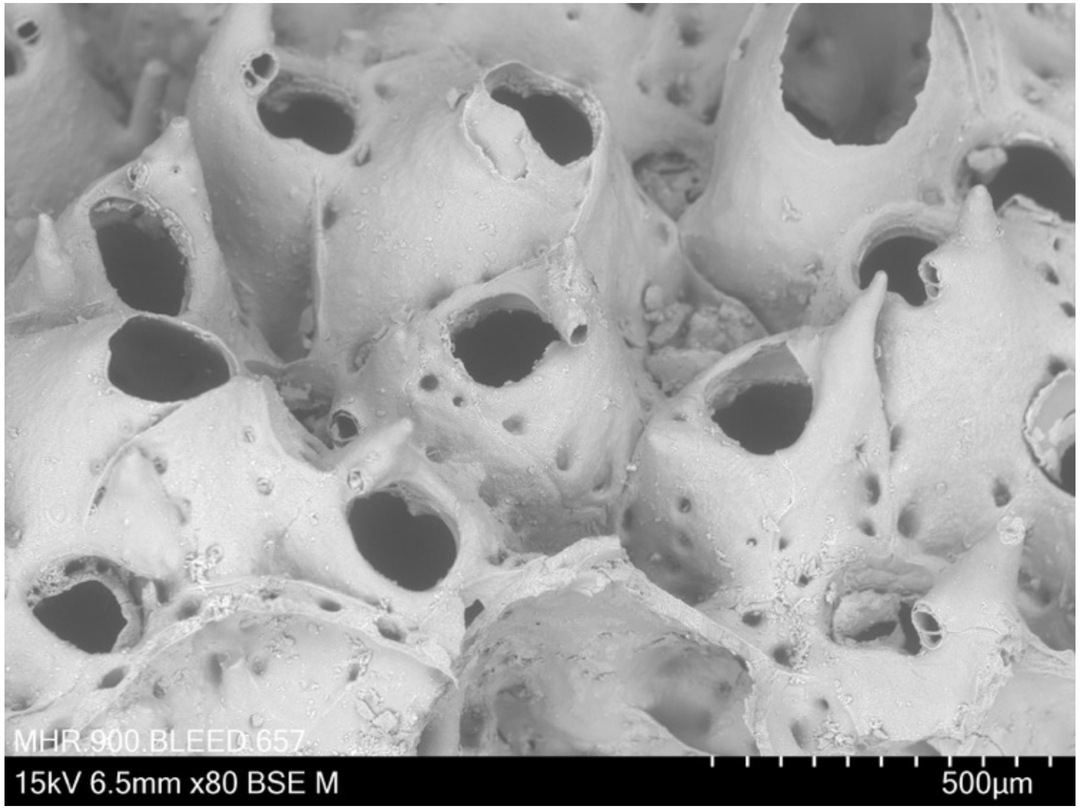


***Celleporaria agglutinans*** **BLEED 657**
Coll: 8.4.2018 (A. M. Smith, D. P. Gordon, H. Mello)
Lat: -47.17361
Long: 167.94583
Depth: 66 m

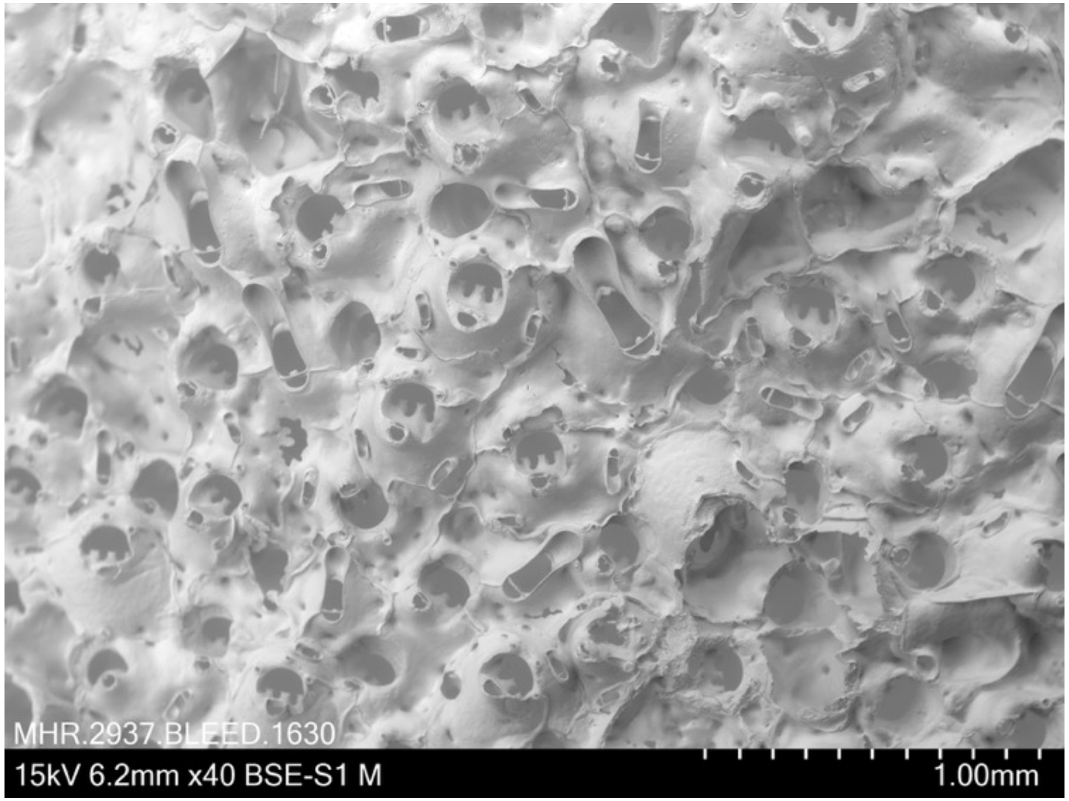


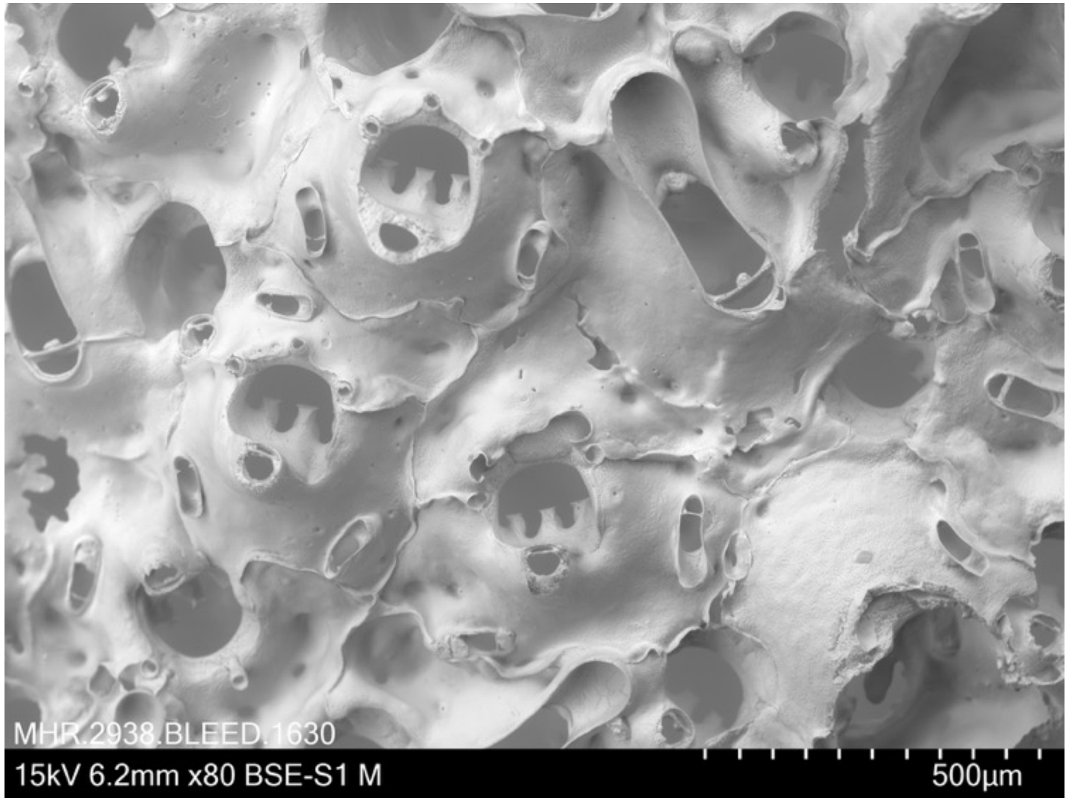


***Celleporaria macrodon*** **BLEED 1630**
Coll: 17.3.2019 (A. M. Smith; D.P. Gordon)
Lat: -34.16
Long: 172.09
Depth: 82.9 m

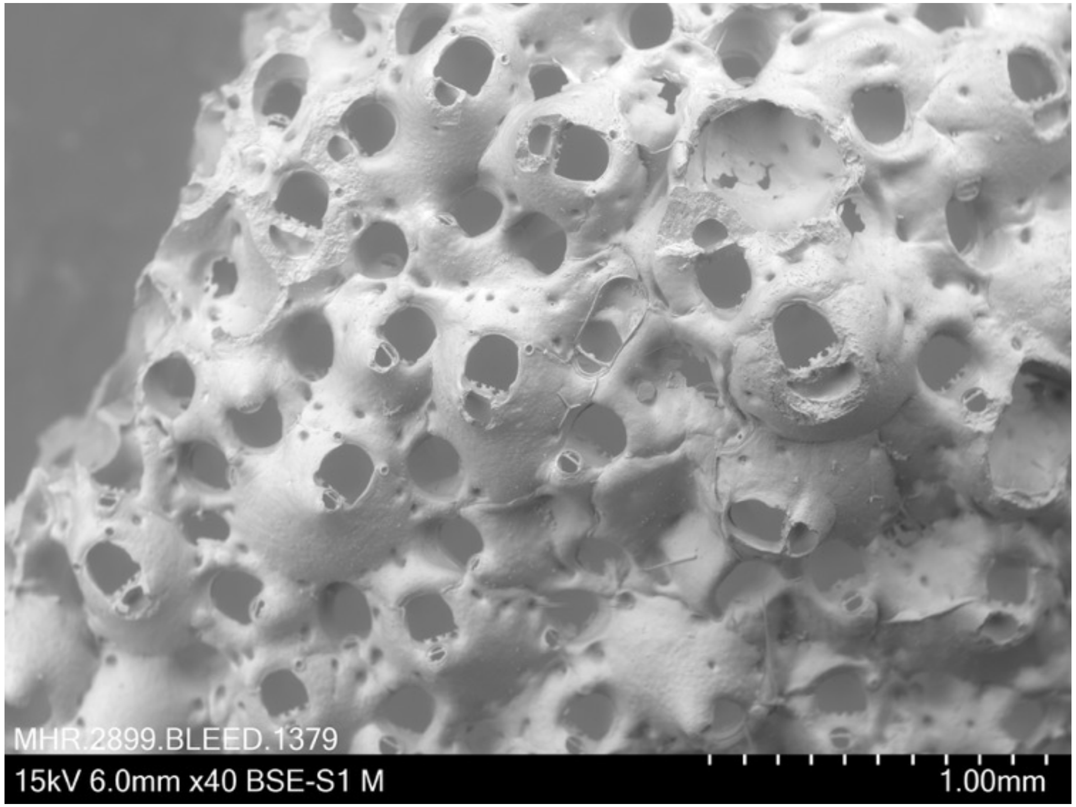


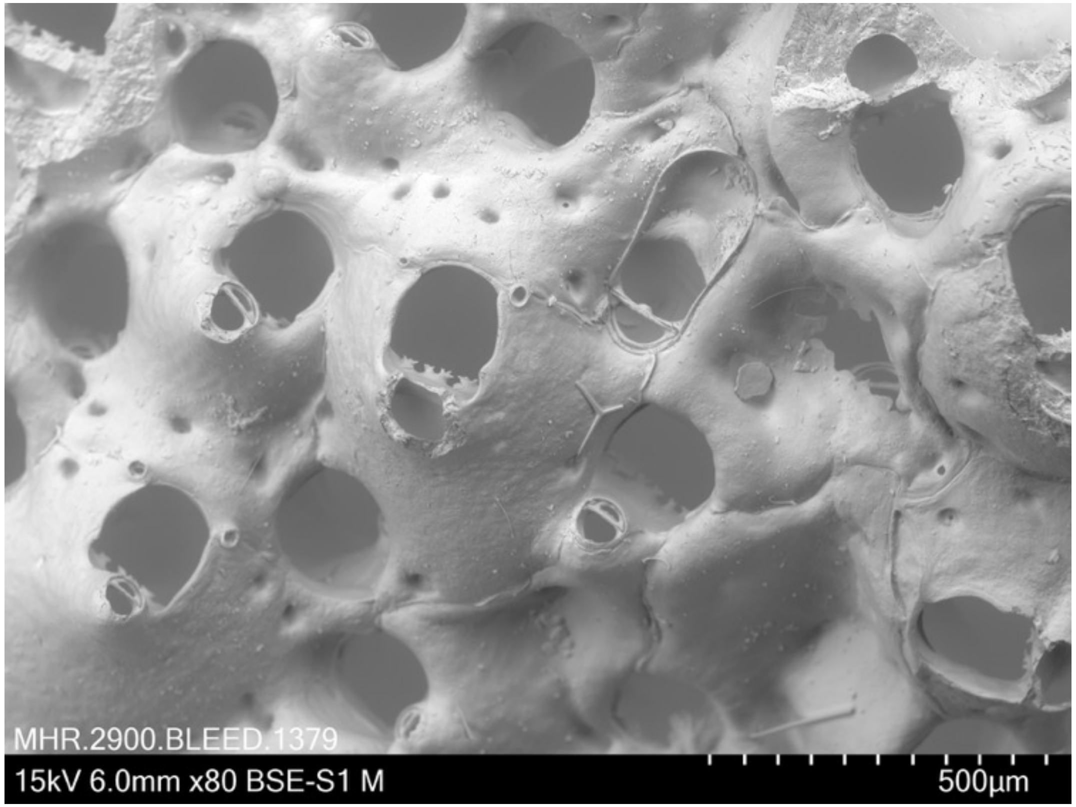


***Celleporaria* cf. *tridenticulata* BLEED 1379**
Coll: 17.3.2019 (A. M. Smith, D. P. Gordon)
Lat: -34.16
Long: 172.09
Depth: 82.9 m

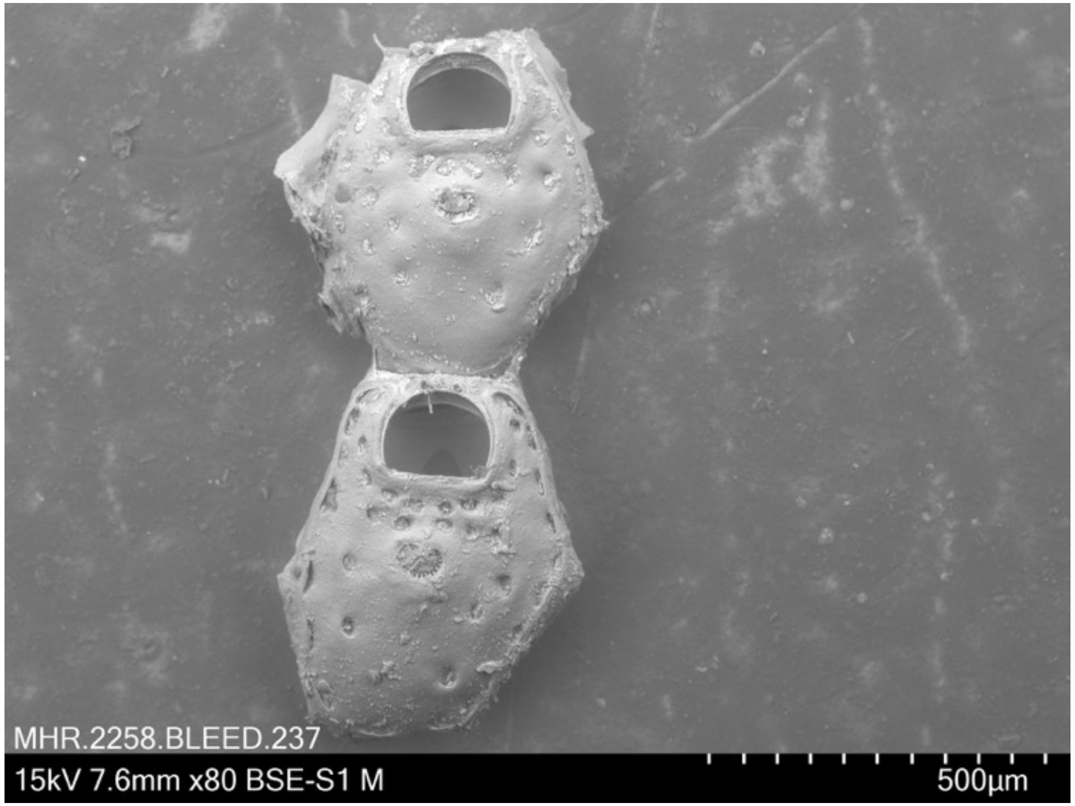


***Fenestrulina* n. sp. 1 BLEED 237**
Coll: 22.3.2009 (A. M. Smith)
Lat: -52.51
Long: 169.28
Depth: 91 m

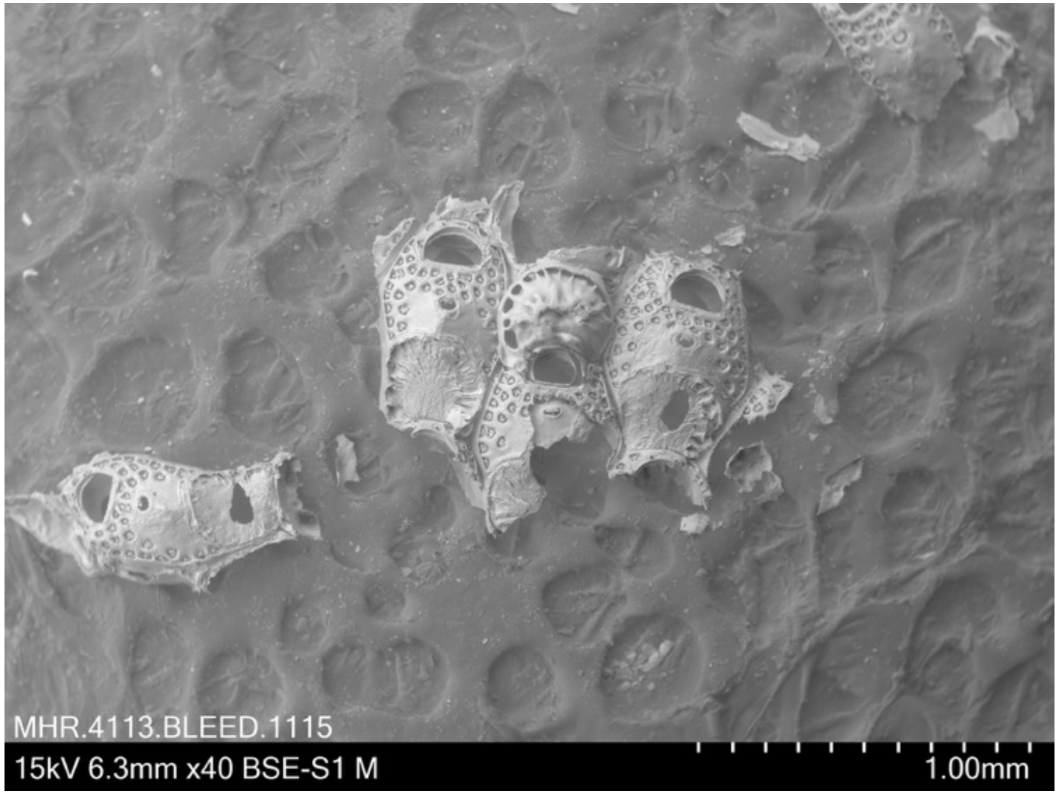


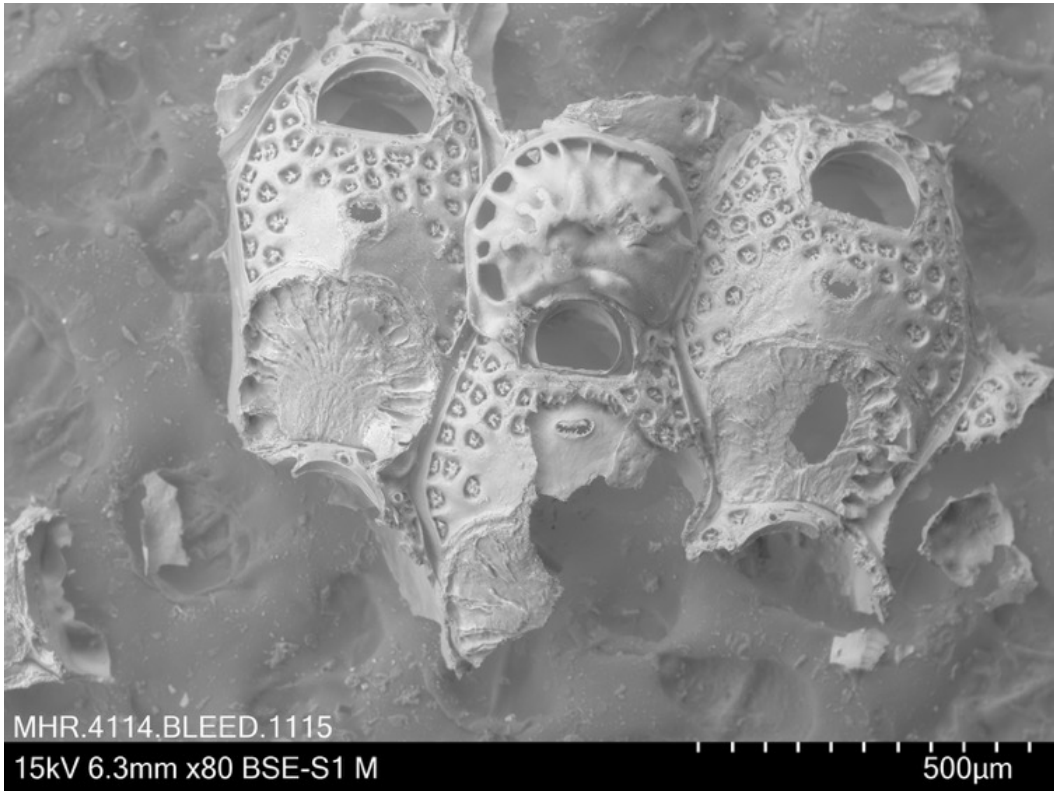


***Fenestrulina* n. sp. 2 BLEED 1115A**
Coll: 10.11.2018 (A. M. Smith)
Lat: -45.835
Long: 170.95
Depth: 111 m

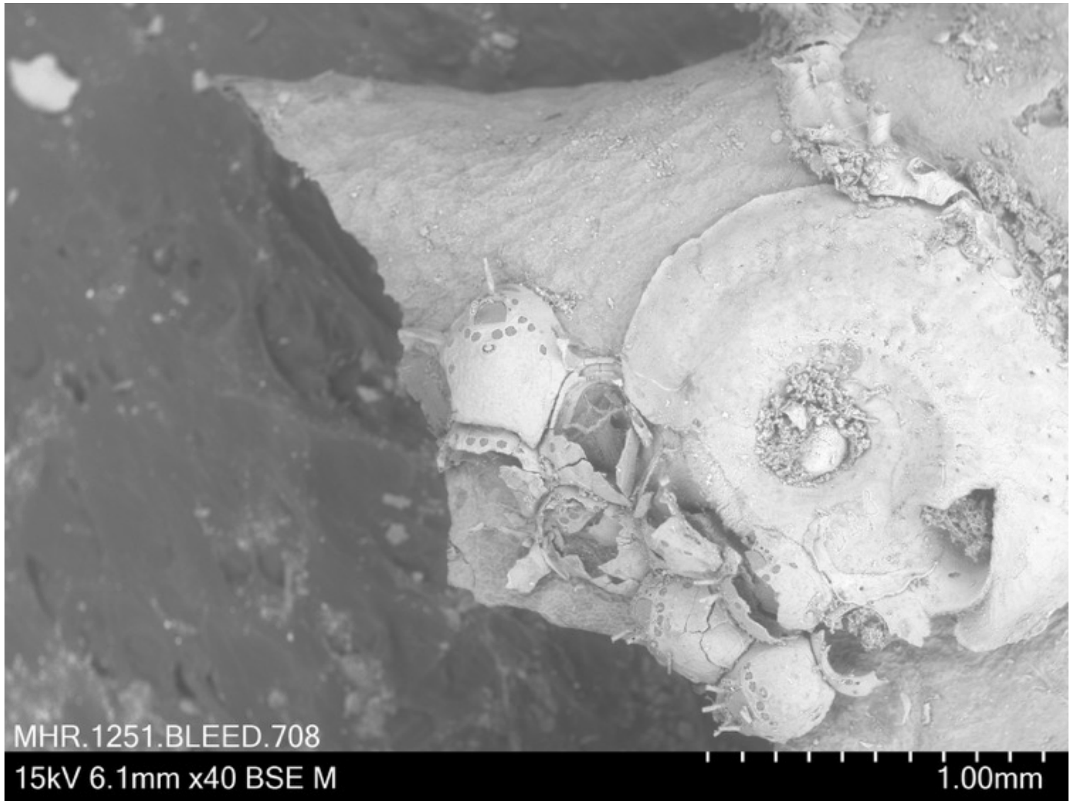


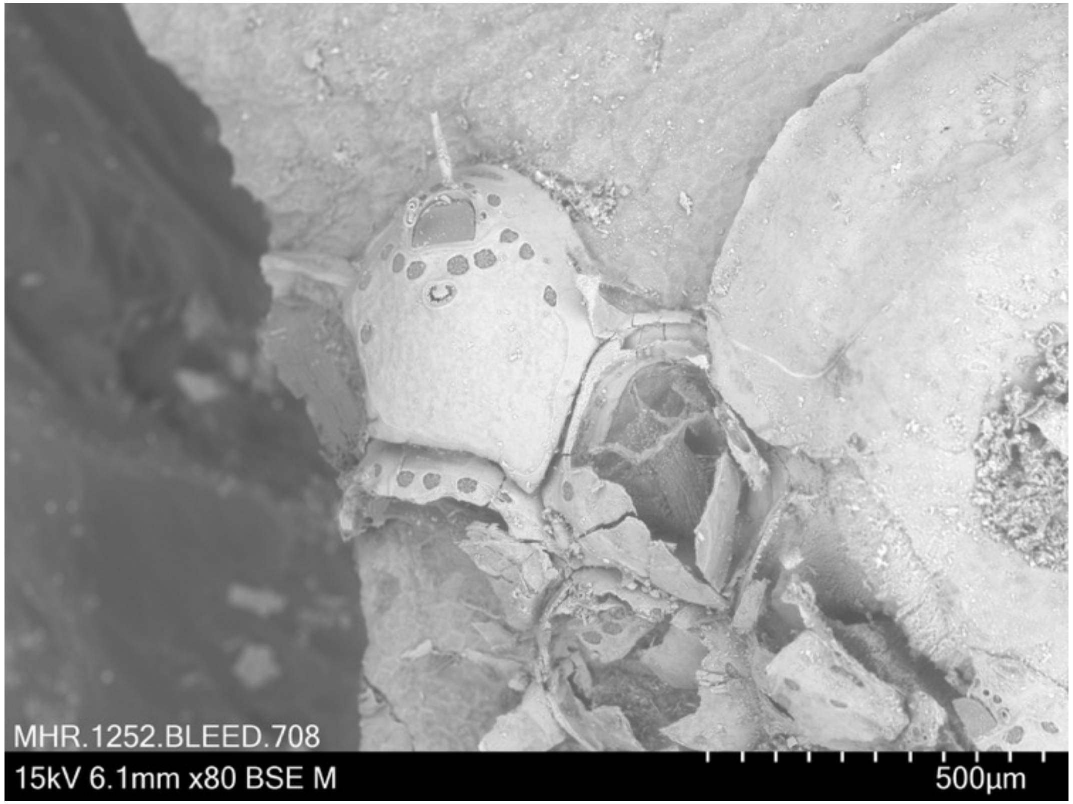


***Fenestrulina* cf. *disjuncta* BLEED 708**
Coll: 10.4.2018 (A. M. Smith, D. P. Gordon, H. Mello)
Lat: -46.97417
Long: 168.16694
Depth: 13 m

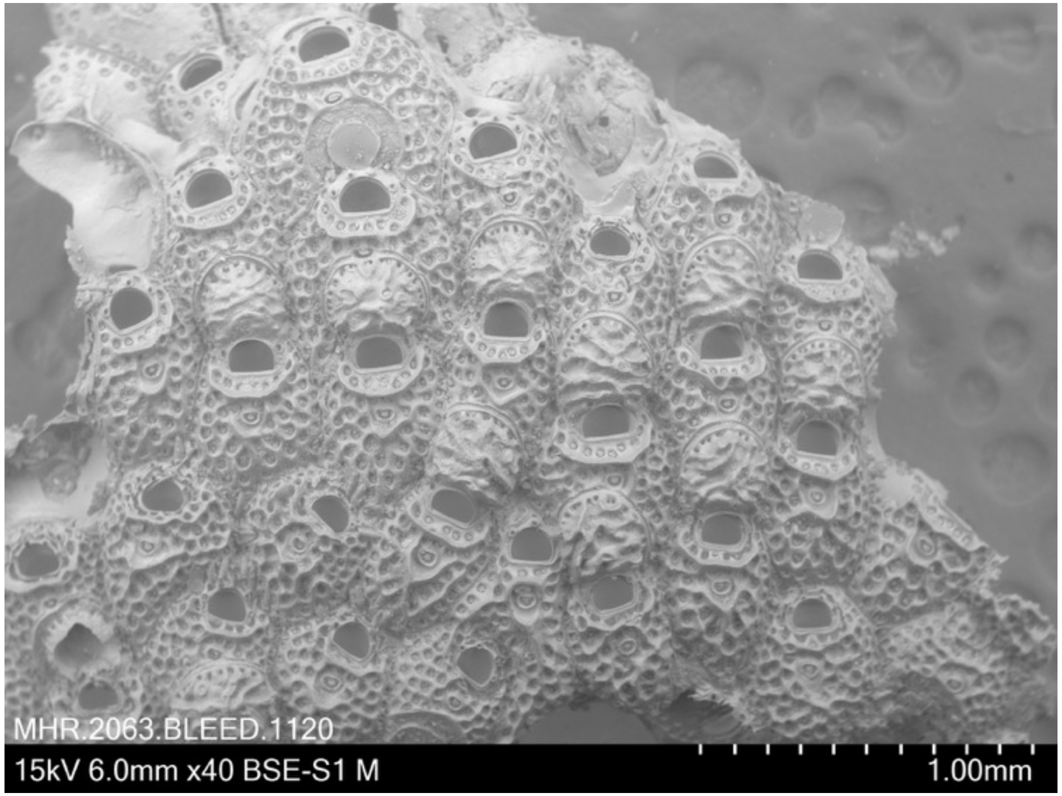


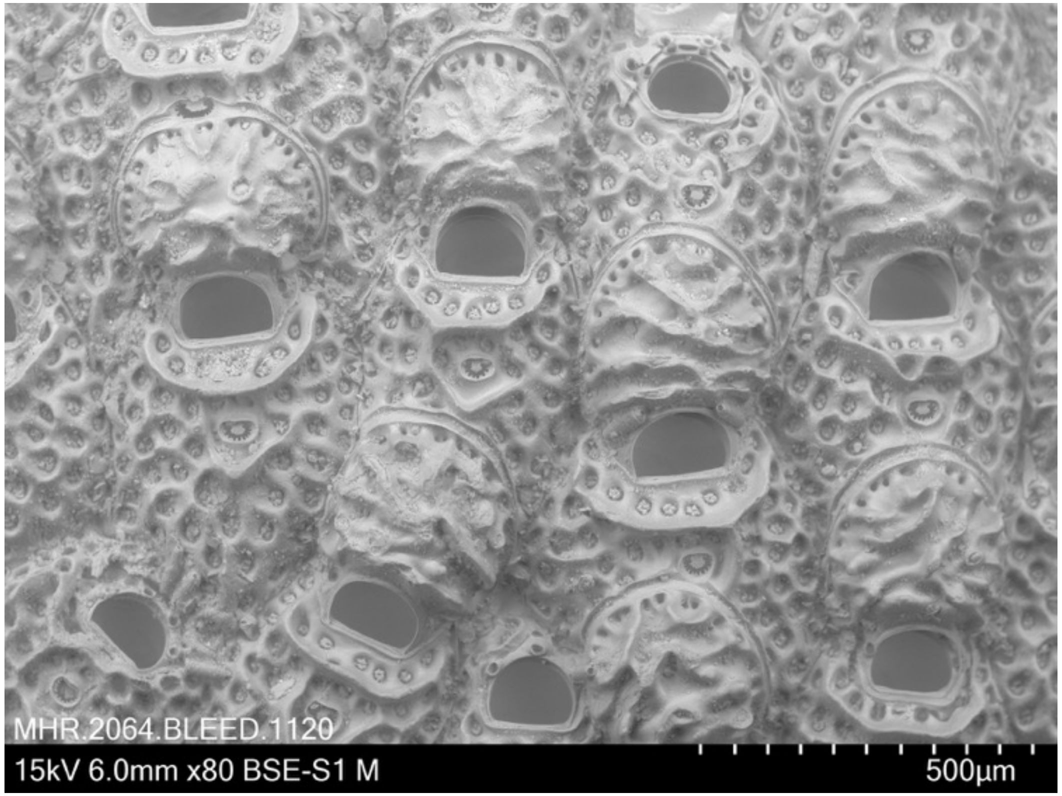


***Fenestrulina reticulata* BLEED 1120**
Coll: 10.11.2018 (A. M. Smith)
Lat: -45.8406
Long: 170.7961
Depth: 56 m

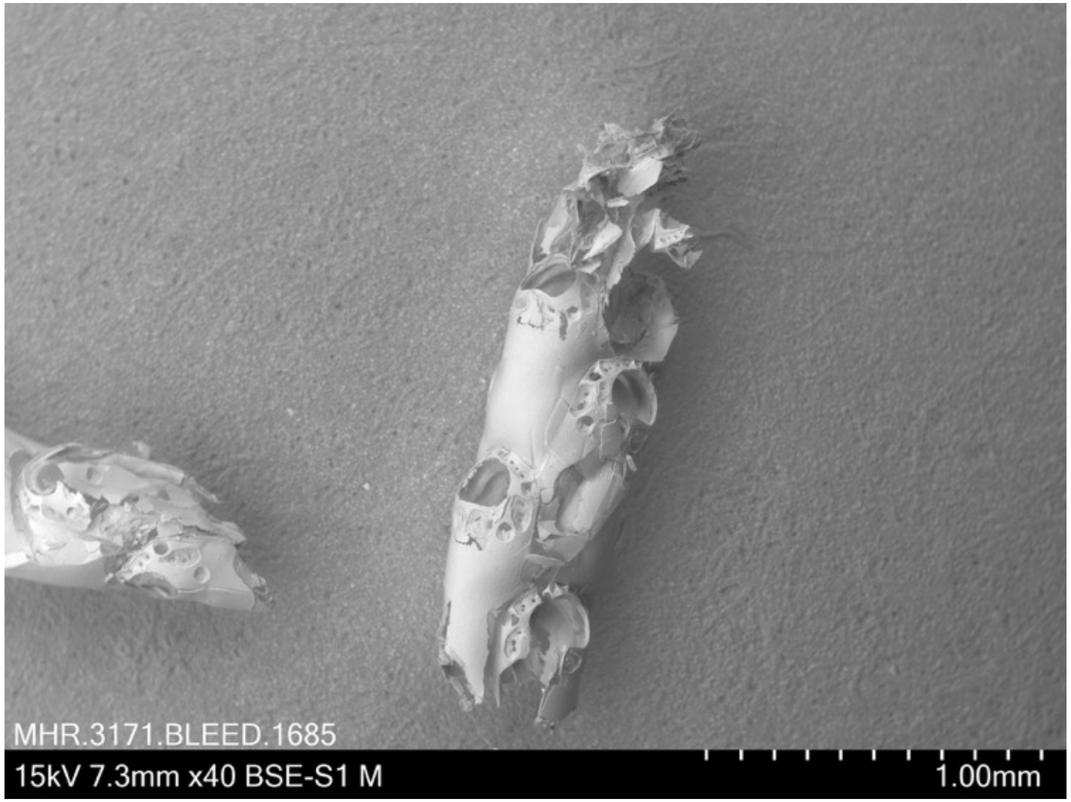


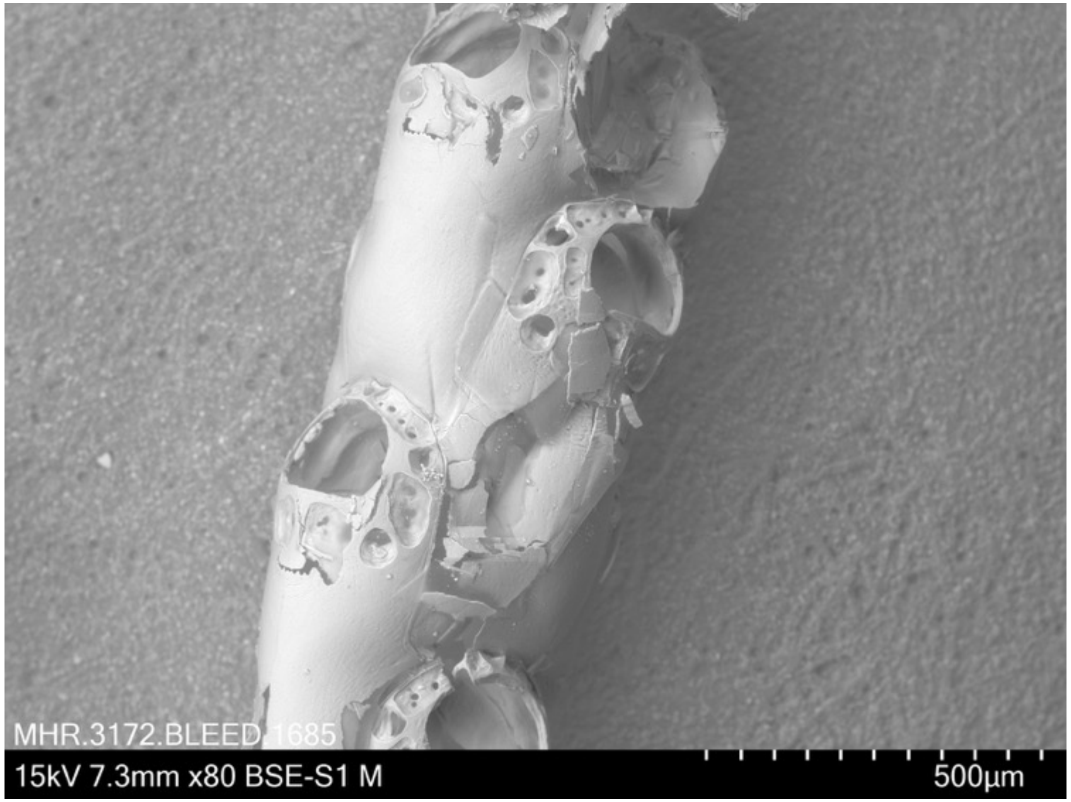


***Malakosaria sinclairii* BLEED 1685**
Coll: 18.3.2019 (A. M. Smith, D. P. Gordon)
Lat: -34.380
Long: 172.69
Depth: 52.5 m

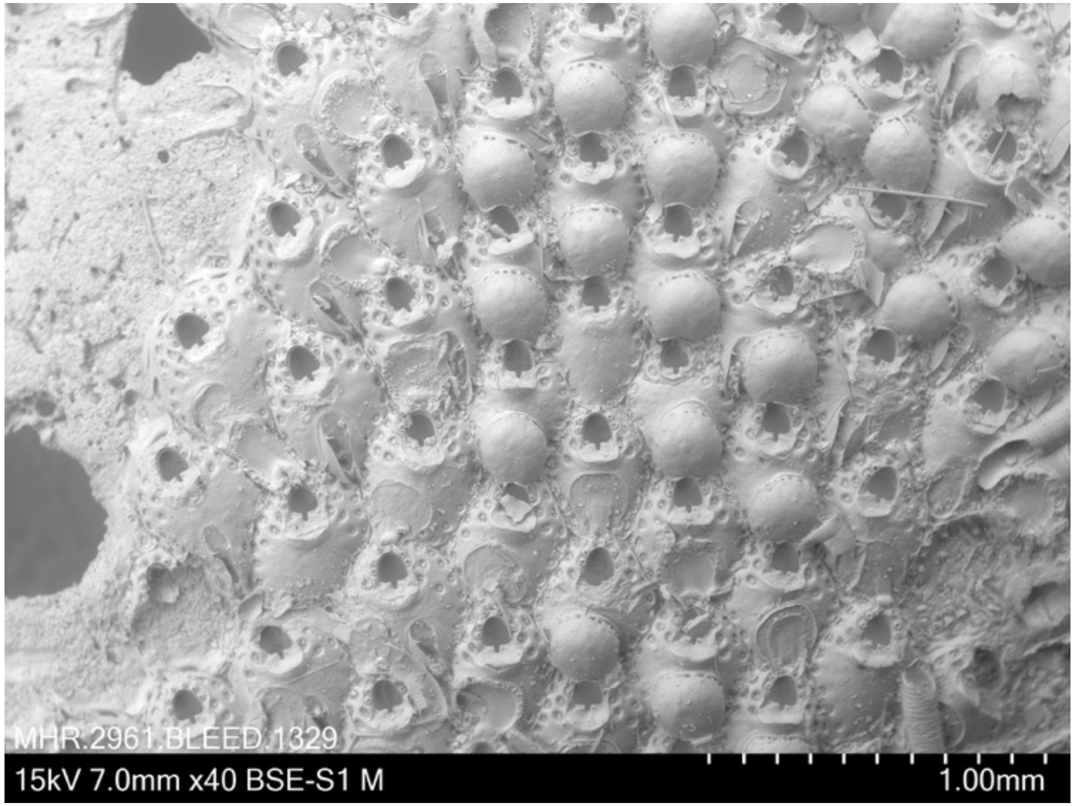


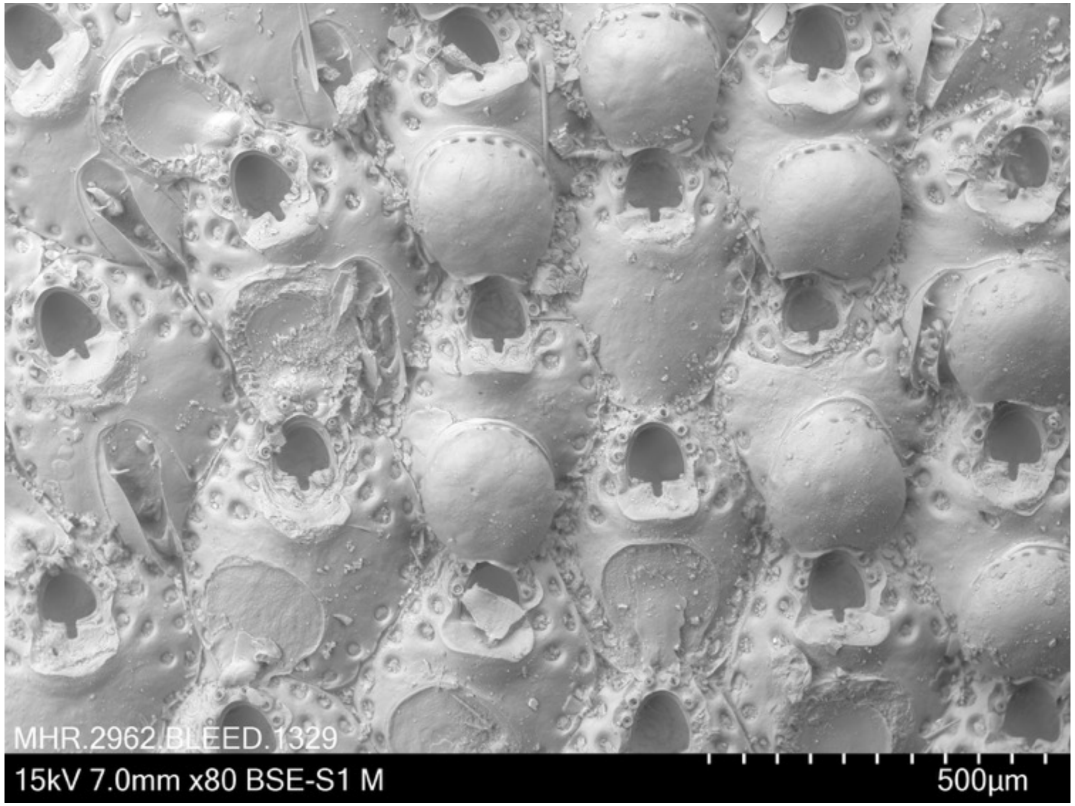


***Phonicosia* n. sp. 1 BLEED 1329**
Coll: 19.3.2019 (A. M. Smith, D. P. Gordon)
Lat: -34.99
Long: 173.97
Depth: 61.6 m

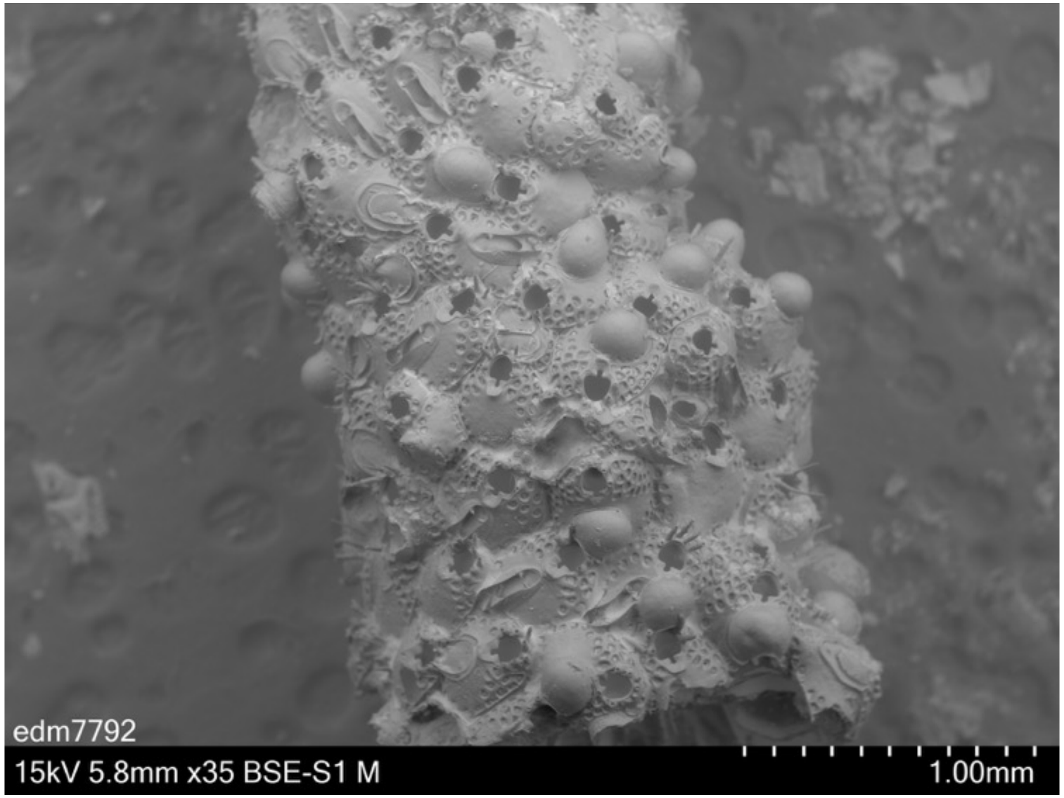


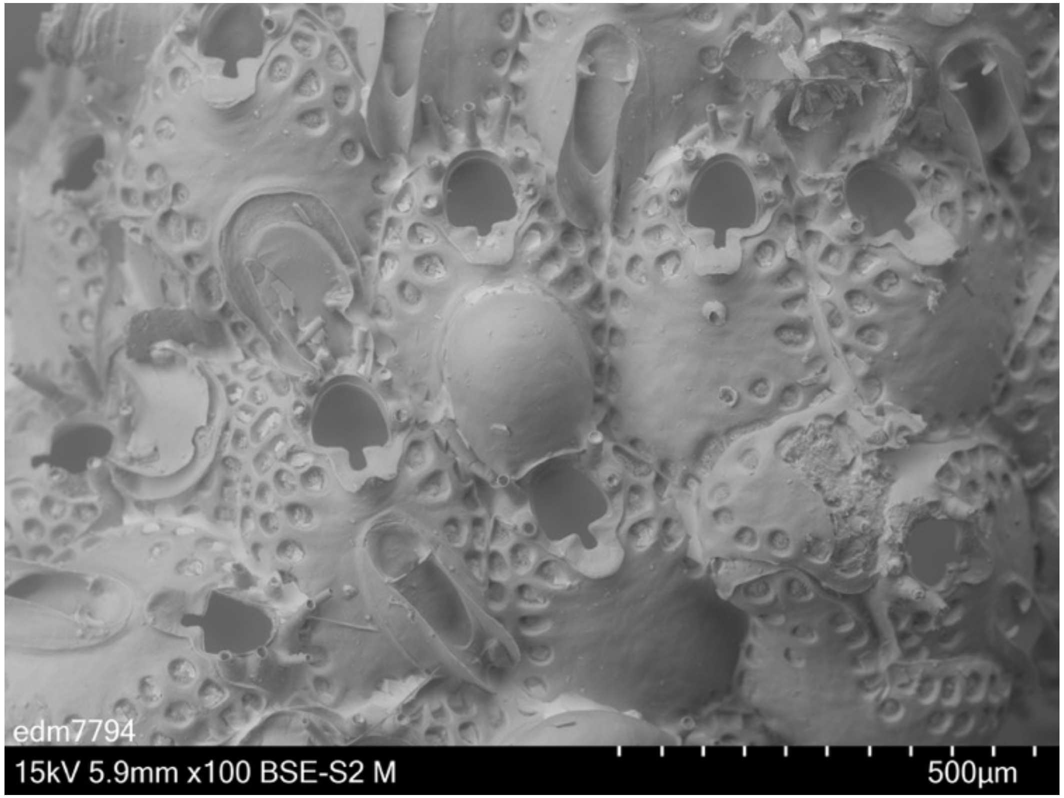


***Phonicosia* n. sp. 2 BLEED 594**
Coll: 10.4.2018 (A. M. Smith, D. P. Gordon, H. Mello)
Lat: -46.924166666666665
Long: 168.21833333333333
Depth: 58 m

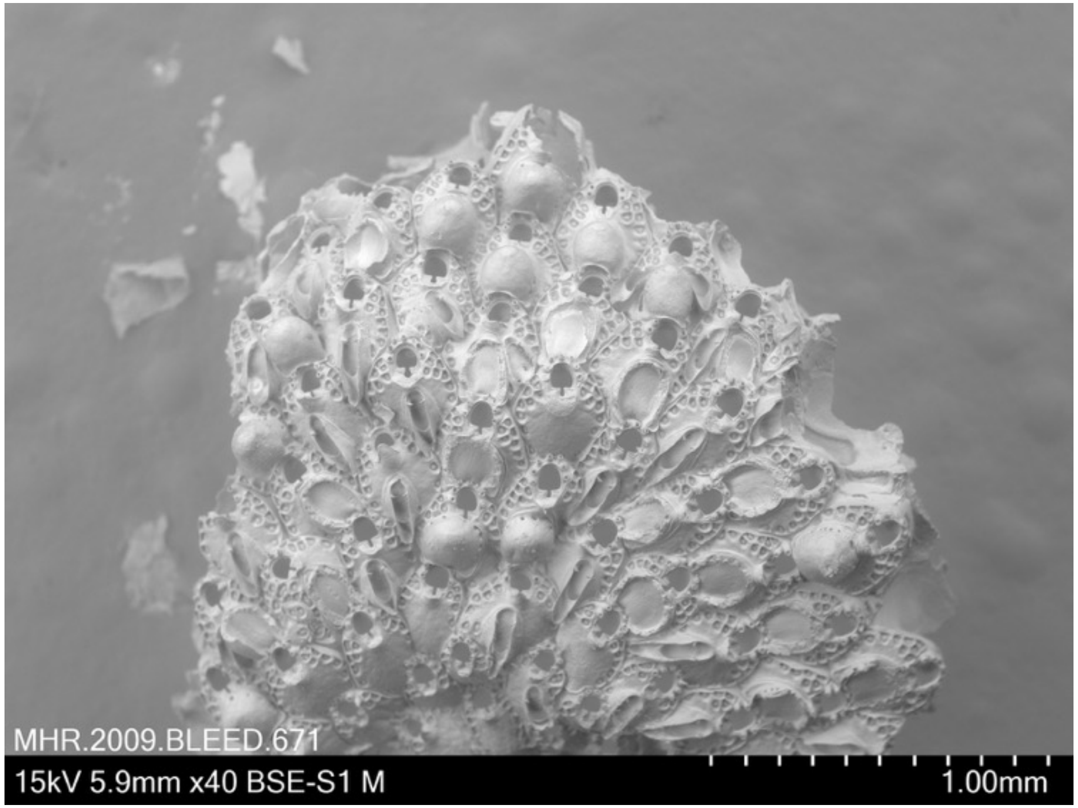


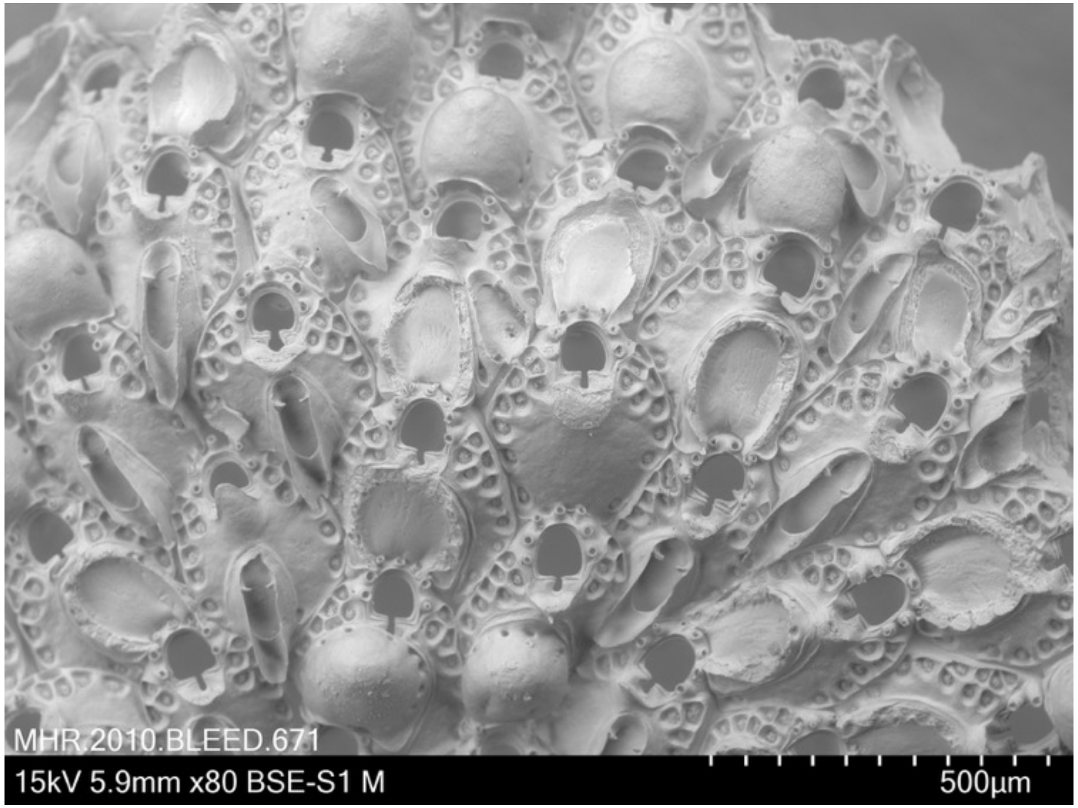


***Phonicosia*** **n. sp. 2 BLEED 671**
Coll: 8.4.2018 (A. M. Smith, D. P. Gordon, H. Mello)
Lat: -47.1372
Long: 168.12583
Depth: 80-85 m

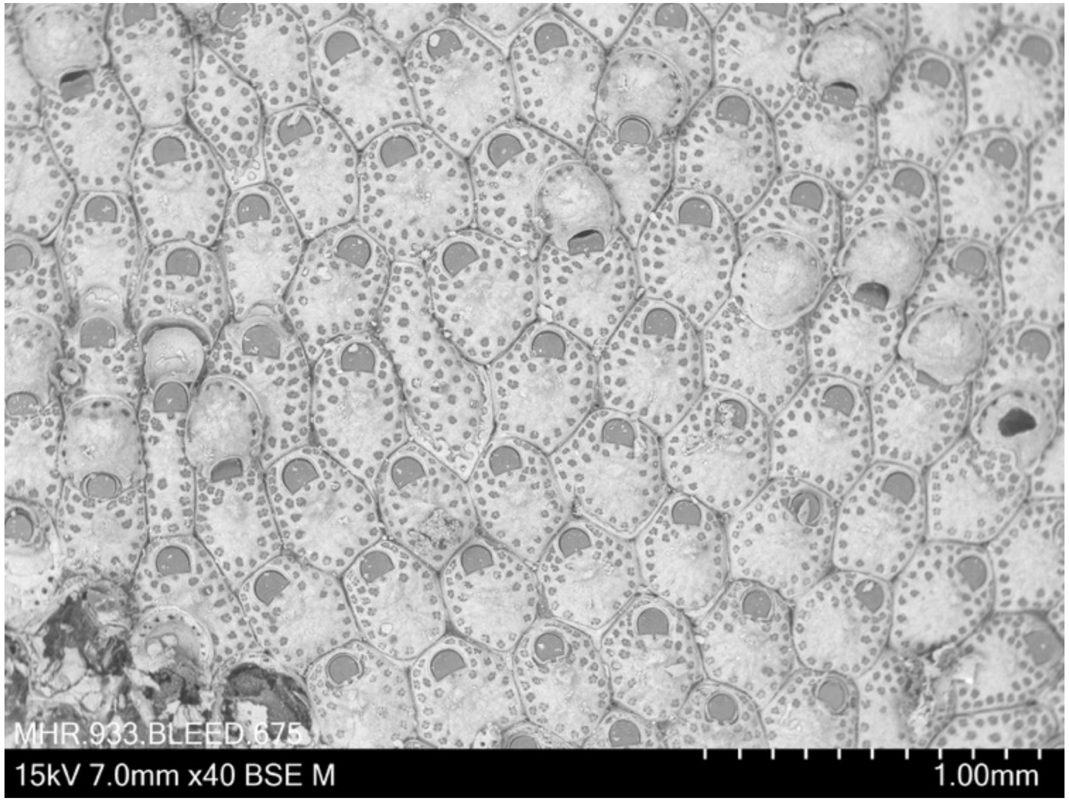


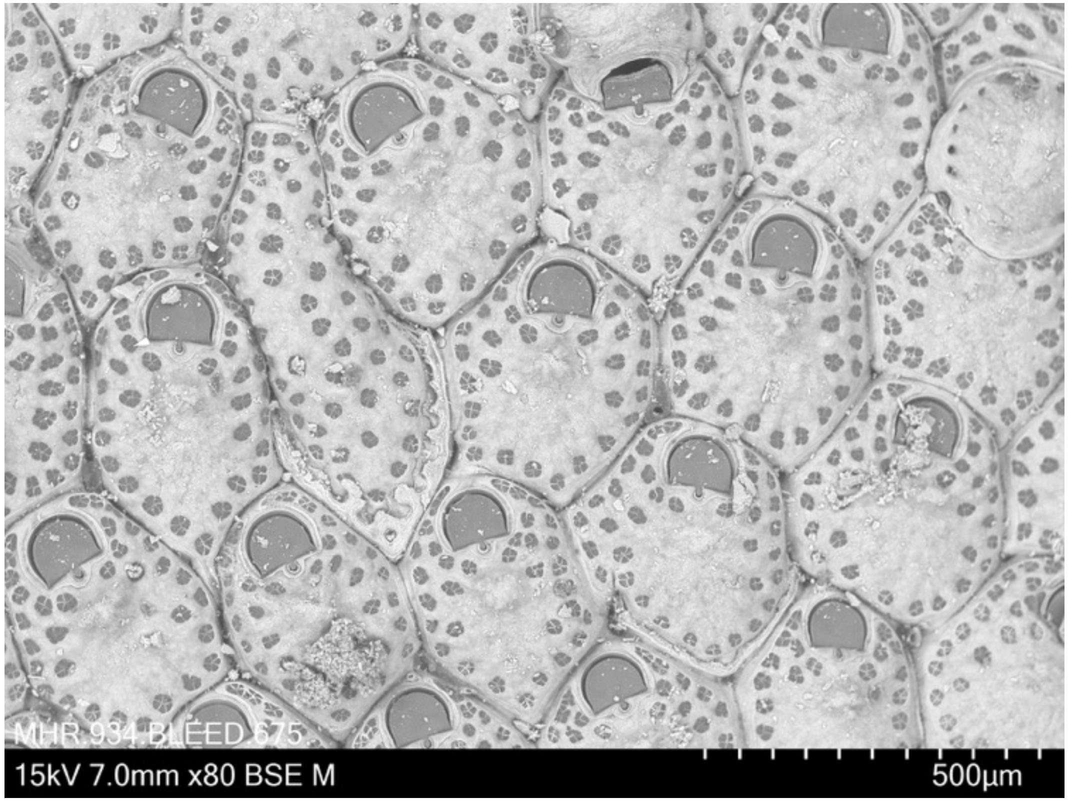


***Cribellopora napi*** **BLEED 675**
Coll: 8.4.2018 (A. M. Smith, D. P. Gordon, H. Mello)
Lat: -47.13722
Long: 168.12583333333333
Depth: 80-85 m

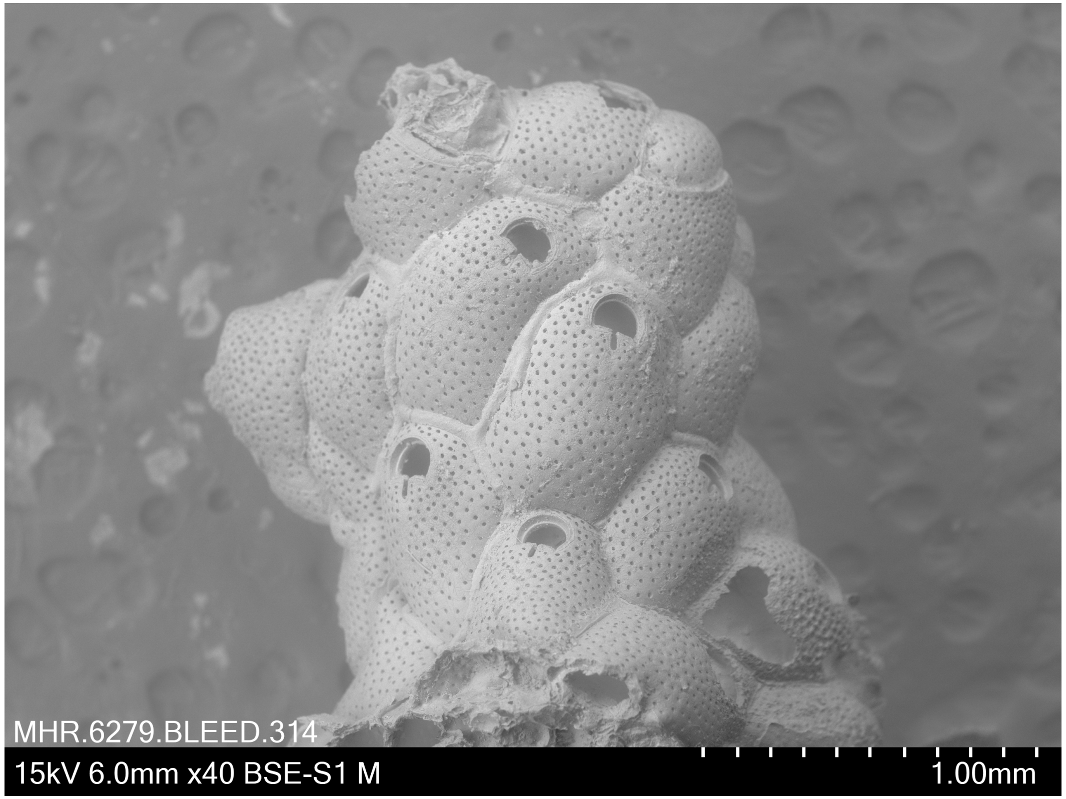


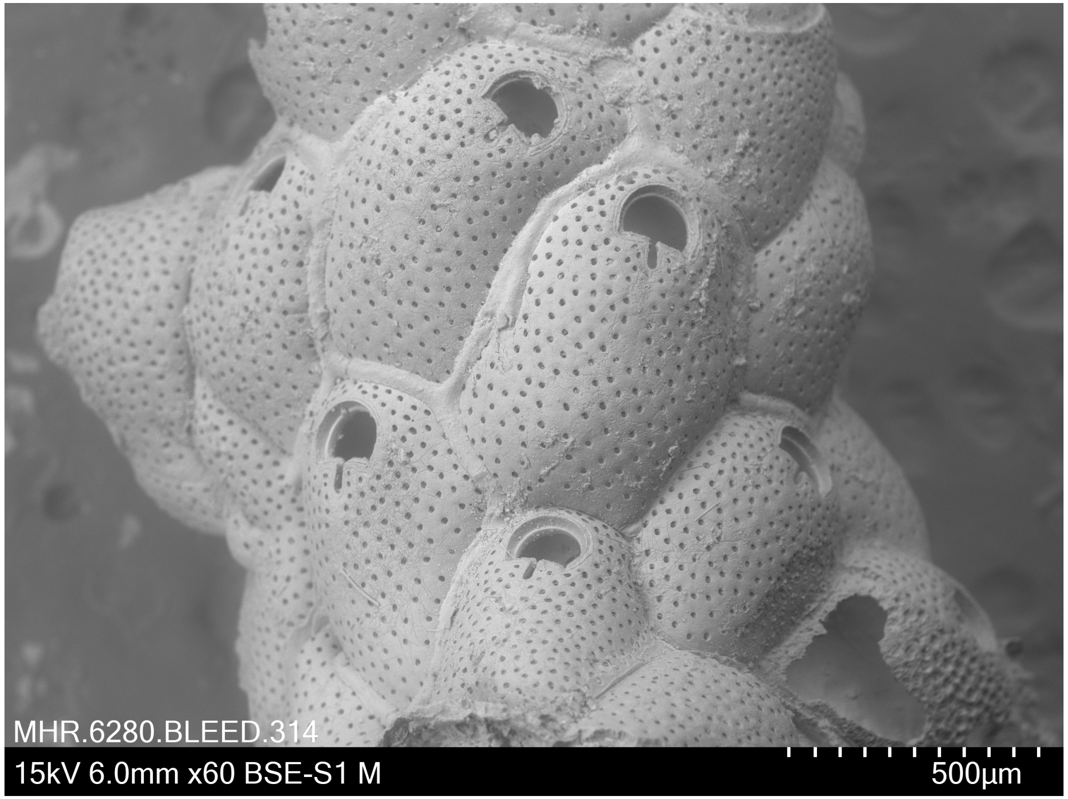


***Arthropoma cecilii* BLEED 177**
NIWA 55263 TAN0906/65
Coll: 8.7.2009
Lat: -35.00383
Long: 174.05567
Depth: 105 m
Notes: BLEED 314 NIWA 55264 is used as a voucher as it is from exactly the same sample as the one sequenced (BLEED 177)

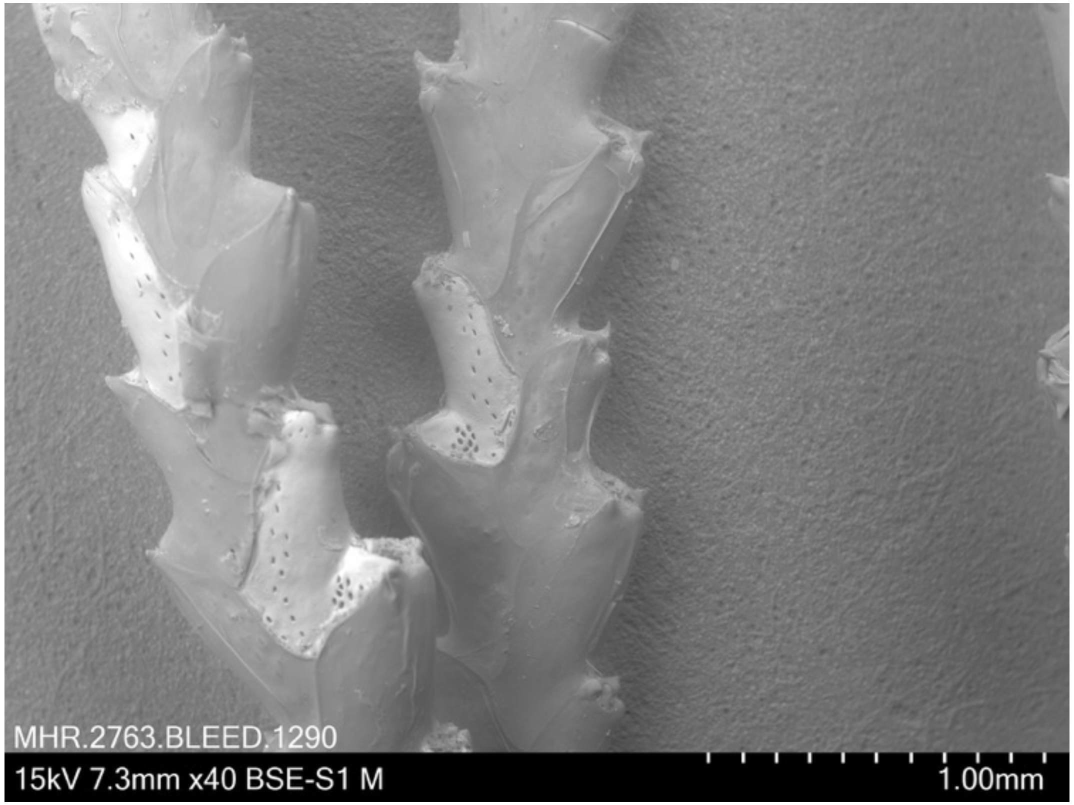

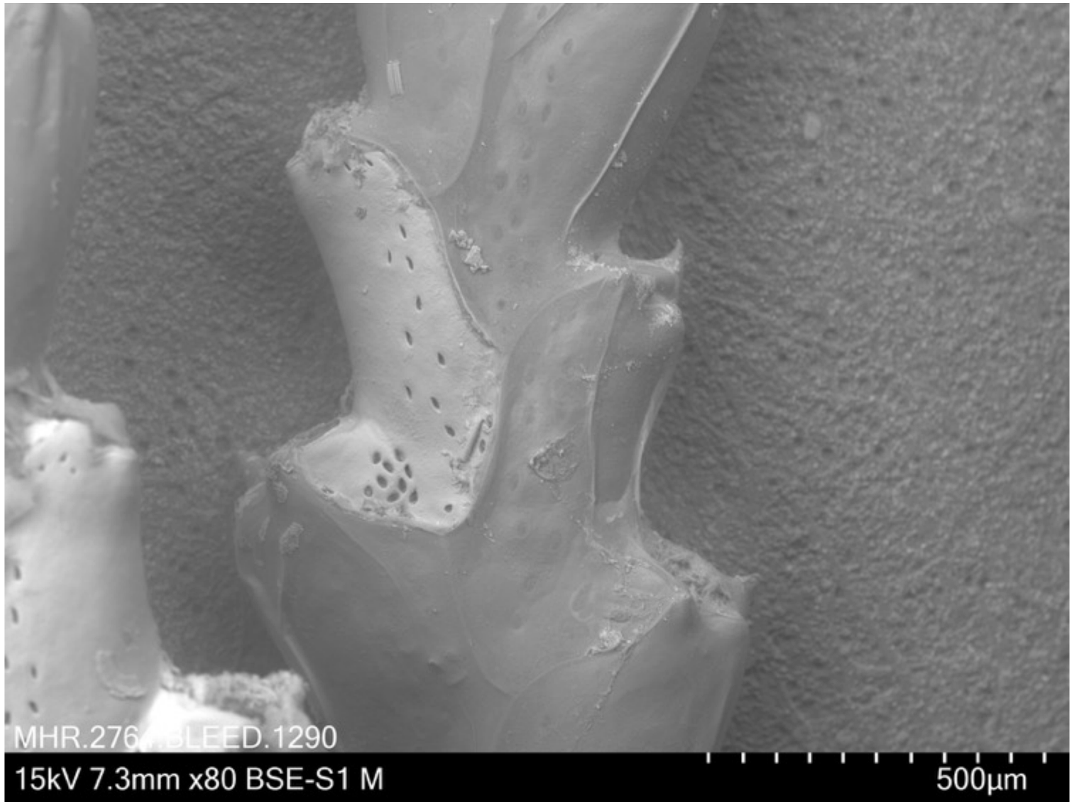


***Reciprocus regalis* BLEED 1290**
Coll: 17.3.2019 (A. M. Smith, D. P. Gordon)
Lat: -34.17
Long: 172.04
Depth: 85.4 m

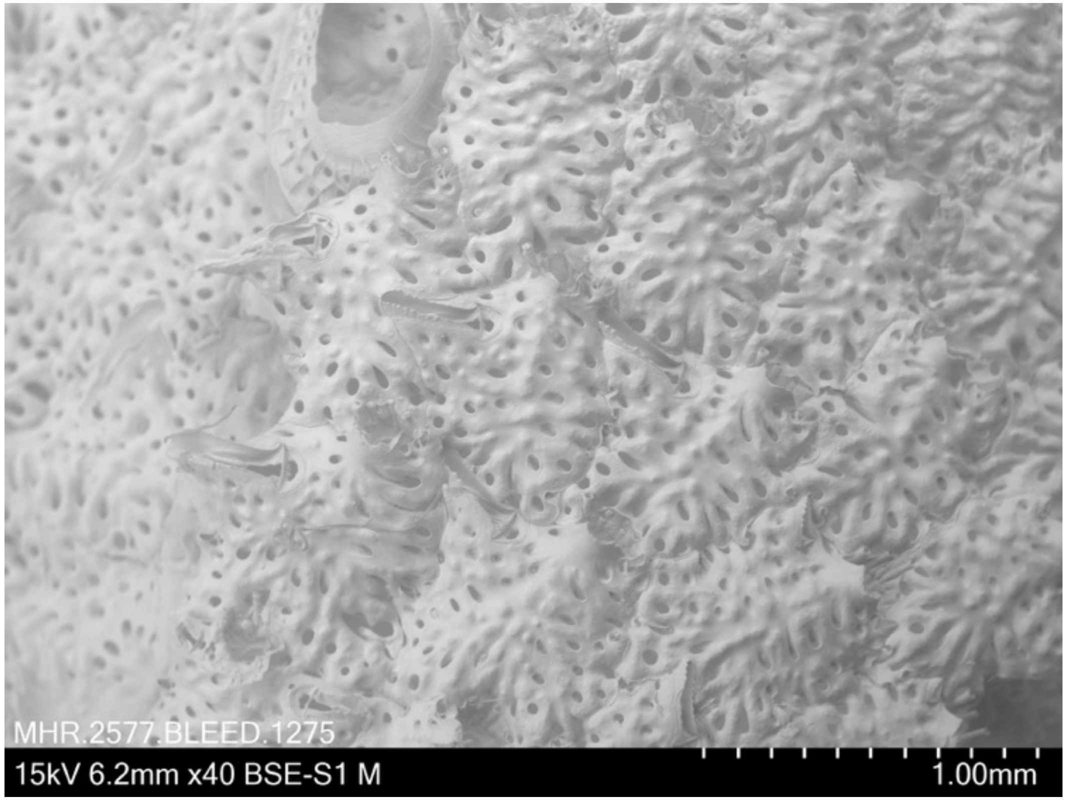


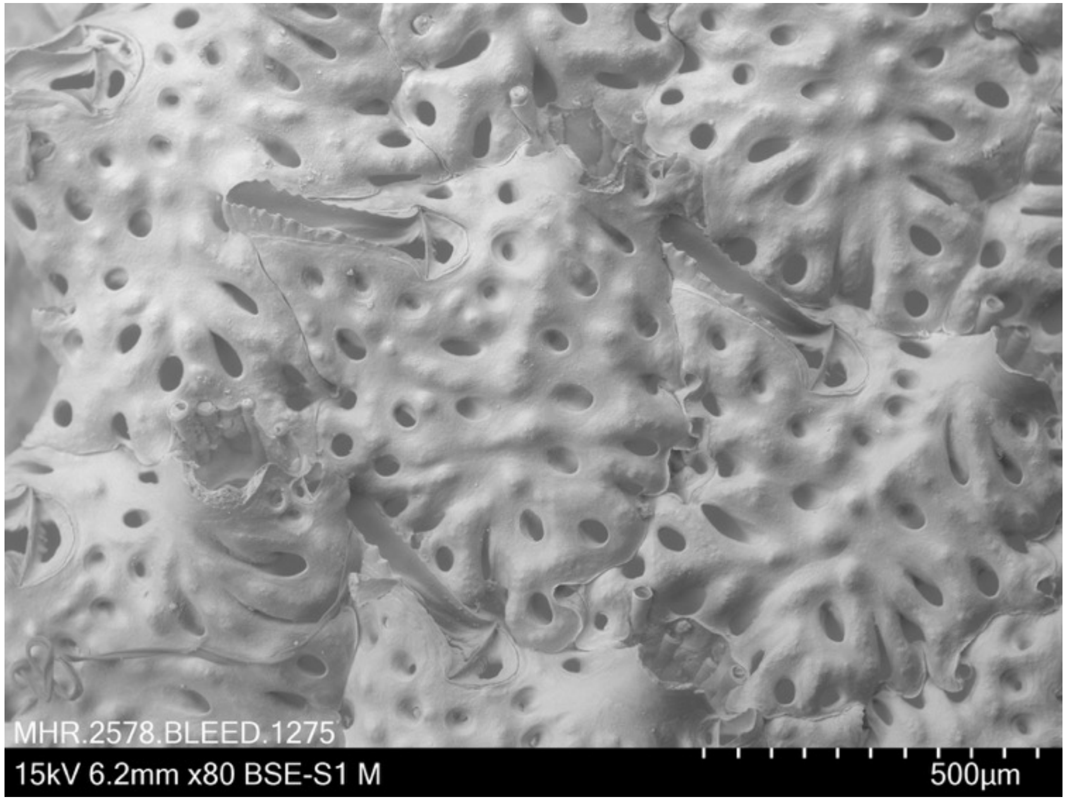


***Escharoides* cf. *excavata* BLEED 1275**
Coll: 16.3.2019 (A. M. Smith, D. P. Gordon)
Lat: -34.99
Long: 173.99
Depth: 81 m

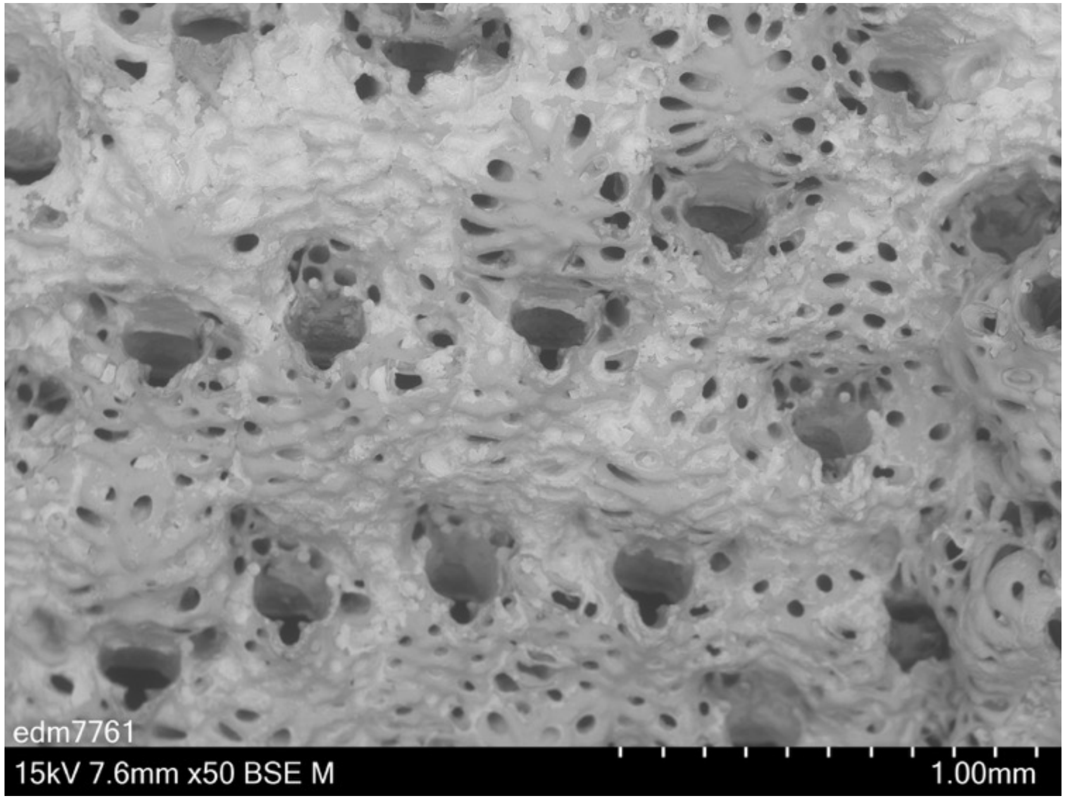


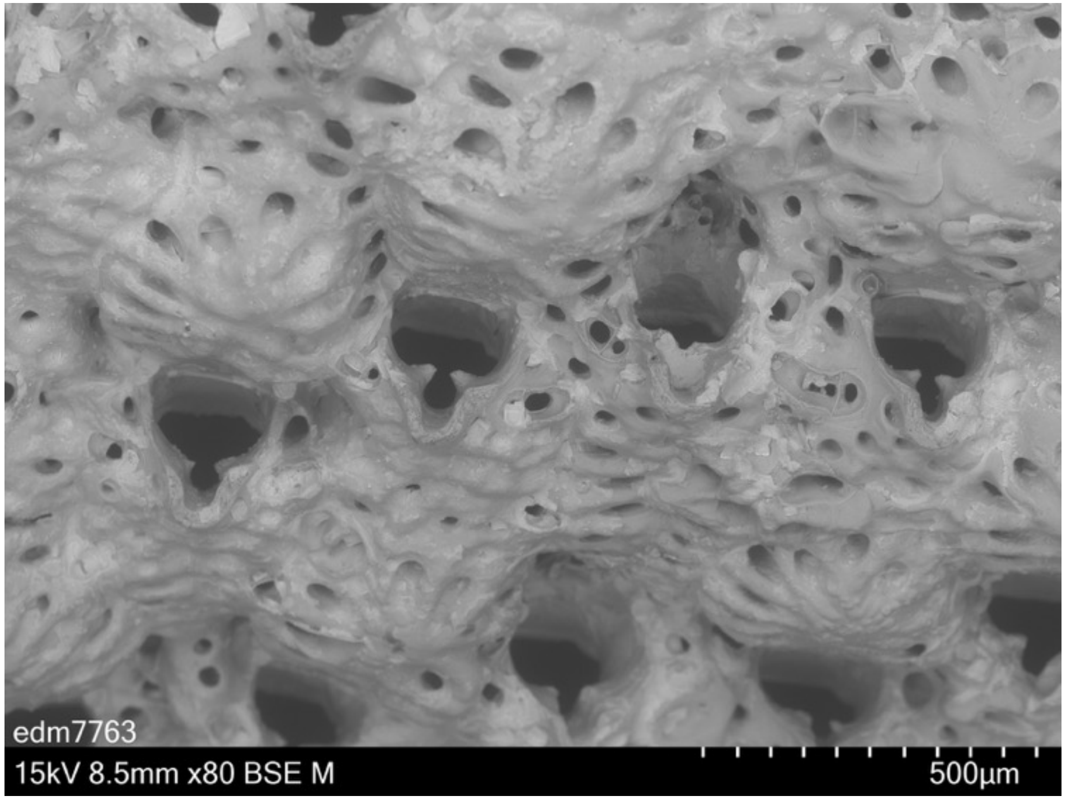


***Escharoides angela* BLEED 636**
Coll: 10.4.2018 (A. M. Smith, D. P. Gordon, H. Mello)
Lat: -46.945
Long: 168.1583
Depth: 42 m

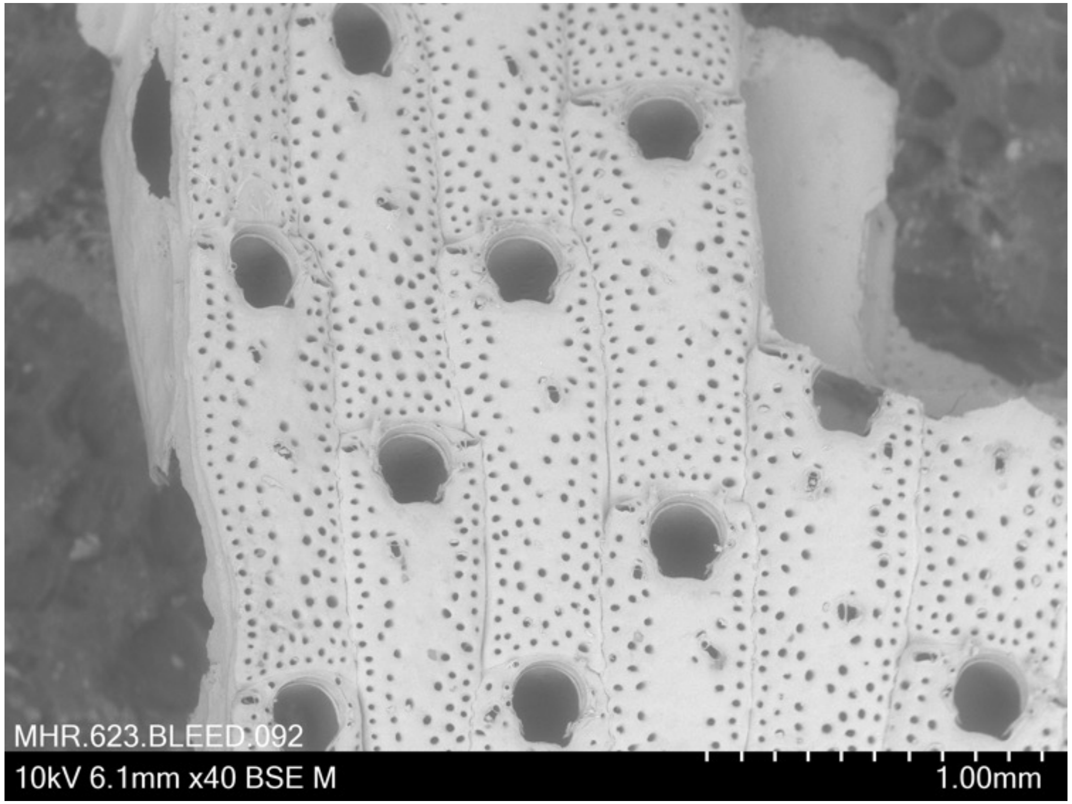


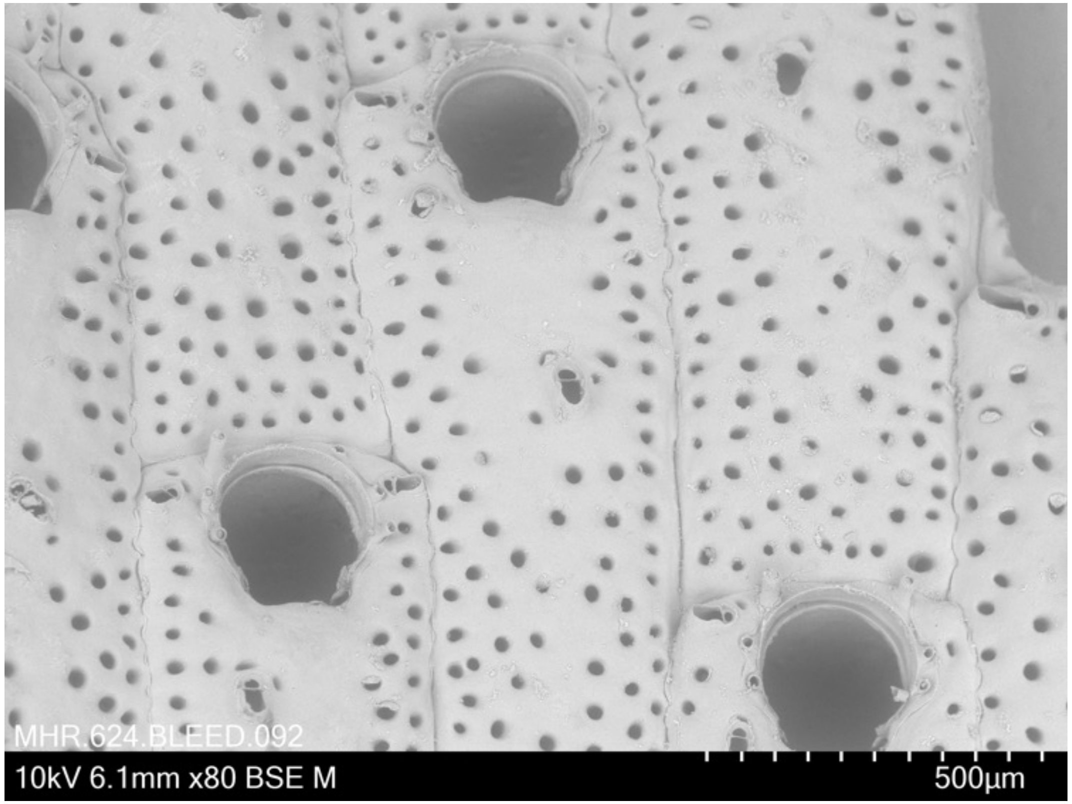


***Hippomenella* cf. *vellicata* BLEED 92**
NIWA 75874 (TRIP3465/17)
Coll: 24.3.2012
Lat: -46.365
Long: 170.433
Depth: 122-145 m

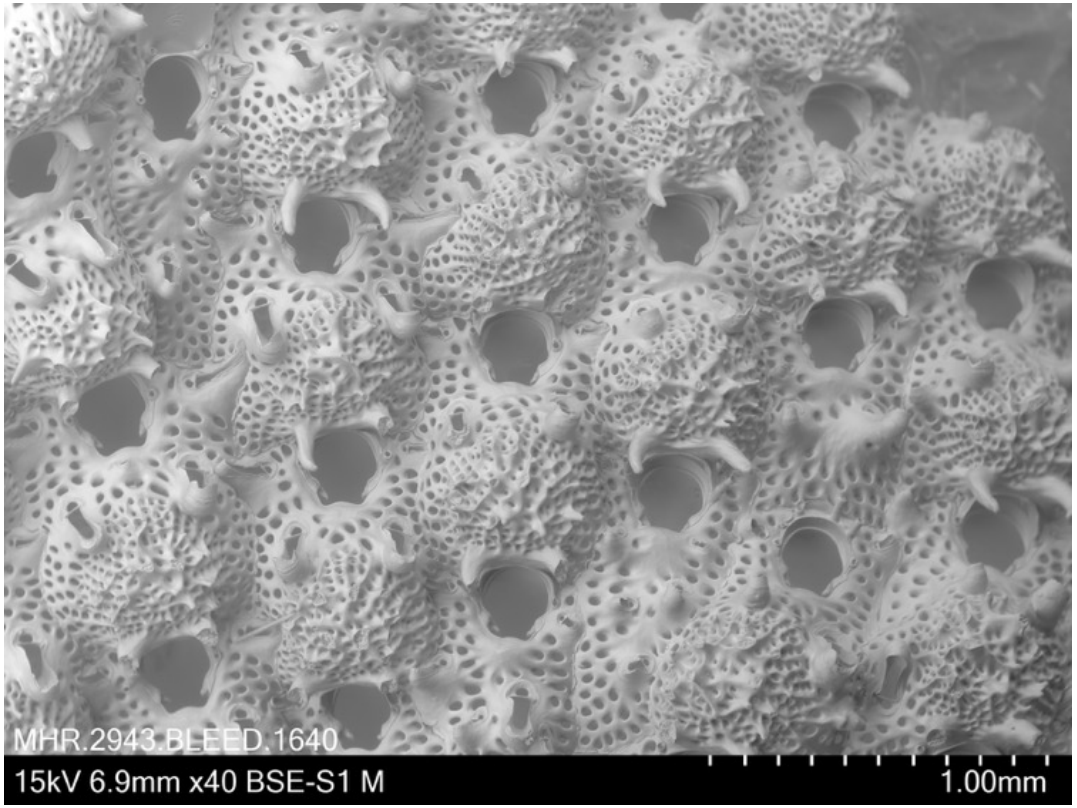


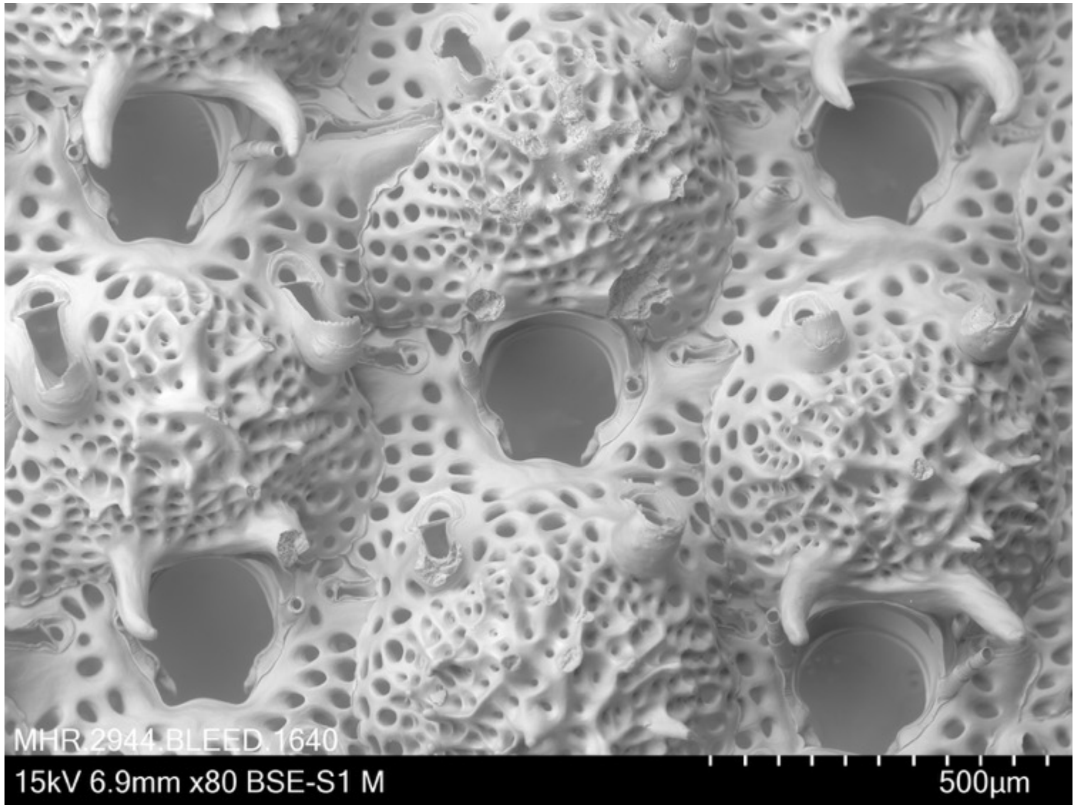


***Hippomenella* n. sp. BLEED 1640**
Coll: 17.3.2019 (A. M. Smith, D. P. Gordon)
Lat: -34.156
Long: 172.09
Depth: 82.9 m

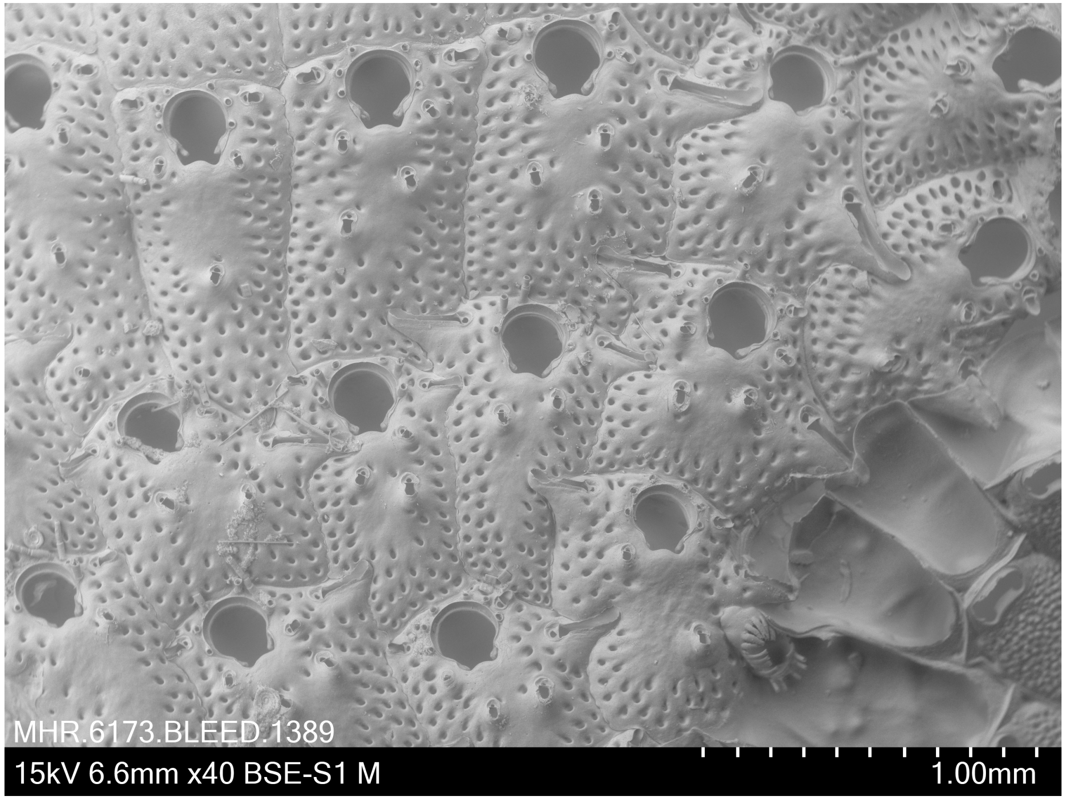


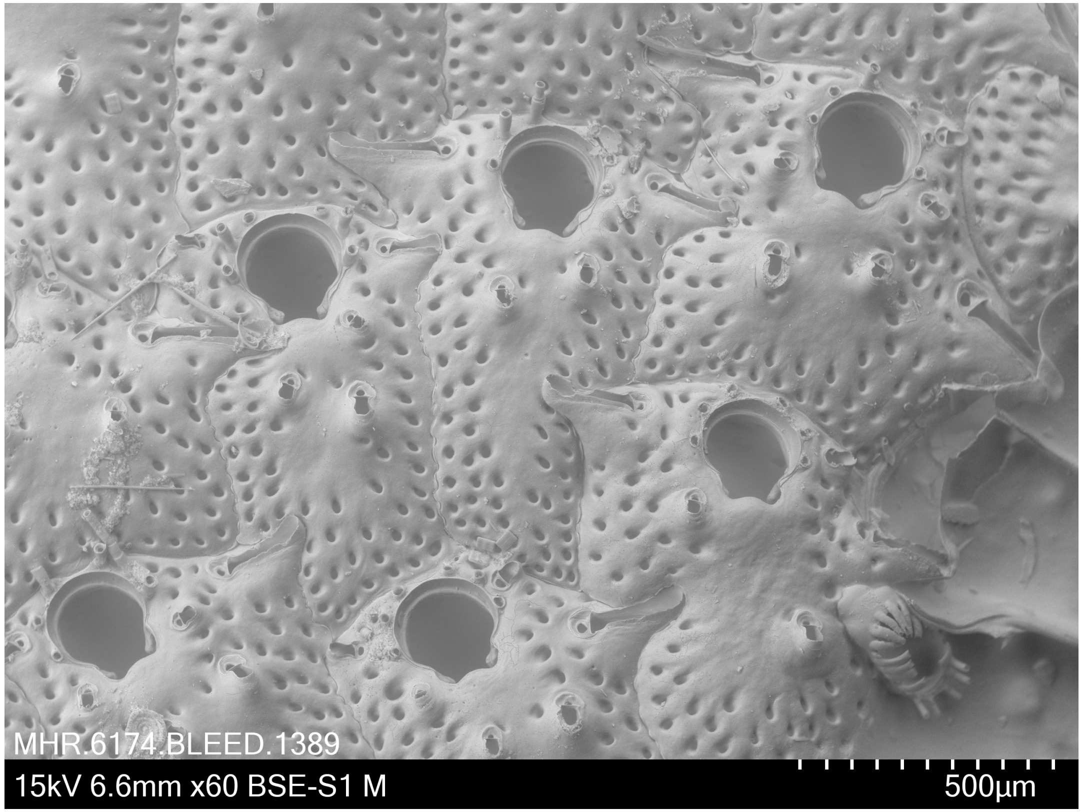


***Hippomenella* cf. *vellicata* BLEED 1389B**
Coll: 17.3.2019 (A. M. Smith; D. P. Gordon)
Lat: -34.159999999999997
Long: 172.09
Depth: 82.9 m

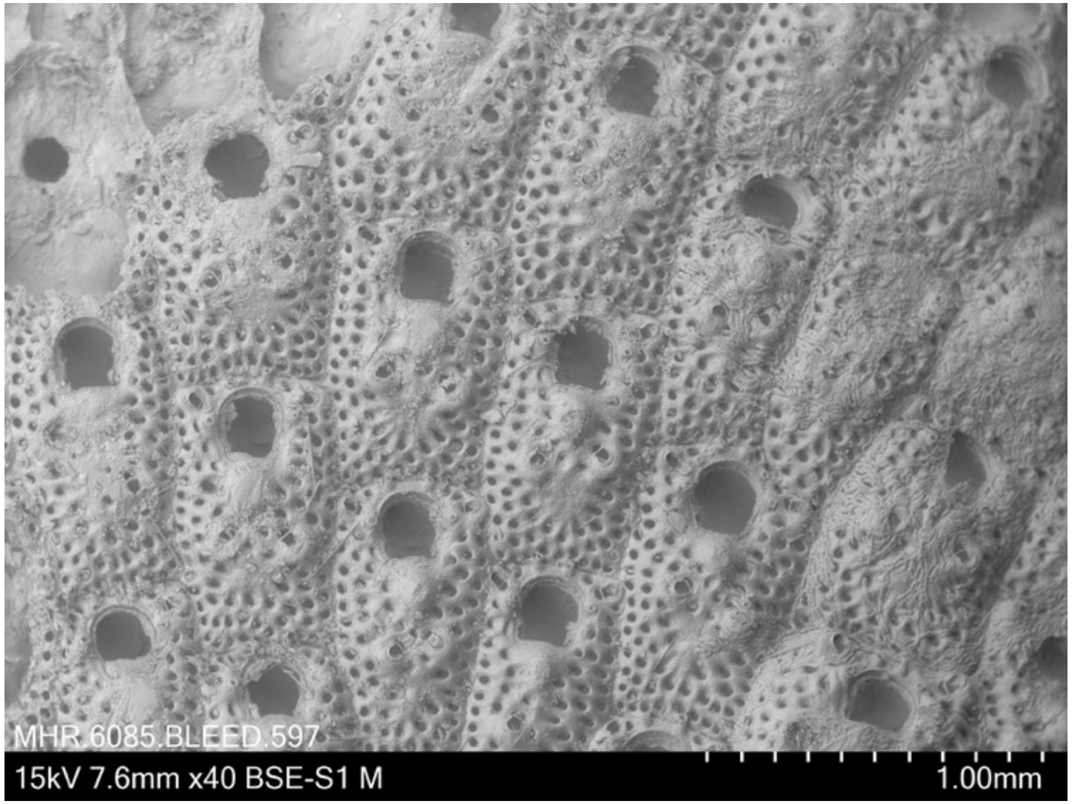


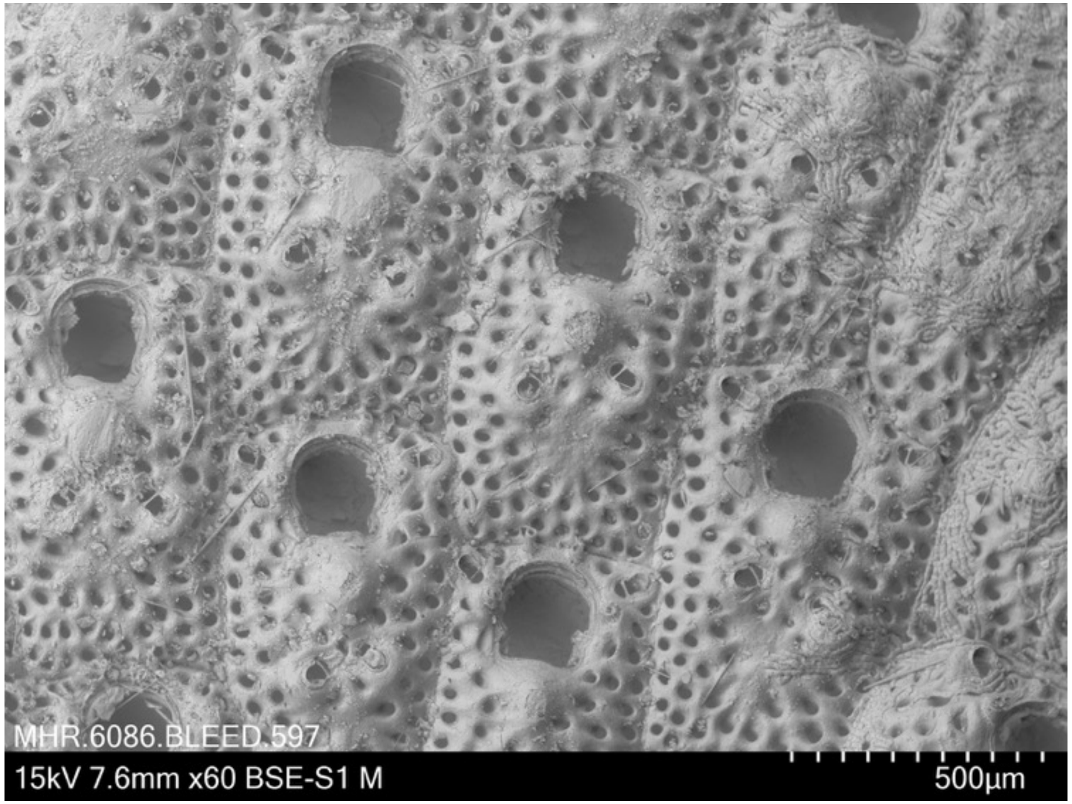


***Hippomenella vellicata*** **BLEED 597**
Coll. 8.4.2018 (A. M. Smith, D.P. Gordon, H. Mello)
Lat: - 46.662
Long: 168.386
Depth: 22 m

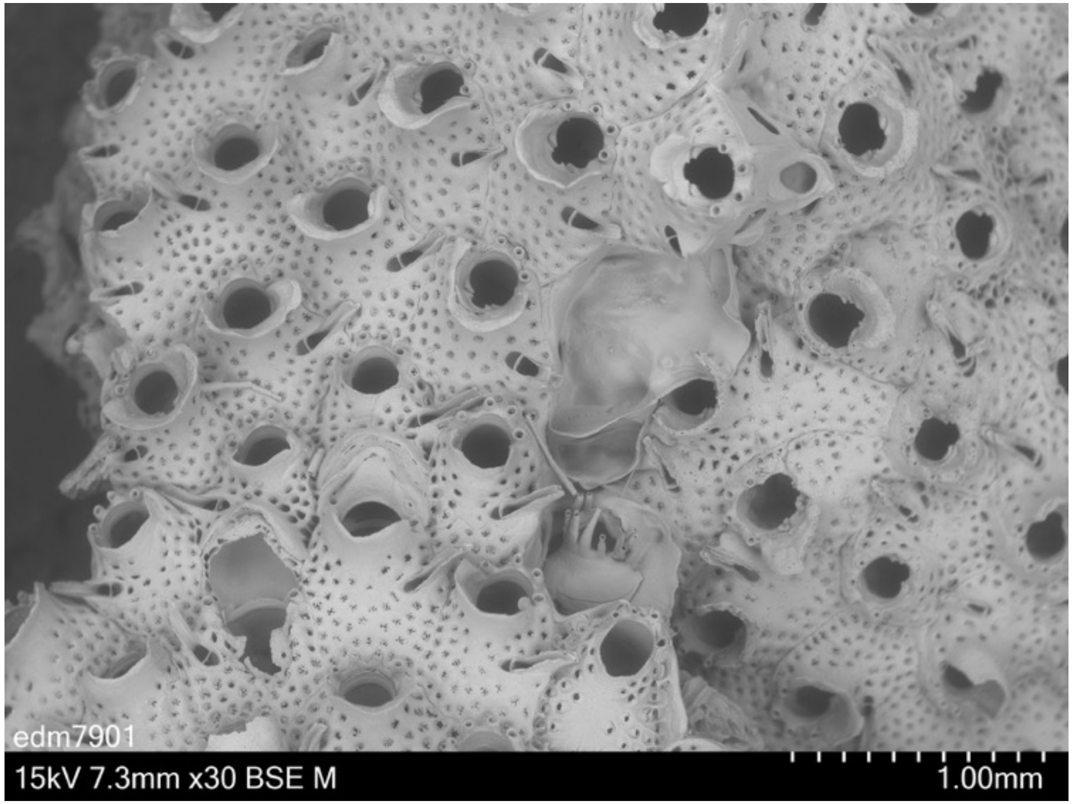


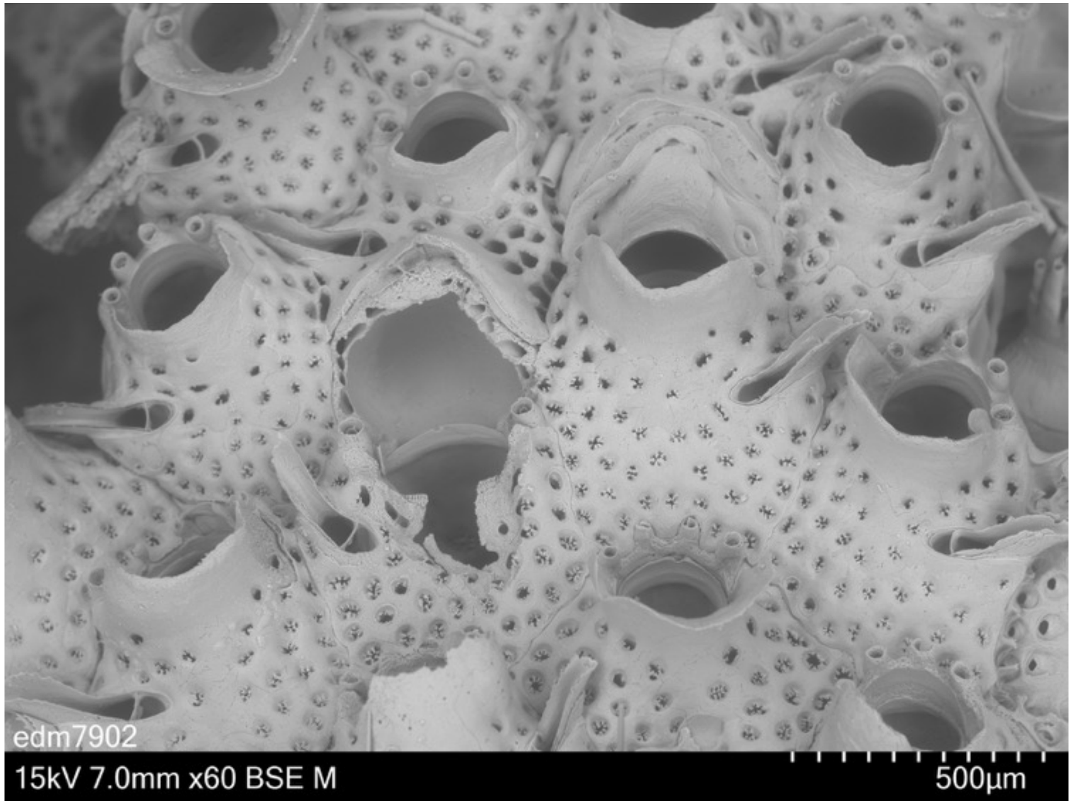


***Chiastosella longaevitas* BLEED 1367**
Coll: 17.3.2019 (A. M. Smith; D. P. Gordon)
Lat: -34.21
Long: 172.11
Depth: 104 m

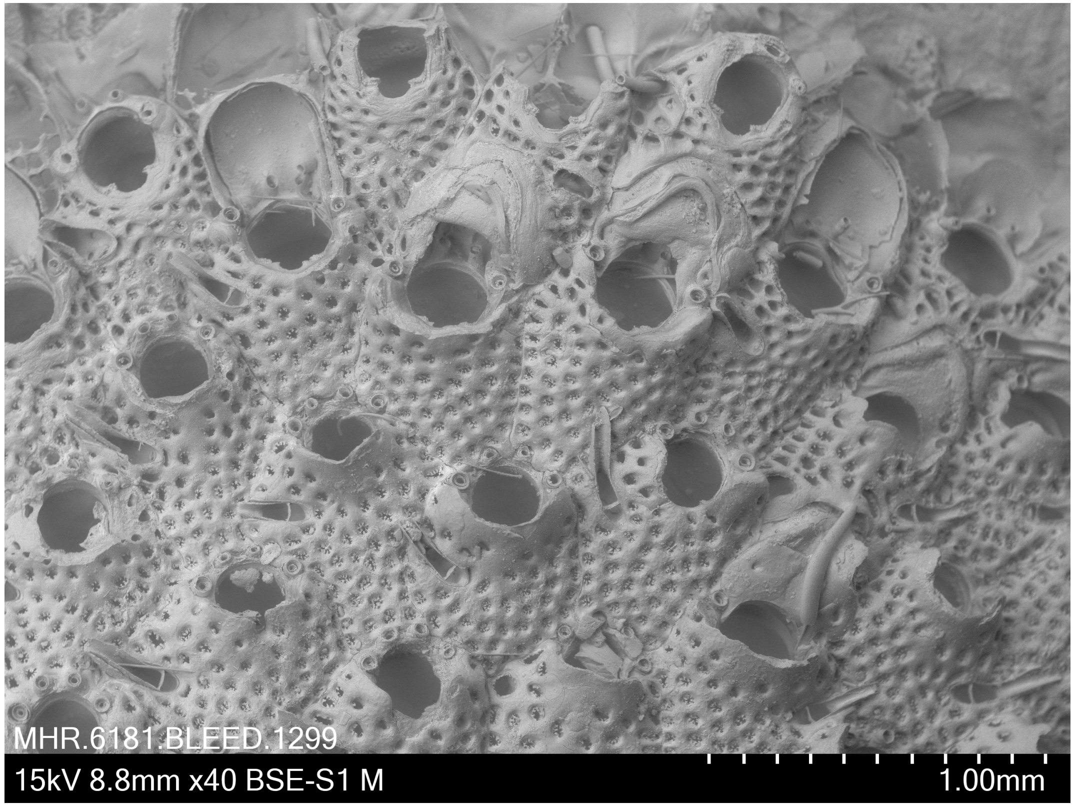


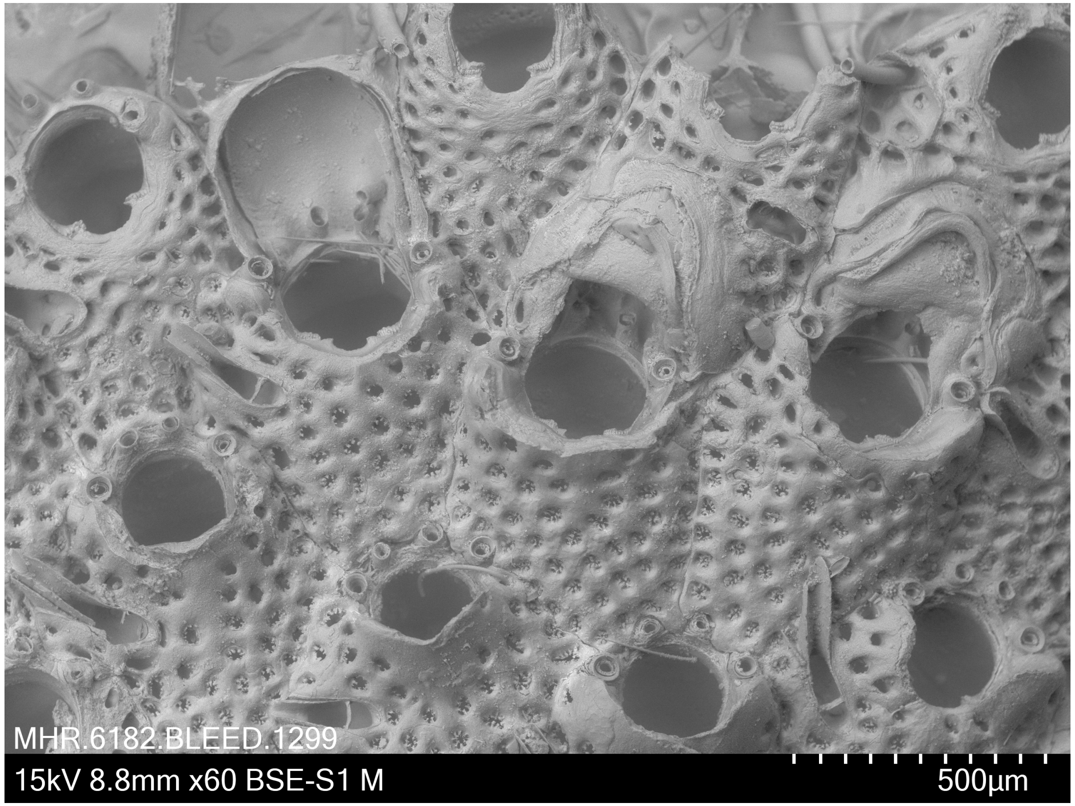


***Chiastosella longaevitas* BLEED 1299A**
Coll: 17.3.2019 (A. M. Smith, D. P. Gordon)
Lat: -34.21
Long: 172.11
Depth: 104 m

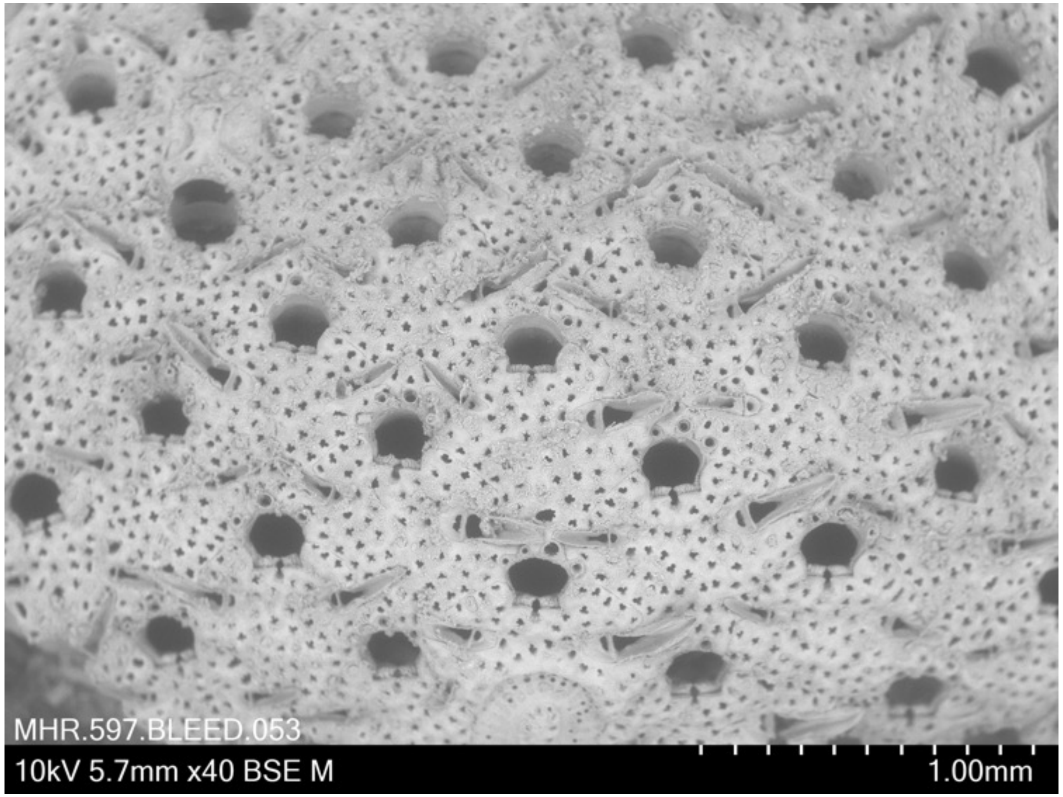


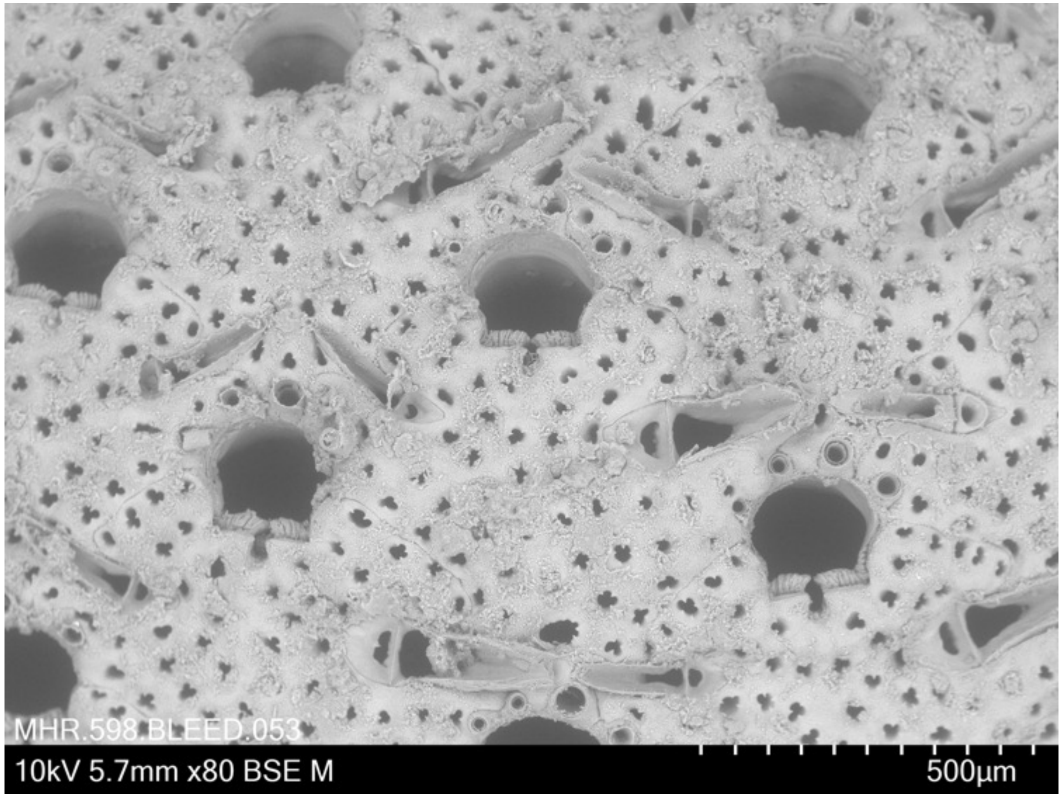


***Chiastosella splendida*** **BLEED 53**
NIWA 56945 (TAN0906/170)
Coll: 15.7.2009
Lat: -34.3781667
Long: 173.0265
Depth: 65 m

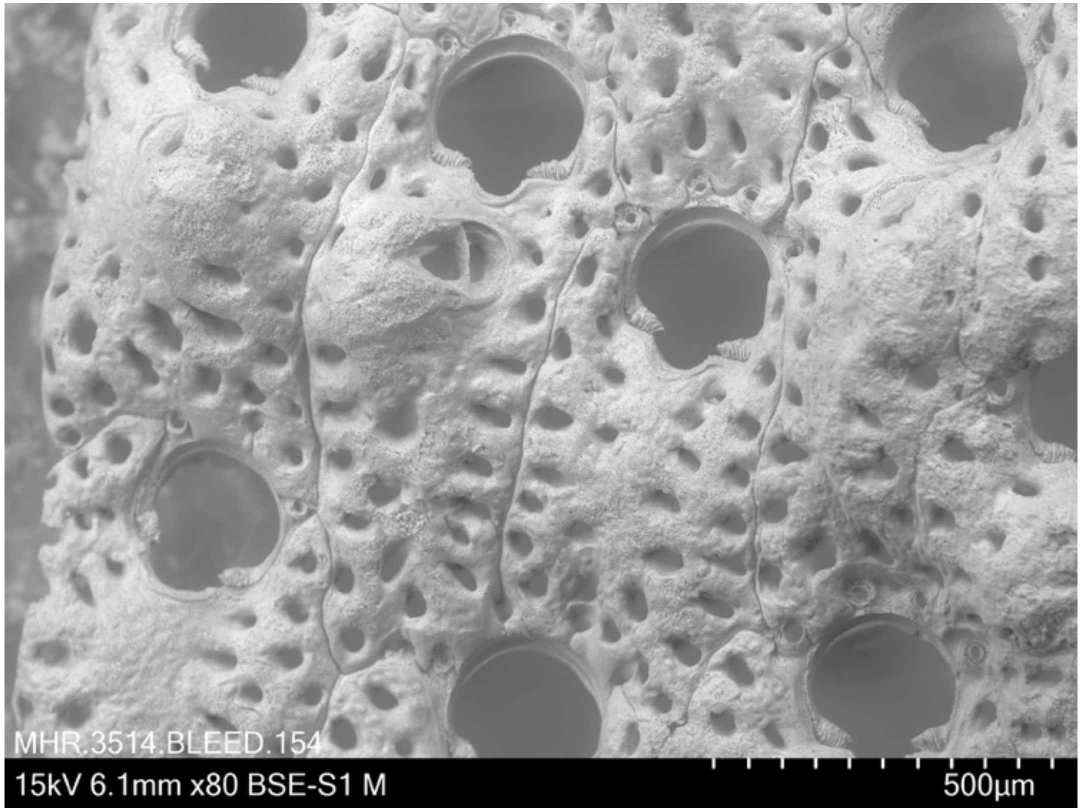


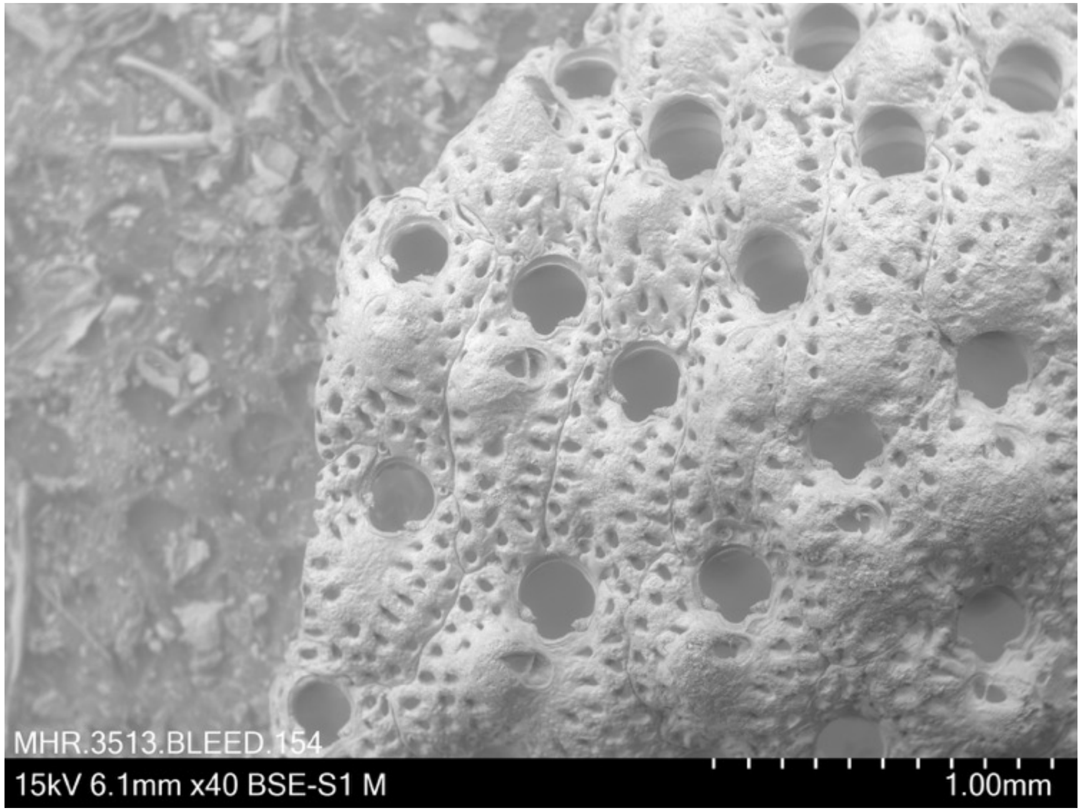


***Chiastosella watersi* BLEED 154**
Coll: 25.01.2010 (A. M. Smith)
Lat: -46.93
Long: 168.16
Depth: 38 m

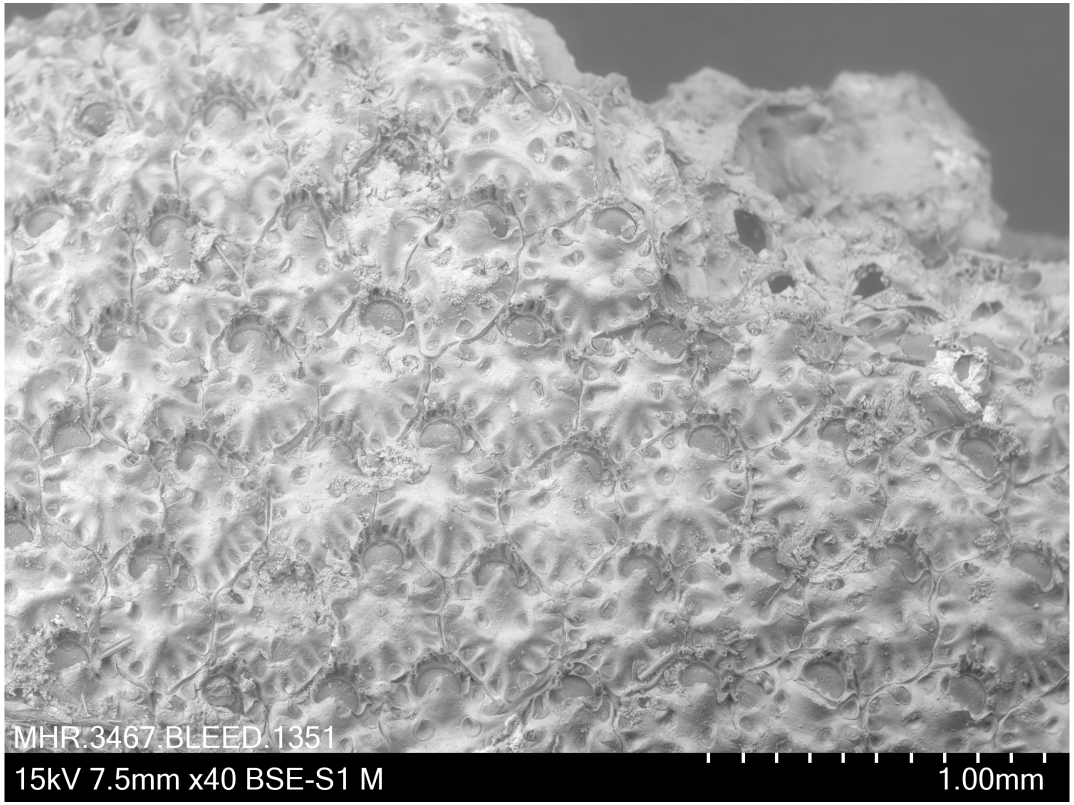


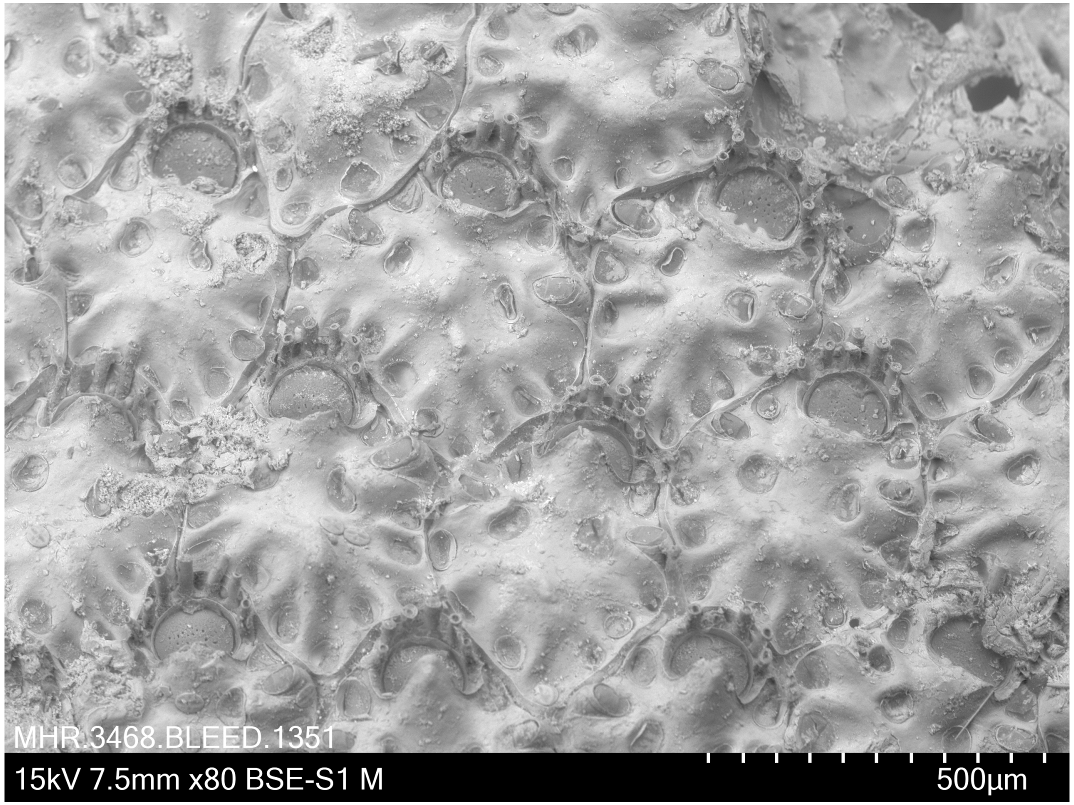


***Chiastosella jellyae*** **BLEED 1351**
Coll: 17.3.2019 (A. M. Smith, D. P. Gordon)
Lat: -34.21
Long: 172.11
Depth: 104 m

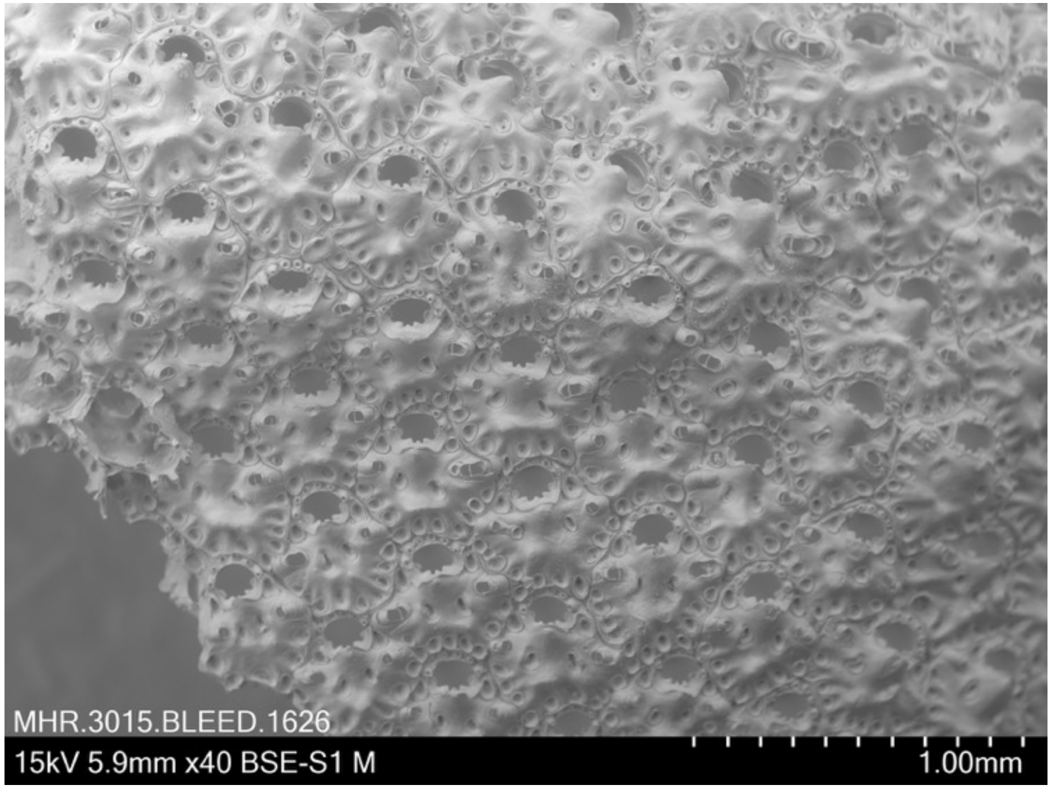


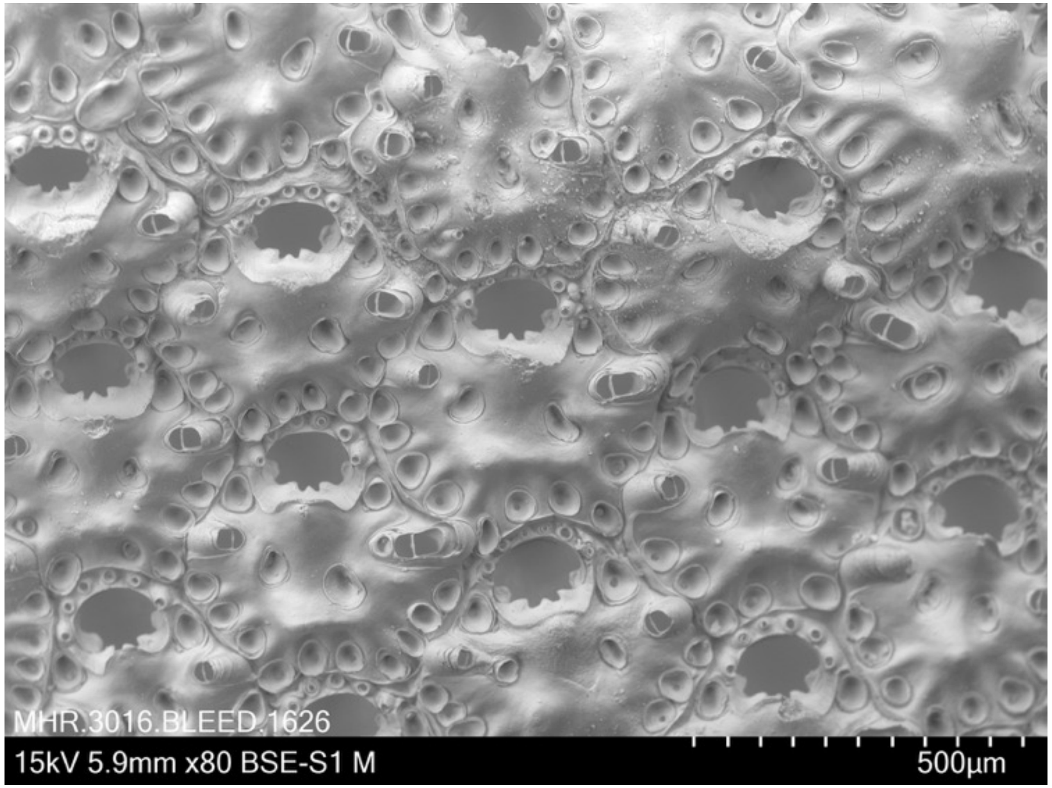


***Chiastosella jellyae* BLEED 1626**
Coll: 17.3.2019 (A. M. Smith, D. P. Gordon)
Lat: -34.159999999999997
Long: 172.09
Depth: 82.9 m
**Notes:** This is formally still *Exochella jellyae* Brown 1952, as of 15.10.2020, but it is morphologically and molecularly both *Chiastosella*

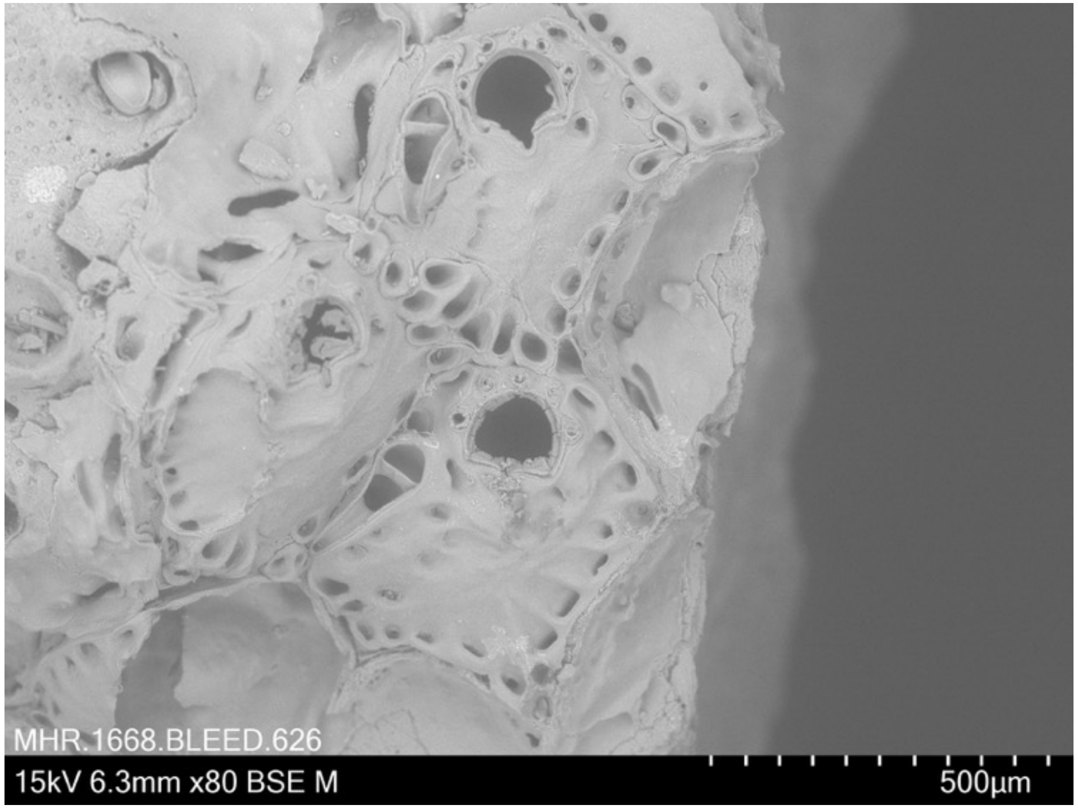


***Chiastosella enigma*** **BLEED 626B**
Coll: 10.4.2018 (A. M. Smith, D. P. Gordon, H. Mello)
Lat: -46.9449999999999
Long: 168.158333333333
Depth: 42 m

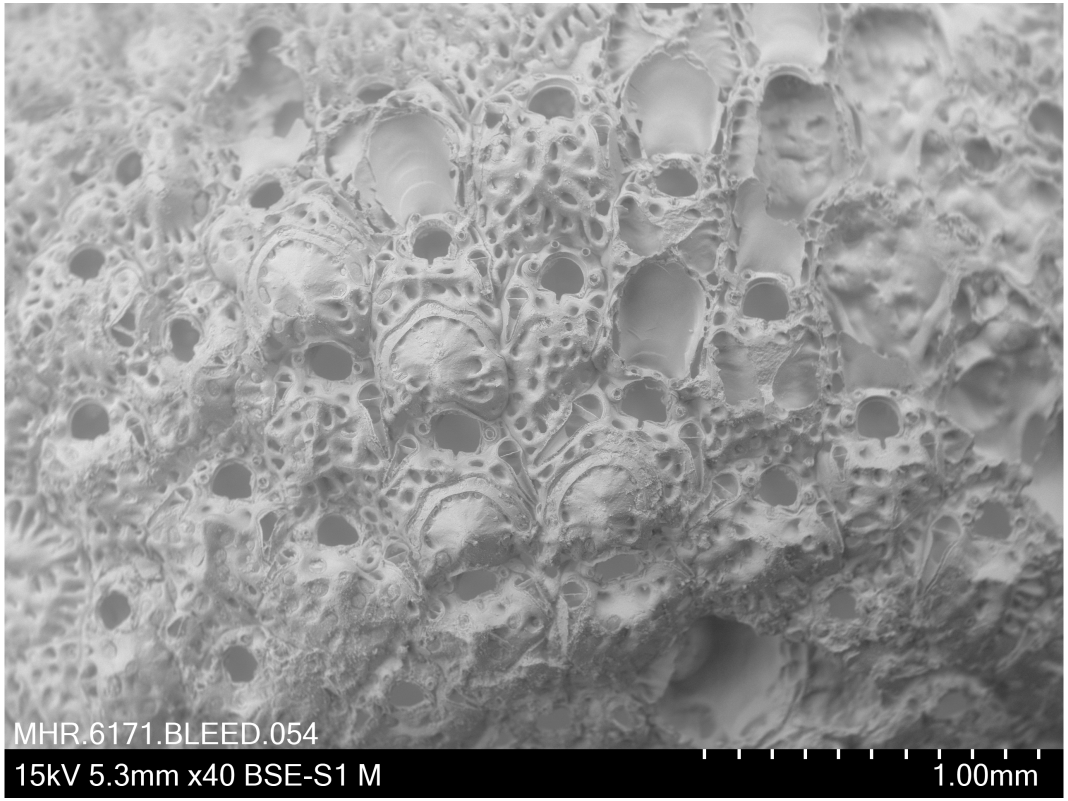


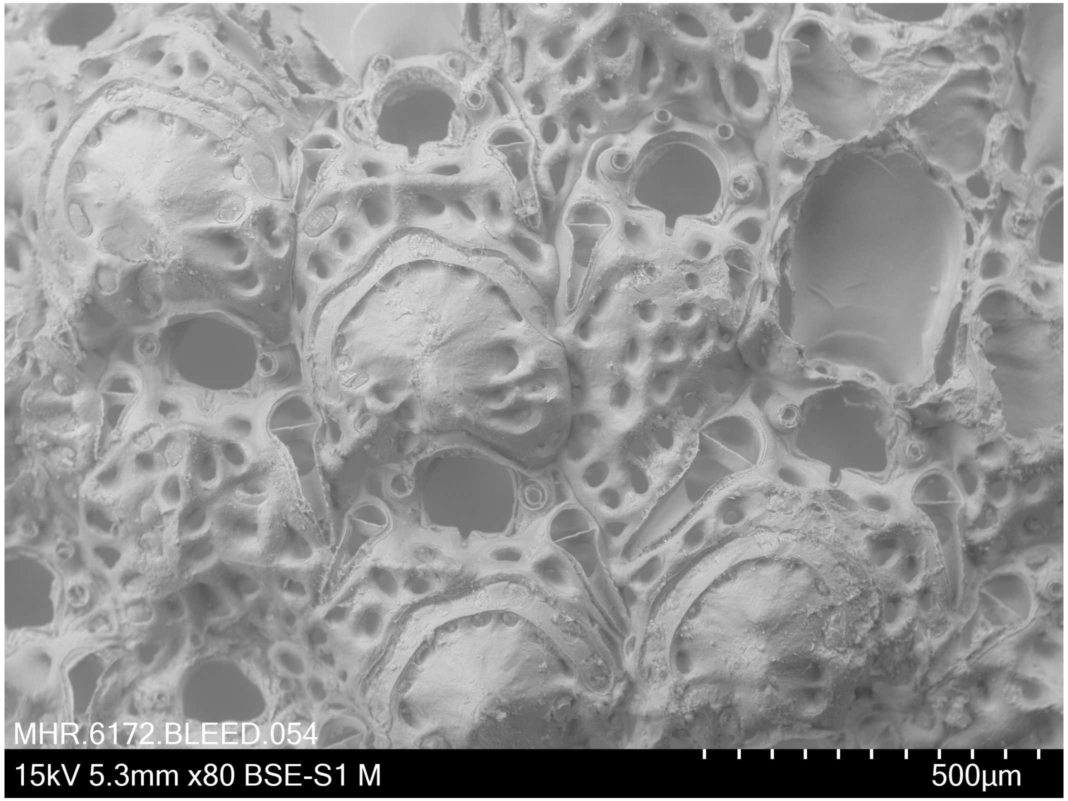


***Chiastosella enigma* BLEED 54**
NIWA 74935 (TAN1108/138)
Coll: 24.05.2011
Lat: -45.5893333
Long: 171.0598333
Depth: 103 m

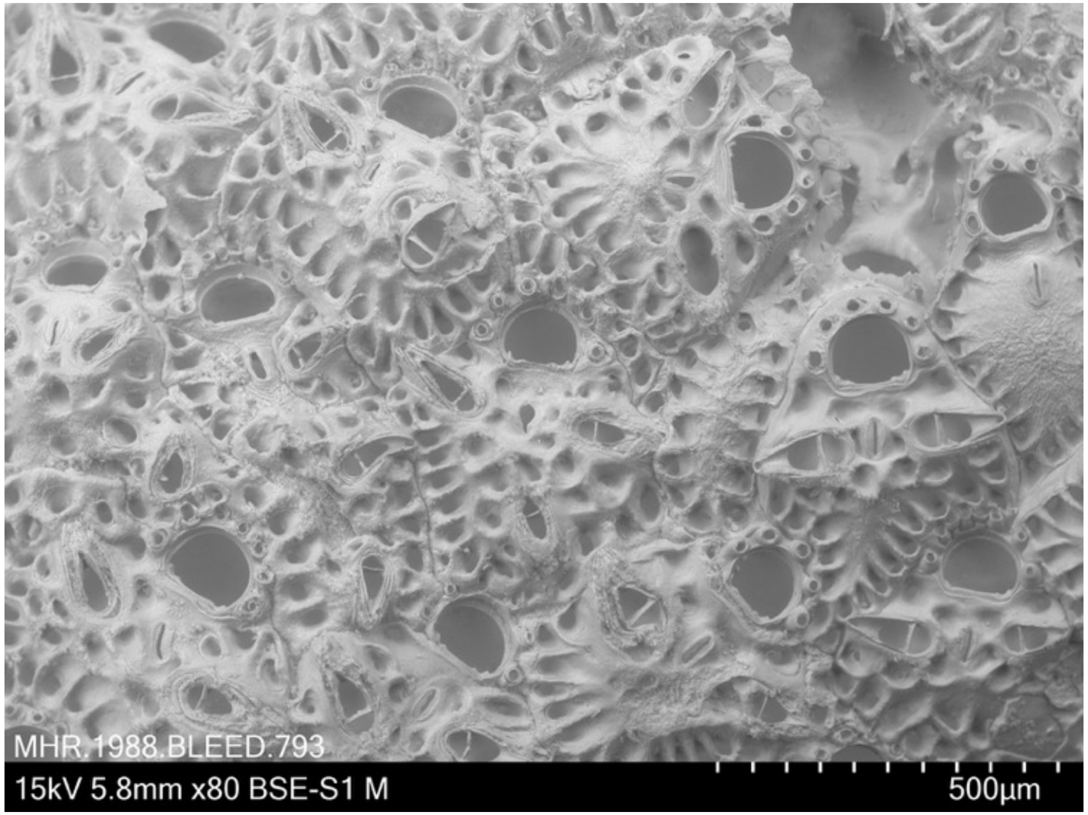


***Calloporina angustipora* BLEED 793**
Coll: 4.3.2018 (D. P. Gordon, L. H. Liow and C. Schack)
Lat: -39.639352S
Long: 177.008705
Depth: intertidal

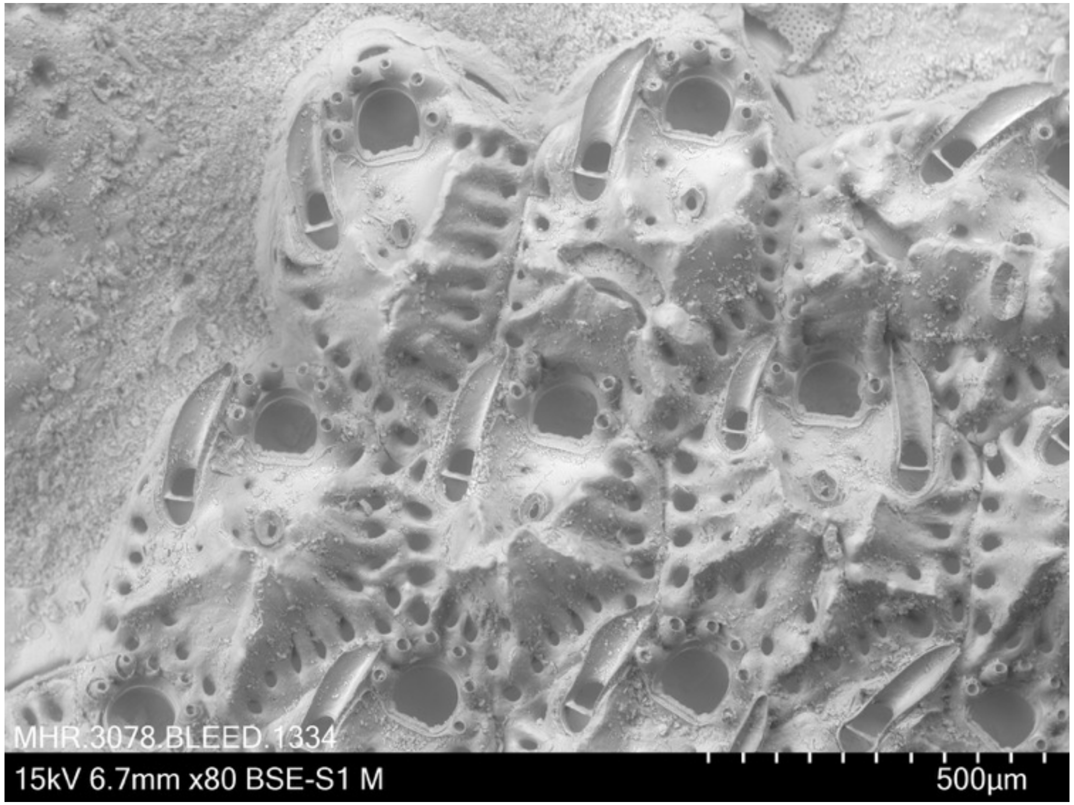


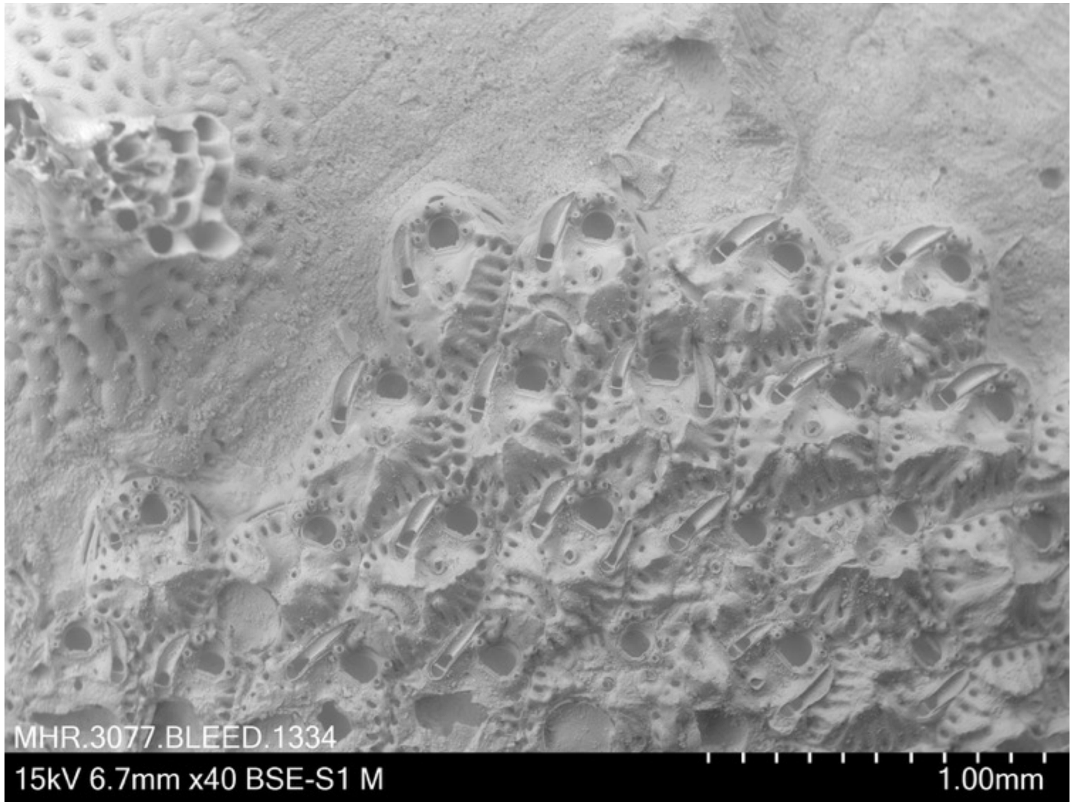


***Calloporina triporosa*** **BLEED 1334**
Coll: 19.3.2019 (A. M. Smith, D. P. Gordon)
Lat: -34.99
Long: 173.97
Depth: 61.6 m

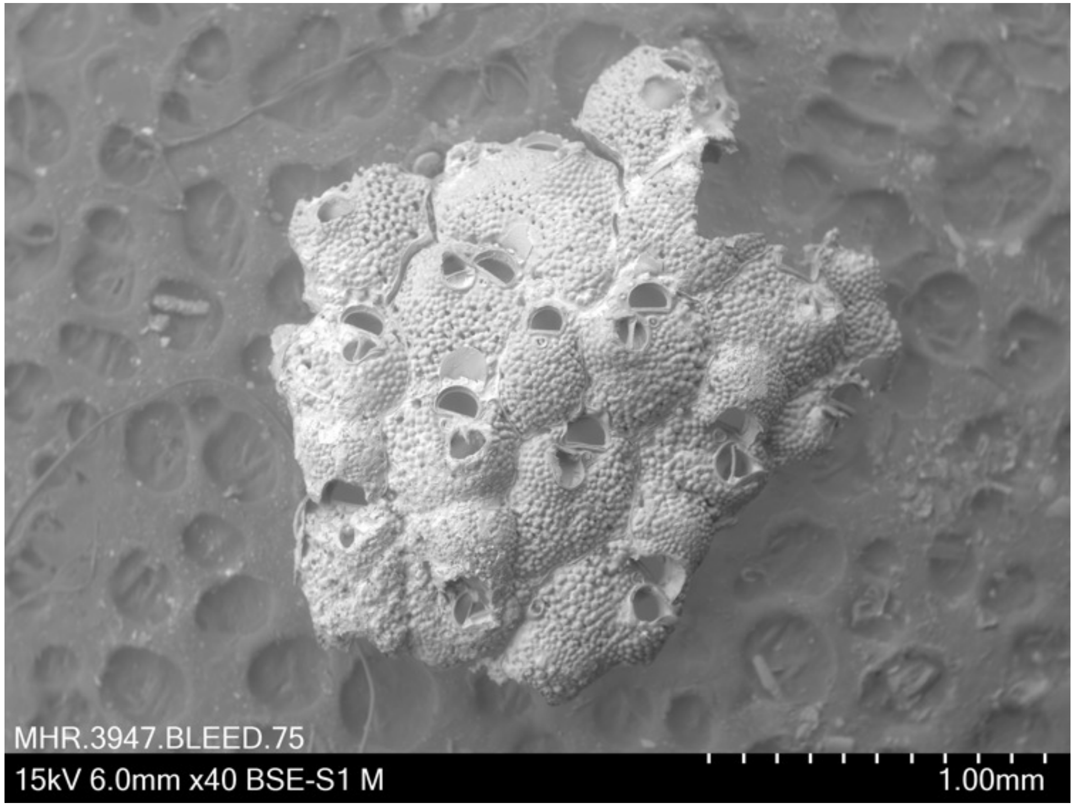


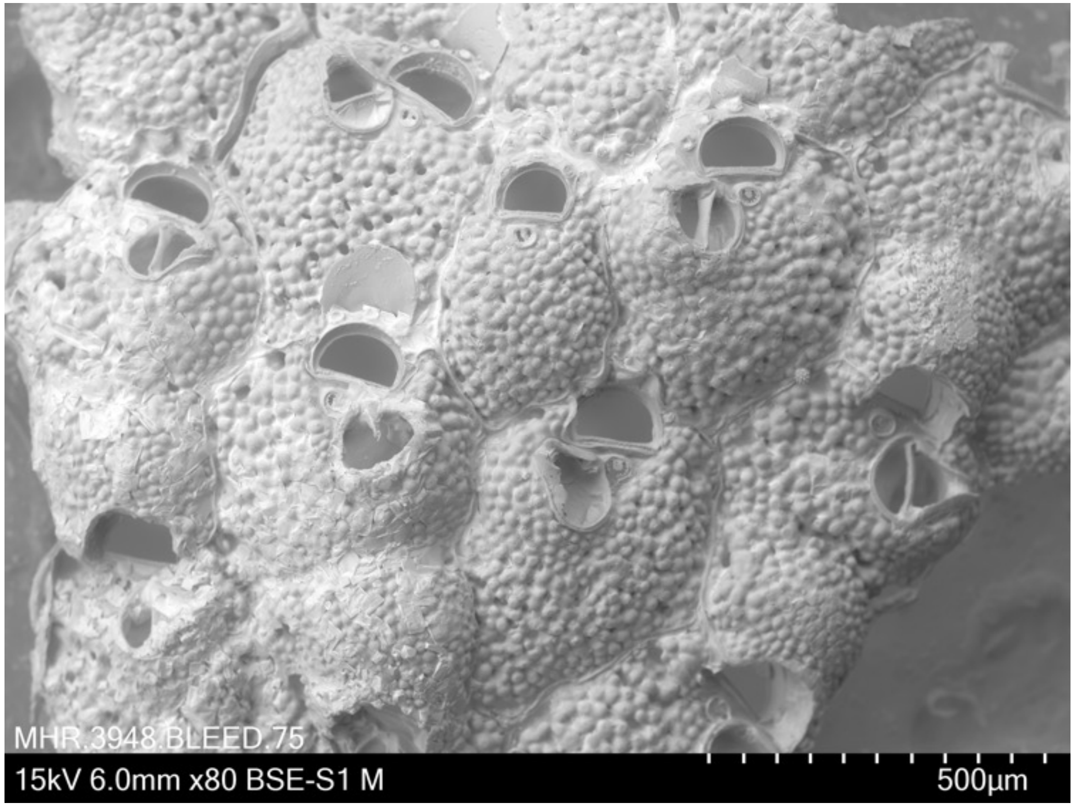


***Microporella* cf. *agonistes* BLEED 75**
NIWA 74426 (TAN1108/66)
Coll: 17.5.2011
Lat: -45.9315
Long: 170.7888333
Depth: 75 m

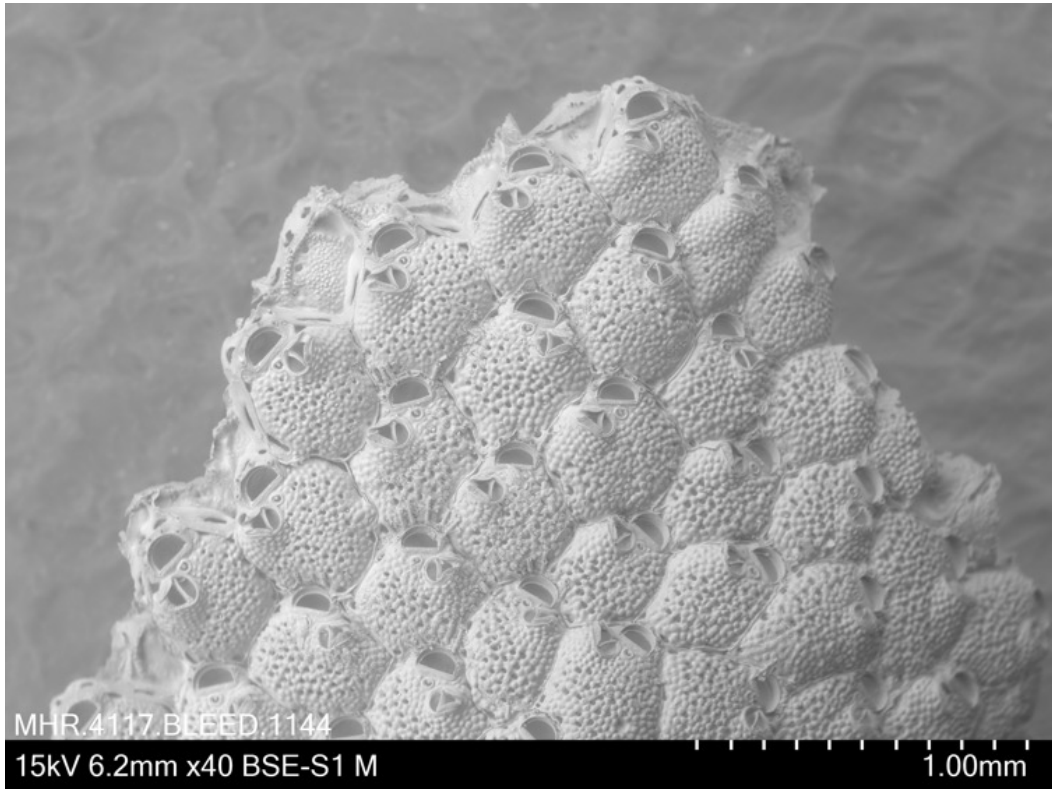


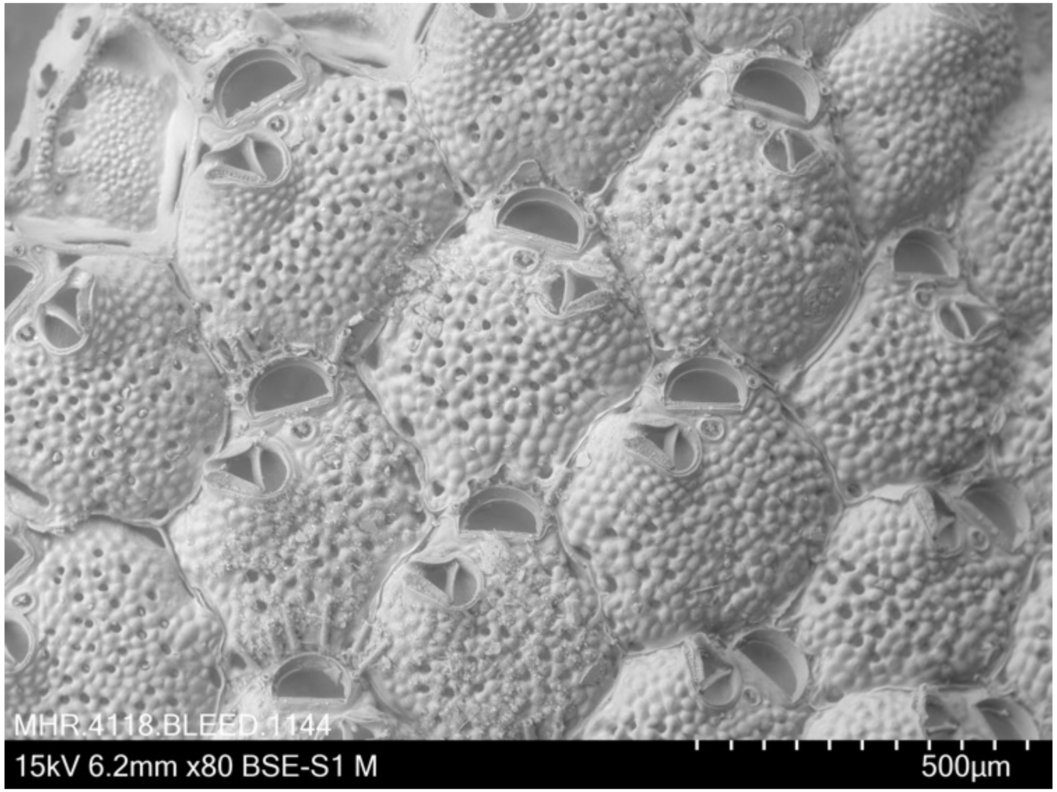


***Microporella* cf. *agonistes* BLEED 1144A**
Coll: 10.11.2018 (A. M. Smith)
Lat: -45.835833000000001
Long: 170.96166700000001
Depth: 131 m

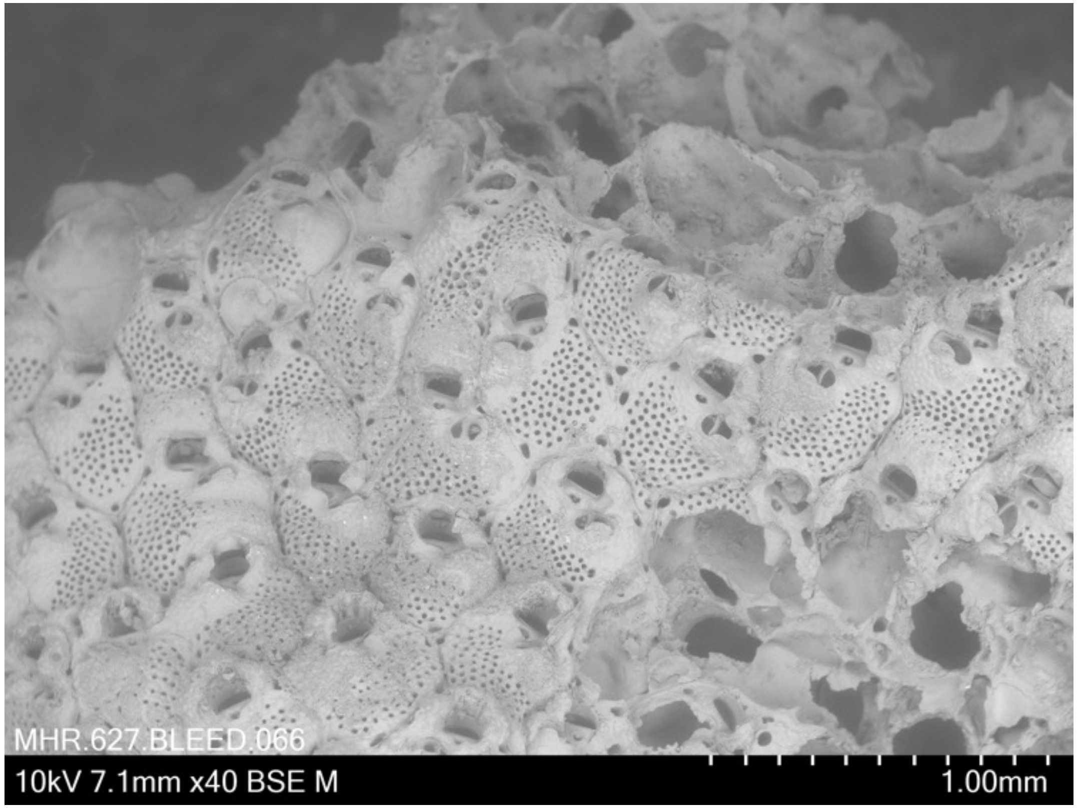


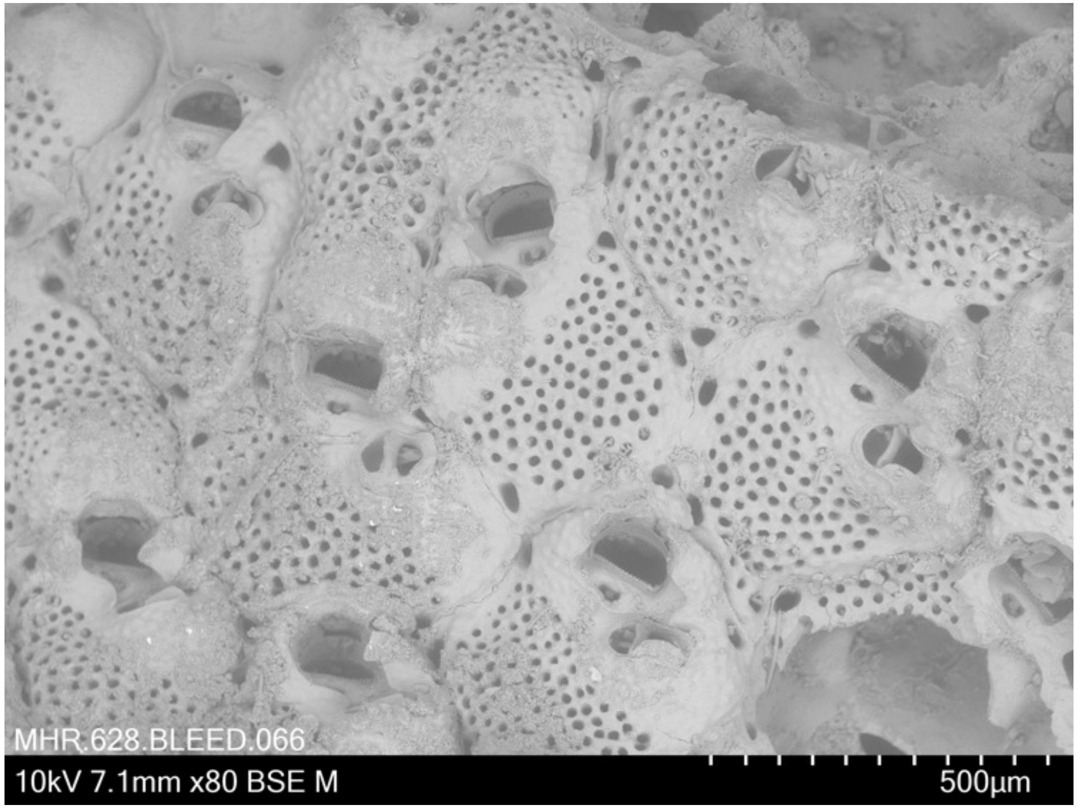


***Microporella* cf. *intermedia* BLEED 66**
NIWA 56622 (TAN0906/159)
Coll: 14.7.2009
Lat: -34.3875999
Long: 173.041
Depth: 59 m

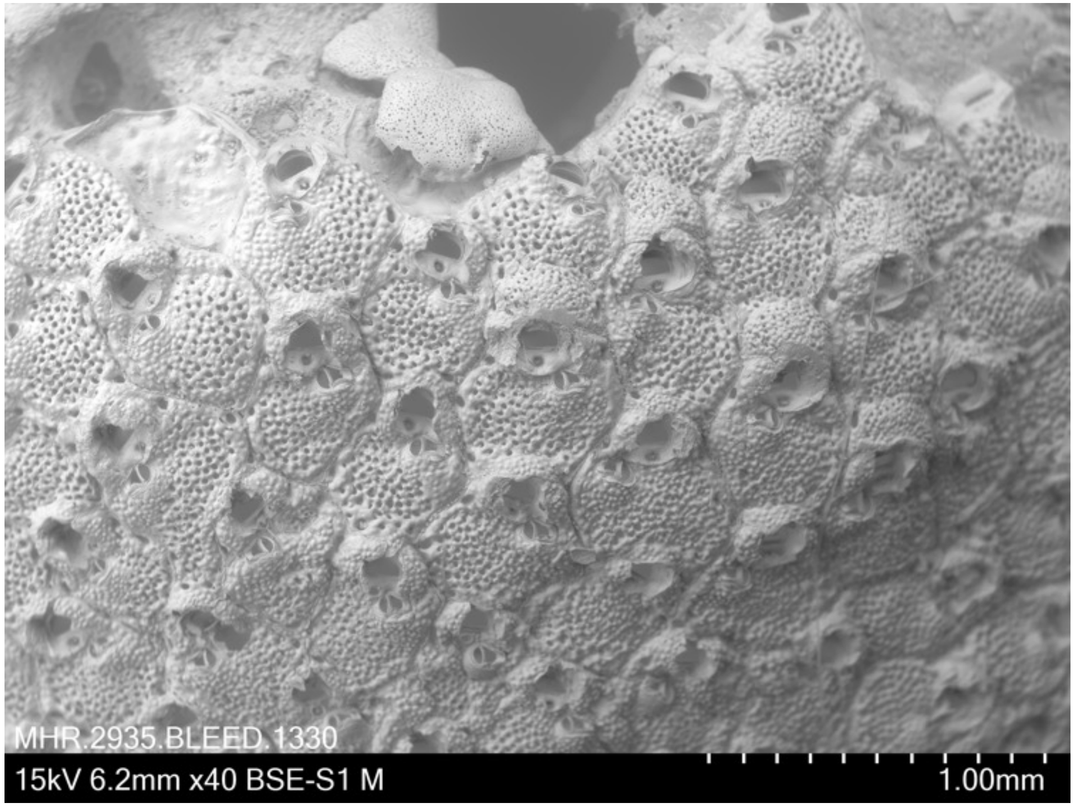


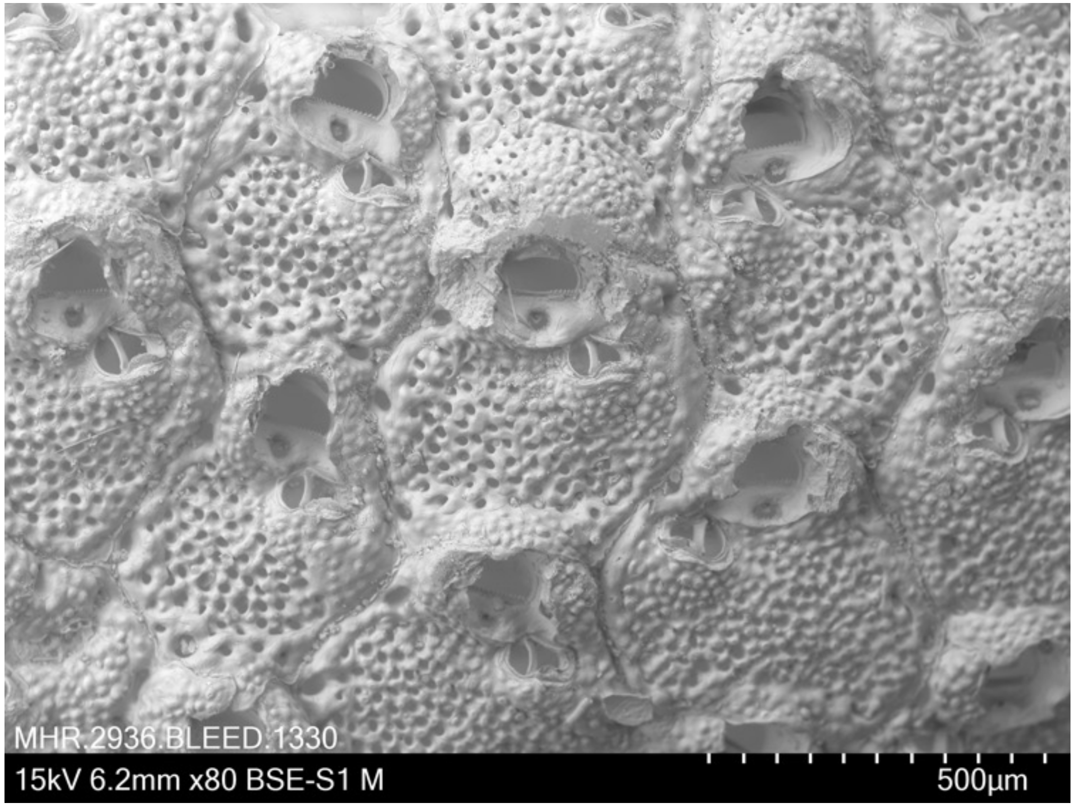


***Microporella* cf. *intermedia* BLEED 1330**
Coll: 19.3.2019 (A. M. Smith; D. P. Gordon)
Lat: -34.99
Long: 173.97
Depth: 61.6 m

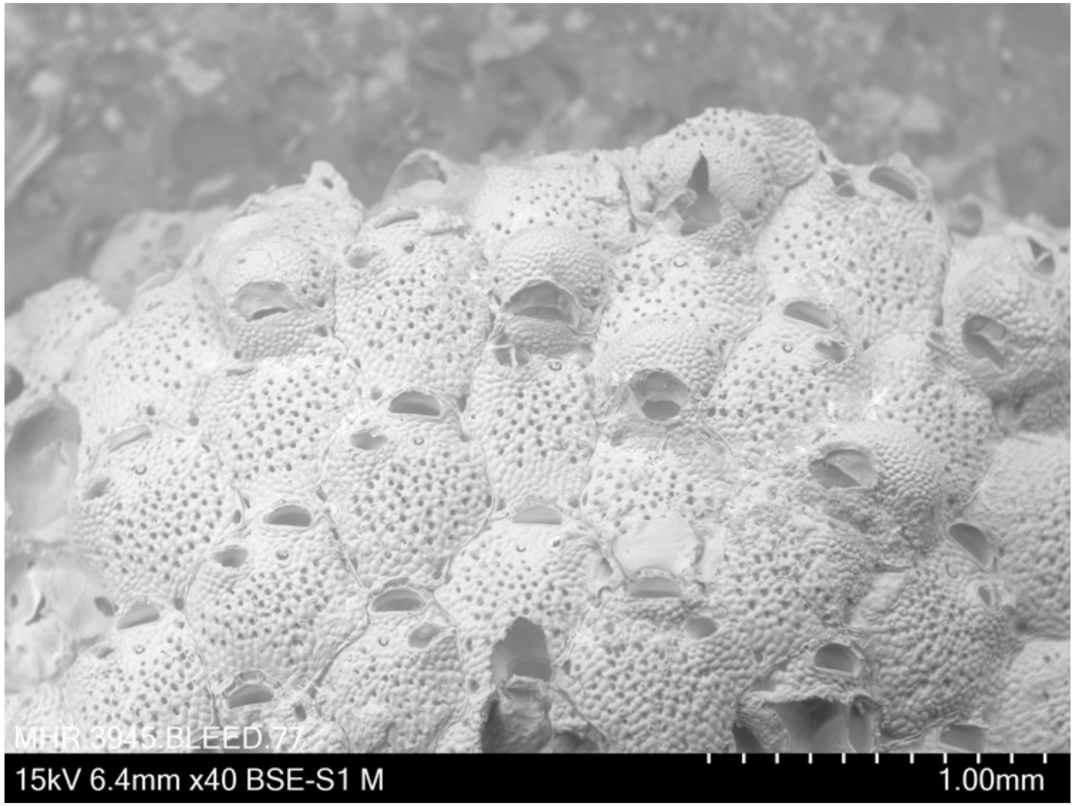


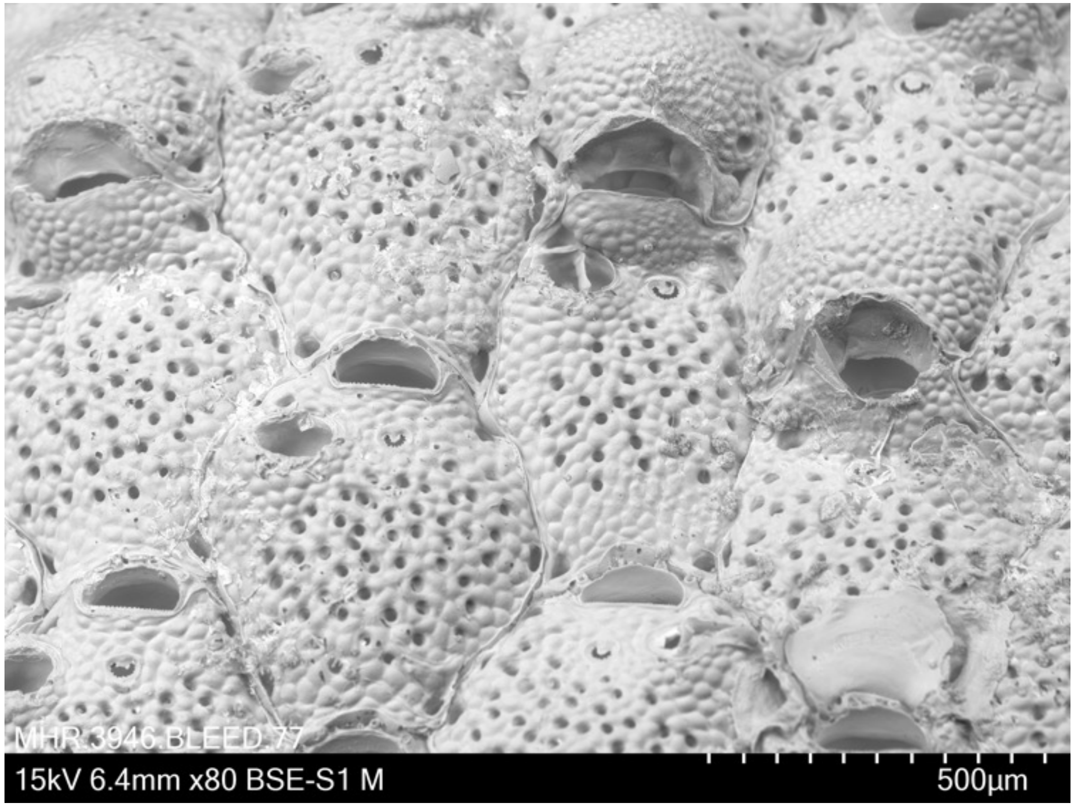


***Microporella* aff. *diademata* BLEED 77**
NIWA 39690 (TAN0803/63)
Coll: 09.04.2008
Lat: -52.487333
Long: 160.415
Depth: 350 m

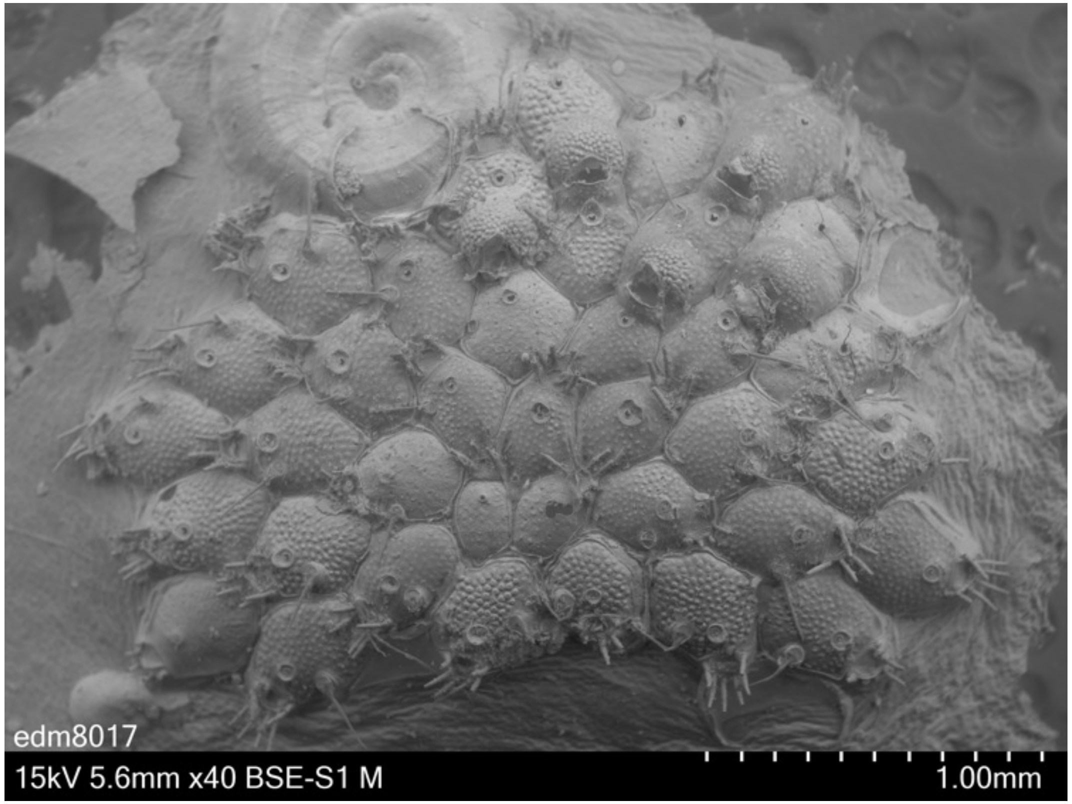


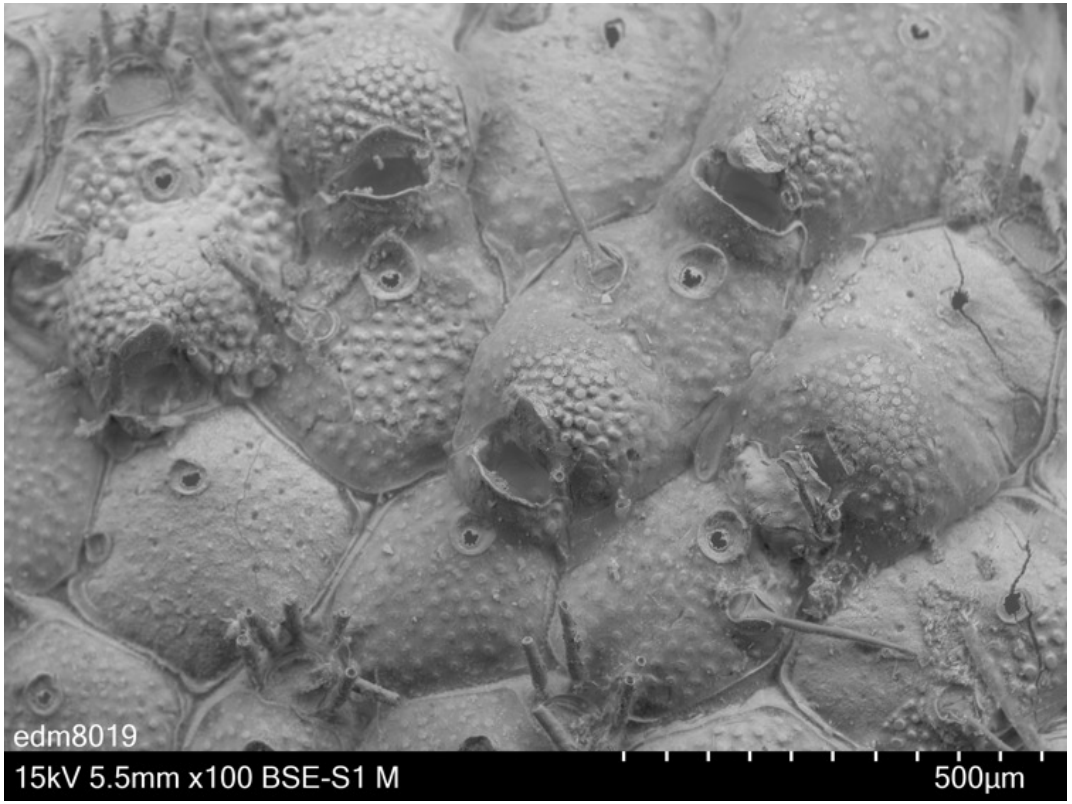


***Microporella* n. sp. BLEED 709**
Coll: 10.4.2018 (A. M. Smith, D. P. Gordon, H. Mello)
Lat: -46.9742
Long: 168.16694
Depth: 13 m

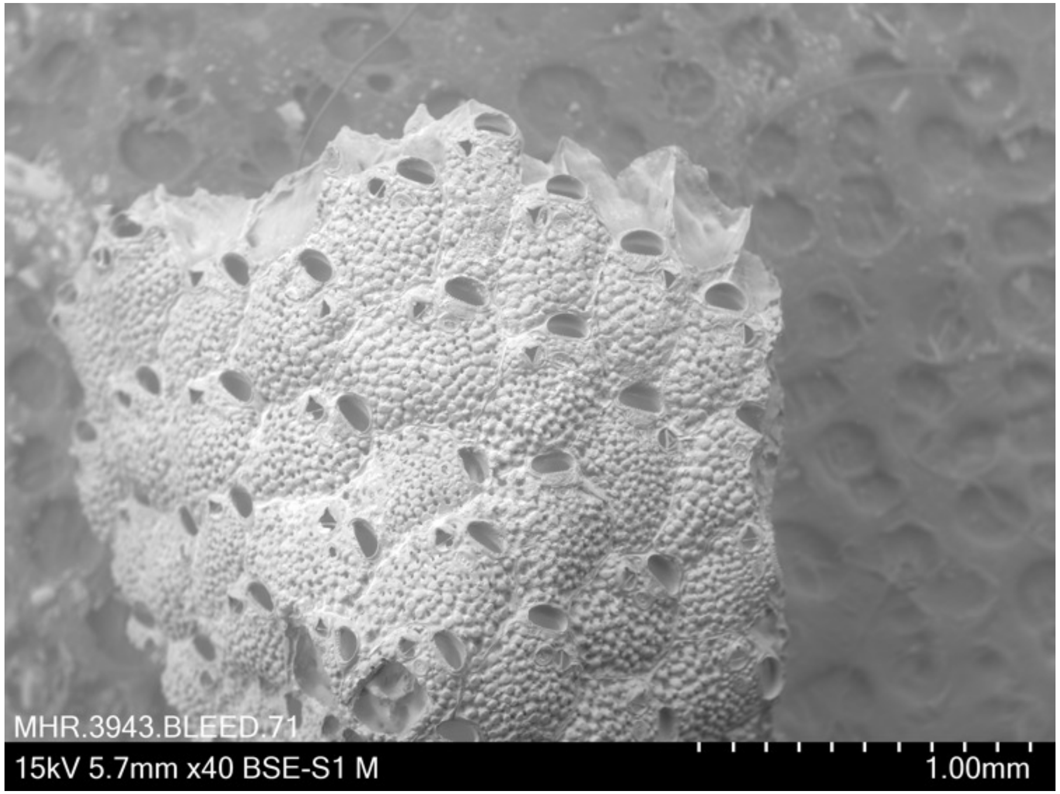


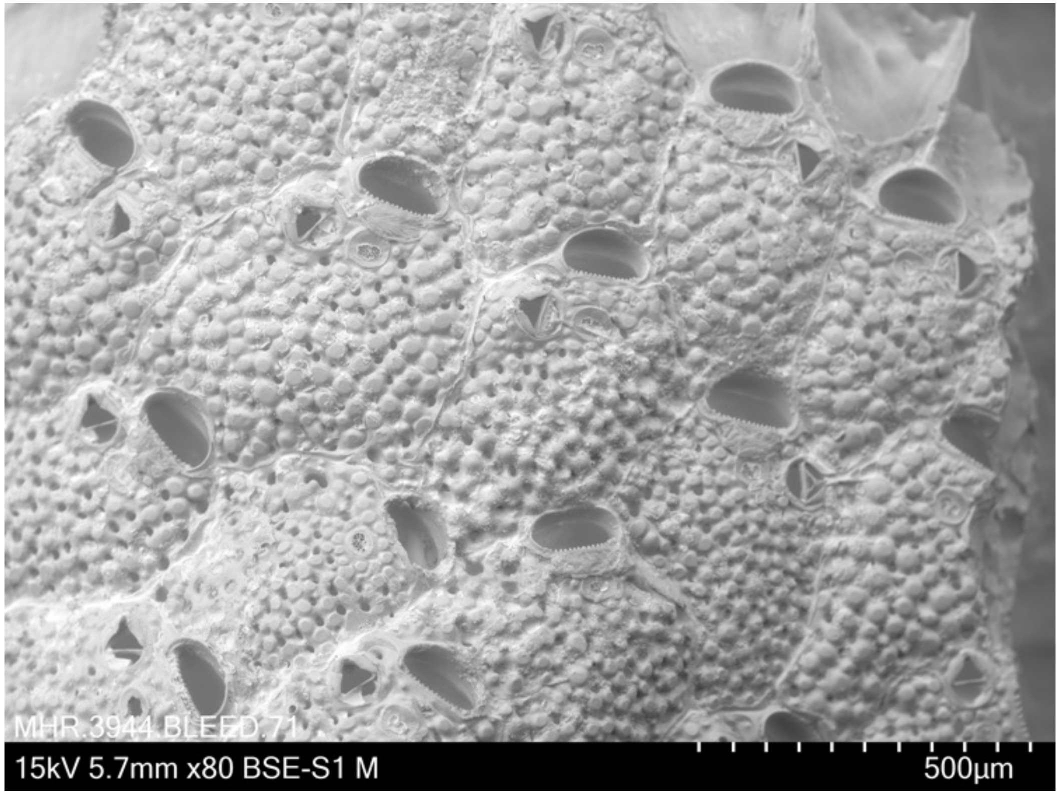


***Microporella ordo*** **BLEED 71**
NIWA 73295 (TAN1105/43)
Coll: 28.3.2011
Lat: -33.9875
Long: 171.75083
Depth: 170-174 m

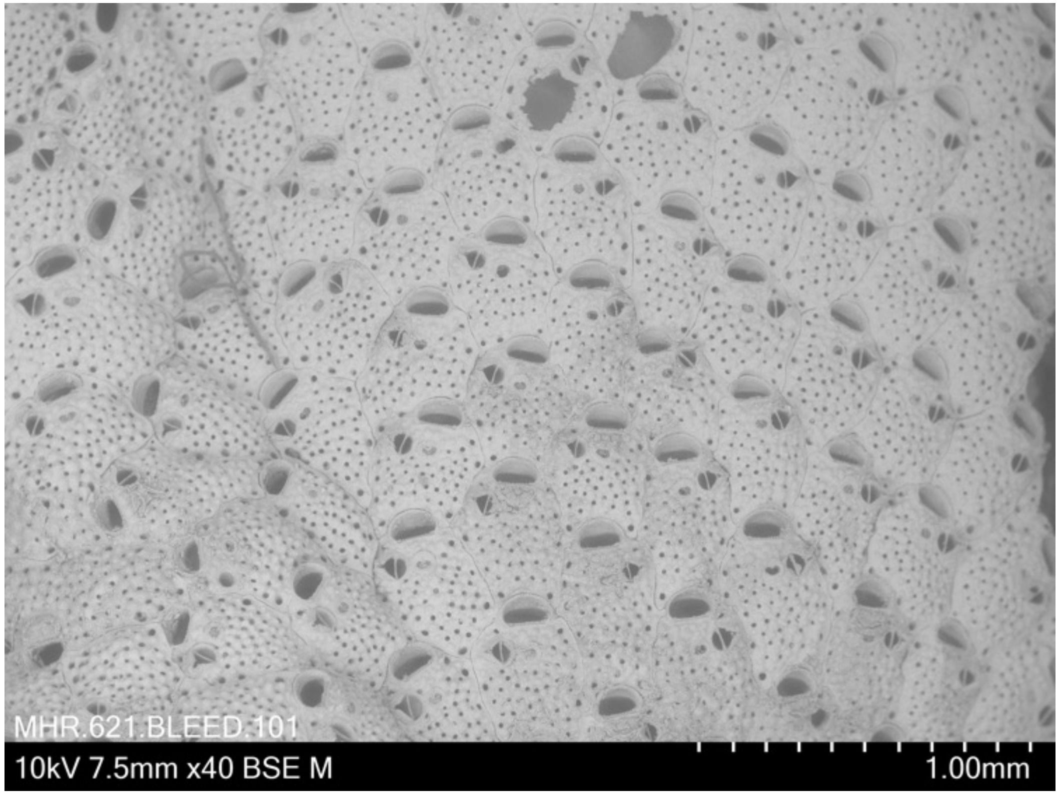


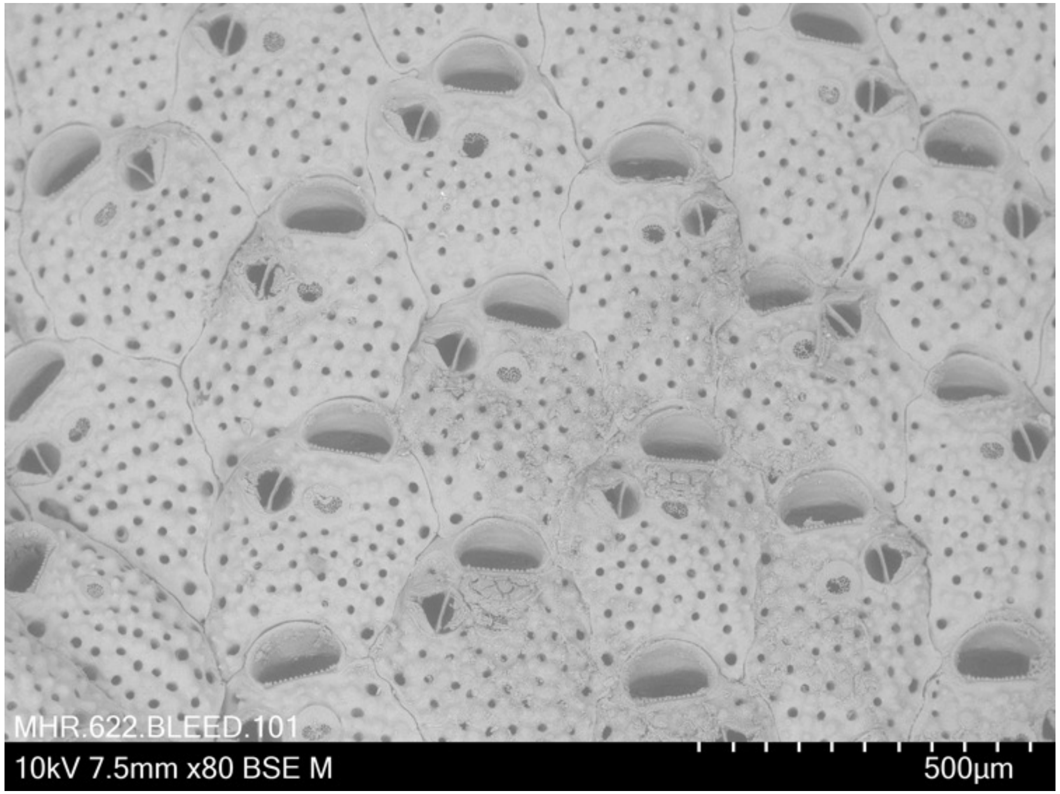


***Microporella ordo* BLEED 101A**
NIWA 56945 (TAN0906/170)
Coll: 15.7.2009
Lat: -34.3781667
Long: 173.0265
Depth: 65 m

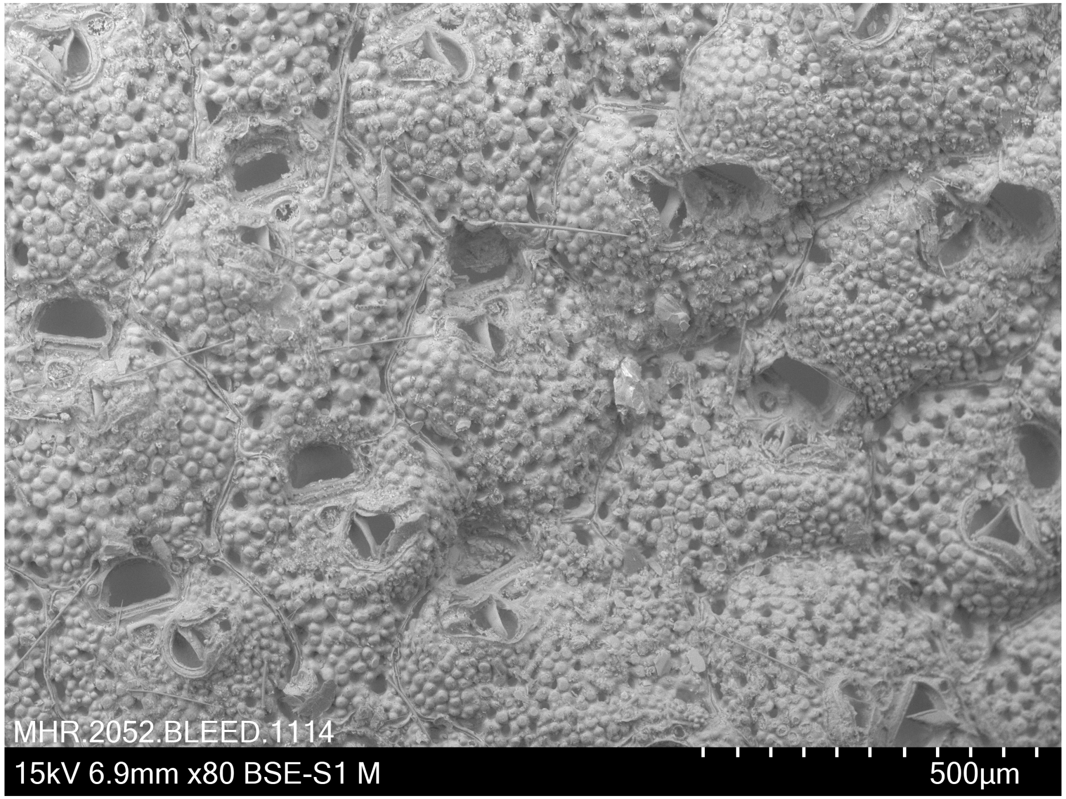


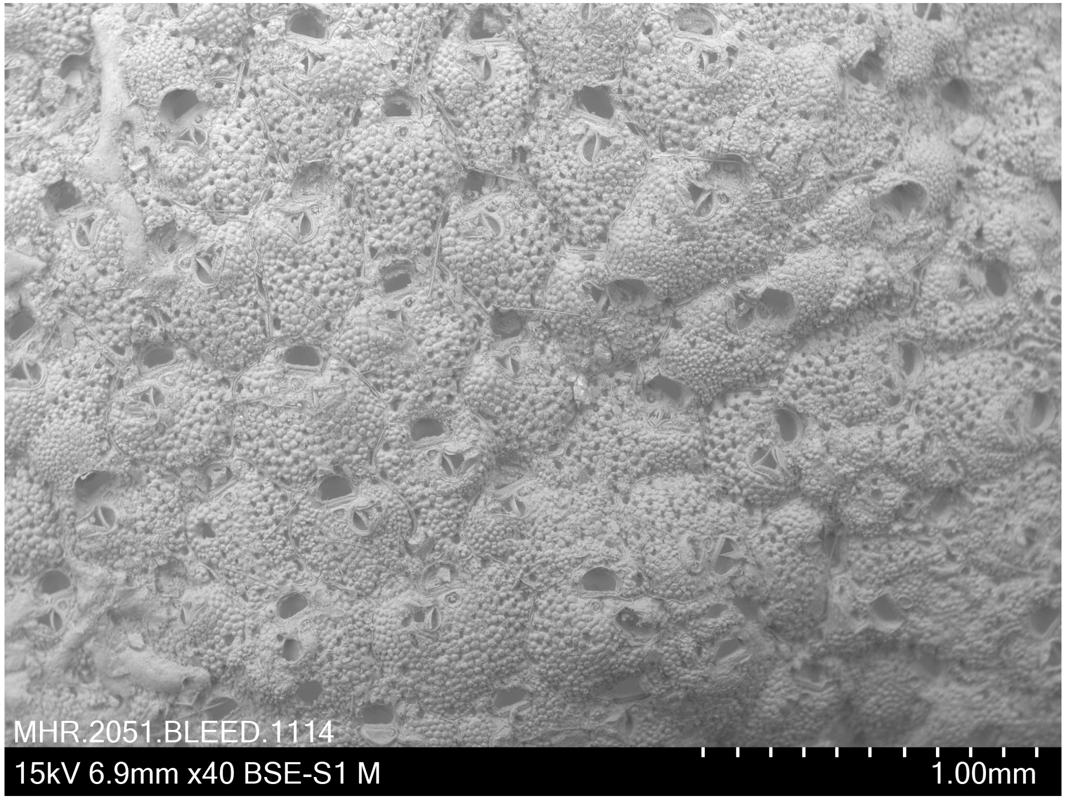


***Microporella* cf. *agonistes* BLEED 1114**
Coll: 10.11.2018 (A. M. Smith)
Lat: -45.822221999999996
Long: 170.880833
Depth: 80.4 m

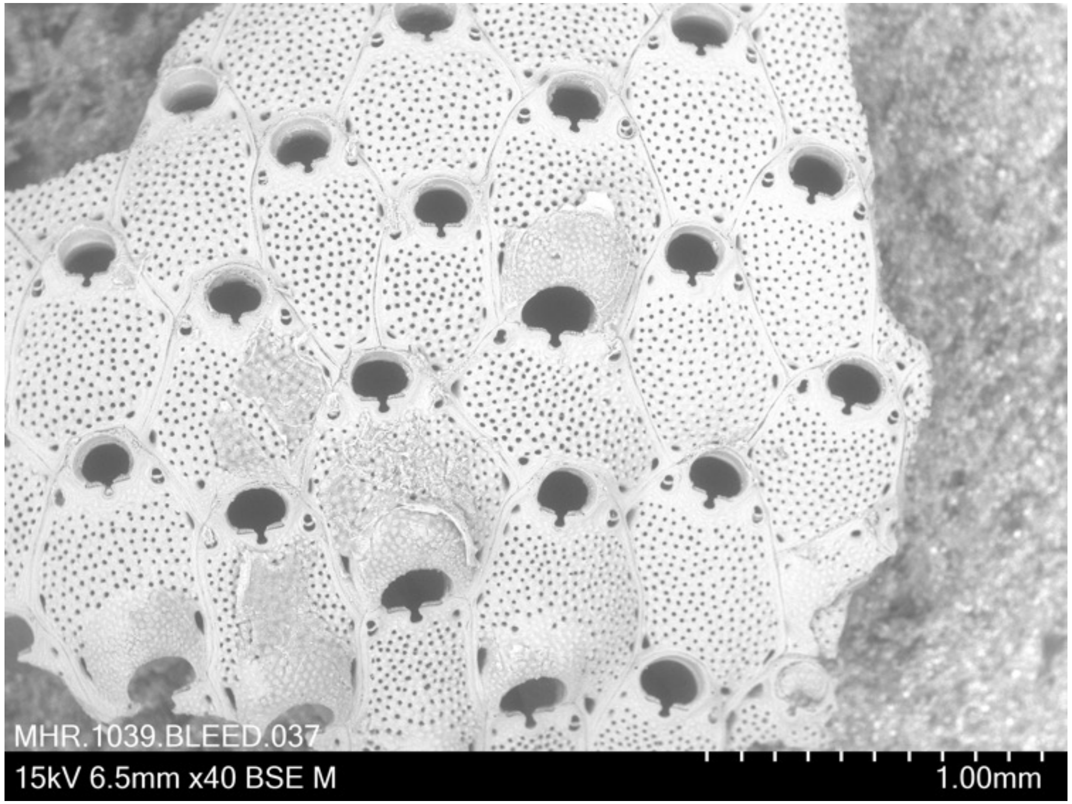


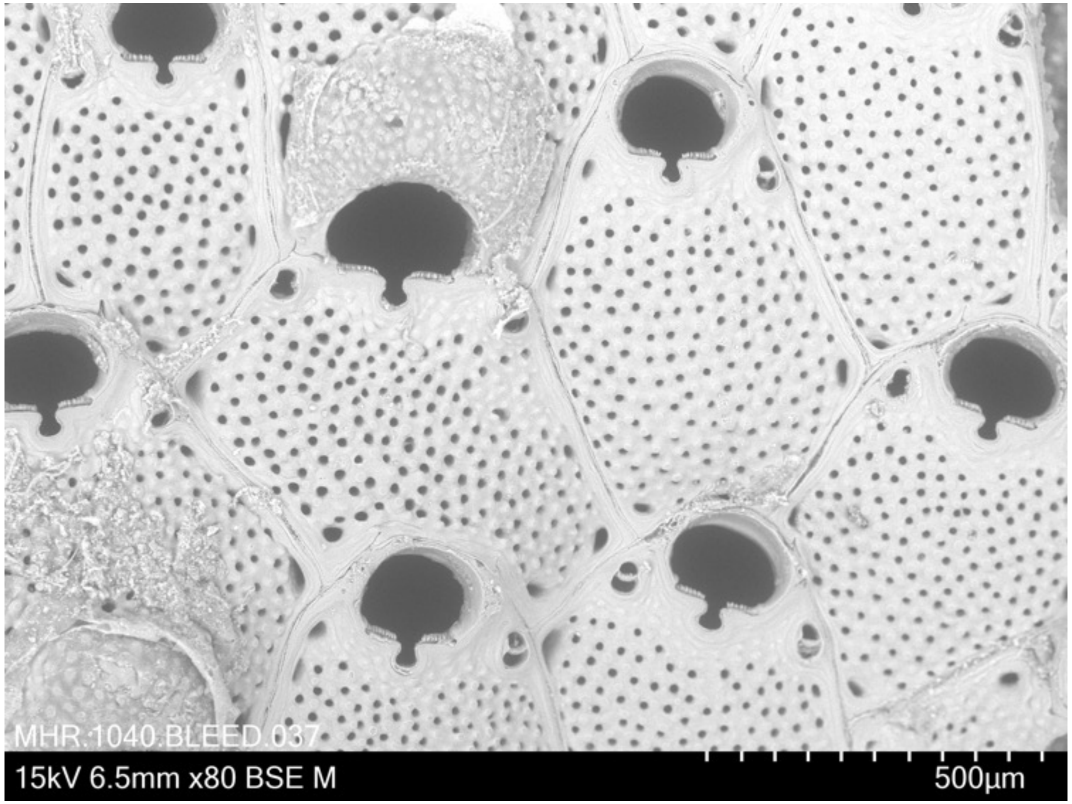


***Taylorus arcuatus* BLEED 37**
Coll: 28.1.2010 (A. M. Smith)
Lat: -47.27
Long: 167.3
Depth: 125 m

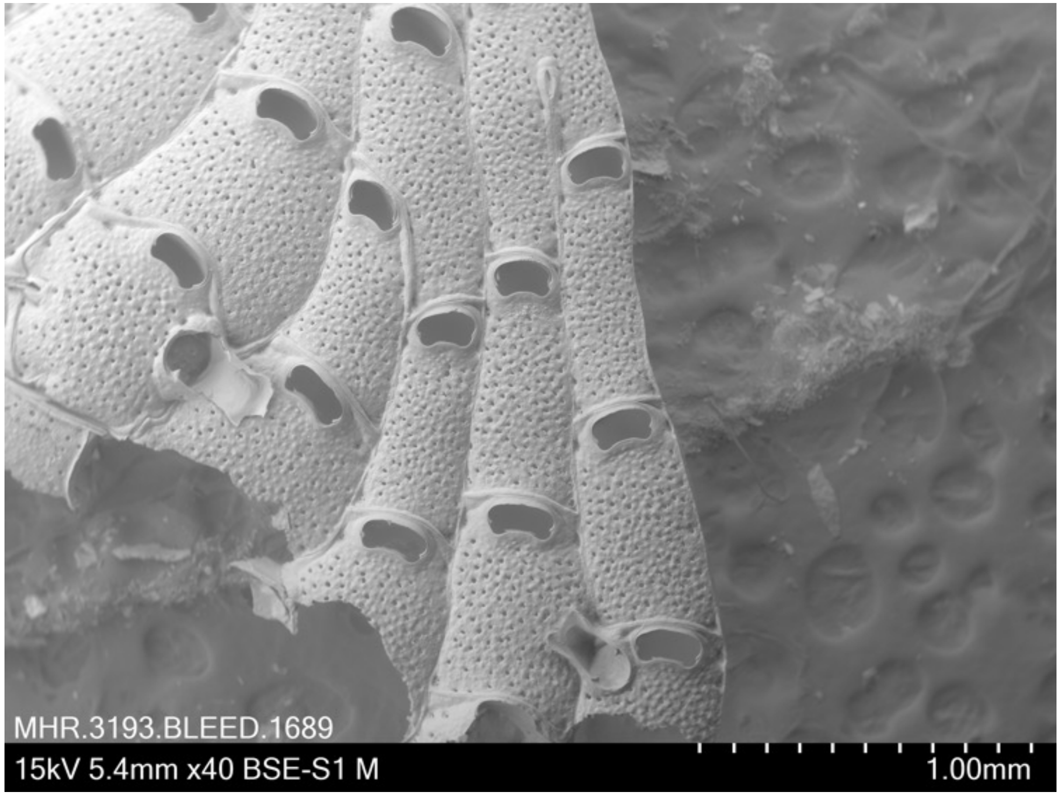


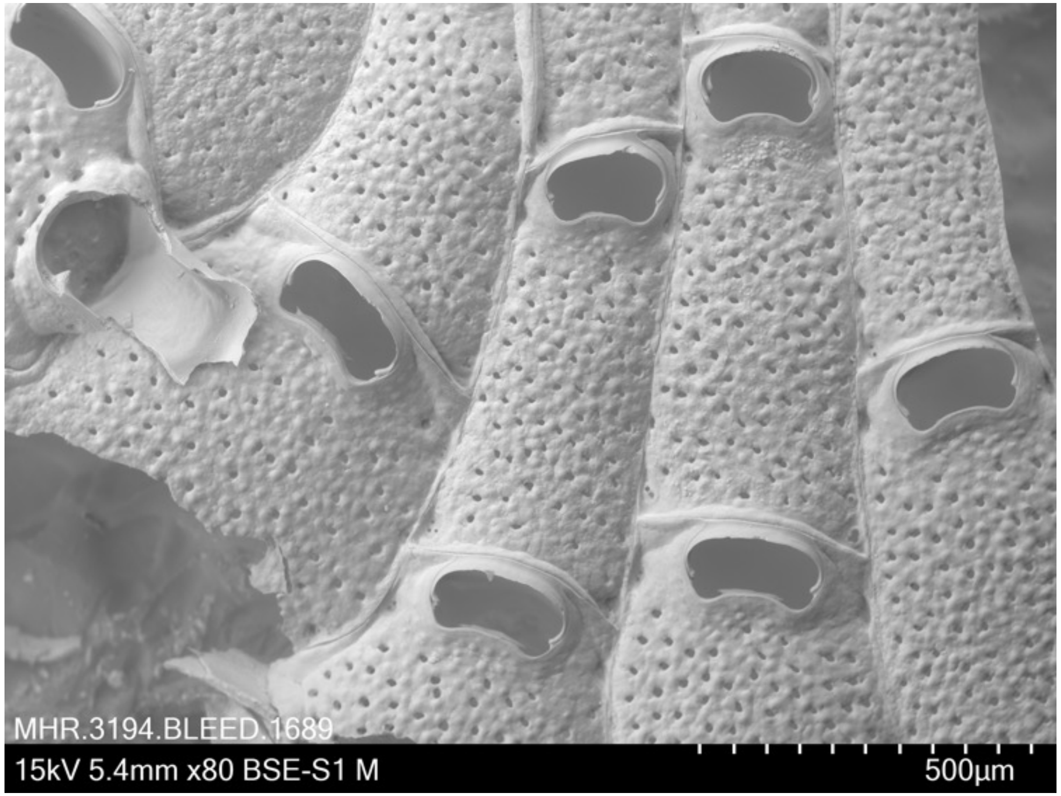


***Powellitheca* n. sp. BLEED 1689**
Coll: 17.3.2019 (A. M. Smith, D. P. Gordon)
Lat: -34.21
Long: 172.11
Depth: 104 m

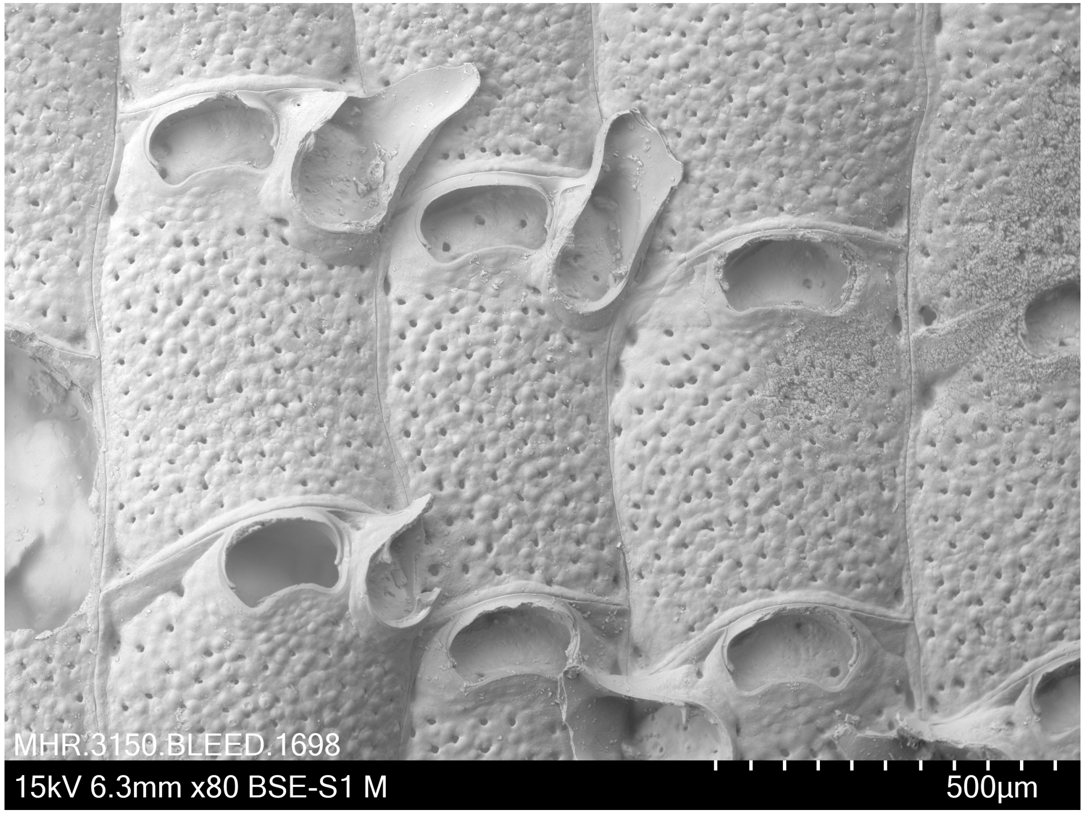


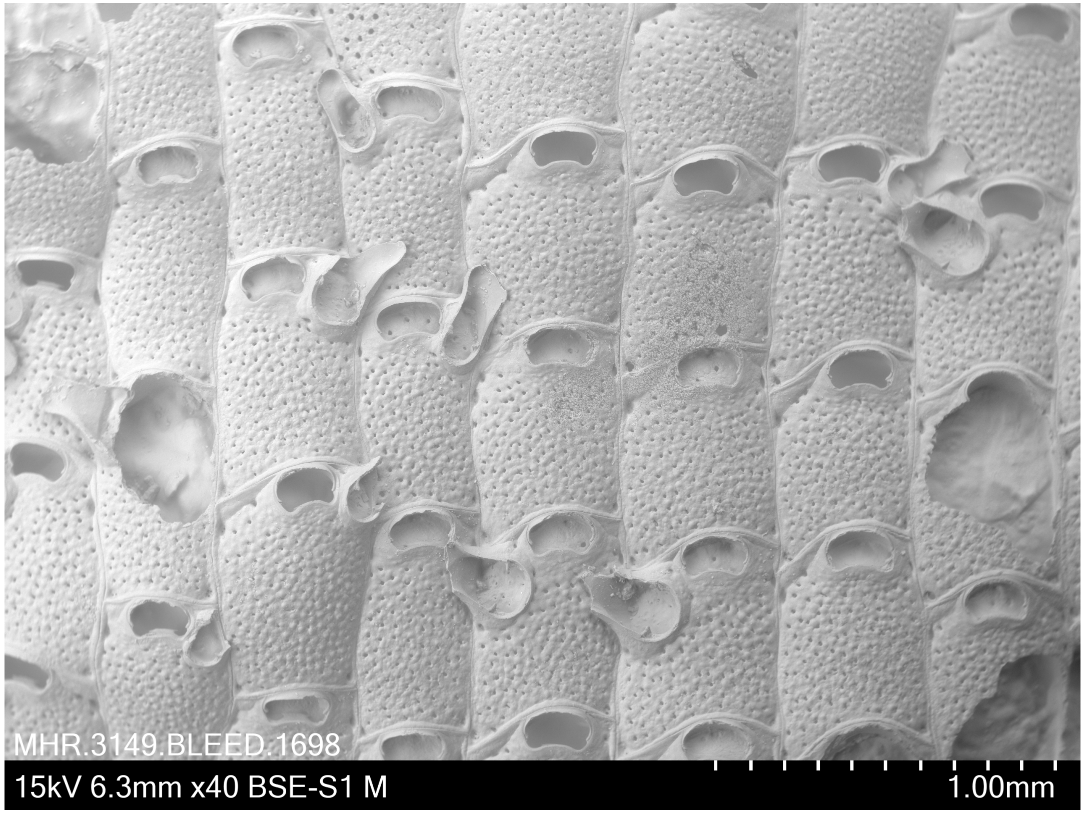


***Powellitheca* n. sp. BLEED 1697**
Coll: 17.3.2019 (A. M. Smith, D. P. Gordon)
Lat: -34.21
Long: 172.11
Depth: 104 m
**Notes:** BLEED 1697 was sequenced, its subsample for SEM was destroyed during handling, but BLEED 1698 is used as a voucher as they are from the same substrate

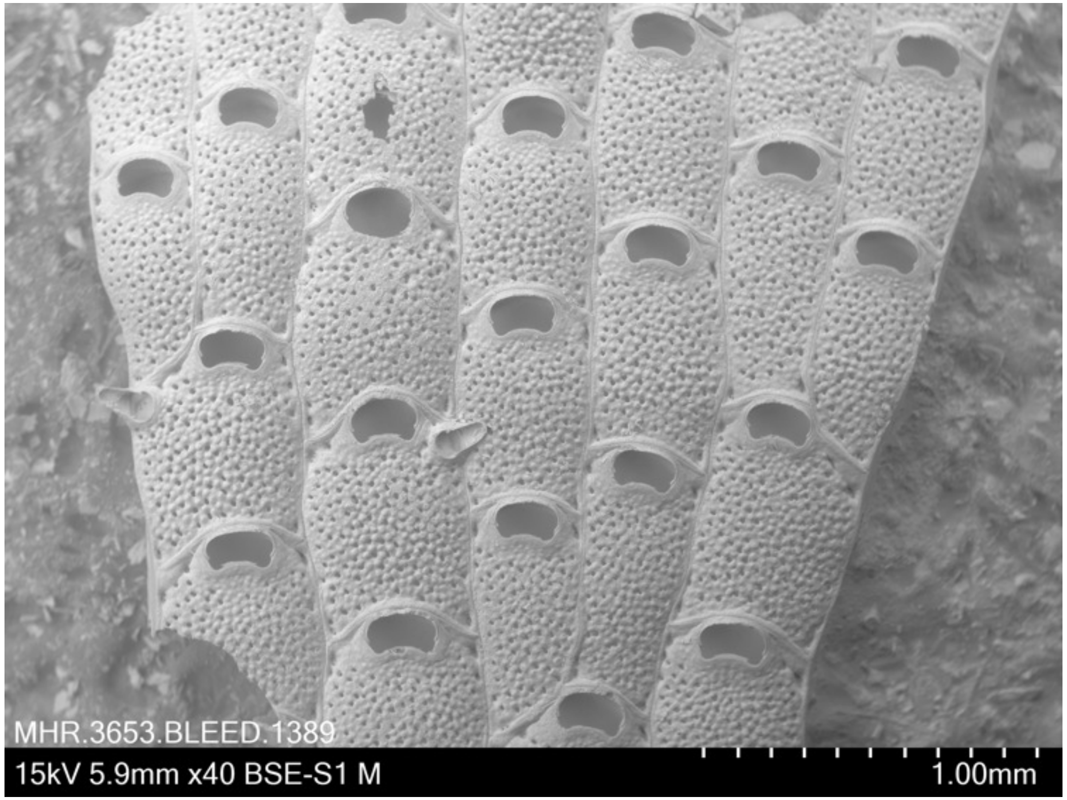


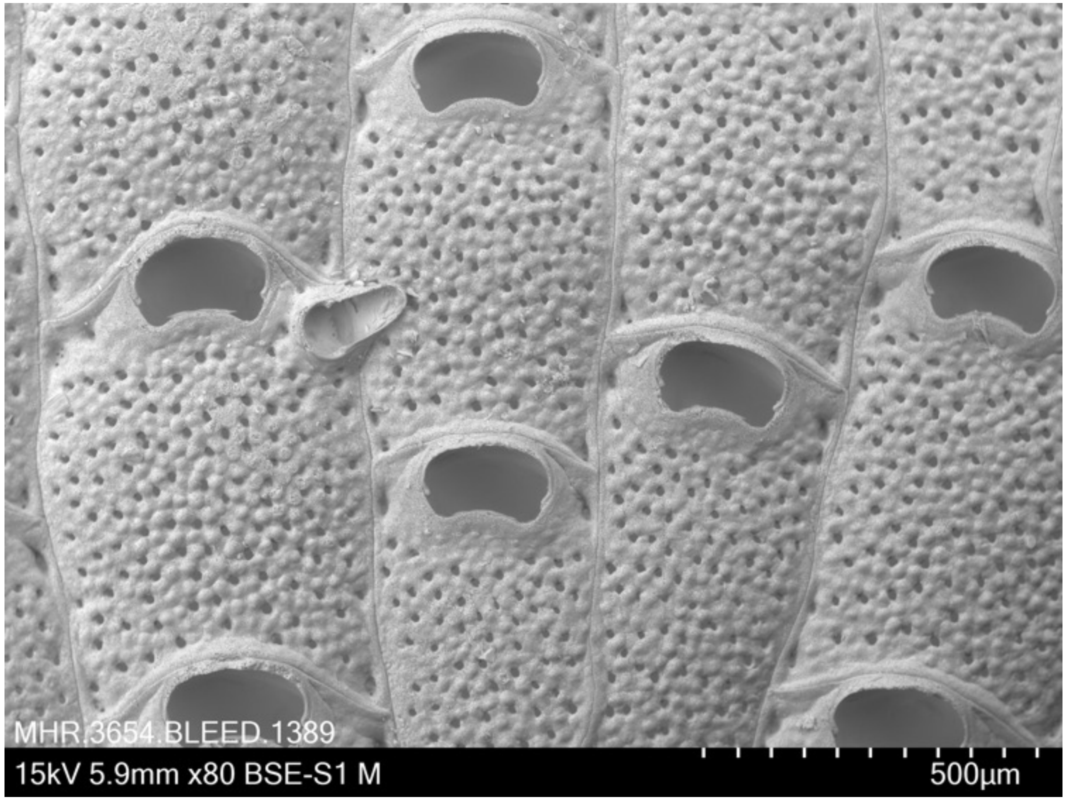


***Powellitheca terranovae* BLEED 1389A**
Coll: 17.3.2019 (A. M. Smith, D. P. Gordon)
Lat: -34.16
Long: 172.09
Depth: 82.9 m

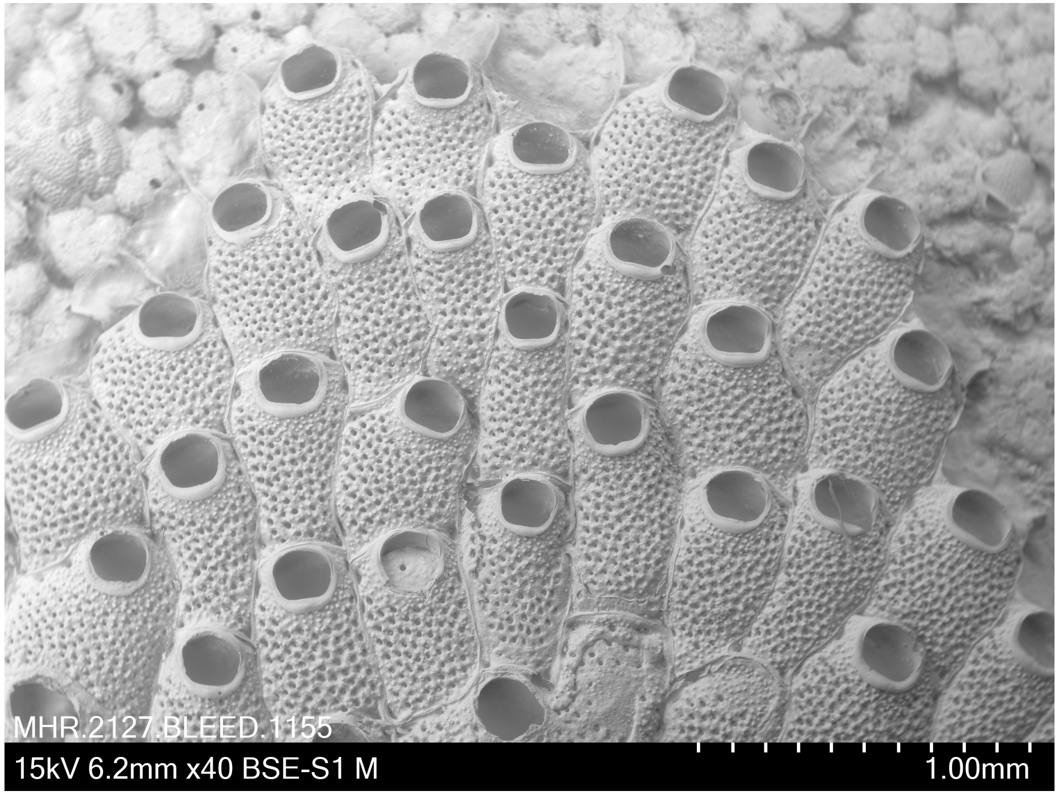


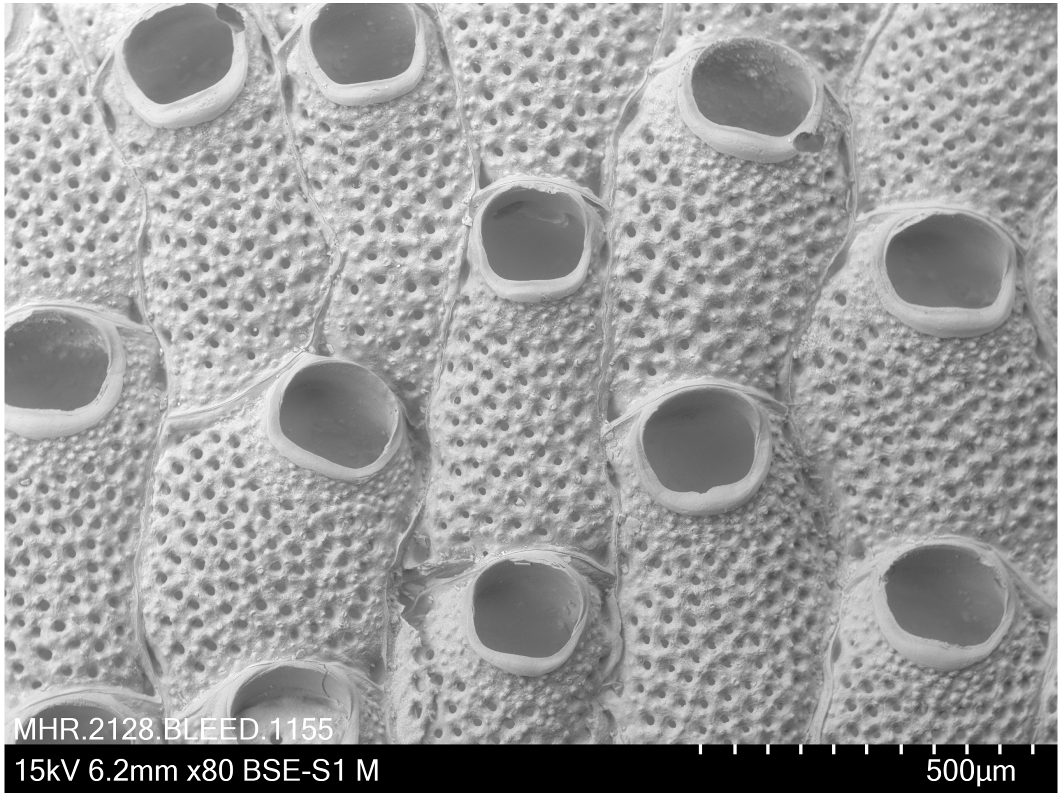


***Cyclicopora?* sp. BLEED 1155**
Coll: 10.11.2018 (A. M. Smith)
Lat: -45.822221999999996
Long: 170.880833
Depth: 80.4 m
**Notes:** This species is not well matched to the definition of *Cyclicopora* based on the type species of the genus *C. longipora* (MacGillivray, 1882). It may be more appropriate as a *Powellitheca* (frontal shield is regularly and evenly pseudoporous and orifice is convex proximally with small condyles) or *Saevitella*.

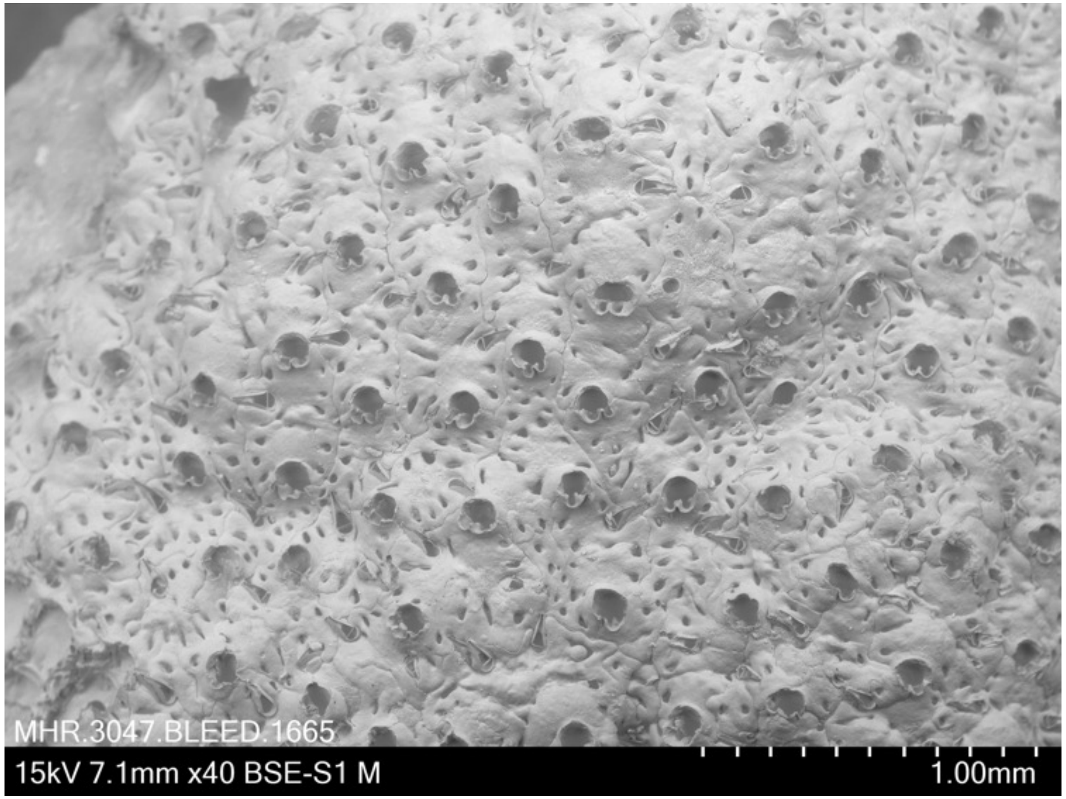


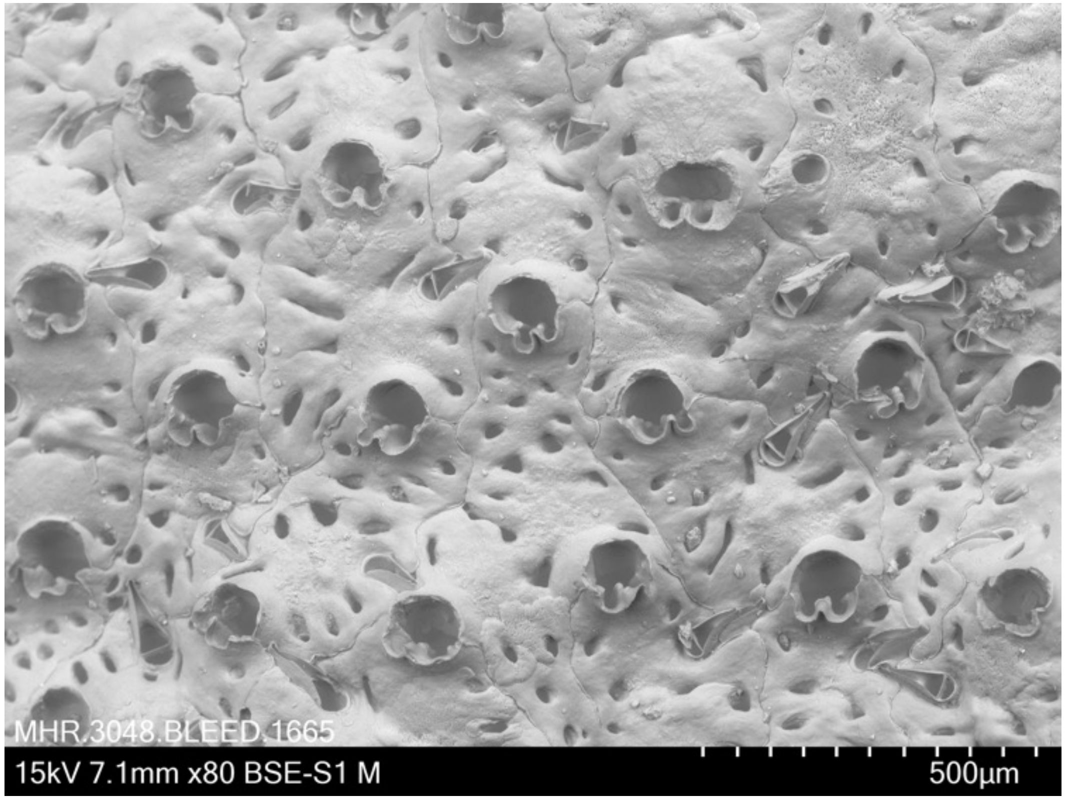


***Exochella* cf. *tricuspis* BLEED 1665**
Coll: 17.3.2019 (A. M. Smith, D. P. Gordon)
Lat: -34.21
Long: 172.11
Depth: 104 m

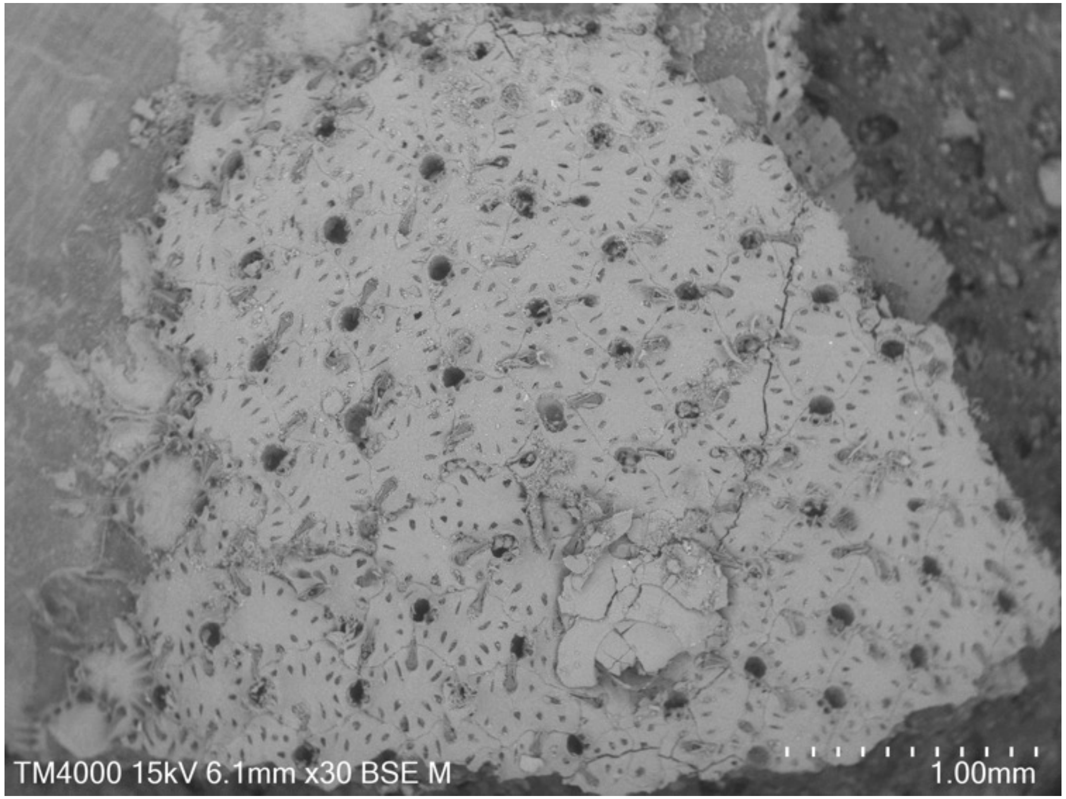


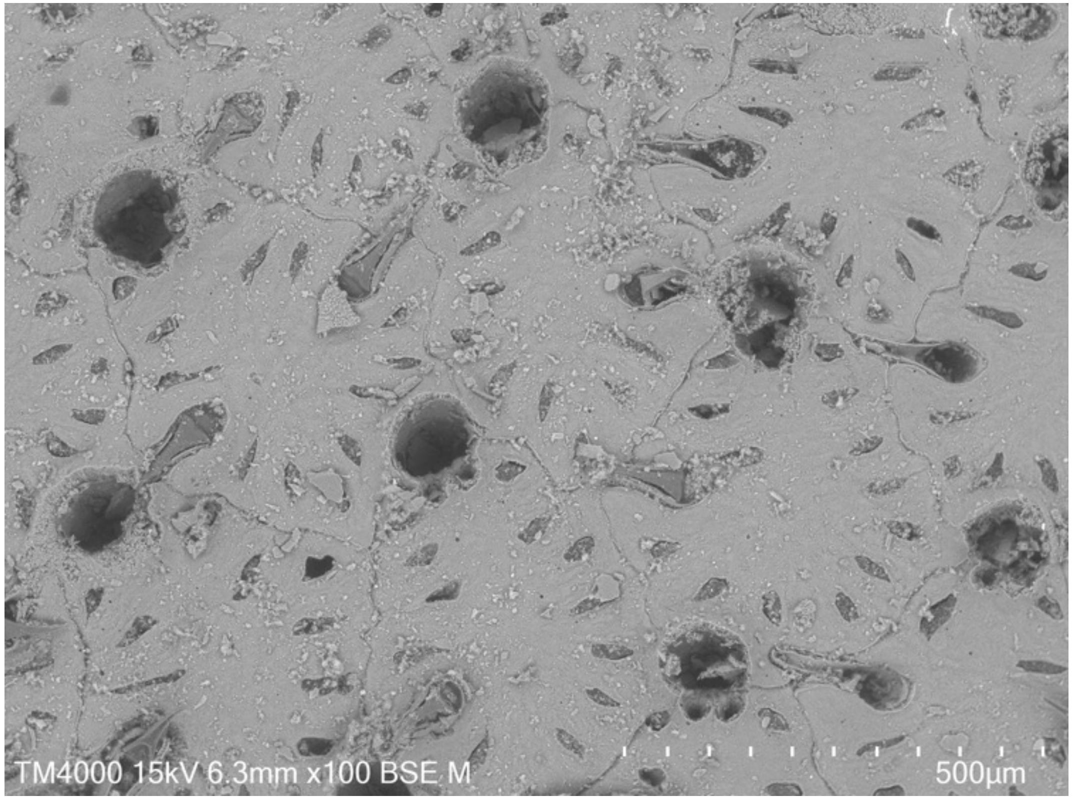


***Exochella jullieni* BLEED 231**
Coll: 1.2.2014 (A. M. Smith)
Lat: -45.6390
Long: 171.116
Depth: 250 m

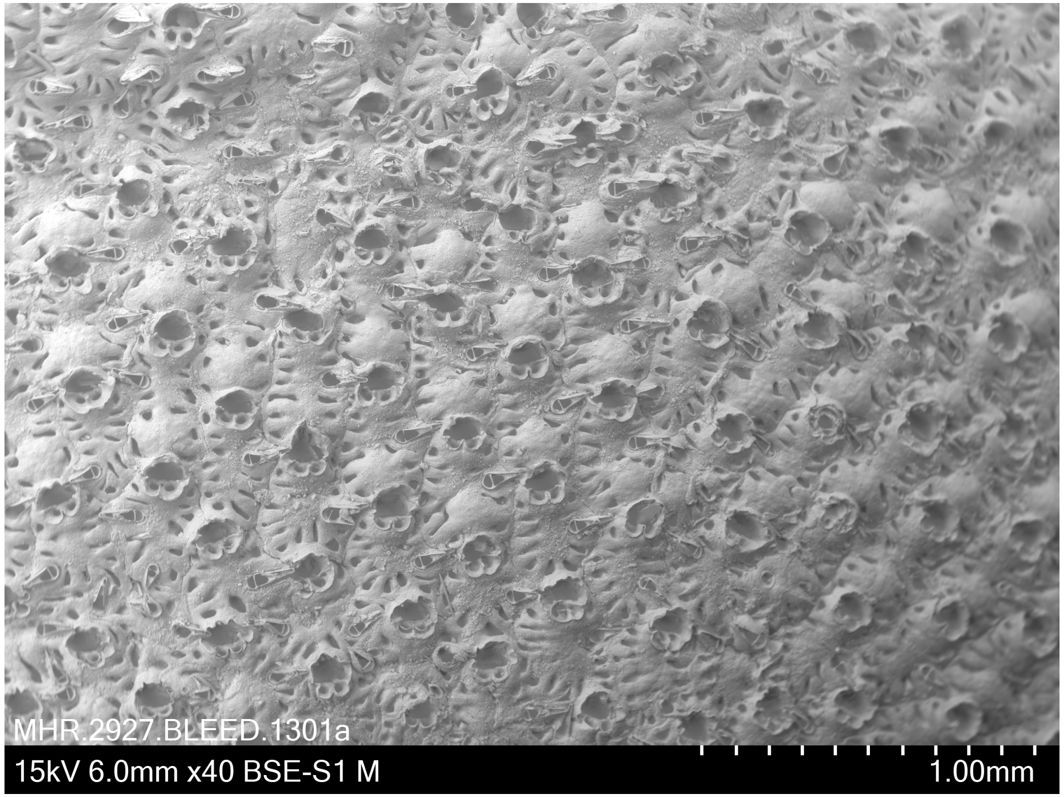


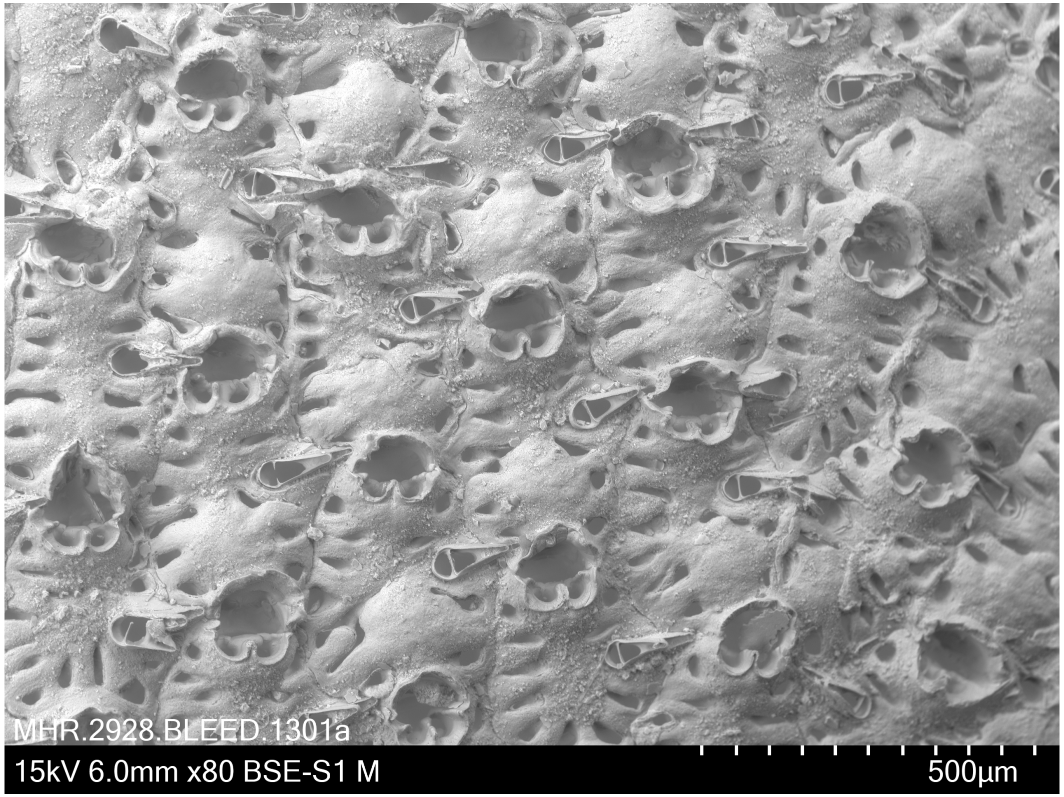


***Exochella* cf. *jullieni* BLEED 1301A**

Coll: 17.3.2019 (A. M. Smith, D. P. Gordon)

Lat: -34.21

Long: 172.11

Depth: 104 m

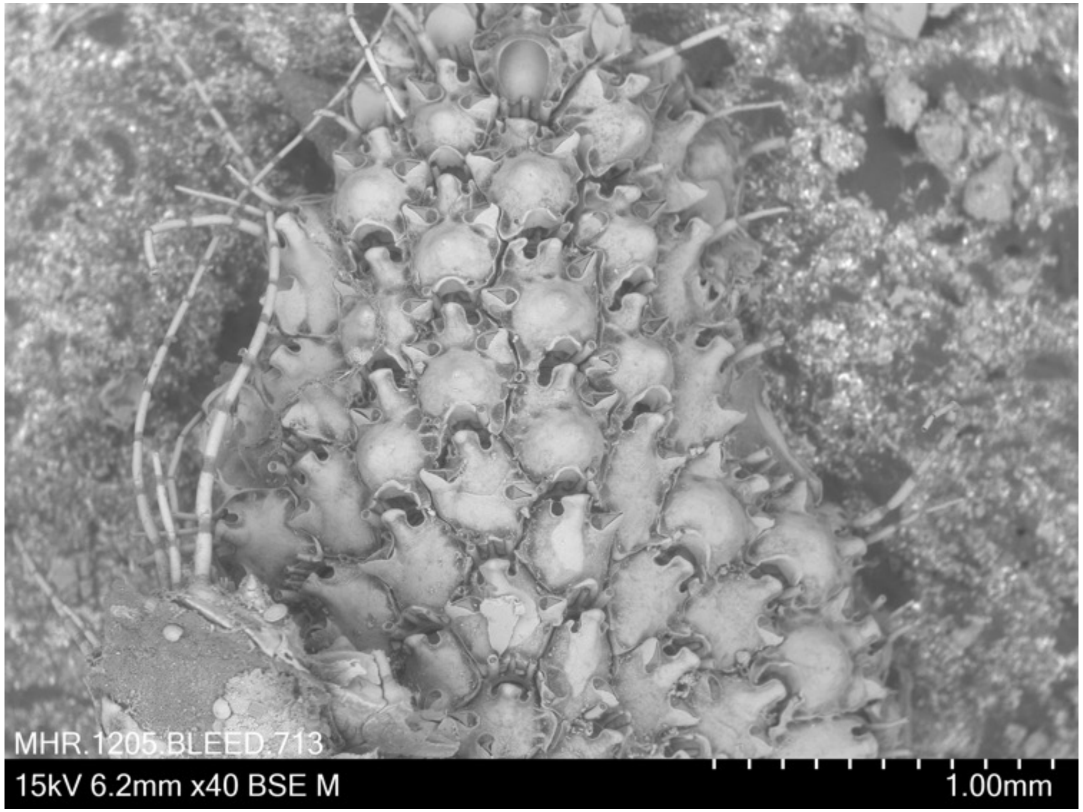


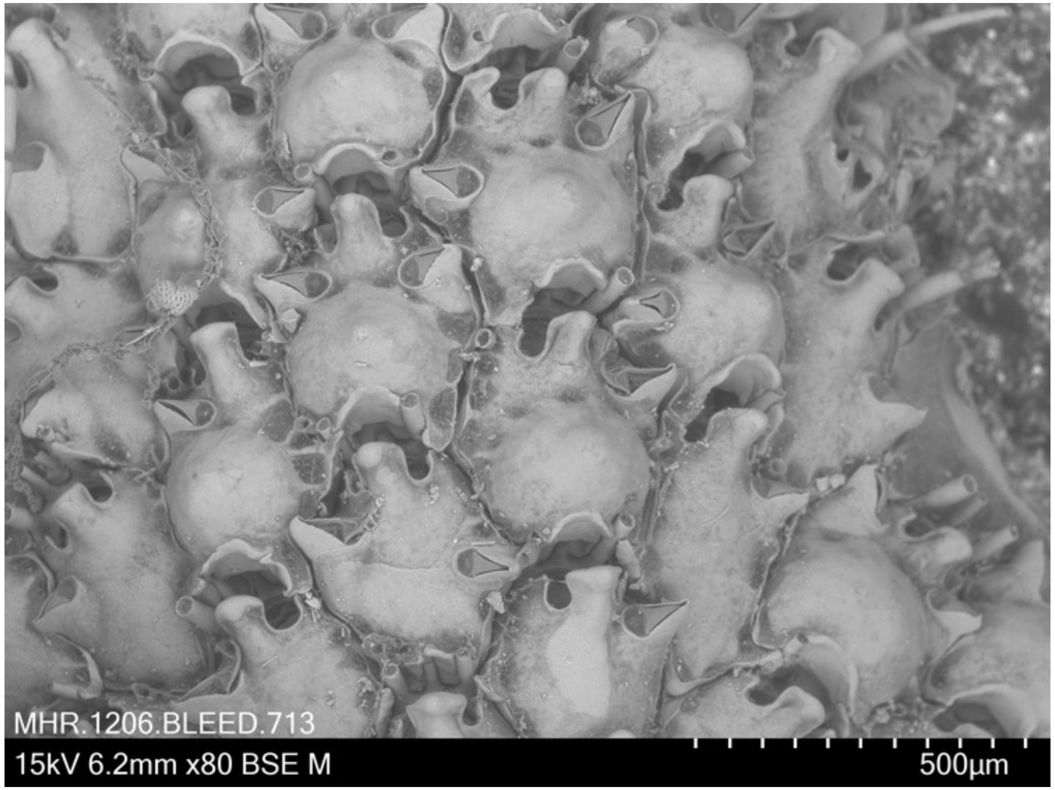


***Exochella armata* BLEED 713**
Coll: 8.4.2018 (A. M. Smith, D. P. Gordon, H. Mello)
Lat: -47.07306
Long: 18.2039
Depth: 18 m

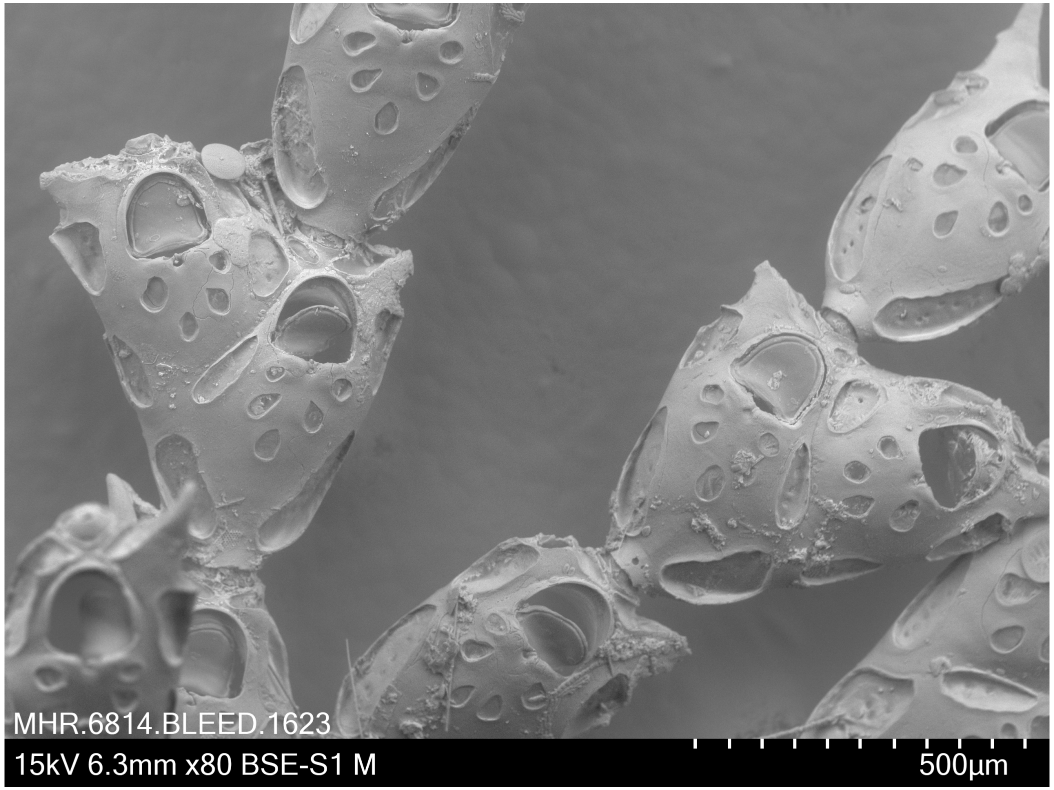


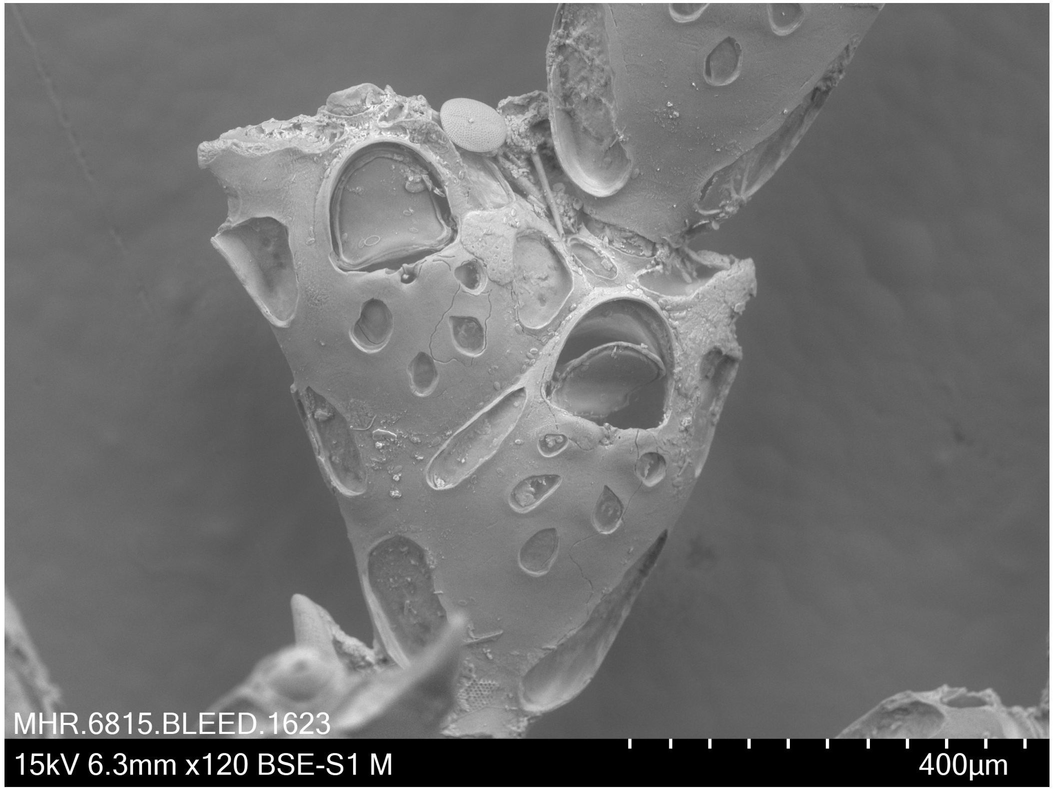


***Orthoscuticella fusiformis*** **BLEED 1623**
Coll: 17.3.2019 (A. M. Smith, D. P. Gordon)
Lat: -34.16
Long: 172.09
Depth: 82.9 m

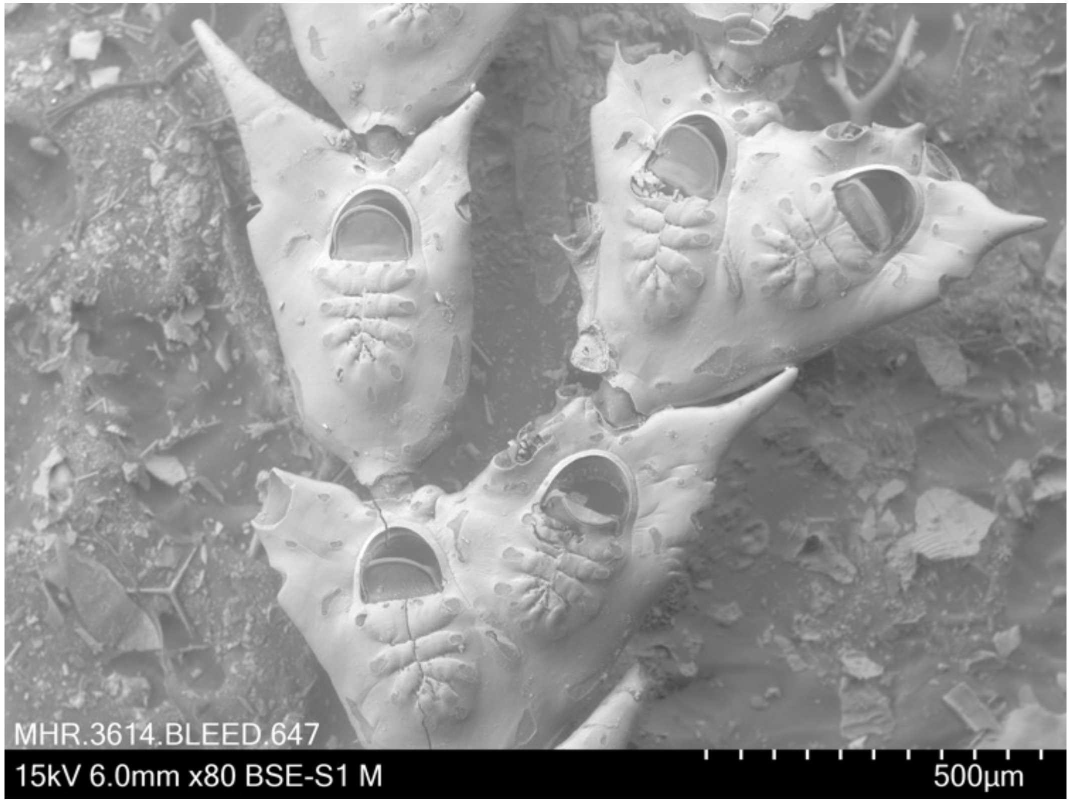


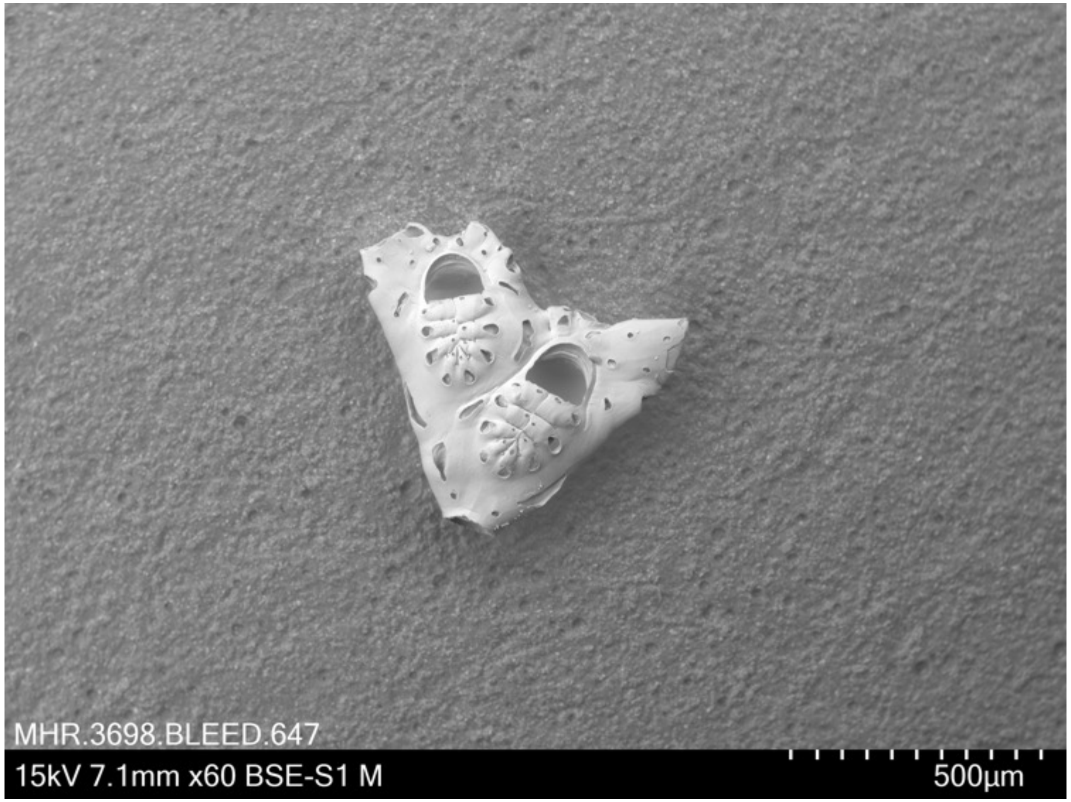


***Costaticella bicuspis* BLEED 647**
Coll: 8.4.2018 (A. M. Smith, D. P. Gordon, H. Mello)
Lat: -47.1736
Long: 167.9558
Depth: 66 m

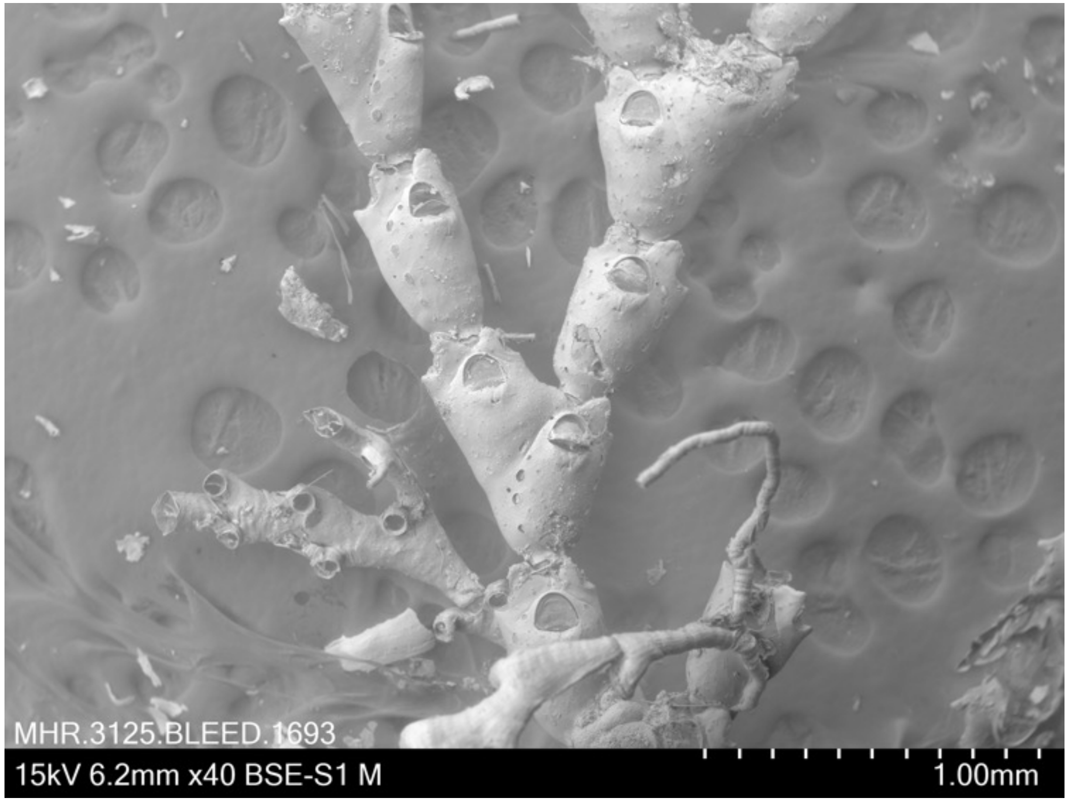


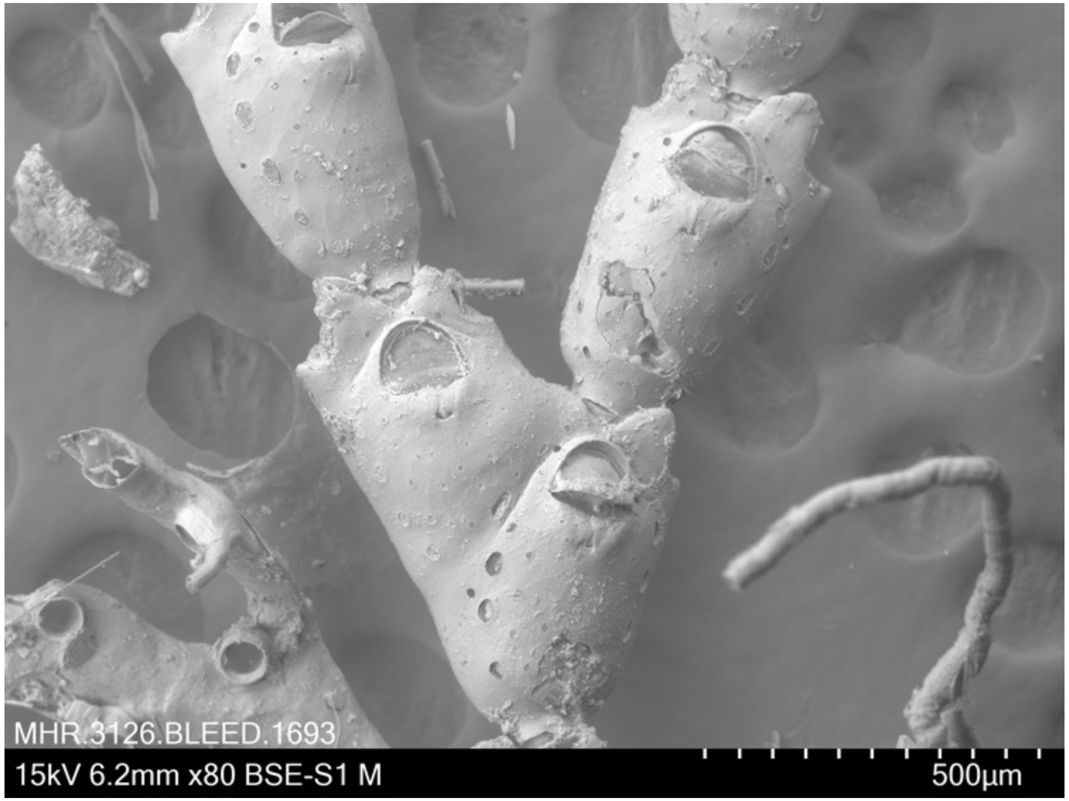


***Paracribricellina cribraria* BLEED 1693**
Coll: 17.3.2019 (A. M. Smith, D. P. Gordon)
Lat: -34.21
Long: 172.11
Depth: 104 m

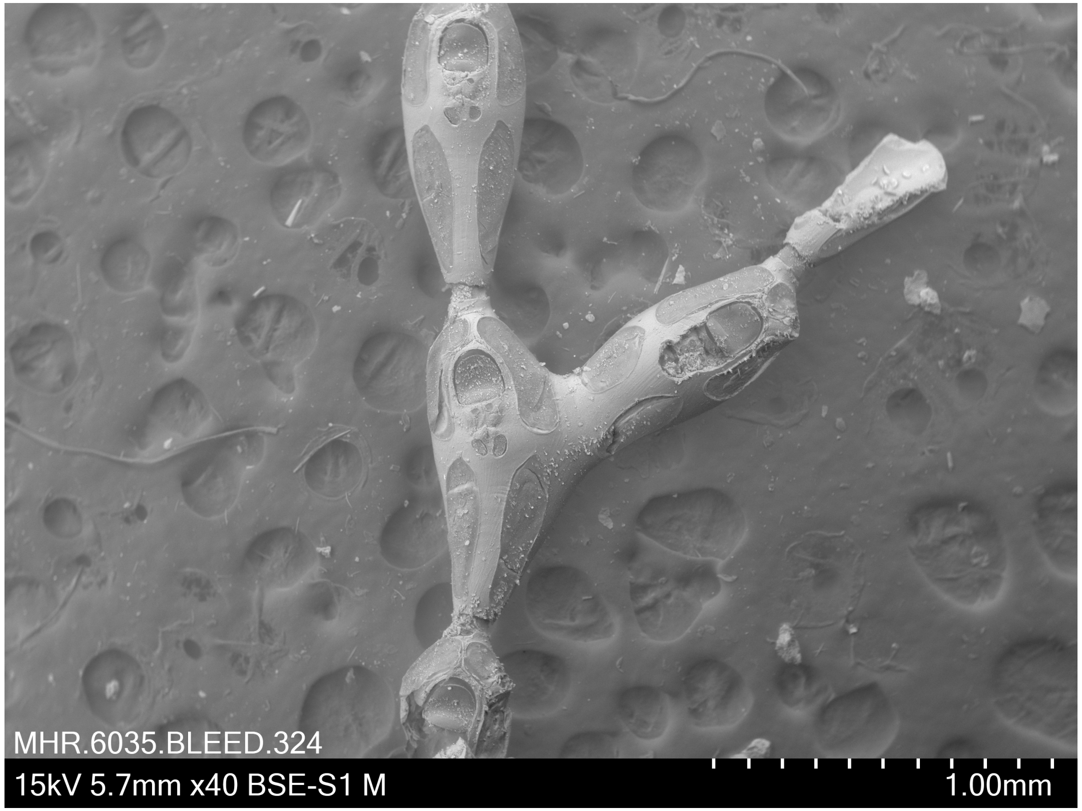


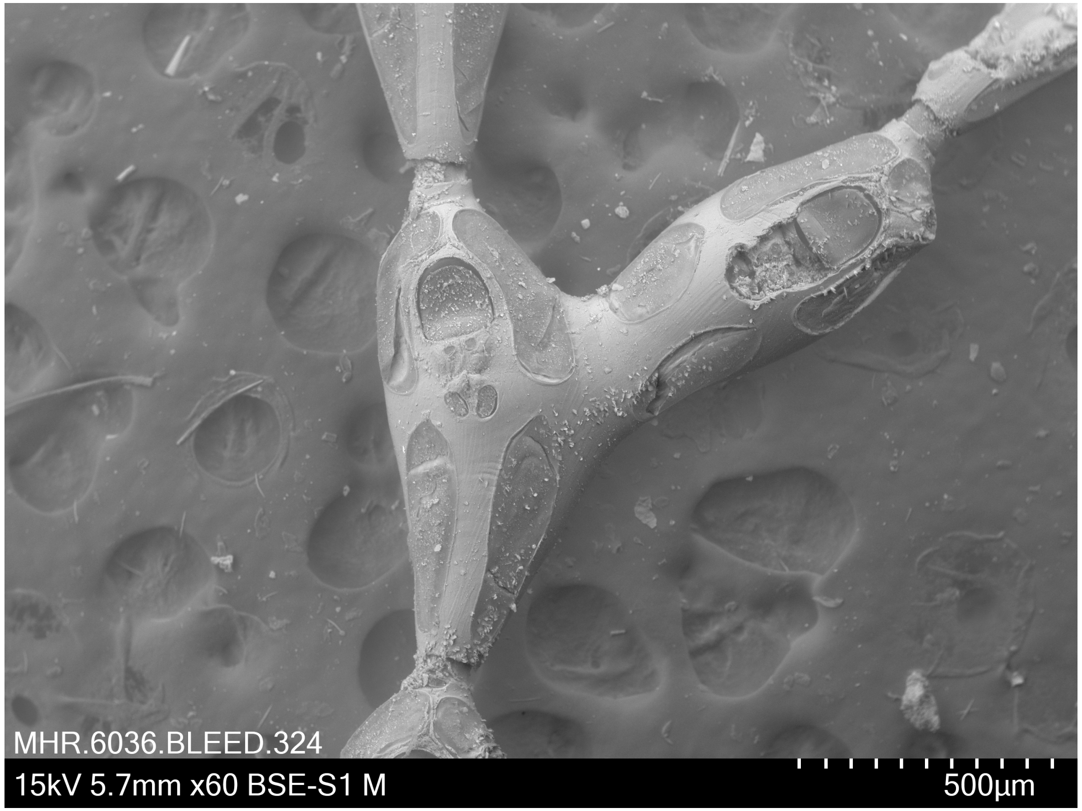


***Talivittaticella problematica* BLEED 324**
NIWA 39687 (TAN 0803/31)
Coll: 1.4.2008
Lat: -50.08767
Long: 163.492
Depth: 1163 m

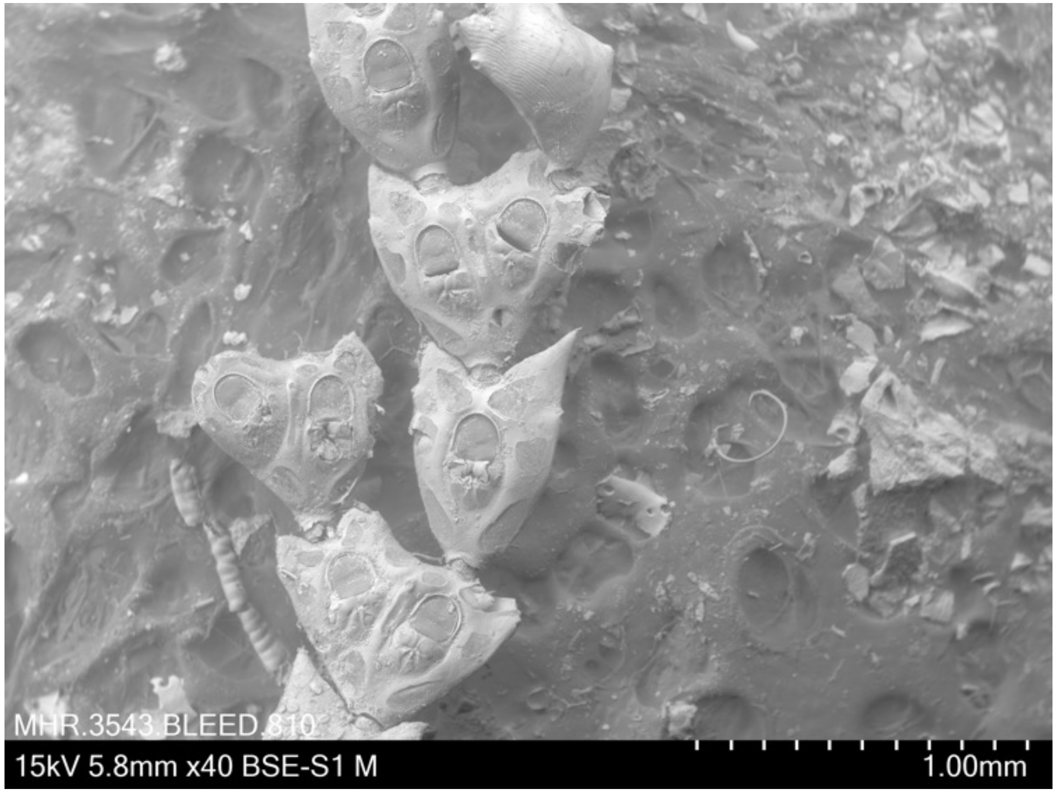


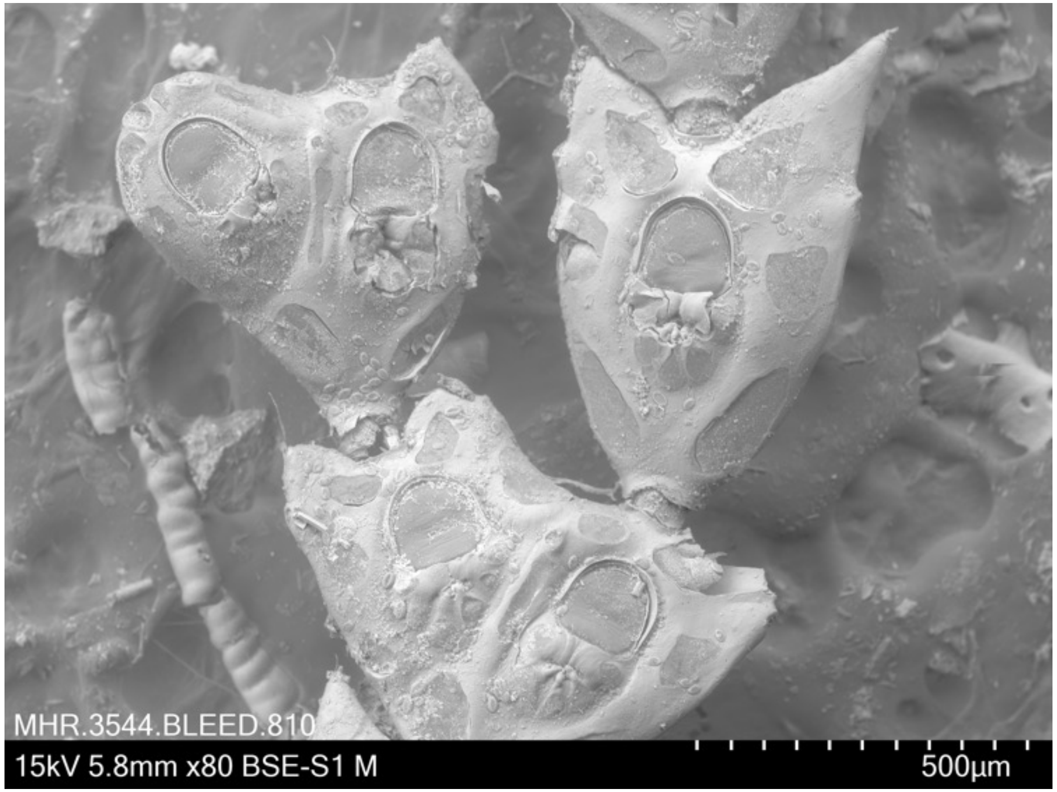


***Pterocella vesiculosa* BLEED 810**
Coll: 3.4.2018 (D. P. Gordon, L. H. Liow and C. Schack)
Lat: -39.639352
Long: 177.008705
Depth: intertidal

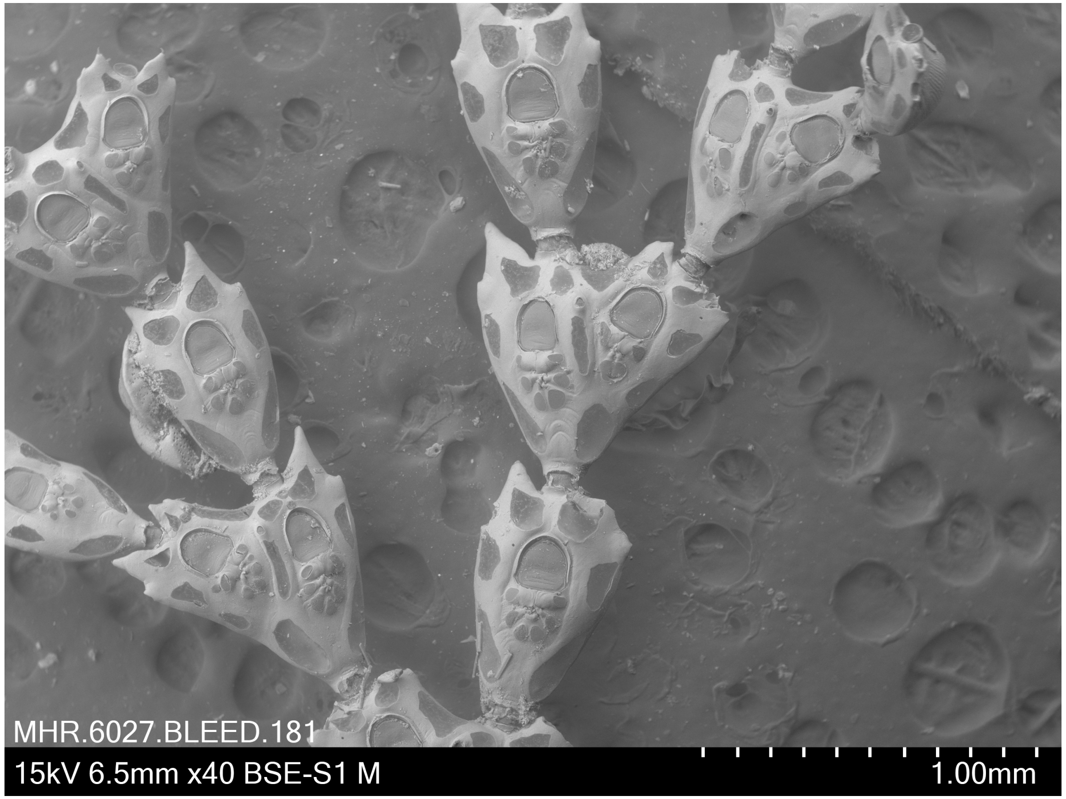


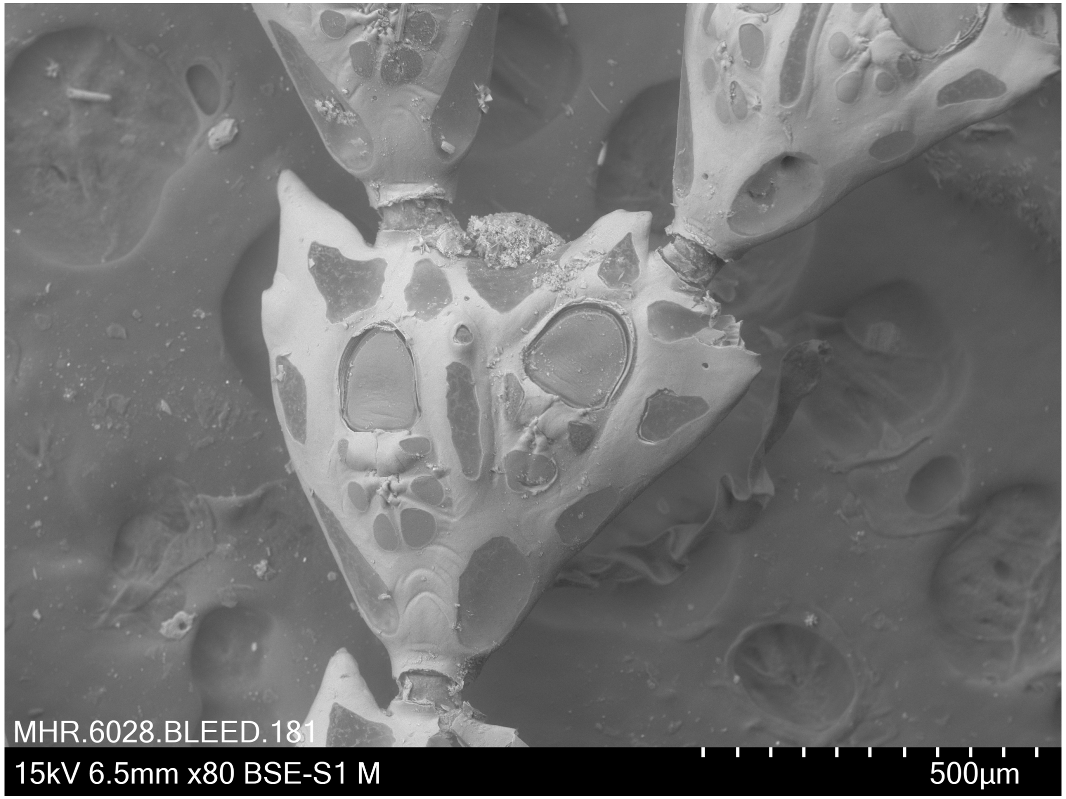


***Pterocella vesiculosa*** **BLEED 181**
NIWA 56624 (TAN0906/159)
Coll: 14.7. 2009
Lat: -34.3875999
Long: 173.041
Depth: 59 m

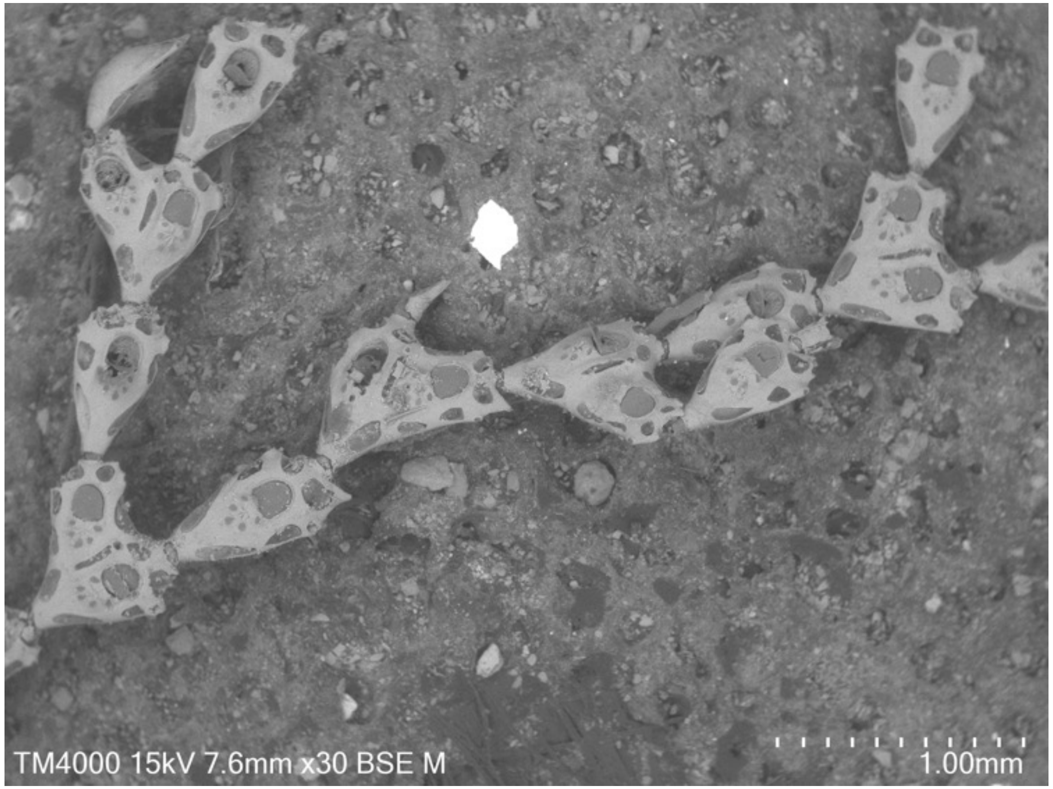


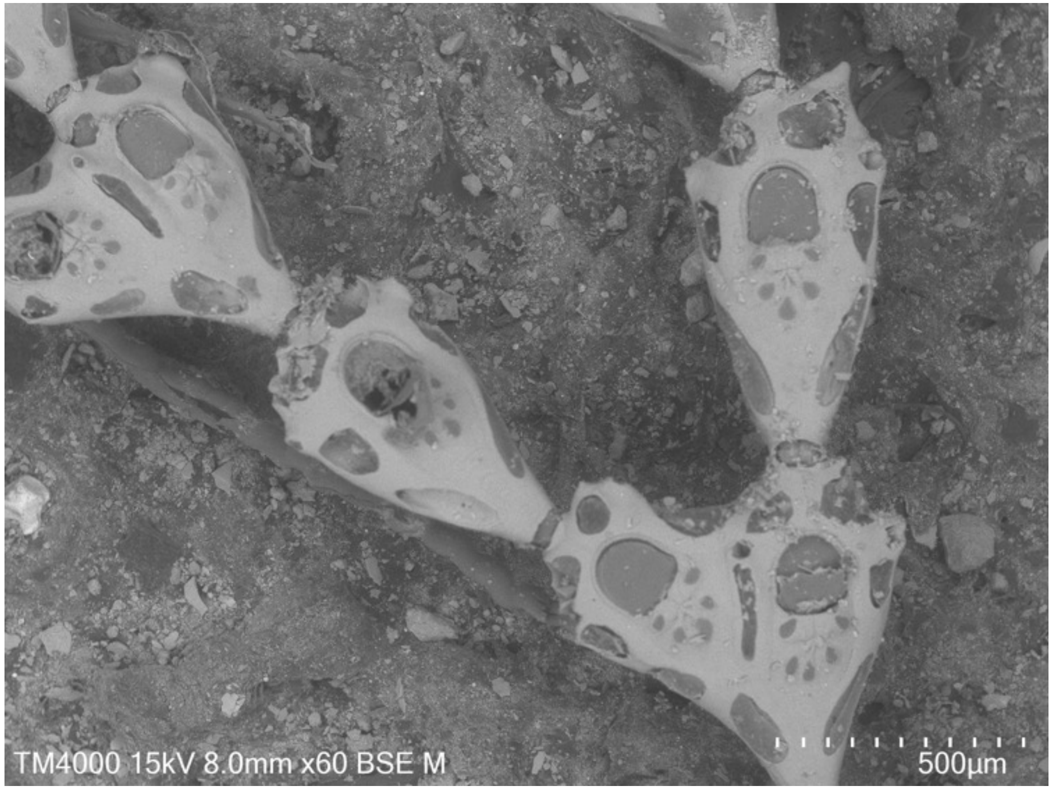


***Pterocella vesiculosa* BLEED 105**
NIWA 73295 **(**TAN1105/43)
Coll: 28.3.2011
Lat: -33.9875
Long: 171.7508333333
Depth: 170-174 m

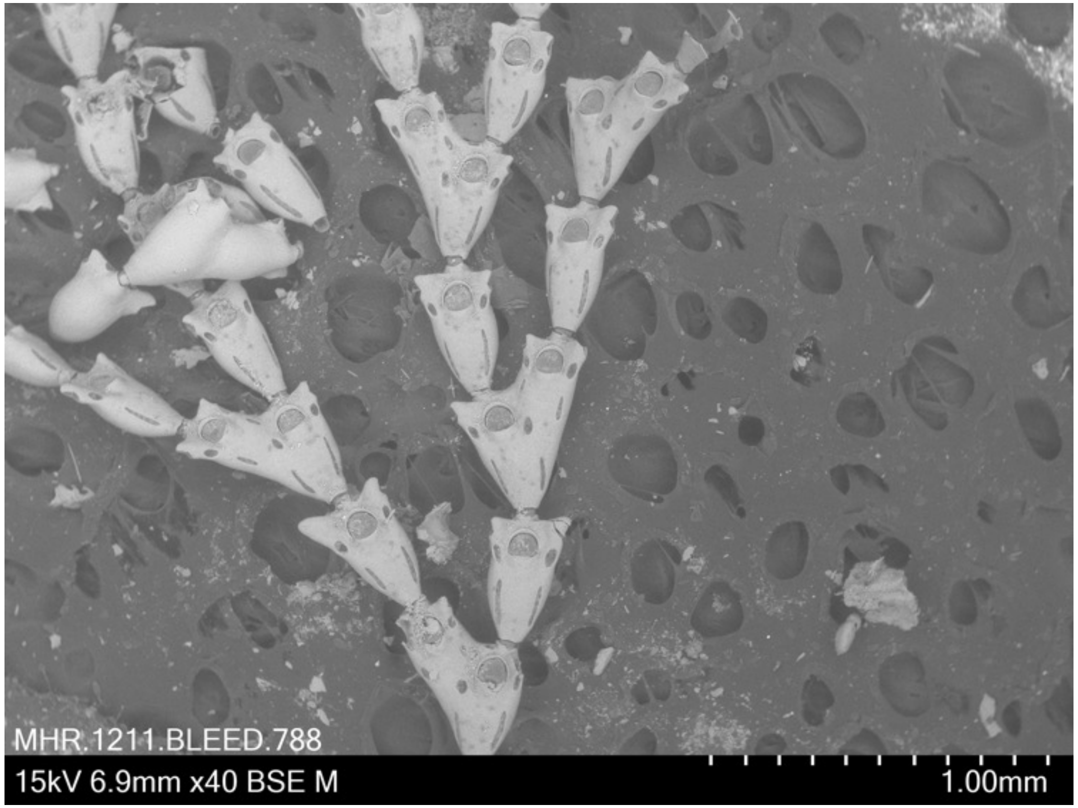


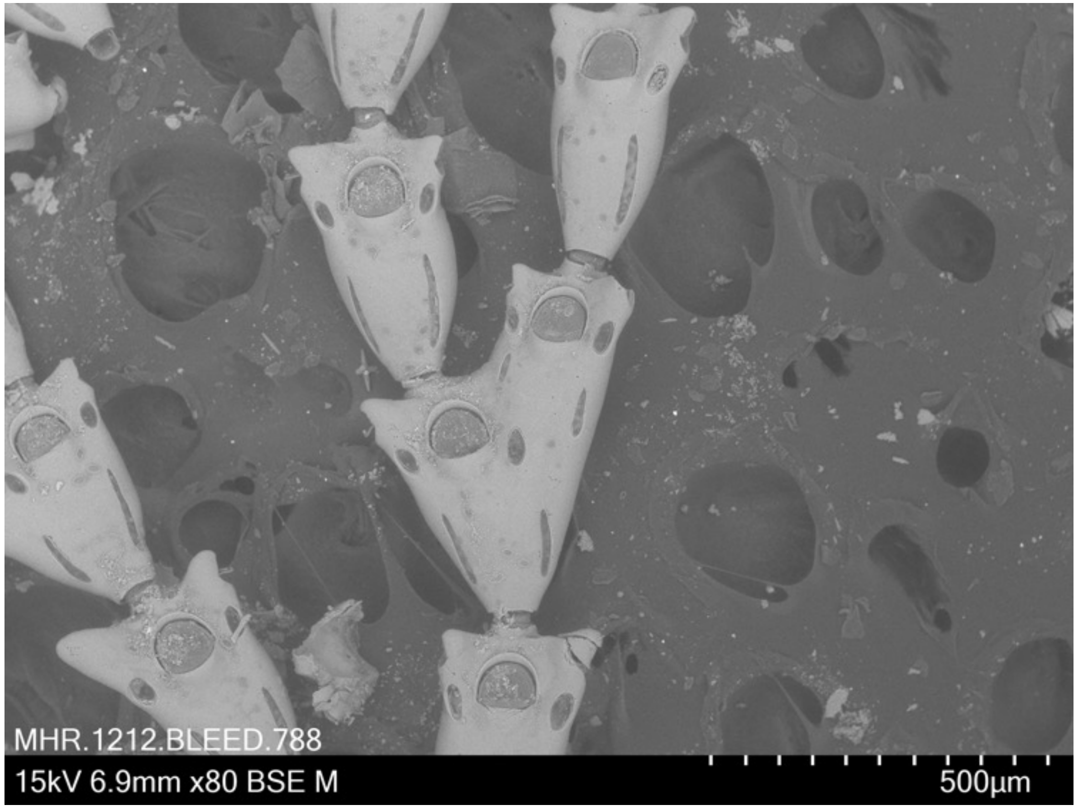


***Catenicella pseudoelegans* BLEED 788**
Coll: 4.3.2018 (D. P. Gordon, L. H. Liow and C. Schack)
Lat: -39.639352
Long: 177.008705
Depth: intertidal

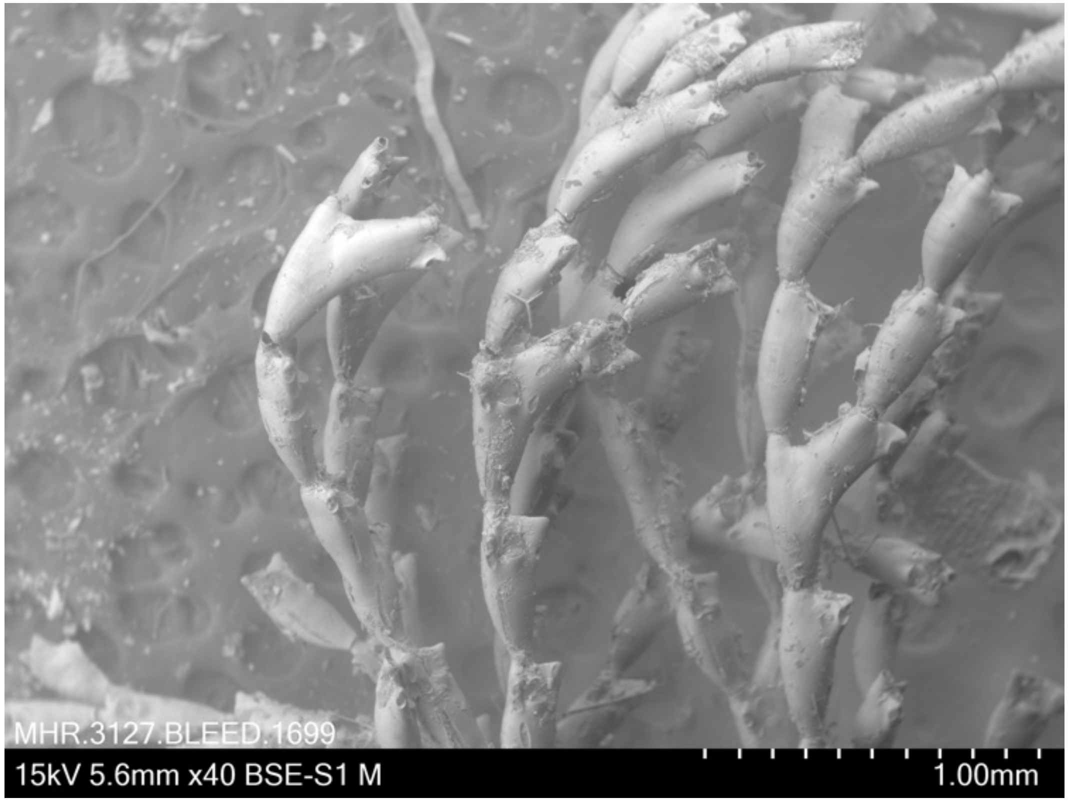


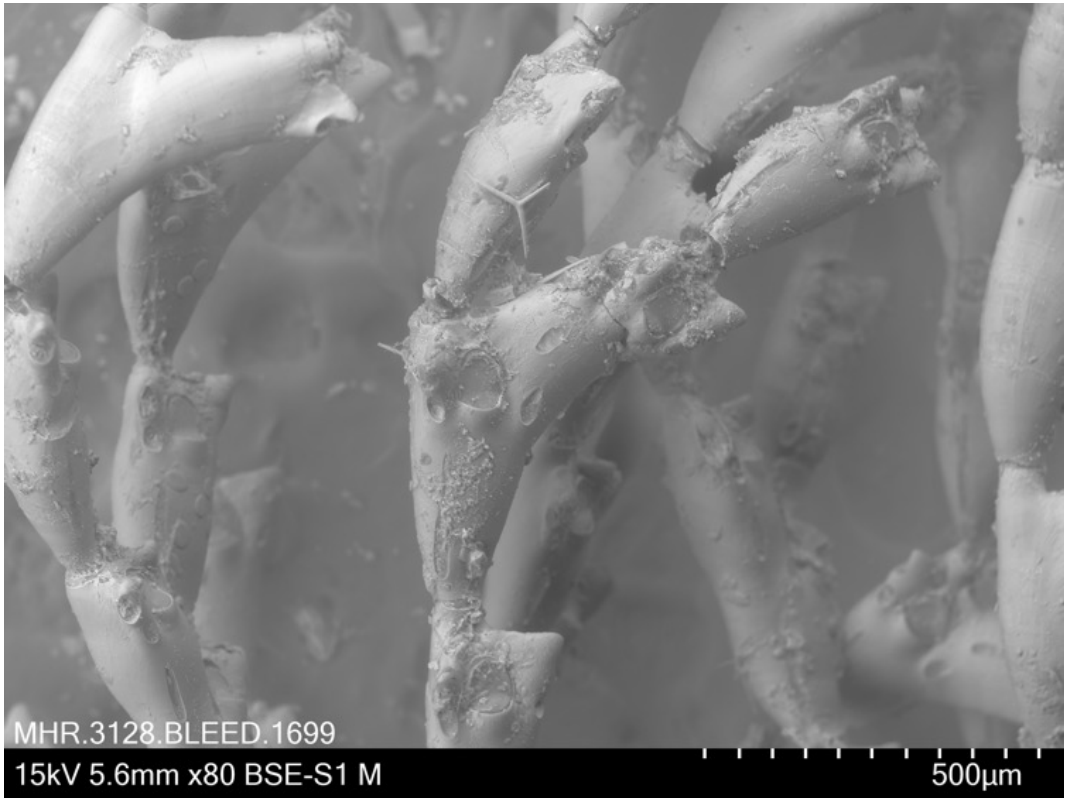


***Catenicella* sp. BLEED 1699**
Coll: 17.3.2019 (A. M. Smith, D. P. Gordon)
Lat: -34.16
Long: 172.07
Depth : 98.9 m

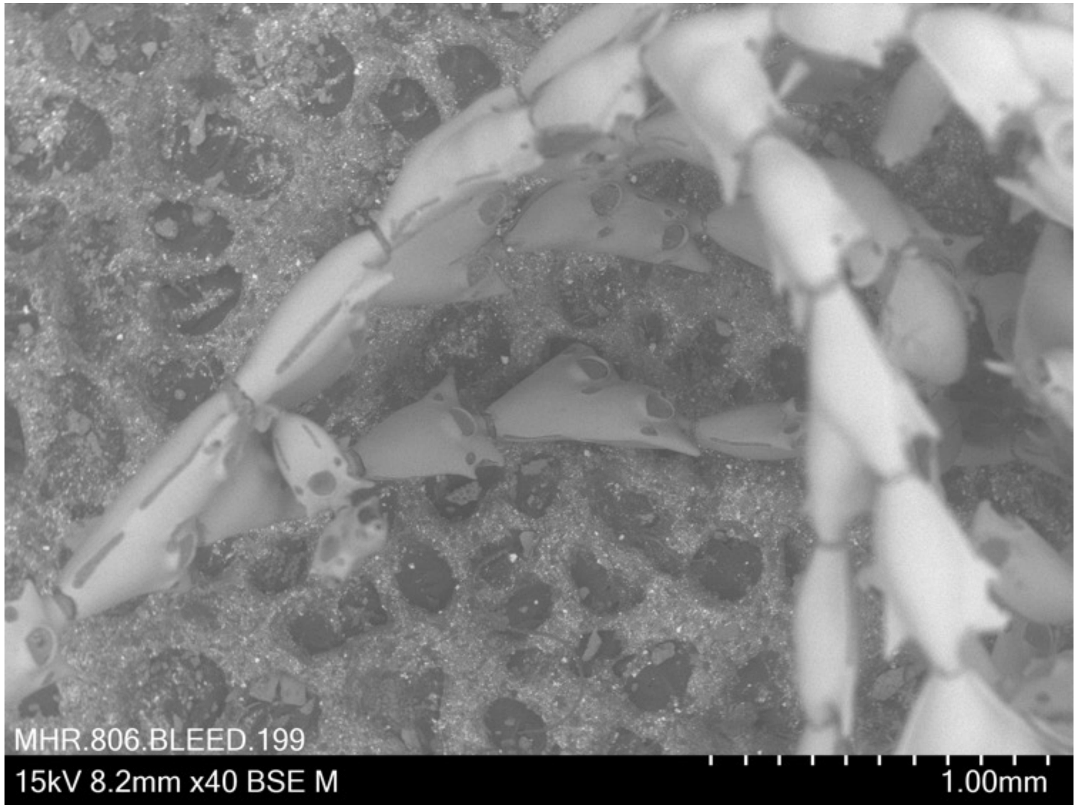


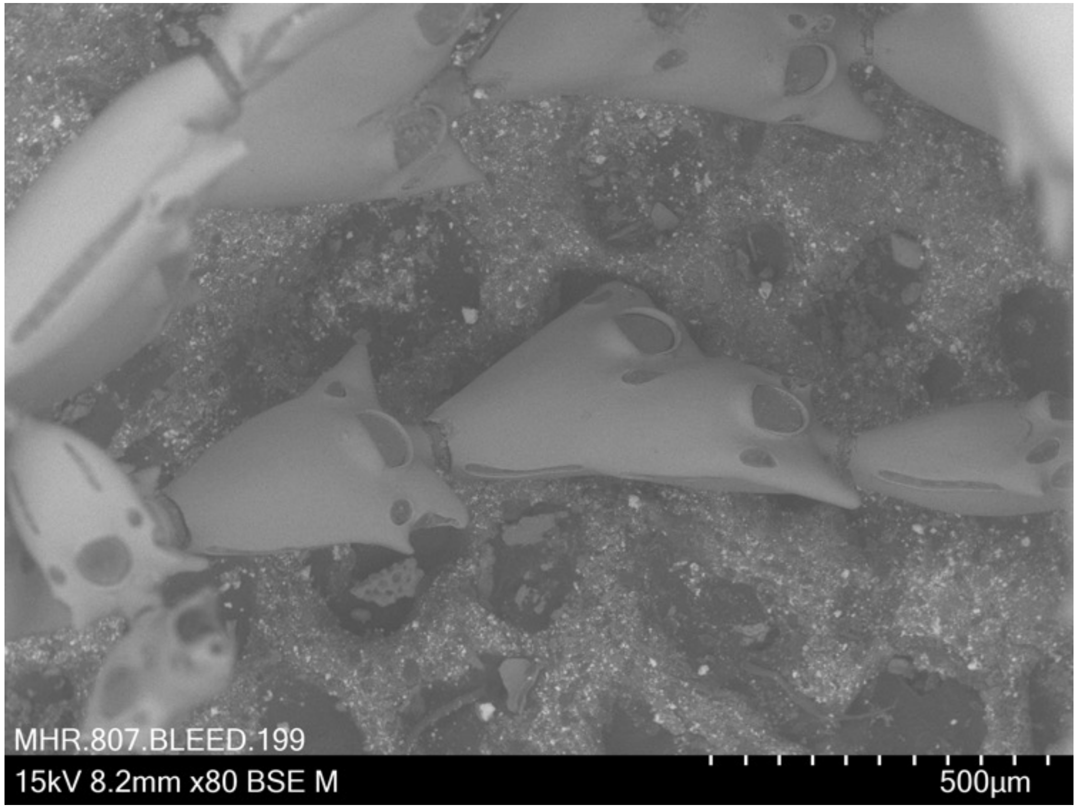


***Cornuticella taurina* BLEED 199**
Coll: 20.5.2015 (D. P. Gordon)
Lat: -41.071039
Long: 176.08894
Depth: Intertidal

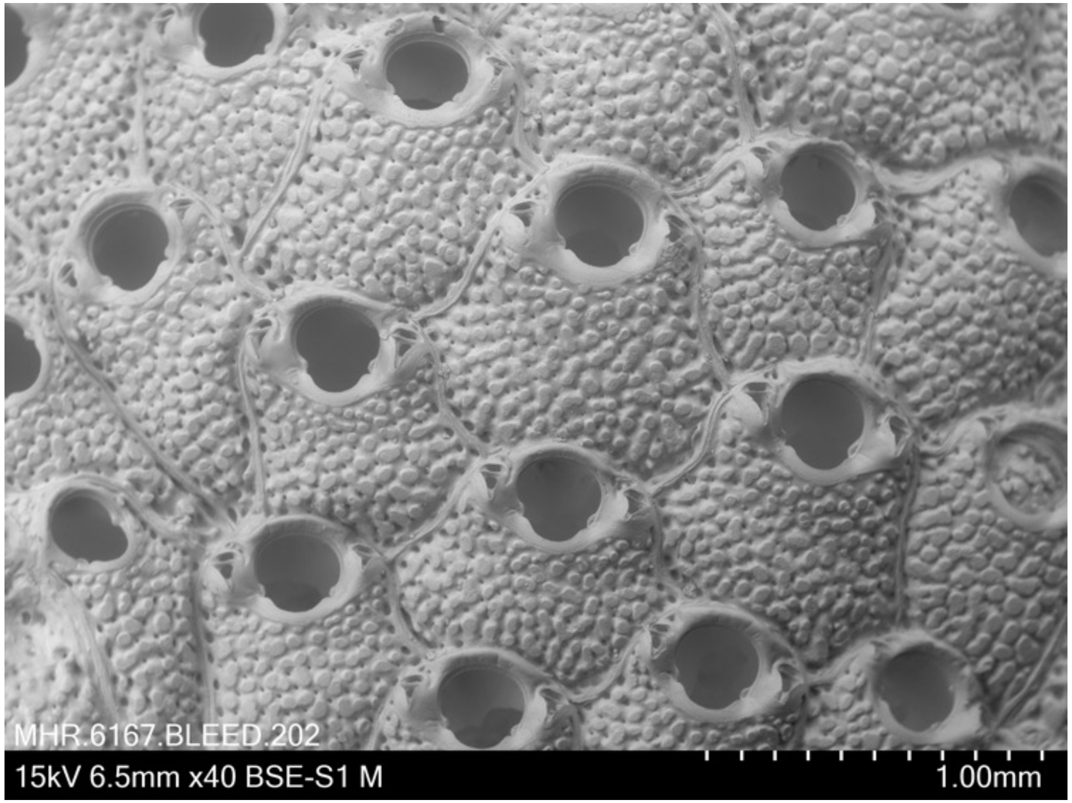


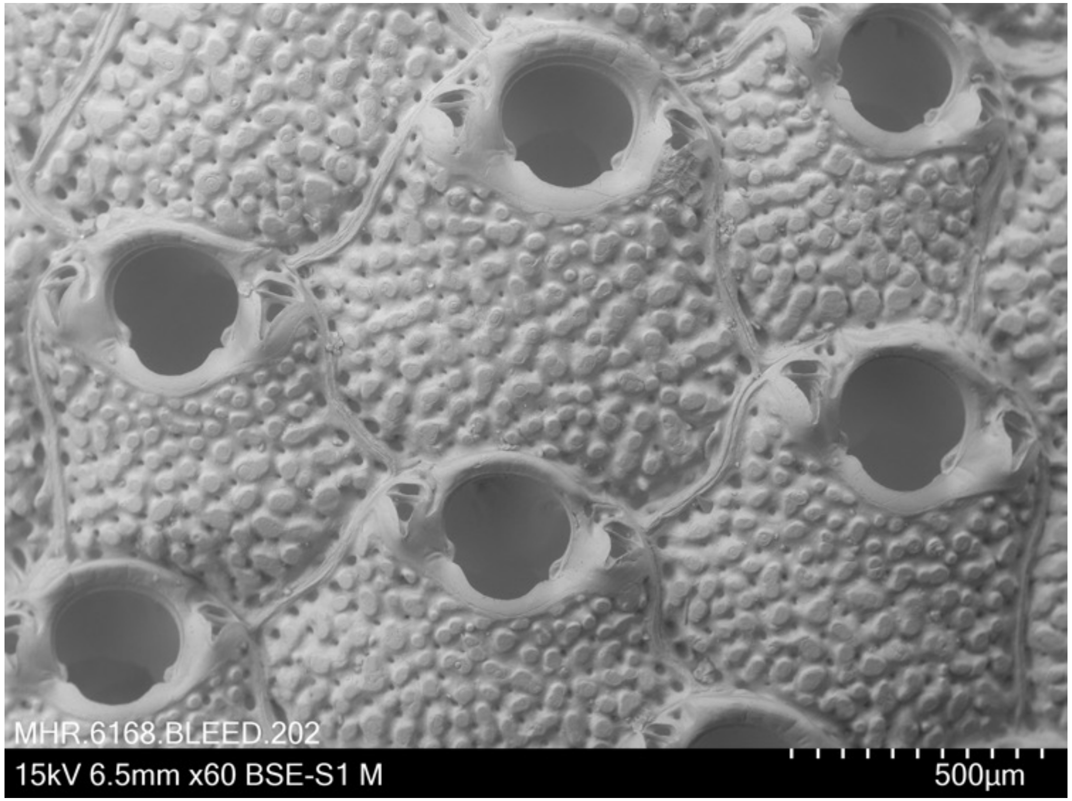


***Cosciniopsis vallata* BLEED 202**
Date: 20.5.2015 (D. P. Gordon)
Lat: -41.071039
Long: 176.088894
Depth: Intertidal

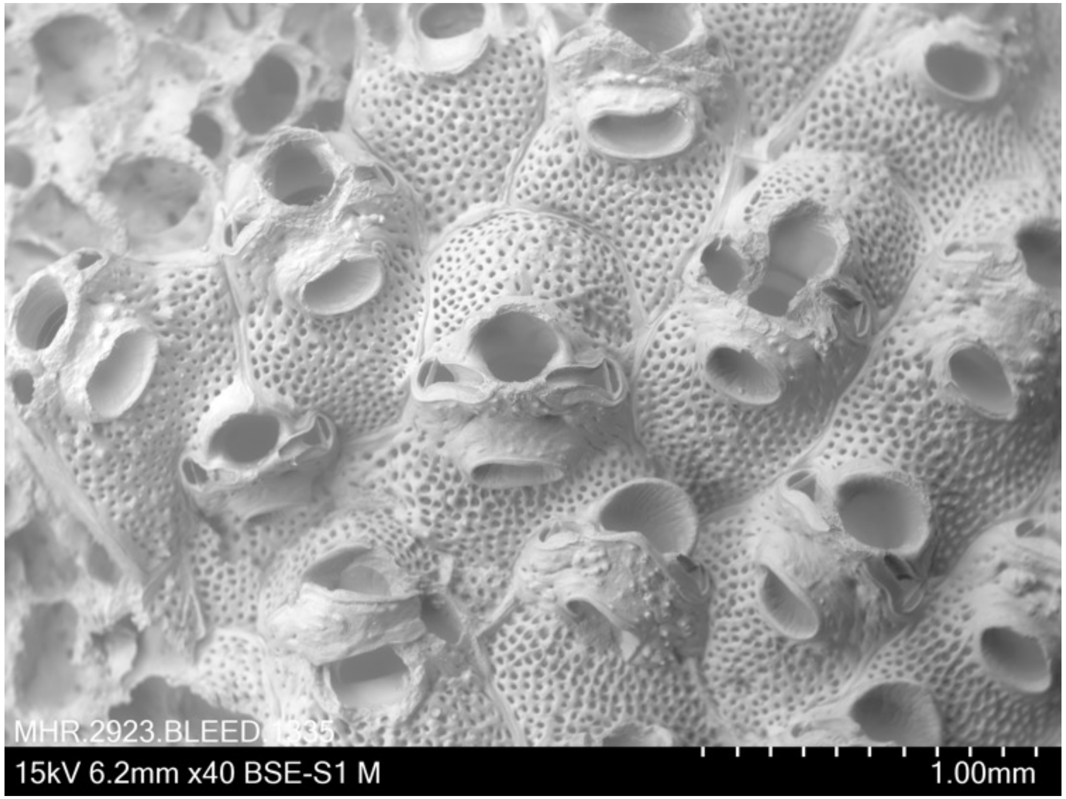


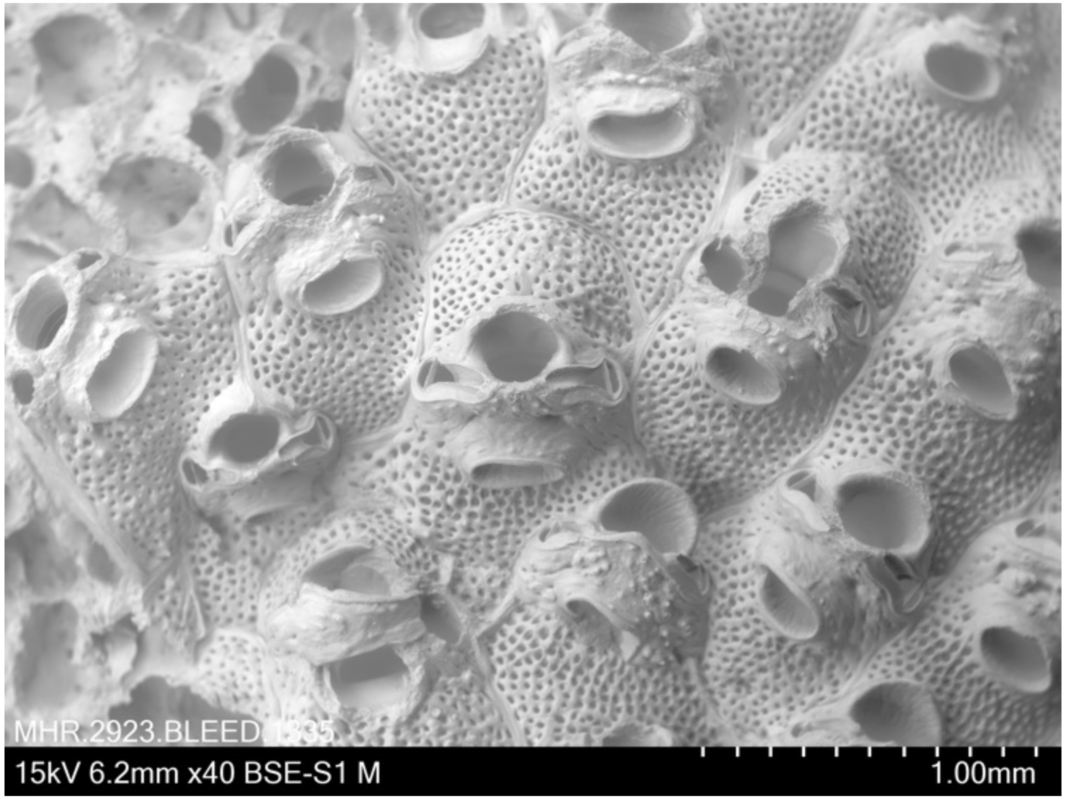


***Gigantopora pupa* BLEED 1335**
Coll: 19.3.2019 (A. M. Smith, D. P. Gordon)
Lat: -34.99
Long: 173.97
Depth: 61.6 m

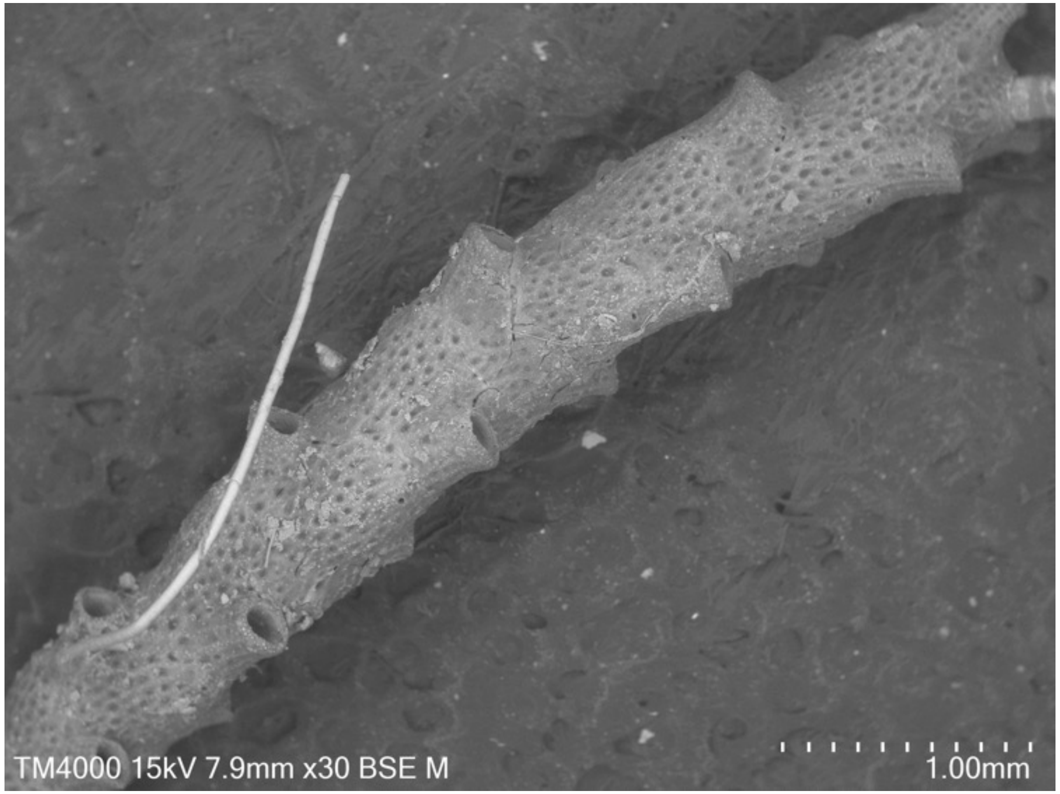


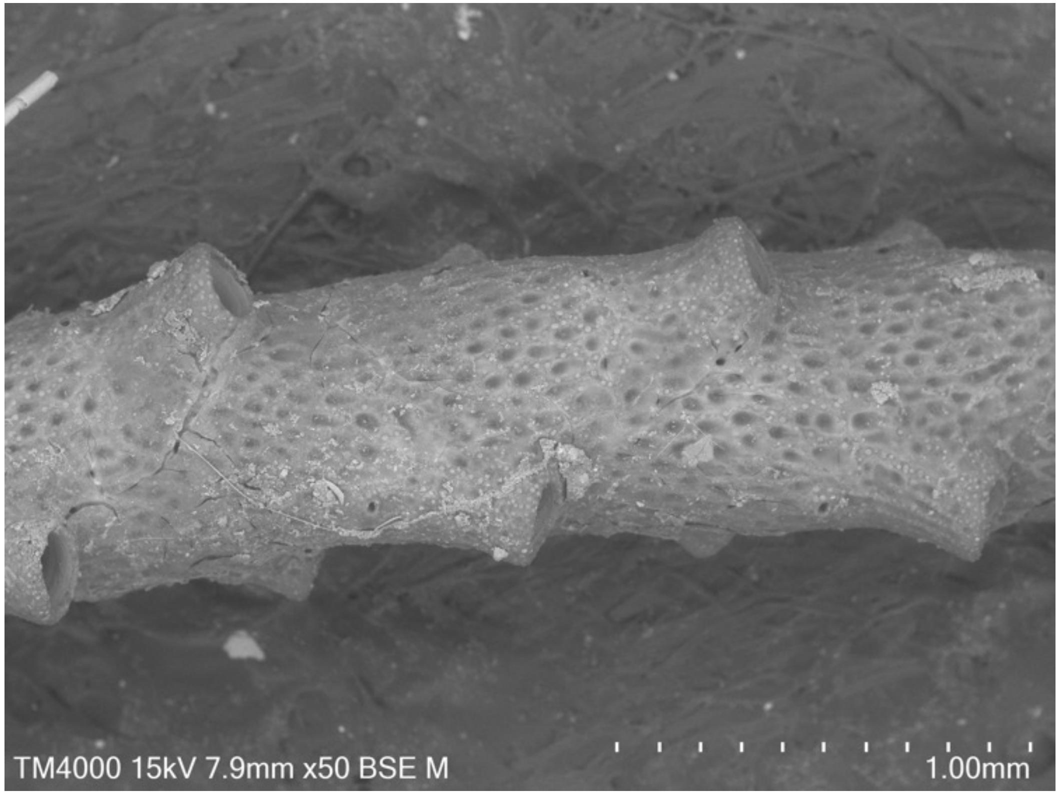


***Margaretta barbata* BLEED 318**
NIWA 56528 (TAN0906/154)
Coll: 14.7.2009
Lat: -34.373833333299999
Long: 172.9231667
Depth: 72 m

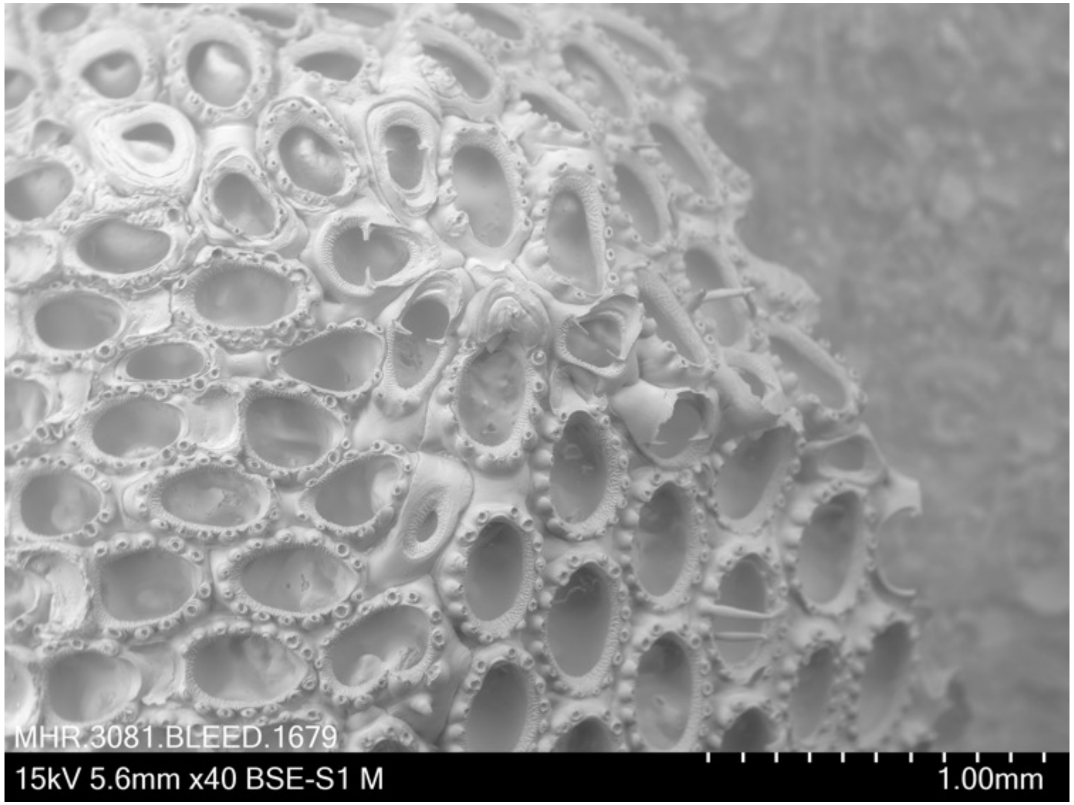


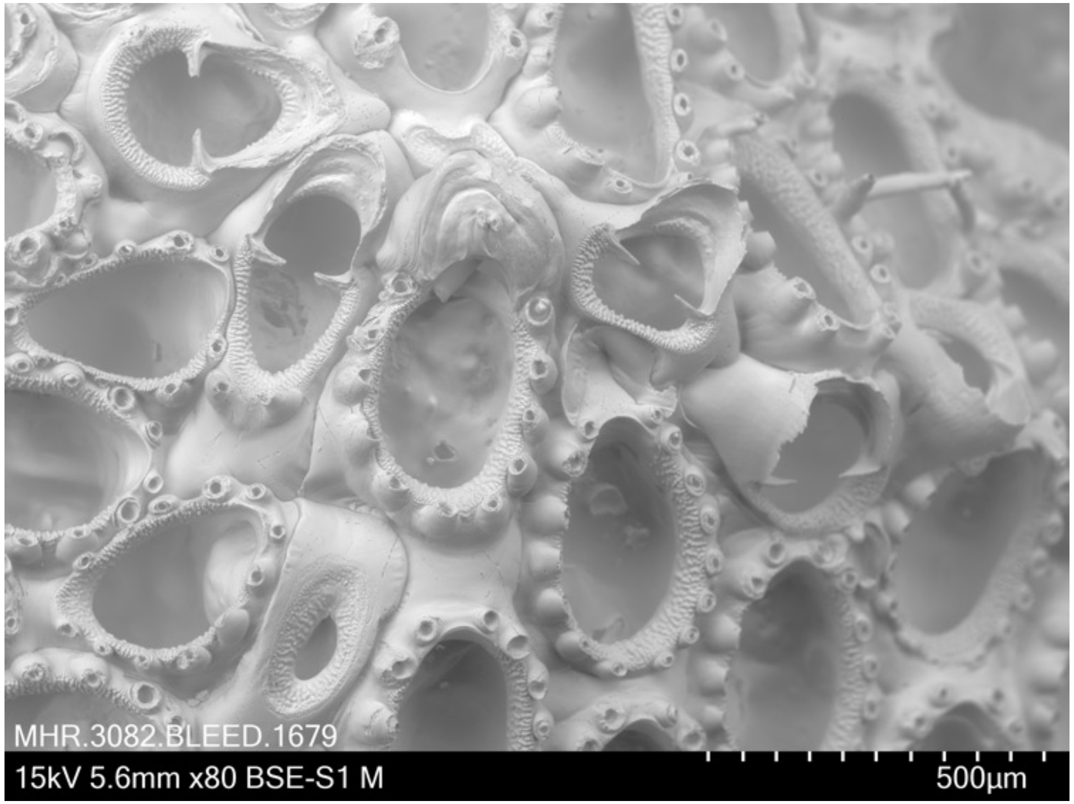


***Valdemunitella* n. sp. BLEED 1679**
Coll: 18.3.2019 (A. M. Smith, D. P. Gordon)
Lat: -34.380000000000003
Long: 172.69
Depth: 52.5-56.5m

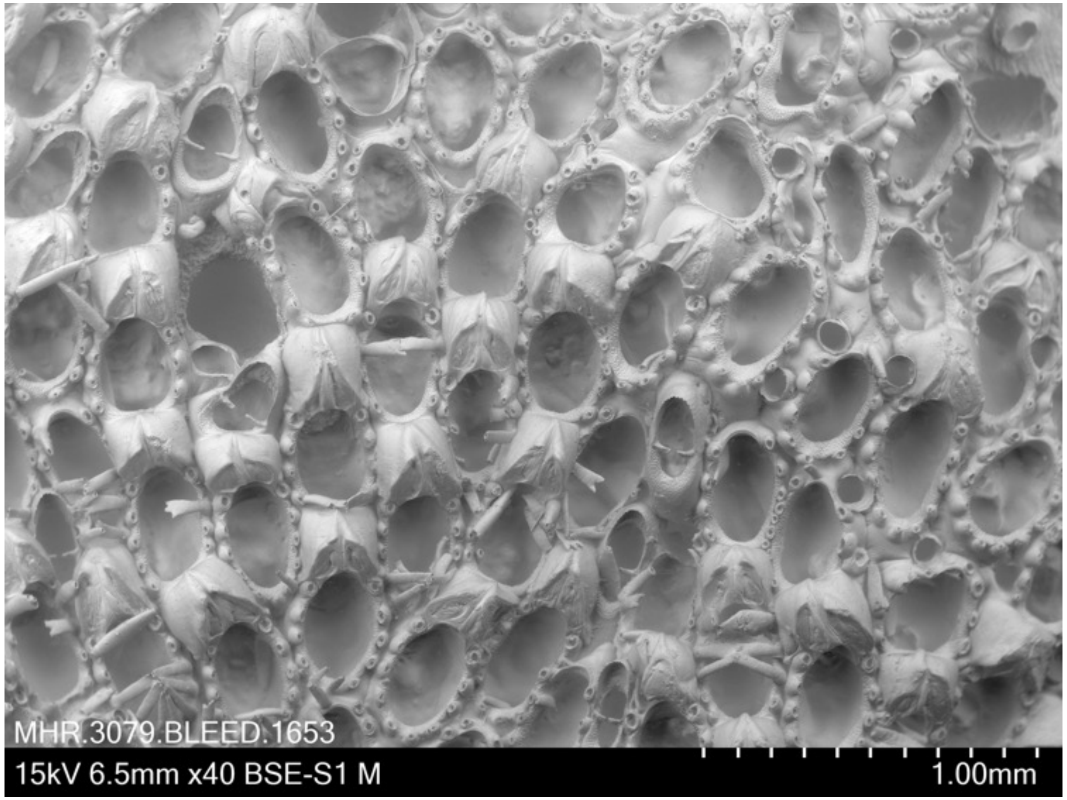


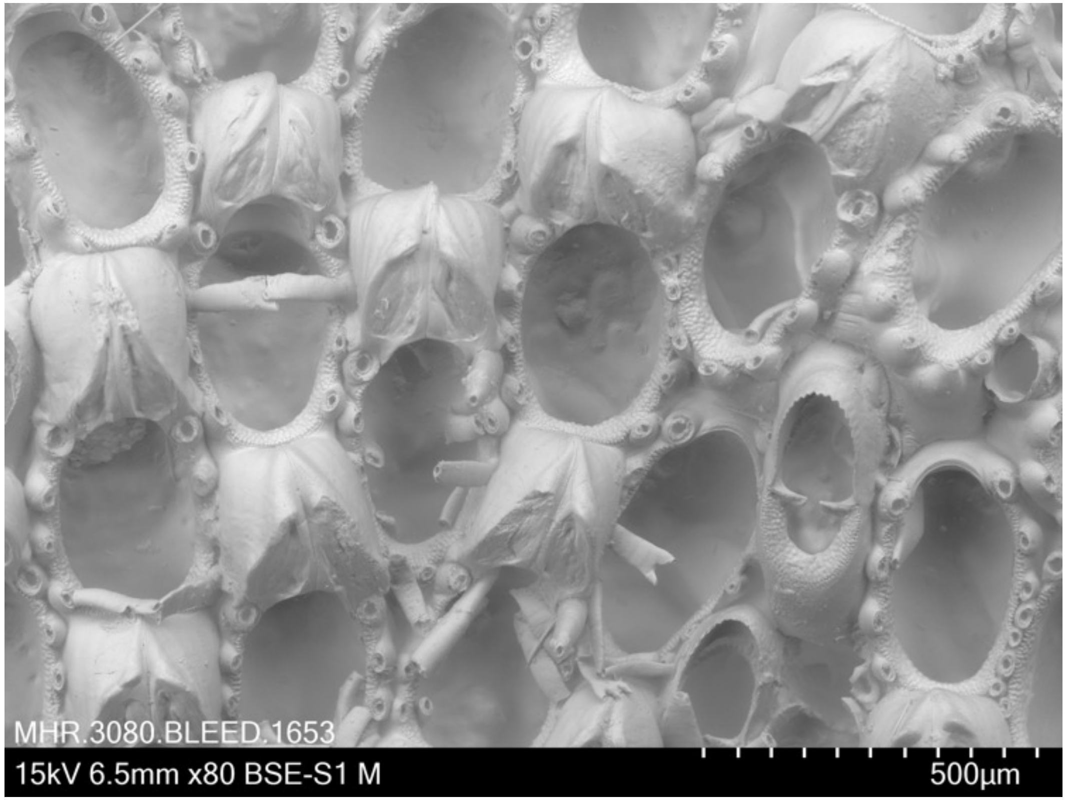


***Valdemunitella* n. sp. BLEED 1653**
Coll: 18.3.2019 (A. M. Smith, D. P. Gordon)
Lat: -34.3800
Long: 172.86
Depth: 54.6 m

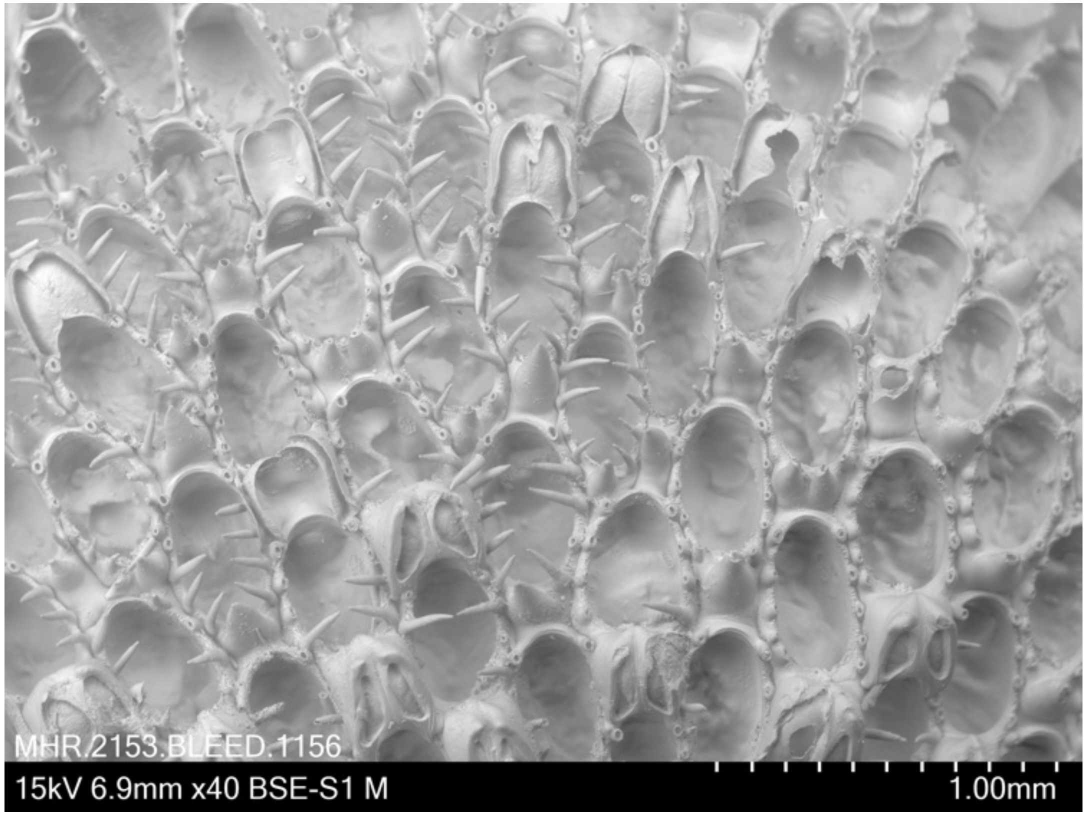

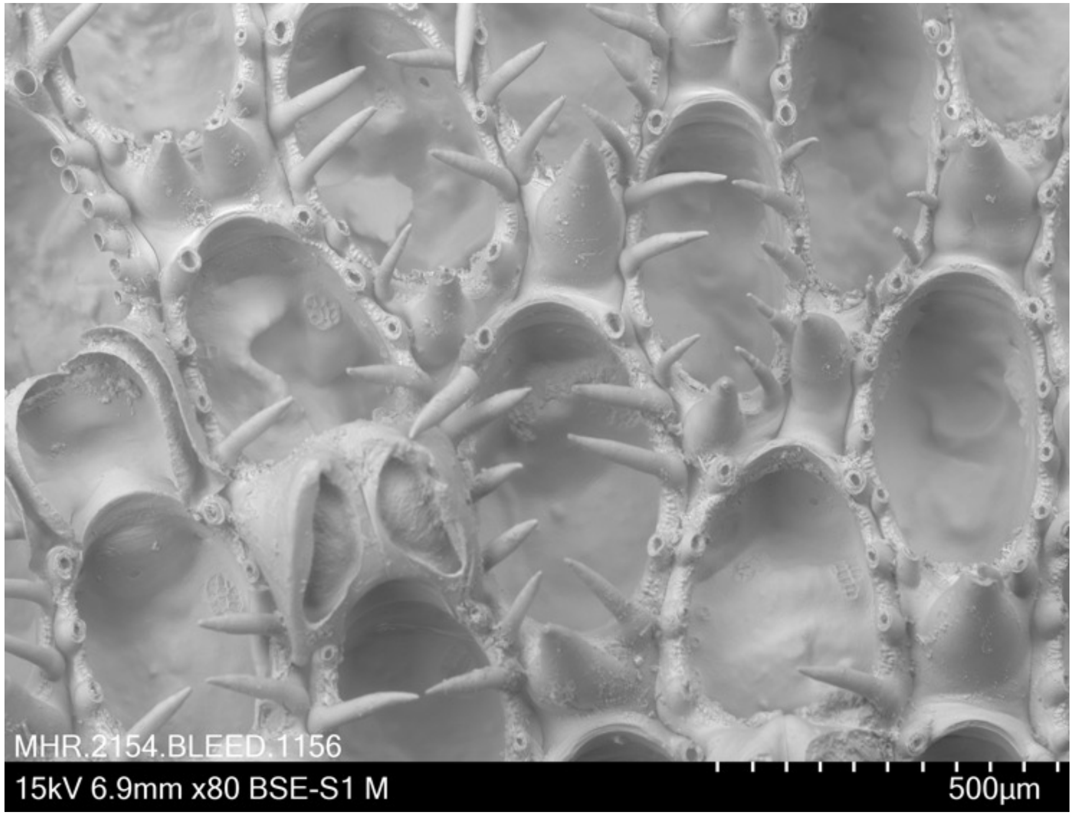


***Valdemunitella pyrula* BLEED 1156**
Coll: 10.11.18 (A. M. Smith)
Lat: -45.840555999999999
Long: 170.796111
Depth: 56 m

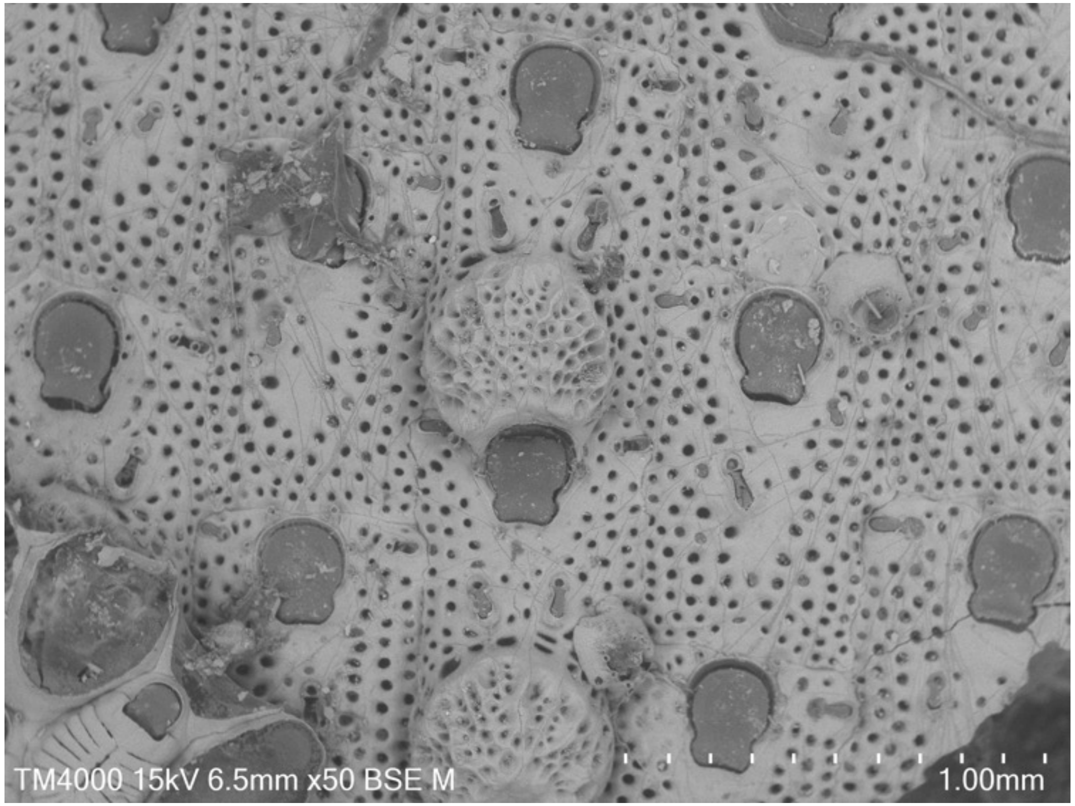


***Valdemunitella huttoni* BLEED 195**
Coll: 3.5.2012
Lat: -43.5166667
Long: 173.6736667
Depth: 85 m
**Notes:** just visible on the left corner; previously *Figularia huttoni*

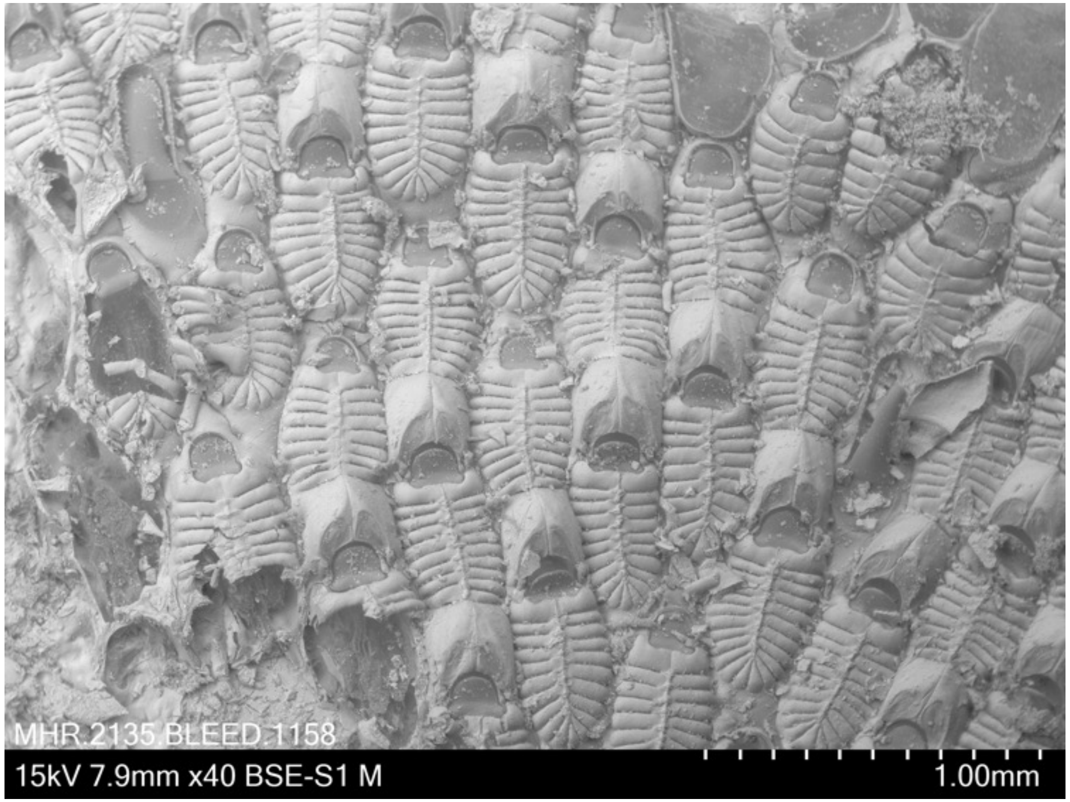


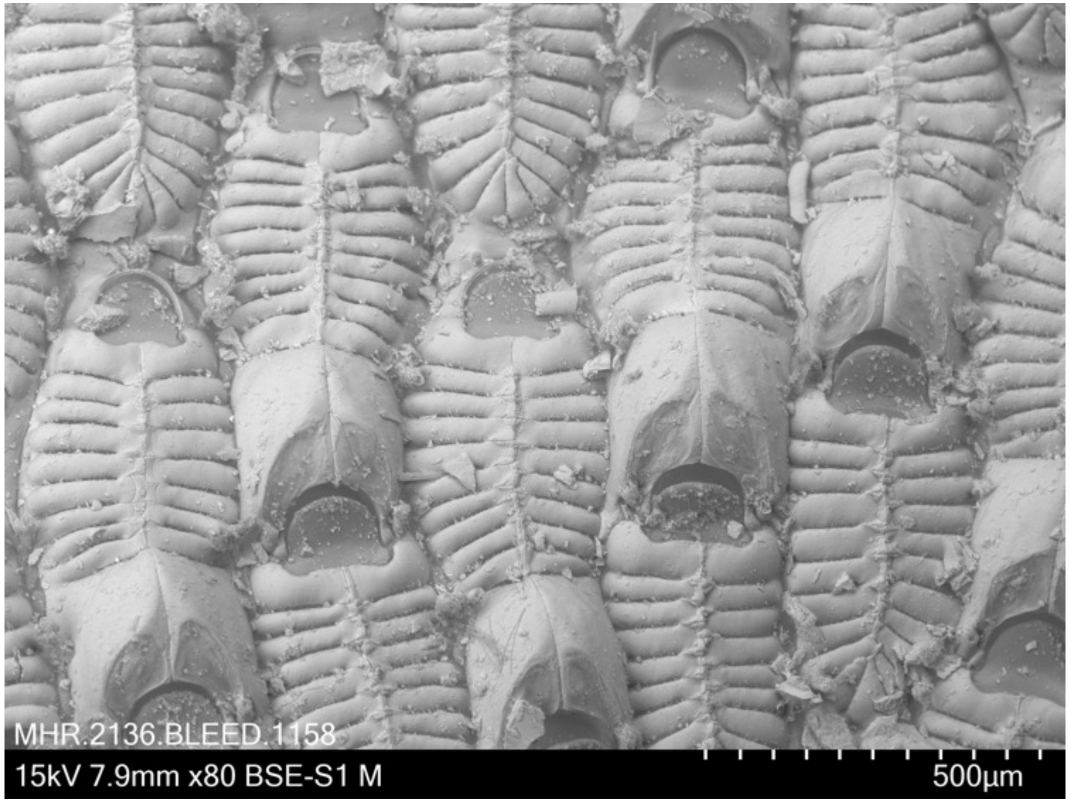


***Valdemunitella huttoni* BLEED 1158**
Coll: 10.11.2018 (A. M. Smith)
Lat: -45.849167000000001
Long: 170.86111099999999
Depth: 78.2 m
<u>**Notes:**</u> previously *Figularia huttoni*

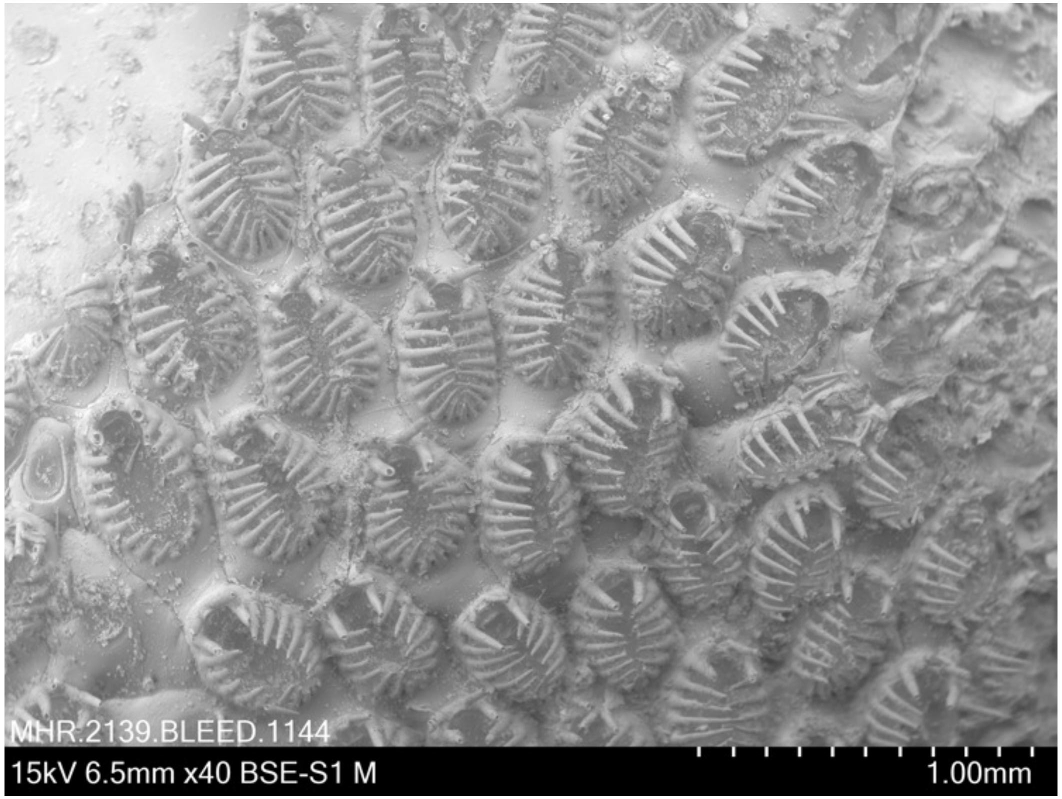


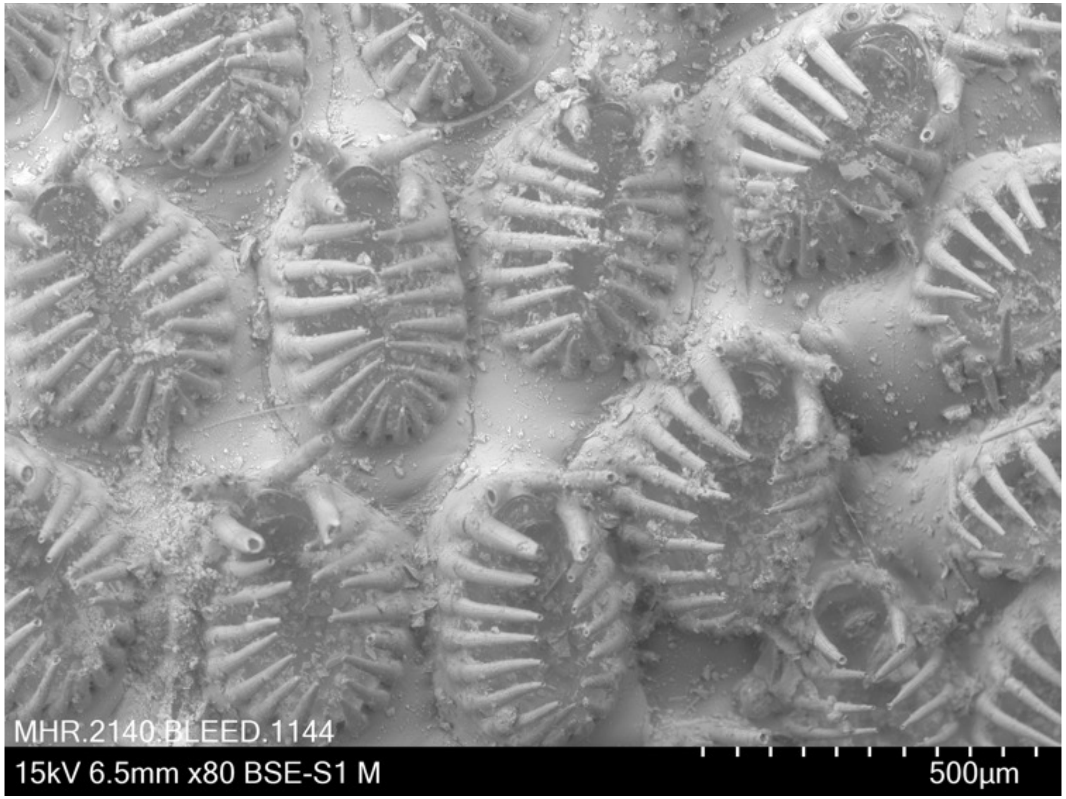


***Valdemunitella* cf. *fraudatrix* BLEED 1144B**

Coll: 10.11.2018 (A. M. Smith)

Lat: -45.835833000000001

Long: 170.96166700000001

Depth: 131 m

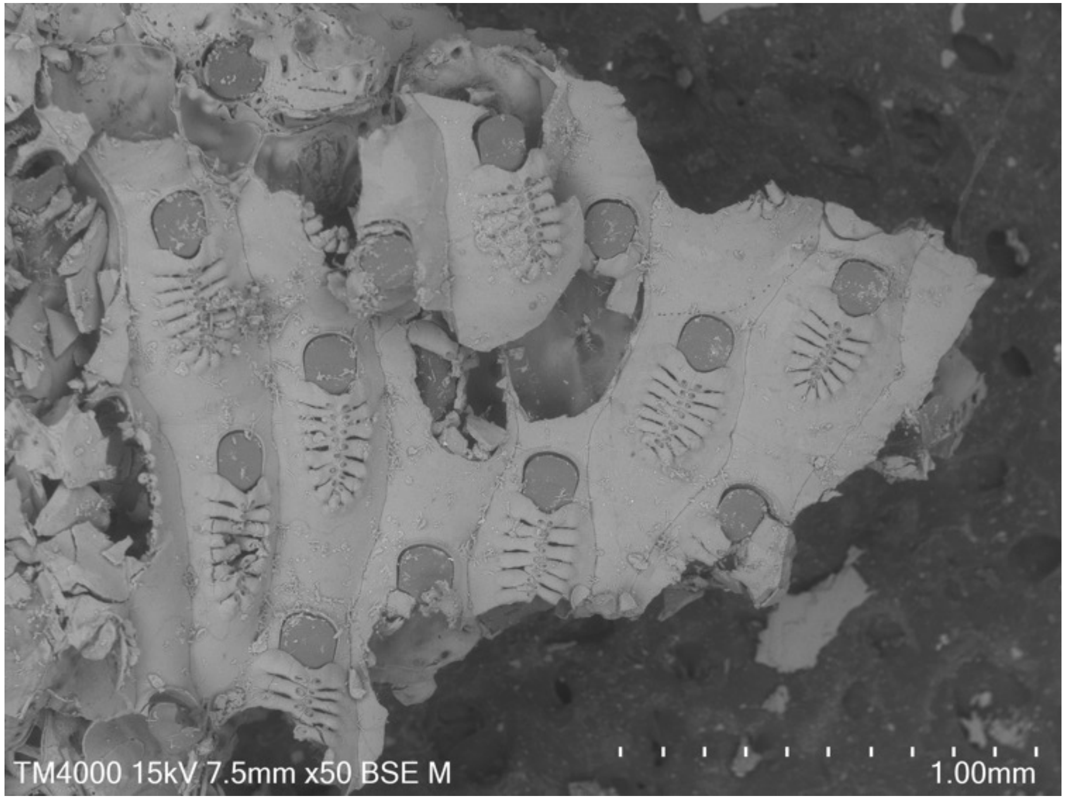

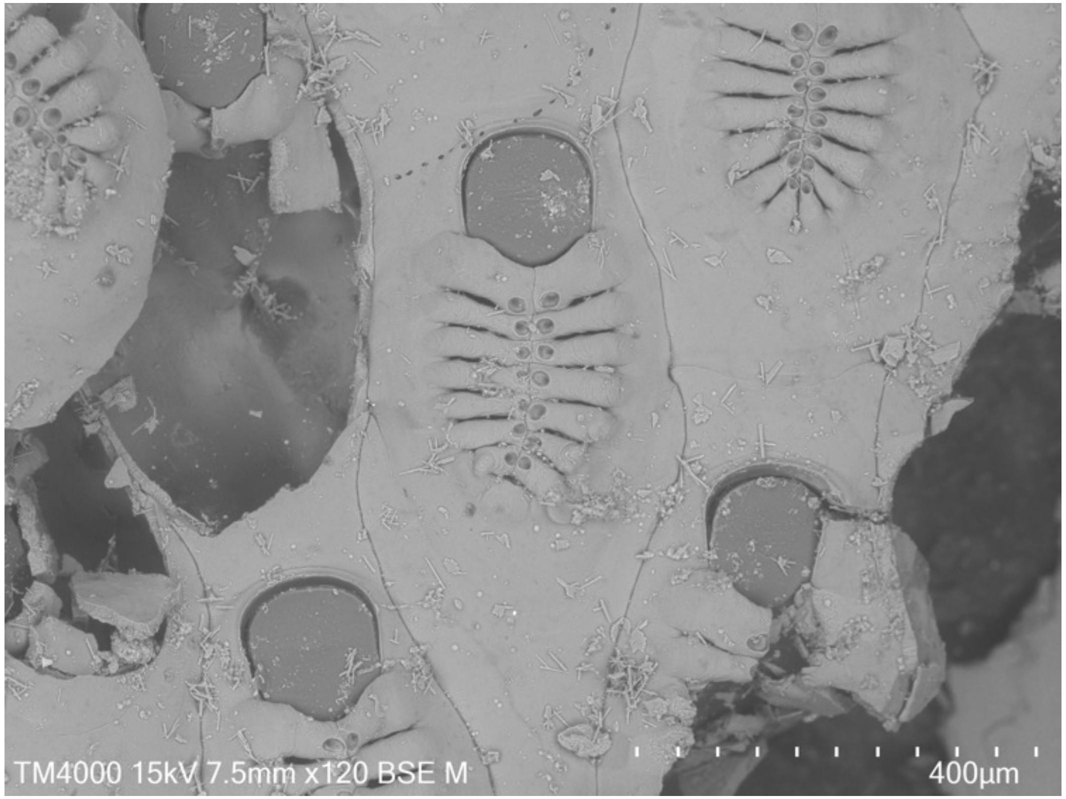


***Figularia mernae*** **BLEED 179**
NIWA 39690 (TAN0803/63)
Coll: 9.4.2008
Lat: -52.487333
Long: 160.415
Depth: 350 m

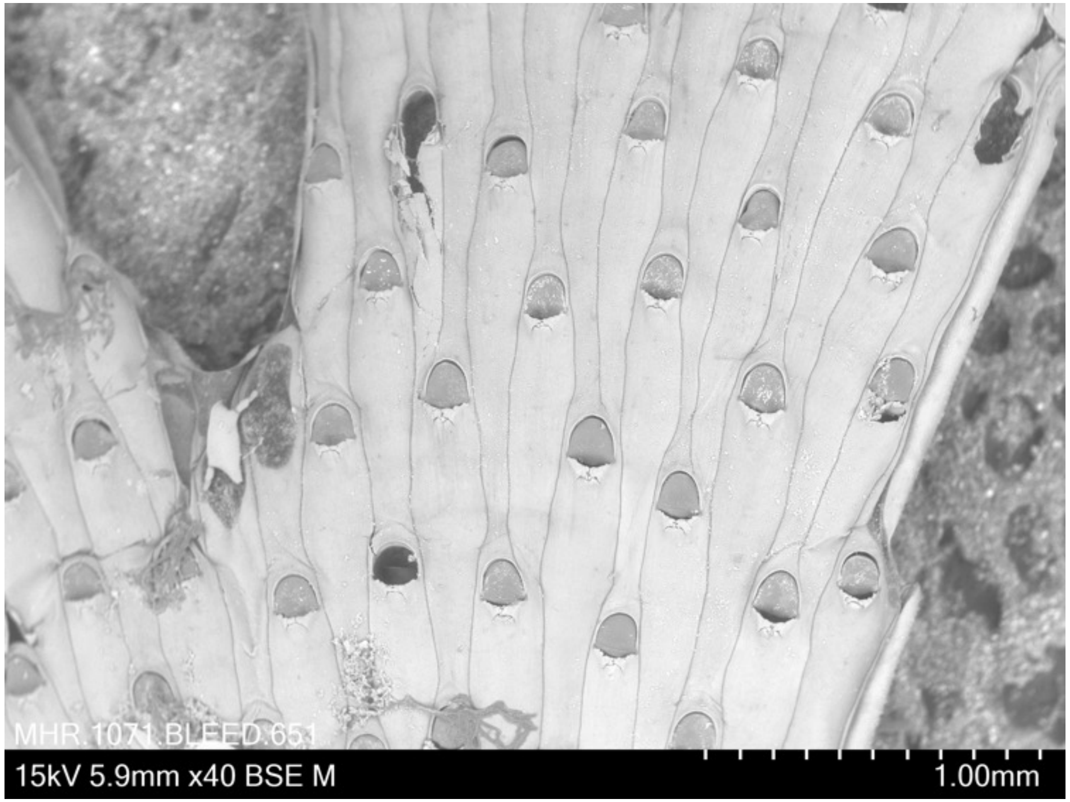


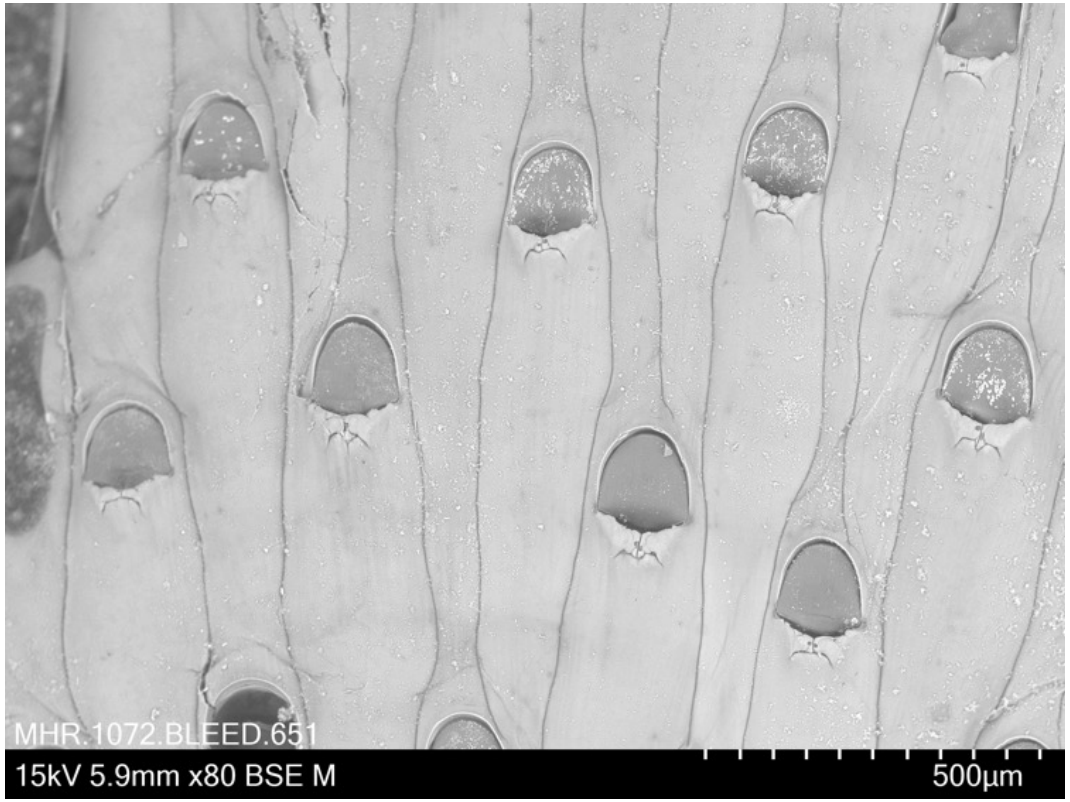


***Euthyroides episcopalis* BLEED 651**
Coll: 8.4.2018 (A. M. Smith, D. P. Gordon, H. Mello)
Lat: -47.17361
Long: 167.94583
Depth: 66 m

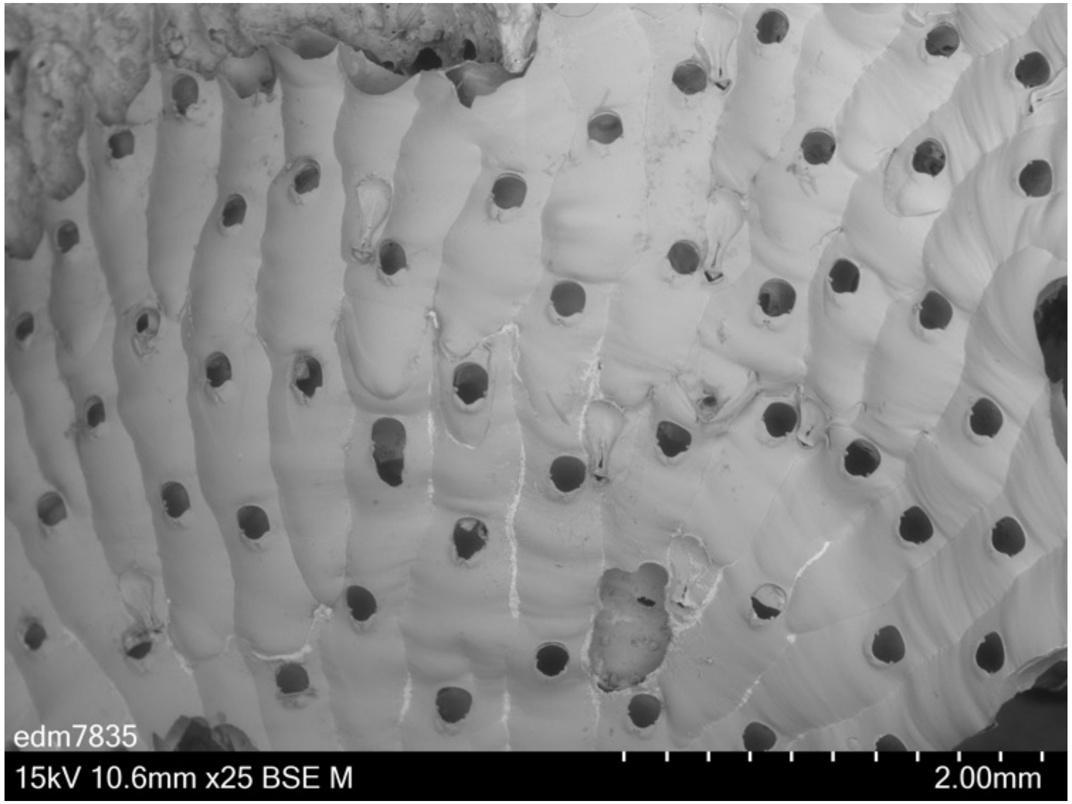


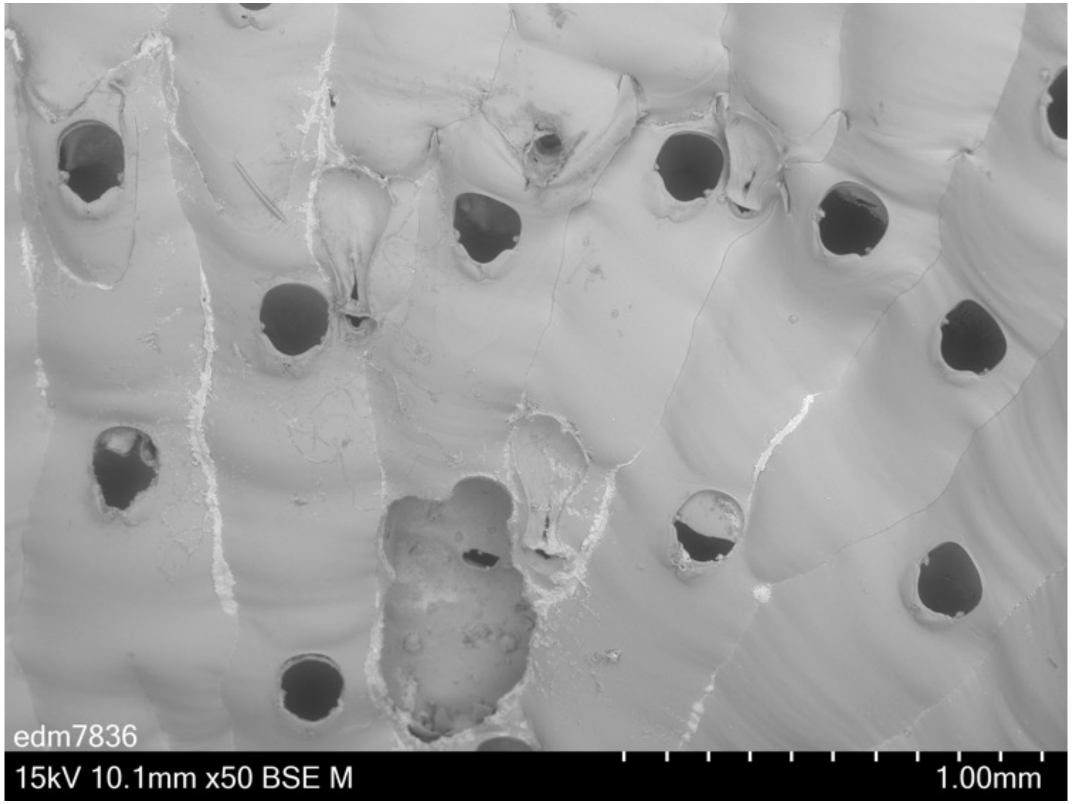


***Euthyroides* n. sp. BLEED 1344**
Coll: 19.3.2019 (A. M. Smith, D. P. Gordon)
Lat: -34.979999999999997
Long: 174.02
Depth: 106 m

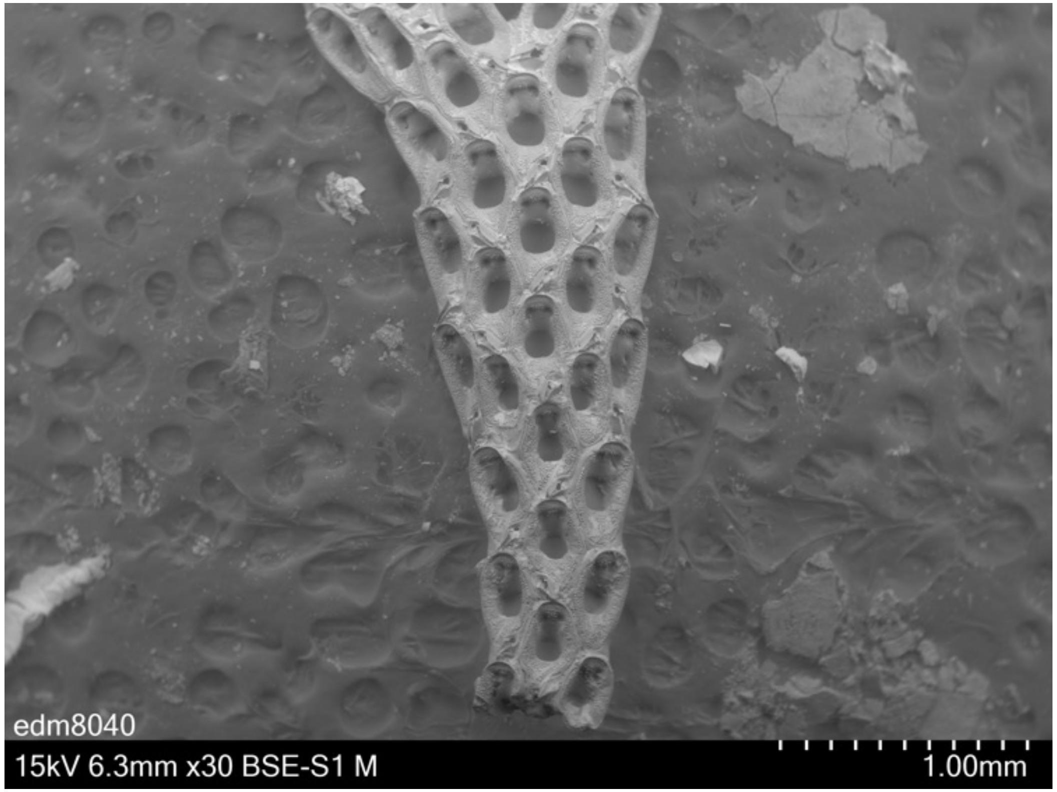


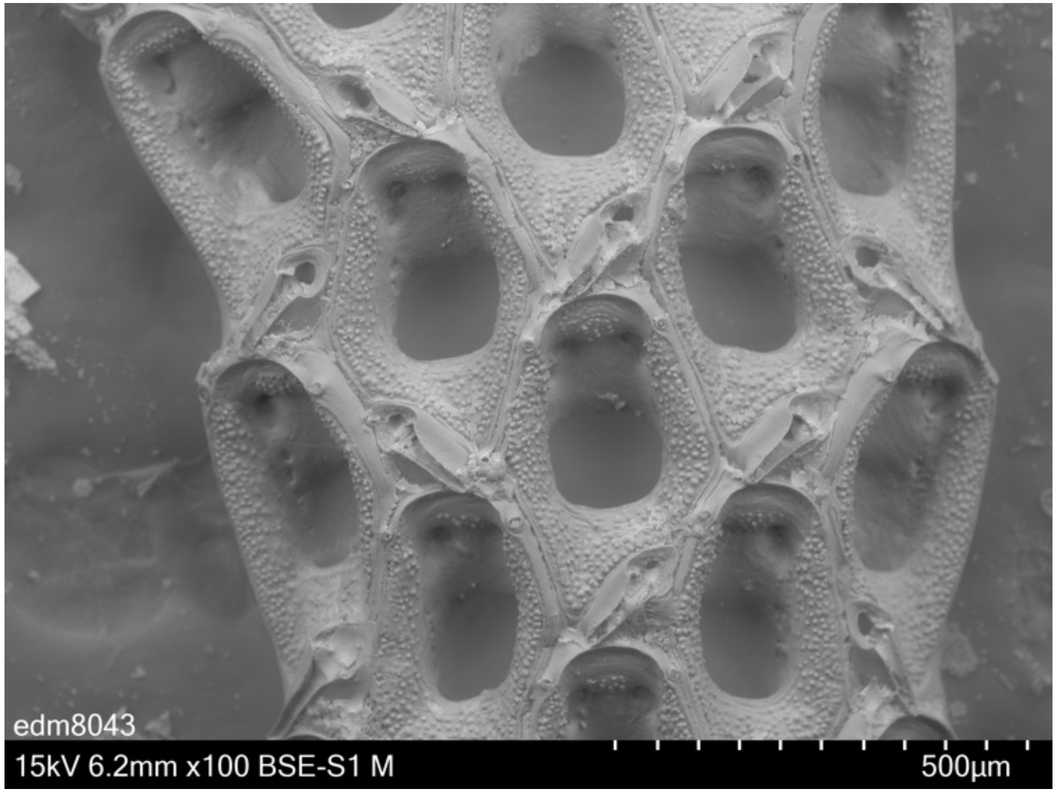


***Menipea* n. sp. 1 BLEED 1340**

Coll: 19.3.2019 (A. M. Smith, D. P. Gordon)

Lat: -34.979999999999997

Long: 174.02

Depth: 106 m

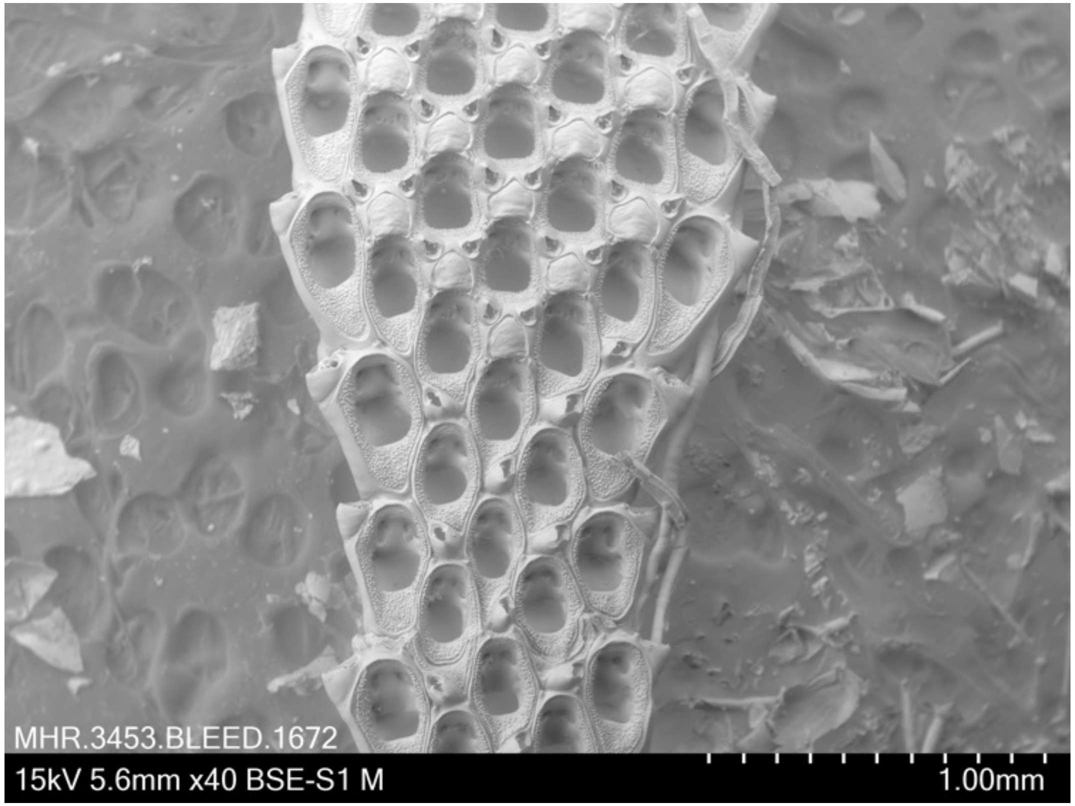

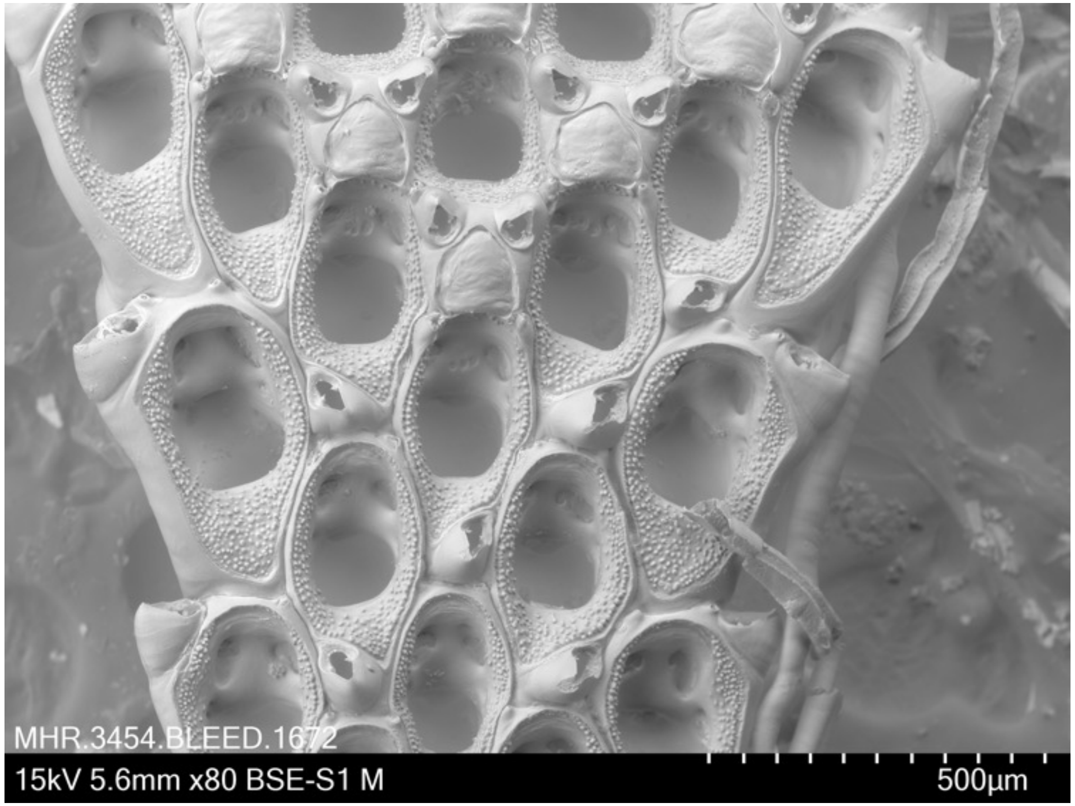


***Menipea* n. sp. 2 BLEED 1672**
Coll: 18.3.2019 (A. M. Smith, D. P. Gordon)
Lat: -34.380000000000003
Long: 172.69
Depth: 52.5-56.5 m

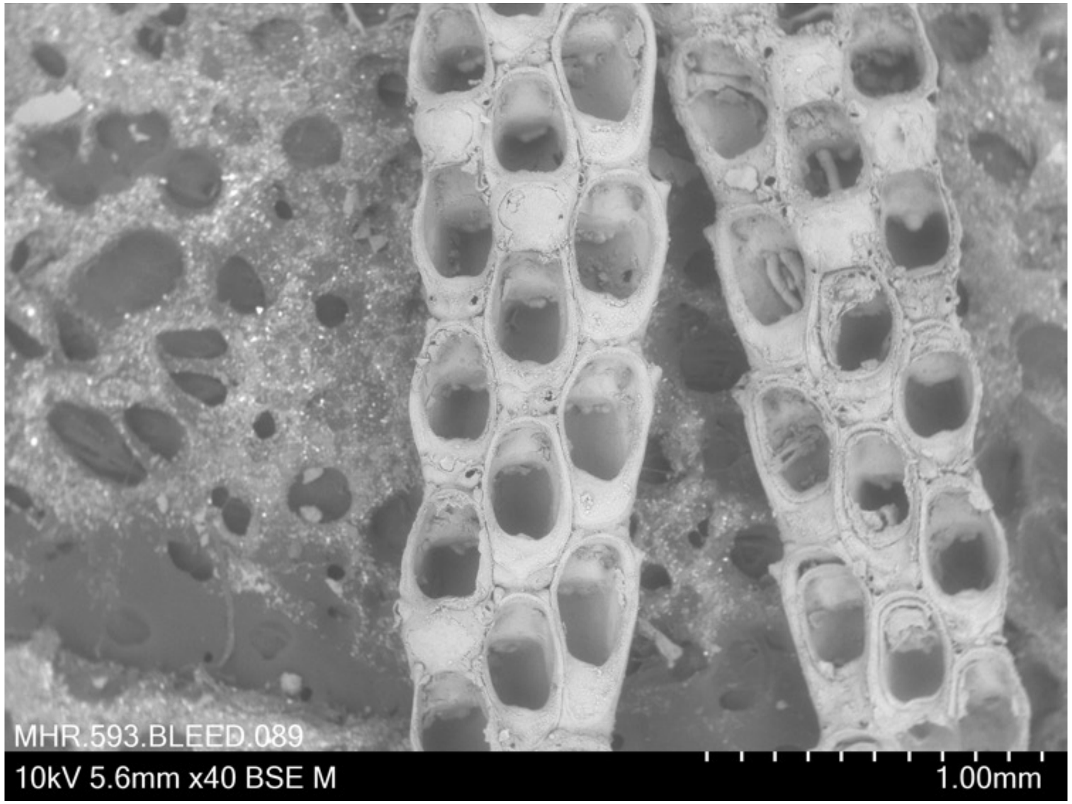


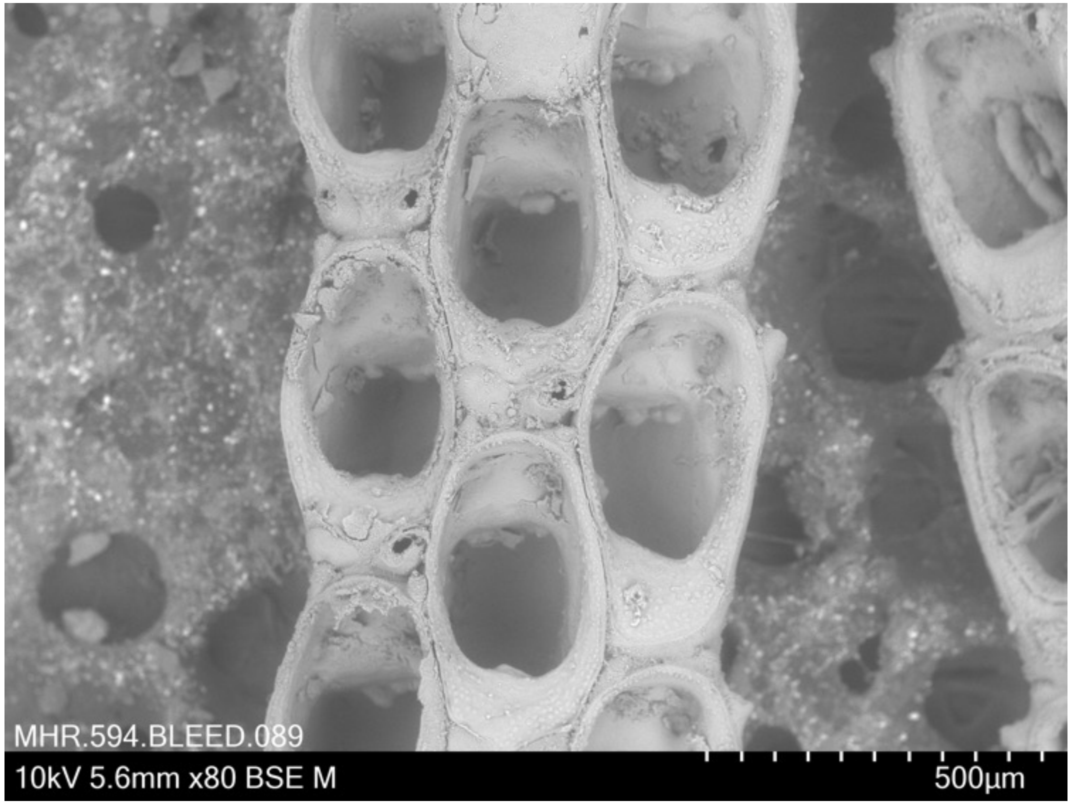


***Menipea vectifera* BLEED 89**
Coll: D. P. Gordon
Location: New Zealand
**Notes:** other meta data missing

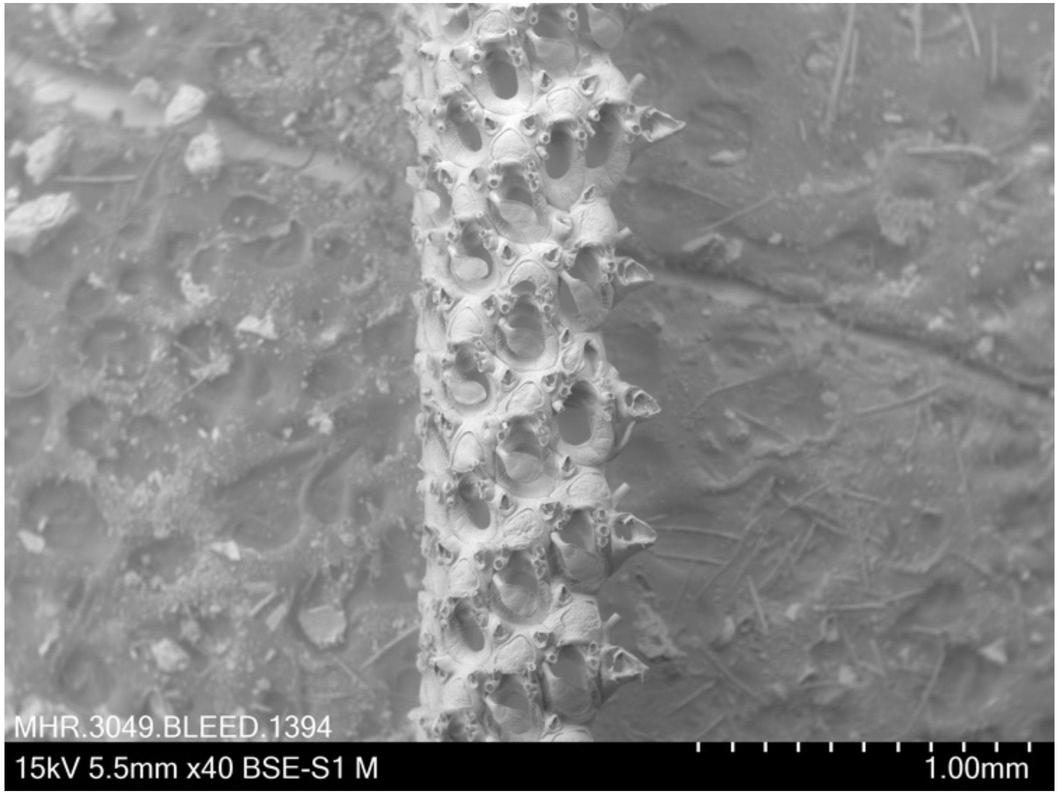


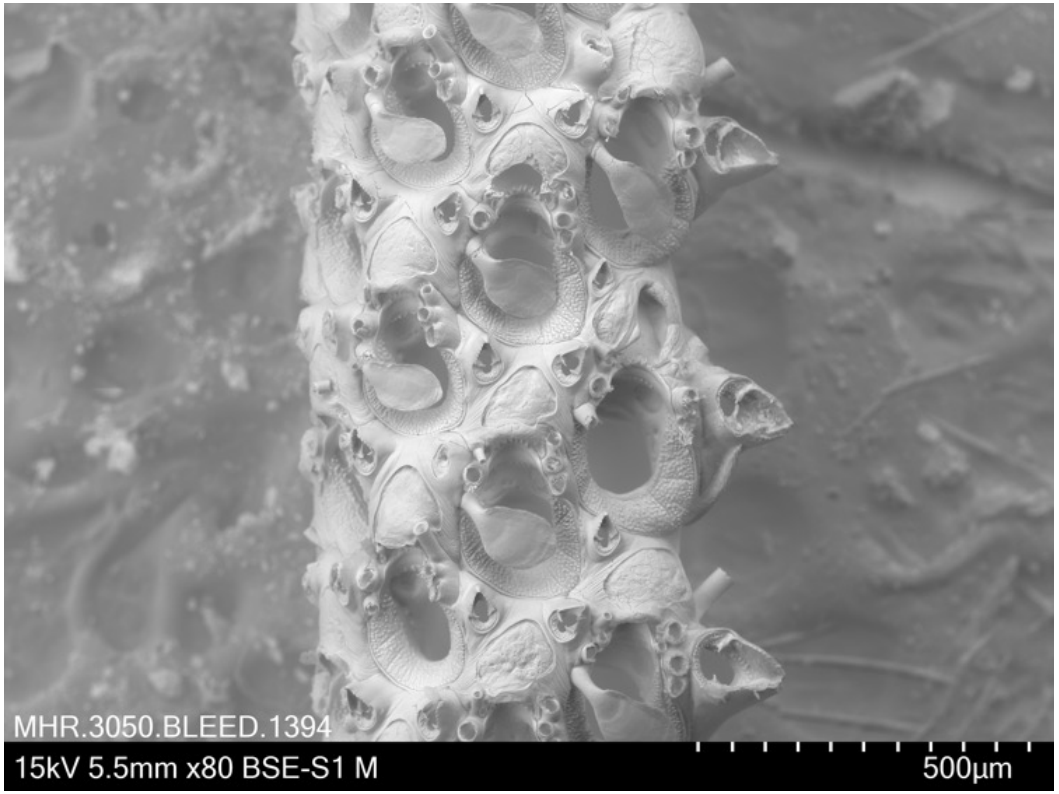


***Amastigia harmeri* BLEED 1394**
Coll: 17.3.2019 (A. M. Smith, D. P. Gordon)
Lat: -34.159999999999997
Long: 172.09
Depth: 82.9 m

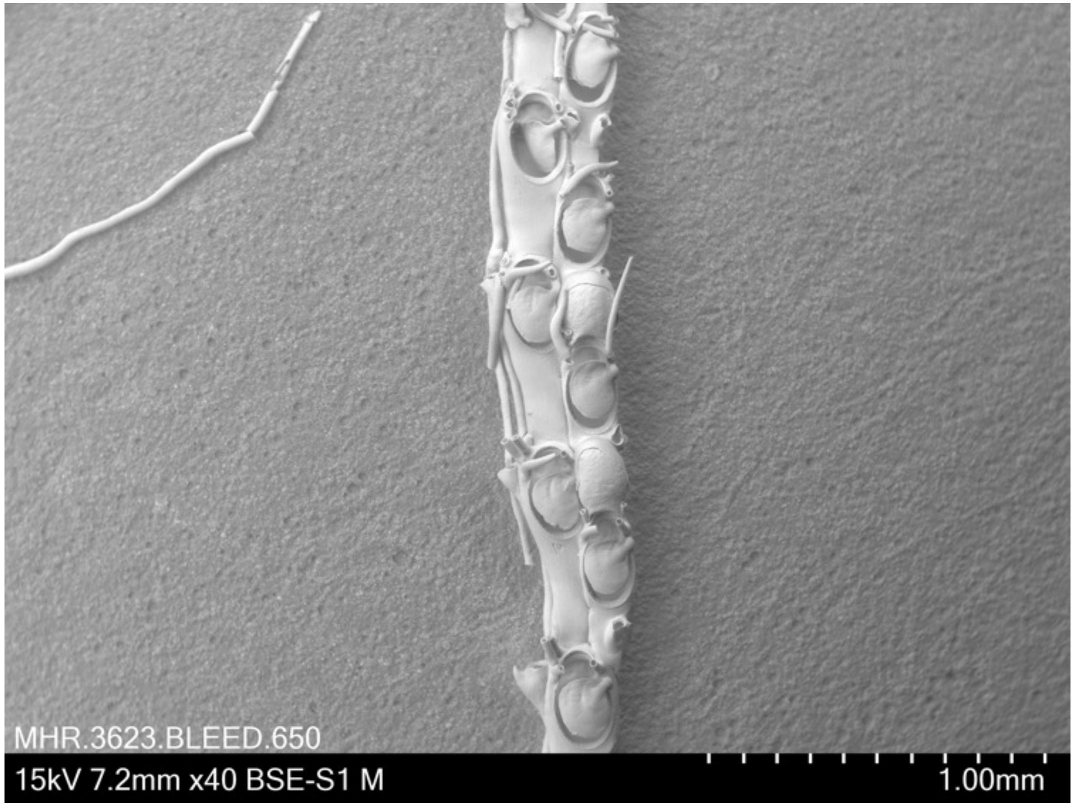


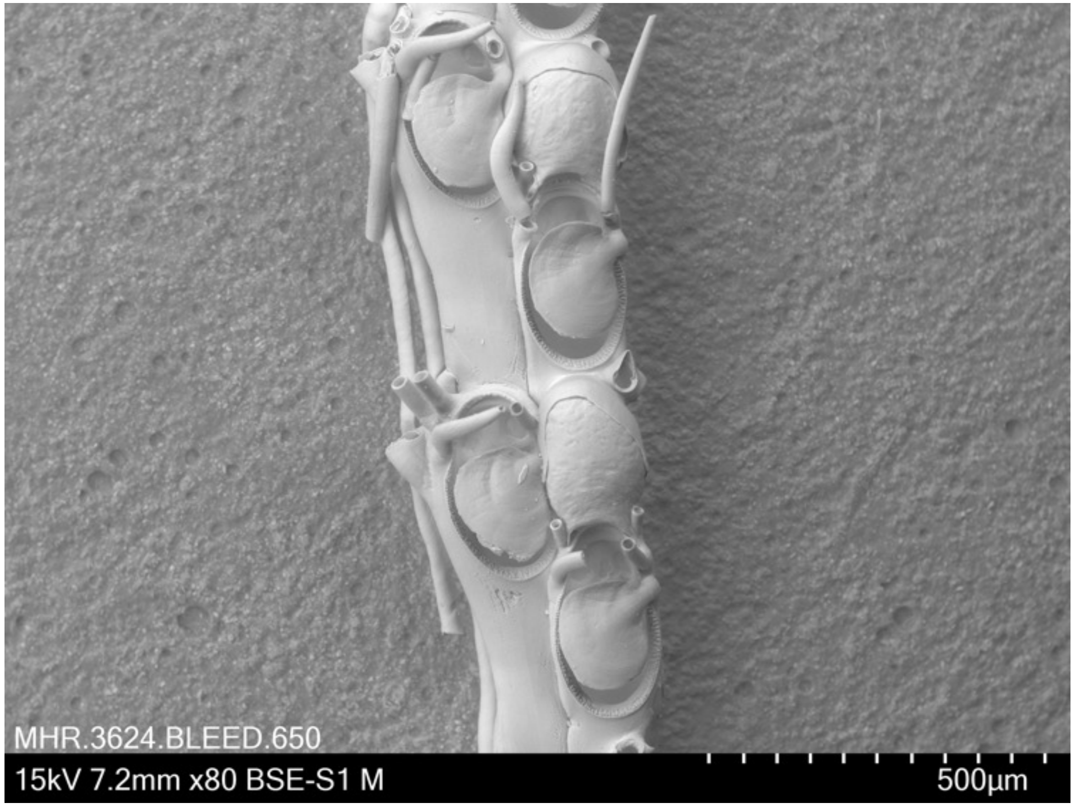


***Amastigia funiculata* BLEED 650**
Coll: 8.4.2018 (A. M. Smith, D. P. Gordon, H. Mello)
Lat: -47.17361
Long: 167.9458
Depth: 66 m

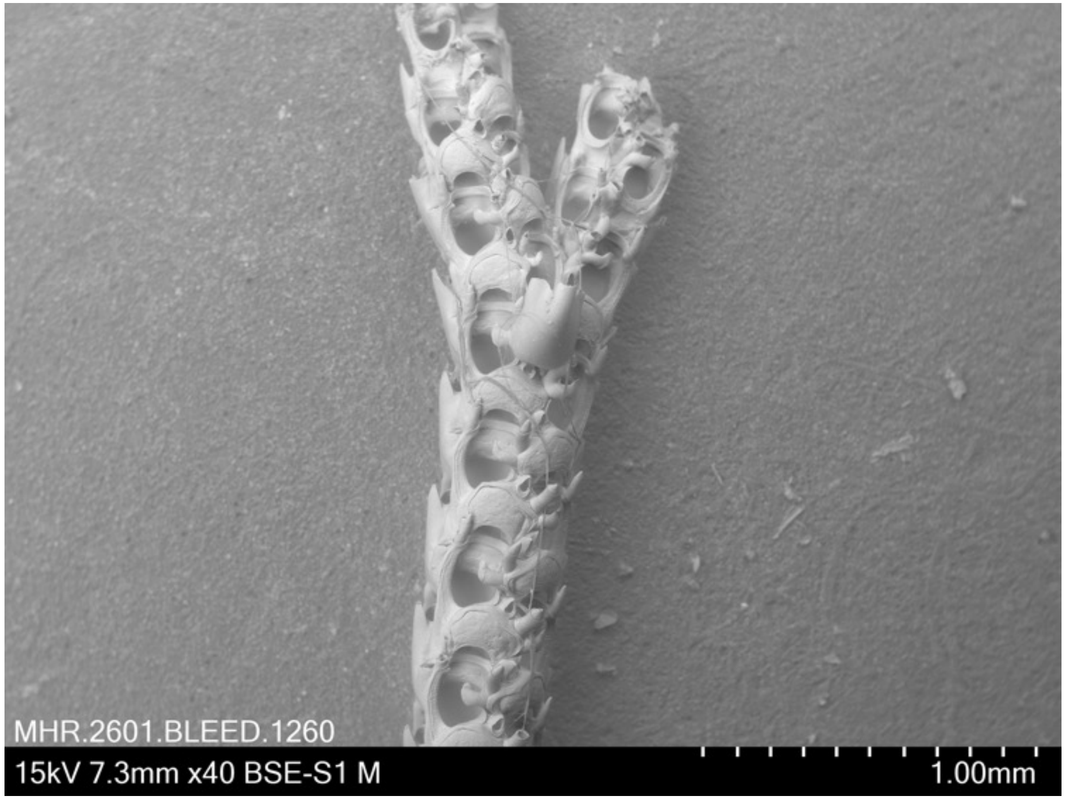


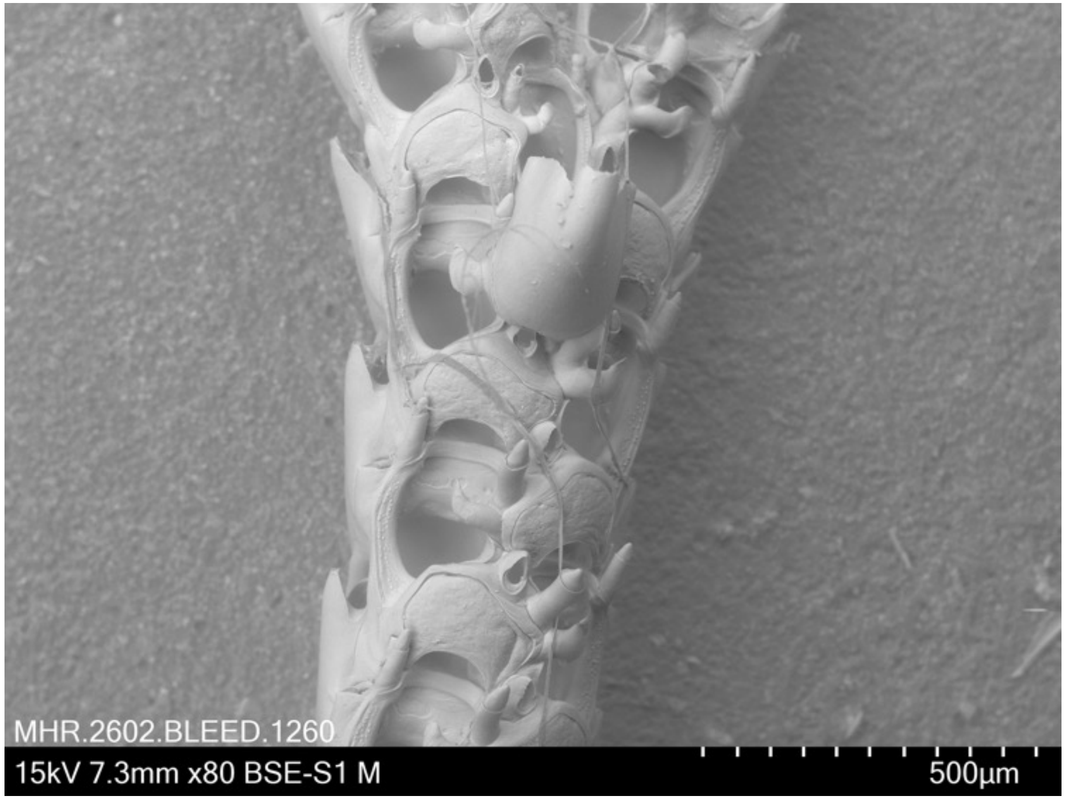


***Caberea rostrata* BLEED 1260**
Coll: 16.3.2019 (A. M. Smith, D. P. Gordon)
Lat: -35.01
Long: 173.92
Depth: 17 m

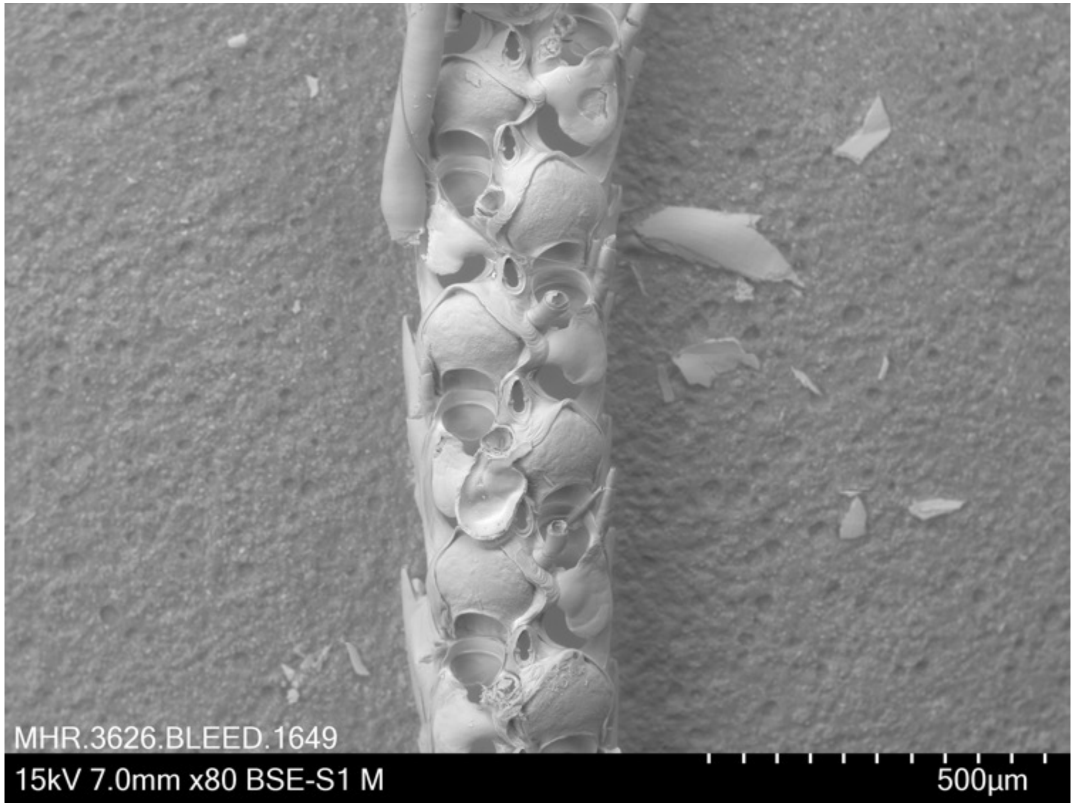


***Caberea* aff. *rostrata* BLEED 1649**
Coll: 17.3.2019 (A. M. Smith; D. P. Gordon)
Lat: -34.159999999999997
Long: 172.09
Depth: 82.9 m

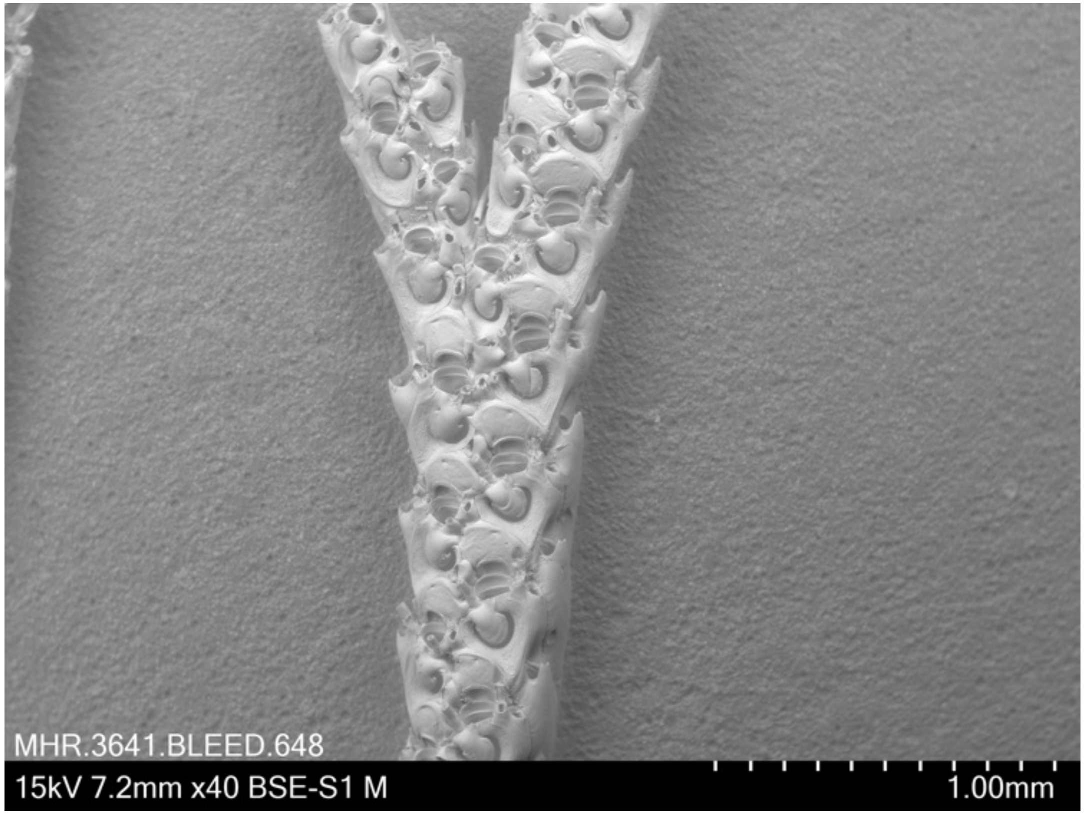


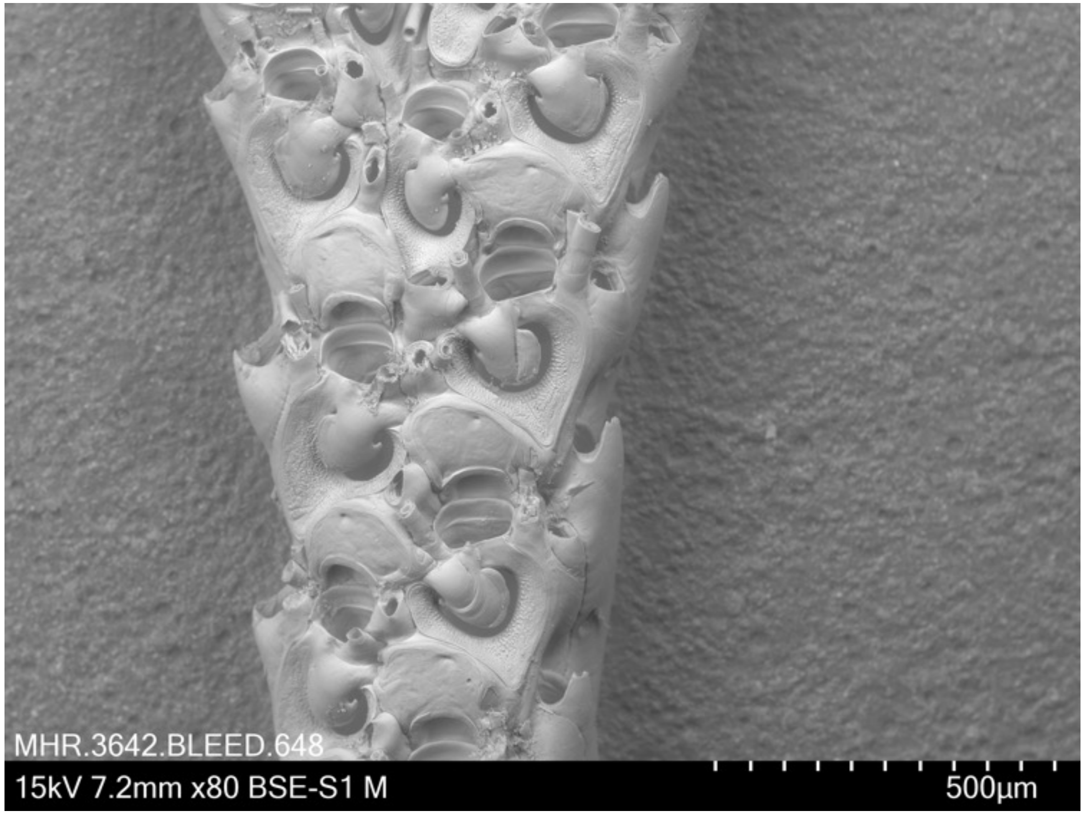


***Caberea zelandica* BLEED 648**
Coll: 8.4.2018 (A. M. Smith, D. P. Gordon, H. Mello)
Lat: -47.17361
Long : 167.94583
Depth: 66 m

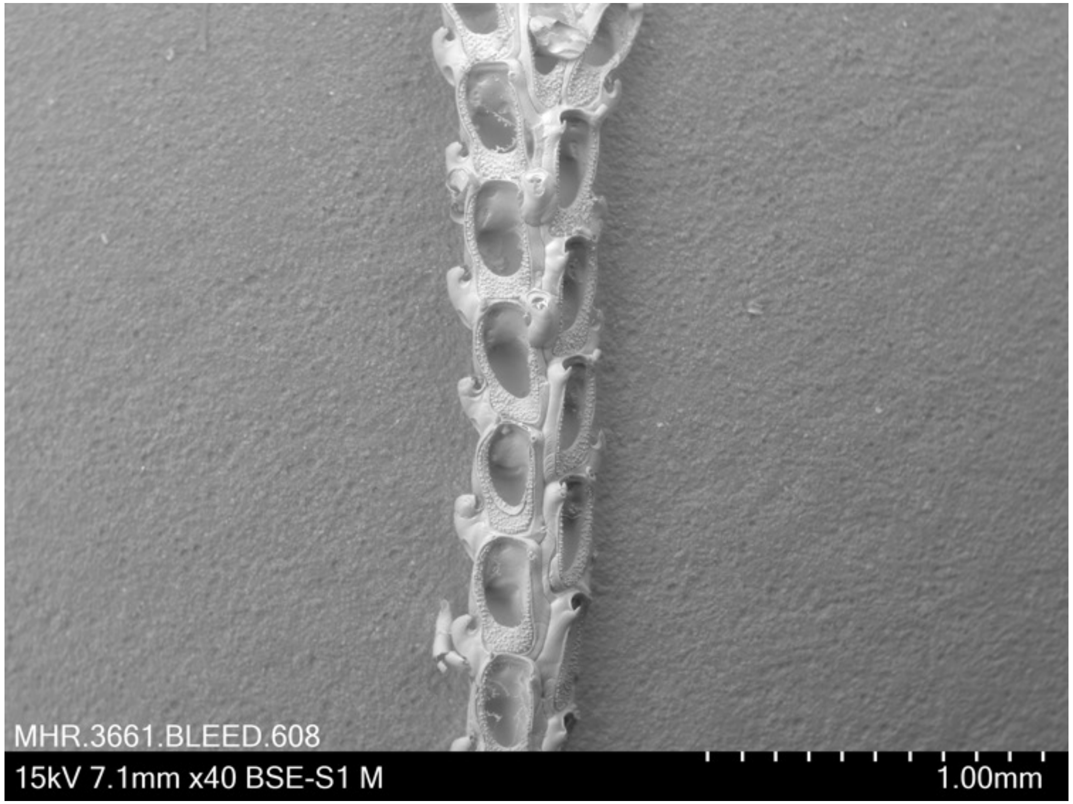


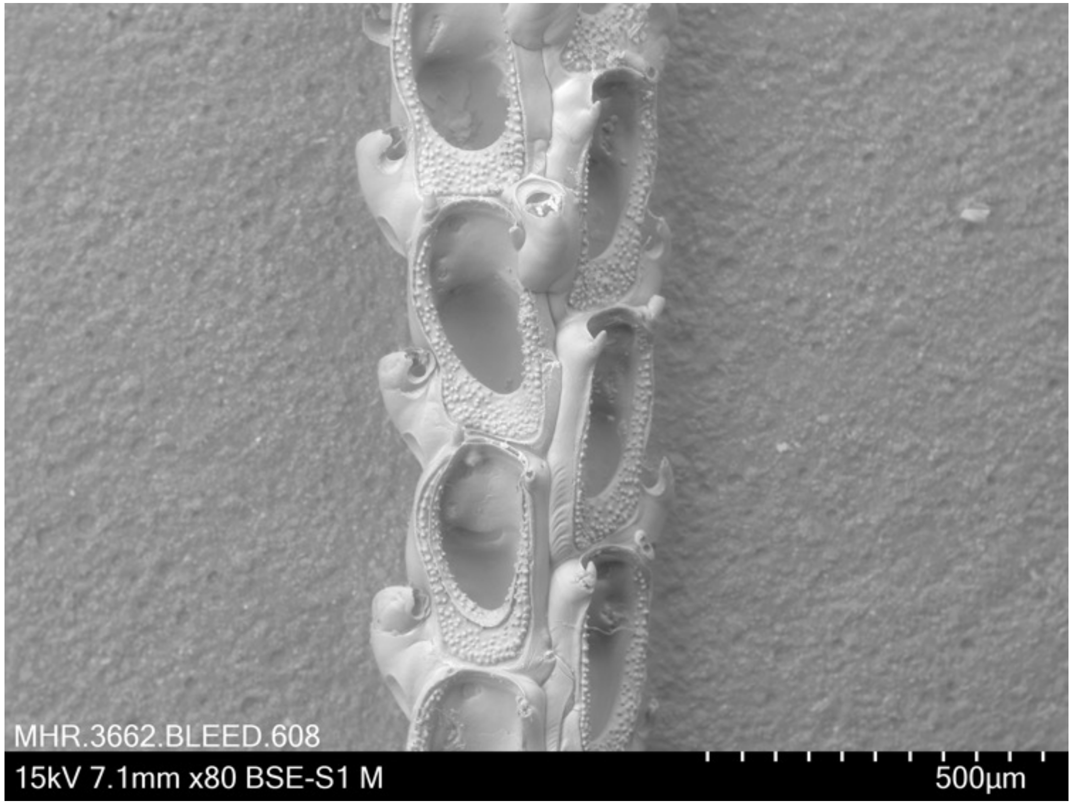


***Canda filifera* BLEED 608**
Coll: 11.4.2018 (A. M. Smith, D. P. Gordon, H. Mello)
Lat: -46.8536
Long: 168.2611
Depth: 24 m

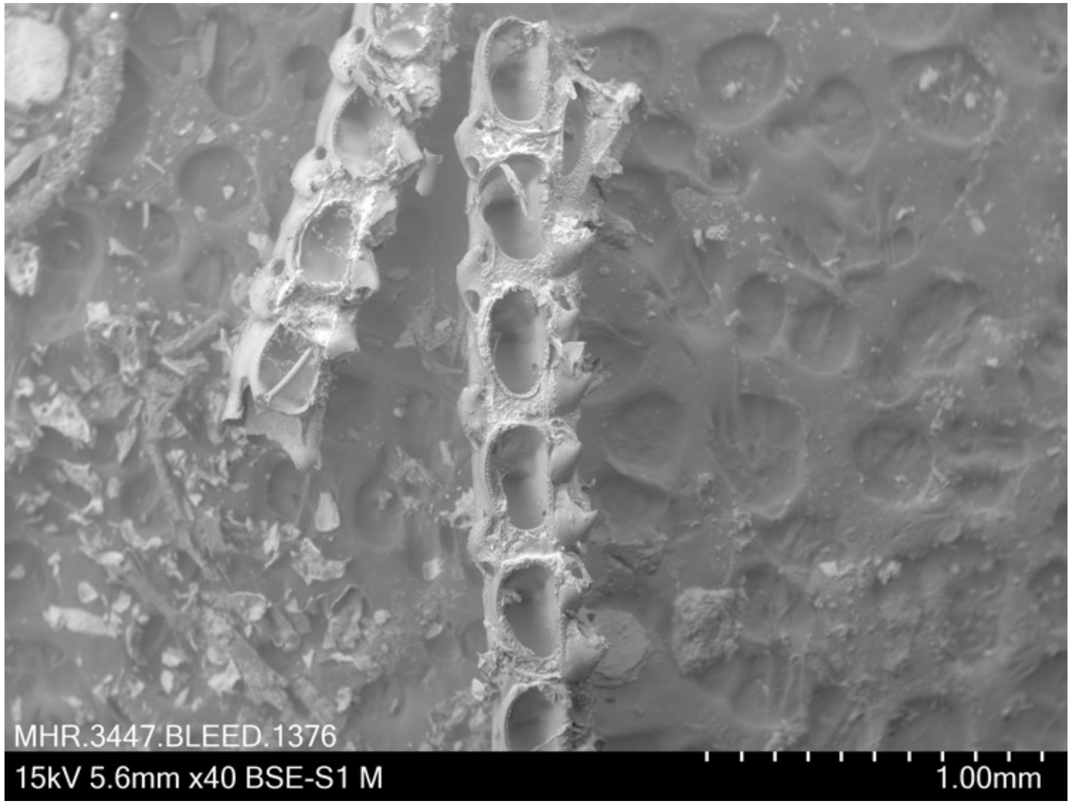


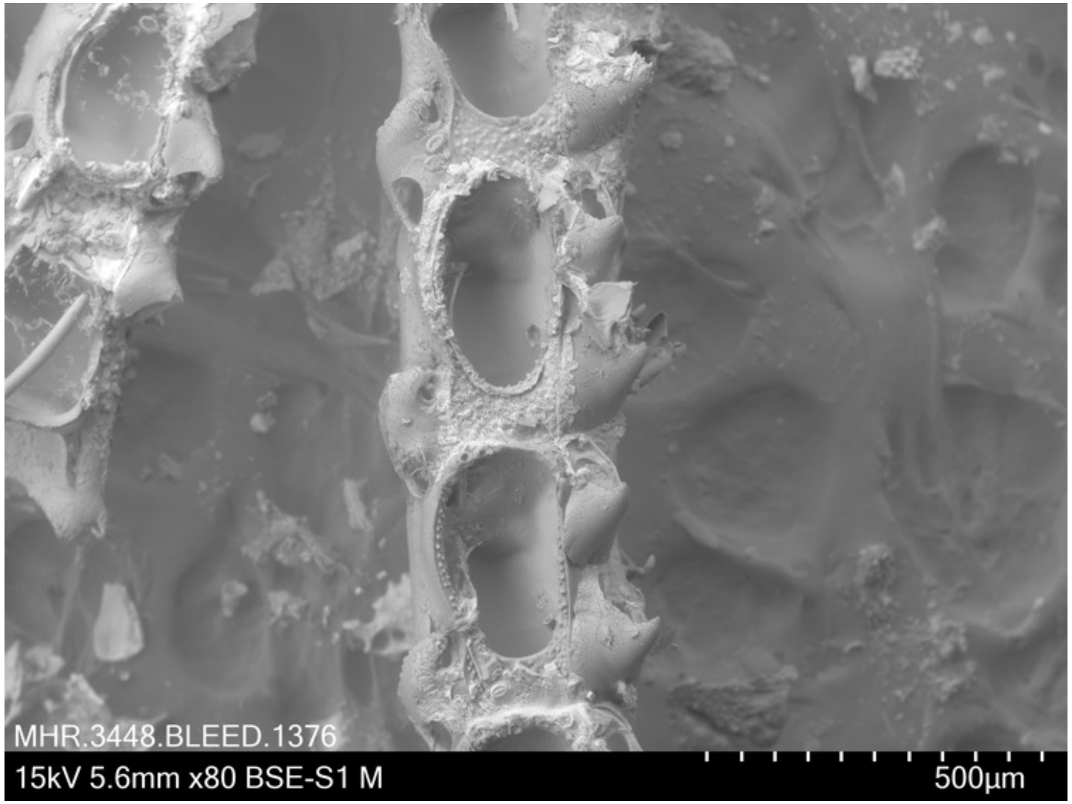


***Canda filifera* BLEED 1376**
Coll: 17.3.2019 (A. M. Smith, D.P. Gordon)
Lat: -34.159999999999997
Long: 172.09
Depth: 82.9 m

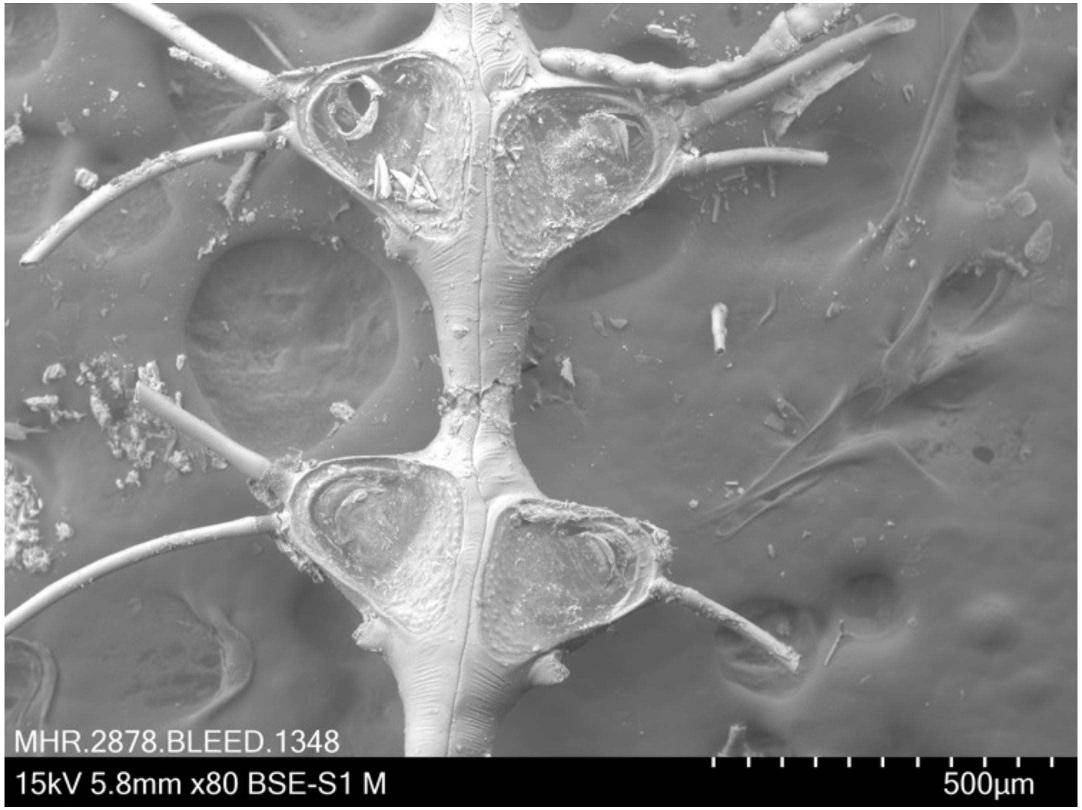


***Emma triangula*** **BLEED 1348**
Coll: 17.3.2019 (A. M. Smith,  D.P. Gordon)
Lat: -34.21
Long: 172.11
Depth: 104 m

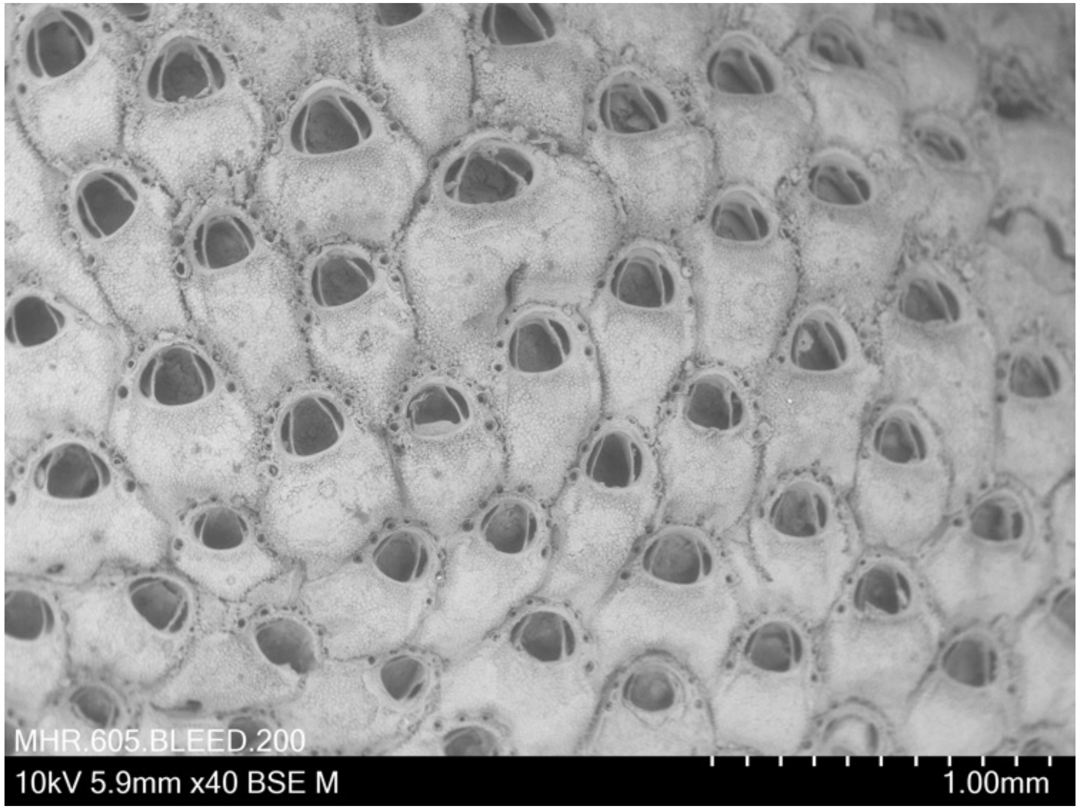


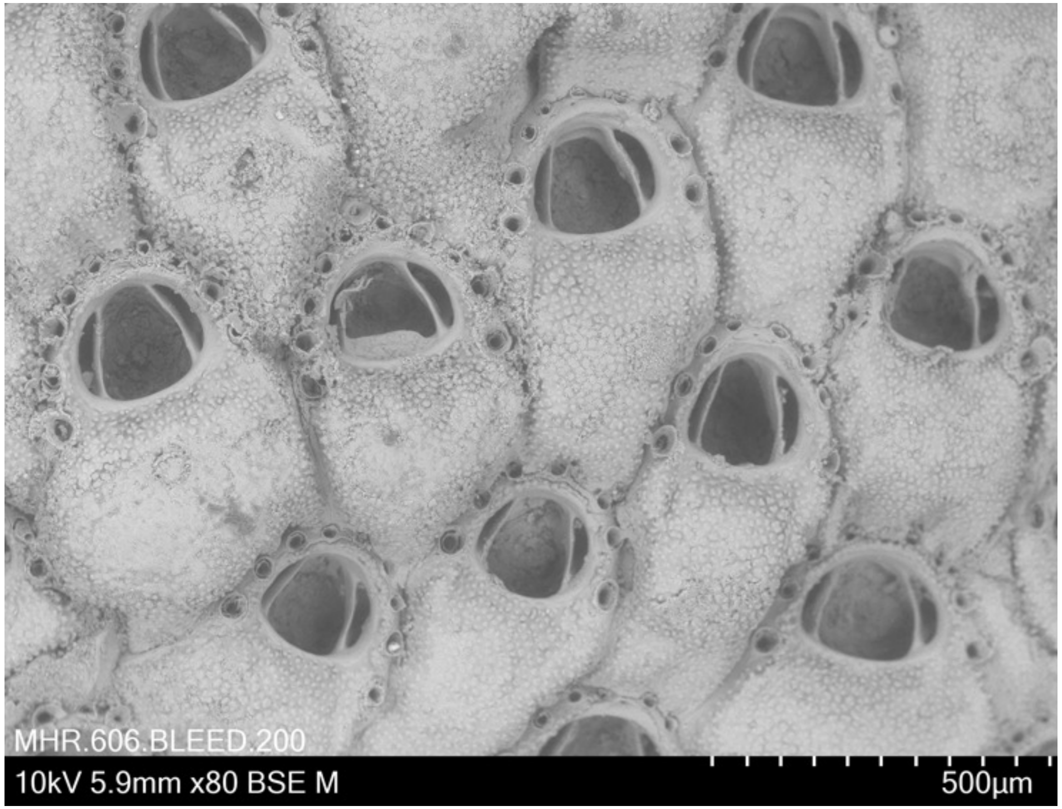


***Chaperia* n. sp. 1 BLEED 200**
Coll: 20.5.2015 (D. P. Gordon)
Lat: -41.0710396
Long: 176.088894
Depth: Intertidal

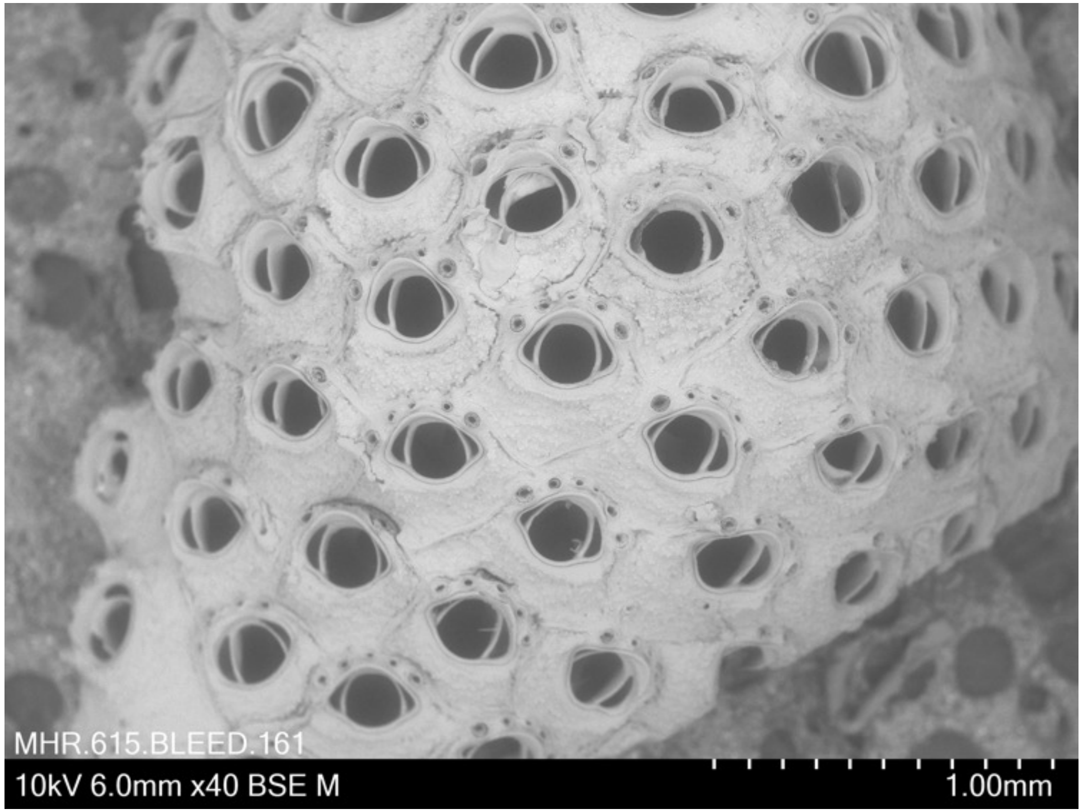


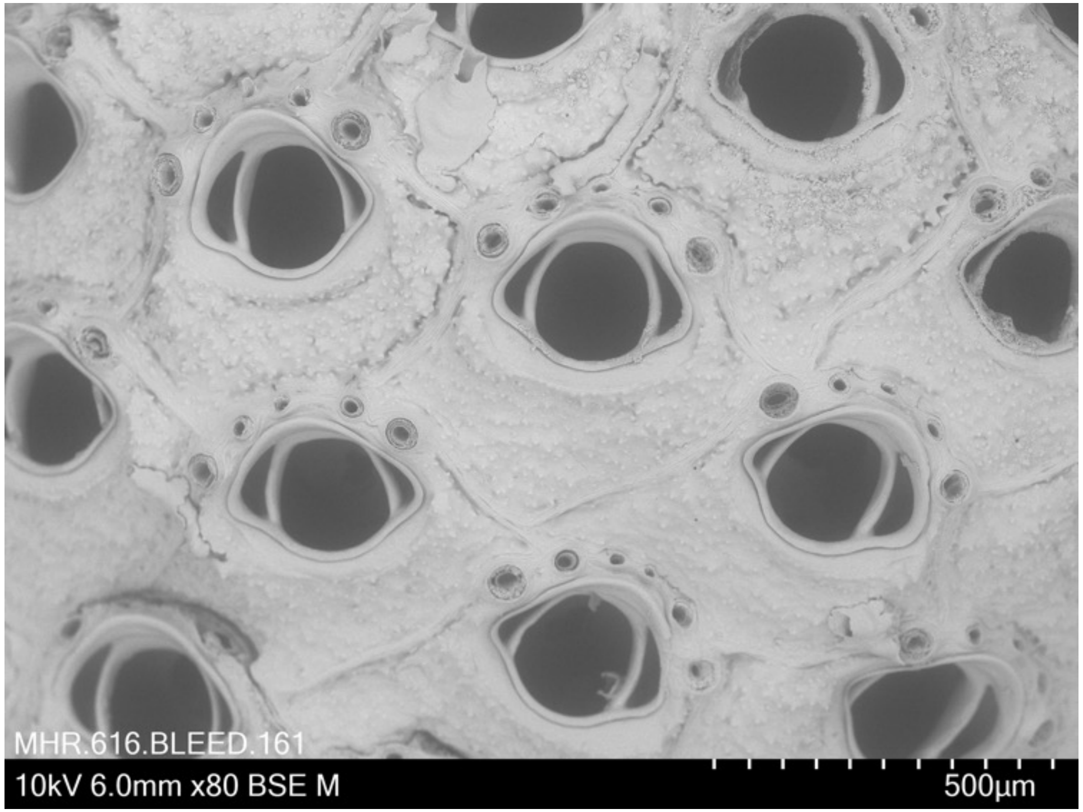


***Chaperia* n. sp. 2 BLEED 161**
NIWA 54782 (TAN0906/25)
Coll: 5.7.2009
Lat: -35.5483333333
Long: 174.55250000000001
Depth: 57 m

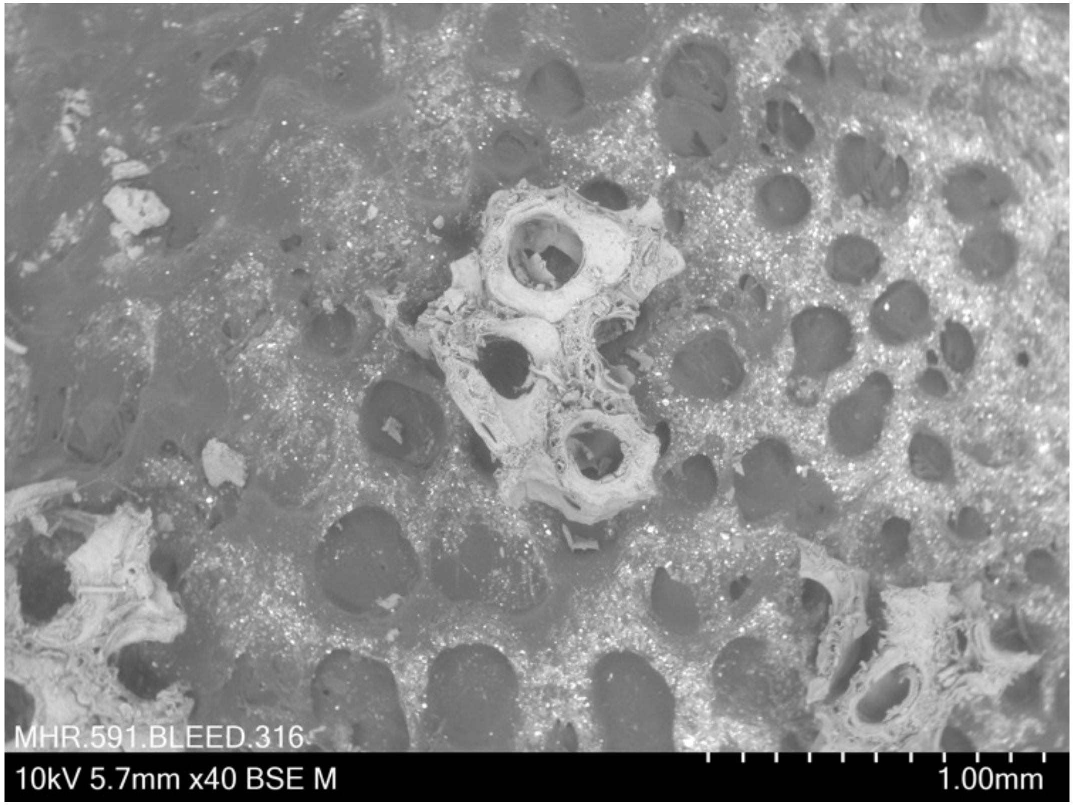

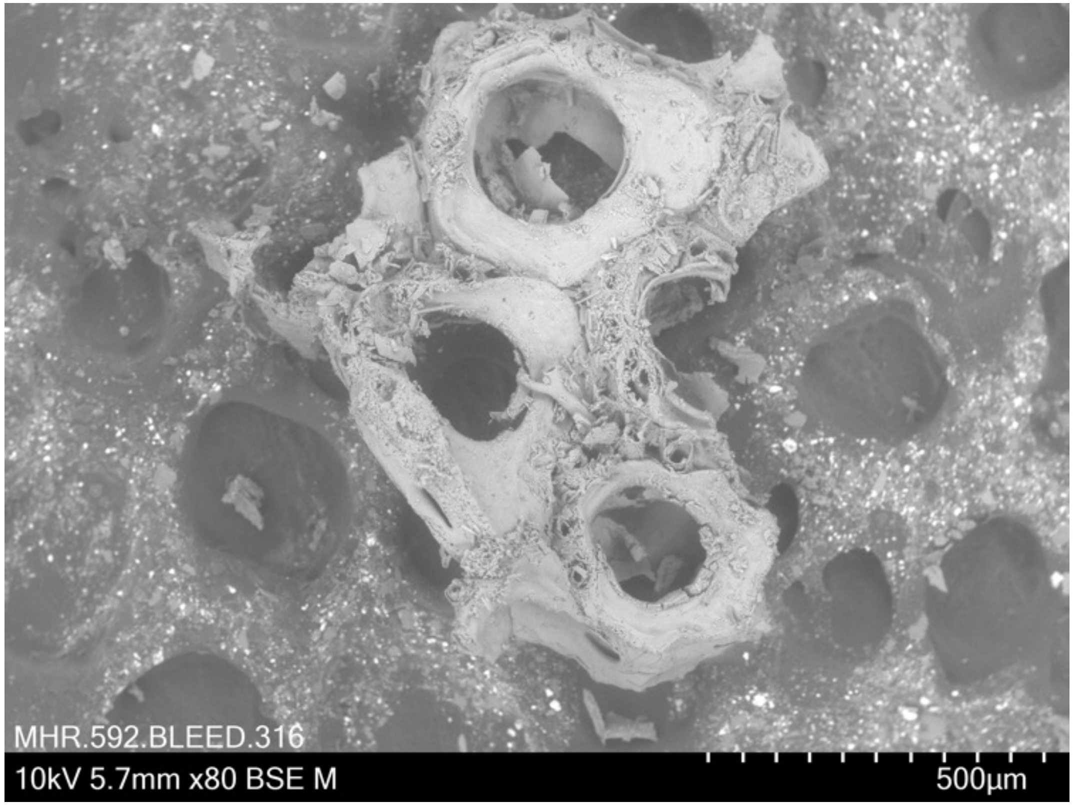


***Chaperiopsis* aff. *cervicornis* BLEED 316**
Coll: 2.2015 (D. P. Gordon)
Lat: -41.2833333333
Long: 174.76666666669999
Depth: intertidal

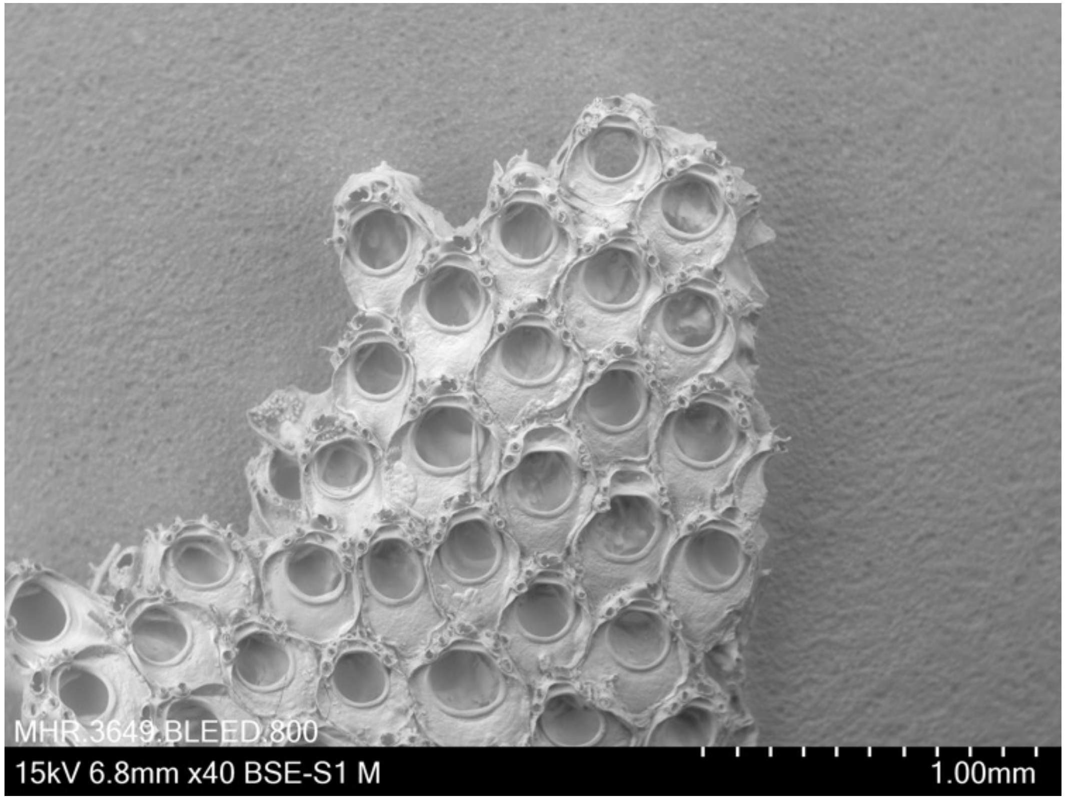


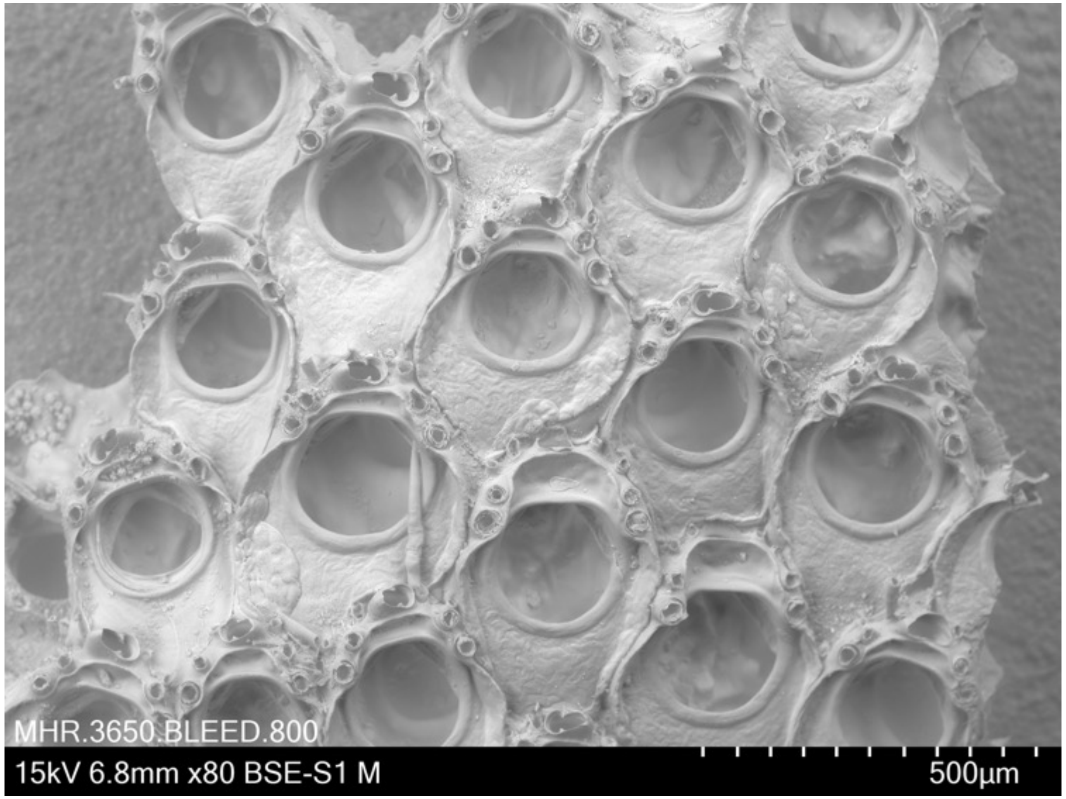


***Chaperiopsis cervicornis* BLEED 800**
Coll: 4.3.2018 (D. P. Gordon, L. H. Liow and C. Schack)
Lat: -39.639352
Long: 177.008705
Depth: intertidal

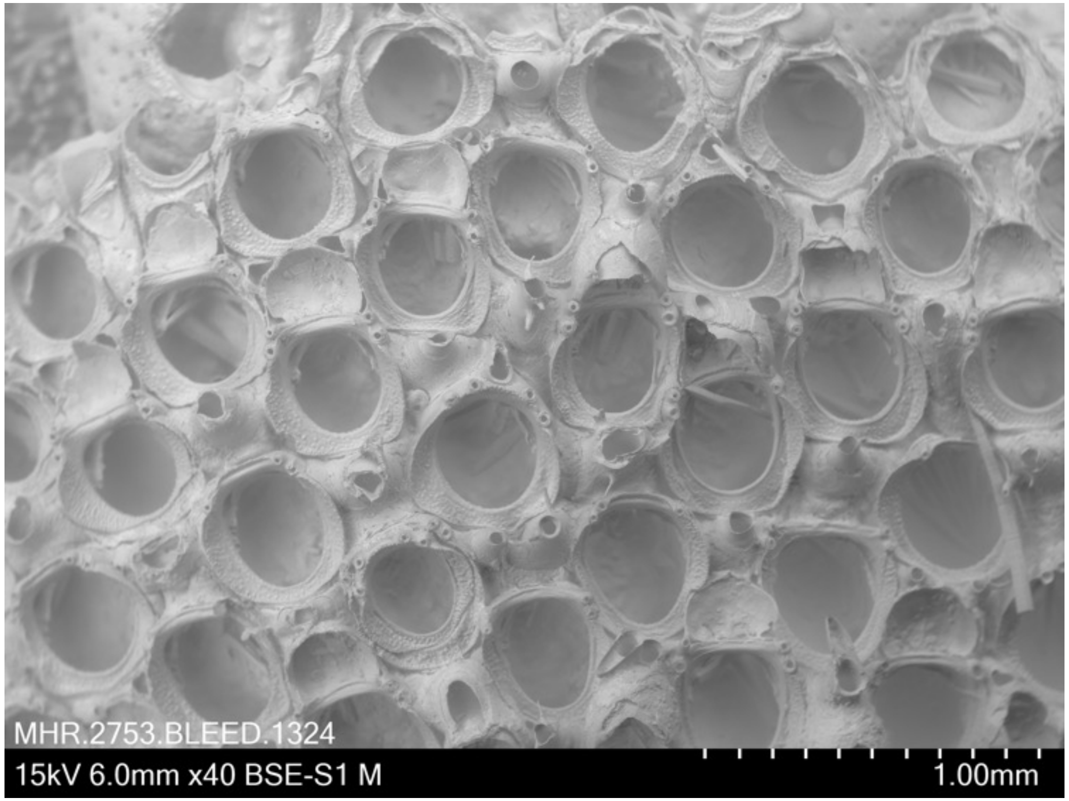


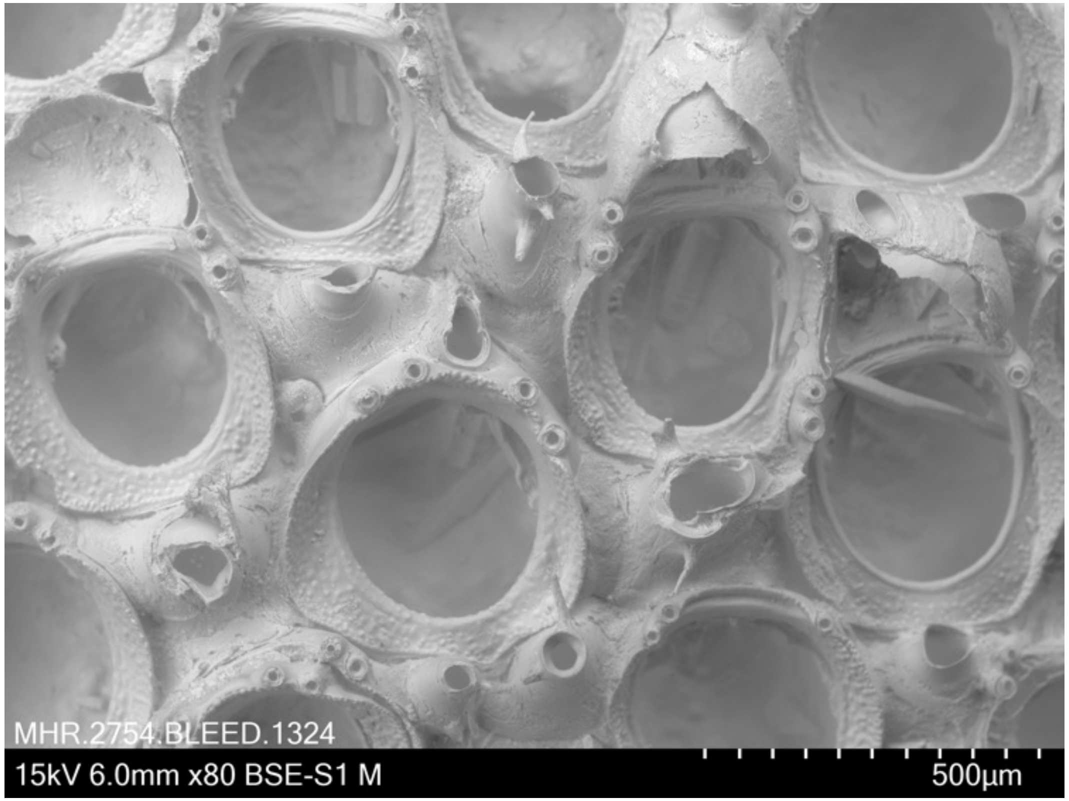


***Chaperiopsis* cf. *rubida* BLEED 1324**
Coll: 15.3.2019 (A. M. Smith; D. P. Gordon)
Lat: -36.46
Long: 174.9
Depth: 37 m

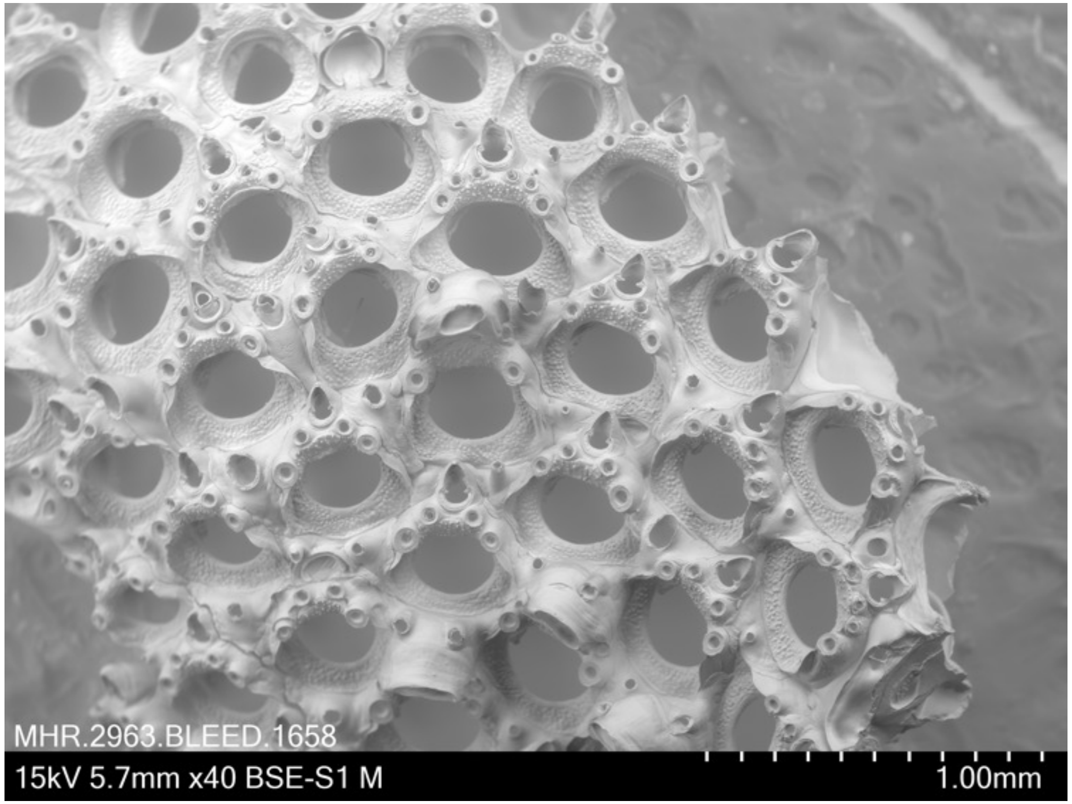


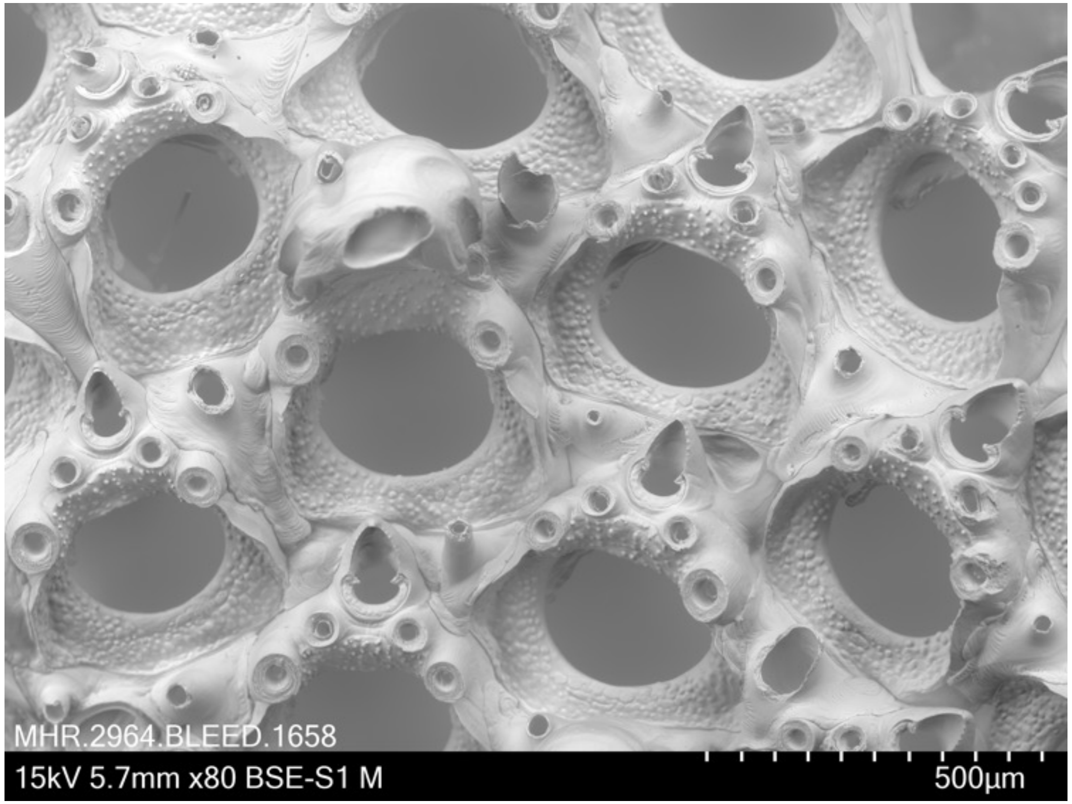


***Chaperiopsis* aff. *rubida* BLEED 1658**
Coll: 17.3.2019 (A. M.  Smith, D. P. Gordon)
Lat: -34.159999999999997
Long: 172.07
Depth: 98.9 m

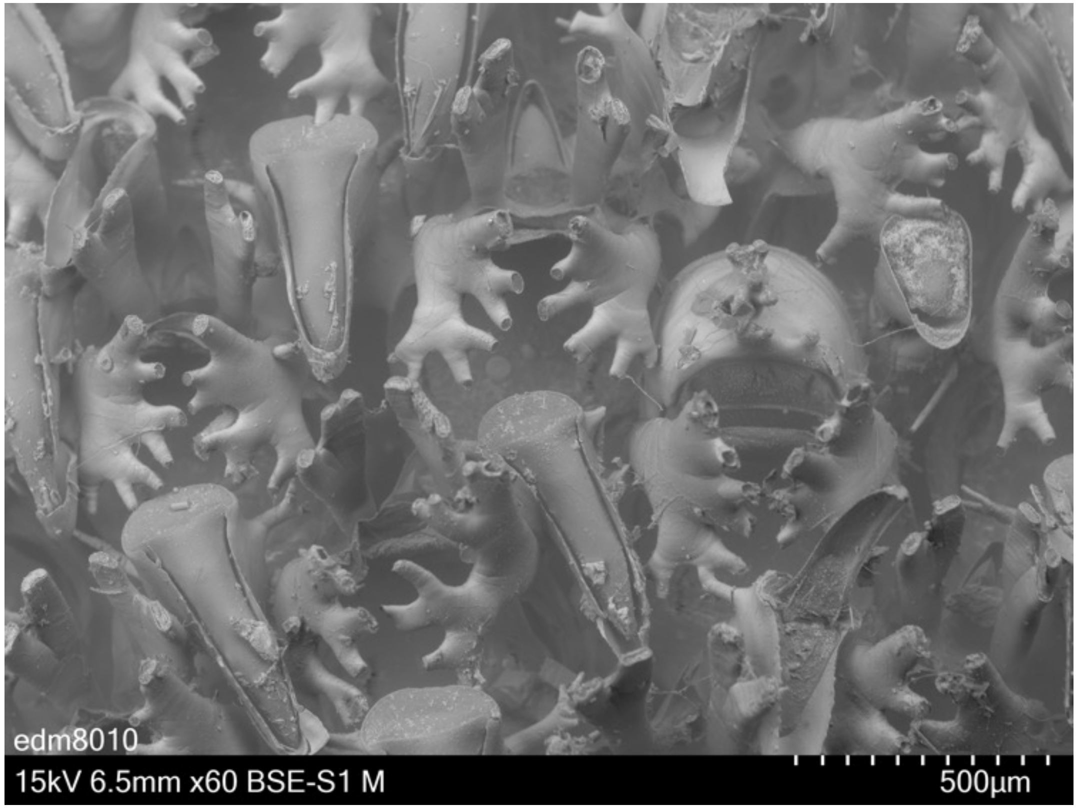


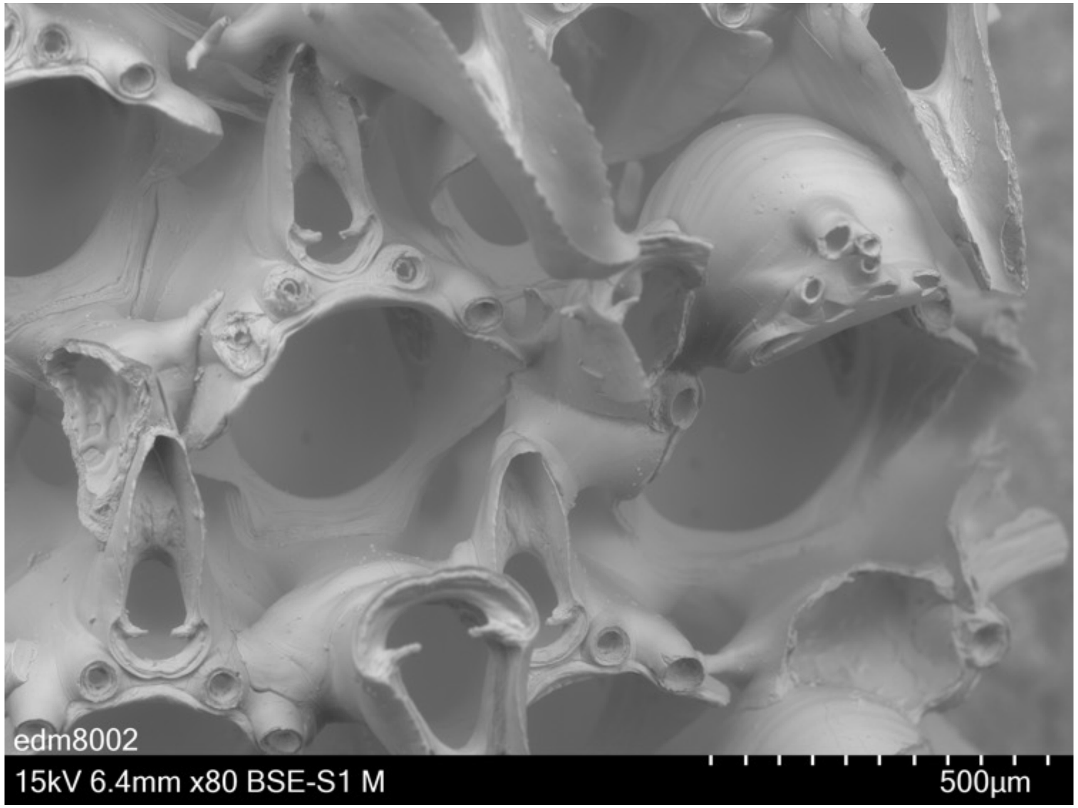


***Chaperiopsis* n. sp. BLEED 1616**
Coll: 17.3.2019 (A. M. Smith, D. P. Gordon)
Lat: -34.159999999999997
Long: 172.09
Depth: 82.9 m

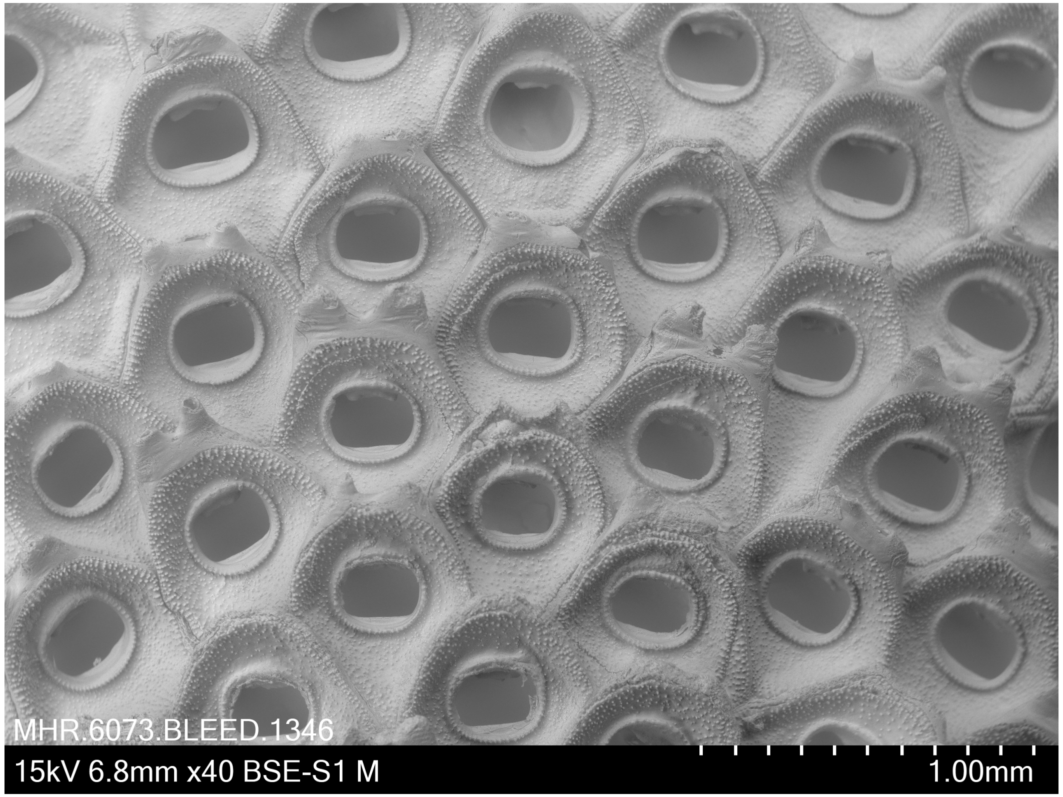


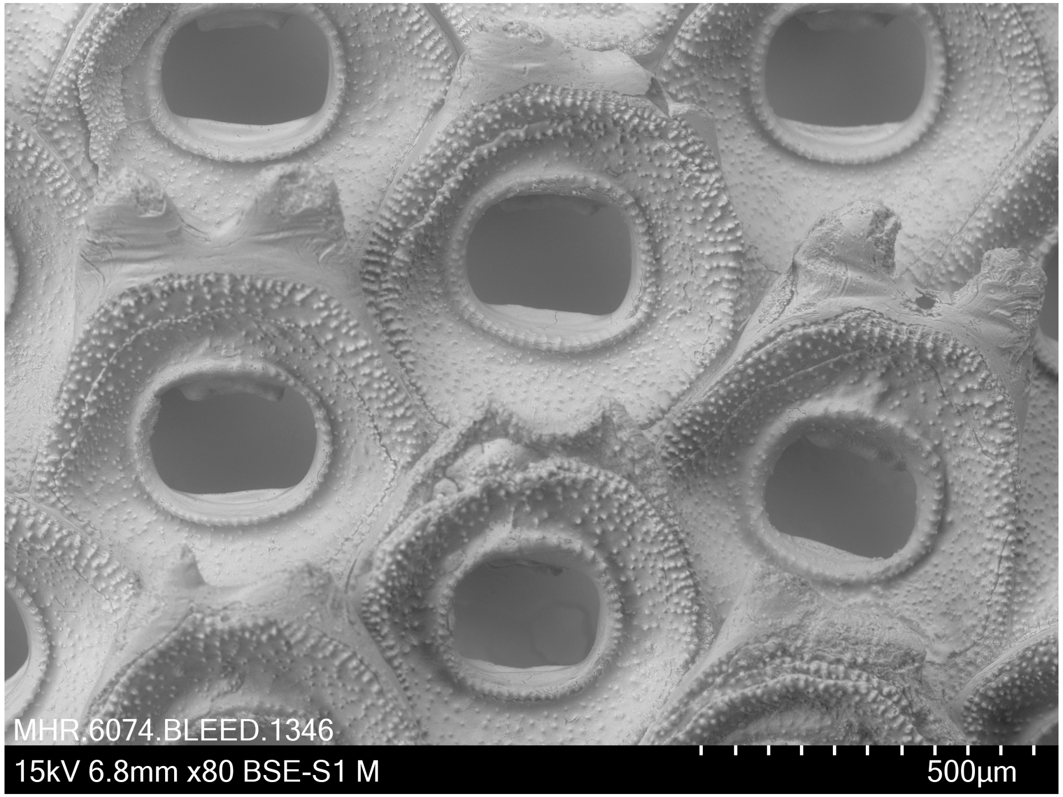


***Patsyella* n. sp. BLEED 1346**
Coll: 17.3.2019 (A. M. Smith, D. P. Gordon)
Lat: -34.21
Long: 172.11
Depth: 104m

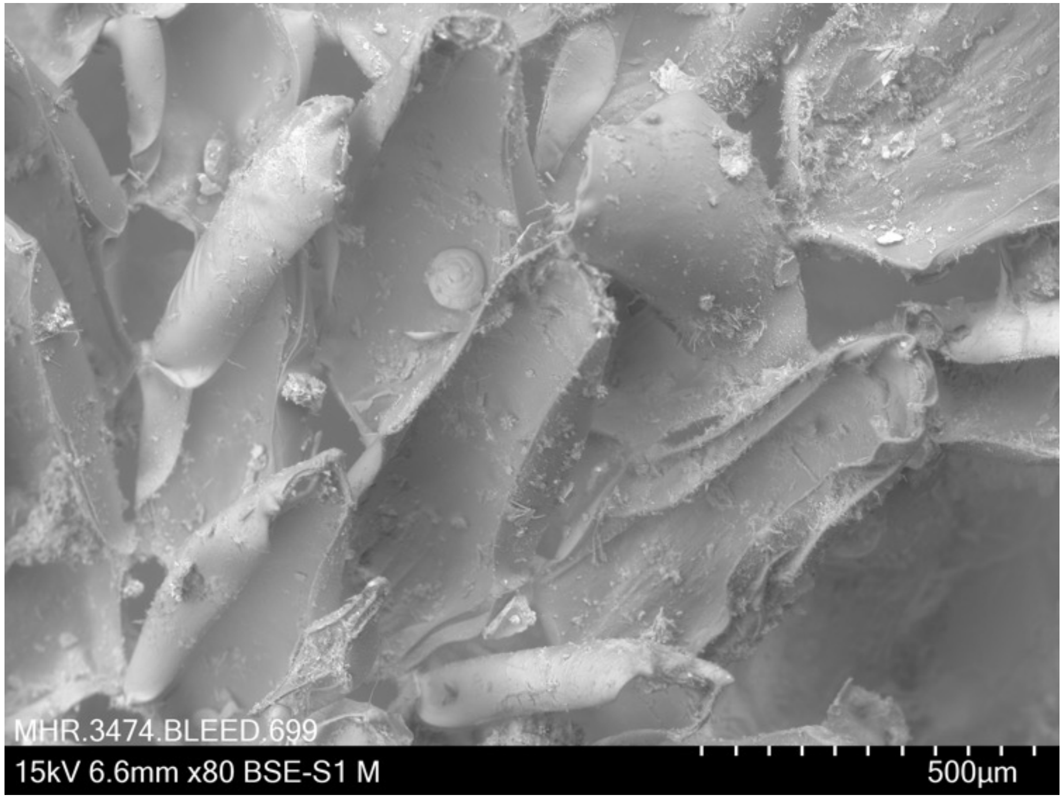


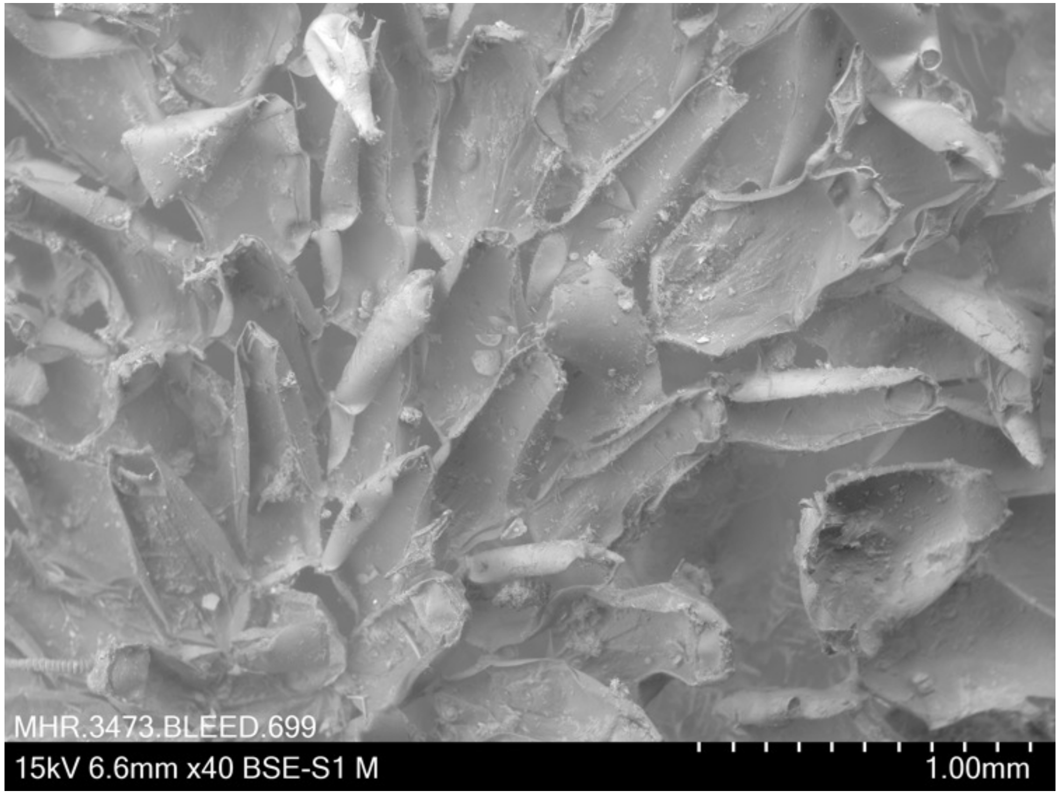


***Beania bilaminata* BLEED 699**
Coll: 8.4.2018 (A. M. Smith, D. P. Gordon, H. Mello)
Lat: -47.135833
Long: 168.01055
Depth: 72 m

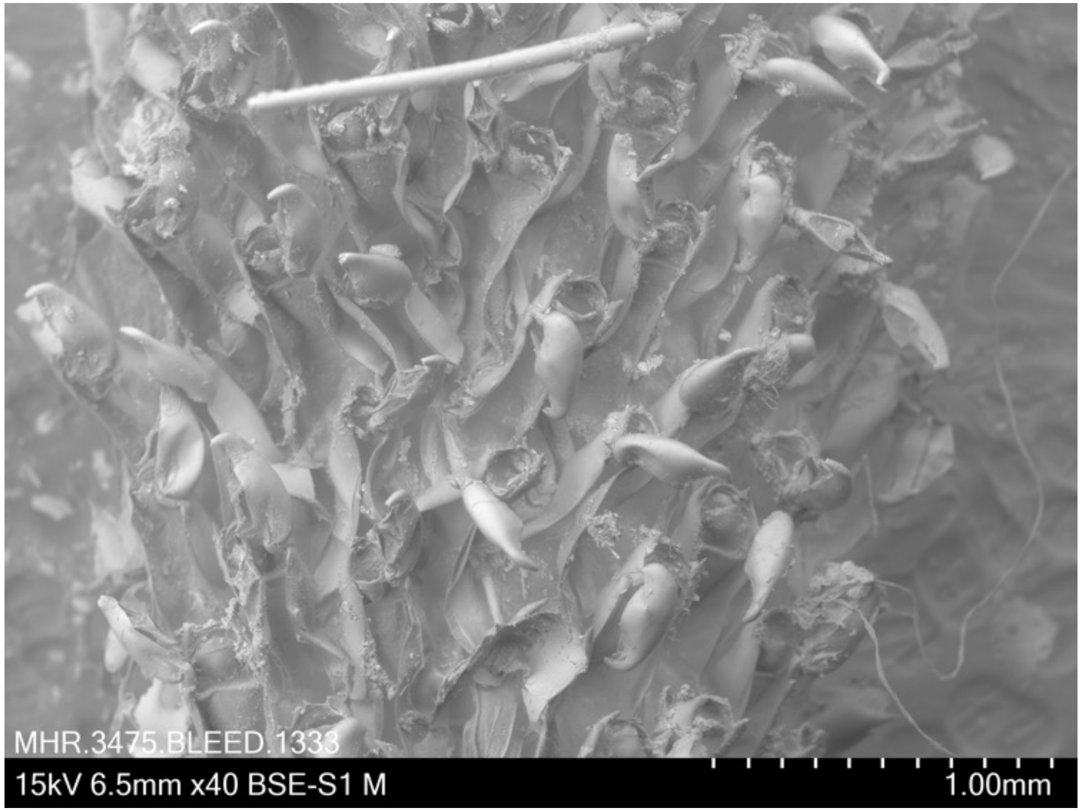

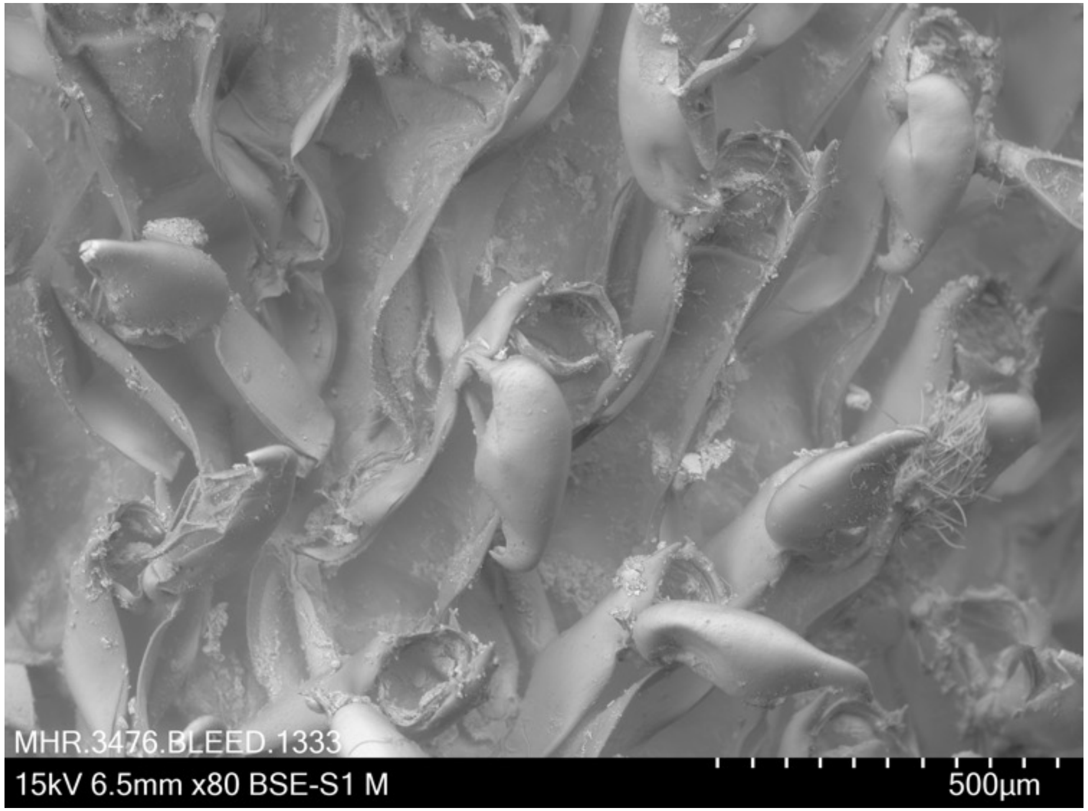


***Beania* sp. BLEED 1333**
Coll: 19.3.2019 (A. M. Smith, D. P. Gordon)
Lat: -34.99
Long: 173.97
Depth: 61.6 m

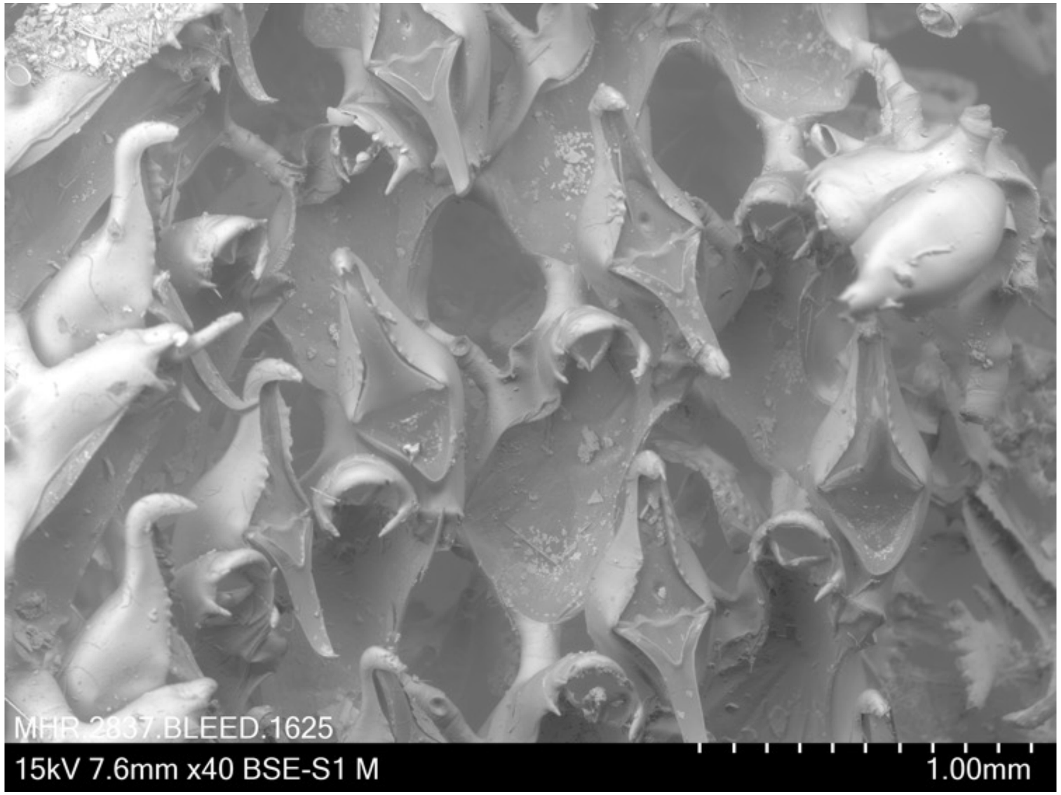


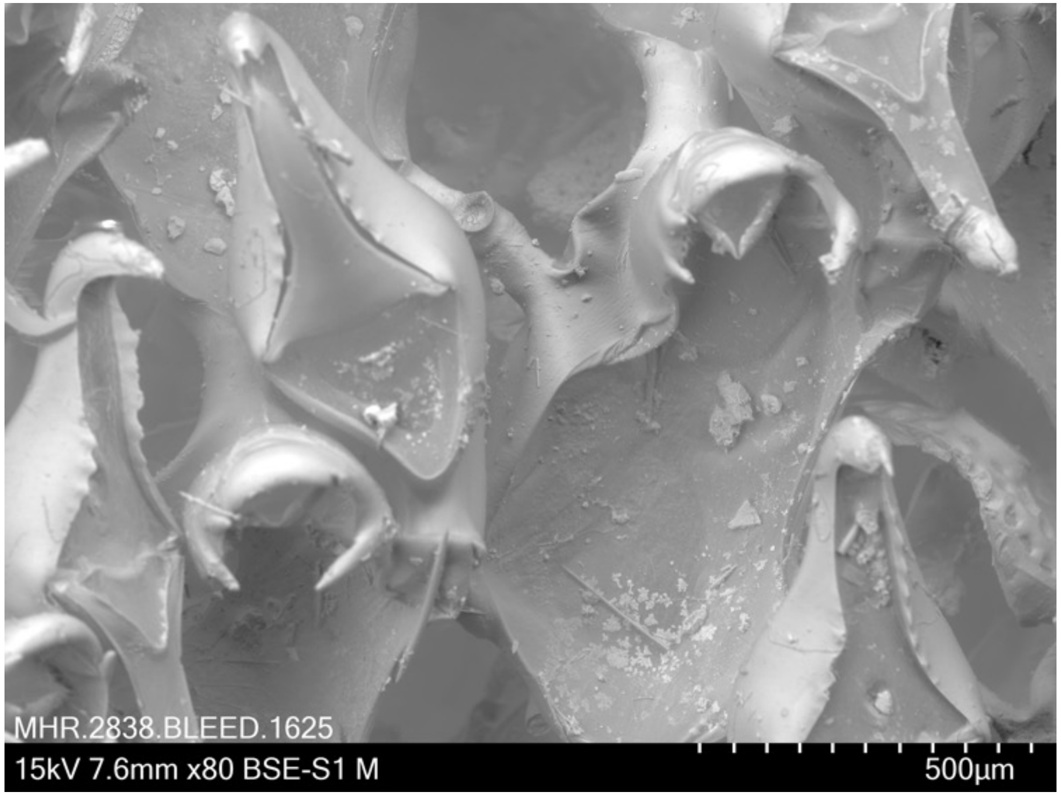


***Beania serrata* BLEED 1625**
Coll: 17.3.2019 (A. M. Smith, D. P. Gordon)
Lat: -34.159999999999997
Long: 172.09
Depth: 82.9 m

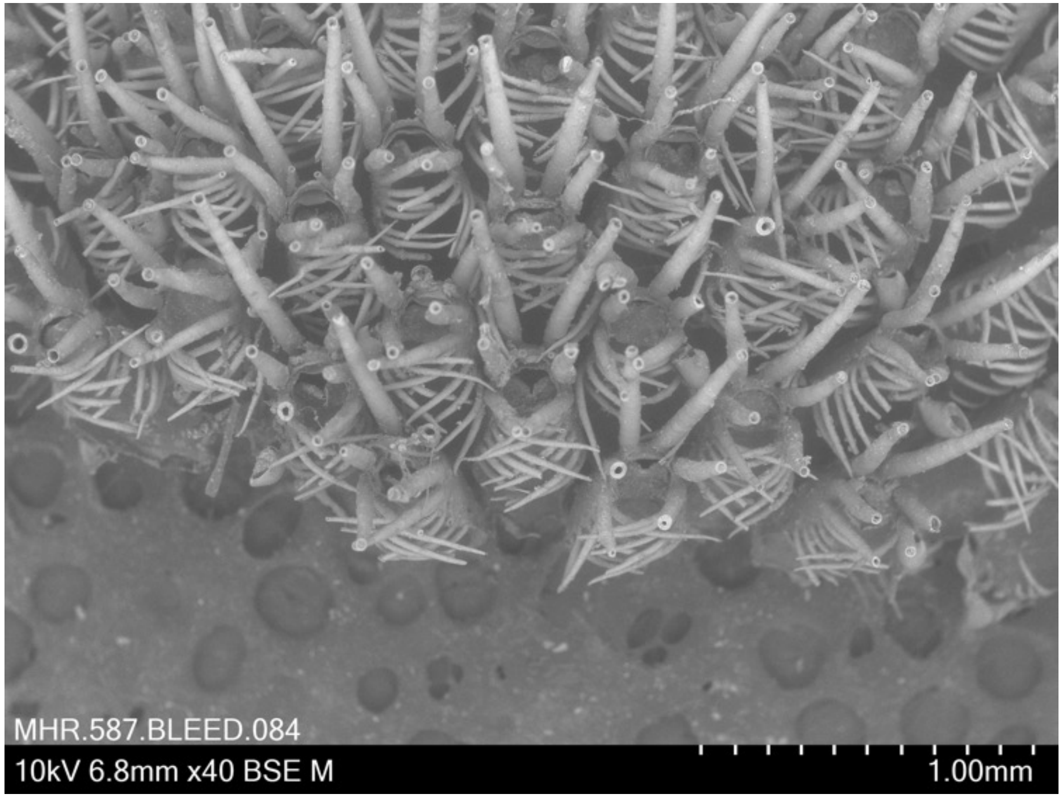


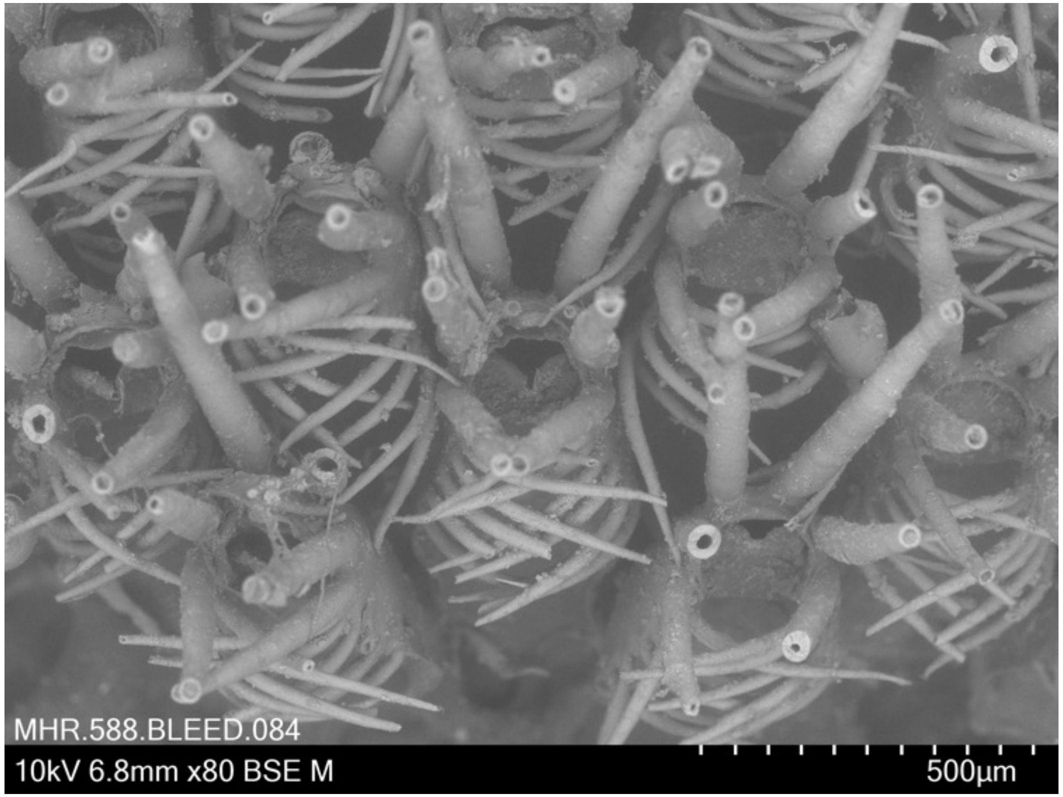


***Beania stonycha*** **BLEED 84**
NIWA 69631 (TRIP3298/36)
Coll: 12.04.2011
Lat: -46.283332999999999
Long: 166.133333
Depth: 270 m

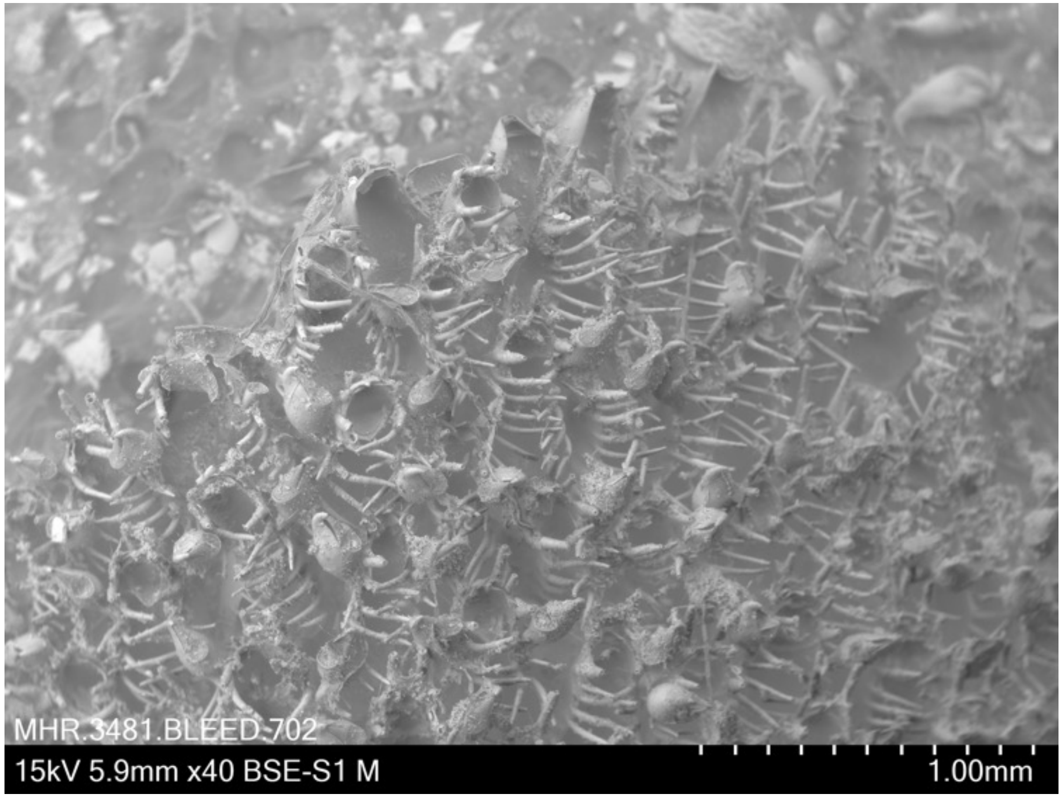


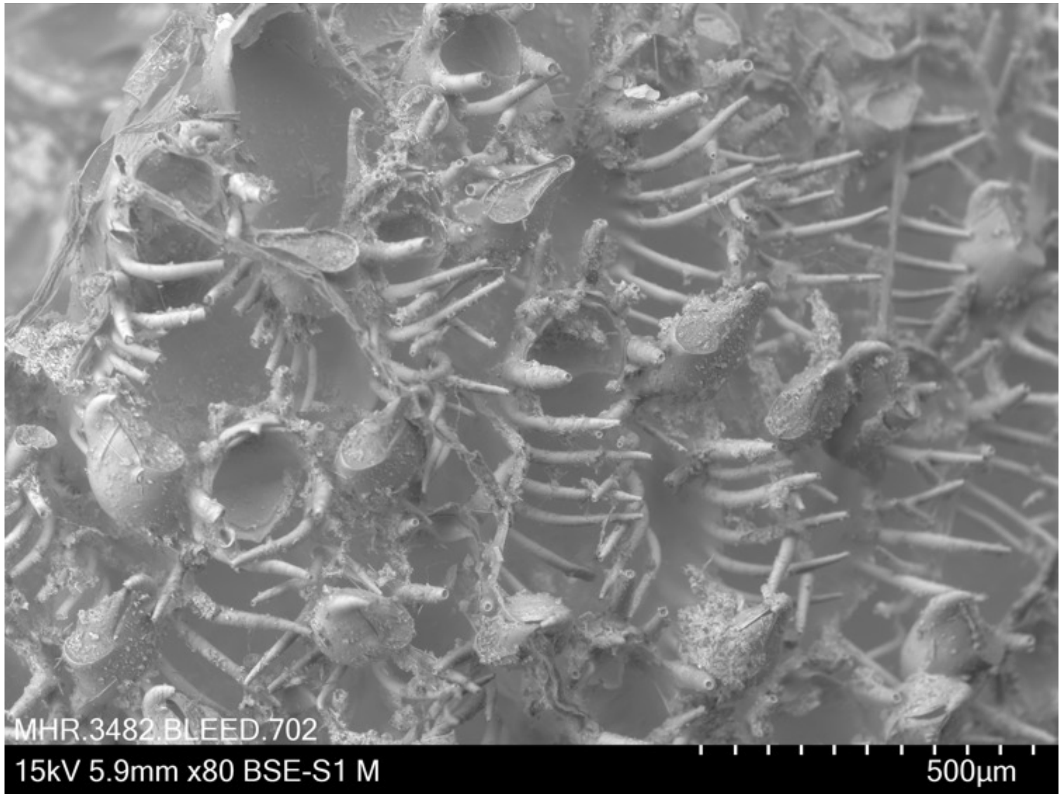


***Beania elongata* BLEED 702**
Coll: 9.4.2018 (A. M. Smith, D. P. Gordon, H. Mello)
Lat: -47.21666667
Long: 167.6
Depth: intertidal

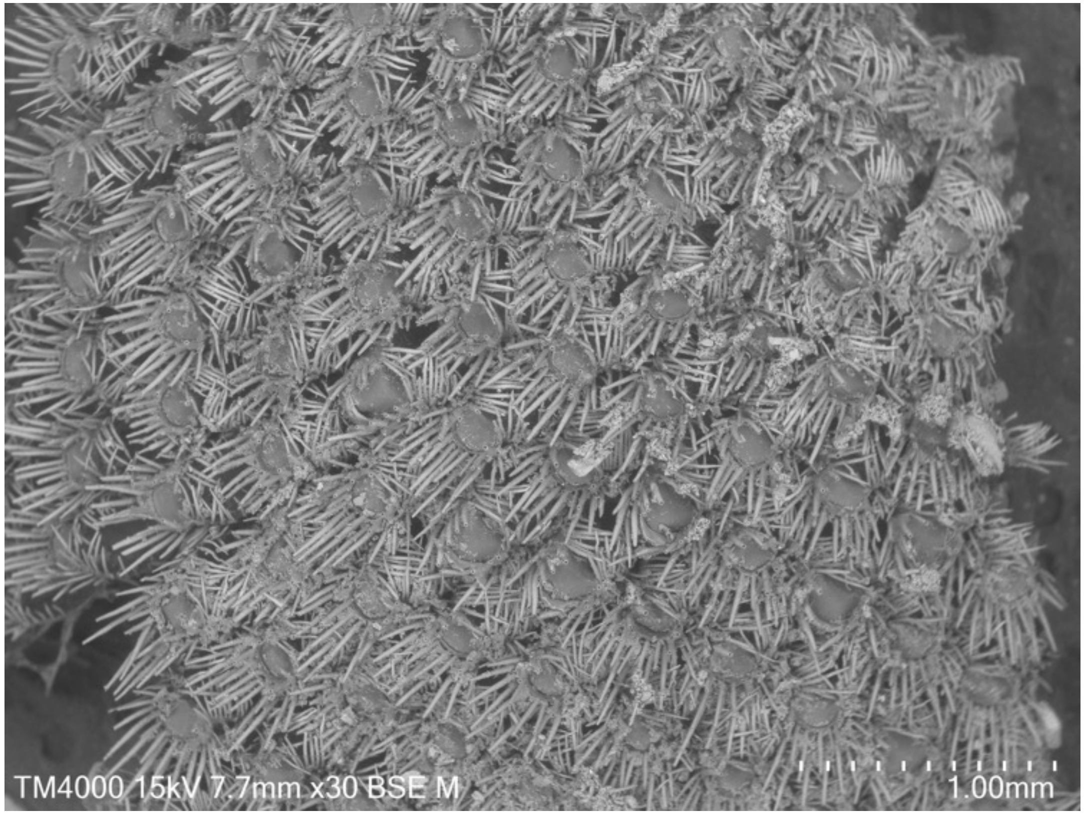


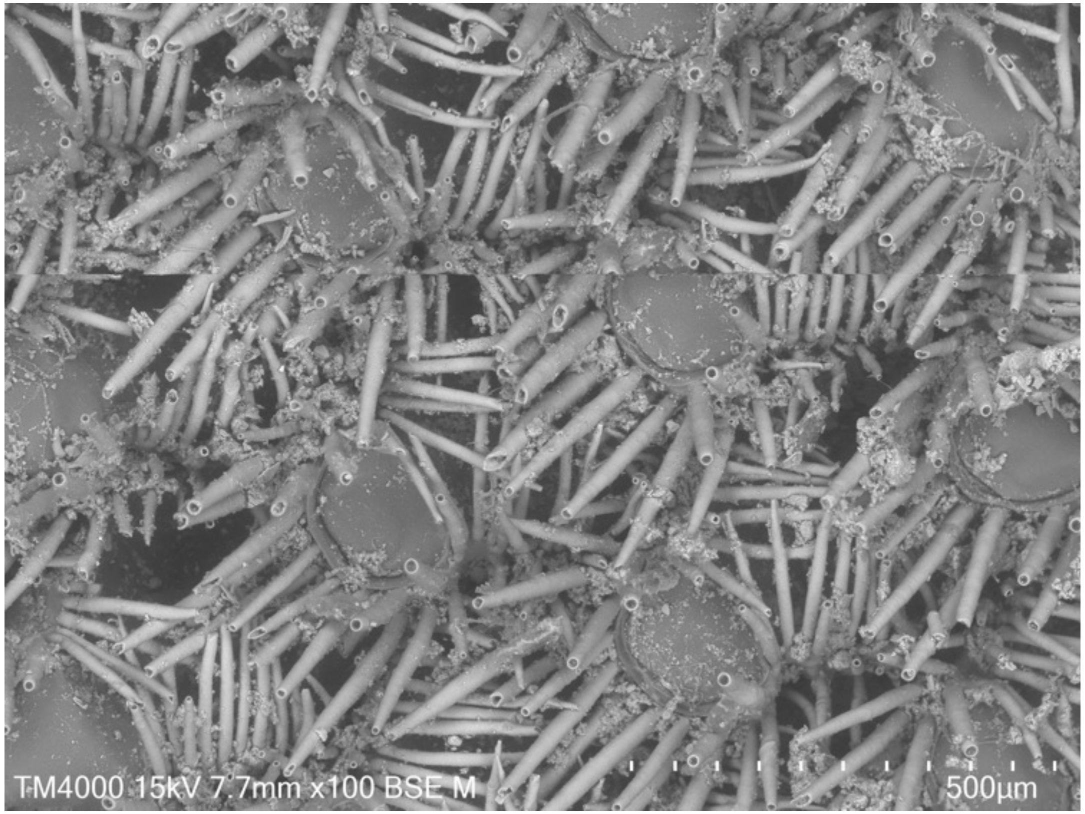


***Beania plurispinosa* BLEED 203**
Coll: 20.5.2015 (D. P. Gordon)
Lat: -41.071039
Long: 176.088894
Depth: intertidal

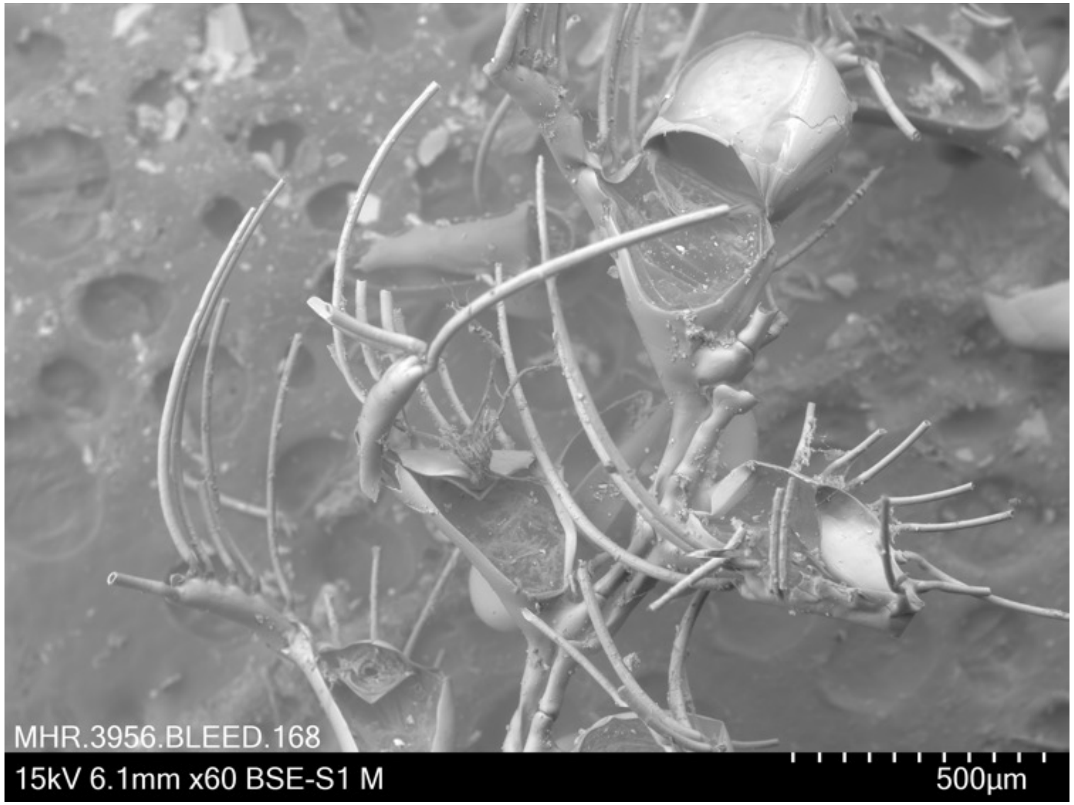


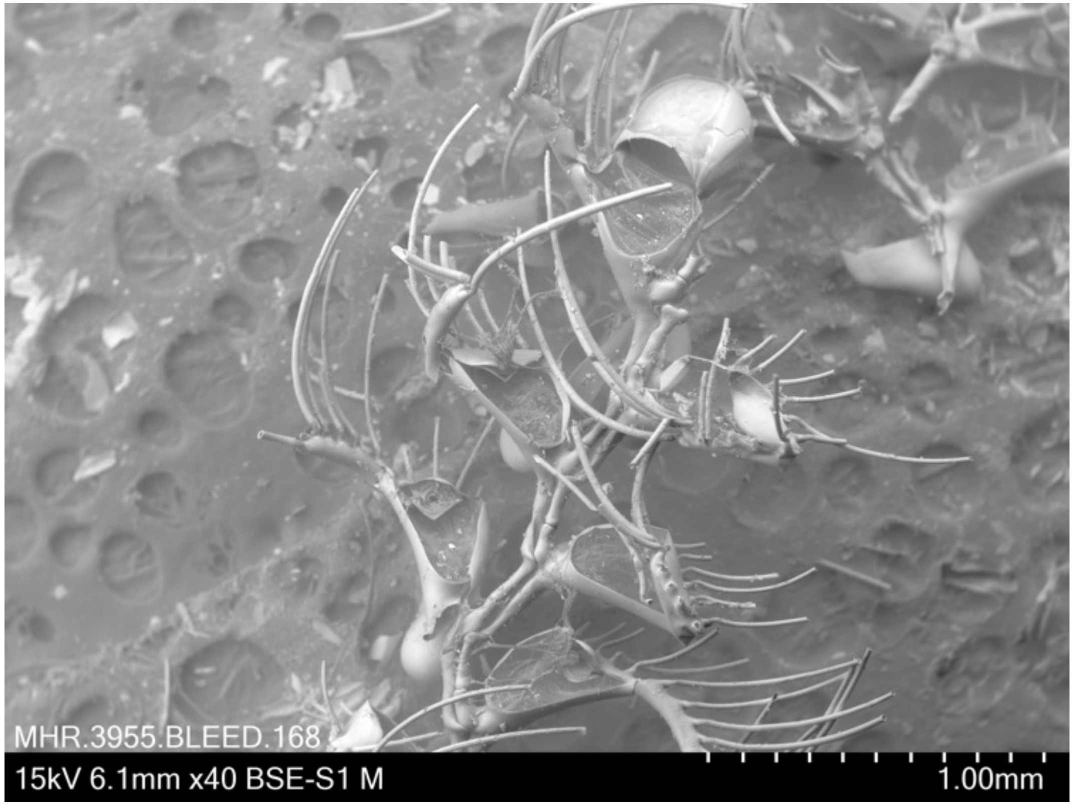


***Cornucopina* n. sp. BLEED 168**
NIWA 95862 (TAN1313/DR3)
Coll: 28.11.2013
Lat: -34.7698333
Long: 177.7121667
Depth: 1420 m

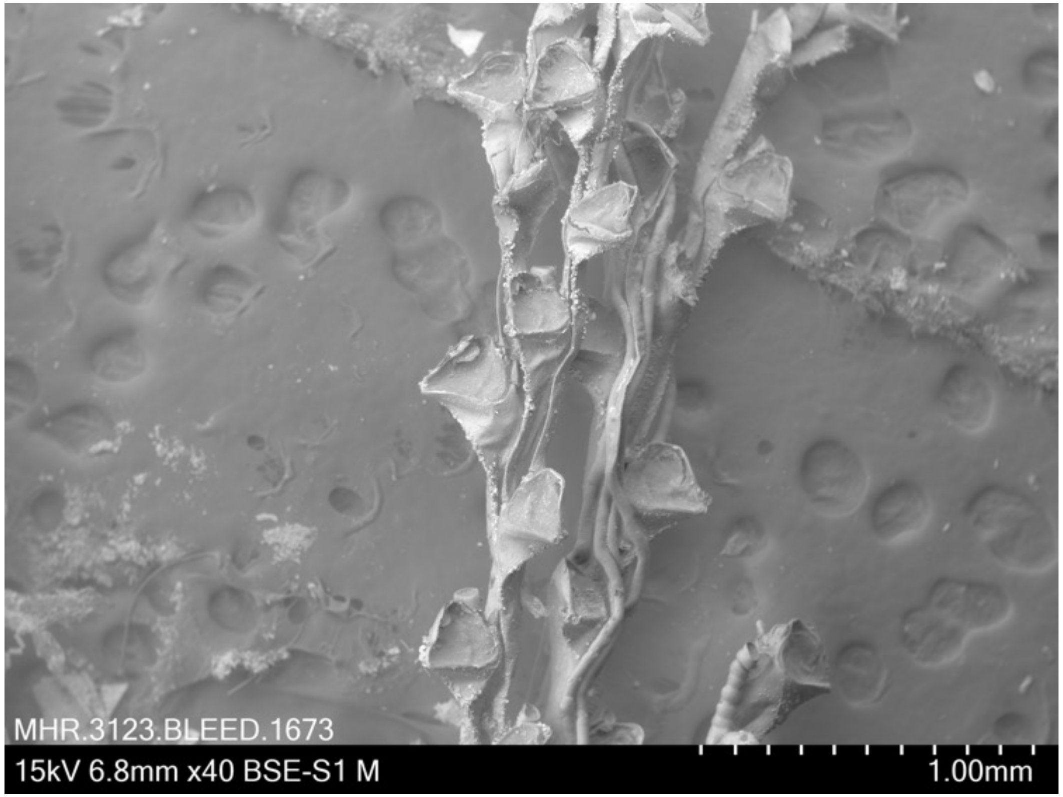


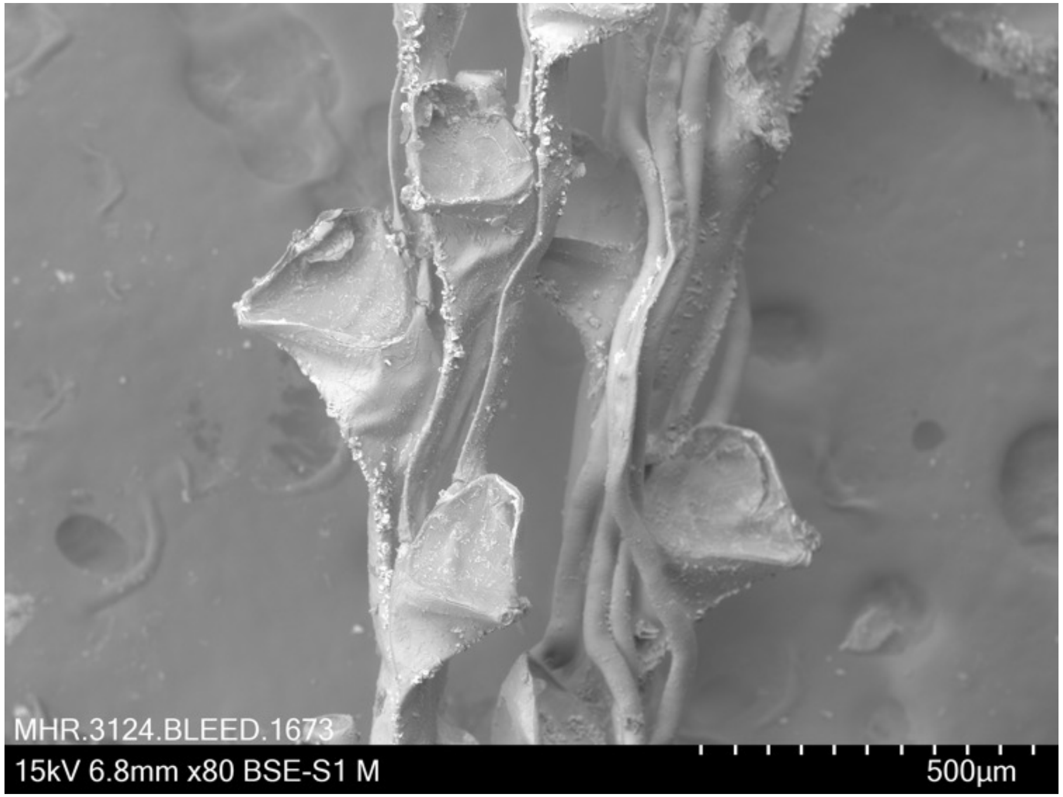


***Cornucopina* sp. BLEED 1673**
Coll: 18.3.2019 (A. M. Smith, D. P. Gordon)
Lat: -34.380000000000003
Long: 172.69
Depth: 52.5-56.5m

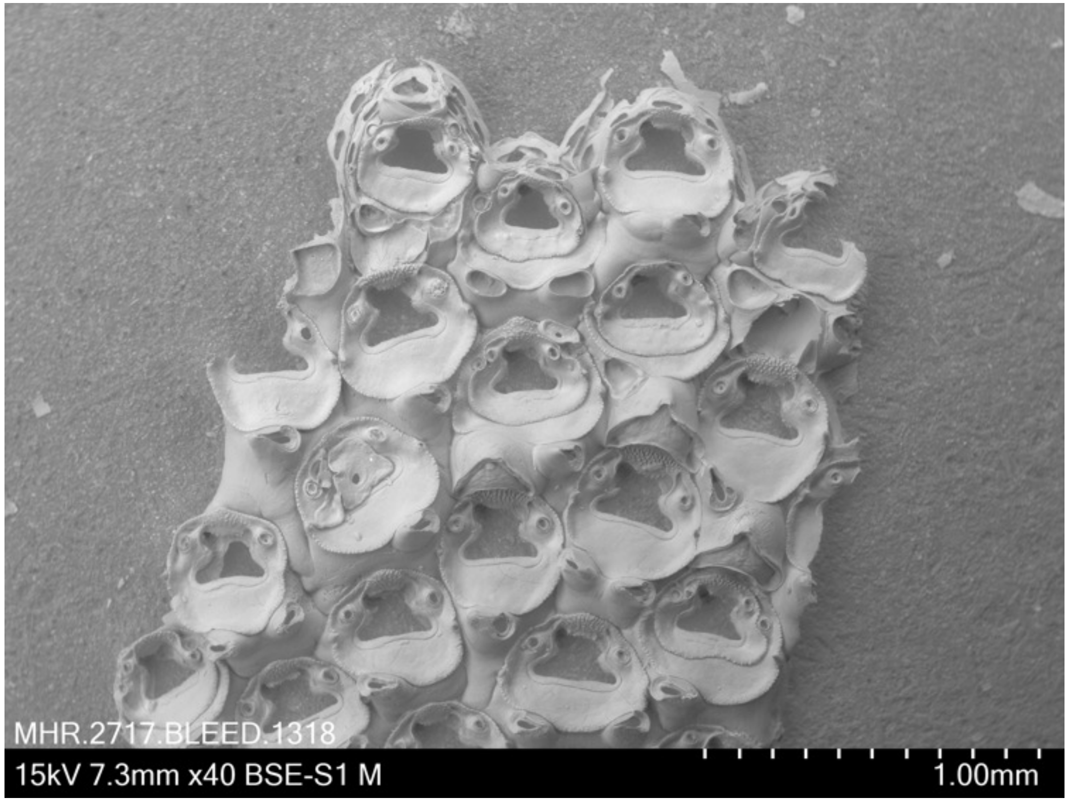


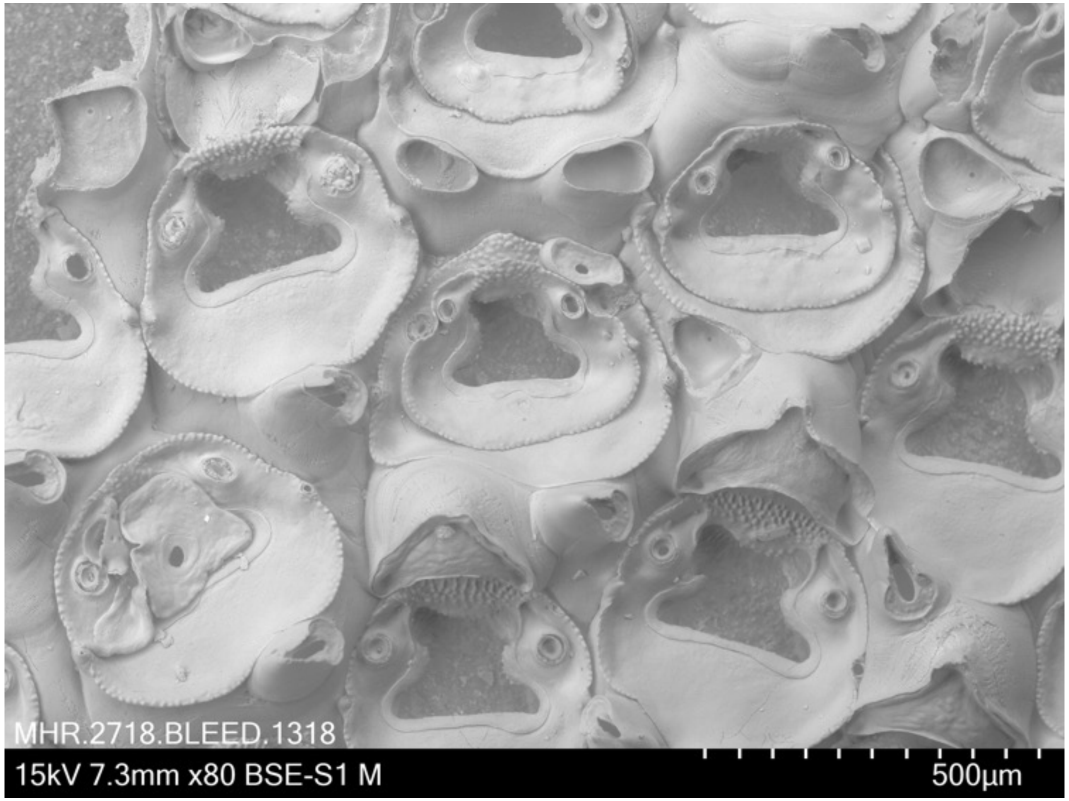


***Amphiblestrum contentum* BLEED 1318**
Coll: 15.3.2019 (A. M. Smith, D. P. Gordon)
Lat: -36.46
Long: 174.9
Depth: 37 m

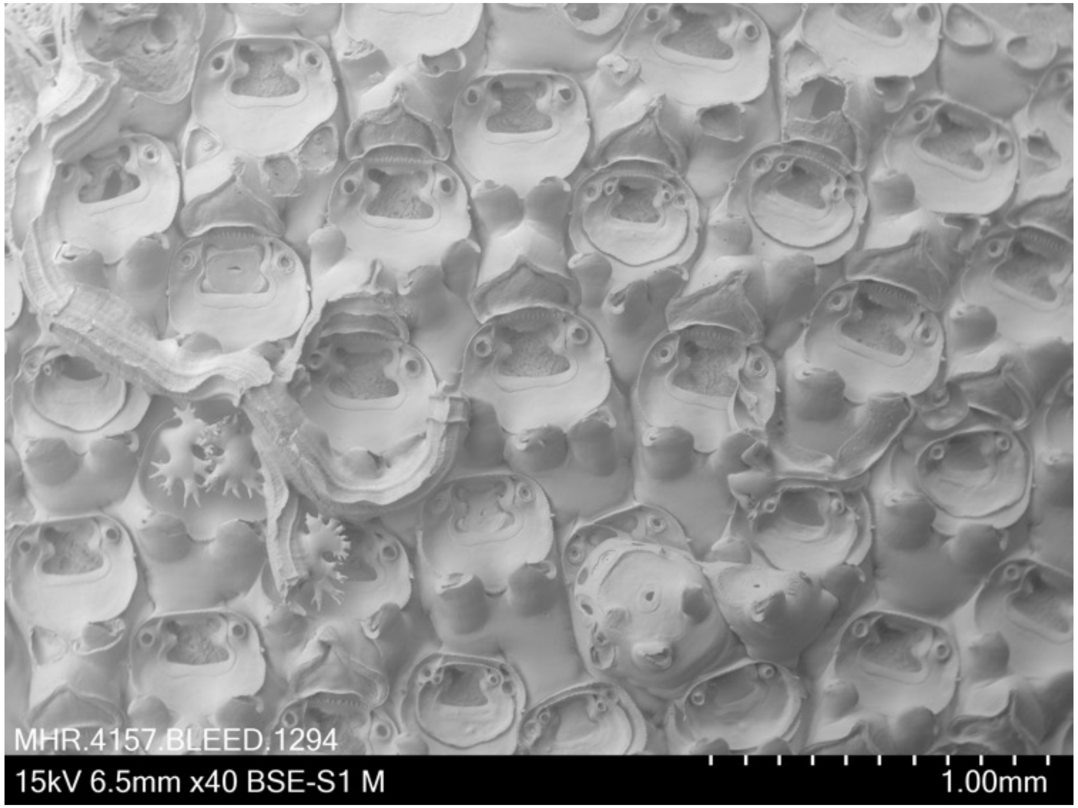


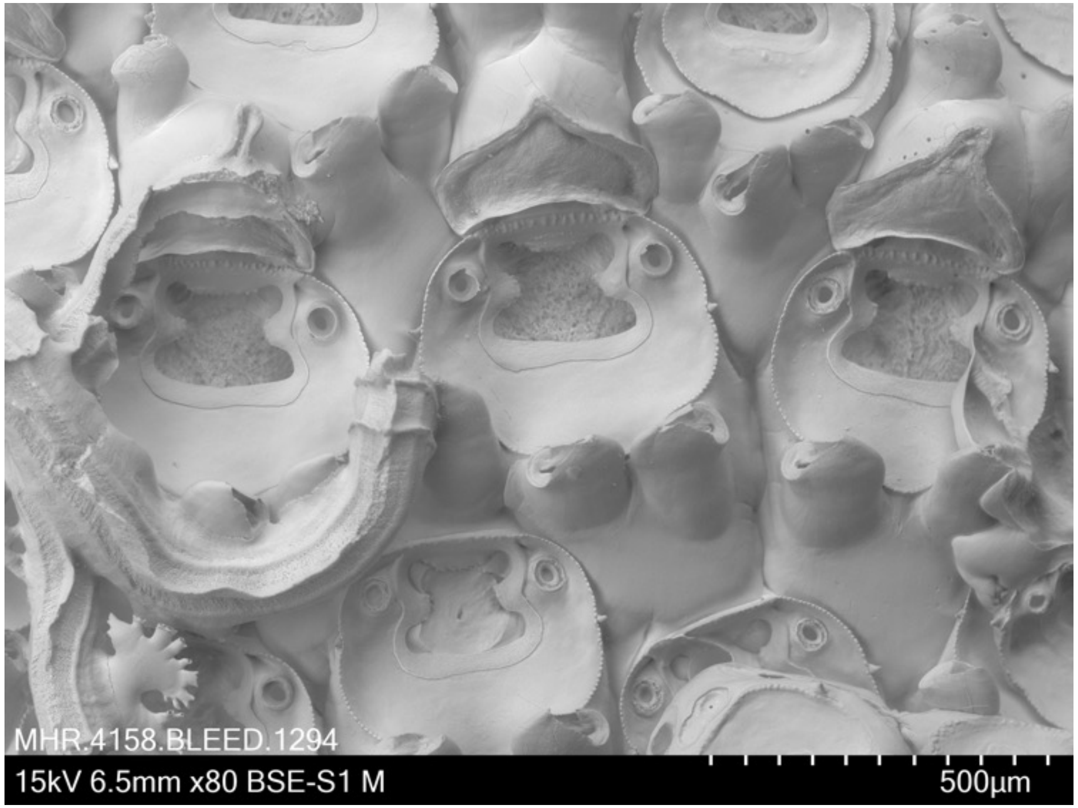


***Amphiblestrum* n. sp. 1 BLEED 1294**
Coll: 17.3.2019 (A. M. Smith; D. P. Gordon)
Lat: -34.21
Long: 172.11
Depth: 104 m

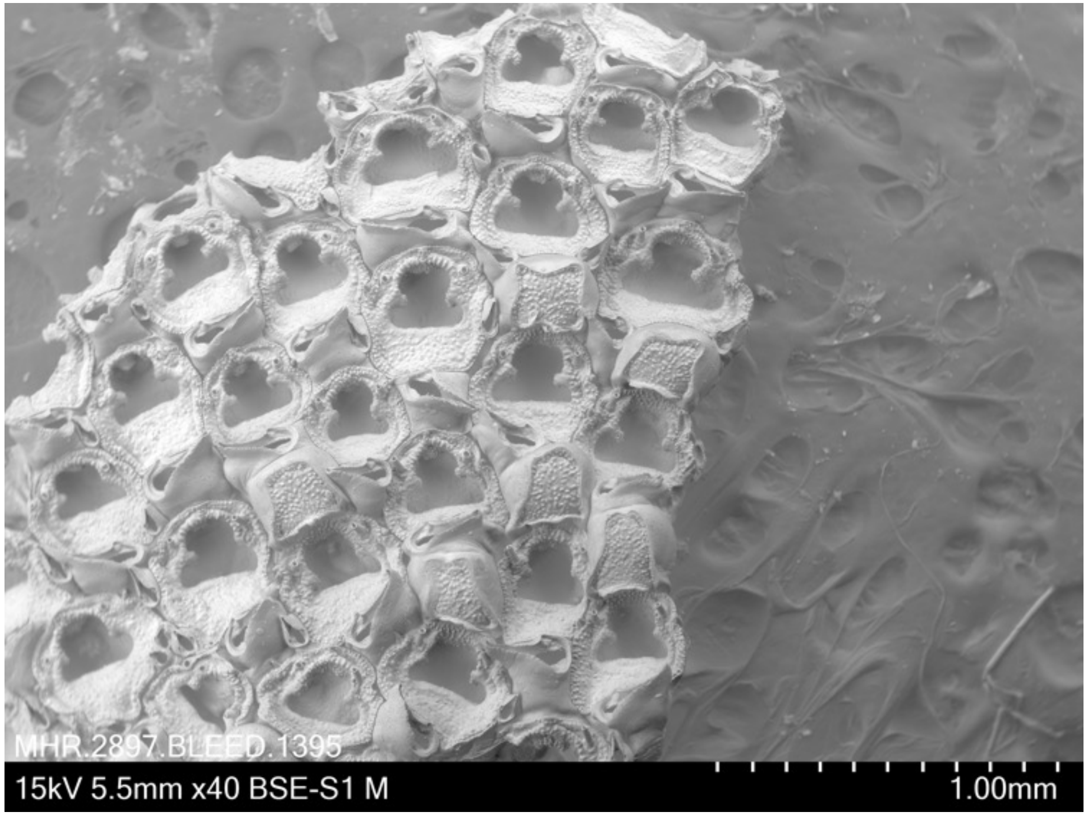


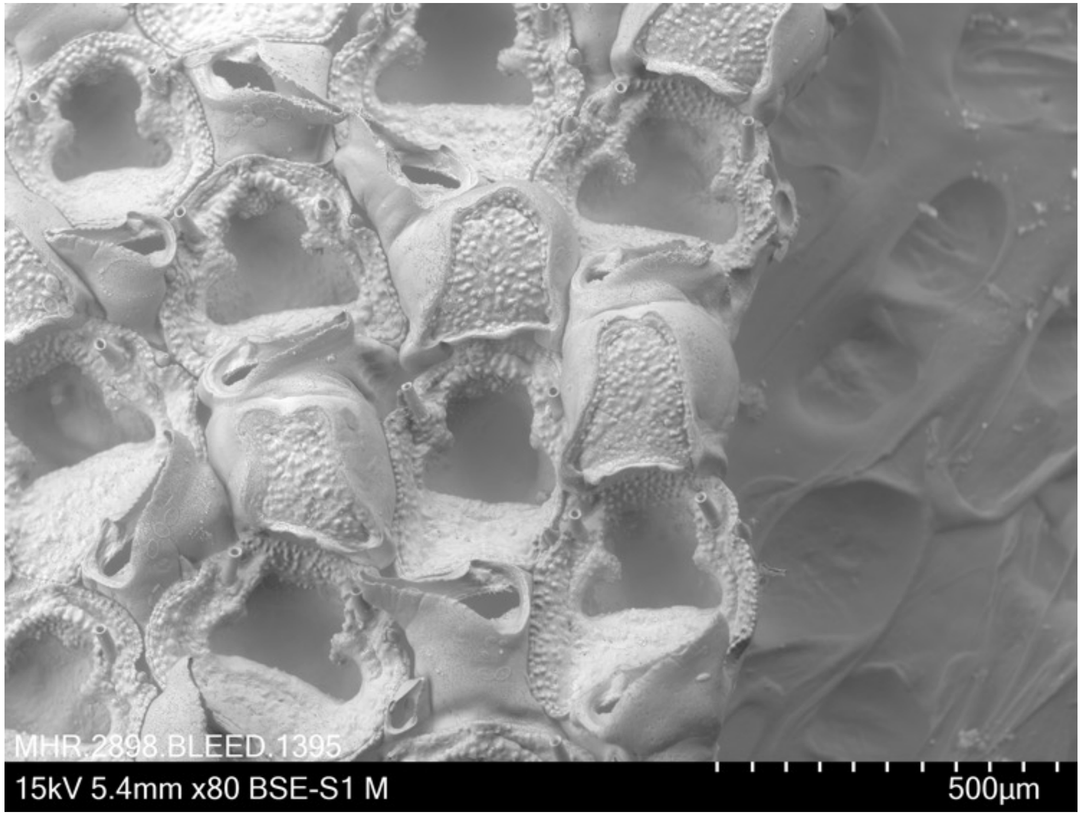


***Amphiblestrum* n. sp. 2 BLEED 1395**
Coll: 17.3.2019 (A. M. Smith, D. P. Gordon)
Lat: -34.159999999999997
Long: 172.09
Depth: 82.9 m

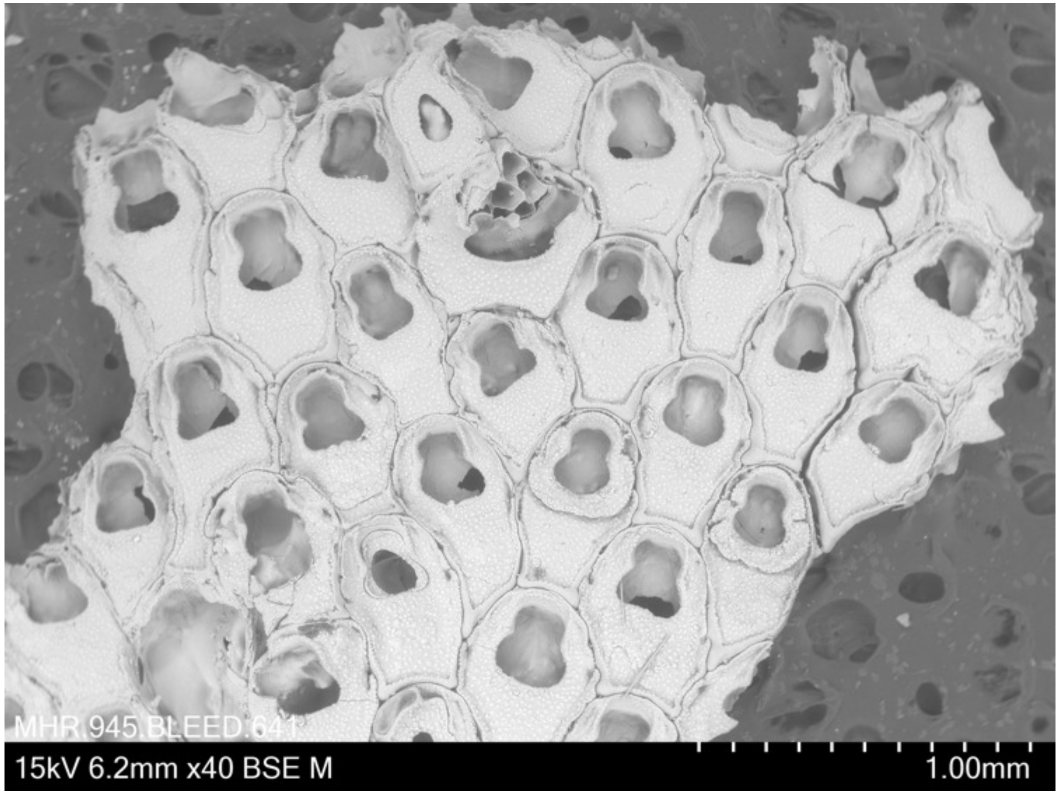


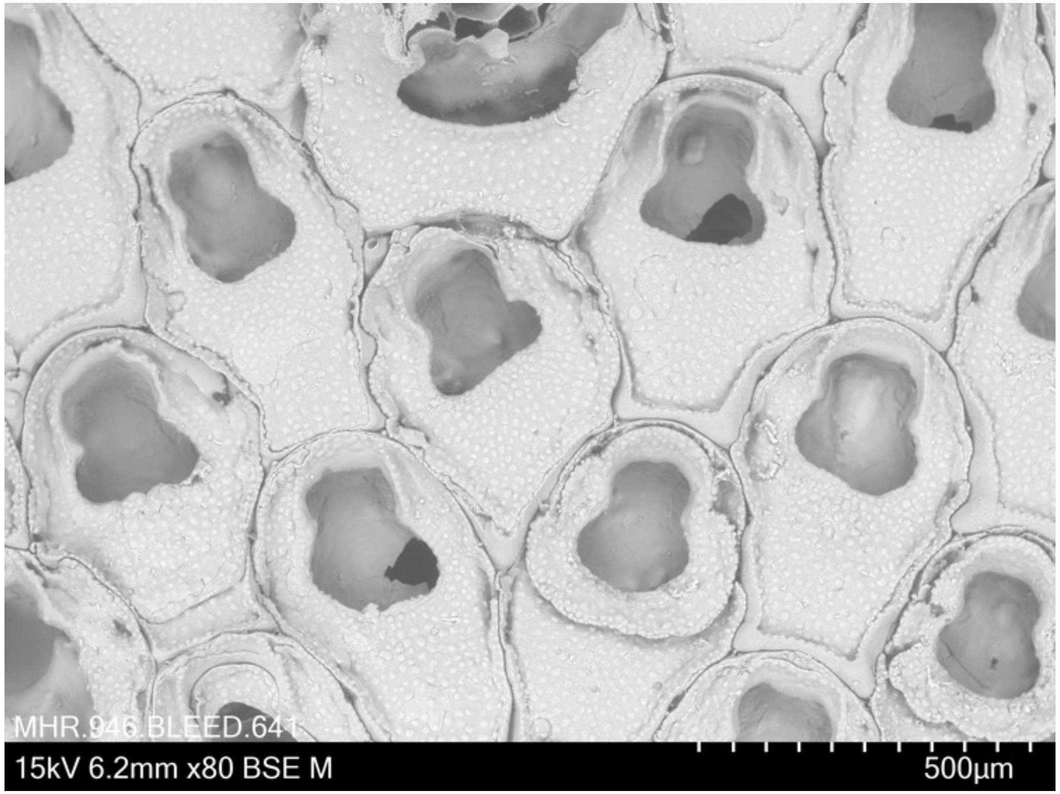


***Amphiblestrum blandum* BLEED 641**
Coll: 10.4.2018 (A. M. Smith, D. P. Gordon, H. Mello)
Lat: -46.924166666666665
Long: 168.2183
Depth: 58 m

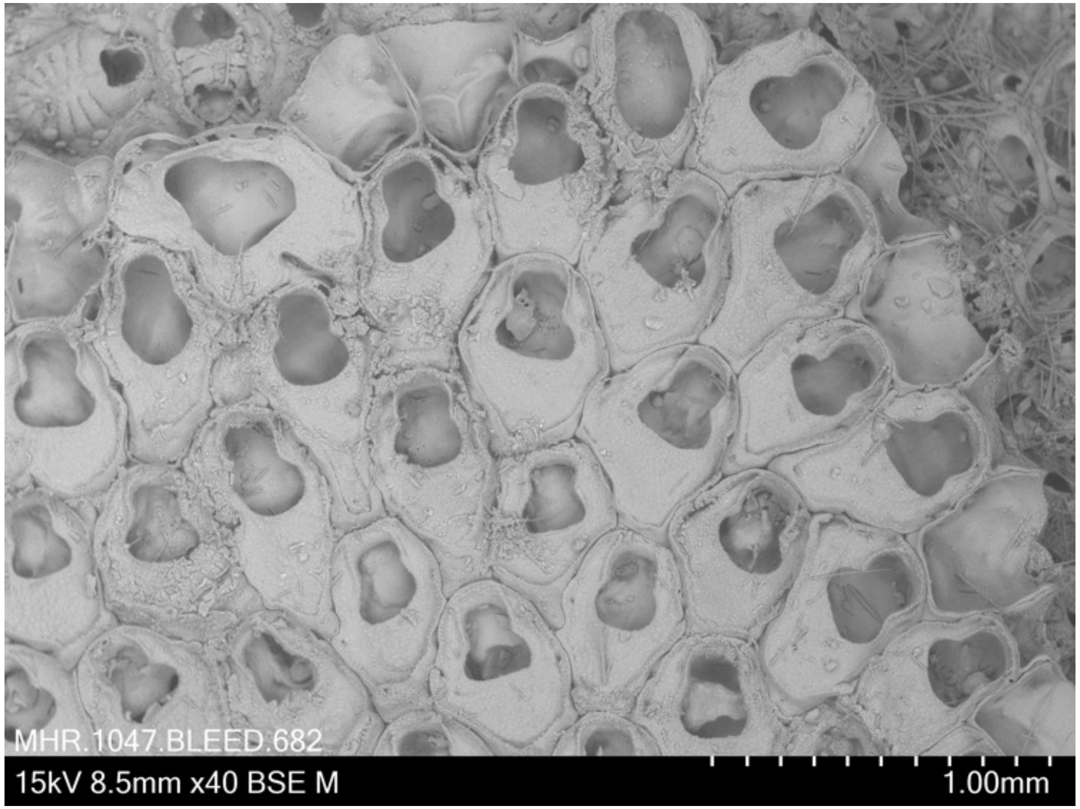

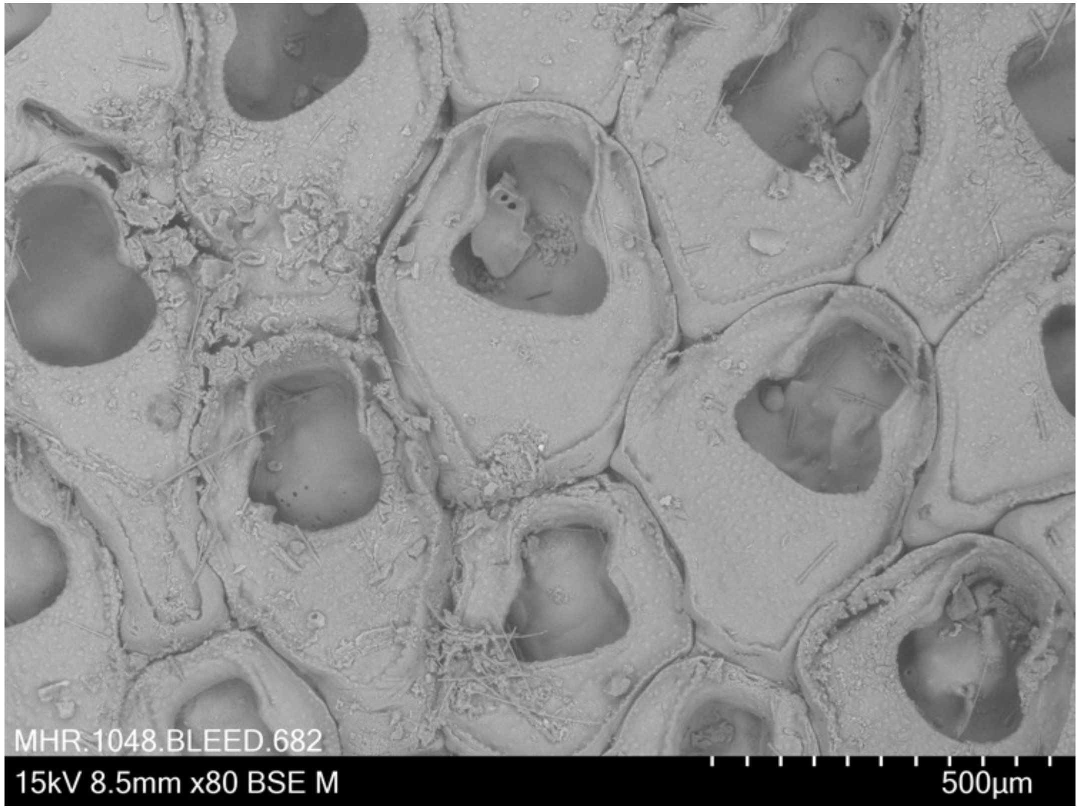


***Amphiblestrum blandum* BLEED 682**
Coll: 8.4.2018 (A. M. Smith, D. P. Gordon, H. Mello)
Lat: -47.136111111
Long: 168.188611
Depth: 77 m

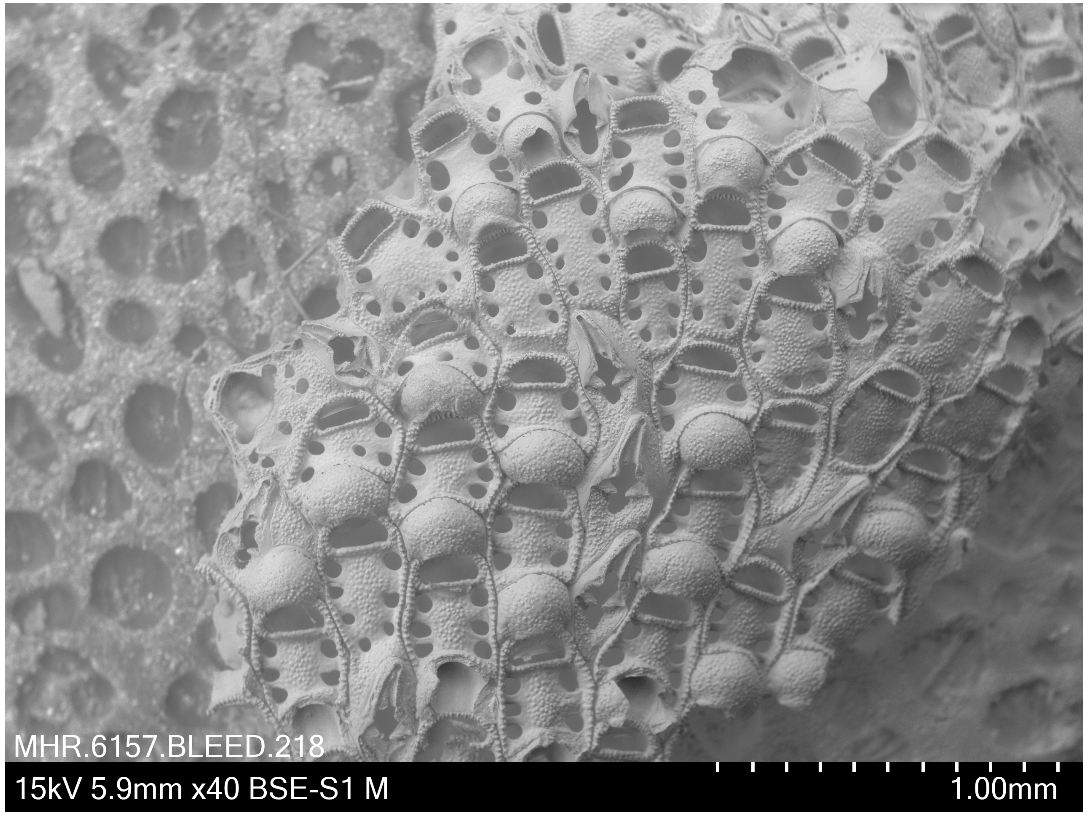


***Opaeophora lepida* BLEED 218**
Coll: 3.2.2008 (A. M. Smith)
Lat: -46.92
Long: 67.34
Depth: 150 m

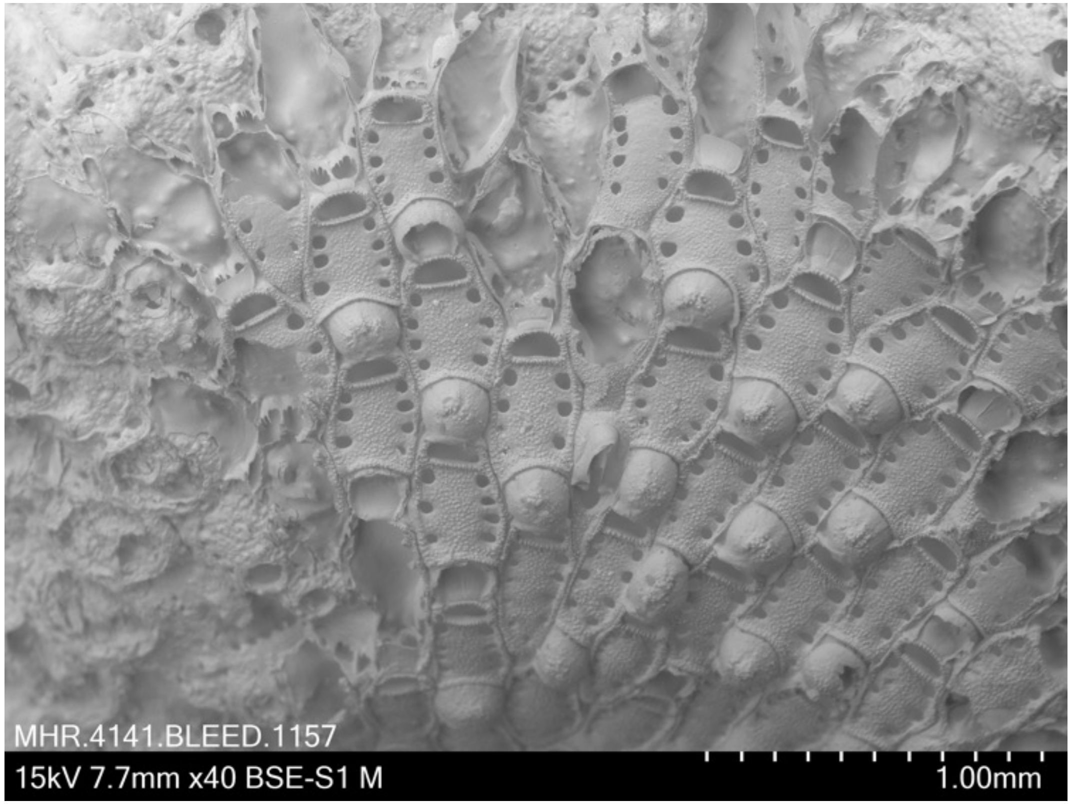


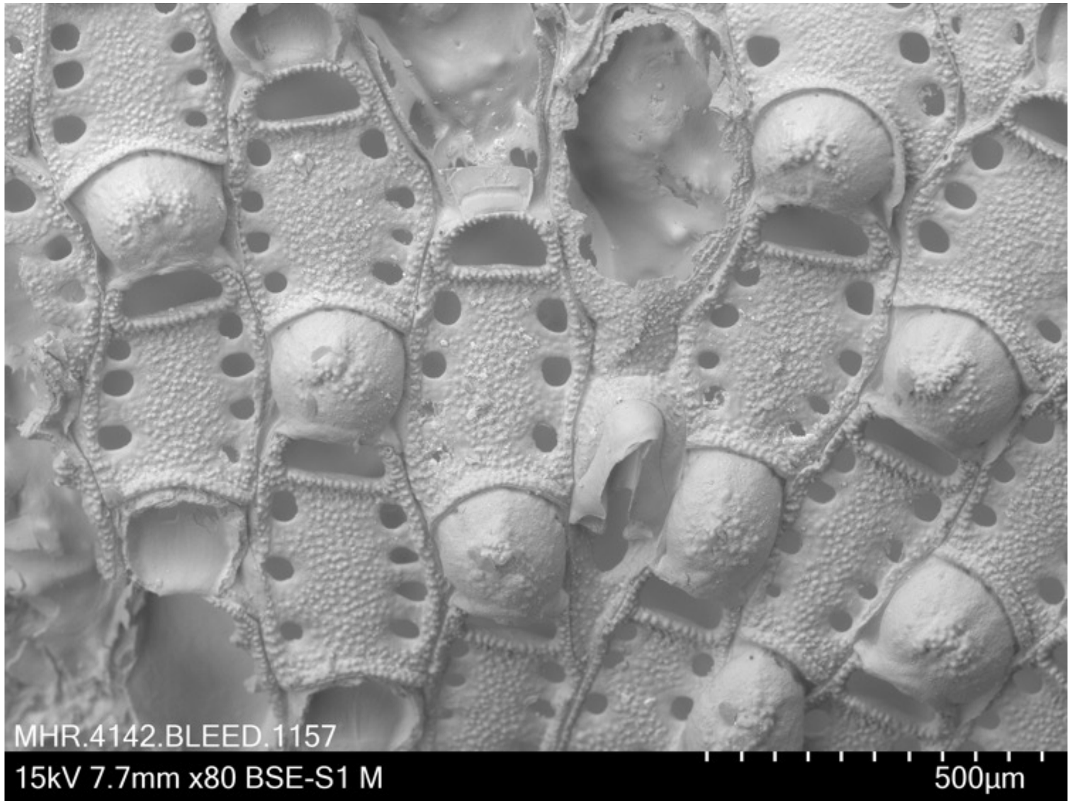


***Opaeophora* n. sp. BLEED 1157**
Coll: 10.11.2018 (A. M. Smith)
Lat: -45.840555999999999
Long: 170.796111
Depth: 56 m

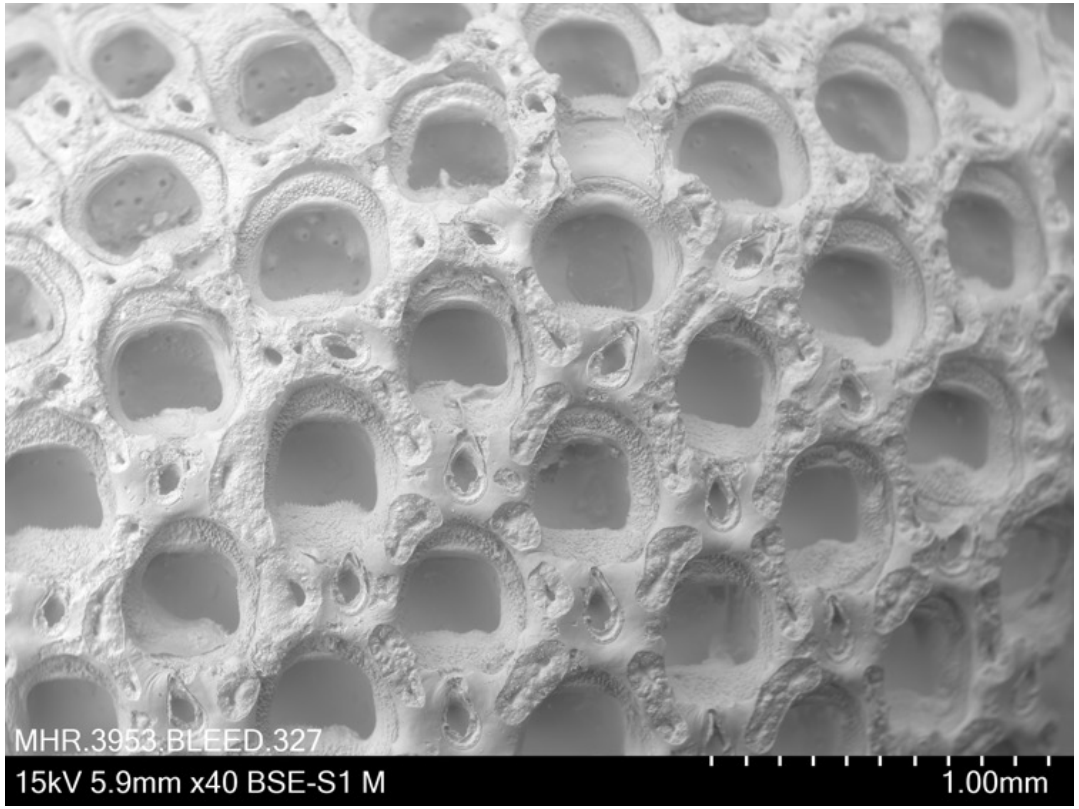

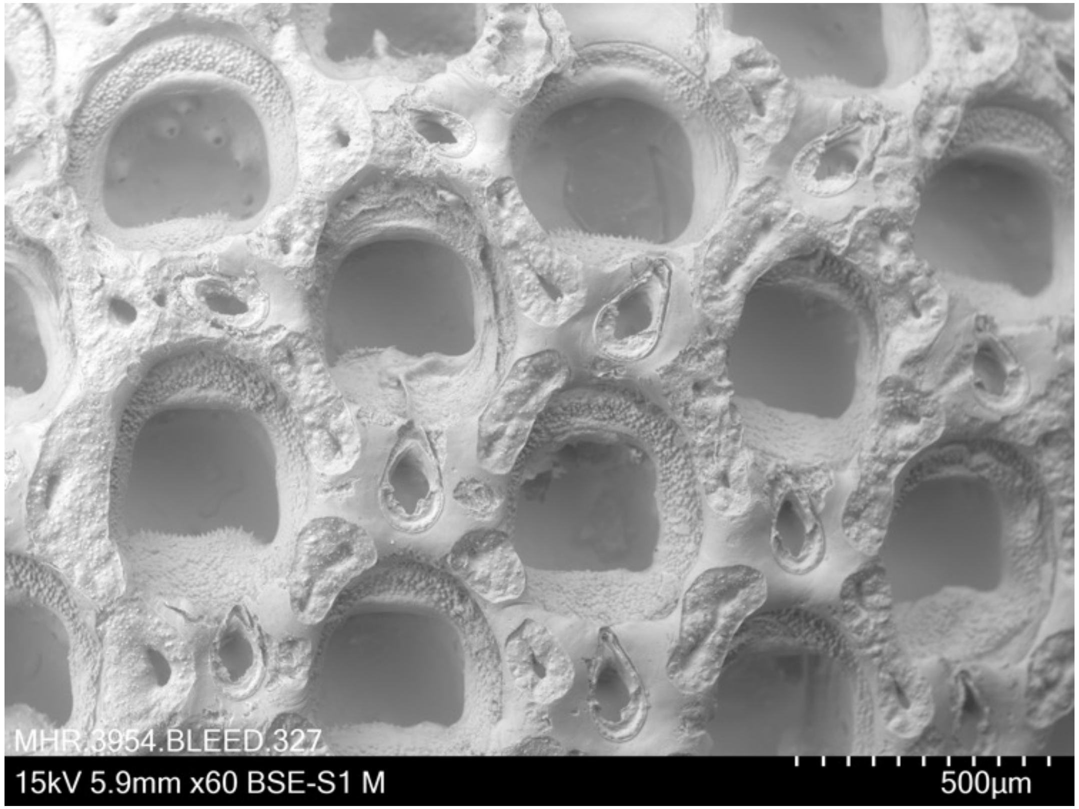


***Odontionella cyclops* BLEED 327**
NIWA 74073 (TAN1108/6)
Coll: 12.5.2011
Lat: -42.796
Long: 173.5876667
Depth: 187 m

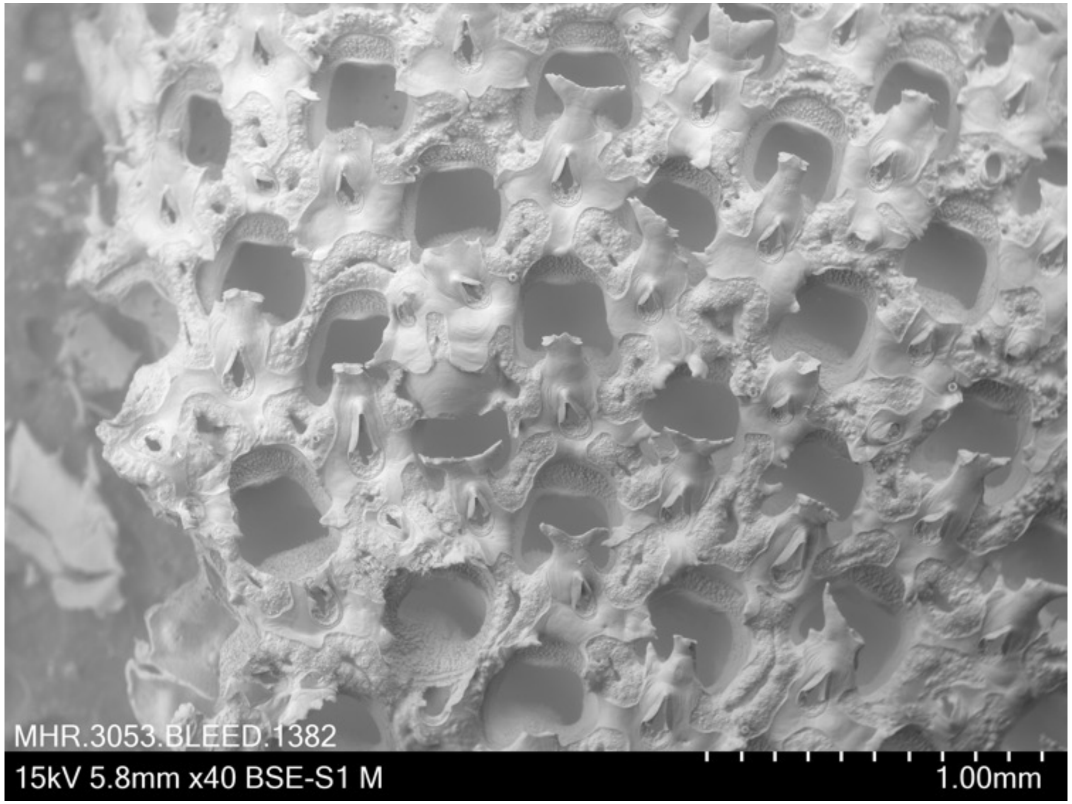


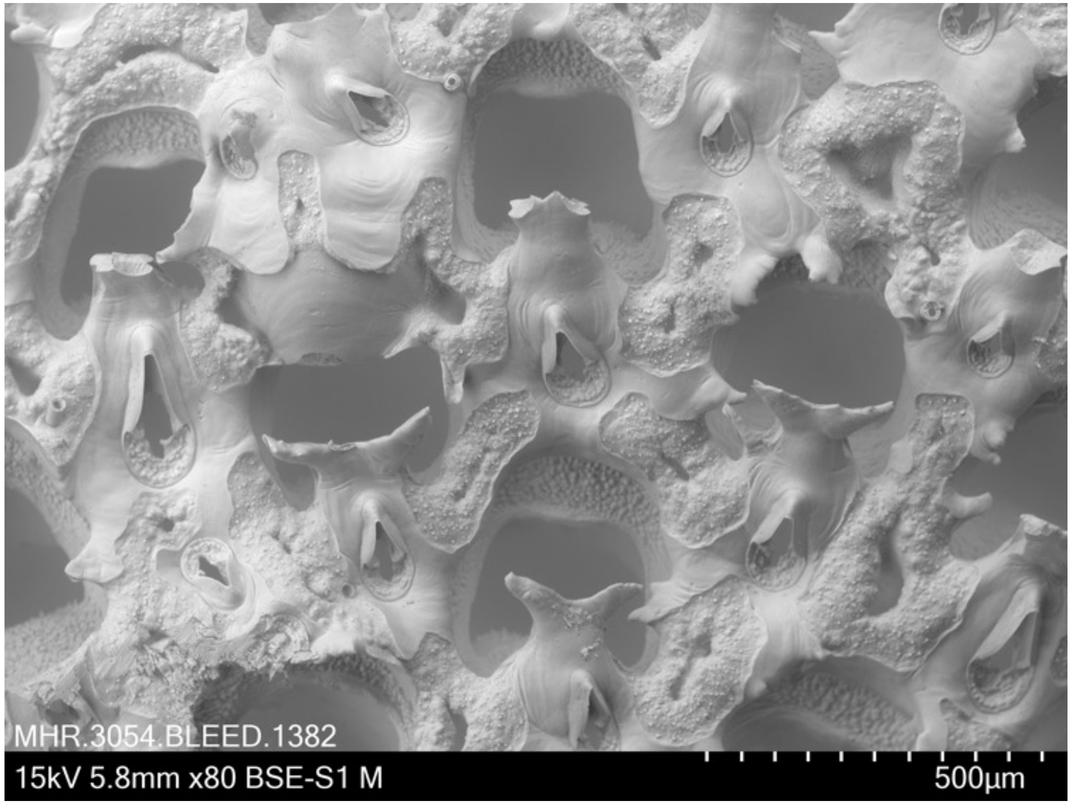


***Odontionella* n. sp. BLEED 1382**
Coll: 17.3.2019 (A. M. Smith, D. P. Gordon)
Lat: -34.159999999999997
Long: 172.09
Depth: 82.9 m

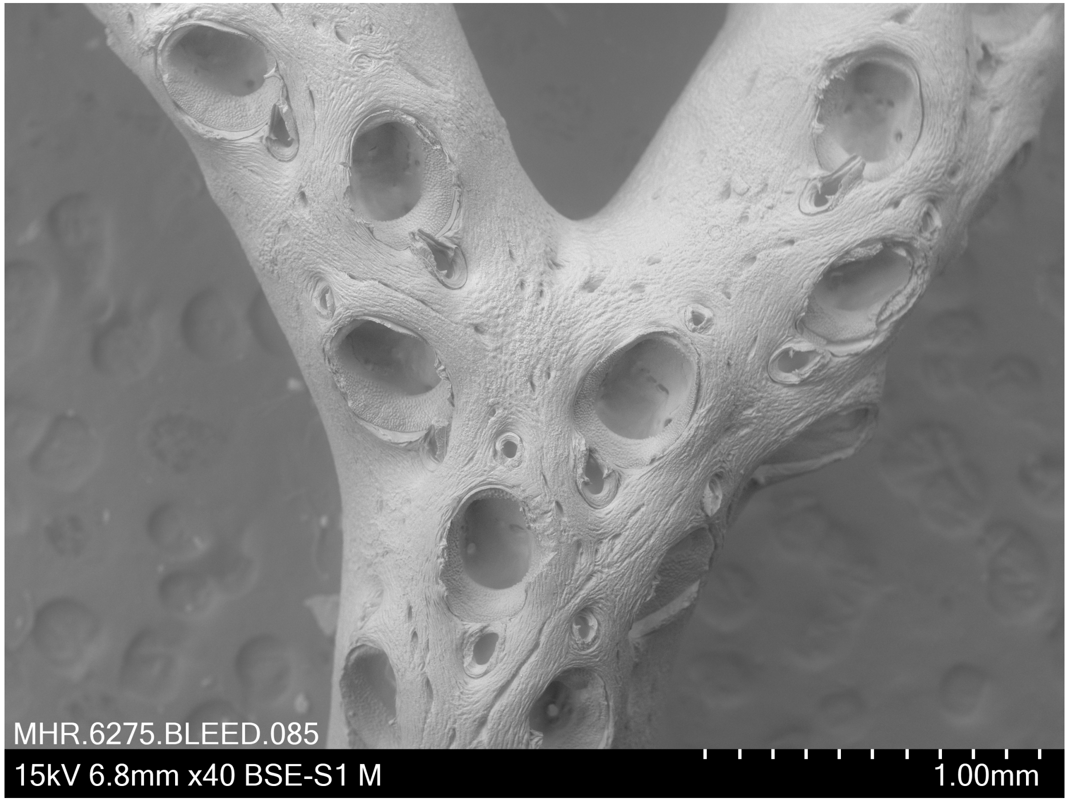


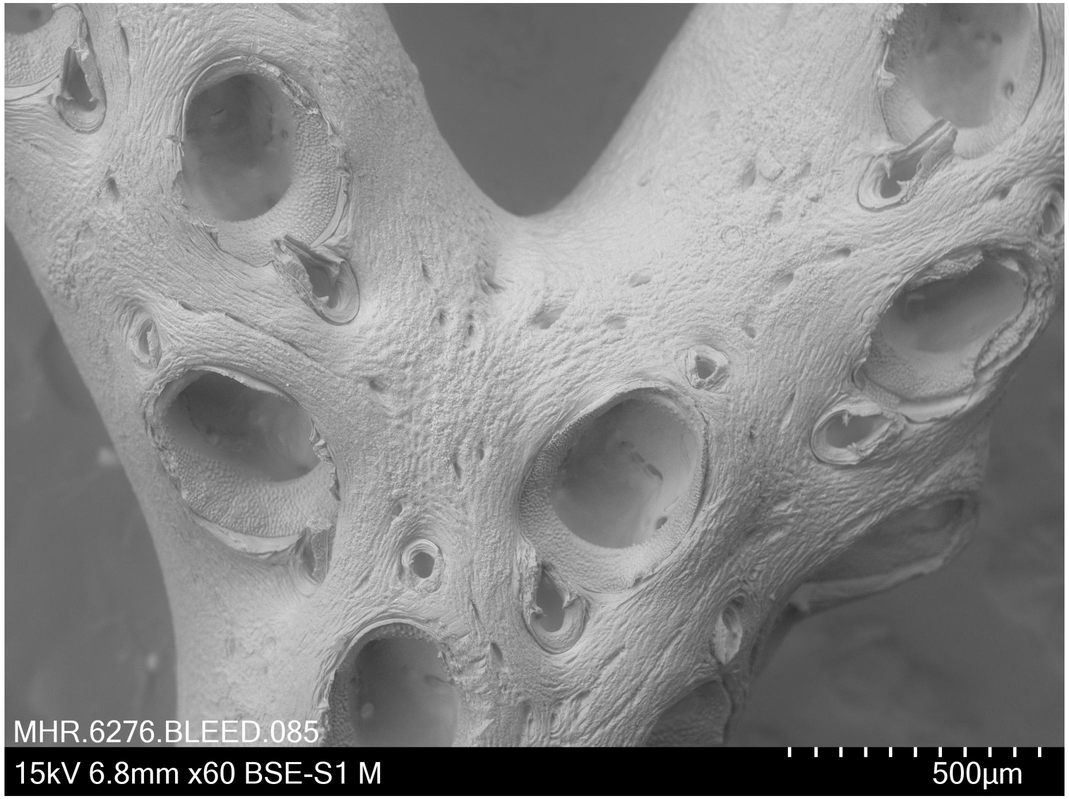


***Foveolaria* n. sp. BLEED 85**
NIWA 65823 (TRIP2938/5)
Coll: 24.09.2009
Lat: -55.616666666699999
Long: -135.6
Depth: 1241 m

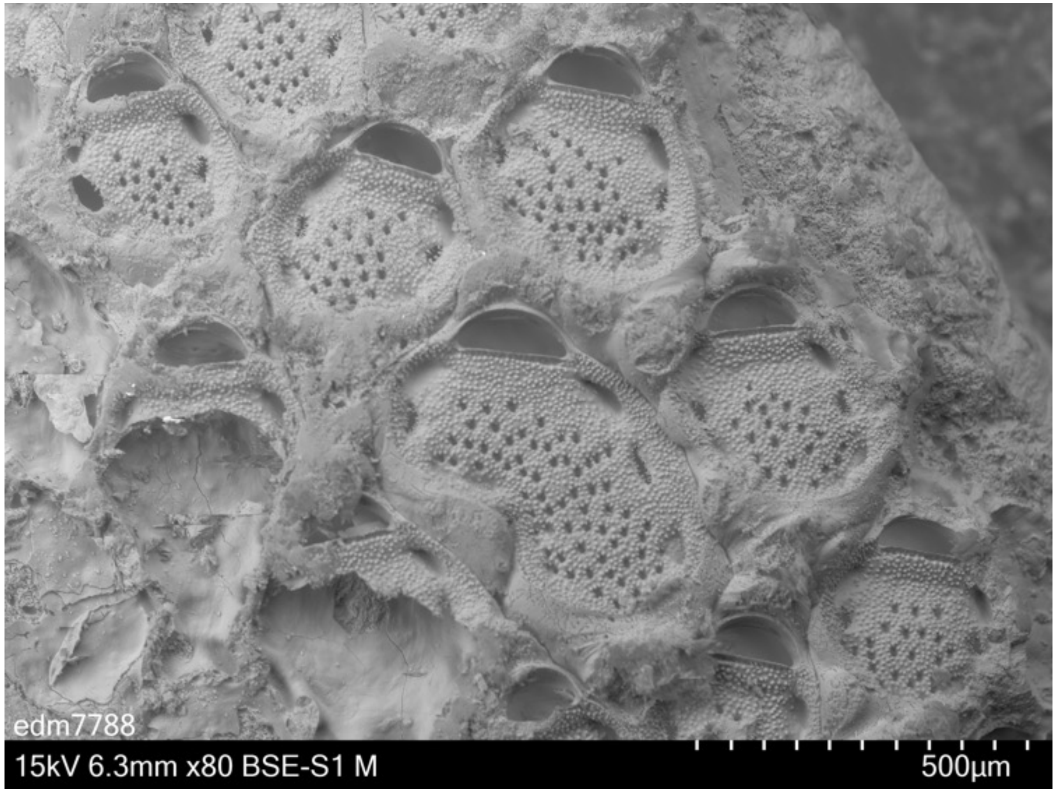


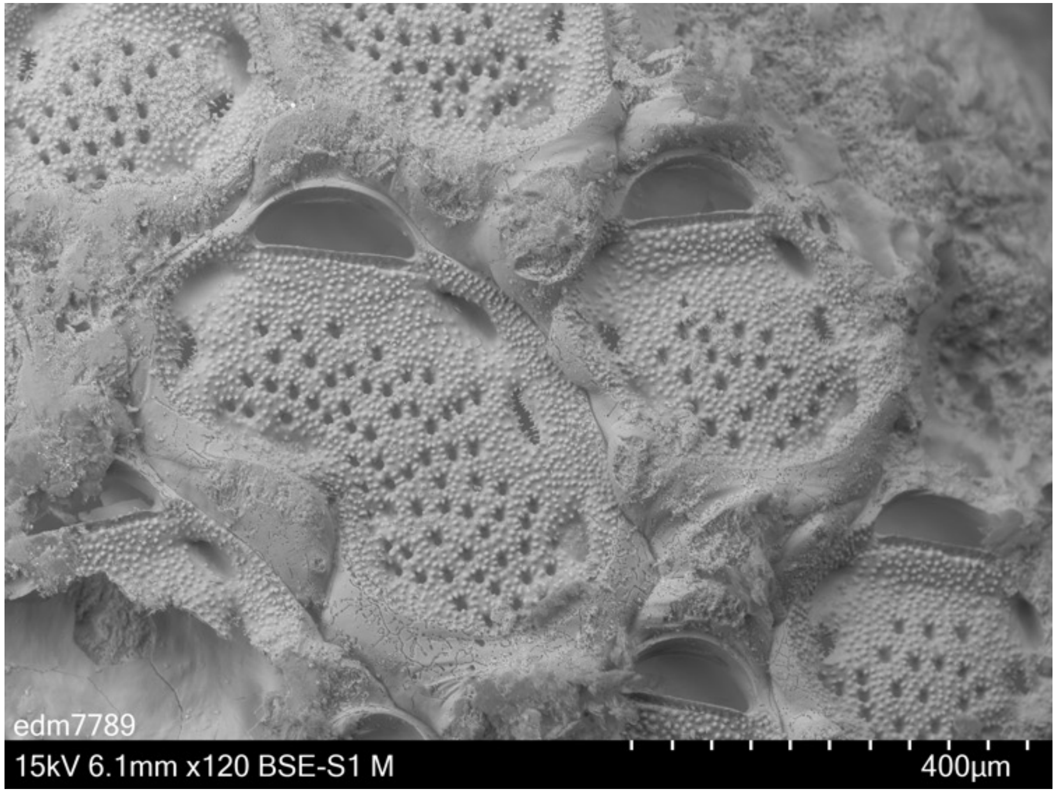


***Micropora* n. sp. 1 BLEED 607**
Coll: 11.4.2018 (A. M. Smith, D. P. Gordon, H. Mello)
Lat: -46.853611111111114
Long: 168.26111111111112
Depth: 24 m

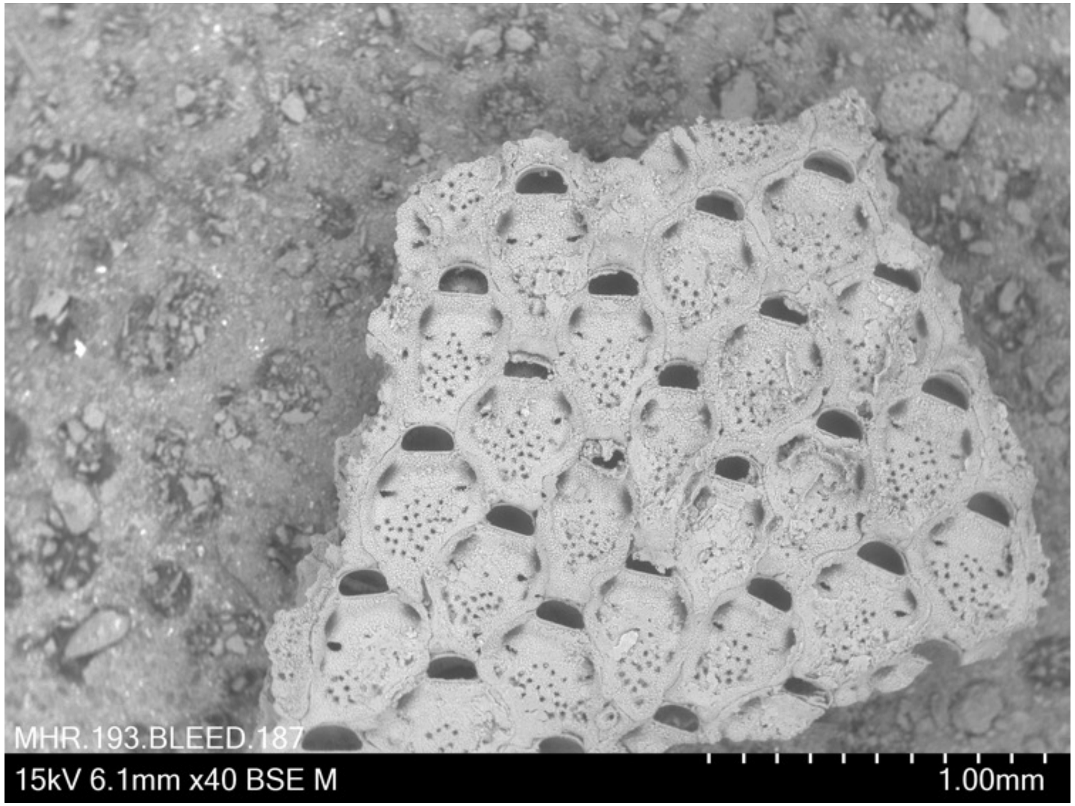


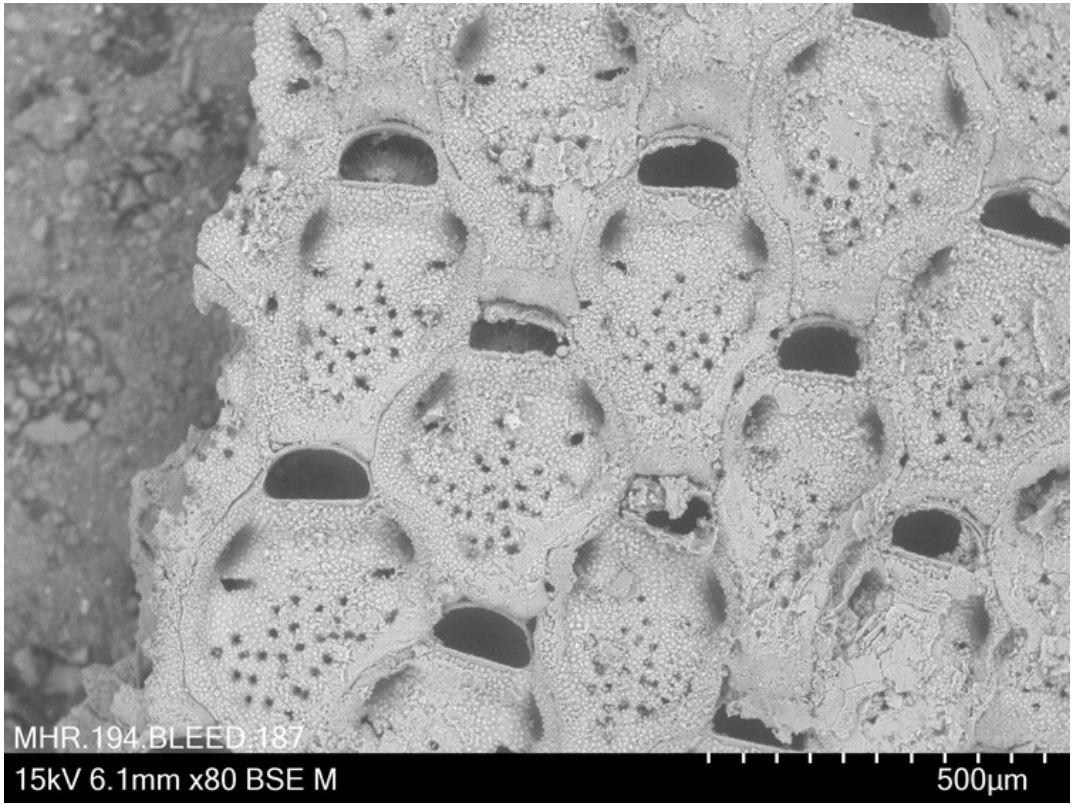


***Micropora* n. sp. 2 BLEED 187**
NIWA 56317 (TAN0906/143)
Coll: 13.7.2009
Lat: -34.7071667
Long: 173.0295
Depth: 77 m

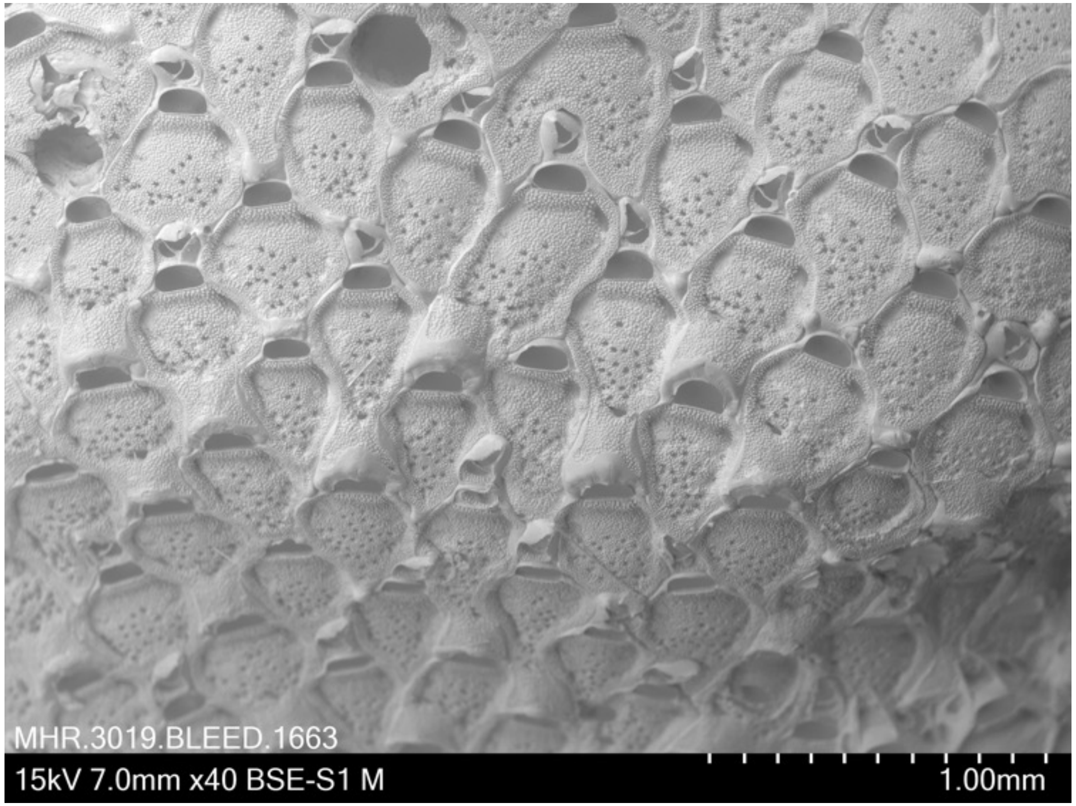


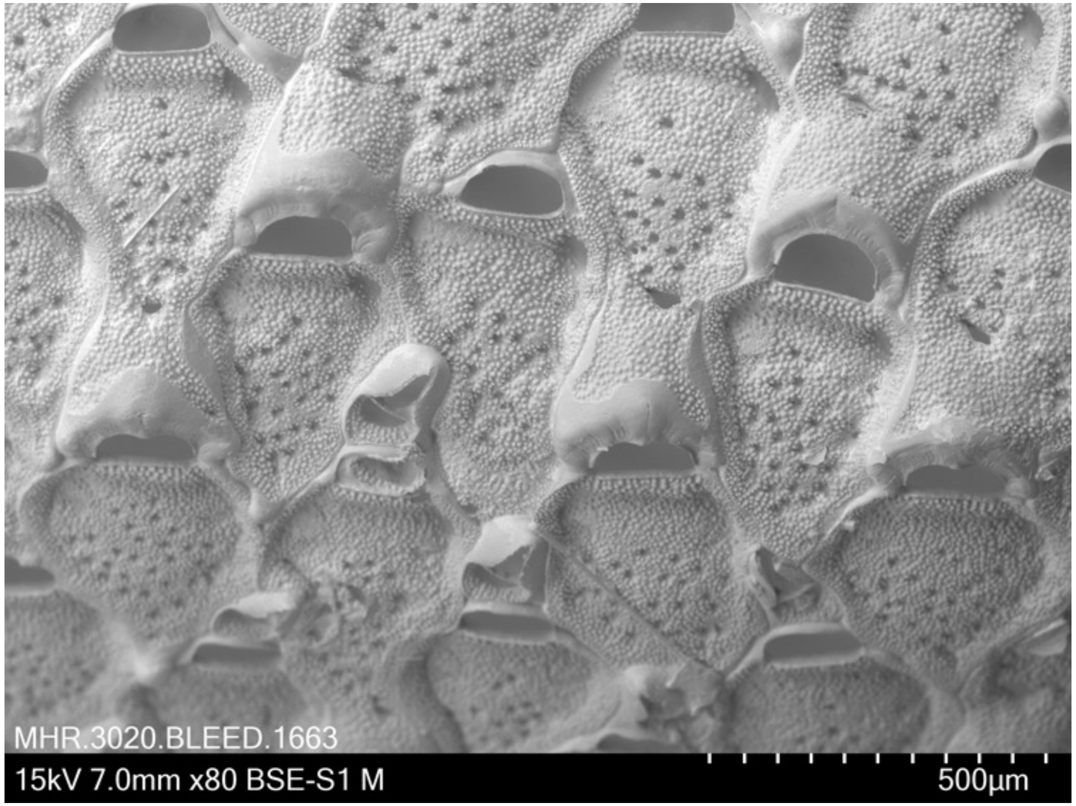


***Micropora* n. sp. 2 BLEED 1663**
Coll: 17.3.2019 (A. M. Smith, D. P. Gordon)
Lat: -34.21
Long: 172.11
Depth: 104 m

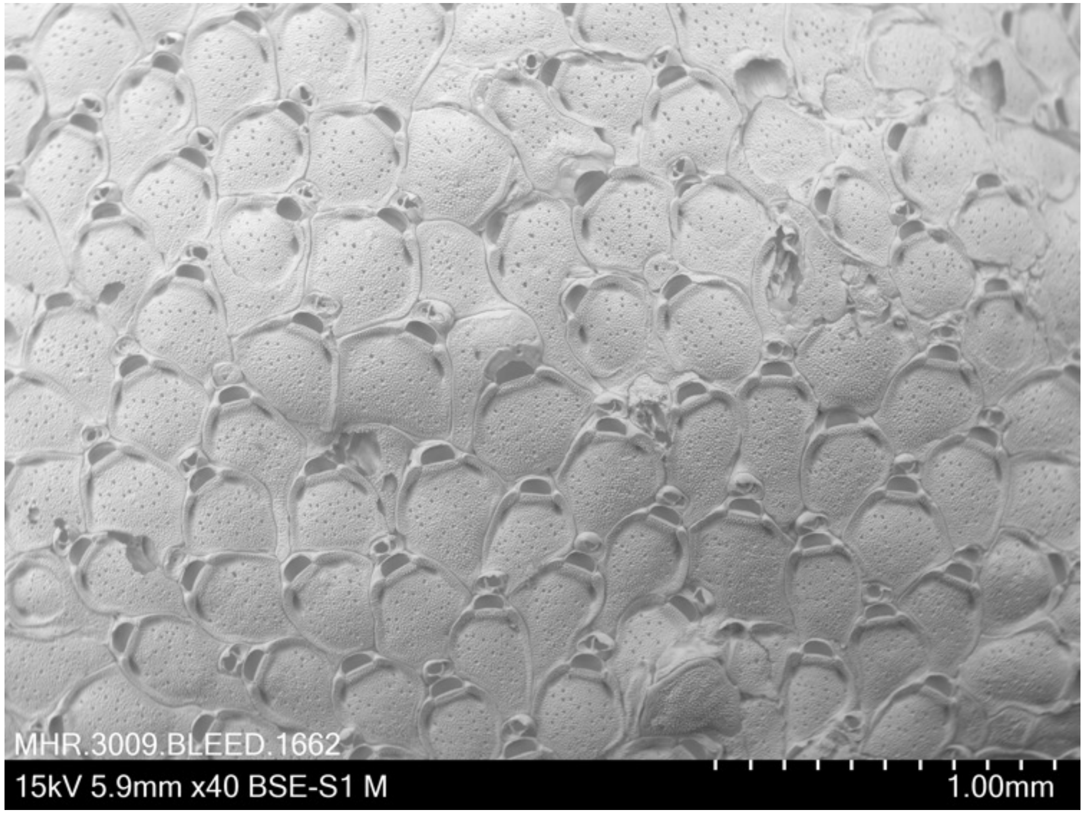

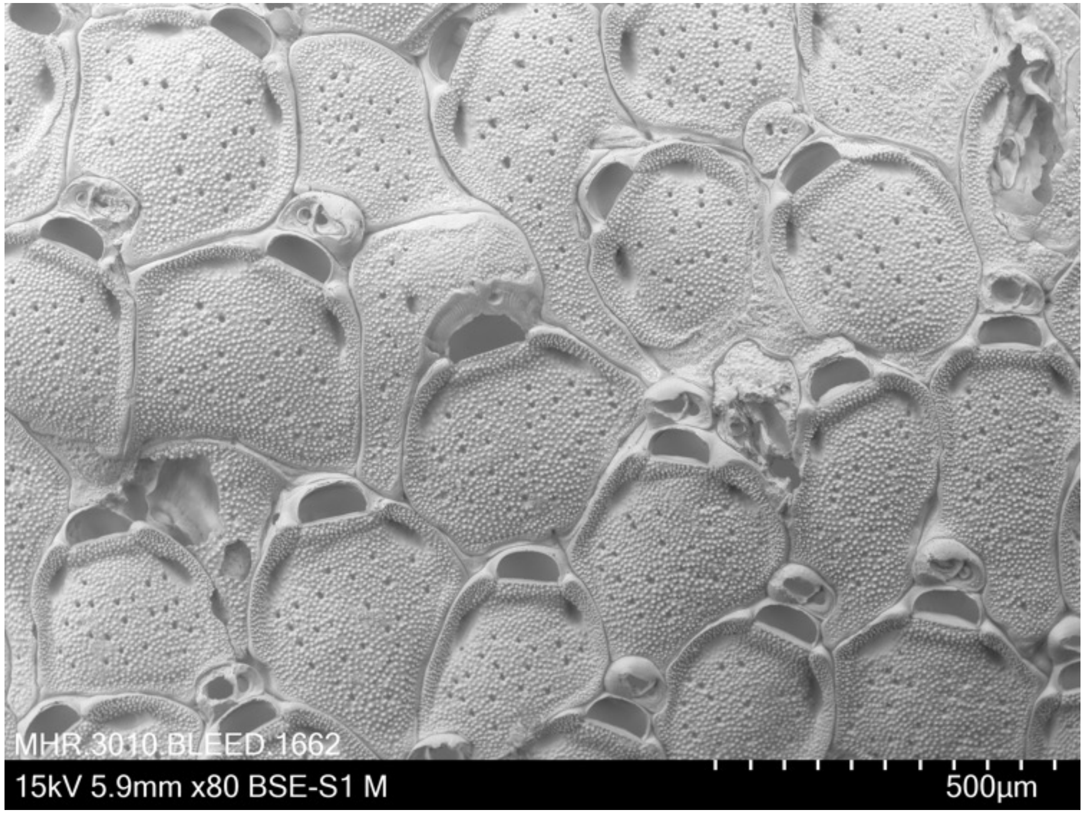


***Micropora* n. sp. 3 BLEED 1662**
Coll: 17.3.2019 (A. M. Smith; D. P. Gordon)
Lat: -34.21
Long: 172.11
Depth: 104 m

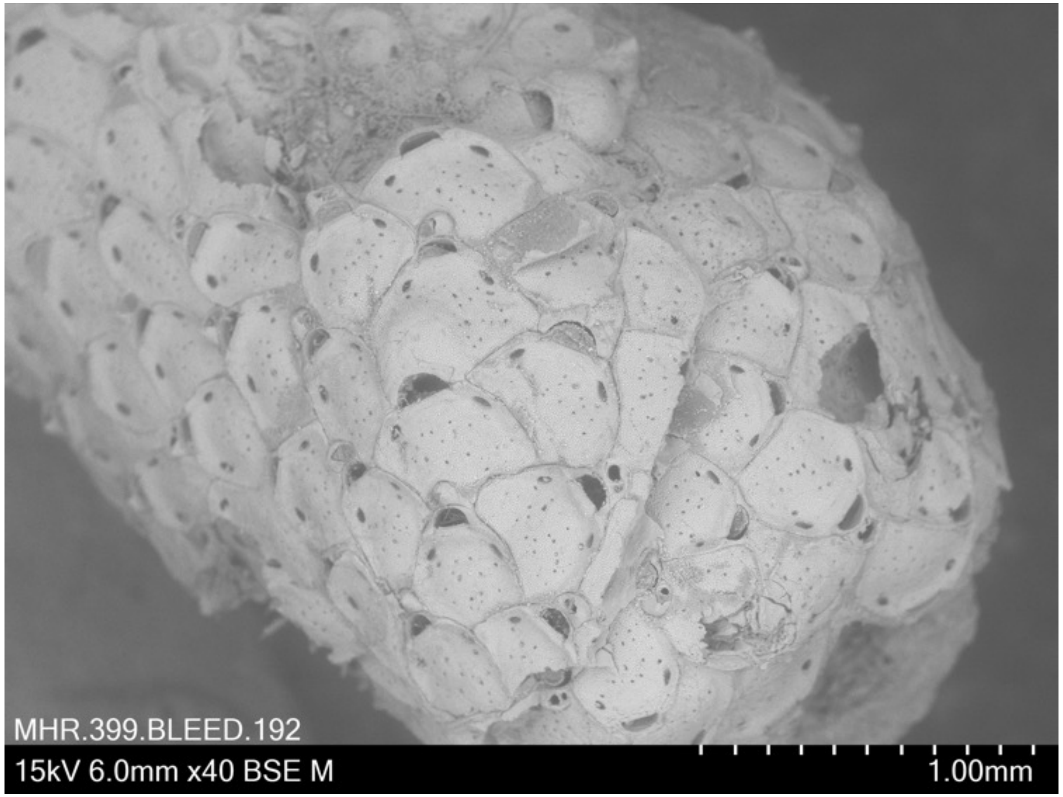


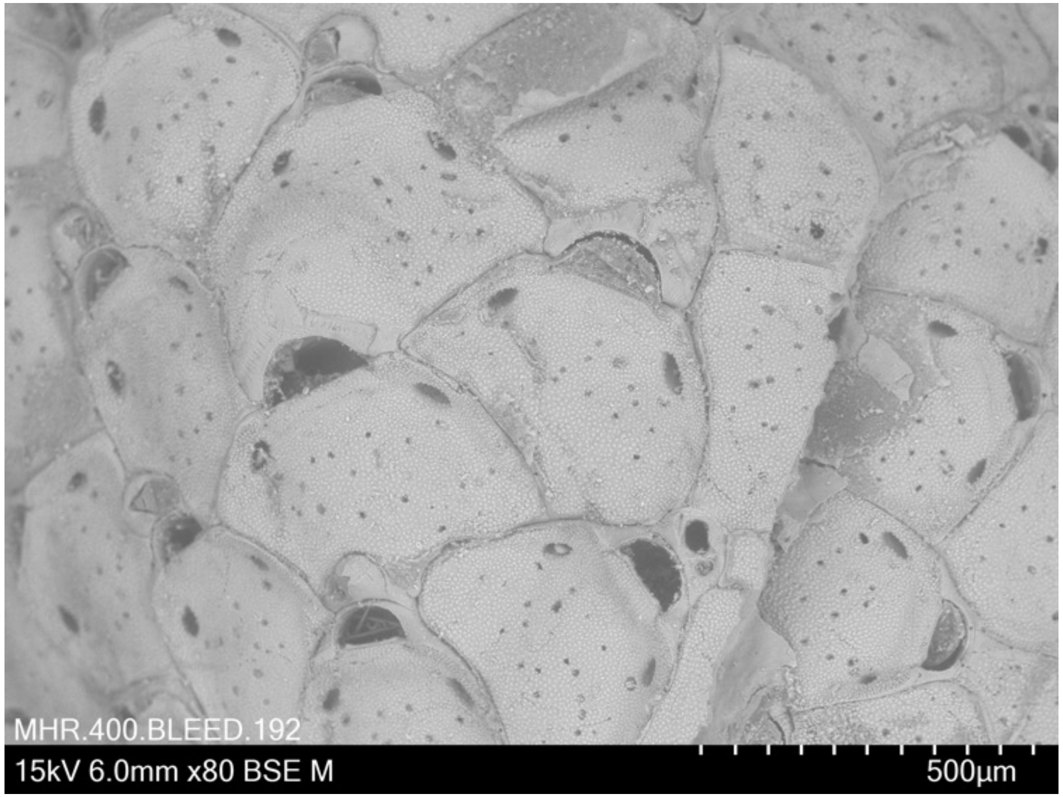


***Micropora* n. sp. 4 BLEED 192**
NIWA 102491 (TAN1503/116)
Coll: 11.04.2015
Lat: -44.1596667
Long: -174.5548333
Depth: 497 m

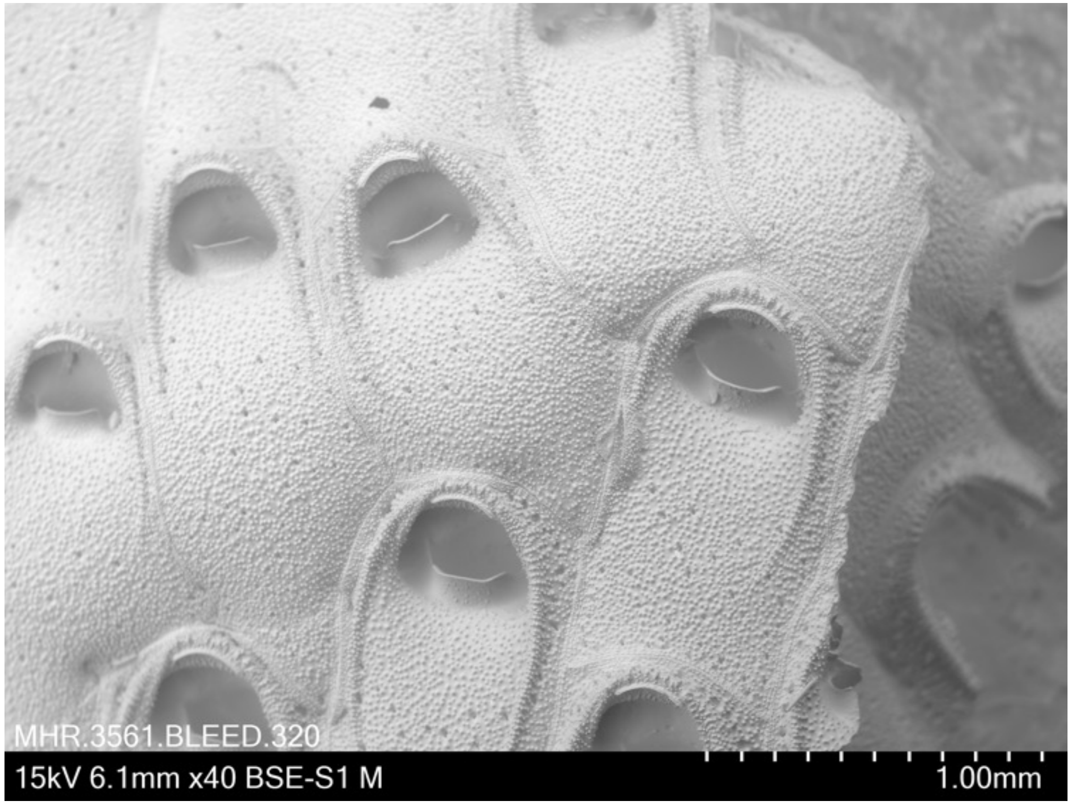


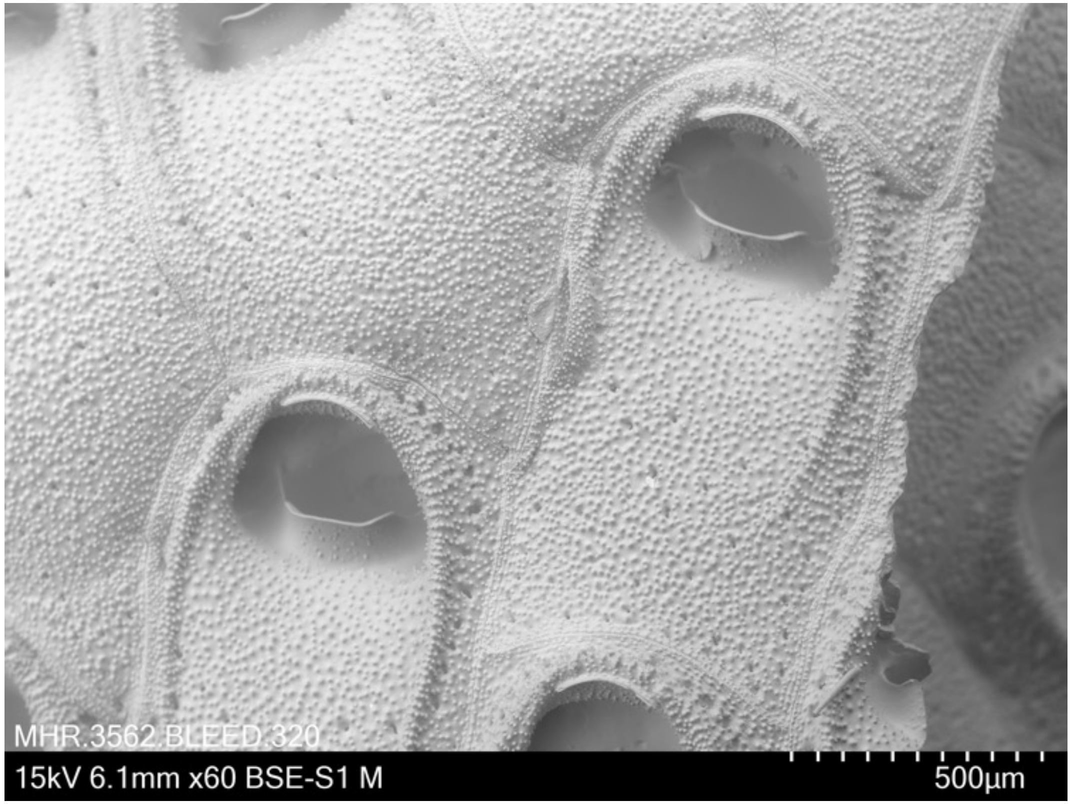


***Crateropora styliformis* BLEED 320**
NIWA 55264 (TAN0906/65)
Coll: 8.7.2009
Lat: -35.003833333300001
Long: 174.05566666670001
Depth: 105-108m

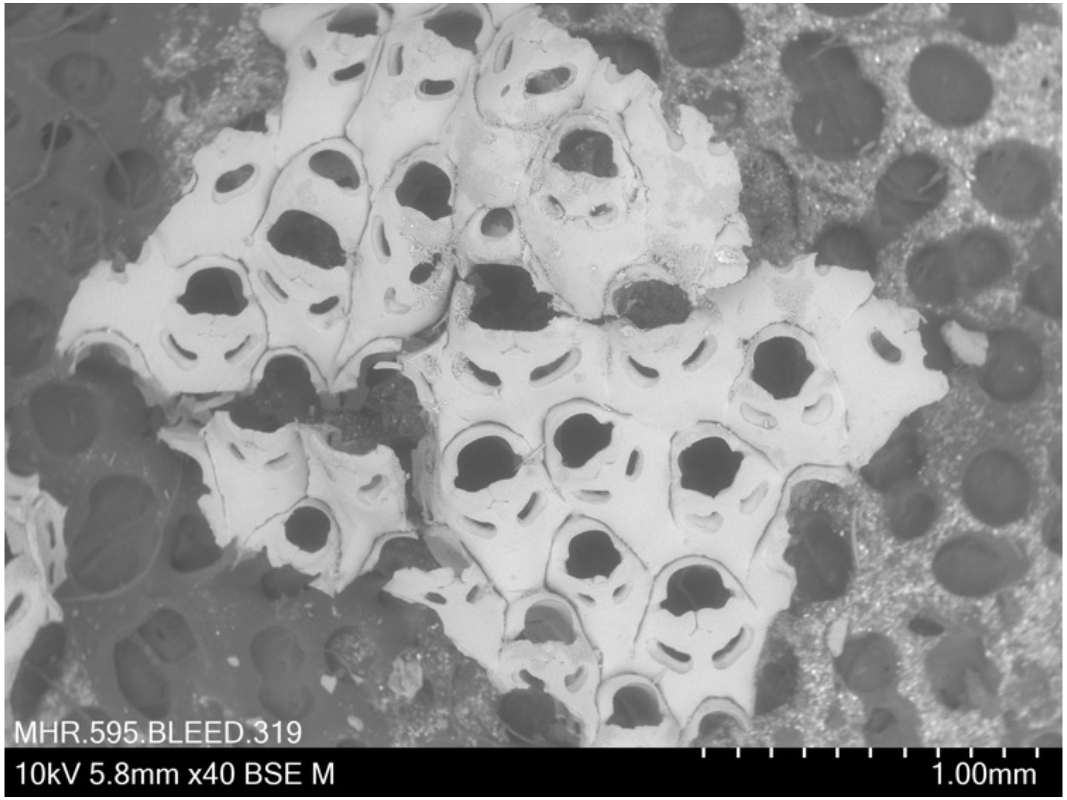


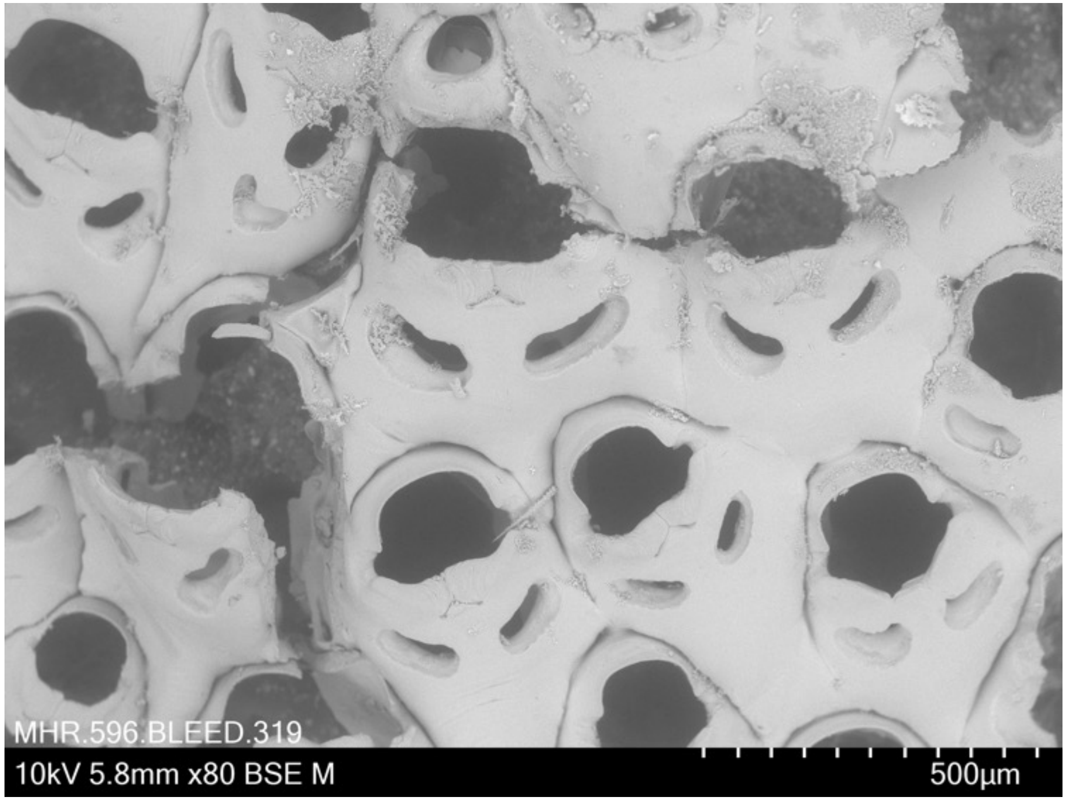


***Eurystomella aupouria* BLEED 319**
NIWA 54468 (TAN0906/2)
Coll: 7.4.2009
Lat: -35.55515
Long: 174.5601667
Depth: 60 m

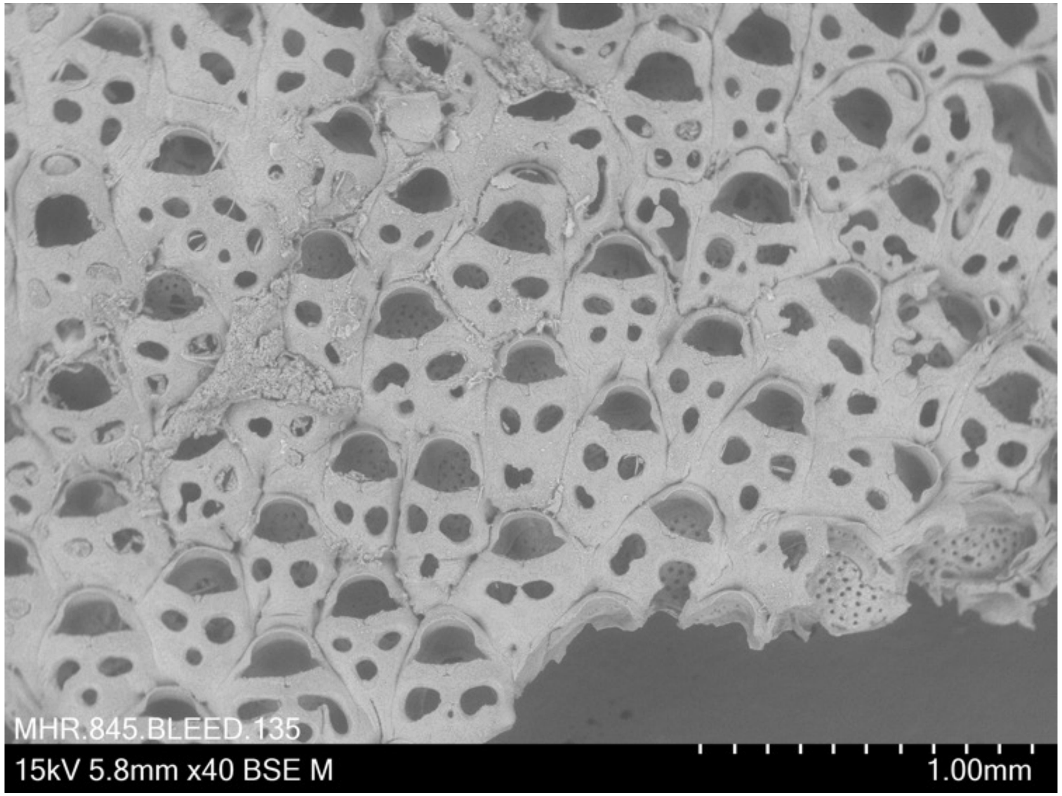


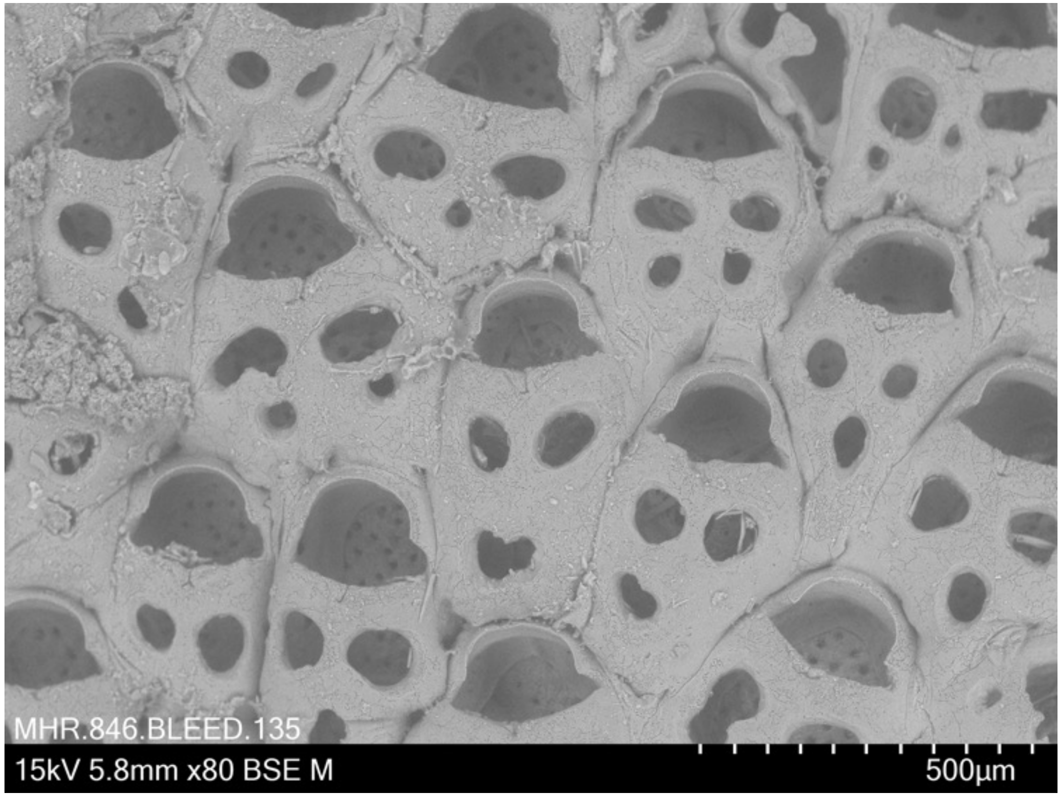


***Eurystomella foraminigera* BLEED 135**
NIWA 99666 (Z16029)
Coll: 02.2015 (D. P. Gordon)
Lat: -41.2833333333
Long: 174.76666666669999
Depth: Intertidal

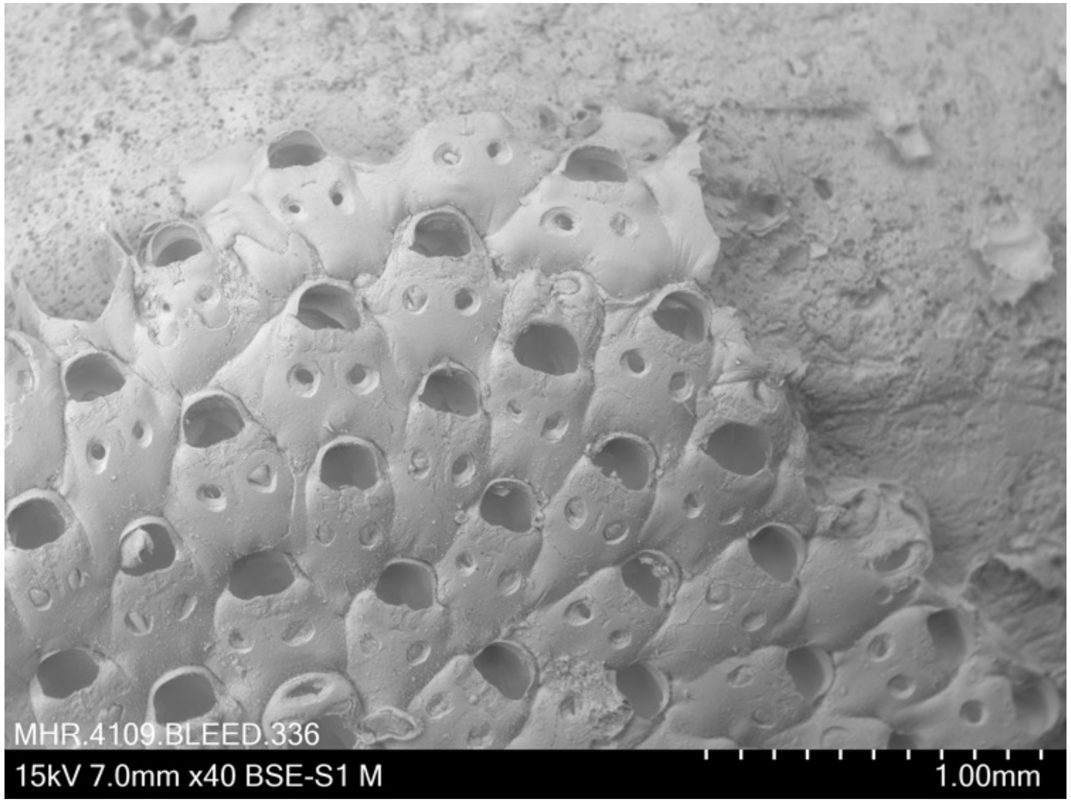


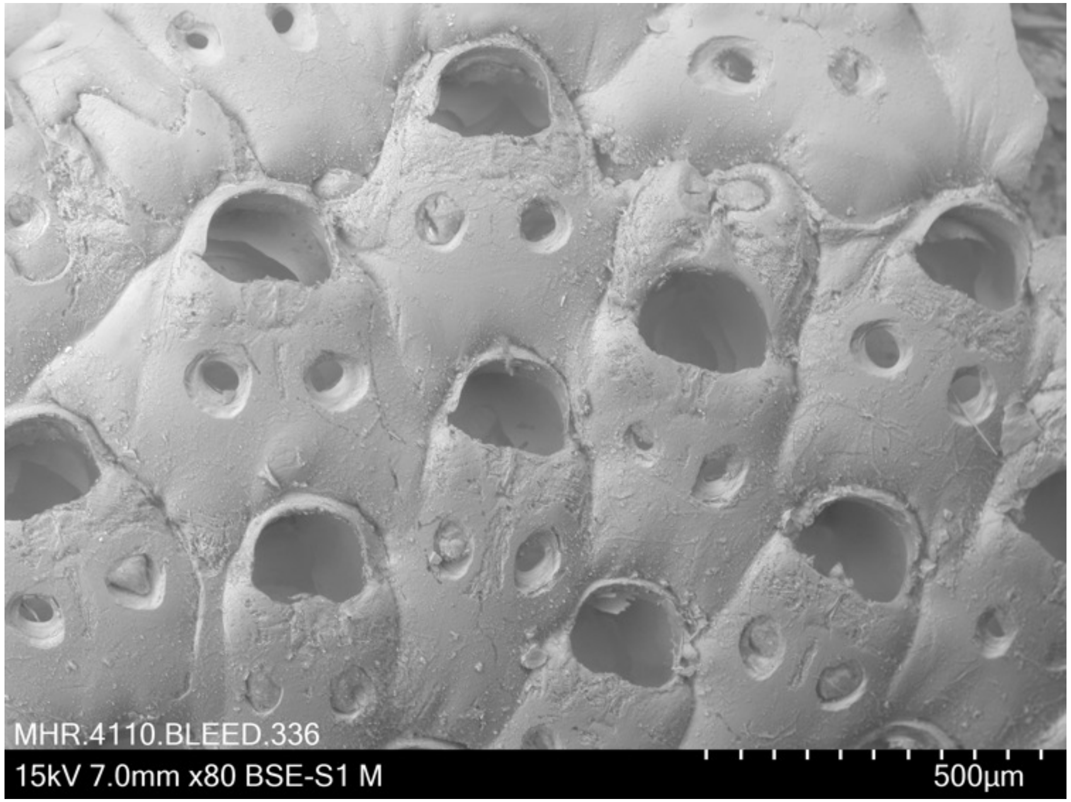


***Eurystomella biperforata* BLEED 336**
Coll: 28.1.2010 (A. M. Smith)
Lat: -47.27
Long: 167.3
Depth: 125m

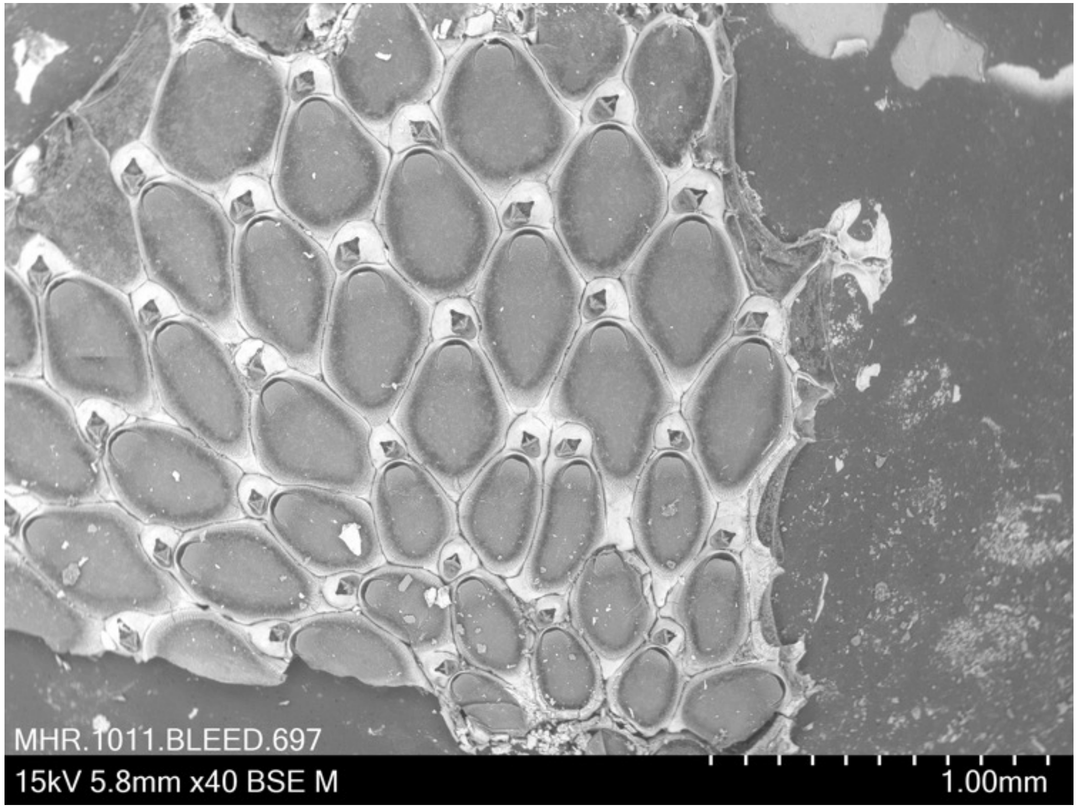


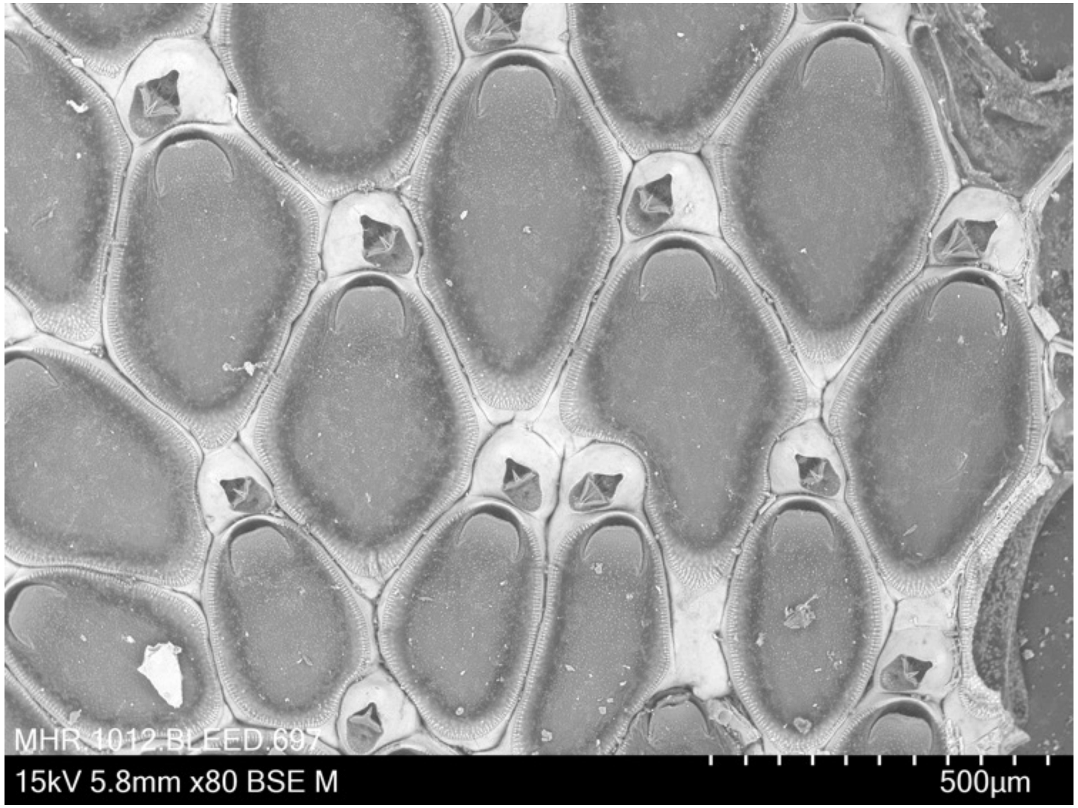


***Ellisina sericea* BLEED 697**
Coll: 8.4.2018 (A. M. Smith, D. P. Gordon, H. Mello)
Lat: -47.1358333333333
Long: 168.010555555556
Depth: 72 m

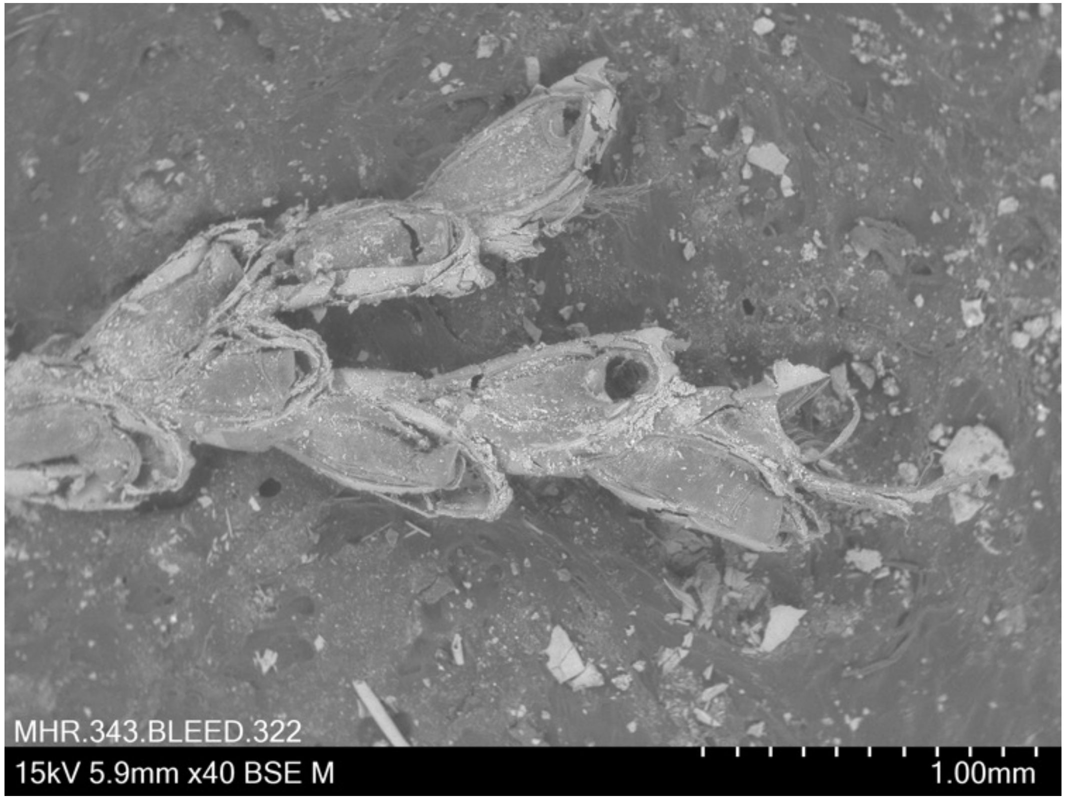


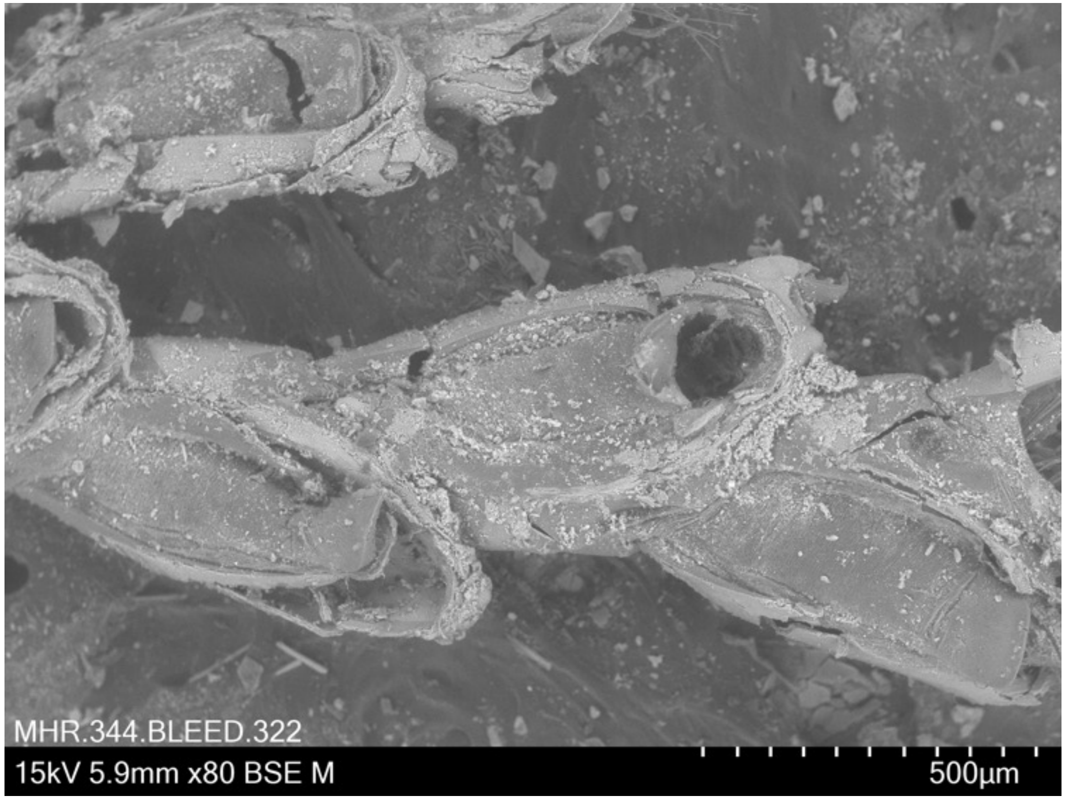


***Euoplozoum* n. sp. BLEED 322**
NIWA 72127 (TAN1104/11)
Coll: 3.3.2011
Lat: -36.459
Long: 177.8473
Depth: 1172 m

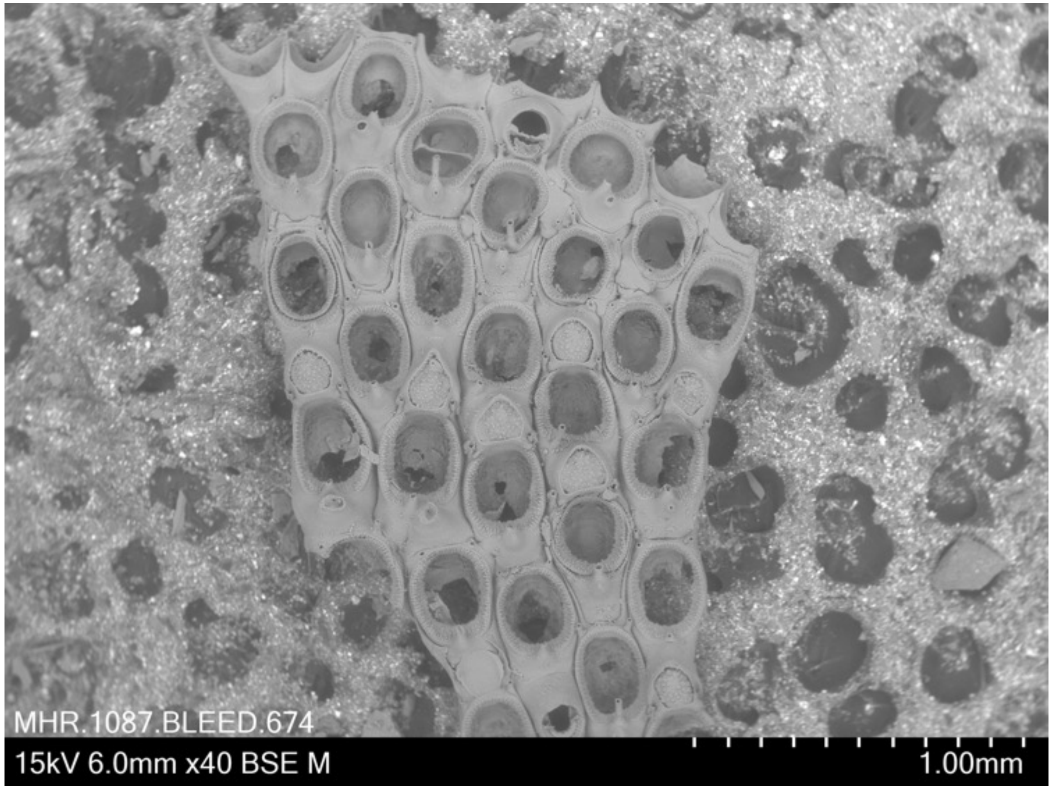


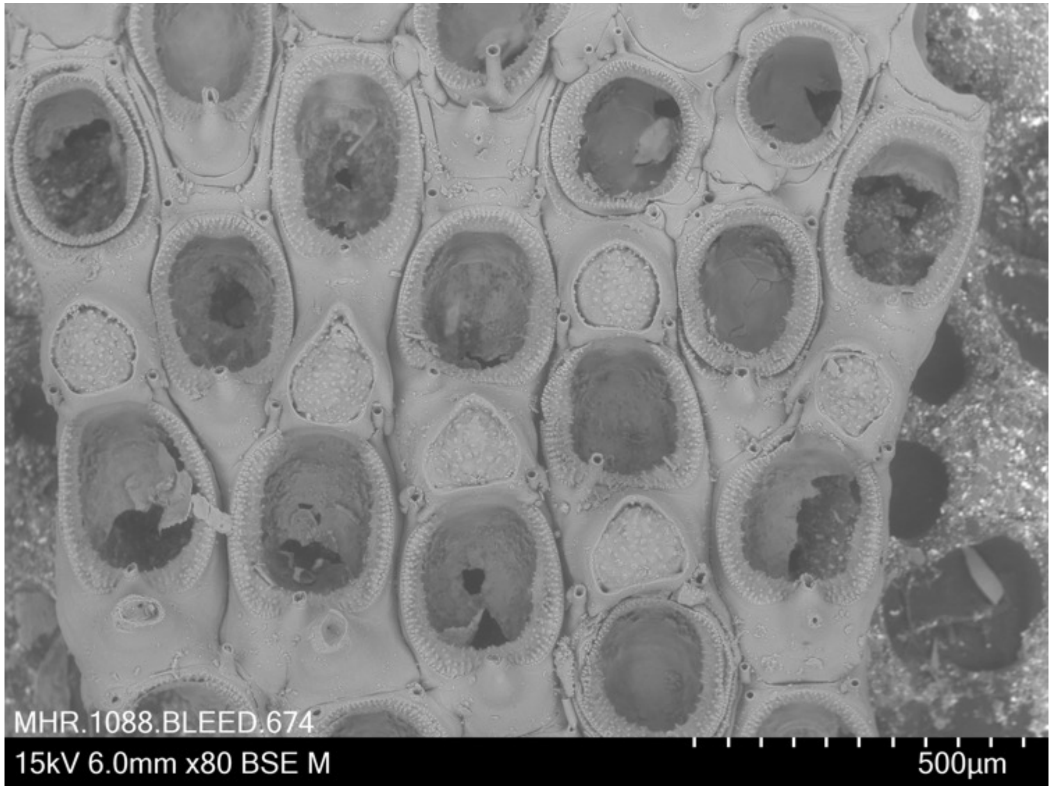


***Crassimarginatella cucullata*** **BLEED 674**
Coll: 8.4.2018 (A. M. Smith, D. P. Gordon, H. Mello)
Lat: -47.13722222222222
Long: 168.12583333333333
Depth: 80-85m

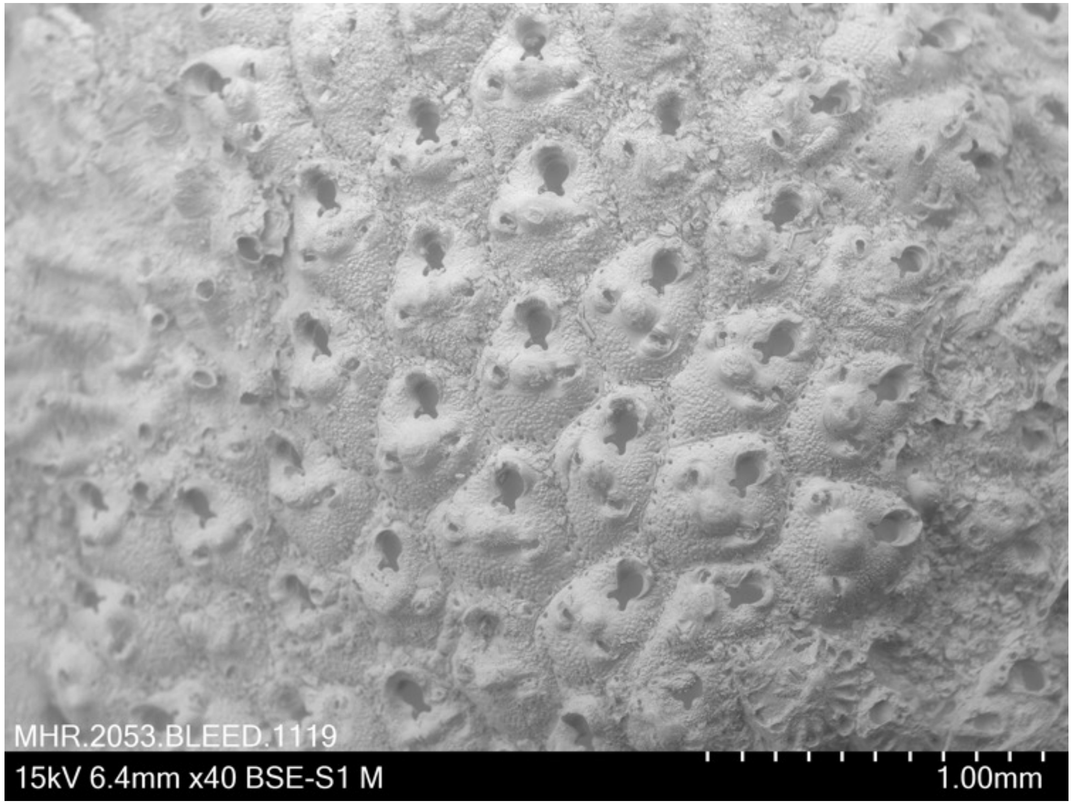


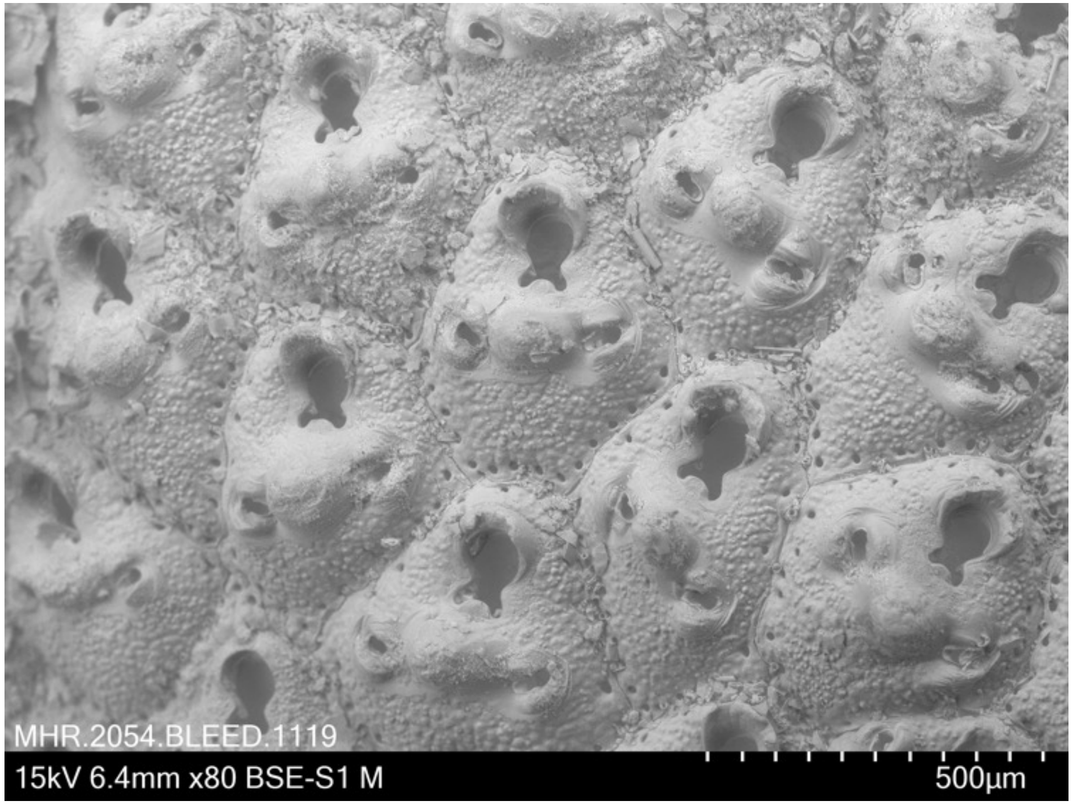


***Crepidacantha zelanica* BLEED 1119**
Coll: 10.11.2018 (A. M. Smith)
Lat: -45.840555999999999
Long: 170.796111
Depth: 56 m

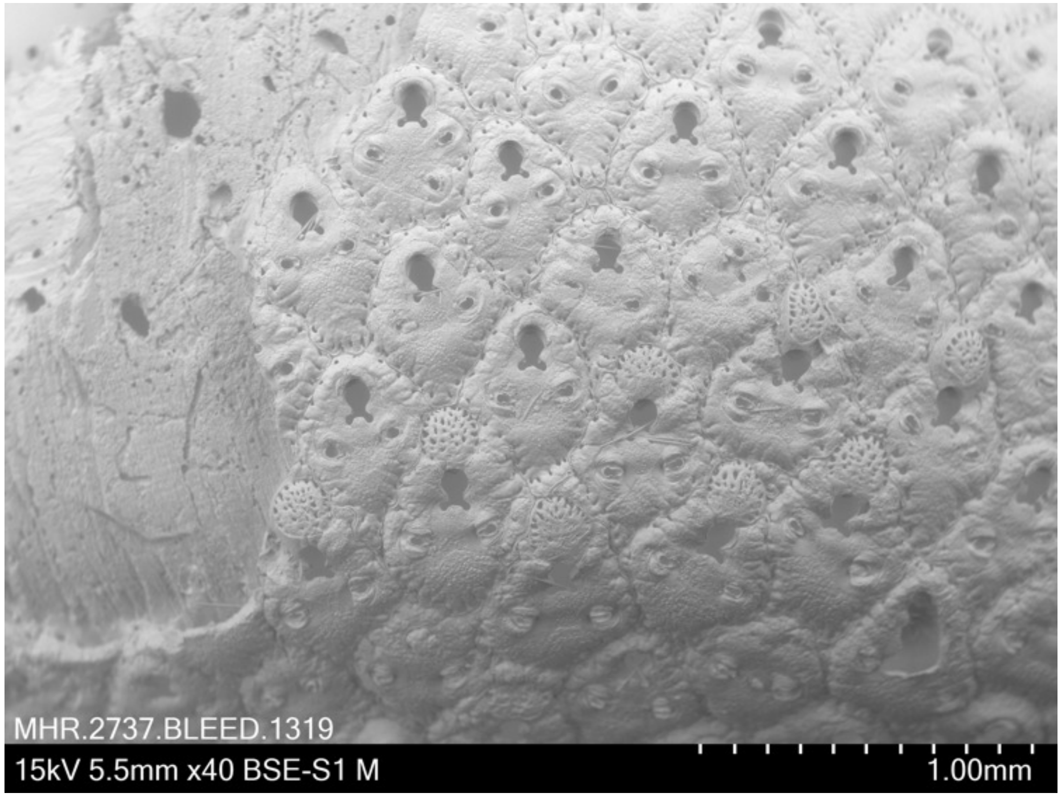


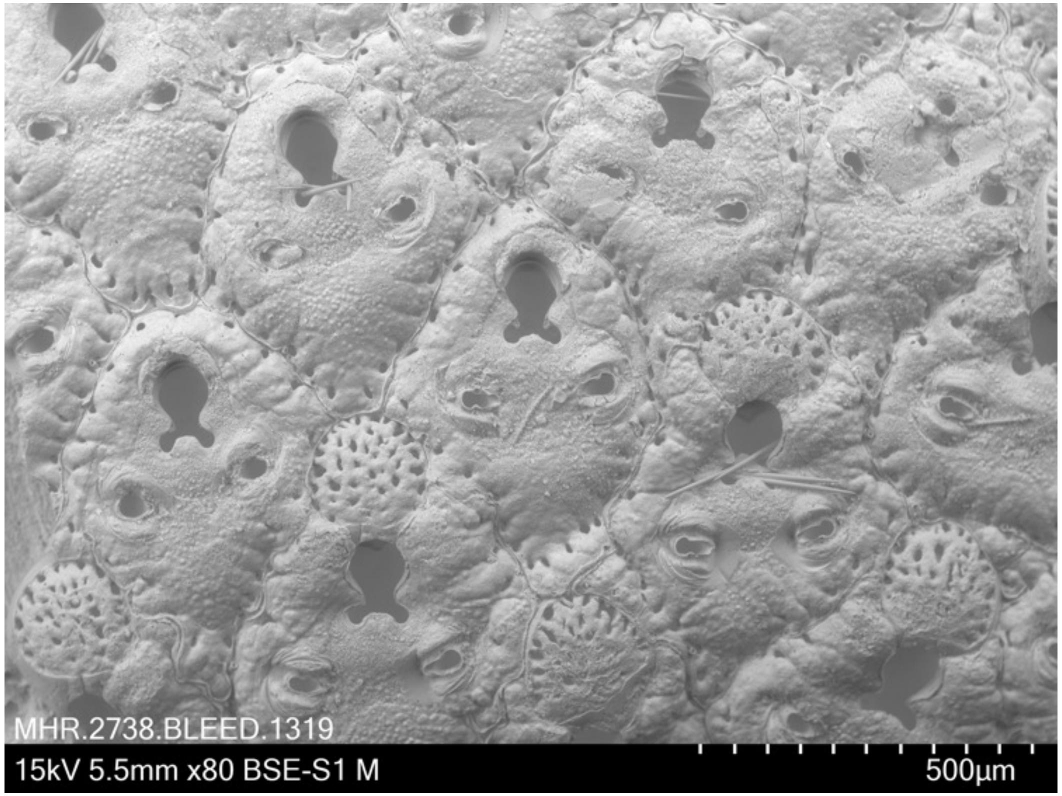


***Crepidacantha zelanica* BLEED 1319**
Coll: 15.3.2019 (A. M. Smith; D. P. Gordon)
Lat: -36.46
Long: 174.9
Depth: 37 m

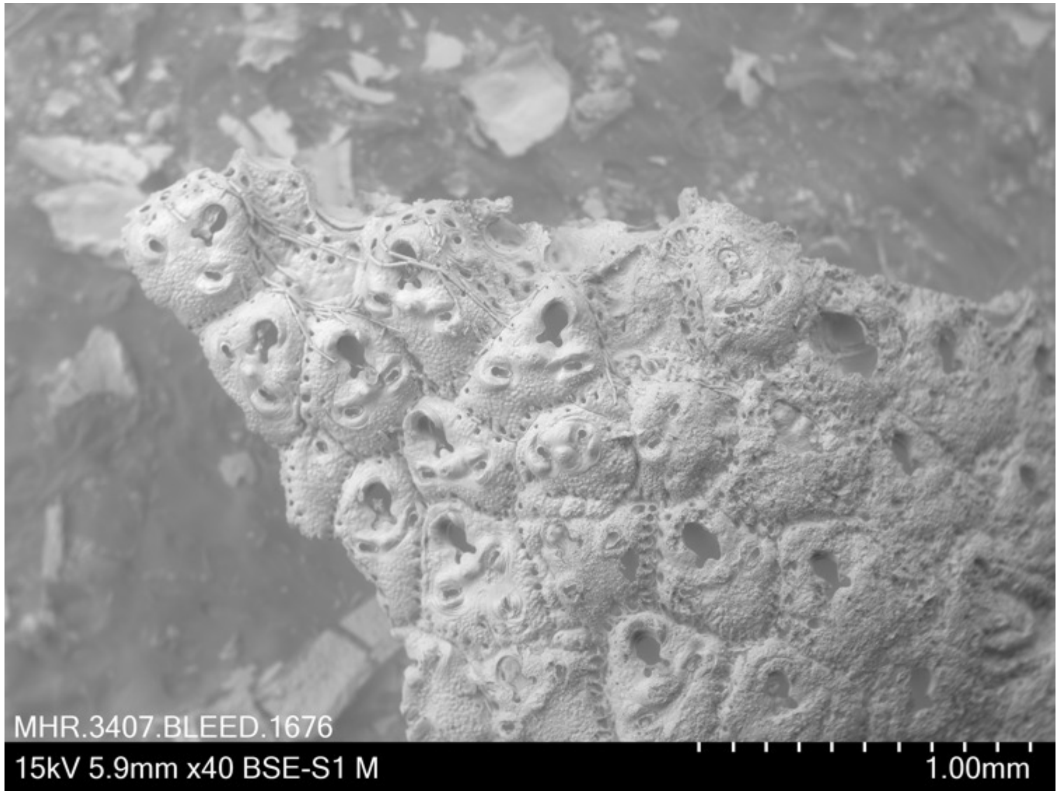


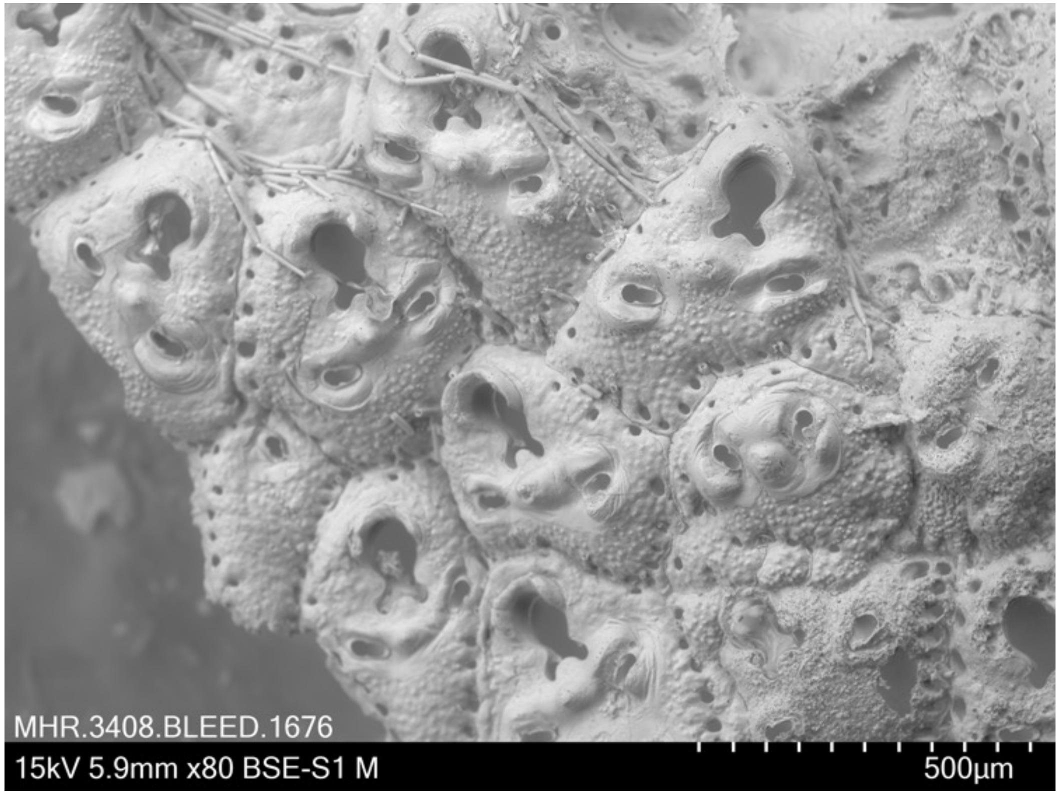


***Crepidacantha kirkpatricki*** **BLEED 1676**
Coll: 18.3.2019 (A. M. Smith, D. P. Gordon)
Lat: -34.380000000000003
Long: 172.69
Depth: 52.5-56.5m

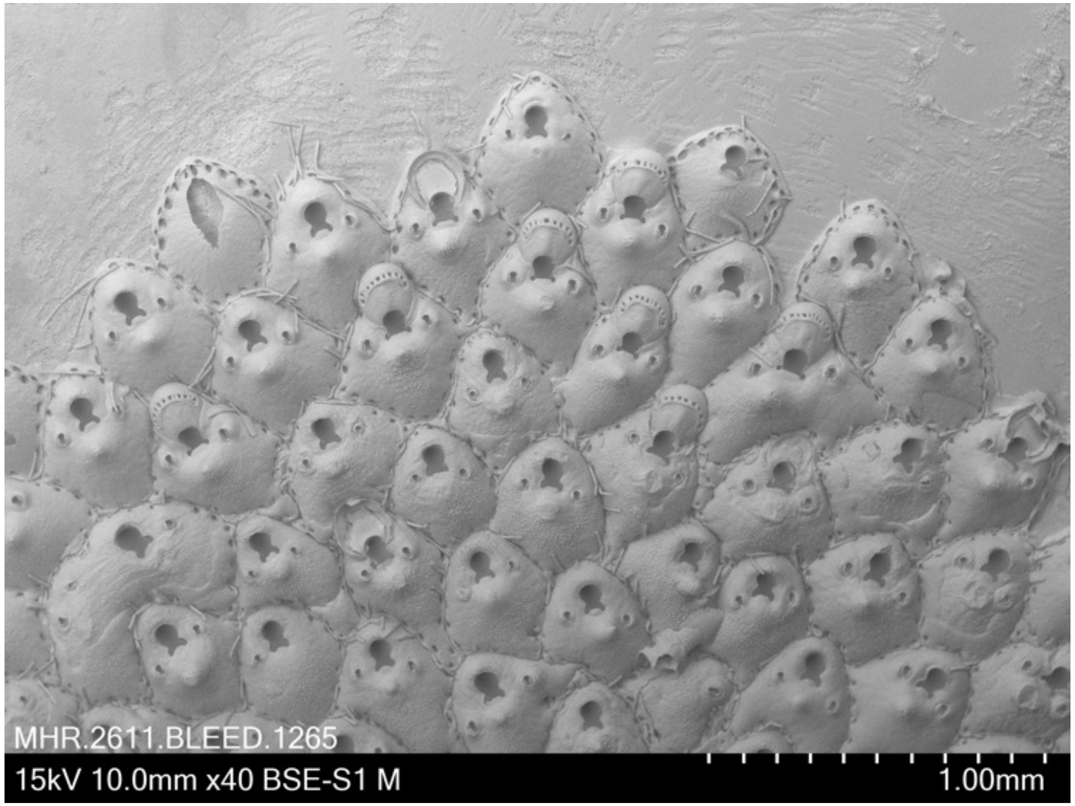


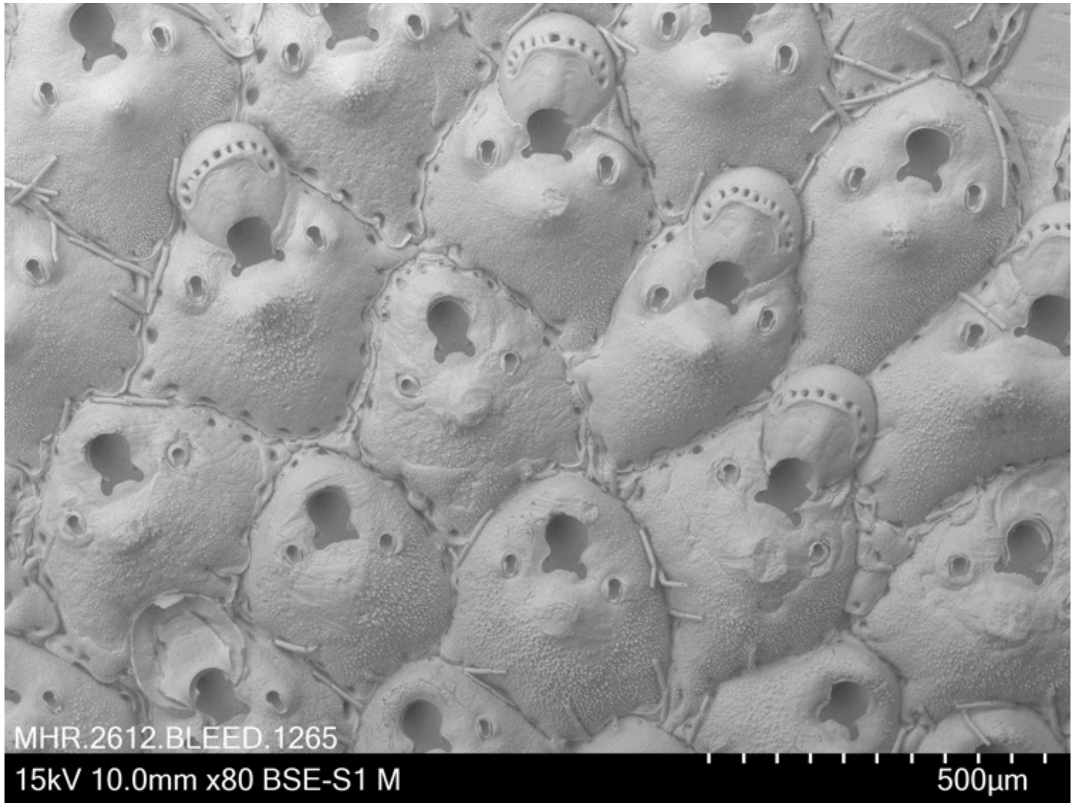


***Crepidacantha crinispina* BLEED 1265**
Coll: 16.3.2019 (A. M. Smith, D. P. Gordon)
Lat: -35.01
Long: 173.92
Depth: 17 m

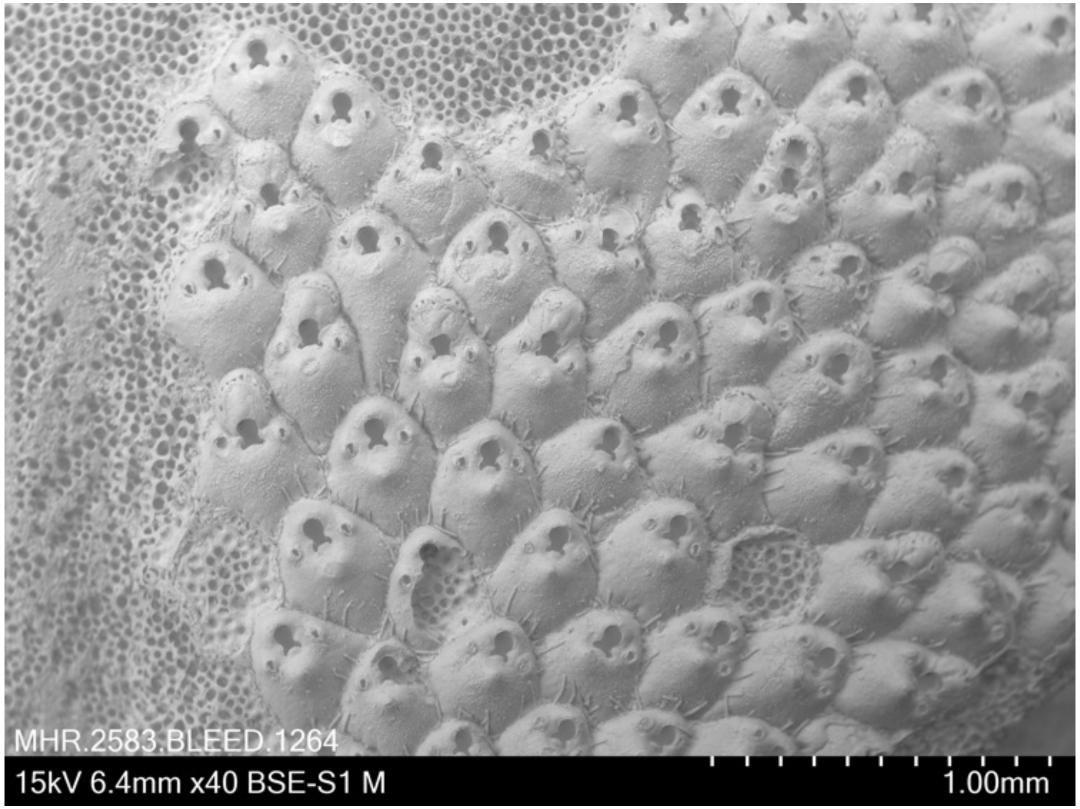


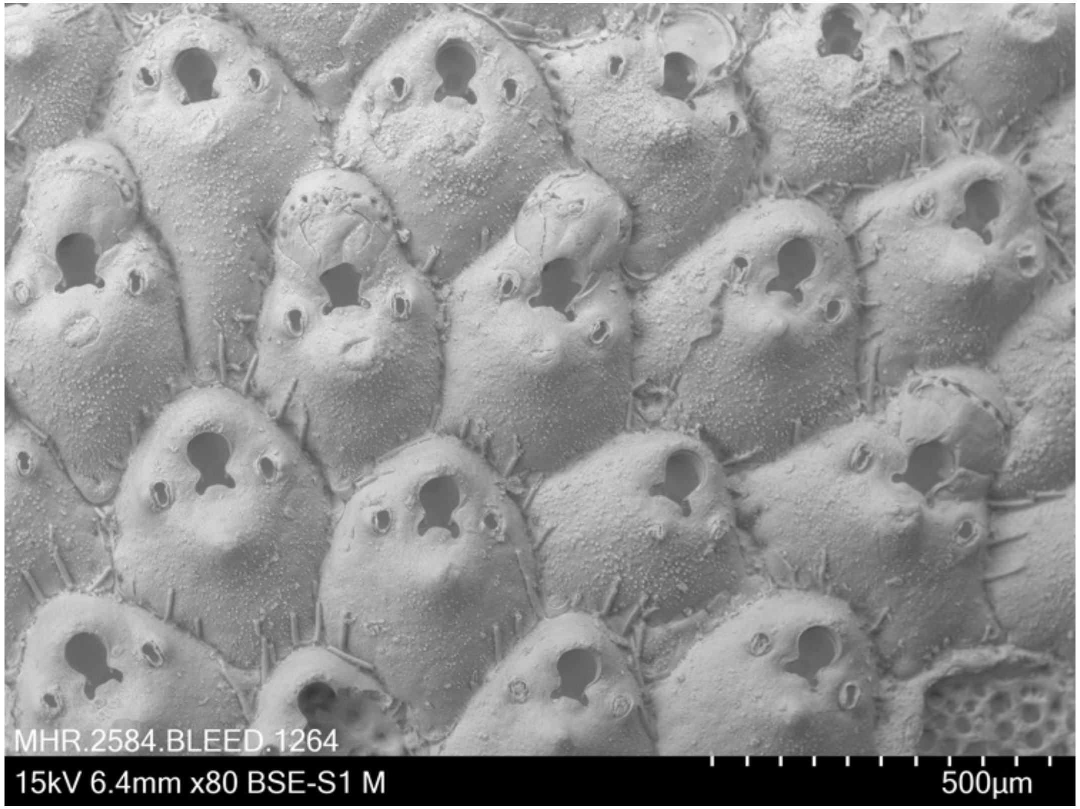


***Crepidacantha crinispina* BLEED 1264**
Coll: 16.3.2019 (A. M. Smith, D. P. Gordon)
Lat: -34.99
Long: 173.99
Depth: 81 m

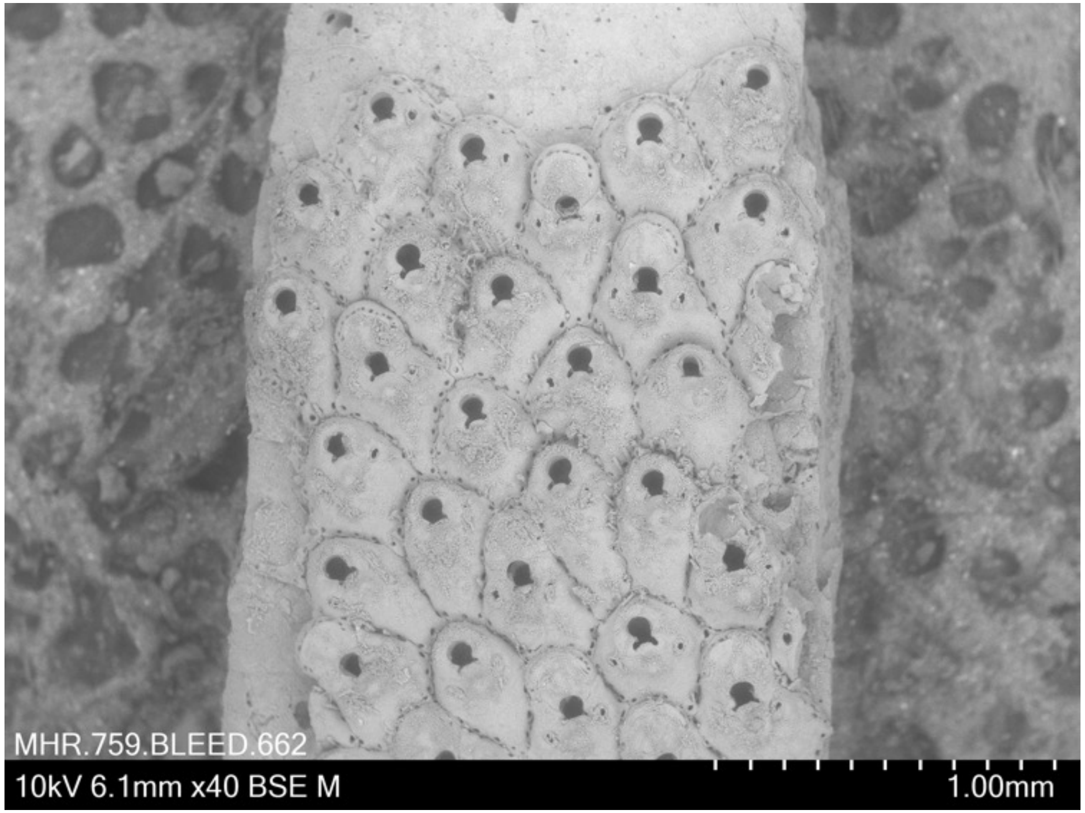

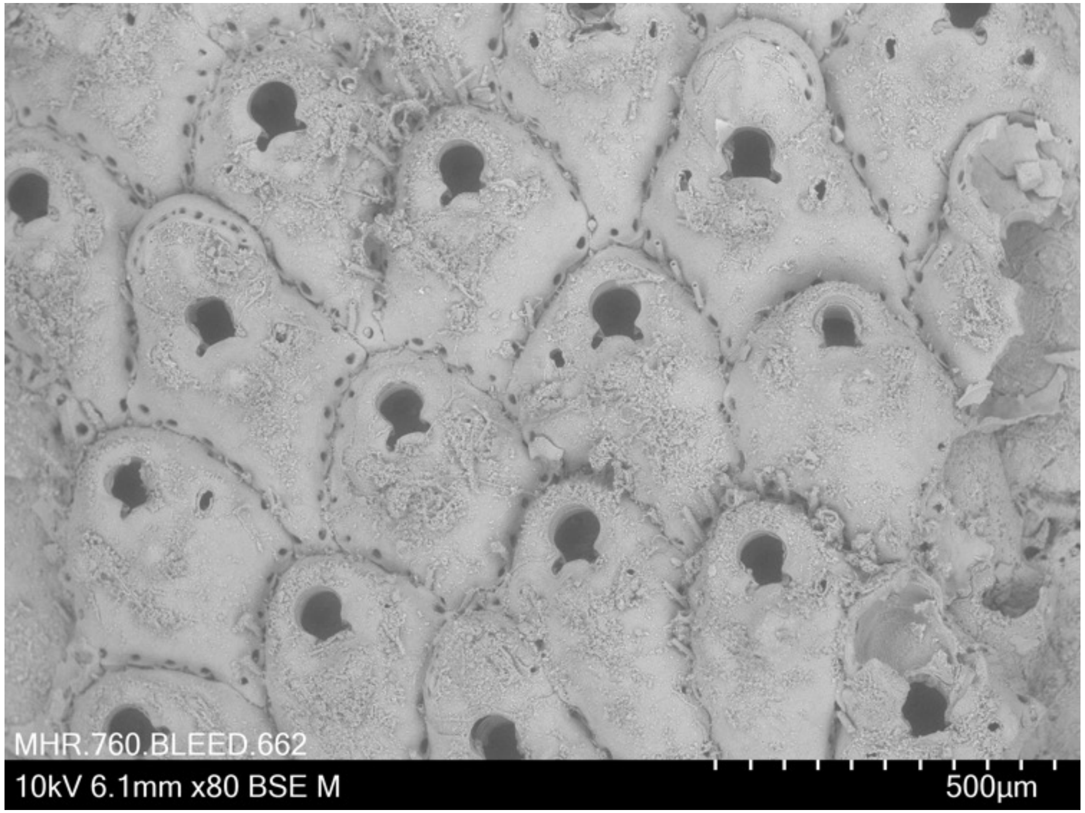


***Crepidacantha crinispina*** **BLEED 662**
Coll: 8.4.2018 (A. M. Smith, D. P. Gordon, H. Mello)
Lat: -47.13722222222222
Long: 168.12583333333333
Depth: 80-85 m

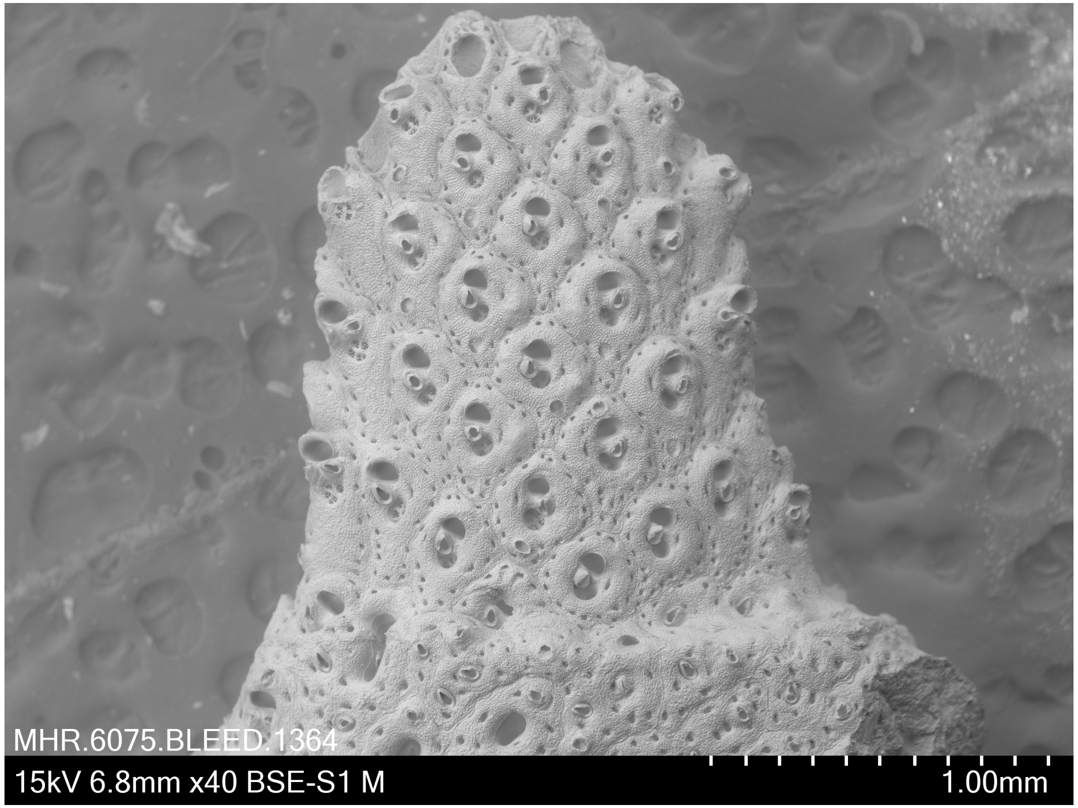


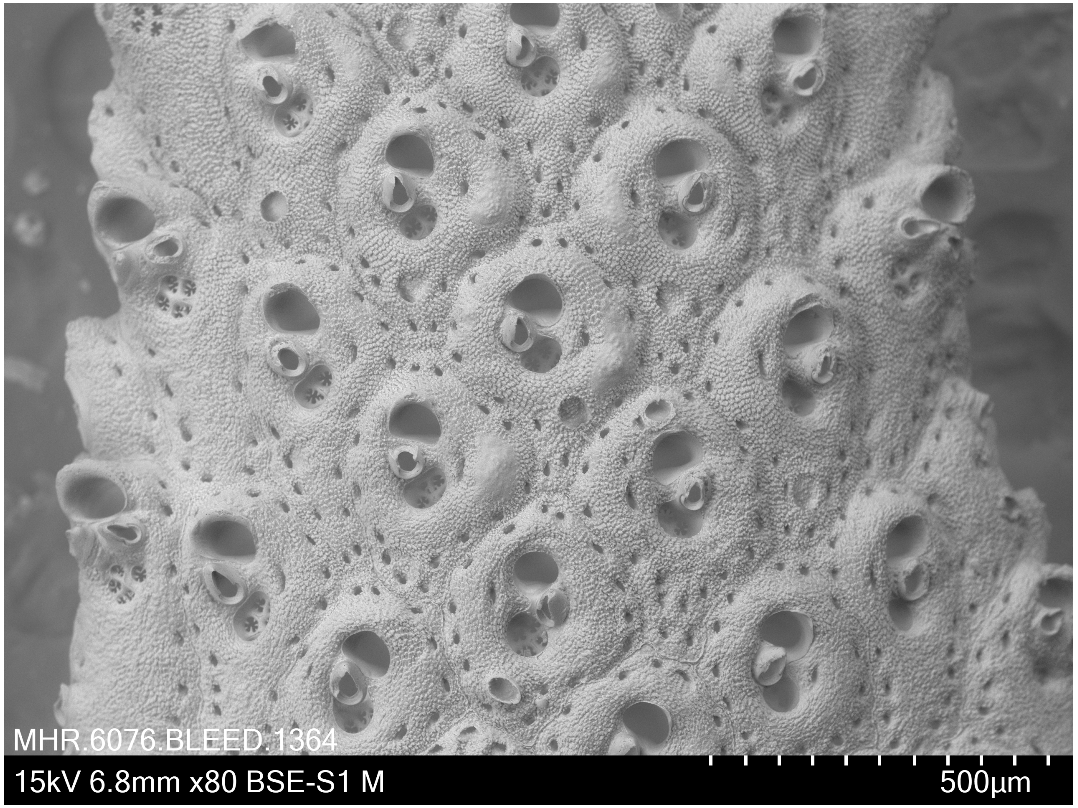


***Adeonellopsis gemina* BLEED 1364**
Coll: 17.3.2019 (A. M. Smith, D. P. Gordon)
Lat: -34.21
Long: 172.11
Depth: 104 m

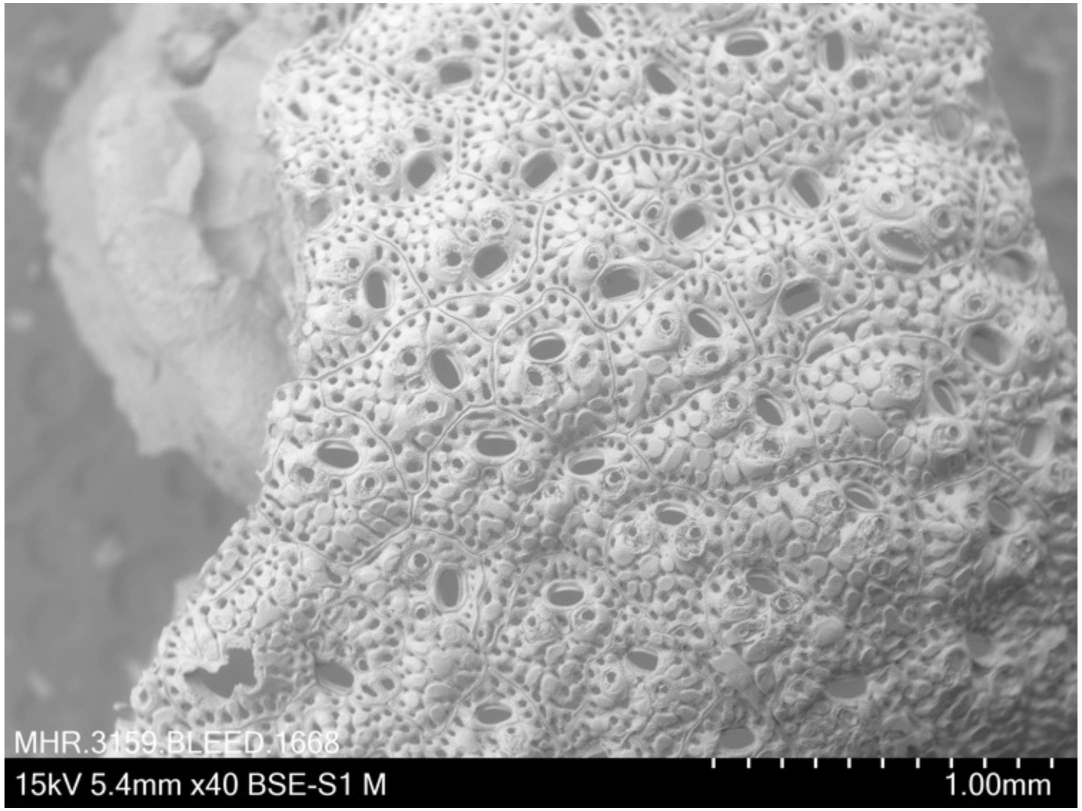

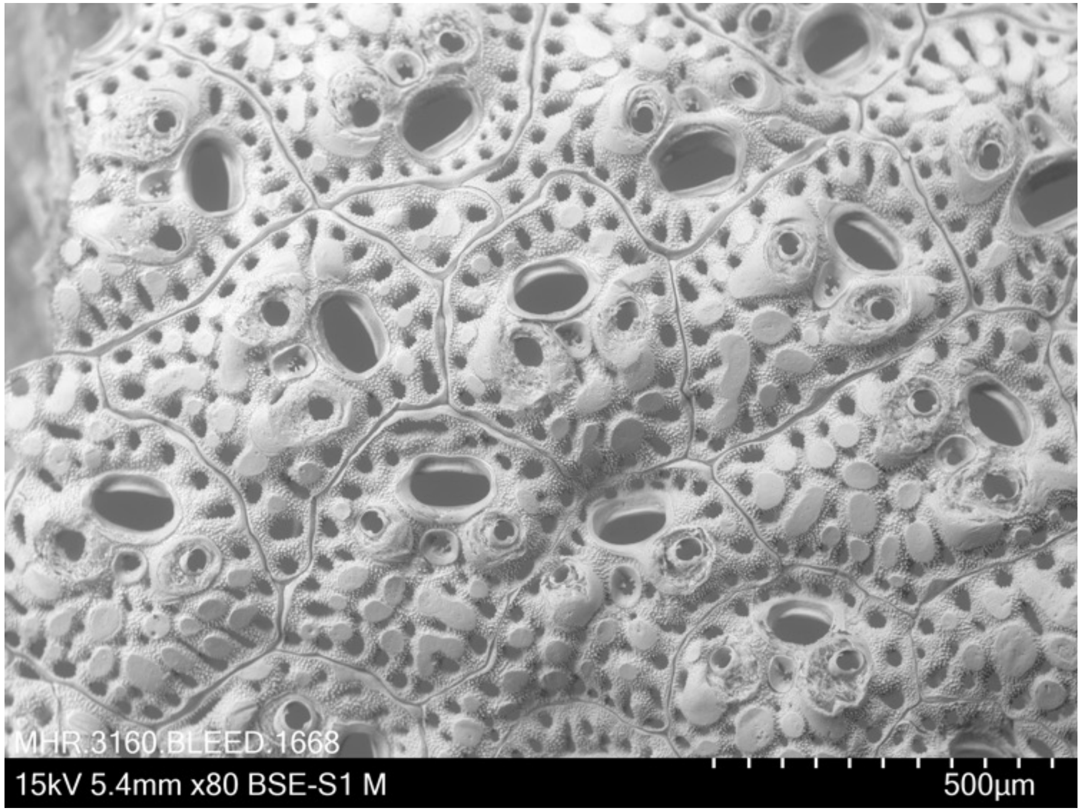


***Inversiula fertilis* BLEED 1668**
Coll: 18.3.2019 (A. M. Smith. D. P. Gordon)
Lat: -34.380000000000003
Long: 172.69
Depth: 52.5-56.5m

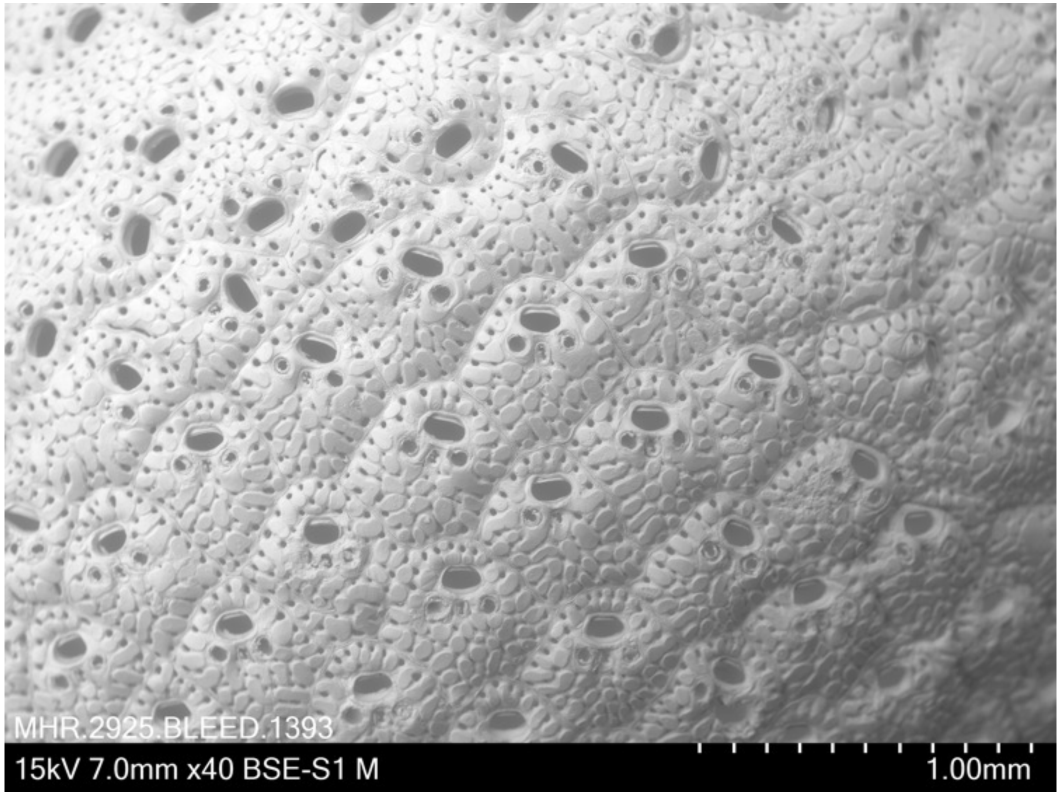


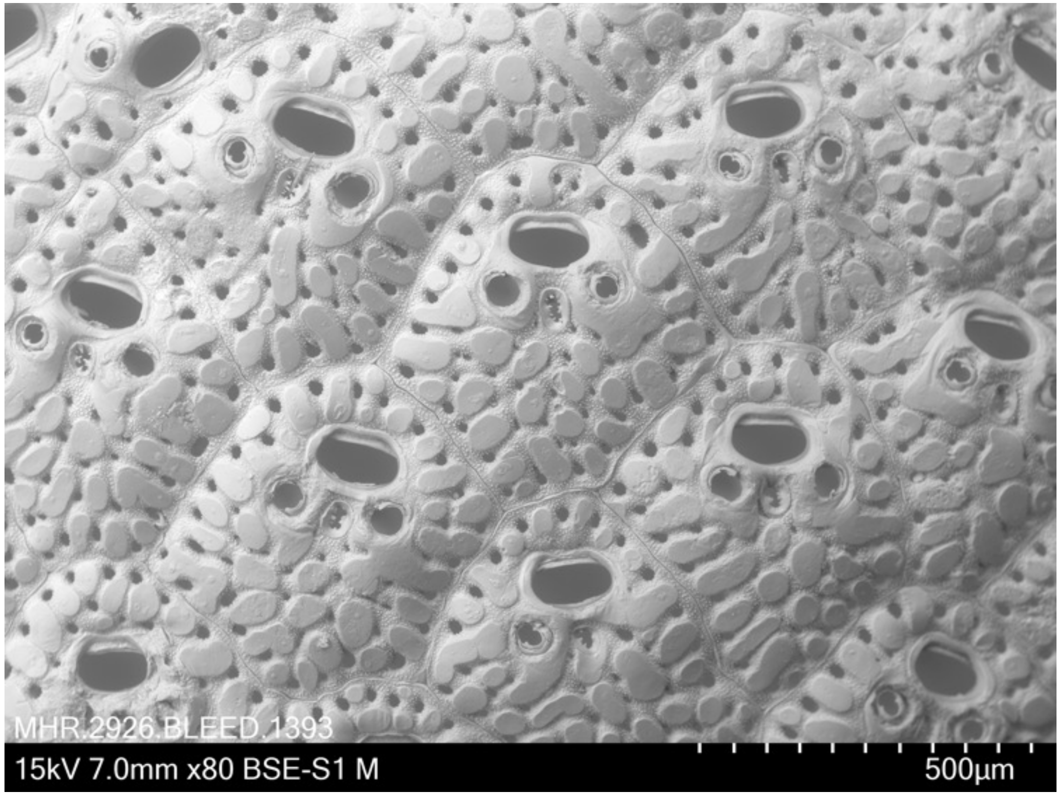


***Inversiula fertilis* BLEED 1393**
Coll: 17.3.2019 (A. M. Smith; D. P. Gordon)
Lat: -34.159999999999997
Long: 172.09
Depth: 82.9 m

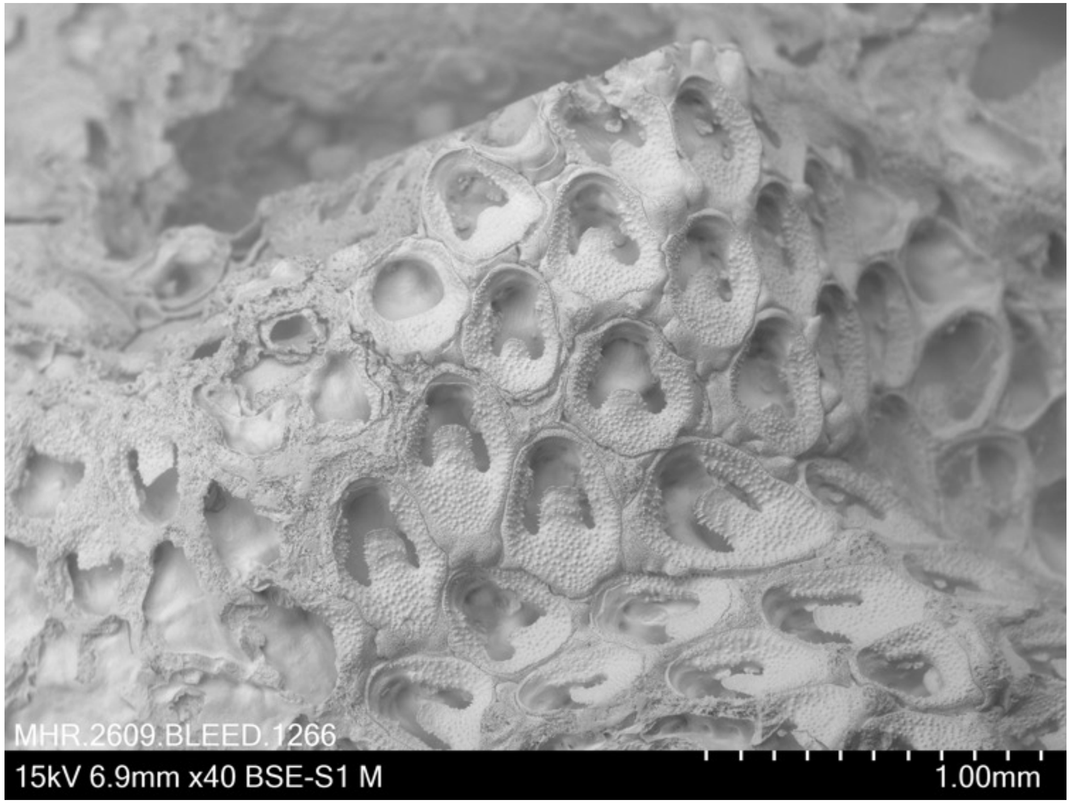


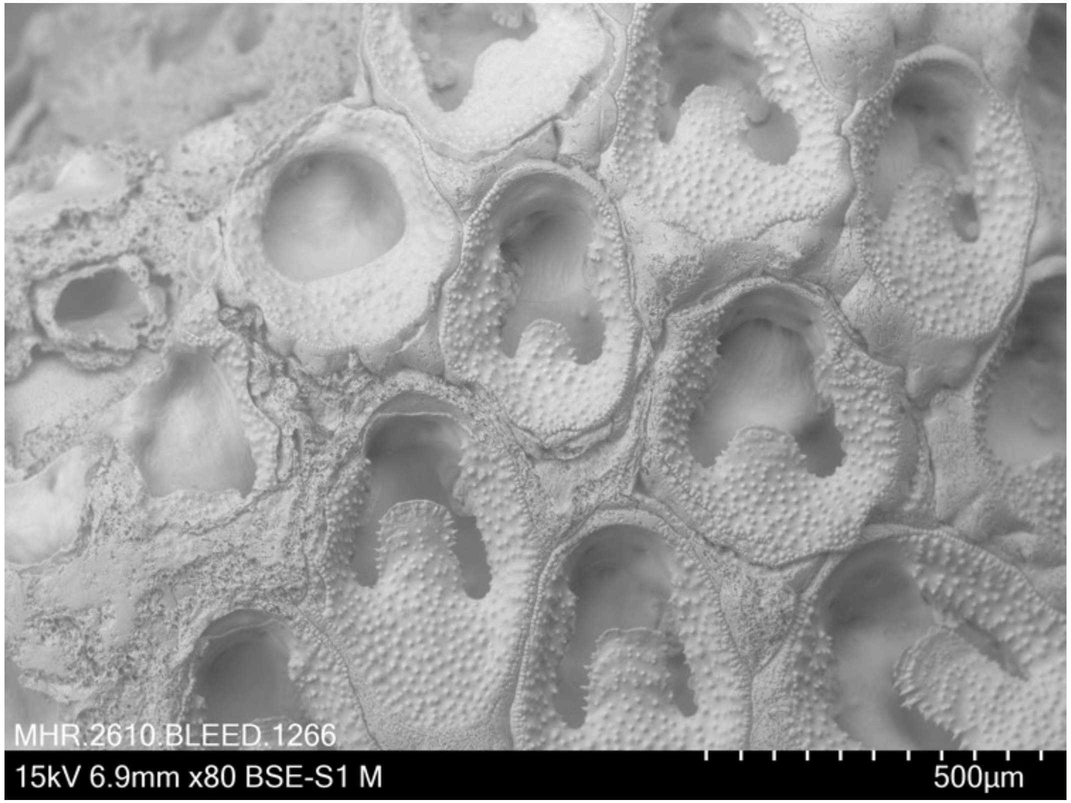


***Caleschara minuta* BLEED 1266**
Coll: 16.3.2019 (A. M. Smith, D. P. Gordon)
Lat: -35.01
Long: 173.92
Depth: 17 m

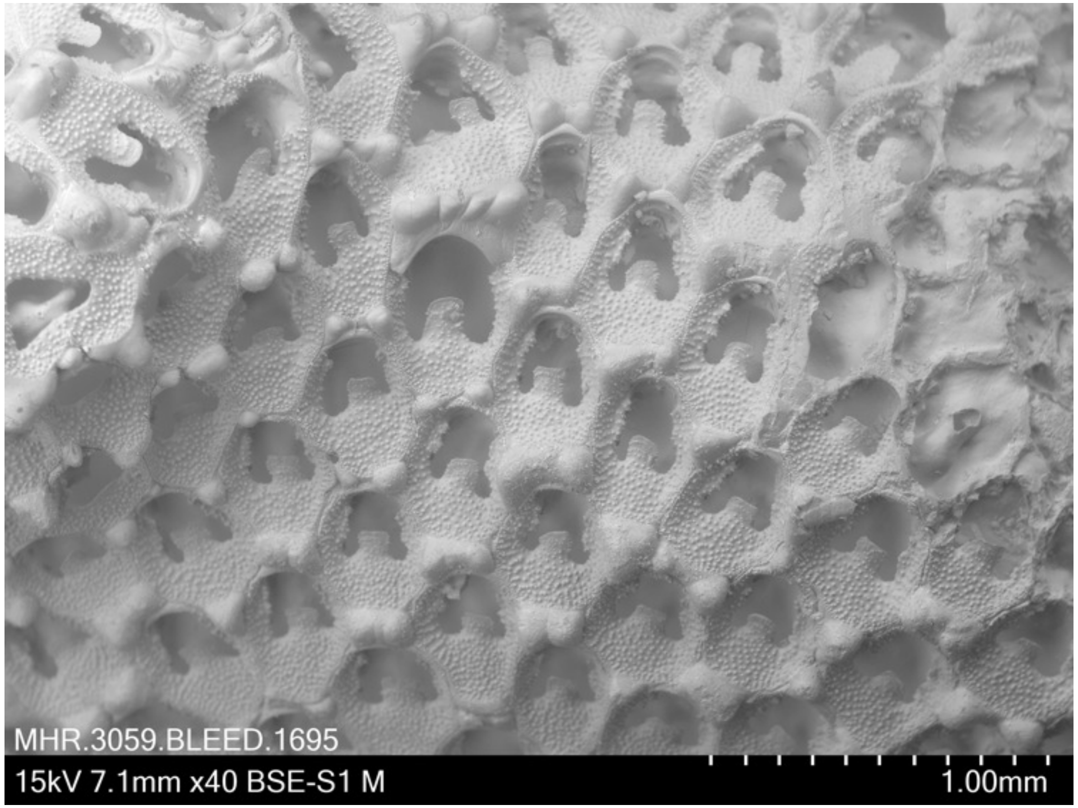


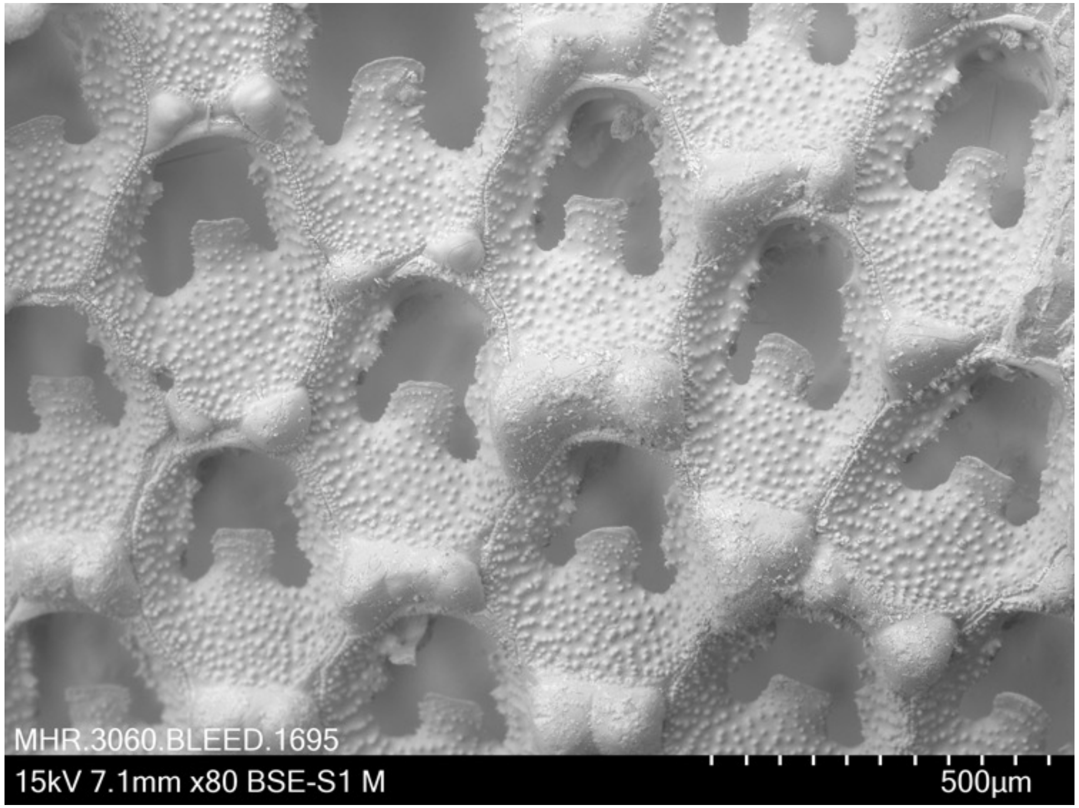


***Caleschara minuta* BLEED 1695**
Coll: 19.3.2019 (A. M. Smith. , D. P. Gordon)
Lat: -34.99
Long: 173.97
Depth: 61.6 m

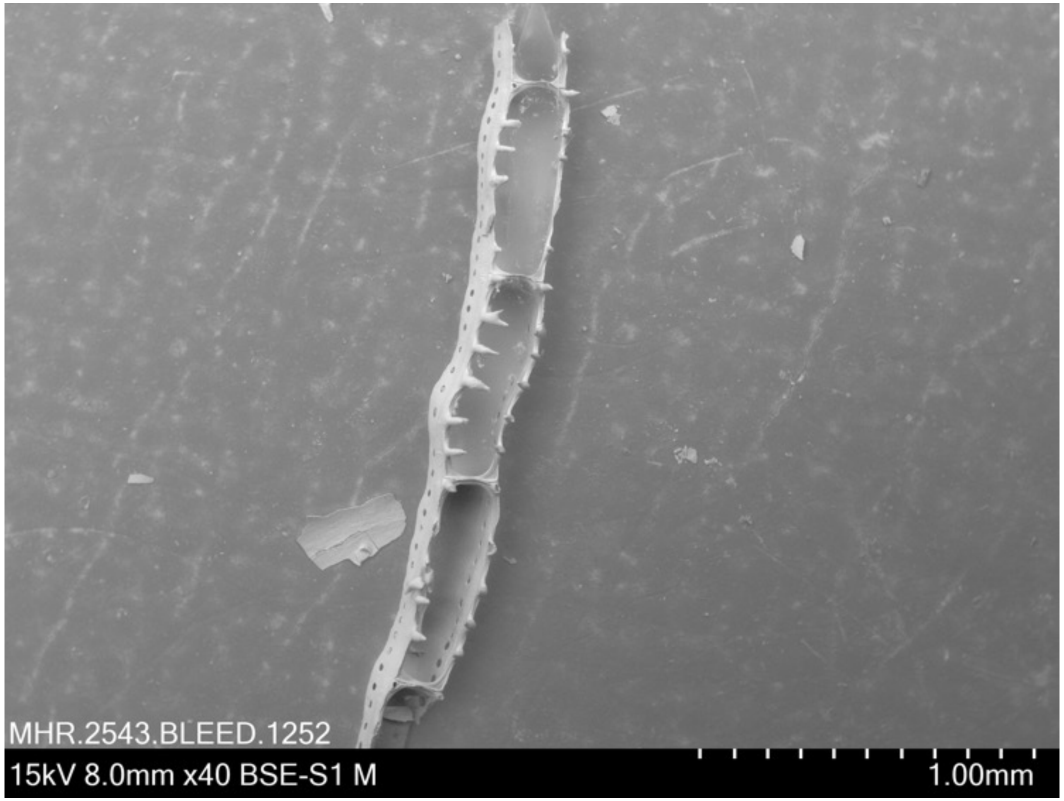


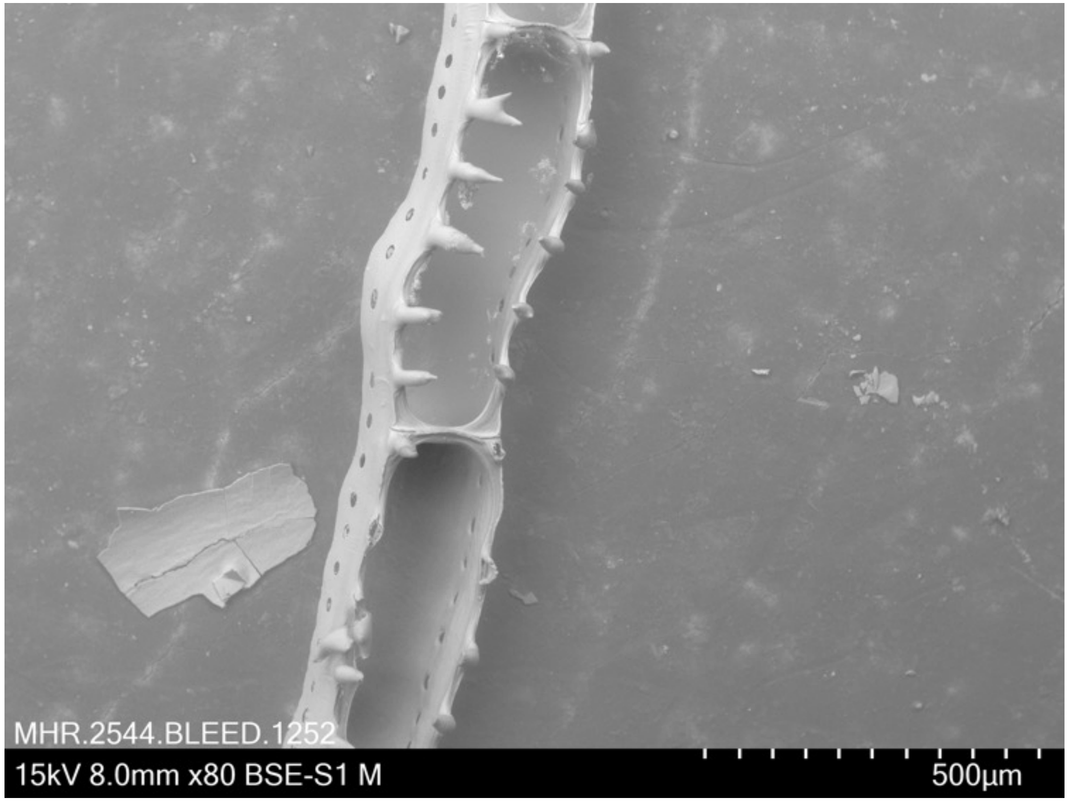


***Hincksina* n. sp. BLEED 1252**
Coll: 15.3.2019 (A. M. Smith, D. P. Gordon)
Lat: -36.19
Long: 174.99
Depth: 52.3 m
**Notes:** This is the same species as BLEED 61 published in Orr *et al.* 2020. Compare *Hincksina* Norman, 1903 with *Gregarinidra* Barroso, 1949 with which it may be synonymized.

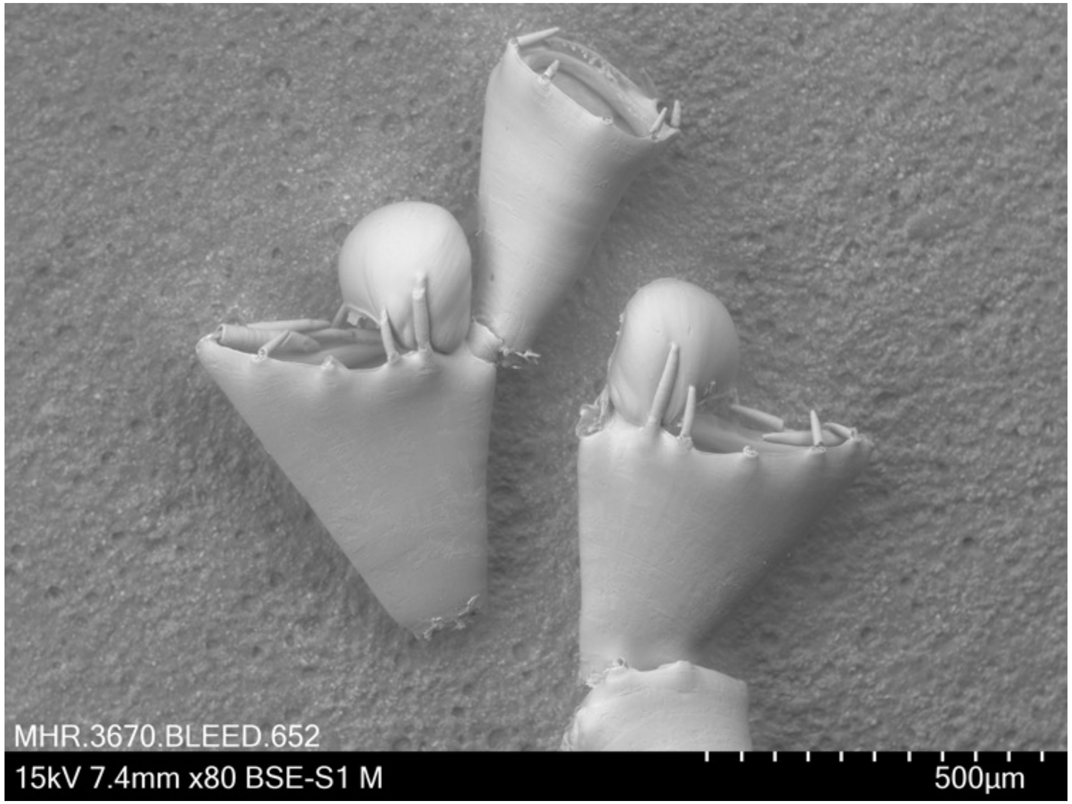


***Dimetopia barbata*** **BLEED 652**
Coll: 8.4.2018 (A. M. Smith, D. P. Gordon, H. Mello)
Lat: -47.17361111111111
Long: 167.94583333333333
Depth: 66 m

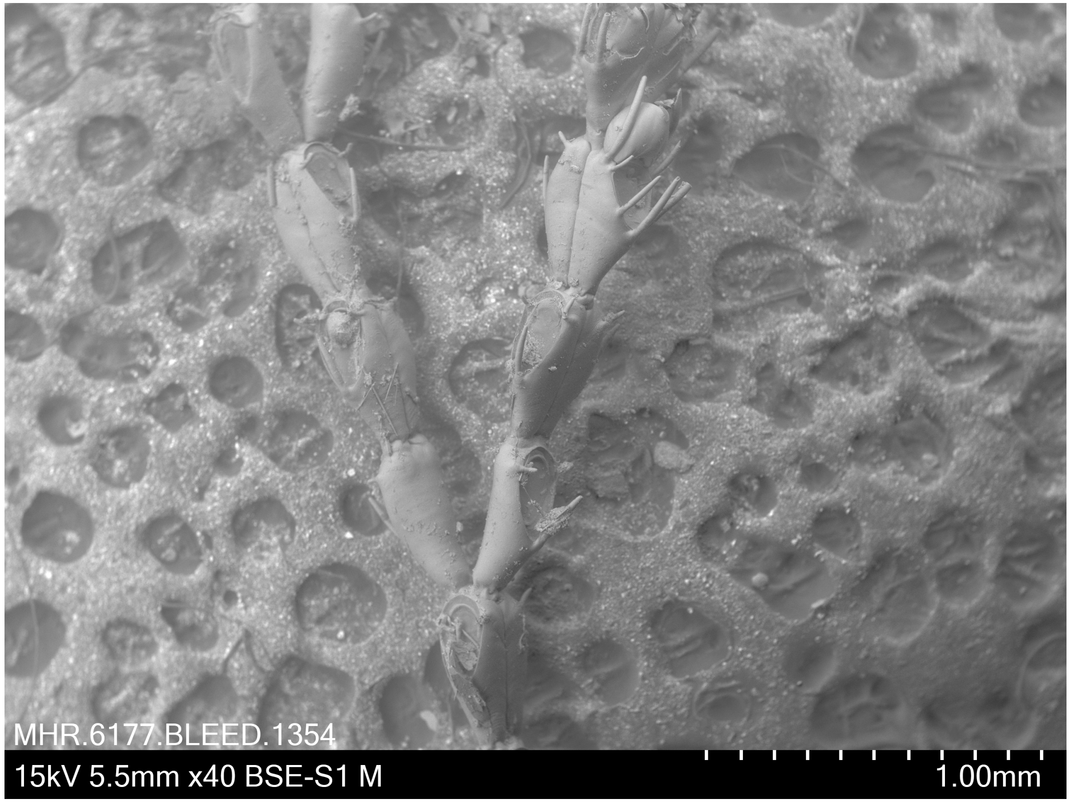


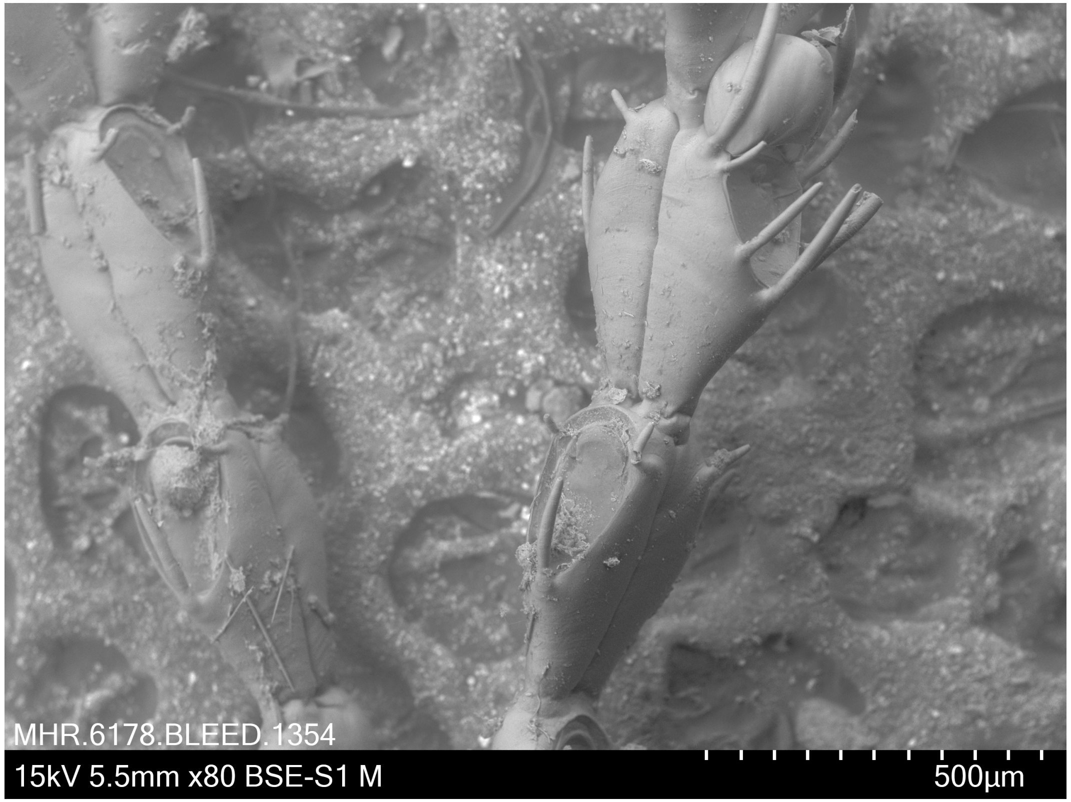


***Dimetopia cornuta* BLEED 1354**
Coll: 17.3.2019 (A. M. Smith; D. P. Gordon)
Lat: -34.21
Long: 172.11
Depth: 104 m

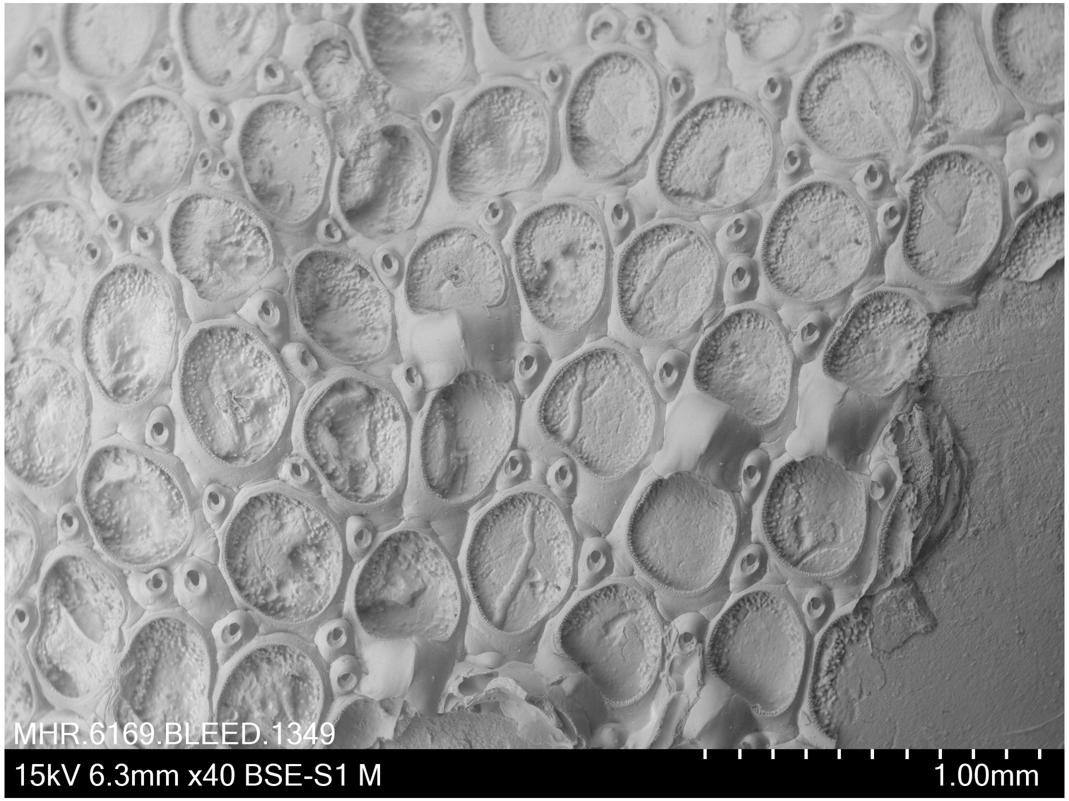


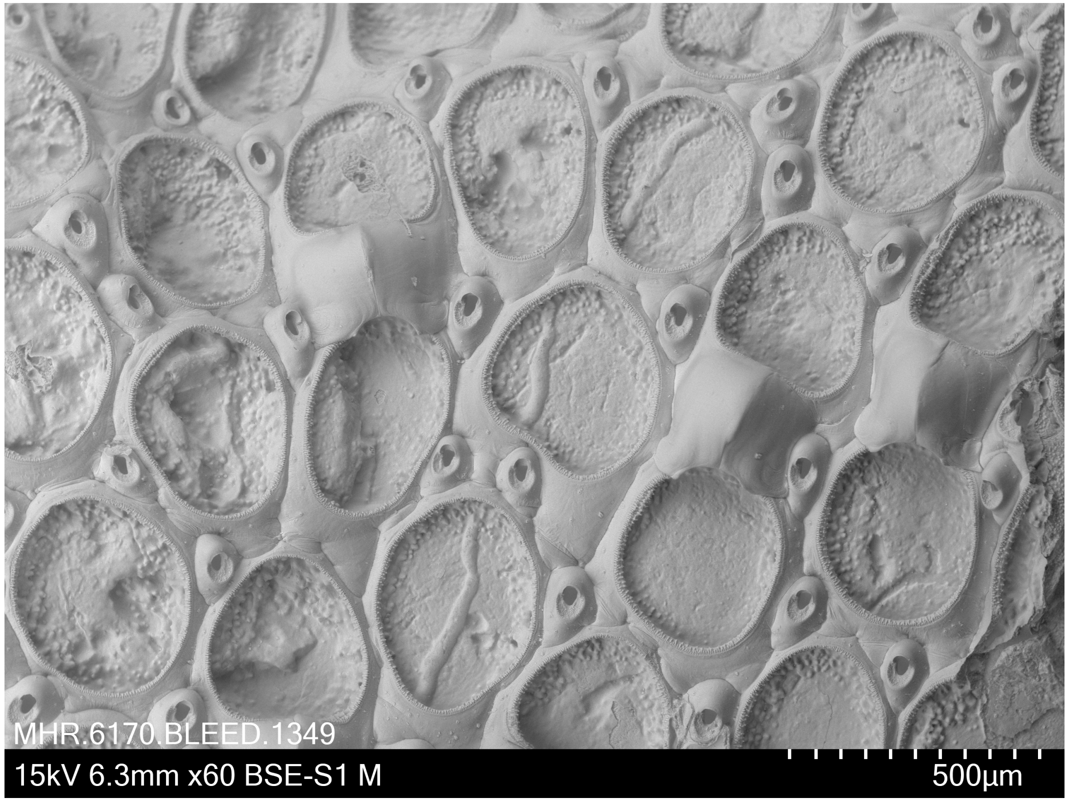


**'*Retevirgula*' *aggregata* BLEED 1349**
Coll: 17.3.2019 (A. M. Smith; D. P. Gordon)
Lat: -34.21
Long: 172.11
Depth: 104 m

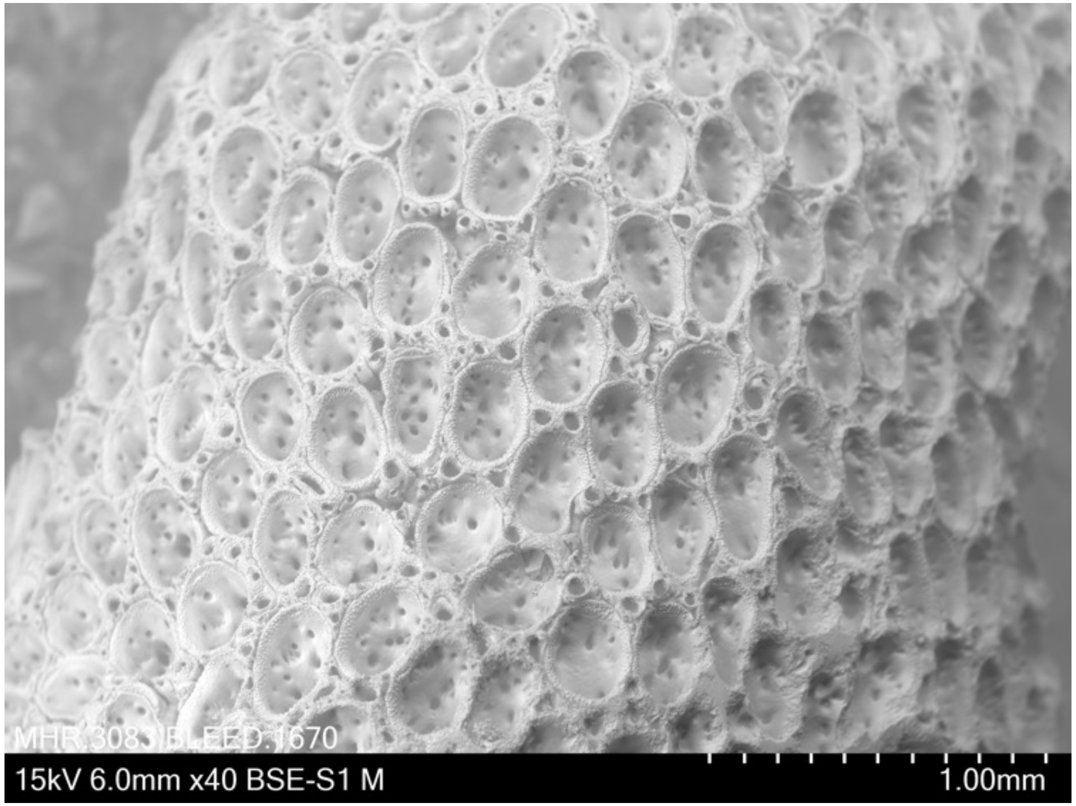


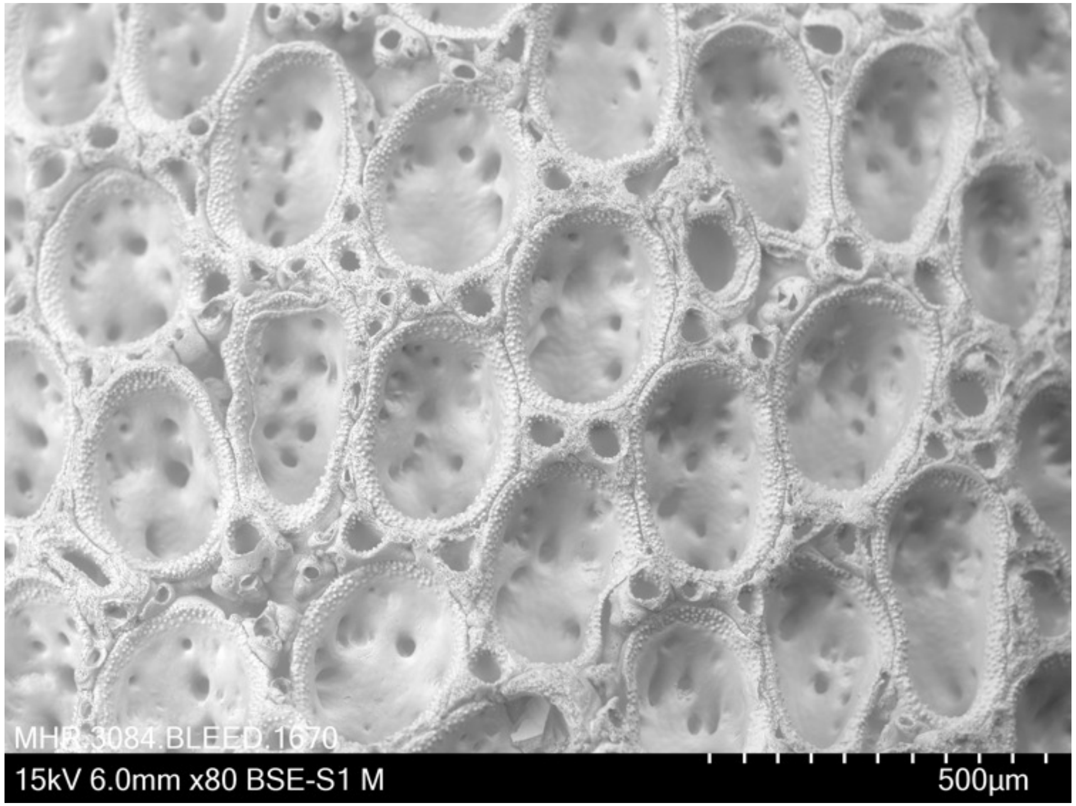


***Akatopora circumsaepta*** **BLEED 1670**
Coll: 18.3.2019 (A. M. Smith, D. P. Gordon)
Lat: -34.380000000000003
Long: 172.69
Depth: 52.5-56.5m

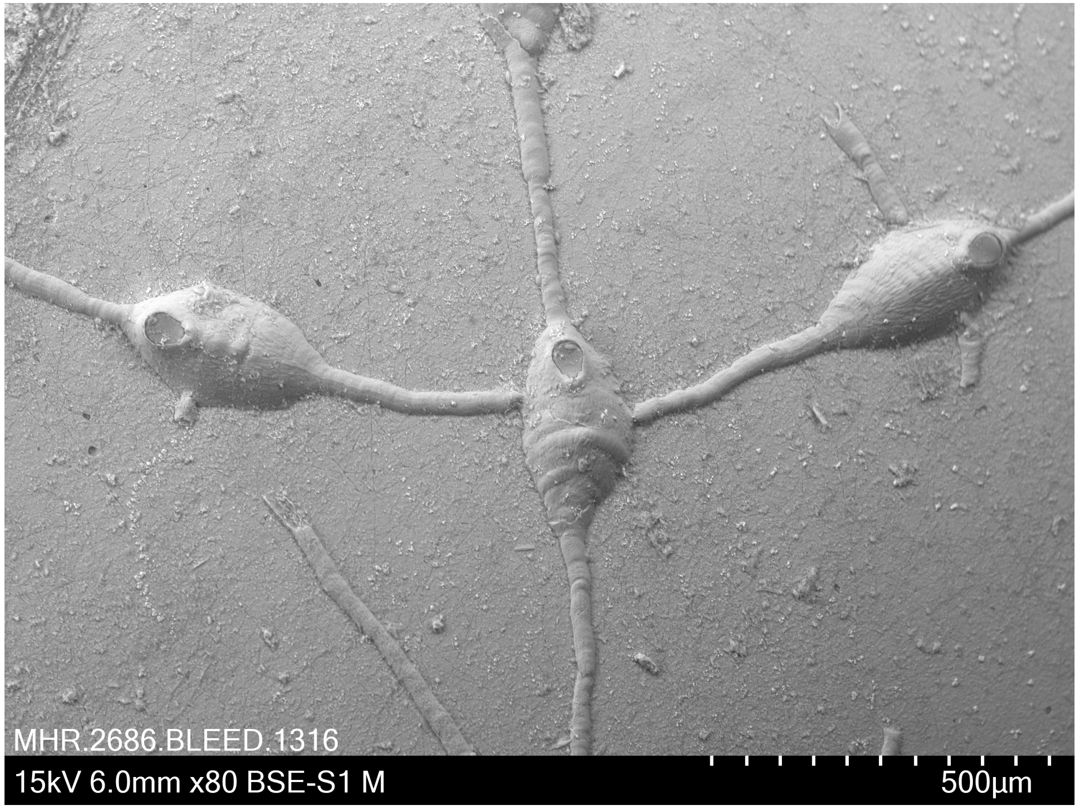


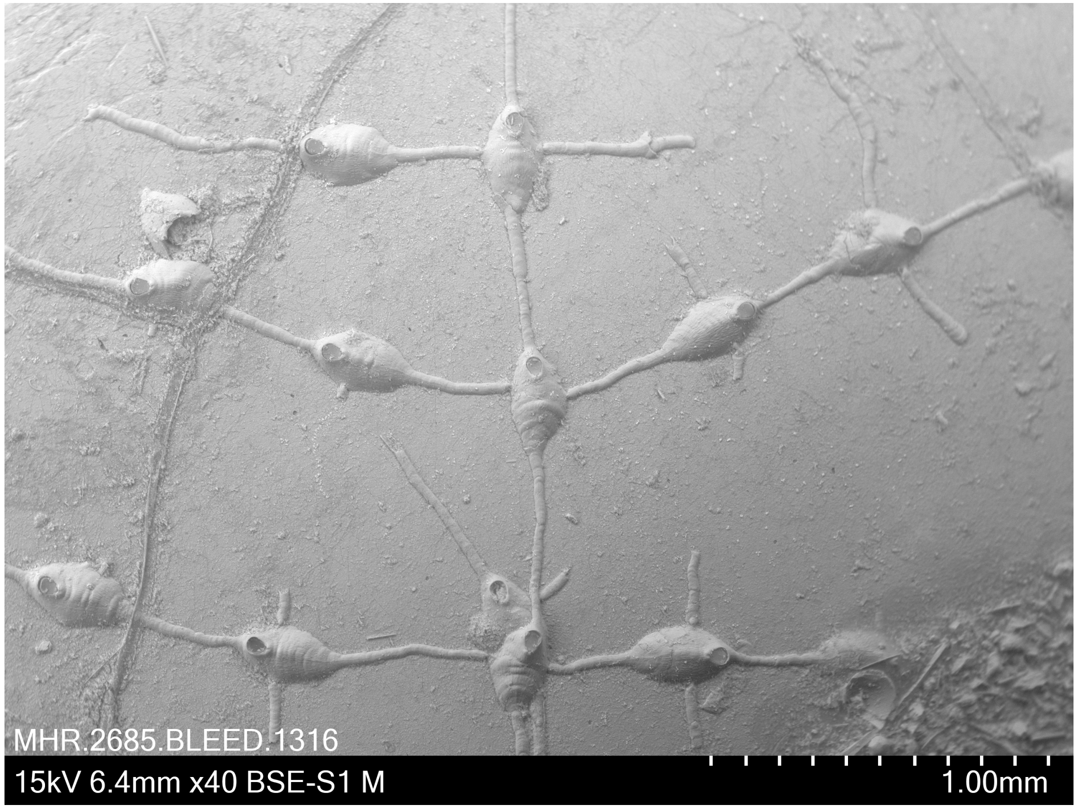


***Hippothoa watersi* BLEED 1316**
Coll: 15.3.2019 (A. M. Smith, D. P. Gordon)
Lat: -36.46
Long: 174.9
Depth: 37 m

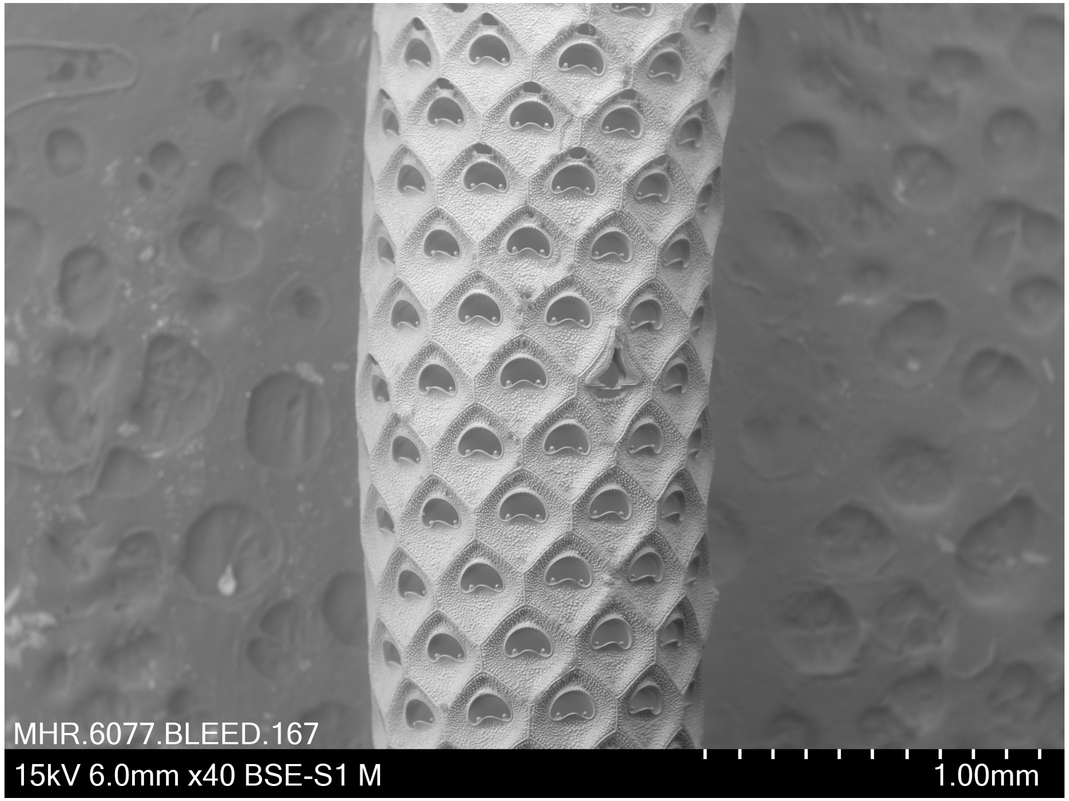


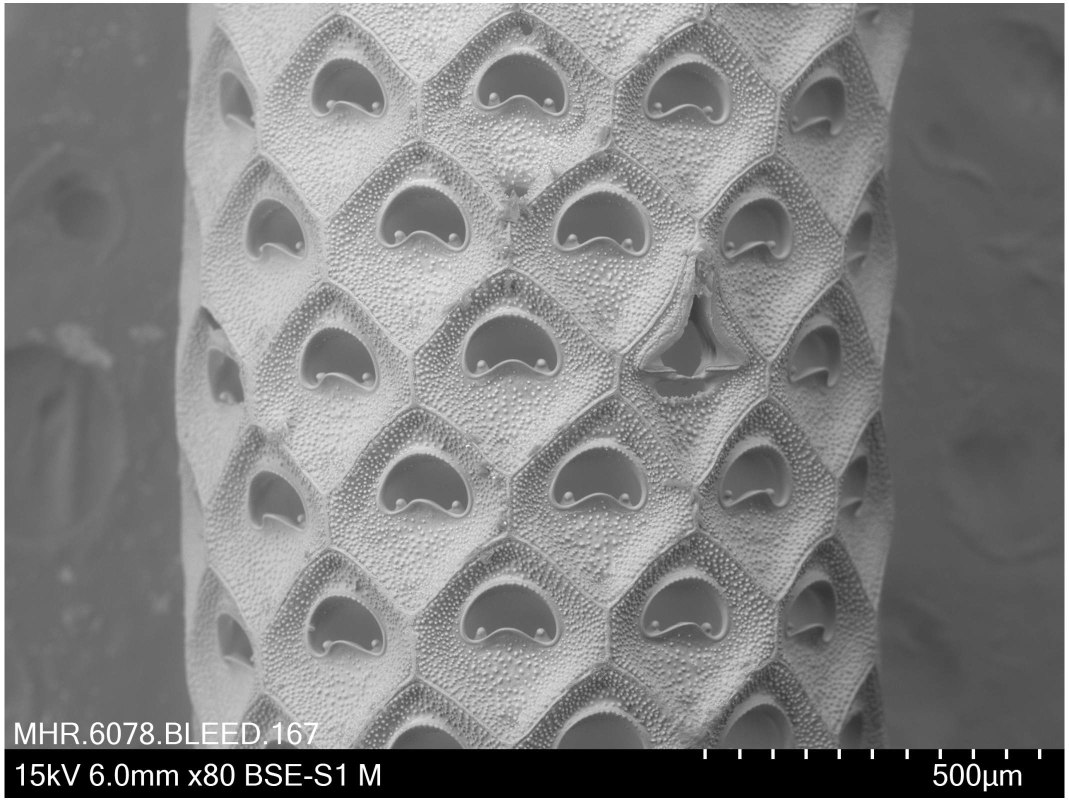


***Cellaria immersa*** **BLEED 167**
NIWA 103458 (KAH1503/8)
Coll: 30.03.2015
Lat: -40.7146667
Long: 173.457333
Depth: 51 m

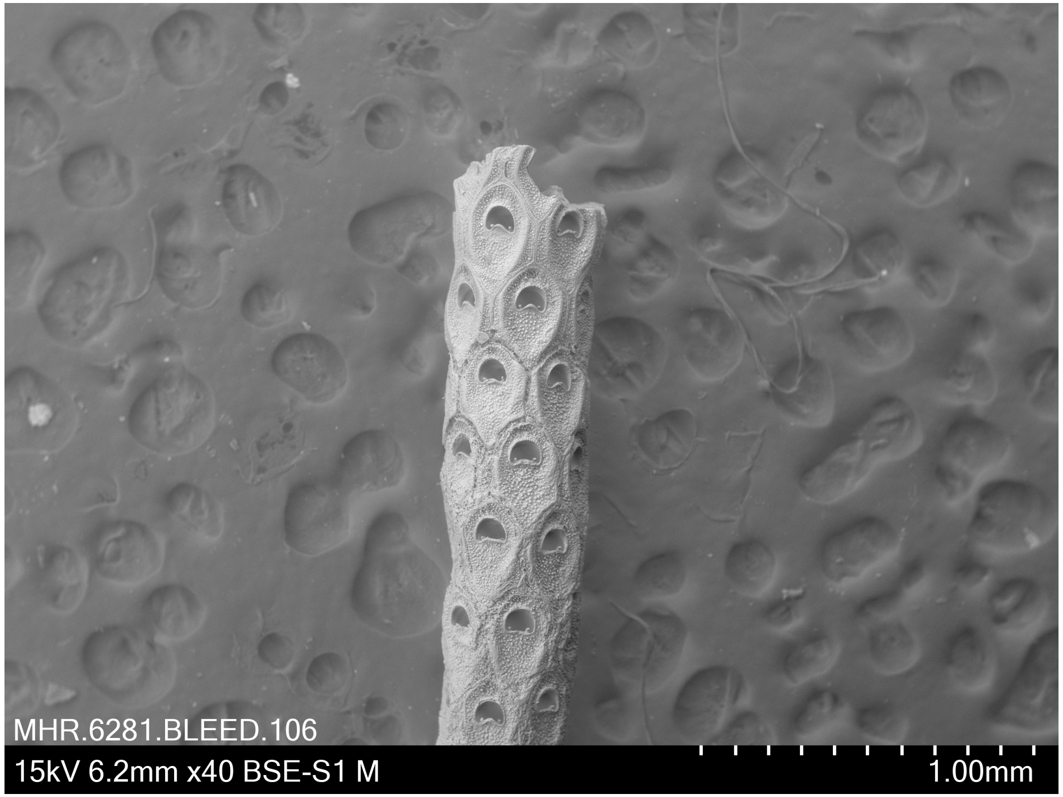


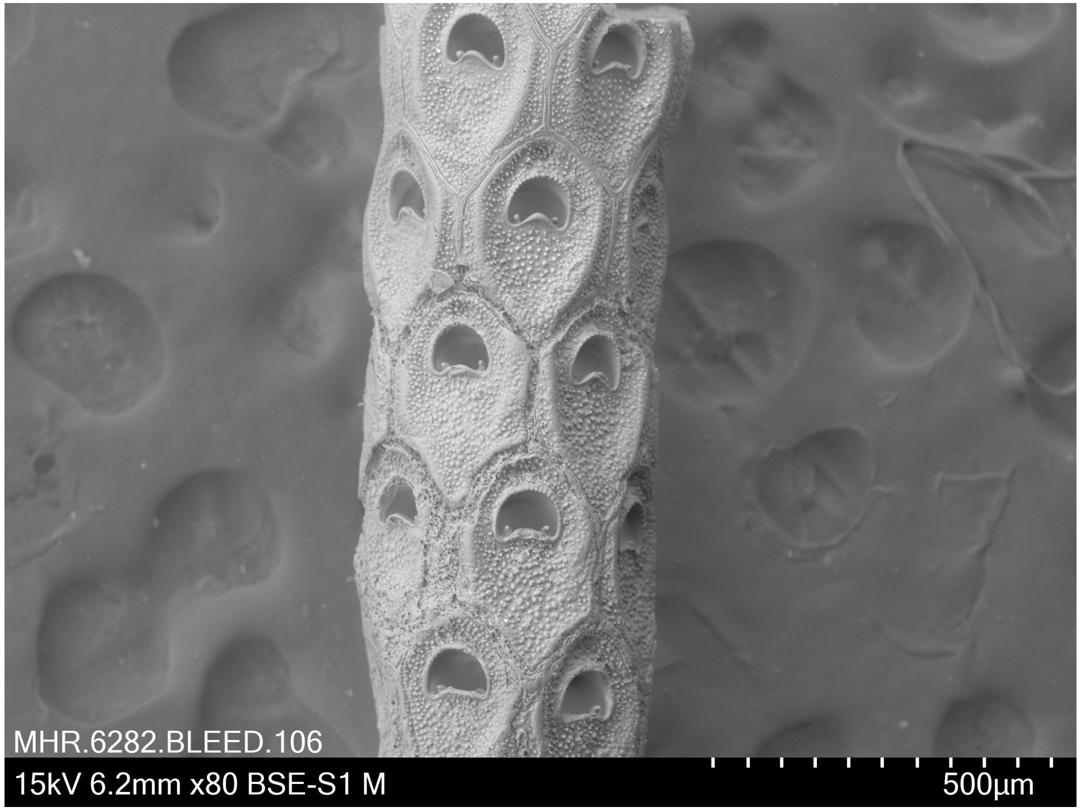


***Cellaria tenuirostris* BLEED 106**
Coll: 20.5.2015 (D. P. Gordon)
Lat: -40.900
Long: 176.217
Depth: intertidal

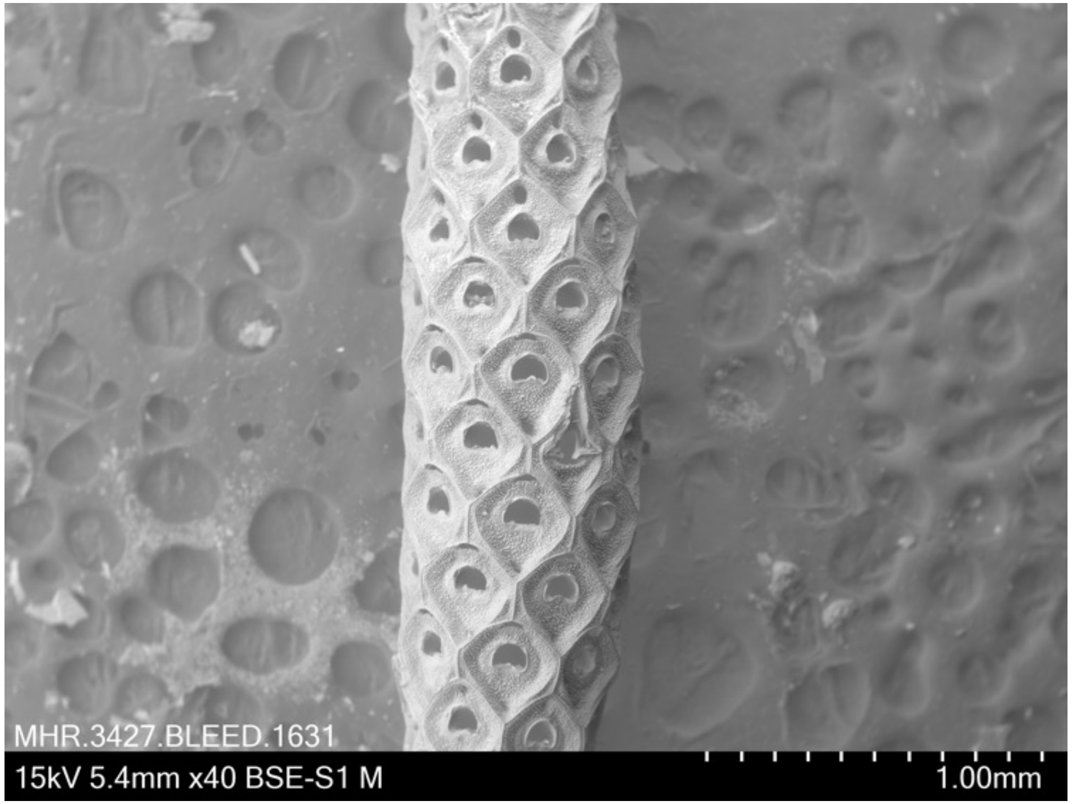


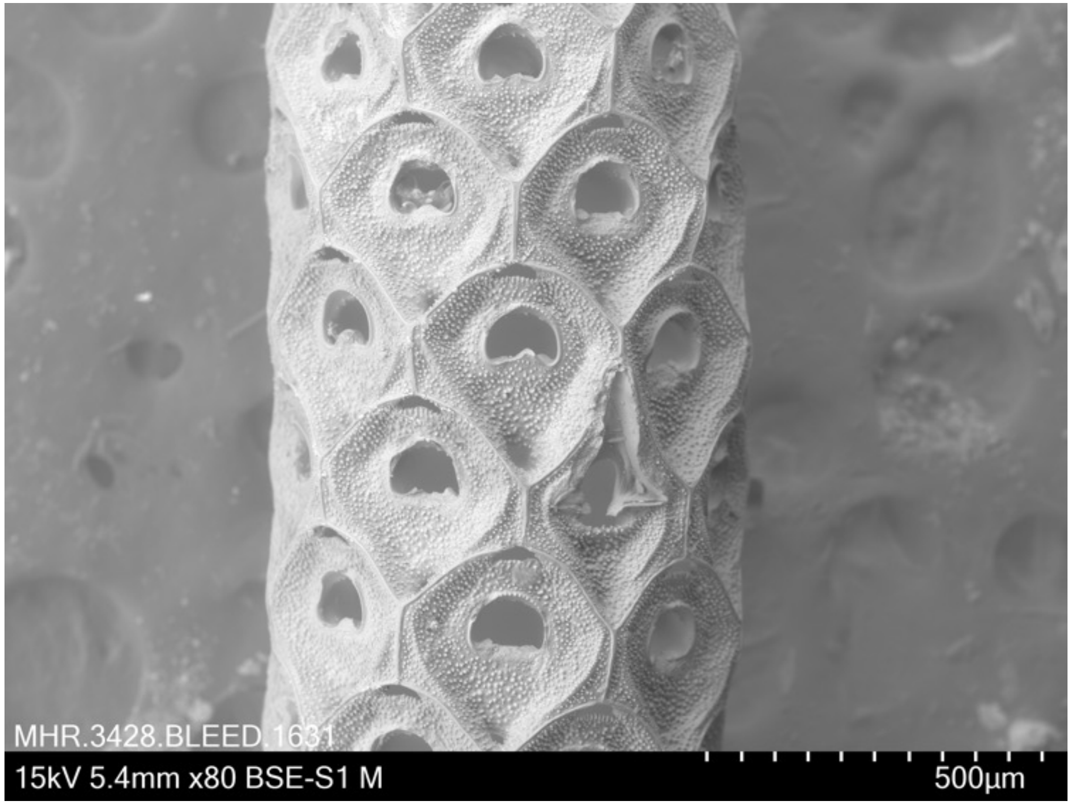


***Cellaria immersa* BLEED 1631**
Coll: 17.3.2019 (A. M. Smith; D. P. Gordon)
Lat: -34.159999999999997
Long: 172.09
Depth: 82.9 m

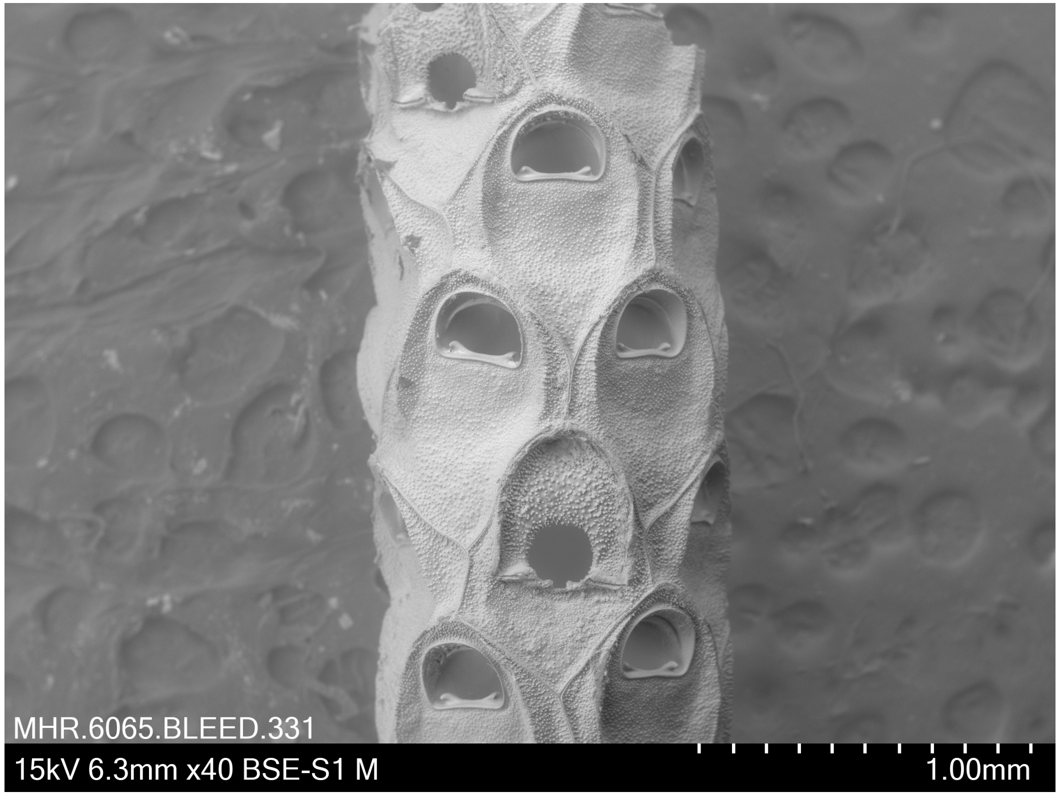


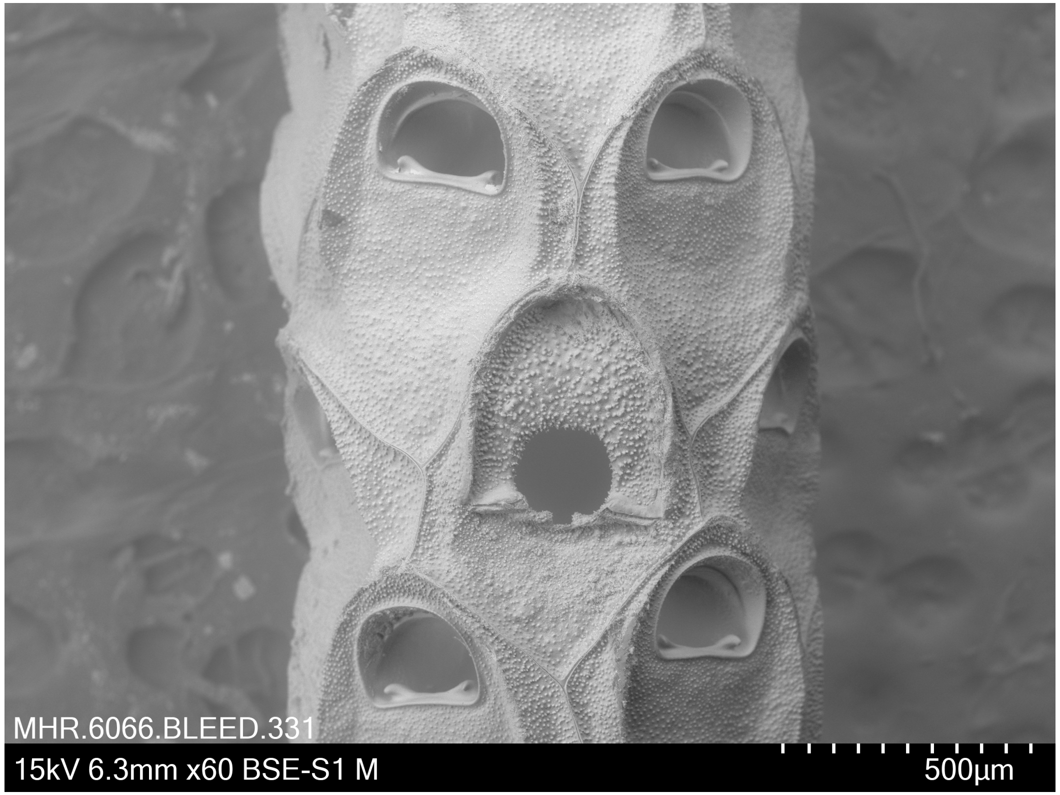


***Steginocellaria magnimandibulata* BLEED 331**
NIWA 95863 TAN 1312/D21
Coll: 11.11.2013
Lat: -33.4463
Long: 169.327
Depth: 1925 m

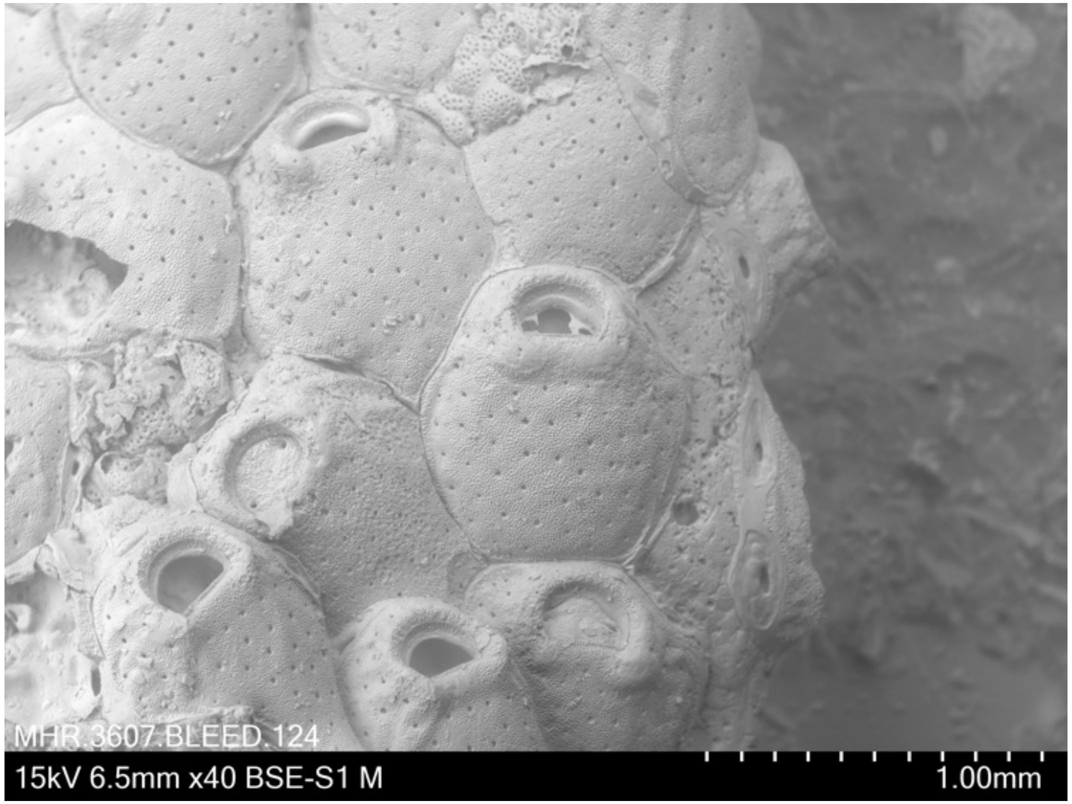


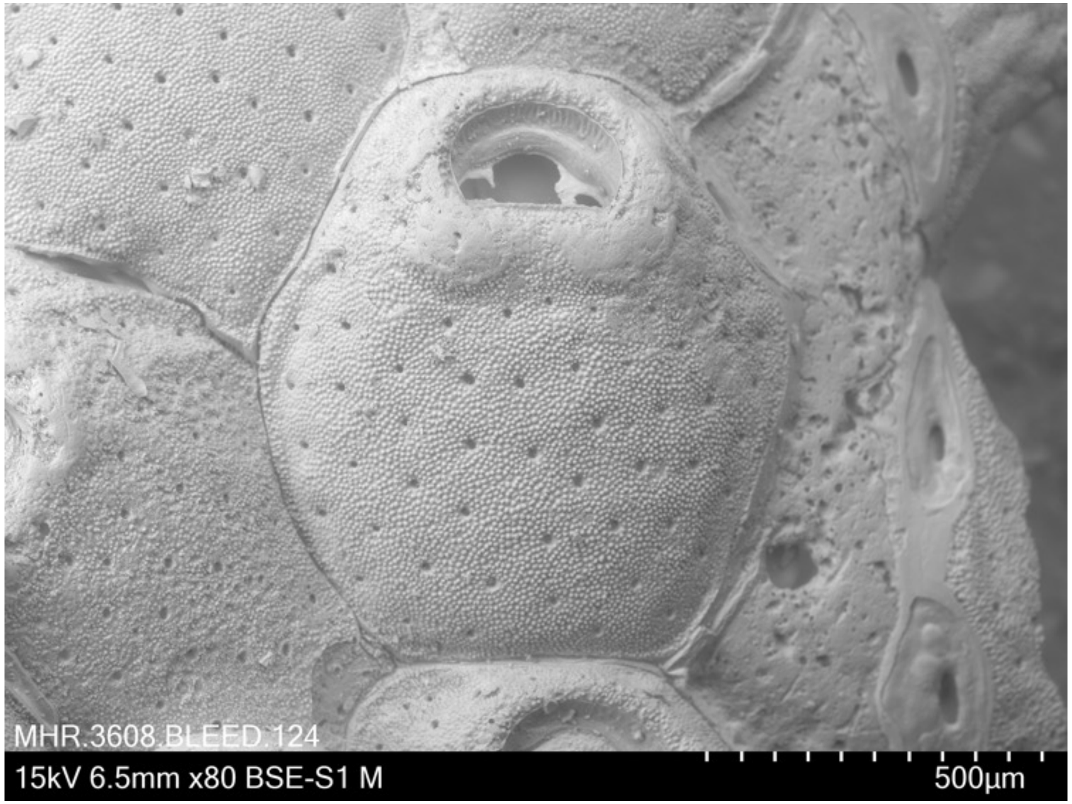


***Macropora nodulosa*** **BLEED 124**
Coll: 26.1.2010 (A. M. Smith)
Lat: -47.28
Long: 167.67
Depth: 100 m

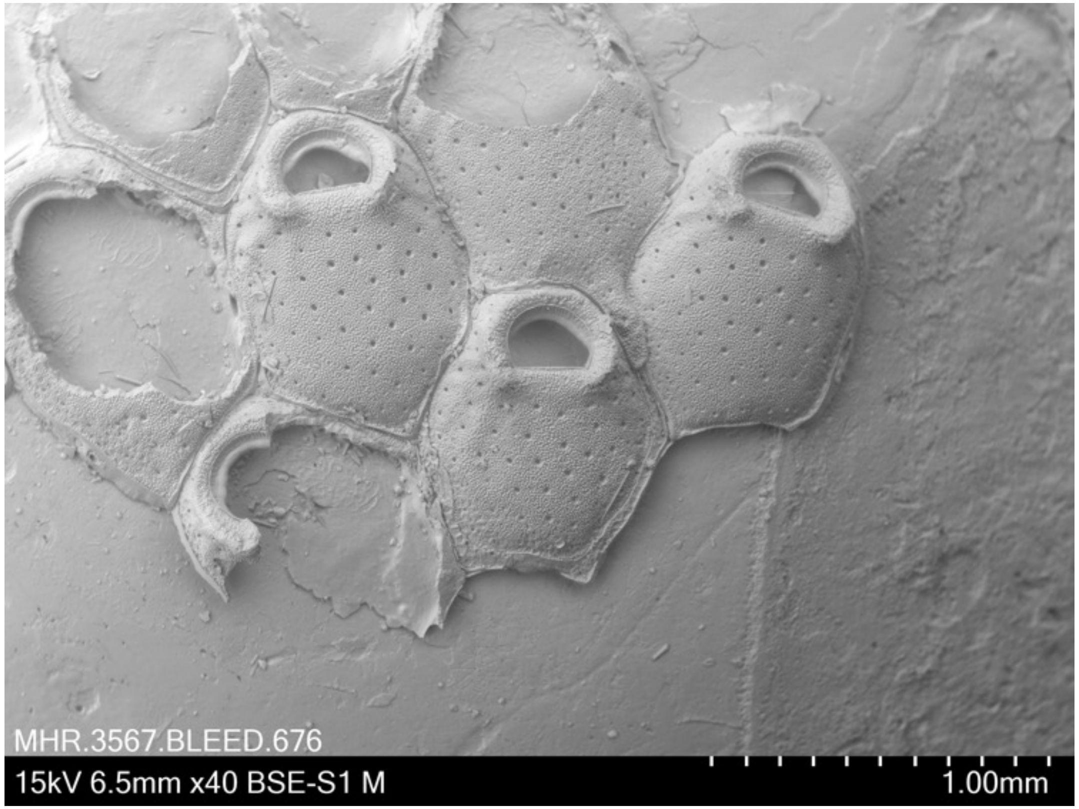


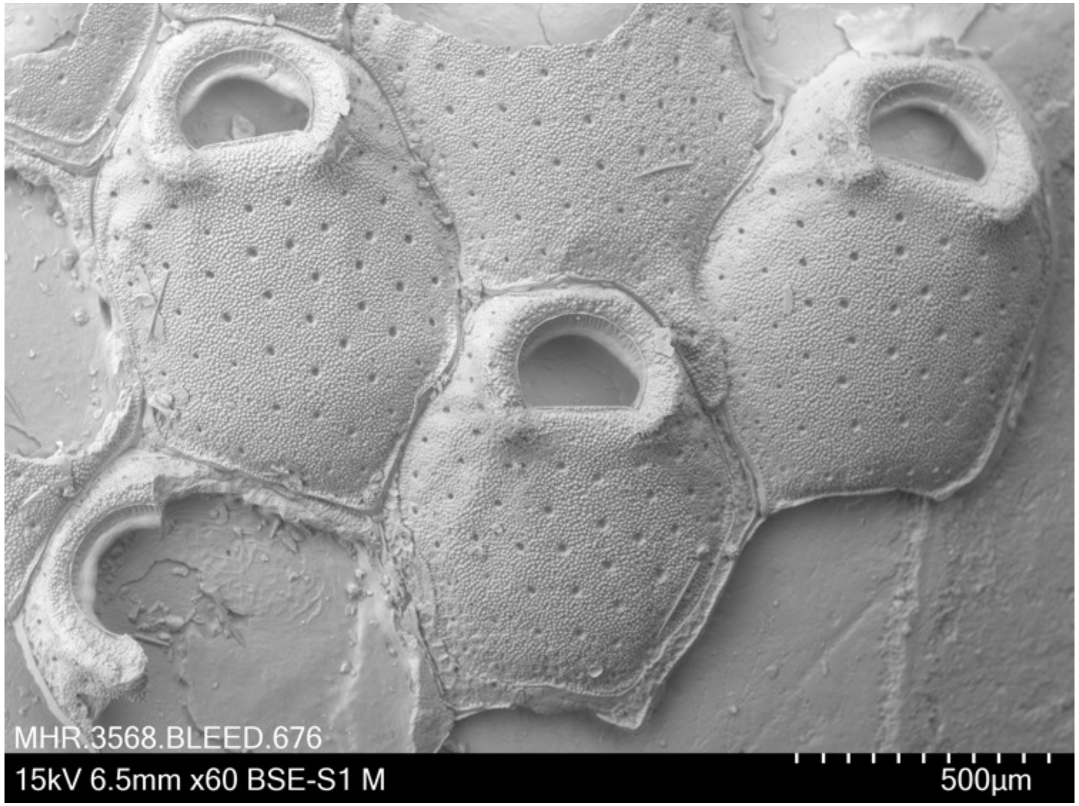


***Macropora nodulosa* BLEED 676**
Coll: 8.4.2018 (A. M. Smith, D. P. Gordon, H. Mello)
Lat: -47.137222222222222
Long: 168.12583333333333
Depth: 80-85 m

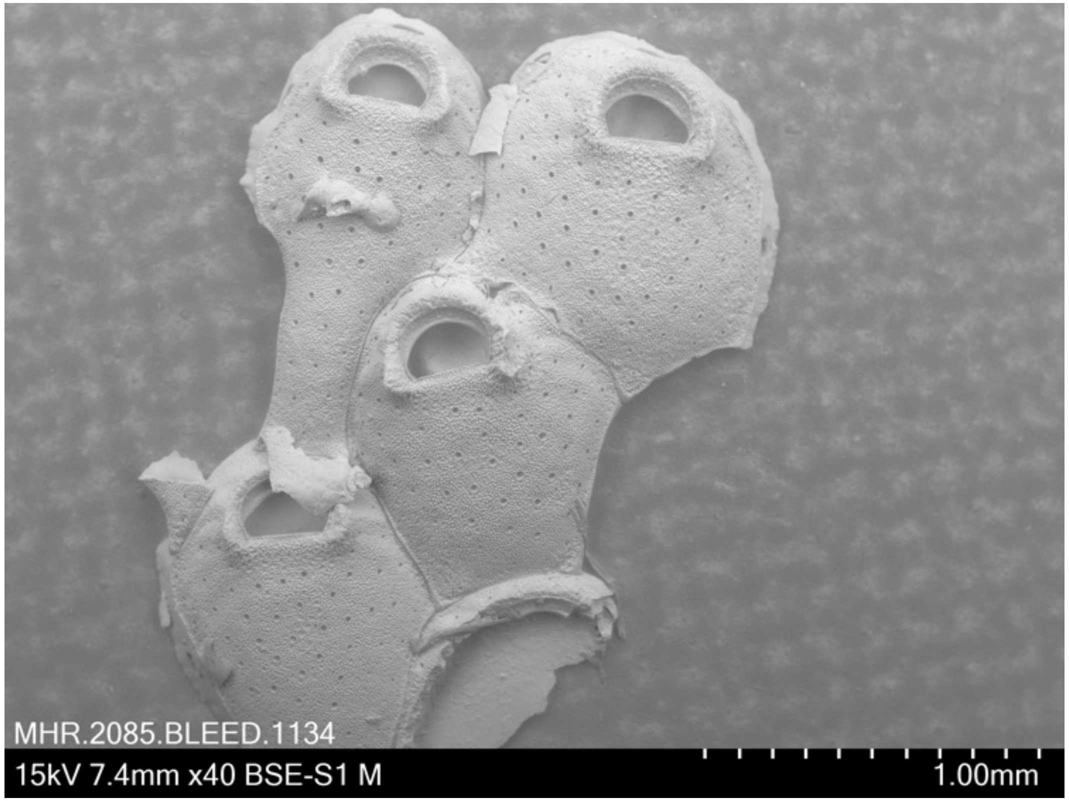


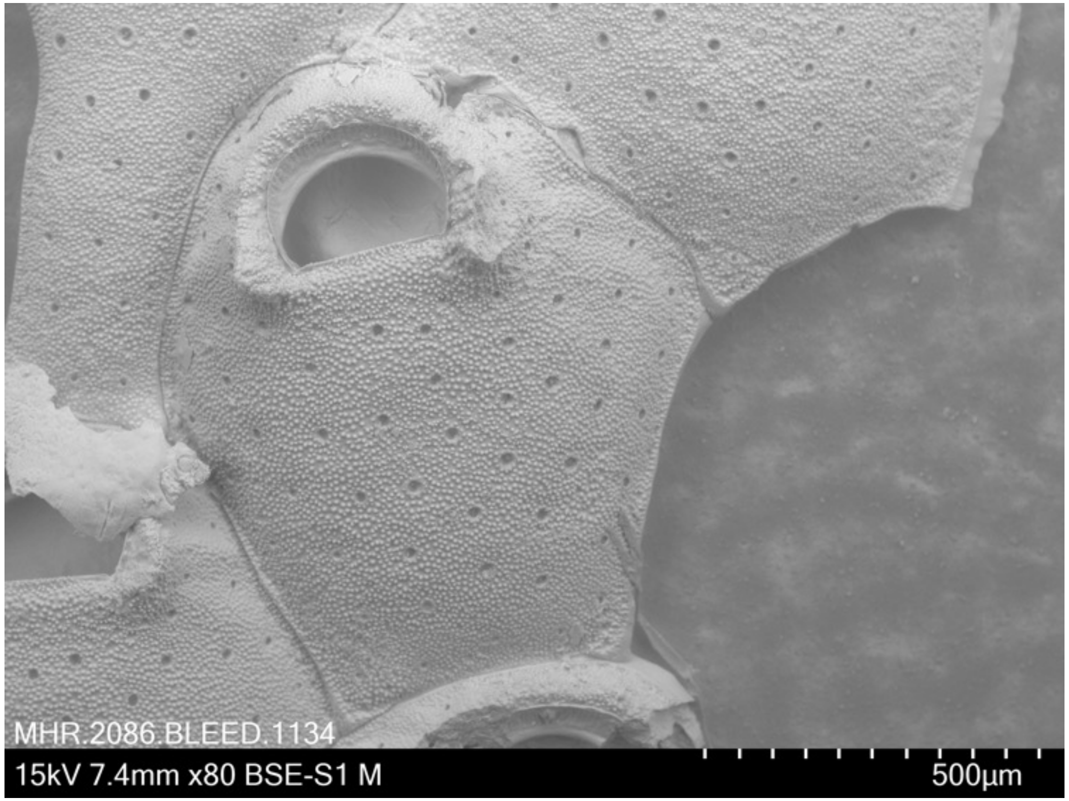


***Macropora nodulosa* BLEED 1134**
Coll: 10.11.2018 (A. M.  Smith)
Lat: -45.835833000000001
Long: 170.96166700000001
Depth: 131 m

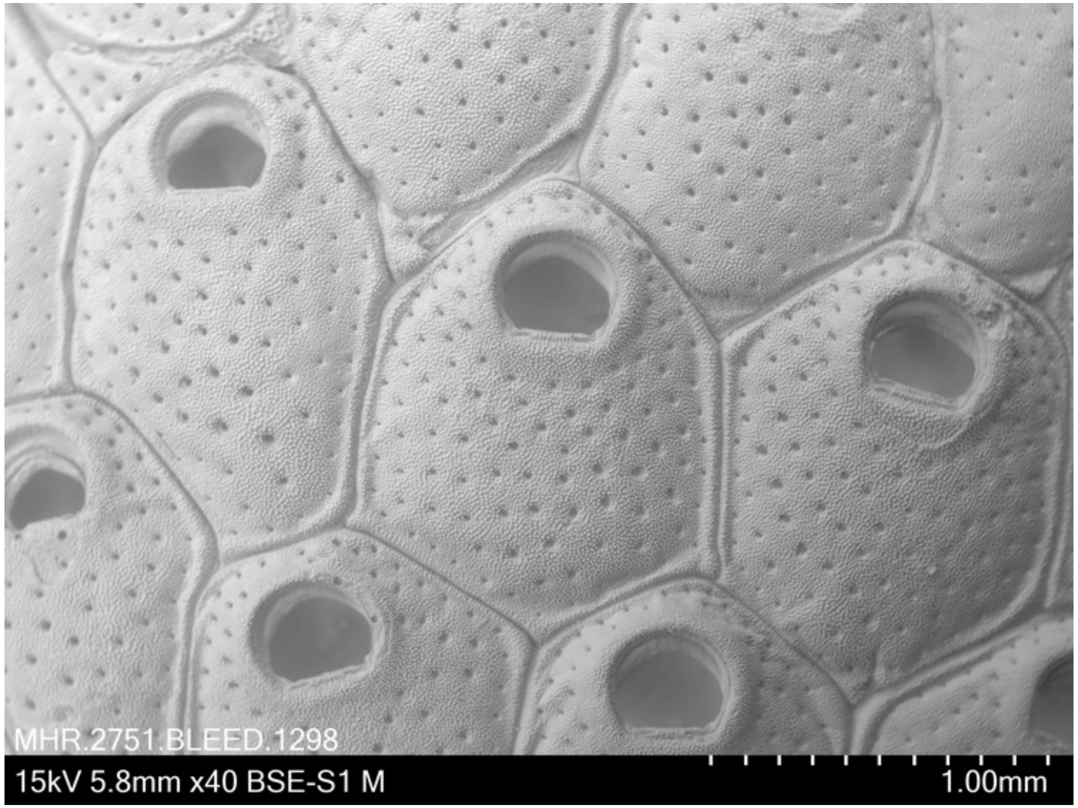


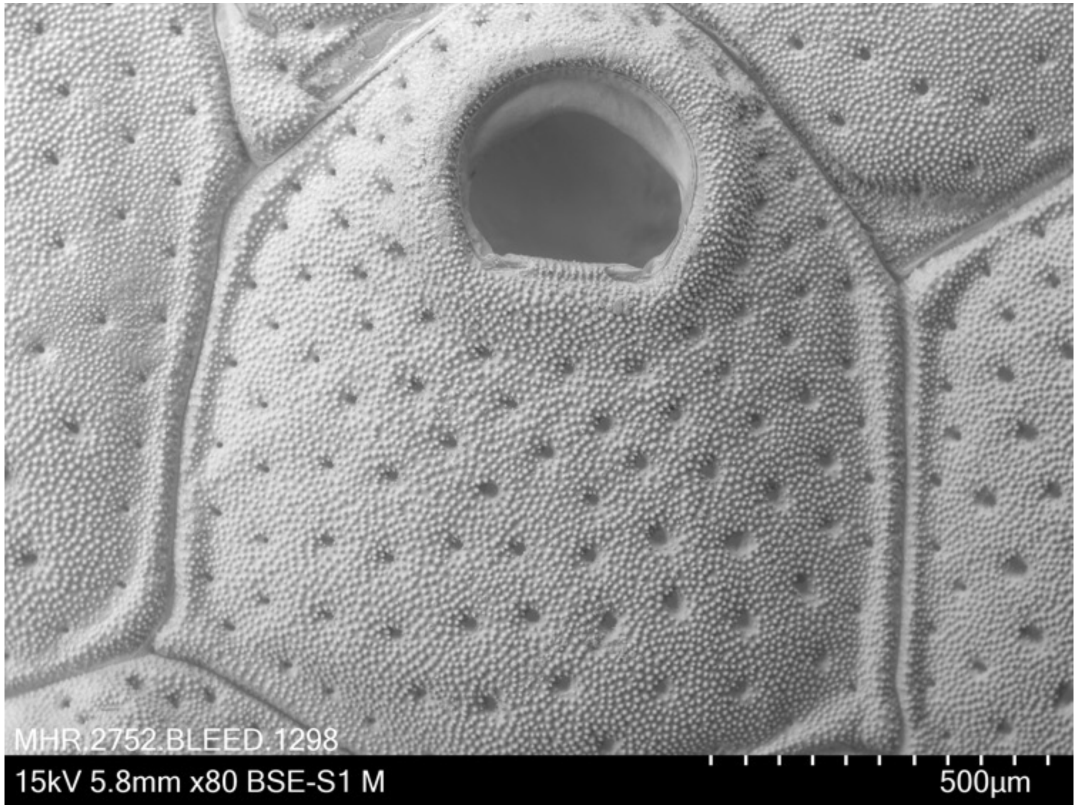


***Macropora levinseni* BLEED 1298**
Coll: 17.3.2019 (A. M. Smith, D. P. Gordon)
Lat: -34.21
Long: 172.11
Depth: 104 m

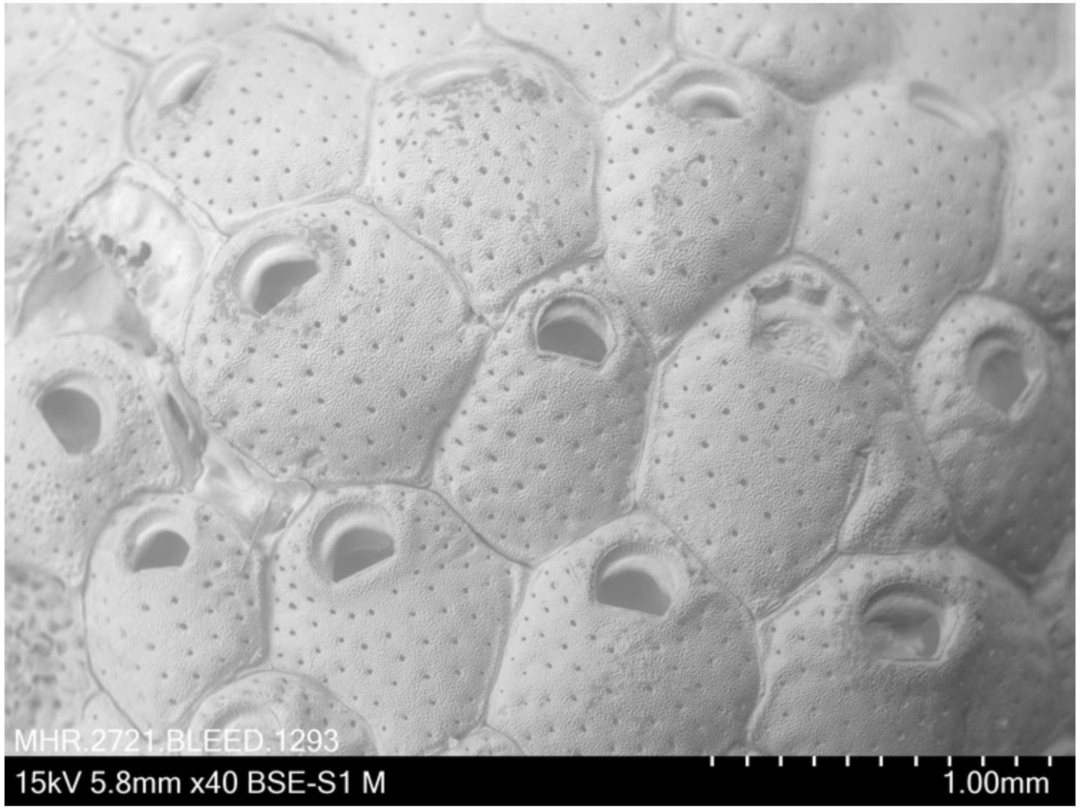


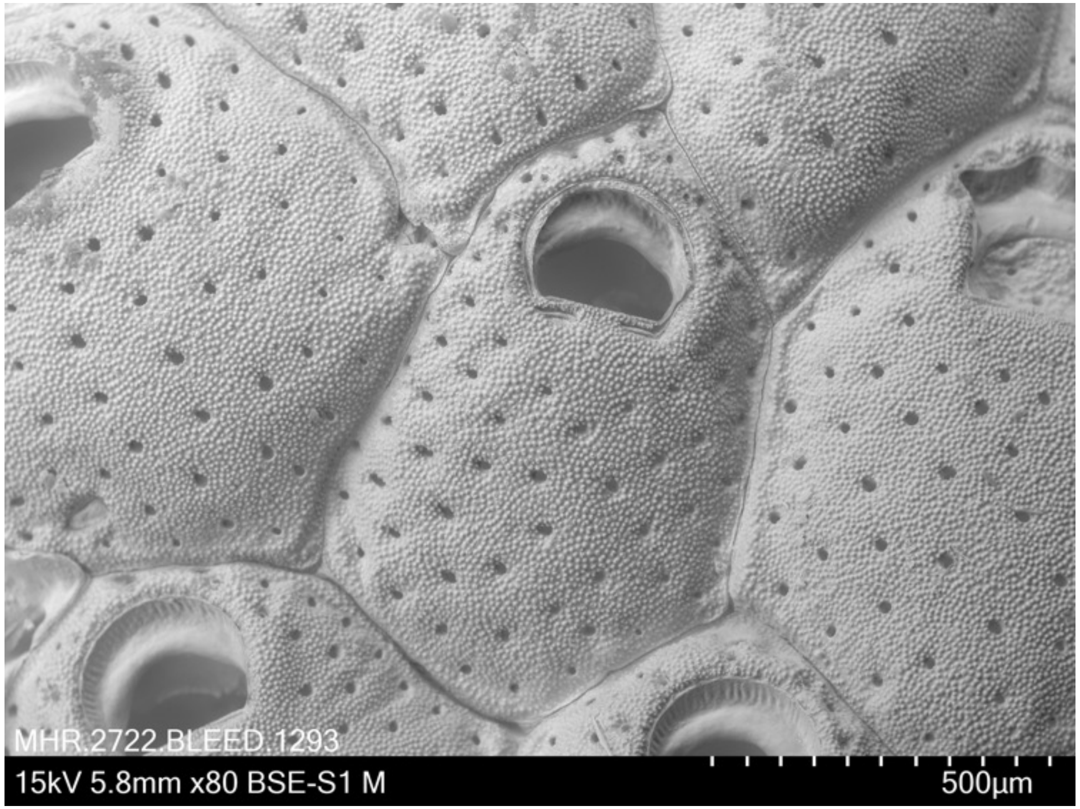


***Macropora grandis* BLEED 1293**
Coll: 17.3.2019 (A. M. Smith, D. P. GoOrdon)
Lat: -34.21
Long: 172.11
Depth: 104 m

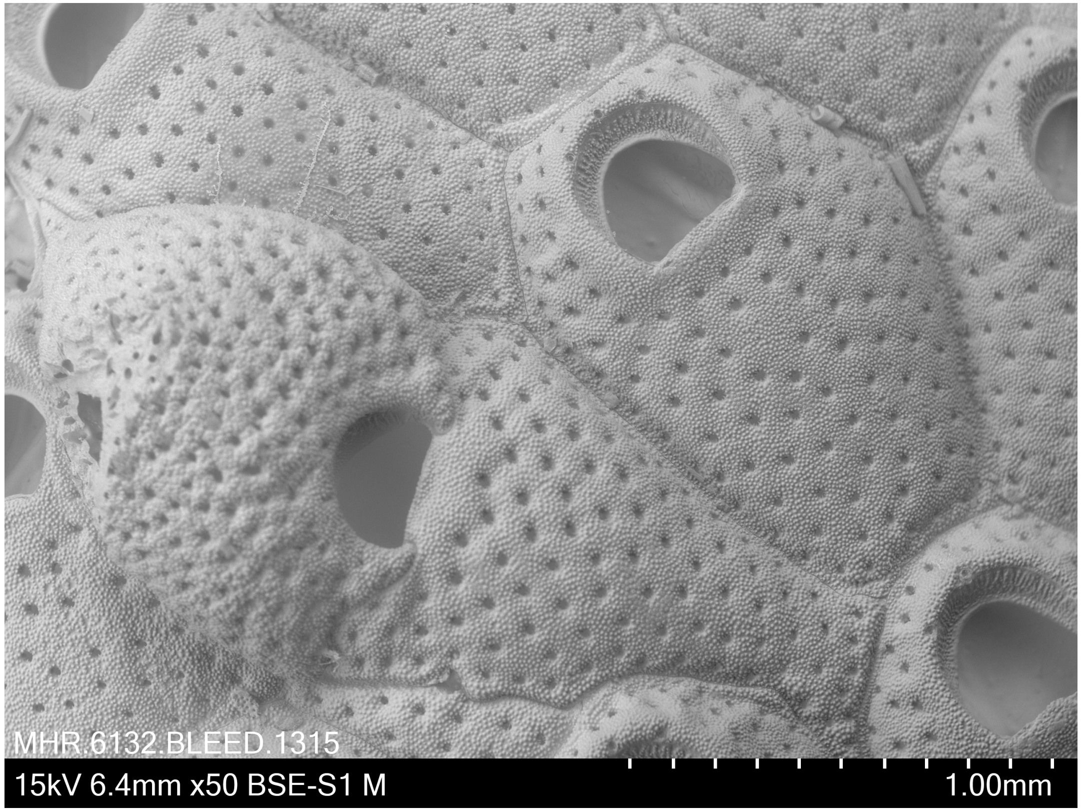


***Macropora grandis*** **BLEED 1315**
Coll: 15.3.2019 (A. M.  Smith; D. P. Gordon)
Lat: -36.46
Long: 174.9
Depth: 37 m

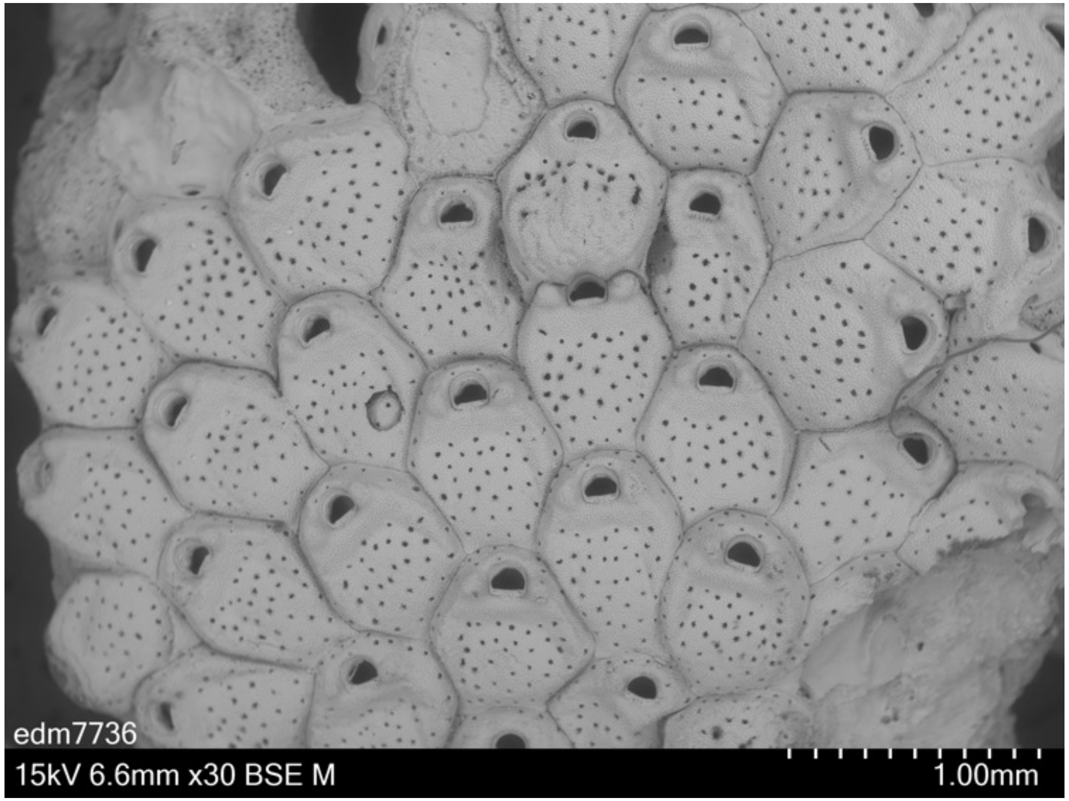


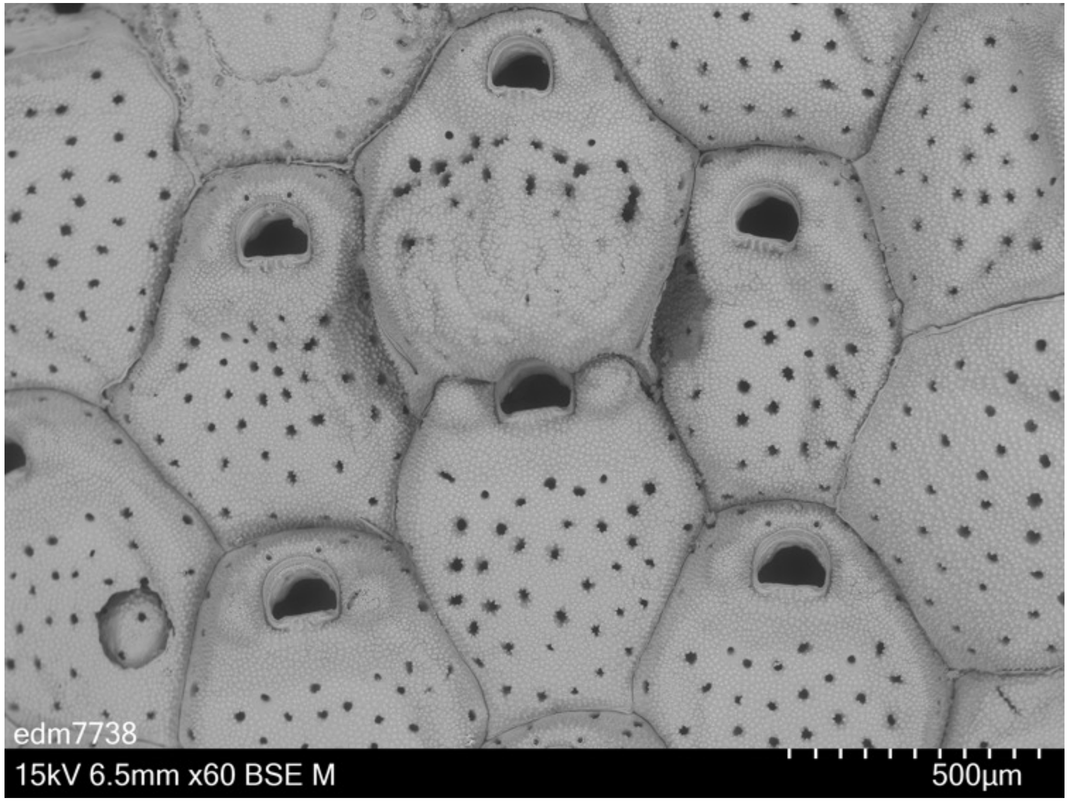


***Monoporella* n. sp. BLEED 1360**
Coll: 17.3.2019 (A. M. Smith, D. P. Gordon)
Lat: -34.21
Long: 172.11
Depth: 104 m

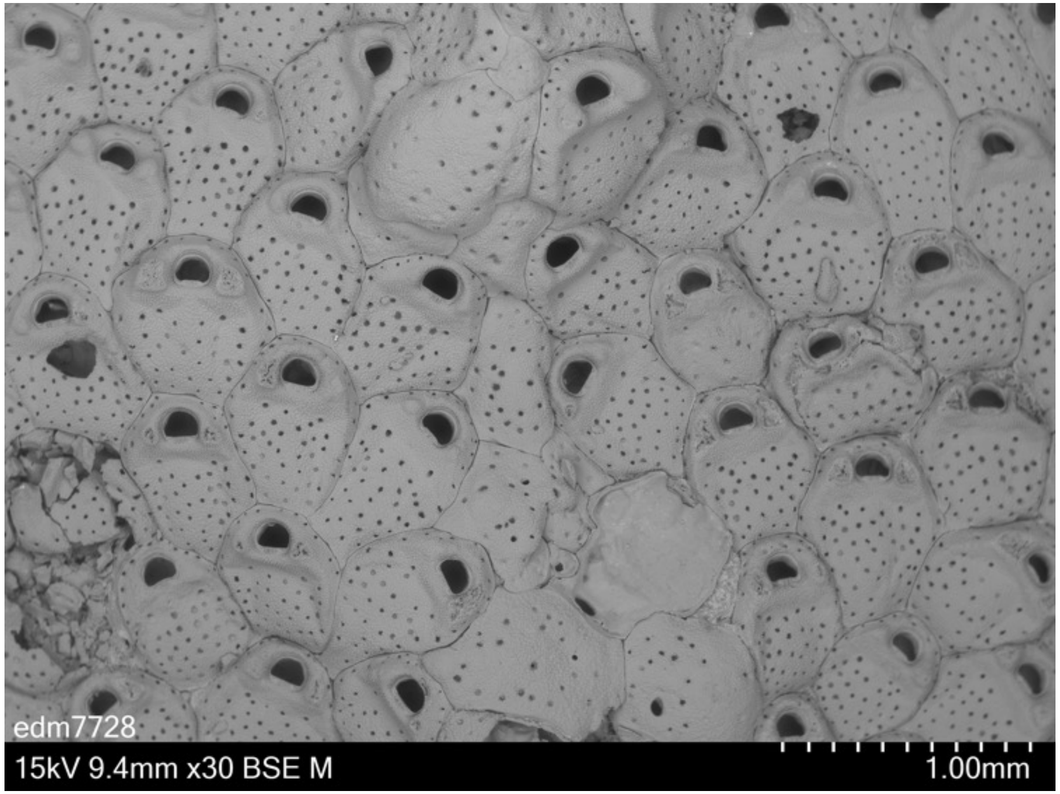


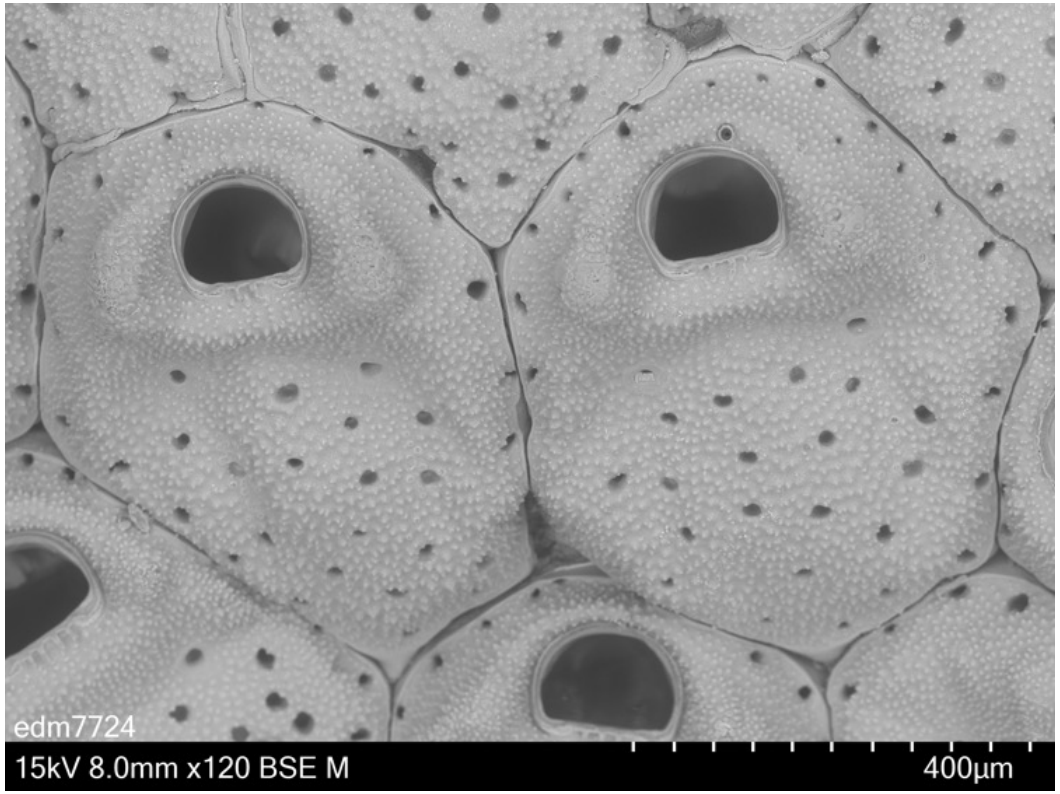


***Monoporella*** **n. sp. BLEED 1690**
Coll: 18.3.2019 (A. M. Smith, D. P. Gordon)
Lat: -34.3800
Long: 172.86
Depth: 54.6 m

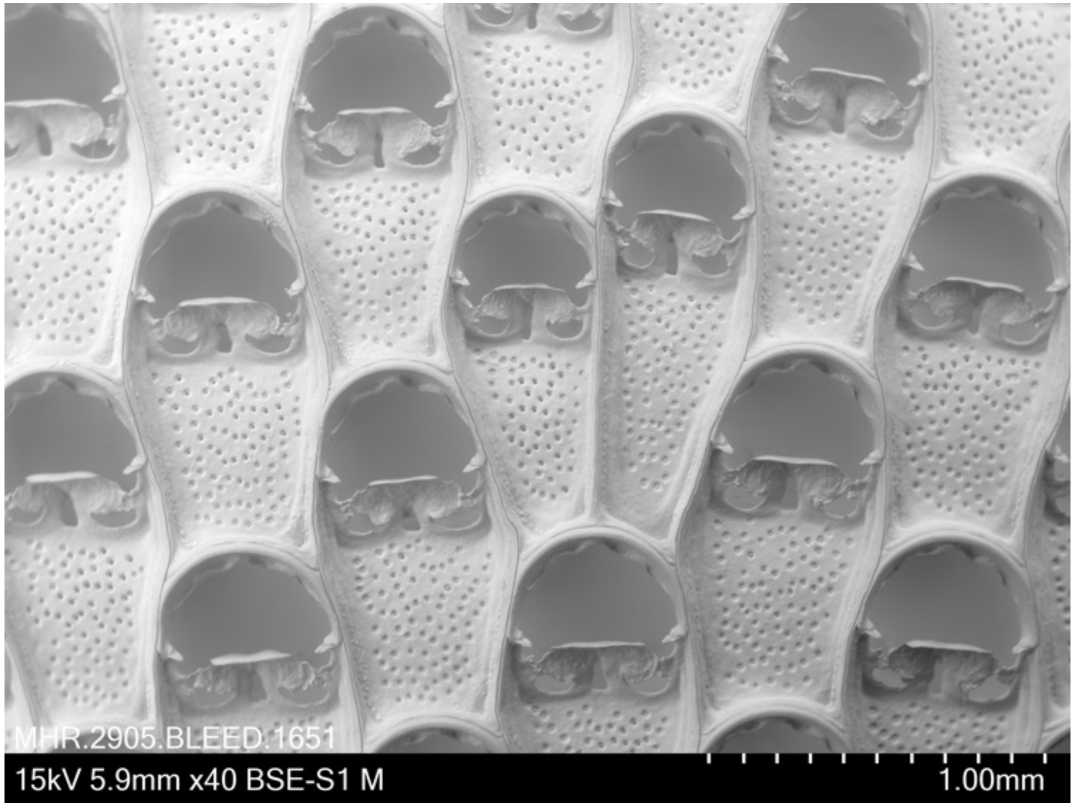


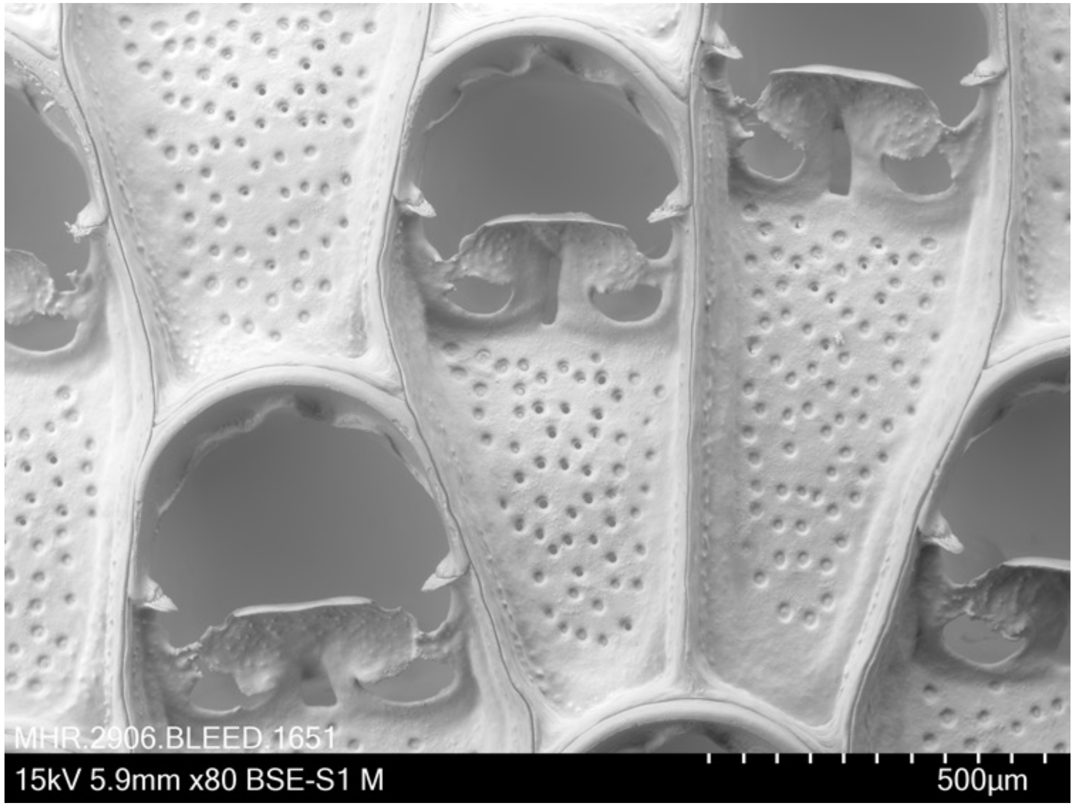


***Steginoporella perplexa*** **BLEED 1651**
Coll: 17.3.2019 (A. M.  Smith, D. P. Gordon)
Lat: -34.159999999999997
Long: 172.07
Depth: 98.9 m

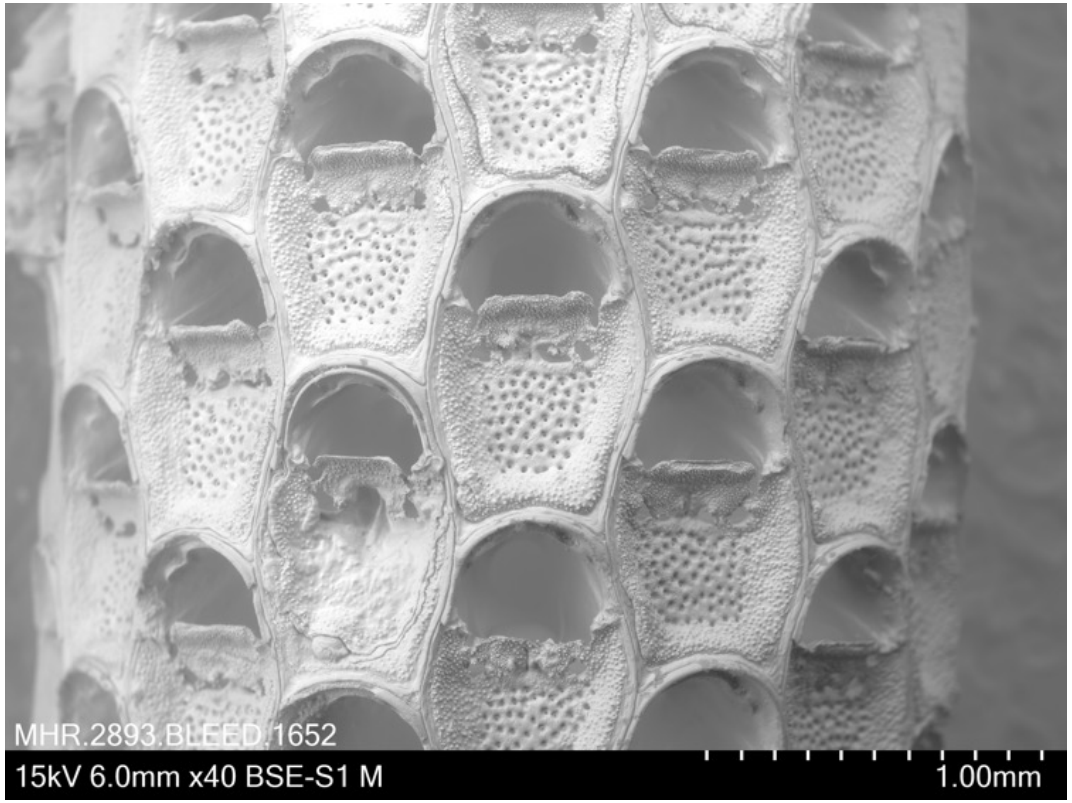


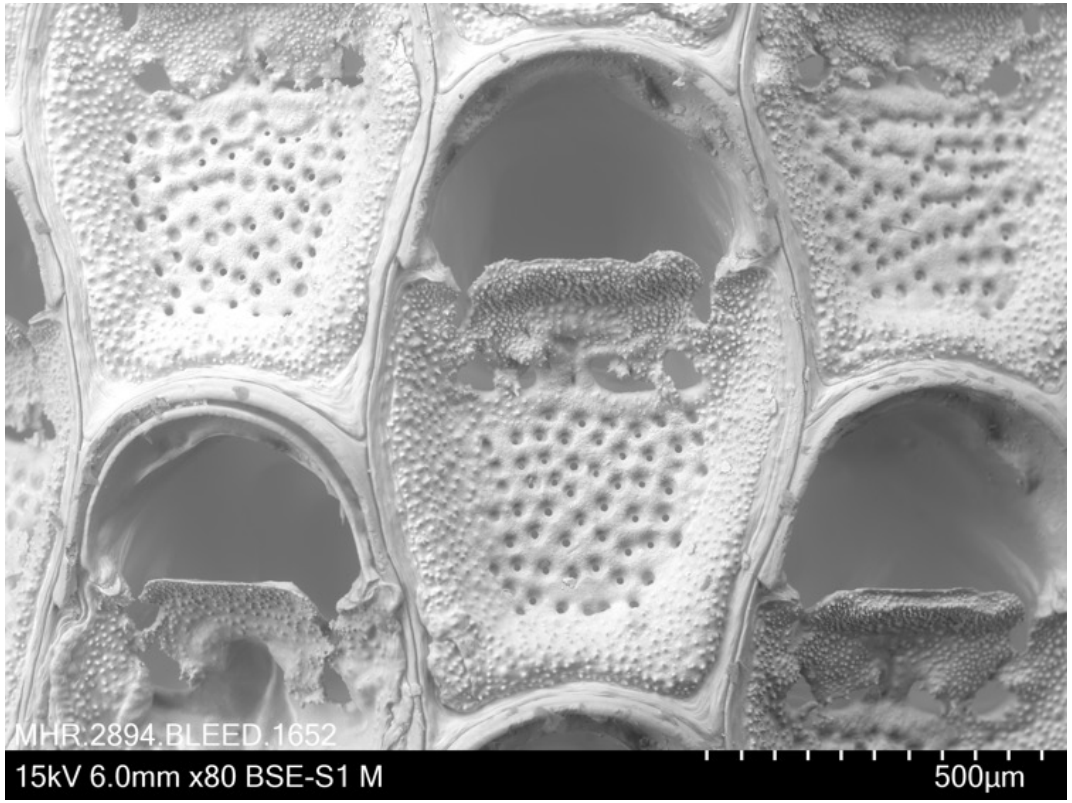


***Steginoporella marcusi* BLEED 1652**
Coll: 17.3.2019 (A. M. Smith, D. P. Gordon)
Lat: -34.159999999999997
Long: 172.07
Depth: 98.9 m

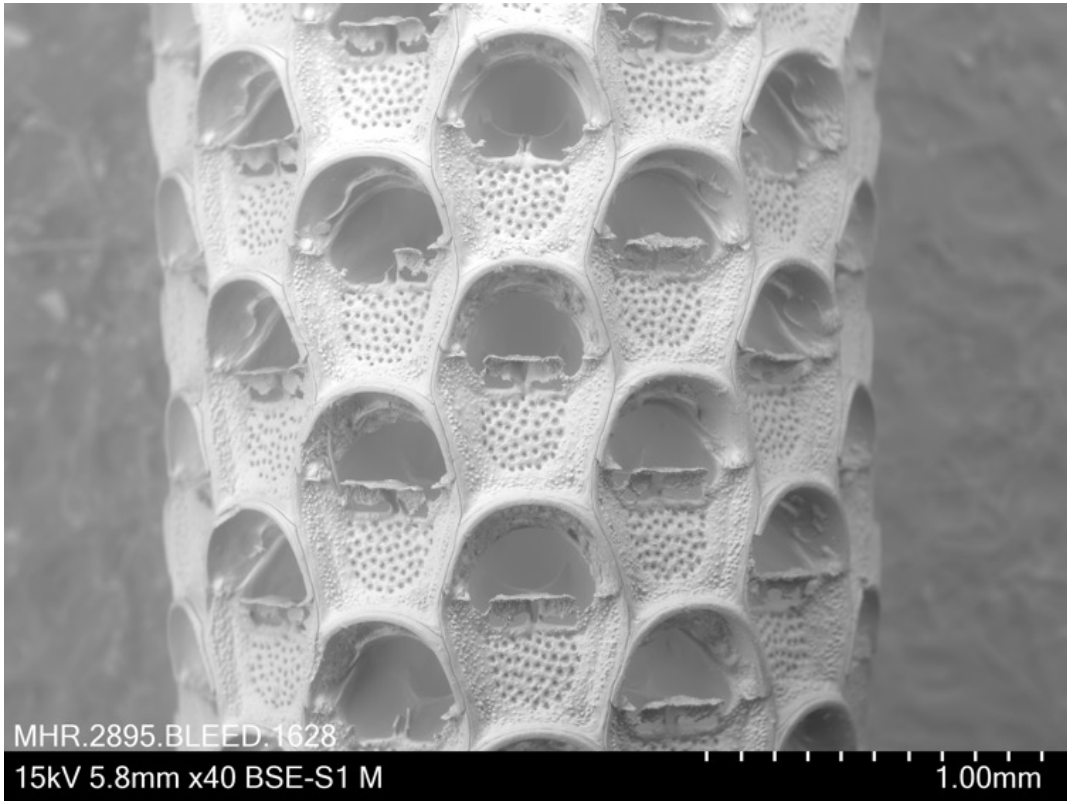


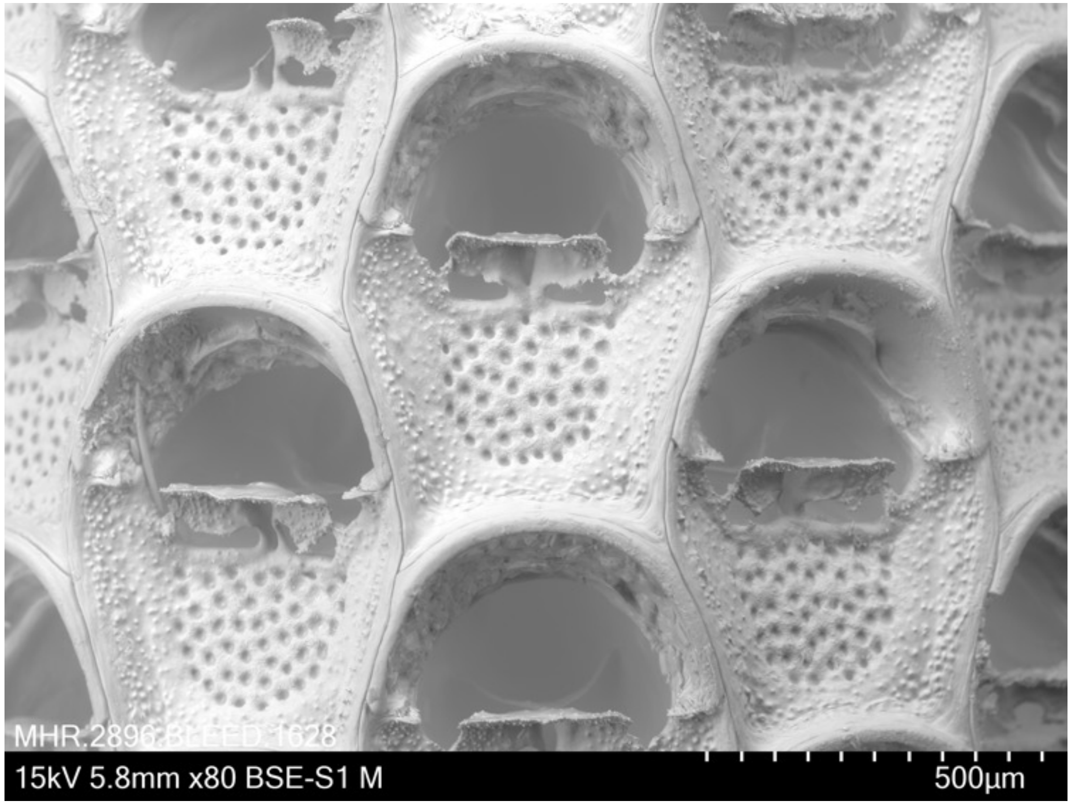


***Steginoporella neozelanica* BLEED 1628**
Coll: 17.3.2019 (A. M. Smith, D.P. Gordon)
Lat: -34.159999999999997
Long: 172.09
Depth: 82.9 m

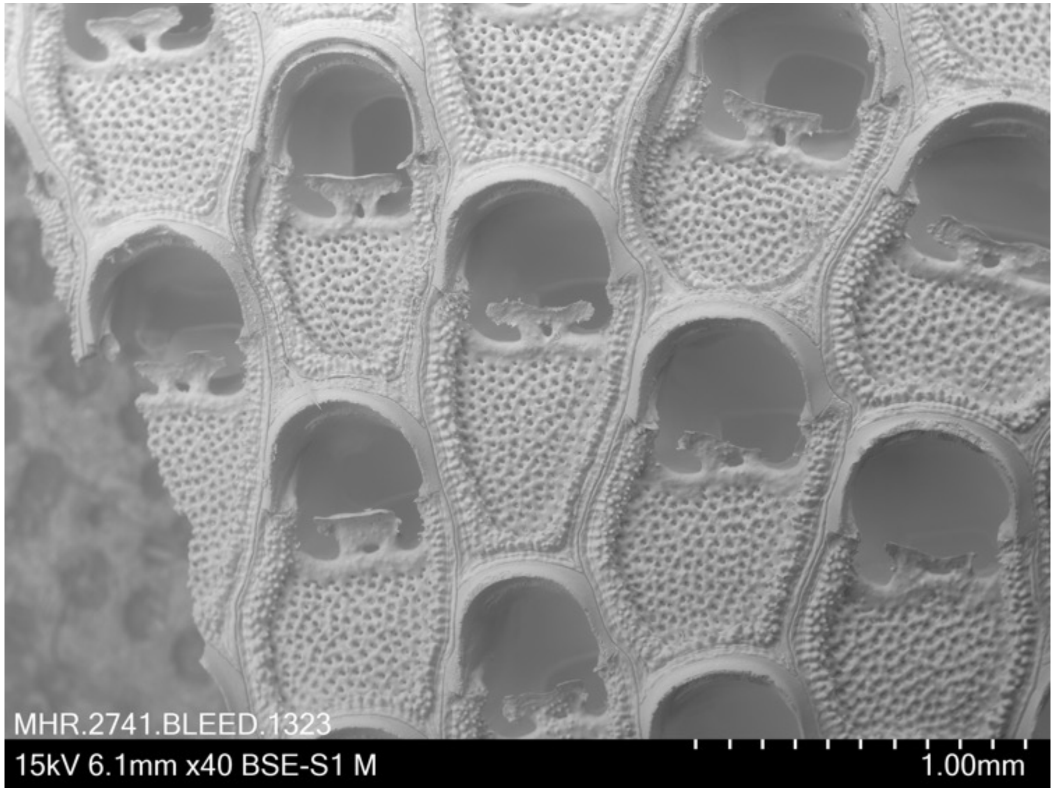


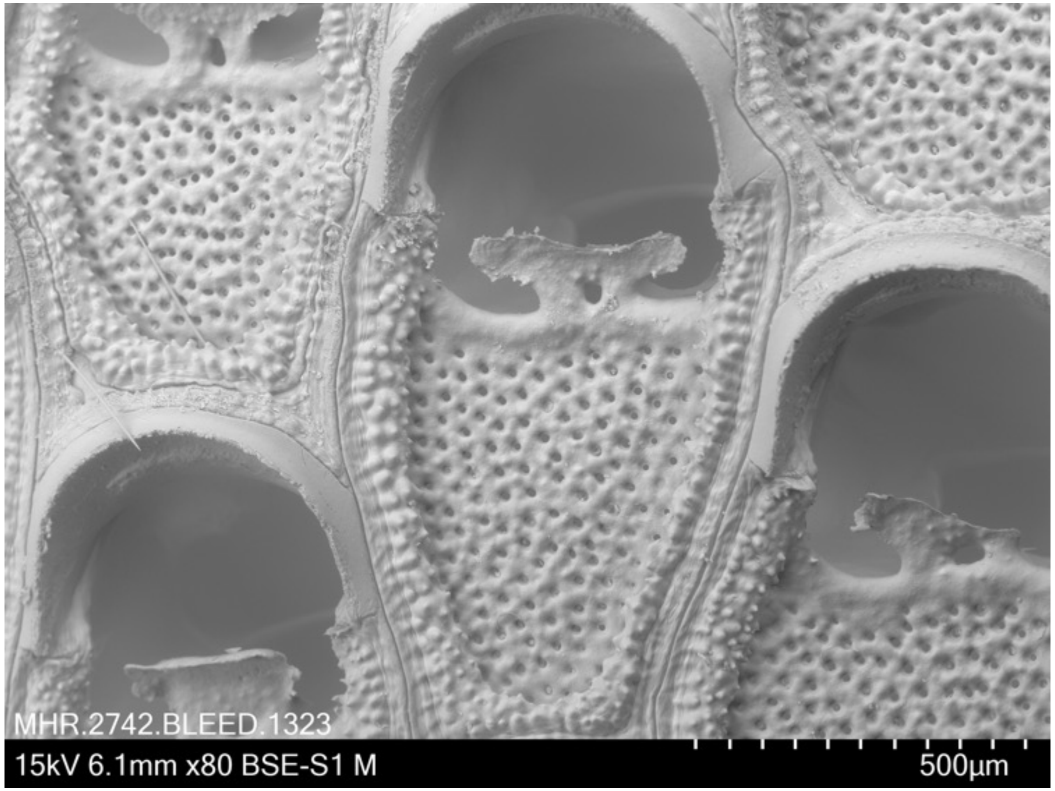


***Steginoporella magnifica* BLEED 1323**
Coll: 15.3.2019 (A. M. Smith, D. P. Gordon)
Lat: -36.46
Long: 174.9
Depth: 37 m

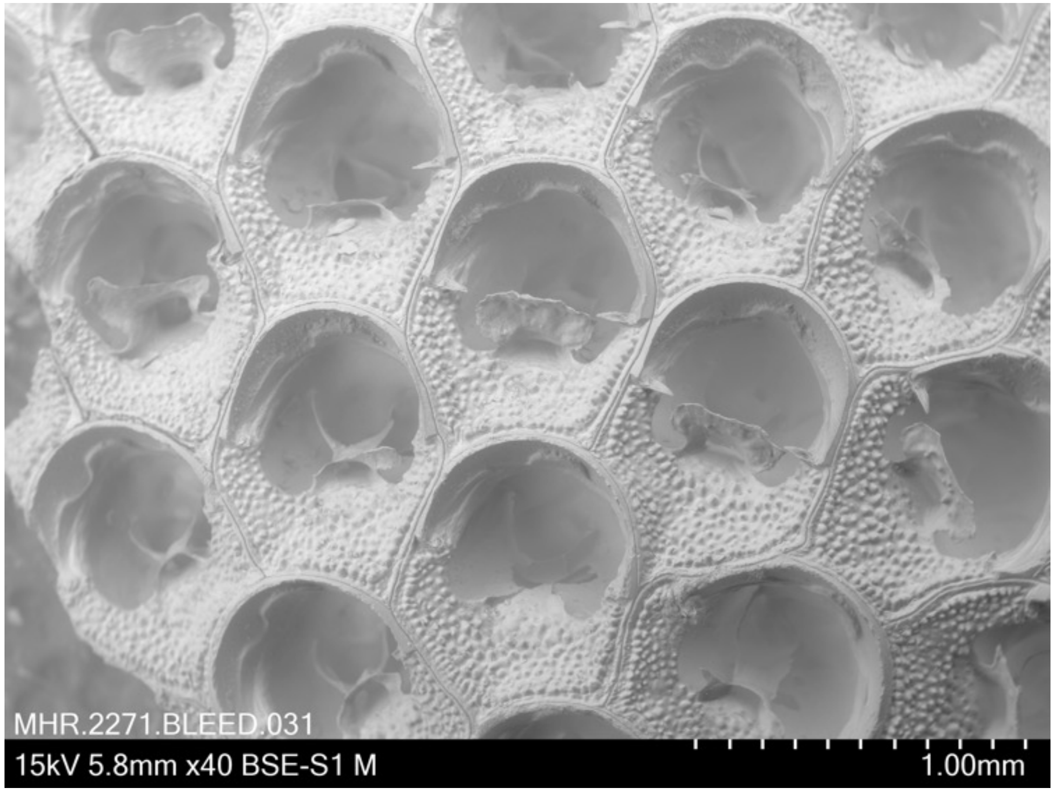


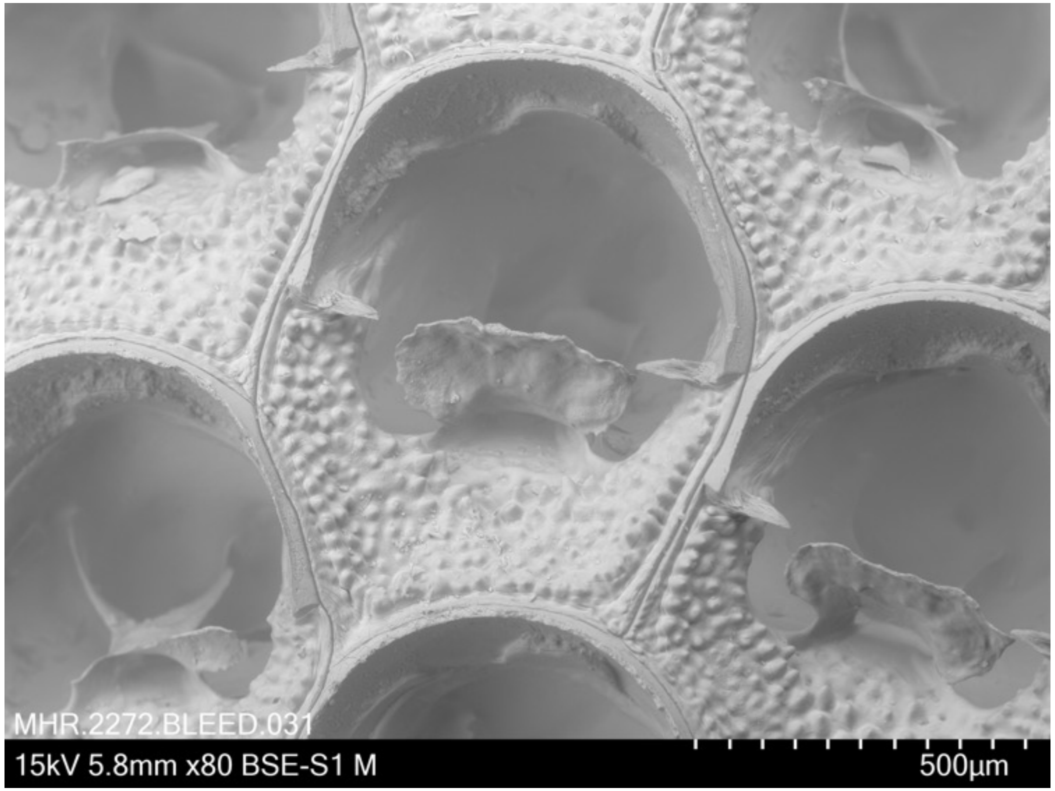


***Steginoporella discors* BLEED 31**
NIWA 54418 (TAN0906/2)
Coll: 4.7.2009
Lat: -35.55515
Long: 174.5601667
Depth: 60 m

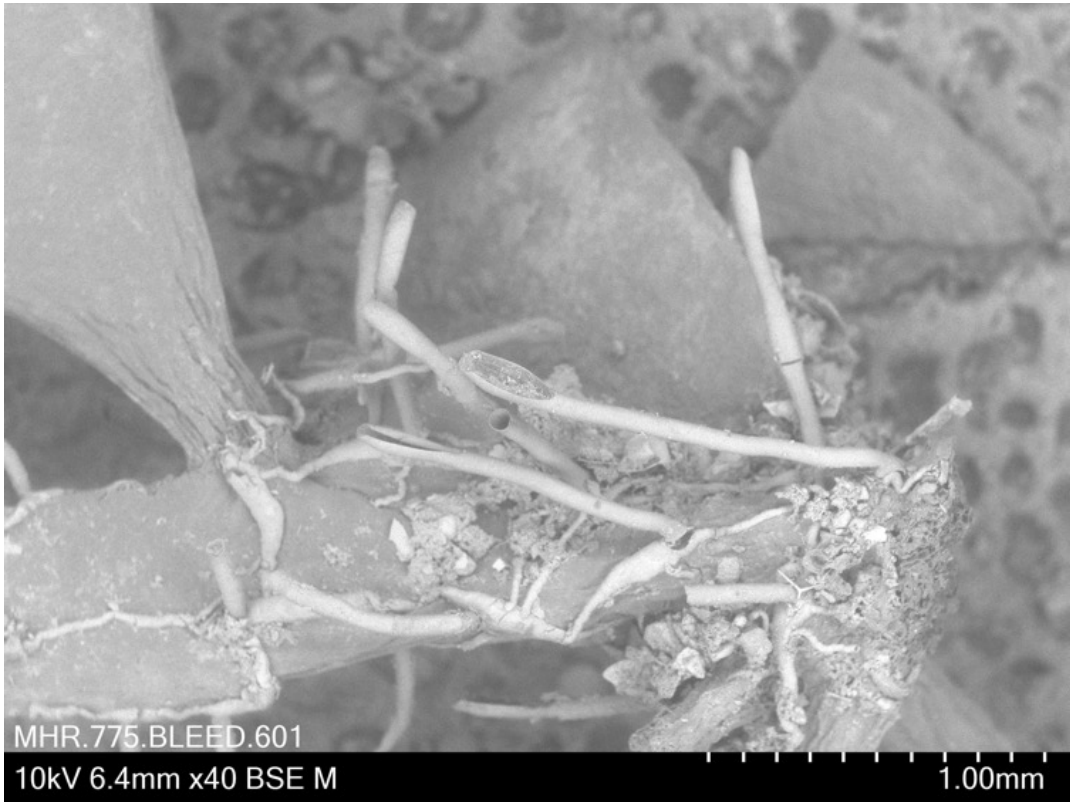


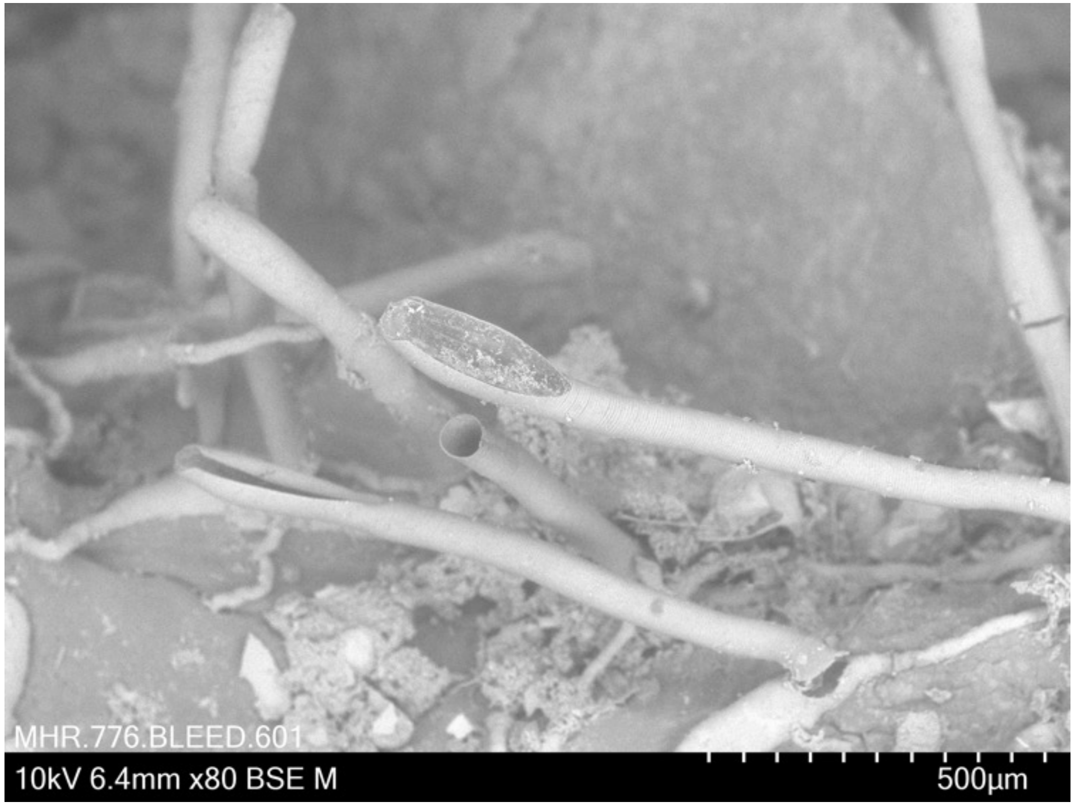


***Aetea* sp. BLEED 601**
Coll: 11.4.2018 (A. M. Smith, D. P. Gordon, H. Mello)
Lat: -46.744166666
Long: 168.325555555555
Depth: 29 m
